# R-package Jsmm: Joint species movement modelling of mark-recapture data

**DOI:** 10.64898/2026.02.24.707702

**Authors:** Luisa F. Rodriguez, Otso Ovaskainen

## Abstract

1. With small-bodied species, it is difficult to directly track individual movements, leaving mark-recapture as the most feasible method for collecting movement data. Mark-recapture data are challenging to analyse because they are indirect: many individuals are never seen after release, and for recaptured individuals there is no information on the movements between release and recapture locations. This makes it difficult to apply many statistical approaches that have been developed for continuous movement data. Among the statistical methods targeted specifically to mark-recapture data, most are focused on the estimation of population sizes or vital parameters rather than the estimation of movement behaviours.
2. We present the R-package Jsmm that expands and implements the earlier published Joint Species Movement Modelling (JSMM) framework with Bayesian inference. Jsmm estimates parameters related to habitat selection (behaviour at edges between habitat types), diffusion (random component of movement), advection (directional component of movement) and reaction (mortality rate), and their dependence on spatial, temporal or spatiotemporal covariates. Jsmm implements both instantaneous capture process and cumulative capture process, enabling its applications to a broad range of studies. If applying Jsmm to data on multiple species, it can estimate how species-specific parameters depend on species traits and/or phylogenetic relationships.
3. We use real and simulated case studies to demonstrate the workflow of Jsmm: (1) defining the model through importing the spatial domain, the spatiotemporal covariates, and the capture-recapture data; (2) fitting the model with Bayesian inference and evaluating model fit through posterior predictive checks; and (3) using the fitted model for inference and/or prediction. The simulated example validates the technical implementation by showing that the estimated parameters match with the assumed values. The real data example on moth light-trapping illustrates the practical utility of the package.
4. The R-package Jsmm offers a flexible resource for analysing capture-recapture data in a model-based framework that explicitly accounts for the spatiotemporal study design of where and when captures are attempted. By analysing data jointly on multiple species, the approach facilitates analyses of sparse datasets where the low number of recaptures would not allow fitting species-specific models separately for each species.

## 1 Introduction

Animal movement is a key driver for population and community dynamics, and hence a central problem in ecology is to understand why, how, when and where organisms move (Nathan,2008). Our ability to comprehend how animal populations respond behaviourally to environmental variations is crucial for conservation and monitoring programs, and for assessing possible economic risks related to biodiversity loss (Alagador & Cerdeira, 2022; Elsner et al., 2025; Schultz et al., 2019).

With current technology, it is possible to directly track the movements of many animal species in a continuous fashion using e.g., GPS loggers (Forrest et al., 2025; Quaglietta & Porto, 2019). However, especially with small-bodied species such as many insects, making direct observations of movements is still not feasible, whereas conducting mark-recapture studies is typically possible (Schultz et al., 2019). Compared to direct observations on movement, analyses of mark-recapture data are challenging because they are indirect: many individuals are marked and never seen again, and for others that are recaptured at specific locations and times, there is no information about how the individual moved between the observations. For this reason, mark-recapture methods are difficult to analyse with statistical approaches that have been developed for direct movement data, such as step selection models (Efford & Schofield, 2022; Florko et al., 2025).

While there is a large body of literature on statistical methods for analysing mark-recapture data, most of this literature has focused on using the data to estimate population sizes or vital parameters such as fecundity and mortality rates rather than to estimate movement behaviours (Tourani, 2022; White, 2008; Williams et al., 2002). The existing approaches for inferring movement behaviours from mark-recapture data can be categorized into two different groups. First, multi-state models (Neil Arnason, 1973; Quaglietta & Porto, 2019) assume that observations are made in a discrete number of sites (such as breeding colonies of a bird species), and they focus on estimating the pairwise transition rates at which individuals move between those sites. Second, spatially explicit models, such as search-encounter models (Royle & Young, 2008; Royle et al., 2014) and diffusion-advection-reaction models (Ovaskainen, 2008), consider how individuals move in a continuous landscape (e.g., following a random walk with tendency to remain within a home range). Here, we focus on this second class of models, specifically the Joint Species Movement Modelling (JSMM) framework developed by Ovaskainen et al., 2019. This framework enables parametrizing habitat selection, movement, mortality, and detection rates as a function of spatial predictors, and it can be applied both for single species and for multi species in mark-recapture studies.

A bottleneck for the parameterization of spatially explicit movement models to mark-recapture data has been the lack of user-friendly and generally applicable software (Lagrange et al., 2014). Here we resolve this bottleneck by presenting the R-package Jsmm. In addition to providing a user-friendly implementation of the JSMM framework, we extend the original approach of Ovaskainen et al., 2019 in three ways. First, we incorporate not only spatial but also temporal and spatio-temporal covariates. Second, we incorporate not only habitat selection, diffusion, and mortality but also advection that can be used to model e.g., homerange movements. Third, we enable one to utilize data collected not only by an instantaneous capture process (Ovaskainen, 2004) but also by a continuously accumulating capture process, such as light-trapping of moths (Gray et al., 2022). To verify the technical validity of our implementation, we apply it to simulated data with known parameter values. To illustrate the functionality of Jsmm in the analysis of real data, we apply it to a case study of moths, the data including movements of 21 species within a heterogeneous landscape consisting of forest fragments and the surrounding agricultural areas (Slade et al., 2013).

## 2 The scope and workflow of Jsmm

The R-package Jsmm implements the fitting of diffusion-advection-reaction models to multispecies mark-recapture data in a Bayesian framework. The model includes species-specific parameters of habitat selection, random (diffusion) and directional (advection) components of movement, mortality (reaction), and the capture process (Fig. 1A). Each of these model parameters can be assumed to depend on spatially, temporally, or spatiotemporally varying covariates, which dependencies are incorporated through generalized linear models. Diffusion, mortality, and capture rate (in case of continuous capture process) are modelled through log-normal models, advection through a normal model, and capture probability (in case of instantaneous capture process) through logistic regression. The approach estimates for each species i, a vector of parameters *θ*_*i*_ that characterize its movement behaviour, mortality and capture, and their dependency on covariates. The model involves a hierarchical structure that connects the species-specific parameters through a multivariate normal model. This enables borrowing information among the species, and asking how movements, mortality and capture, and their dependency on covariates, relate to species traits and phylogenetic relationships. The package Jsmm is developed in R (R Core Team, 2024) with some core functions developed in C++ to improve computational performance (Stroustrup, 2013). Technical specifications are provided in Supporting Information S1. The three steps of applying Jsmm are defining the model, fitting and validating the model, and using the model for inference and prediction (Fig. 2B):

**Figure 1:**
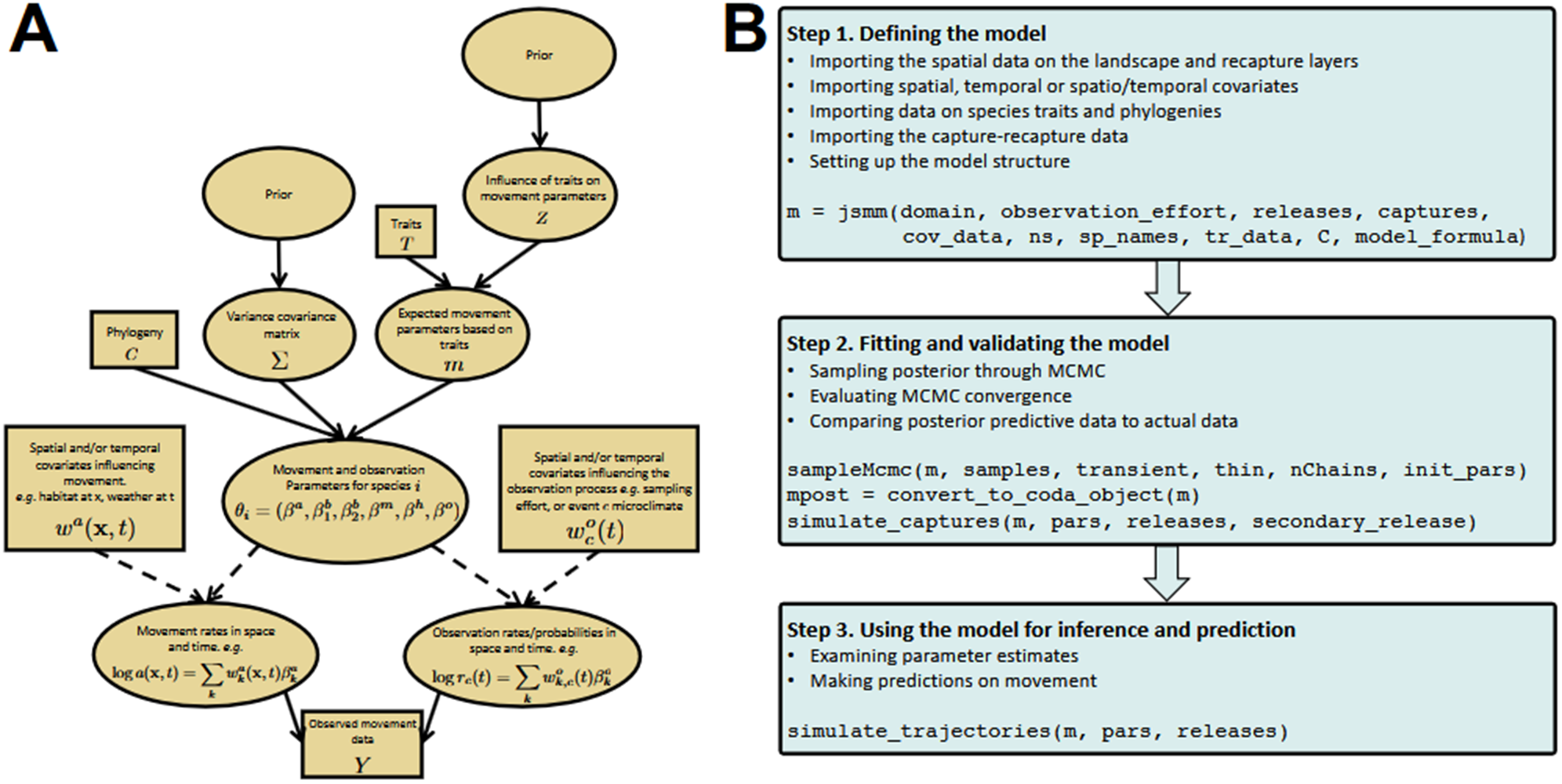
The scope and workflow of the R-package Jsmm. Panel A shows the Directed Acyclic Graph (DAG) of the statistical approach implemented in the R-package Jsmm. The model includes species-specific parameters *θ*_*i*_ that describe rates of diffusion (random component of movement), advection (directional component of movement), reaction (mortality) and catchability, and their dependence on spatiotemporal predictors. These species-specific parameters are modelled as a function of species traits and phylogenetic or taxonomic relationships. Panel B shows the three steps of the Jsmm workflow: defining the model, fitting it to data and validating the model through evaluation of model fit, and using the model for inference and prediction.

**Figure 2:**
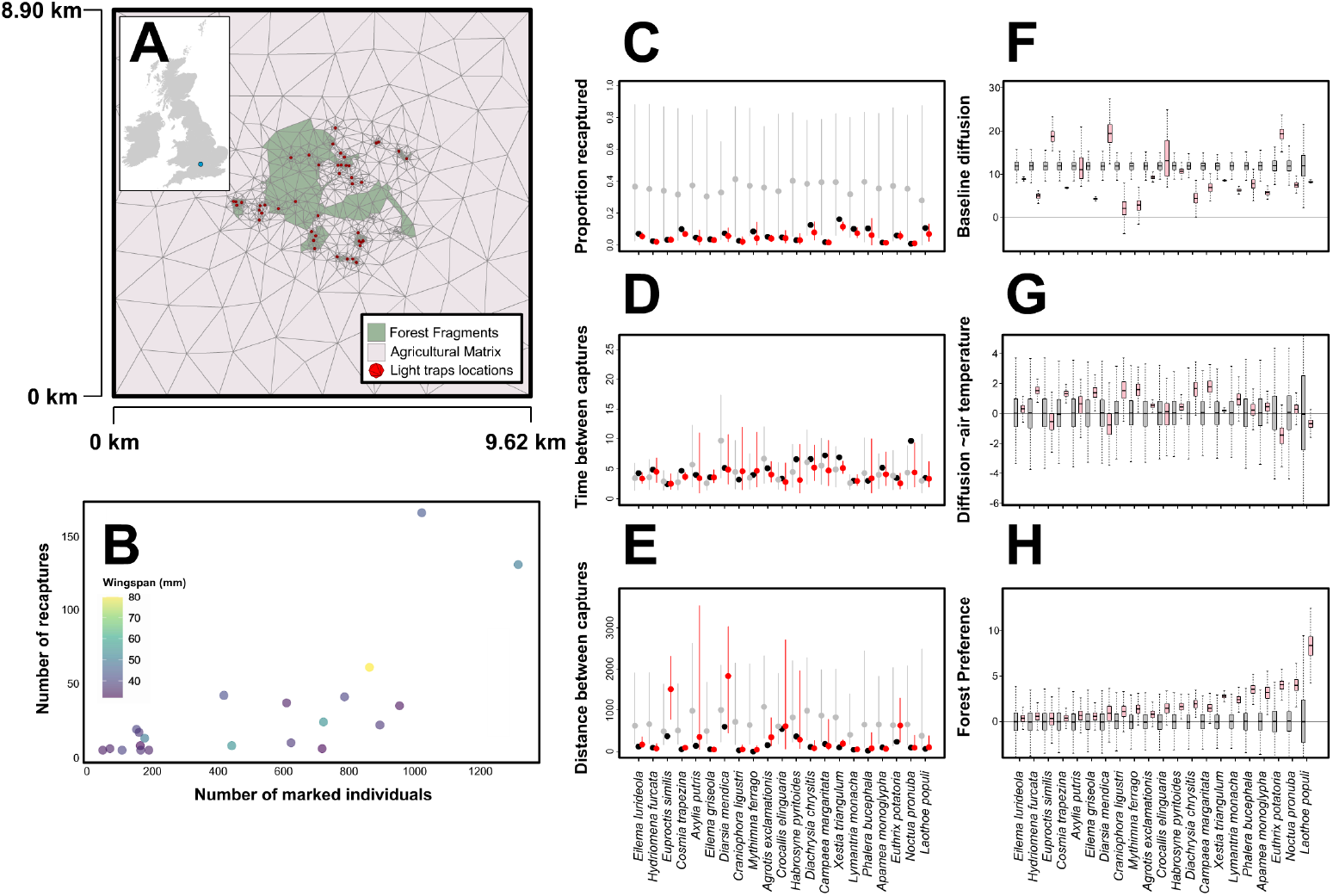
Case study on moth mark-release-recapture data analyzed with the R-package Jsmm. Panel A shows the study area where moths were trapped with a network of 44 light traps. The colors refer to habitat type, and the triangulation was used to implement the numerical computations. ¬¬Panel B shows that the 21 species included in the study varied markedly in the numbers of released and recaptured individuals and in their wingspan. Panels CDE illustrate evaluation of model fit, implemented by examining how well posterior predictive data (red dots and error bars show means and 95% quantiles) are able to replicate relevant summary statistics of the real data (black dots), in comparison to prior predictive data (grey dots and error bars show means and 95% quantiles). The species-specific summary statistics shown are the proportion of recaptured individuals (C), the mean time between captures (D), and the mean distance between captures (E). Panels FGH exemplify species-specific parameter estimates by showing a comparison between posterior (pink) and prior (grey) distributions. The boxes show medians and 75% quantiles, and the whiskers 95% quantiles. In panels C-H, the species have been ordered from smallest (left) to largest (right).

### Step 1. Defining the model

In this step, the user imports the data files and specifies the model assumptions. The input data consist of a description of the spatial domain (named list domain), describing the locations and the times of releases and recaptures (named list `observation_effort`), the mark-recapture data (named list `data`), the spatial, temporal and spatio-temporal covariates (named list `cov_data`), and optionally species traits and phylogenetic relationships (named list `tr_data` and matrix C). The user specifies assumptions on how model parameters are assumed to depend on covariates and species traits using model formula notation (named list `model_formula`, including e.g., `diffusion` *∼* `air_temperature` and `traits` *∼* `body_size`). The model constructor function jsmm then creates an unfitted model object that includes all the data and information about the model structure and the assumed prior distribution (for information on the default prior, see Supporting Information S1).

### Step 2. Fitting and validating the model

In this step, the user fits the model through posterior sampling via Markov chain Monte Carlo (MCMC) methods (function `sampleMcmc`, with parameters describing the desired number, length, transient, and thinning of the chains). The function `convert_to_coda_object` exports the posterior distribution to the format used by the R-package coda (Plummer et al., 2006) to facilitate examining MCMC convergence. To test if the structural model assumptions are in line with the data, the user can evaluate model fit by comparing the actual data with a predictive posterior distribution (function `simulate_captures`) in terms of relevant summaries, such as numbers and spatial and temporal characteristics of recaptures (see examples on Supporting Information S2 and R-package scripts).

### Step 3. Using the model for inference and prediction

In this step, the user postprocesses results from the fitted model to addresses the questions that motivated the study. This typically involves examining parameter estimates of interest, such as the effects of covariates on the model parameters or the effect of species traits on these parameters. The user may also perform scenario simulations to ask how the species would move, e.g., under modified landscape structure (see Gray et al., 2022 for an example).

## 3 Case studies

### Simulated case studies

To verify the functionality of the R-package, we applied it to simulated data sets with known parameter values. These case studies suggested the technical correctness of our implementation, as the estimated posterior distributions were concentrated near the true values. For full details on the simulated case studies, we refer to the Supporting Information S2: Demos simulated ICP and CCP and their corresponding R scripts on GitHub, which illustrate both cumulative and instantaneous capture processes.

### Case study on moths mark-recapture data

To illustrate the use of the R-package in actual data analysis, we used it to reanalyse the mark-recapture data originally collected by Slade et al., 2013 and used as a case study in Ovaskainen et al., 2019. The study was conducted in a 400-ha fragmented forest around Wytham Woods in southern England, where 44 light traps were placed within forest fragments and at solitary oak trees located within agricultural matrix (Fig. 2A). The light traps were activated before dusk and visited at dawn for 31 days. We included in our analyses those 21 species that had at least five recaptured individuals, including in total 10593 individuals of which 666 were recaptured at least once, with substantial variation among the species (Fig. 2B). When defining the model, we assumed that the species may show habitat selection between the forest fragments and the agricultural matrix, that diffusion may depend on the air temperature, that mortality rate is constant, and that capture is of cumulative nature, reflecting the process of light trapping. We further assumed that all species-specific parameters may depend on the wingspan of the species and on taxonomical relationships among the species.

Evaluation of model fit through posterior predictive simulation showed that the model successfully captured the generally very low recapture rate (Fig. 2C) and that most recaptures were made soon after (Fig. 2D) and near (Fig. 2E) the release. The model successfully captured the variation among the species, i.e., which species had especially high capture rate or were recaptured after a long time or far away from the release (Fig. 2CDE). These predictions reflect the variation in the specie-specific parameter estimates (Fig. 2FGH). Consistent with previous analyses (Ovaskainen et al., 2019), we estimated that most species preferred forests over agricultural areas (Fig. 2G). While previous analyses could not incorporate temporally varying predictors, we found that the movement rate increased for some and decreased for some other species with air temperature (Fig. 2H). At the community level, we found that larger species showed a stronger preference for the forests over agricultural areas (posterior probability Pr=0.95) and that larger species may have higher mortality rates (Pr=0.89). Beyond the variation explained by the traits, we did not find support for taxonomically structured responses (Pr=0.19). For full details on this case study, see Supporting information S3 and scripts_moths on R-package scripts.

## 4 Discussion

The Jsmm software presented in this work provides a flexible framework and structured workflow for fitting diffusion-advection-reaction models to mark-recapture data. Our implementation allows analysing mark-recapture data acquired either by instantaneous or cumulative capture, and incorporating spatial, temporal, and spatiotemporal predictors of movement and capture parameters. If analysing movement data jointly for many species, the Jsmm framework enables asking how movement and capture parameters relate to species traits and phylogenies and improving parameter estimation by borrowing information across the species. The Bayesian model-based framework brings tools for model validation, e.g., through posterior predictive checks. The diffusion-advection-reaction framework facilitates evaluating the outcome of movements through multiple metrics, e.g., occupancy times or movement probabilities (Ovaskainen, 2008). The movement model can also be used to run scenario simulations, e.g., to ask how movement barriers or corridors other landscape modifications would influence movement behaviour (Ovaskainen et al., 2008). To support FAIR (Findable, Accessible, Interoperable, and Reusable) data principles, we provide the Jsmm software as an open-source repository with detailed vignettes and example scripts guiding step-by-step analyses.

We illustrated the application of the Jsmm software with a case study of moth movements. Our results show that the approach can effectively extract ecological signals from typical mark-recapture datasets. In line with previous studies on the influence of species traits on dispersal behavior (Betzholtz et al., 2019), we found species with larger wingspans to show a preference for forest fragments over agricultural regions (Fig. 2H). We observed that for all species, the movement rates were sensitive to changes in the air temperature, with many species increasing their movements at higher temperatures (Fig 2G). We did not observe a clear relationship between temperature-dependency of movement rates and wingspan, possibly due to the multiple thermoregulation mechanisms that exist in moths (Heinrich, 2012).

While we consider the Jsmm package to be valuable resource for analyzing mark-recapture data, it has three limitations that suggest important areas for future development. First, mark-recapture are often (mostly) used to estimate population dynamic parameters such as fecundity and mortality rather than movement parameters (Tourani, 2022). While Jsmm estimates movement and mortality parameters, it does not address population dynamic parameters. How to utilize spatially explicit mark-recapture data to jointly estimate both movement and population dynamic parameters is an important future challenge that could be addressed by integrated population modelling (McClintock et al., 2022). Second, the present Jsmm approach considers both movement parameters and morphological traits at the species level, yet accounting for their within-species variation would be highly relevant to address questions related to e.g., evolution of dispersal. While within-species variation could be incorporated to present framework by modelling individual-level parameters through species-level mean and variance parameters, successfully parameterizing such a model would require a more extensive mark-recapture dataset than typically available in the multispecies context. Third, the continuous-time formulation of Jsmm brings the advantage of the estimated parameters being scale-invariant with respect to the frequency of data collection, but it also brings a computational cost, especially when fitting the model with continuous capture process, and when time-dependent variables are included. To optimize computations, we have implemented the computational most intensive functions in C++, while keeping a user-friendly R-interface through the Rcpp package (Eddelbuettel & François, 2011). Further substantial optimization could be possibly obtained e.g. using GPU computation or different parallelization techniques.

A key applied aim of ecological research is to gain understanding about how individuals, populations, communities, and ecosystems respond to the ongoing global changes such as land use change and climate change. One important aspect of such responses is the ability of individuals to move across the changing landscapes, e.g., to reach new areas where climate will be suitable for them in the future. We hope that the software developed here enables researchers and practitioners to gain a better understanding of the role of movement in spatial population dynamics, both in basic research as well as in applications in conservation and management.

## Supporting information

S1. Mathematical and technical information about the JSMM approach and its implementation into the R-package Jsmm.

S2. Vignettes demonstrating the application of the R-package Jsmm.

S3. Additional information on the moths case study used as an example in this paper.

## Supporting information

S1 Mathematical and technical information about the JSMM approach and its implementation into the R-package Jsmm

S2 Vignettes demonstrating the application of the R-package Jsmm

S3 Additional information on the moths case study

## Acknowledgments

LFR was funded by the University of Helsinki-HIIT PhD Fellowship on multidisciplinary applications of AI. OO was funded by the Research Council of Finland (grants no. 336212 and 345110).

## Authors’ contributions

LFR and OO jointly designed the Jsmm 1.0 software. LFR implemented the simulated and real data case studies and prepared the vignettes. LFR and OO jointly prepared the manuscript.

**AI tools:** No AI tools were used to produce the present manuscript, Jsmm software nor documentation.

## Data Availability Statement

The R-package Jsmm as well as the data and scripts needed to reproduce the analyses presented in this paper and the vignettes are available in [GitHub](https://github.com/lufrodriguezca/Jsmm).

## Notes

https://github.com/lufrodriguezca/Jsmm

### Competing Interest Statement

The authors have declared no competing interest.

### Summary of Updates

1. In supplementary material S1, a minor notation on page six has been corrected: K and M represent stiffness and mass matrix, respectively. 2. In supplementary material S2 (vignette book), deprecated function simulate_movement() was replaced with function simulate_captures().

## References

Alagador, D., & Cerdeira, J. O. (2022). Operations research applicability in spatial conservation planning. Journal of Environmental Management, 315, 115172. 10.1016/j.jenvman.2022.115172

Betzholtz, P.-E., Forsman, A., & Franzén, M. (2019). Inter-individual variation in colour patterns in noctuid moths characterizes long-distance dispersers and agricultural pests. Journal of Applied Entomology, 143 (9), 992–999. 10.1111/jen.12670

Eddelbuettel, D., & François, R. (2011). Rcpp: Seamless R and C++ Integration. Journal of Statistical Software, 40 (8), 1–18. 10.18637/jss.v040.i08

Efford, M. G., & Schofield, M. R. (2022). A review of movement models in open population capture–recapture. Methods in Ecology and Evolution, 13 (10), 2106–2118. 10.1111/2041-210X.13947

Elsner, M., Atkinson, G., & Zahidi, S. (2025). The Global Risks Report. World Economic Forum, 2025.

Florko, K. R., Togunov, R. R., Gryba, R., Sidrow, E., Ferguson, S. H., & Yurkowski,D. (2025). An introduction to statistical models used to characterize species-habitat associations with animal movement data. Movement Ecology, 13 (1), 27. 10.1186/s40462-025-00549-2

Forrest, S. W., Pagendam, D., Bode, M., Drovandi, C., Potts, J. R., Perry, J., Vanderduys, E., & Hoskins, A. J. (2025). Predicting fine-scale distributions and emergent spatiotemporal patterns from temporally dynamic step selection simulations. Ecography, 2025 (2), e07421. 10.1111/ecog.07421

Gray, R., Rodriguez, L., Lewis, O., Chung, A., Ovaskainen, O., & Slade, E. (2022). Movement of forest-dependent dung beetles through riparian buffers in Bornean oil palm plantations. Journal of Applied Ecology, 59 (1), 238–250. 10.1111/1365-2664.14049

Heinrich, B. (2012). The Hot-Blooded Insects (1st). Springer Berlin, Heidelberg. 10.1007/978-3-662-10340-1

Lagrange, P., Pradel, R., Bélisle, M., & Gimenez, O. (2014). Estimating dispersal among numerous sites using capture–recapture data. Ecology, 95 (8), 2316–2323. 10.1890/13-1564.1

McClintock, B. T., Abrahms, B., Chandler, R. B., Conn, P. B., Converse, S. J., Emmet,R. L., Gardner, B., Hostetter, N. J., & Johnson, D. S. (2022). An integrated path for spatial capture–recapture and animal movement modeling. Ecology, 103 (10), e3473. 10.1002/ecy.3473

Nathan, R. (2008). An emerging movement ecology paradigm. Proceedings of the National Academy of Sciences, 105 (49), 19050–19051. 10.1073/pnas.0808918105

Neil Arnason, A. (1973). The estimation of population size, migration rates and survival in a stratified population. Population Ecology, 15 (2), 1–8. 10.1007/BF02510705

Ovaskainen, O. (2004). Habitat-specific movement parameters estimated using mark-recapture data and a diffusion model. Ecology, 85 (1), 242–257. 10.1890/02-0706

Ovaskainen, O. (2008). Analytical and numerical tools for diffusion-based movement models. Theoretical Population Biology, 73 (2), 198–211. 1016/j.tpb.2007.11.002

Ovaskainen, O., Luoto, M., Ikonen, I., Rekola, H., Meyke, E., & Kuussaari, M. (2008). An Empirical Test of a Diffusion Model: Predicting Clouded Apollo Movements in a Novel Environment. The American Naturalist, 171 (5), 610–619. 10.1086/587070

Ovaskainen, O., Ramos, D. L., Slade, E. M., Merckx, T., Tikhonov, G., Pennanen, J., Pizo, M. A., Ribeiro, M. C., & Morales, J. M. (2019). Joint species movement modeling: How do traits influence movements? Ecology, 100 (4), 1–8. 10.1002/ecy.2622

Plummer, M., Best, N., Cowles, K., & Vines, K. (2006). CODA: Convergence Diagnosis and Output Analysis for MCMC. R News, 6 (1), 7–11. https://www.r-project.org/doc/Rnews/Rnews_2006-1.pdf

Quaglietta, L., & Porto, M. (2019). SiMRiv: An R package for mechanistic simulation of individual, spatially-explicit multistate movements in rivers, heterogeneous and homogeneous spaces incorporating landscape bias. Movement Ecology, 7:11. 10.1186/s40462-019-0154-8

R Core Team. (2024). R: A Language and Environment for Statistical Computing. R Foundation for Statistical Computing. https://www.R-project.org/

Royle, J. A., Chandler, R. B., Sollmann, R., & Gardner, B. (2014). Spatial Capture-recapture. Elsevier. 10.1016/C2012-0-01222-7

Royle, J. A., & Young, K. V. (2008). A hierarchical model for spatial capture-recapture data. Ecology, 89 (8), 2281–2289. 10.1890/07-0601.1

Schultz, C. B., Haddad, N. M., Henry, E. H., & Crone, E. E. (2019). Movement and Demography of At-Risk Butterflies: Building Blocks for Conservation. Annual Review of Entomology, 64, 167–184. 10.1146/annurev-ento-011118-112204

Slade, E. M., Merckx, T., Riutta, T., Bebber, D. P., Redhead, D., & Riordan, P. (2013). Life-history traits and landscape characteristics predict macro-moth responses to forest fragmentation. Ecology, 94 (7), 1519–1530. 10.1890/12-1366.1

Stroustrup, B. (2013). The C++ Programming Language (4th). Addison-Wesley Professional.

Tourani, M. (2022). A review of spatial capture–recapture: Ecological insights, limitations, and prospects. Ecology and Evolution, 12 (1), e8468. 10.1002/ece3.8468

White, G. C. (2008). Closed population estimation models and their extensions in Program MARK. Environmental and Ecological Statistics, 15 (1), 89–99. 10.1007/s10651-007-0030-3

Williams, B. K., Nichols, J., & Conroy, M. (2002). Analysis and Management of Animal Populations: Modeling, Estimation and Decision Making. Academic Press. https://pubs.usgs.gov/publication/5200256

