## Supplementary material for "R-package Jsmm: Joint species movement modelling of mark-recapture data": S1 Mathematical and technical information about the JSMM approach and its implementation into the R-package Jsmm

Supporting Information S1:

### Mathematical and technical information about the JSMM approach and its implementation into the R-package Jsmm

Luisa F. Rodriguez and Otso Ovaskainen, 2026. Joint Species Movement Modelling with the R-package Jsmm. Preprint available at *biorxiv*.

The Jsmm implementation and the scripts for running analyses can be found in GitHub.

License CC BY-NC-ND 4.0. Rodriguez and Ovaskainen. All rights reserved, including those for text and data mining, AI training, and similar technologies.

#### 1 Mathematical Framework

We use the diffusion-advection-reaction modelling framework to model the movement behavior of individuals in a spatial domain $\Omega$ over a time period $[T_1, T_2]$. While our implementation of the software is specific to two-dimensional domains, we present here the mathematical framework more generally for $d$-dimensional domains.

We consider each individual separately and thus do not account for how the individuals might influence each other's movements. We denote by $u(\mathbf{x}, t)$ the probability density of the individual location ($\mathbf{x} \in \mathbb{R}^d$) at time $t$. The probability density function $u$ is a scalar field $u : \Omega \times [T_1, T_2] \to \mathbb{R}$. The integral of the probability density $u(\mathbf{x}, t)$ over $\Omega$ gives the probability by which the individual is still active (not dead nor trapped) at time $t$. The elliptic partial operator $\mathcal{L}$ that defines the diffusion-advection-reaction model is given by

$$\mathcal{L}u(\mathbf{x}, t) = \sum_{i,j=1}^{d} \partial_{ij}[a_{ij}(\mathbf{x}, t)u(\mathbf{x}, t)] - \sum_{i=1}^{d} \partial_i[b_i(\mathbf{x}, t)u(\mathbf{x}, t)] - (m(\mathbf{x}, t) + r(\mathbf{x}, t))u(\mathbf{x}, t). \quad (1)$$

The diffusion part of the model ($a$) models the random component of the movement, the advection part ($b$) a deterministic tendency to move to a given direction, and the reaction part mortality ($m$) and capture rates ($r$). The diffusion part is the only mandatory part of the model, and thus both advection and mortality can be omitted if they are not relevant to the case study. The capture rate is included if the sampling methodology is based on a continuous capture process, but not if it is based on an instantaneous capture process. We thus, assume that the diffusion rate may vary over both space and time. A key feature of the modelling framework (1) is that all the movement parameters ($a$, $b$, $m$, and $r$) are allowed to depend on space, time, and space-time. As described below, we will model such dependency through a generalized linear modelling approach.

The diffusion tensor is a symmetric and positive-definite matrix defined as

$$a(\mathbf{x}, t) = (a_{ij}(\mathbf{x}, t)), \quad (2)$$

where $a_{ij}$ is a scalar field $a_{ij} : \Omega \times [T_1, T_2] \to \mathbb{R}^+$, $i, j \in 1, \ldots, d$. While it would be possible to extend our framework to model also anisotropic diffusion, the current implementation is restricted to isotropic diffusion, where $a_{ii}(\mathbf{x}, t) = a(\mathbf{x}, t)$ and $a_{ij}(\mathbf{x}, t) = 0$ for $j \neq i$. In other words, we assume that with the random movement component, the individuals move equally in all directions.

The advection vector is defined as

$$b(\mathbf{x}, t) = [b_1(\mathbf{x}, t), \ldots, b_d(\mathbf{x}, t)], \tag{3}$$

where each component $b_i : \Omega \times [T_1, T_2] \to \mathbb{R}$, $i \in \{1, \ldots, d\}$ describes how fast the individual is drifted to the negative direction of the coordinate axis.

The mortality parameter $m(\mathbf{x}, t) : \Omega \times [0, T] \to \mathbb{R}^+$ describes the rate at which the individual dies so that the probability of the individual dying over a short time interval $dt$ is $m(\mathbf{x}, t)dt$. Similarly, the capture parameter $r(\mathbf{x}, t) : \Omega \times [T_1, T_2] \to \mathbb{R}^+$ describes the rate at which the individual is captured by some trap that is located in its neighborhood so that the probability of the individual being trapped within a subdomain $\Omega_c$ over a short time interval $dt$ is $\int_{\Omega_c} r(\mathbf{x}, t) dt d\mathbf{x}$.

In the diffusion model, the probability density of an individual location evolves according to the partial differential equation

$$\partial_t u(\mathbf{x}, t) = \mathcal{L} u(\mathbf{x}, t). \tag{4}$$

Additionally, the modelling framework accounts for habitat preference, which we denote for location $\mathbf{x}$ and time $t$ by $k(\mathbf{x}, t)$ (Ovaskainen & Cornell, 2003; Ovaskainen & Crone, 2009). Habitat preference may vary continuously over space (e.g. due to preference for certain altitude) or discontinuously over space (e.g. due to preference for discretely varying habitat types). The probability density will be locally proportional to the habitat preference. Discontinuously varying habitat preference varying corresponds to edge-mediated behavior at internal edges, e.g., borders between habitat types. The discontinuous probability density at a point $\mathbf{x}'$ located at the common boundary $\Gamma_{ij}$ of two subdomains $\Omega_i$, and $\Omega_j$, for a given time $t'$, can be expressed as

$$k_j(t') \lim_{\mathbf{x} \in \Omega_i \to \mathbf{x}'} u(\mathbf{x}', t') = k_i(t') \lim_{\mathbf{x} \in \Omega_j \to \mathbf{x}'} u(\mathbf{x}', t') \tag{5}$$

where $k_i$ and $k_j$ are the habitat preferences in the two subdomains.

#### 1.1 Capture processes

We consider two kinds of capture processes: the instantaneous capture process (ICP) and the continuous capture process (CCP).

In the case of the ICP, we assume short-term capture events at particular study sites, during which the researcher tries to detect the marked individuals. Each capture event $c$ is described in terms of its location (the capture site $\Omega_c$) and time $t_c$. We assume that, if a marked individual is present in the capture site $\Omega_c$ at the time of the event, it is captured by probability $p_c$. We note that in the case of the ICP, the capture rate term $r$ is not included in the model (1).

In the case of the CCP, the capture process is included in the modelling framework (1) through the rate parameter $r$. In this case, each capture event $c$ is described in terms of its location (the capture site $\Omega_c$) and a time-interval $[t_{c,1}, t_{c,2}]$ during which the trap is open or active. Within each capture event $c$, we assume that the capture rate parameter does not vary spatially over the capture area, but that it can vary over time: $r(\mathbf{x}, t) = r_c(t)$ for $\mathbf{x} \in \Omega_c$ and $t \in [t_{c,1}, t_{c,2}]$. For locations and times that do not correspond to any capture event, the capture rate is set to zero, $r(\mathbf{x}, t) = 0$.

#### 1.2 Simulating capture data

We consider one individual and assume that it is released in a particular release event described in terms of its location (the release site $\Omega_r$) and time $t_r$. The diffusion-advection-reaction model is initiated at time $t_r$ by assuming a constant probability density $u = 1/|\Omega_r|$ within $\Omega_r$, and zero probability density outside $\Omega_r$. How the simulation is performed depends on whether the ICP or the CCP capture process is assumed.

In the case of the ICP, the probability density is evolved according to (4) until the next time that a capture event takes place. Then the location of the individual is sampled according to the probability density, one minus the total integral giving the probability that the individual has died. If the individual is still alive, the probability density is initiated according to the delta distribution centered at the current location. If the current time involves a capture event, it is checked if the location of the individual is within a capture site $\Omega_c$. If not, capture cannot take place. If yes, whether the individual is captured is randomized from the Bernoulli distribution with capture probability $p_c$. If the randomization of the Bernoulli distribution results in the individual being captured, the capture is added to the observed data. The modelling approach allows for study designs where individuals that have been captured are released again, and for study designs where the captured individuals are removed from the system. If the capture event includes the re-release of captured individuals, then the evolution of the probability density $u$ is continued by (4) until the next sampling time.

In the case of the CCP, the trajectories of the individuals are simulated using the same approach as for the ICP. To simulate captures, we note that the probability $p_c$ by which the individual is trapped during the time interval $[t_1, t_2]$ is given by (Gray et al., 2022)

$$p_c = \int_{t_1}^{t_2} r_c(t) \int_{\Omega_c} u(\mathbf{x}, t) d\mathbf{x} dt, \tag{6}$$

where $r_c(t)$ is the capture rate. In the numerical implementation, the model is evaluated over short time intervals $[t_1, t_2]$, with one capture event extending possibly over multiple such time intervals. After each time interval, the Bernoulli distribution is applied to determine if the individual is trapped or not. If the individual is trapped, it stays captured until the end of the capture period. If the individual is not trapped, the probability density for its location is updated as $u(\mathbf{x}, t_2) = \tilde{u}(\mathbf{x}, t_2)/(1 - p)$, where $\tilde{u}$ is the probability density before the Bernoulli randomization. $u(\mathbf{x}, t_2)$ determines if the individual is actually trapped. If the individual becomes trapped at any time during the capture event, in the end of the capture event the identity of the capture event is added to the capture history, and the individual is possibly released again (as in the case of ICP).

#### 1.3 Computing likelihood of data

We next describe how to compute the likelihood of some observed data under the model. We consider one individual and assume that it is released in a particular release event described in terms of its location (the release site $\Omega_r$) and time $t_r$ and that it is either captured in some capture event $c$ that takes place after the release, or that it is never again observed in the study. The more general likelihood related to multiple individuals and multiple recaptures of the same individual can then be easily constructed from this building block.

The diffusion-advection-reaction model is initiated at time $t_r$ by assuming a constant probability density $u = 1/|\Omega_r|$ within $\Omega_r$, and zero probability density outside $\Omega_r$. At this point, the log-likelihood of the observed data is initialized with $L = 0$. The probability density is then evolved according to (4) until the time of the first capture event that takes place after the release time $t_r$. At this point, the procedure of computing the likelihood of the data depends on whether the ICP or the CCP capture process is assumed.

In the case of the ICP, the evolution of the probability density $u$ (4) is interrupted at the time $t_c$ when instantaneous capture is attempted at a certain capture site $\Omega_c$. The probability by which the individual is present at the site of the capture event is given by

$$U_c = \int_{\Omega_c} u(\mathbf{x}, t_c) d\mathbf{x}. \tag{7}$$

Denoting by $p_c$ the capture probability (probability of capturing an individual conditional on it being present) for event $c$, the probability by which the individual is present and captured is then $q_c = U_c p_c$. If the individual was captured in the event $c$, the term $\log(q_c)$ is added to the log-likelihood $L$, and the computation of the log-likelihood is complete. If the individual was not captured in the event $c$, the term $\log(1 - q_c)$ is added to the log-likelihood $L$. If the event $c$ was the last capture event, the computation of the log-likelihood is complete. If not, the probability density for the individual location is updated as described in the section on generating simulated data, after which the diffusion model is evolved until the next capture event.

In the case of the CCP, the evolution of the probability density $u$ (4) is interrupted at the time $t_{c,1}$ when instantaneous capture is initiated at any capture site $\Omega_c$. At this point, the cumulative capture probability for that trapping event $c$ is initiated as $q_c = 0$, and the trapping event is opened by including a positive rate $r(\mathbf{x}, t)$ in the diffusion equation. The time evolution of the probability density $u$ (4) is then continued, and at the same time, the probability of the individual being trapped ($q_c$) is updated with (6). This is continued until some capture event starts, or any trapping event is closed, i.e. the time $t_{c,2}$ is reached for some $c$. If a trapping event starts, that trap is opened and its capture probability is initiated to zero as described above. If a trapping event is closed, then the probability by which the trapping event should have captured the individuals is given by $q_c$, which has been computed while evolving the diffusion equation. If the individual was captured in the event $c$, the term $\log(q_c)$ is added to the log-likelihood $L$, and the computation of the log-likelihood is complete. If the individual was not captured in the event $c$, the term $\log(1-q_c)$ is added to the log-likelihood $L$. If the event $c$ was the last capture event, the computation of the log-likelihood is complete. If not, the individual is either dead, has been captured by some trap $l$ that was open at time $t_{c,2}$ but has not been closed yet, or is still alive and moving. Thus, if the individual was not observed, the probability density $u(\mathbf{x}, t)$ for the individual's location needs to be updated, as well the probability $P_D$ by which the individual is dead, as well as the probabilities $q_l$ by which the individual would have been trapped in some other capture event than the focal capture event $c$. Each of these can be updated by dividing them by $1 - q_c$.

#### 1.4 Model parametrization through generalized linear modelling and spatiotemporal predictors

The parameters of the diffusion-advection-mortality model (1) depend on space ($\mathbf{x}$) and time ($t$). We model such spatial and temporal variation through the dependency of these parameters on some measured predictors using a generalized linear modelling framework.

As the diffusion parameter $a(\mathbf{x}, t)$ is restricted to positive values, we model it assuming the log link-function. We thus assume that

$$\log(a(\mathbf{x}, t)) = \sum_k w_k^a(\mathbf{x}, t)\beta_k^a, \tag{8}$$

where $w_k^a(\mathbf{x}, t)$ is the value of the $k$:th predictor at location $\mathbf{x}$ and time $t$, and $\beta_k^a$ describes how the diffusion rate depends on this predictor. We include the intercept in the model by setting the first predictor to one for all locations and times, $w_1^a(\mathbf{x}, t) = 1$. Interactions among different predictors, as well as non-linear effects of predictors, can be included as usual in the generalized linear modelling framework.

As the advection parameter $b_i(\mathbf{x}, t)$ can obtain any real value, we model it assuming the identity link-function. We thus assume that

$$b_i(\mathbf{x}, t) = \sum_k w_k^b(\mathbf{x}, t)\beta_{k,i}^b, \tag{9}$$

where $w_k^b(\mathbf{x}, t)$ is the value of the $k$:th predictor at location $\mathbf{x}$ and time $t$, and $\beta_{k,i}^b$ describes how the advection to the direction of the $i$ coordinate depends on this predictor.

As the mortality parameter $m(\mathbf{x},t)$ is restricted to positive values, we model it assuming the log link-function. We thus assume that

$$\log(m(\mathbf{x},t)) = \sum_k w_k^m(\mathbf{x},t)\beta_k^m, \tag{10}$$

where $w_k^m(\mathbf{x},t)$ is the value of the $k$:th predictor at location $\mathbf{x}$ and time $t$, and $\beta_k^m$ describes how the mortality rate depends on this predictor.

As described above, we assume that the capture probability parameter $p$ (in the case of the ICP) has event-specific values, $p = p_c$, whereas the capture rate parameter $r$ (in the case of CCP) can vary temporally within each event, $r = r_c(t)$.

We assume that each capture event is described in terms of some predictors related to the observation process, denoted for predictor $k$ and capture event $c$ by $w_{k,c}^o$ for ICP and by $w_{k,c}^o(t)$ for CCP. As the capture rate parameter $r$ is restricted to positive values, we model it assuming the log link-function,

$$\log r_c(t) = \sum_k w_{k,c}^o(t)\beta_k^o, \tag{11}$$

where $\beta_k^o$ describes how the capture rate depend on the predictor $w_{k,c}$.

As the capture probability parameter $p$ is restricted to values between 0 and 1, we model it assuming the logit link-function,

$$\log(p_c/(1-p_c)) = \sum_k w_{k,c}^o\beta_k^o, \tag{12}$$

where $\beta_k^o$ describes how the capture probability depend on the predictor $w_{k,c}$.

### 2 Numerical methods

#### 2.1 Solving the diffusion model with the finite element method

To approximate numerically the solution (4), we apply the finite element method (FEM), which is based on the variational or weak formulation of the model. Following Ovaskainen (2008), we write the variational formulation of Eq. (4) as

$$\int_\Omega \frac{\partial u(\mathbf{x},t)}{\partial t} v(\mathbf{x},t)d\mathbf{x} = \int_\Omega \mathcal{M}(u(\mathbf{x},t), v(\mathbf{x},t))d\mathbf{x} \tag{13}$$

where $v(\mathbf{x},t)$ is an arbitrary test function, and the bilinear form $\mathcal{M}$ is defined as

$$\begin{aligned}\mathcal{M}(u(\mathbf{x},t), v(\mathbf{x},t)) = &-\Sigma_{i,j}\partial_j\left[a(\mathbf{x},t)u(\mathbf{x},t)\right]\partial_i v(\mathbf{x},t)\\ &-\Sigma_i b_i(\mathbf{x},t)u(\mathbf{x},t)\partial_i v(\mathbf{x},t)\\ &-c(\mathbf{x},t)u(\mathbf{x},t)v(\mathbf{x},t)\\ &-r(\mathbf{x},t)u(\mathbf{x},t)v(\mathbf{x},t).\end{aligned} \tag{14}$$

FEM requires a domain discretization of the domain with respect to space and the study period over time. We discretize the spatial domain into triangular elements $\mathcal{T}_e$, which we index by $e = 1, \ldots, n_e$. We index the nodes of the triangles by $l = 1, \ldots, n_l$, and denote their coordinates by $\mathbf{x}_j$. We let $\tilde{N}_j(\mathbf{x})$ denote the set of basis functions, which are defined so that they are piecewise linear, $\tilde{N}_j(\mathbf{x}_j) = 1$ and $\tilde{N}_j(\mathbf{x}_i) = 0$ for $i \neq j$.

To account for the discontinuities through the interior boundaries, we let $N_j(\mathbf{x},t) = k(\mathbf{x},t)\tilde{N}_j(\mathbf{x})$ denote modified shape functions. We denote the approximate solution by

$$\hat{u}(\mathbf{x},t) = \Sigma_{j=1}^{m} u_j(t) N_j(\mathbf{x},t),$$

where $\boldsymbol{u}(t) = \{u_j(t)\}$ is the coefficient vector.

Over a time-step of length $\Delta t$ from $t_1$ to $t_2 = t_1 + \Delta t$, the diffusion model can be numerically solved as Ovaskainen, 2008

$$\boldsymbol{u}(t_2) = D_1^{-1} D_2 \boldsymbol{u}(t_1), \tag{15}$$

where

$$\begin{aligned} D_1 &= M^T - \theta \Delta t K^T, \\ D_2 &= M^T + (1-\theta)\Delta t K^T, \end{aligned} \tag{16}$$

and where $K$ and $M$ denote the stiffness and mass matrices, respectively

$$\begin{aligned} M_{ij} &= \int_\Omega N_i(\mathbf{x},t_1)\tilde{N}_j(\mathbf{x})d\mathbf{x}, \\ K_{ij} &= \int_\Omega \mathcal{M}(N_i(\mathbf{x},t_1),\tilde{N}_j(\mathbf{x}))d\mathbf{x}. \end{aligned} \tag{17}$$

#### 2.2 Representing spatiotemporal covariates through discretizations

As described above, the model parametrization involves estimated responses to spatiotemporally varying covariates, which covariates are denoted by $w_k^a(\mathbf{x},t)$ for diffusion, by $w_k^b(\mathbf{x},t)$ for advection, by $w_k^m(\mathbf{x},t)$ for mortality, by $w_k^o(t)$ for CCP-related capture processes, and by $w_k^o$ for ICP-related capture processes.

For numerical solving, the covariates need to be compatible with the triangular discretization applied in the finite element method. Thus we enable the user to define the covariates either through the nodes or the elements. To define a temporally or spatiotemporally variant covariant, a discretization over time is needed. We denote such a discretization by $t_1, \ldots t_n$, where the first and the last time points must coincide with the beginning and end of the study period $t_1 = T_1$, $t_n = T_2$.

- **Spatial covariates defined through nodes can be used for diffusion, advection, and mortality**. We denote by $w_{k,l}$ the value of the covariate $k$ for node $l$, and assume that the covariate is time invariant and varies over space in a piecewise linear manner, $w_k(\mathbf{x},t) = \Sigma_{j=1}^{m} w_{k,l}\tilde{N}_j(\mathbf{x})$.
- **Spatial covariates defined through elements can be used for habitat preference, diffusion, advection, and mortality**. We denote by $w_{k,e}$ the value of the covariate $k$ for element $e$, and assume that the covariate is invariant of time and is constant within each element, $w_k(\mathbf{x},t) = w_{k,e}$ for $\mathbf{x} \in \mathcal{T}_e$.
- **Spatially invariant temporal covariates can be used for diffusion, advection, and mortality.** We denote by $w_{k,j}$ the value of the covariate $k$ for time point $t_j$ of the discretization, and assume that the covariate is invariant of space and varies over time in a piecewise linear manner.
- **Spatiotemporal covariates defined through nodes can be used for diffusion, advection, and mortality**. We denote by $w_{k,l,j}$ the value of the covariate $k$ for node $l$ at time point $t_j$, and assume that the covariate varies over space in a piecewise linear manner, $w_k(\mathbf{x},t_j) = \Sigma_{j=1}^{m} w_{k,l,j}\tilde{N}_j(\mathbf{x})$, and that it varies over time (within each time period of the temporal discretization) in a piecewise linear manner.

- **Spatiotemporal covariates defined through elements can be used for habitat preference, diffusion, advection, and mortality.** We denote by $w_{k,e,j}$ the value of the covariate $k$ for element $e$ at time point $t_j$, and assume that at each time point the covariate is constant within each element, $w_k(\mathbf{x}, t_j) = w_{k,e,j}$ for $\mathbf{x} \in \mathcal{T}_e$, and that it varies over time (within each time period of the temporal discretization) in a piecewise linear manner.

- **Capture event-specific covariates can be used for ICP and CCP.** We denote by $w_{k,c}$ the value of the covariate for capture event $c$.

- **Temporally varying capture event-specific covariates can be used for CCP.** To define a temporally covariant covariate for capture event $c$, a discretization of time over the trapping period is needed. We denote such a discretization for event $c$ by $t_1, \ldots t_n$, where the first and the last time points must coincide with the beginning and end of the trapping period, $t_1 = t_{c,1}$, $t_n = t_{c,2}$. We denote by $w_{k,c,j}$ the value of the covariate $k$ for time point $t_j$ of trapping event $c$, and assume that the covariate varies over time in a piecewise linear manner.

### 3 Prior distributions of JSMM

Concerning the prior distributions, we assume the same functional forms as in Ovaskainen et al., 2019:

$$\theta_i \sim N(\mathbf{m}, \mathbf{\Sigma} \otimes [\rho \mathbf{C} + (1-\rho)\mathbf{I}_{n_i}])$$

where the likelihood of the movement data, depends on the species $i$ parameters $\theta_i$. The mean $\mathbf{m} = vec(\mathbf{TZ})$ is a vectorized form of the matrix $\mathbf{M} = \mathbf{TZ}$, where the matrix $\mathbf{T}$ consists of the elements $t_{ik}$. We denote by $\mathbf{z} = vec(\mathbf{Z})$ the vectorization of the matrix $\mathbf{Z}$.

The parameters for which prior distributions need to be defined are $\mathbf{z}$, $\mathbf{\Sigma}$ and $\rho$.

- For $\mathbf{z}$ vector it is assumed a multinormal prior $\mathbf{z} \sim N(\mu_z, \mathbf{\Sigma}_z)$
- For $\mathbf{\Sigma}$ was assumed an Inverse-Wishart prior $W^{-1}(\mathbf{\Psi}, \nu)$, where $\nu = n_p$.
- $\rho$, discrete prior. Probability of 0.5 for $\rho = 0$, and the remaining probability of 0.5 uniformly to the range $(0, 1]$, discretized to 100 values.

While the user can modify the parameters of the prior distributions, we have attempted to implement default values that would be applicable for typical case studies. These have been selected with the following criteria:

- *Mortality.* We set the prior mean mortality rate to a value that leads to mortality probability of 0.5 during the study period. This value was selected as the default prior as we assume that often mark-recapture studies span over a time period that is relevant for estimating mortality rate.

- *Diffusion.* We set the prior mean diffusion rate to a value that makes expected lifetime displacement of the individual equal to half of the diameter of the landscape domain. This value was selected as the default prior as we assume that often mark-recapture studies are conducted over a spatial domain the size of which is relevant for estimating lifetime movement distances.

- *Advection.* We set the prior mean advection rate to a value that makes expected lifetime displacement of the individual equal to half of the diameter of the landscape domain. This value was selected as the default prior as we assume that often mark-recapture studies are conducted over a spatial domain the size of which is relevant for estimating lifetime movement distances, and that in case advection is included in the model, we assume that it will have a substantial influence on movements.

- *Capture.* In case of ICP, we set the prior mean capture probability to a value that makes capture probability (conditional on the presence of the individual) to 0.5. In case of CCP, we set the prior mean capture rate to a value that makes capture probability (conditional on the presence of the individual) over average-length capture interval to 0.5. These values were selected as the default prior as we assume that often mark-recapture studies have a capture process that leads to substantial yet not complete success of capture.
- *Covariate effects.* The default prior for covariate effects is compatible with covariates that are scaled to zero mean and unit variance, and hence we recommend the user to scale the covariates before the analyses. We set the prior mean of all covariate effects to zero, which corresponds to the assumption that we do not know if they will have positive or negative effects. We set their prior variance to one, which corresponds to the assumption that the covariates can have substantial effects on the movement and capture parameters.

### 4 Notation

| | |
|---|---|
| $\Omega$ | Spatial domain. |
| $[T_1, T_2]$ | Time period. |
| $u(\mathbf{x}, t)$ | Probability density function of an individual located ($\mathbf{x} \in \mathbb{R}^d$) at time $t$. |
| $\Gamma_{ij}$ | Common boundary between the subdomains $\Omega_i$ and $\Omega_j$. |
| $\mathcal{L}u$ | Elliptic partial operator that defines the diffusion-advection-reaction model. |
| $a(\mathbf{x}, t)$ | Diffusion tensor. |
| $b(\mathbf{x}, t)$ | Advection vector. |
| $m(\mathbf{x}, t)$ | Mortality rate. |
| $k(\mathbf{x}, t)$ | Habitat preference. |
| $r(\mathbf{x}, t)$ | Capture rate parameter. |
| ICP | Instantaneous capture process. |
| CCP | Continuous capture process. |
| $r$ | Release event. |
| $\Omega_r$ | Release site corresponding to the release event $r$. |
| $t_r$ | Release time corresponding to the release event $r$. |
| $c$ | Capture event. |
| $\Omega_c$ | Capture site corresponding to the capture event $c$. |
| $t_c$ | Capture time corresponding to the capture event $c$. |
| $U_c$ | Probability by which an individual is present for the capture event $c$. |
| $p_c$ | Capture probability for the event $c$ |
| $q_c$ | Probability individual present and captured for the event $c$ |
| $[t_{c,1}, t_{c,2}]$ | Time interval in which a capture event $c$ span. |
| $w_k^a(\mathbf{x}, t)$ | Value of the $k$:th predictor at location $\mathbf{x}$ at time t for diffusion |
| $\beta_k^a$ | Diffusion dependence on $k$:th predictor. |
| $w_k^b(\mathbf{x}, t)$ | Value of the $k$:th predictor at location $\mathbf{x}$ at time t for advection |
| $\beta_{k,i}^b$ | Advection to the direction of the $i$ coordinate dependence on $k$:th predictor. |
| $w_k^m(\mathbf{x}, t)$ | Value of the $k$:th predictor at location $\mathbf{x}$ at time t for mortality |
| $\beta_k^m$ | Mortality rate dependence on $k$:th predictor. |
| $w_{k,c}^o$ | Value of the $k$:th predictor for capture probability parameter (ICP) |
| $w_{k,c}^o(t)$ | Value of the $k$:th predictor at time $t$ for capture rate parameter (CCP) |
| $\beta_k^o$ | Capture dependence on $k$:th predictor. |
| $\mathcal{M}$ | Bilinear form. |
| $v$ | Test Function. |
| FEM | Finite Element Method. |
| $\mathcal{T}_e$ | Set of non overlapping triangles in $\Omega$. |
| $\tilde{N}_j(\mathbf{x})$ | Set of basis functions |
| $N_j = k(\mathbf{x}, t)\tilde{N}_j(\mathbf{x})$ | Modified basis functions |

| | |
|---|---|
| $M$ | Mass matrix FEM. |
| $K$ | Stiffness matrix FEM. |
| $\otimes$ | Kronecker (outer) product. |
| $\theta_i$ | Vector of model parameters for the species $i$. |
| $t_{ik}$ | Trait $k$ of the species $i$. |
| $\mathbf{T}$ | Matrix that consists of elements $t_{ik}$. |
| $\zeta_{kp}$ | Parameter that measures the influence of trait on movement parameter $p$. |
| $\mathbf{Z}$ | Matrix with the regression parameters $\zeta_{kp}$. |
| $\rho$ | Parameter that measures the strength of the phylogenetic signal. |
| $\mathbf{C}$ | Phylogenetic correlation matrix. |
| $\mathbf{I}_{n_i}$ | Identity matrix of dimension $n_i$. |
| $\mathbf{Z}$ | Matrix with the parameters $\zeta_{kp}$. |
| $\mathbf{z}$ | Vector form of the matrix $\mathbf{Z}$. |
| $\mu_z$ | Mean parameter for the Multivariate normal distribution. |
| $\mathbf{\Sigma}_z$ | Variance-covariance matrix for the multivariate normal distribution. |
| $\mathbf{\Sigma}$ | Covariance matrix that describes the dependence of the model parameters $\theta_i$. |
| $\mathbf{\Psi}$ | Scale matrix for the Inverse-Wishart distribution. |
| $\nu$ | Degrees of freedom for the Inverse-Wishart distribution. |
| $n_p$ | Number of parameters. |
| $\mathbf{m}$ | Mean vector of the multivariate normal distribution. |

### References


Gray, R. E. J., Rodriguez, L. F., Lewis, O. T., Chung, A. Y. C., Ovaskainen, O., & Slade, E. M. (2022). Movement of forest-dependent dung beetles through riparian buffers in Bornean oil palm plantations. *Journal of Applied Ecology*, *59*(1), 238–250. https://doi.org/10.1111/1365-2664.14049

Ovaskainen, O. (2008). Analytical and numerical tools for diffusion-based movement models. *Theoretical Population Biology*, *73*(2), 198–211. https://doi.org/10.1016/j.tpb.2007.11.002

Ovaskainen, O., & Cornell, S. J. (2003). Biased movement at a boundary and conditional occupancy times for diffusion processes. *Journal of Applied Probability*, *40*(3), 557–580. http://www.jstor.org/stable/3215936

Ovaskainen, O., & Crone, E. E. (2009). Modeling animal movement with diffusion. In S. A. Cantrell, C. Cosner, & S. Ruan (Eds.), *Spatial ecology* (pp. 63–83). Chapman; Hall/CRC.

Ovaskainen, O., Ramos, D. L., Slade, E. M., Merckx, T., Tikhonov, G., Pennanen, J., Pizo, M. A., Ribeiro, M. C., & Morales, J. M. (2019). Joint species movement modeling: How do traits influence movements. *Ecology*, *100*(4), e02622. https://doi.org/10.1002/ecy.2622
