## Supplementary material for "R-package Jsmm: Joint species movement modelling of mark-recapture data": S2 Vignettes demonstrating the application of the R-package Jsmm: README.pdf

Luisa F. Rodriguez and Otso Ovaskainen, 2026. Joint Species Movement Modelling with the R-package Jsmm. Preprint available at *biorxiv*.

The Jsmm implementation and the scripts for running analyses can be found in GitHub.

License CC BY-NC-ND 4.0. Rodriguez and Ovaskainen. All rights reserved, including those for text and data mining, AI training, and similar technologies.

---

#### Jsmm R-package installation

To install the R package, run the following commands on R:

```
install.packages("remotes")
remotes::install_github("lufrodriguezca/Jsmm")
```

#### scripts_simulated folder

Contains scripts for performing all the workflow of the Jsmm R-package (Fig. 1B main manuscript) for a simple example with a square domain. The Vignettes explaining step by step this example can be found in the Supporting Information S2.

#### scripts_moths folder

Contains scripts for performing all the workflow of the Jsmm R-package (Fig. 1B main manuscript) for the main example in the manuscript.

The empirical data used in the main example in the manuscript was originally published by Slade et al., 2013 and used as a case study in Ovaskainen et al., 2019.

The air temperature covariate data used in the main example in this manuscript were originally published by Rennie et al., 2017 :

#### Supporting_information_S2.pdf file

Vignettes demonstrating the application of the R-package Jsmm.

### References

Ovaskainen, O., Ramos, D. L., Slade, E. M., Merckx, T., Tikhonov, G., Pennanen, J., Pizo, M. A., Ribeiro, M. C., & Morales, J. M. (2019). Joint species movement modeling: How do traits influence movements. *Ecology*, *100*(4), e02622. https://doi.org/10.1002/ecy.2622

Rennie, S., Adamson, J., Anderson, R., Andrews, C., Bater, J., Bayfield, N., Beaton, K., Beaumont, D., Benham, S., Bowmaker, V., Britt, C., Brooker, R., Brooks, D., Brunt, J., Common, G., Cooper, R., Corbett, S., Critchley, N., Dennis, P., . . . Wood, C. (2017). Uk environmental change network (ecn) meteorology data: 1991-2015. https://doi.org/10.5285/fc9bcd1c-e3fc-4c5a-b569-2fe62d40f2f5

Slade, E. M., Merckx, T., Riutta, T., Bebber, D. P., Redhead, D., & Riordan, P. (2013). Life-history traits and landscape characteristics predict macro-moth responses to forest fragmentation. *Ecology*, *94*(7), 1519–1530. https://doi.org/10.1890/12-1366.1
