## Supplementary material for "R-package Jsmm: Joint species movement modelling of mark-recapture data": S2 Vignettes demonstrating the application of the R-package Jsmm: 1a_define_model_moths.R

```
library(ggcorrplot)
library(mapview)
library(here)
library(Jsmm)

#---------------------------------------------------------------------------
# LOAD domain AND ITS TRIANGULATION
#---------------------------------------------------------------------------

load(file = here("data_moths","polygonal_domain.Rdata"))

# load optional parameters for producing plot:
load(file = here("data_moths","customize_domain_plot.Rdata"))

max_t_area  = 820000 # max area and min angle for the triangulation (See R-package RTriangle)
min_t_angle = 20

domain = Jsmm::jsmm_add_triangulation(domain = domain_p, max_t_area = max_t_area , min_t_angle = min_t_angle)


Jsmm::plot_domain(domain = domain, customize_plot = customize_plot)


mapview(domain$polygon, zcol = "id") # visualize triangulation with zoom


#---------------------------------------------------------------------------
# CREATE observation_effort
#---------------------------------------------------------------------------

# Adding triangulation information to the observation_effort list

load(file = here("data_moths","observation_effort.Rdata"))

plot_observation_effort(observation_effort , xlab = "time (days)", ylab = "Location",
                        by_x = 1, cex.axisx = 1, cex.axisy = 1, lasy = 2, tcky = 0)

#---------------------------------------------------------------------------
# Spatial and temporal covariates
#---------------------------------------------------------------------------

air_temperature_data = read.csv(file = "data_moths/air_temperature_data.csv")

cov_data = list()

vegetation_type = rep("forest_fragment", length(domain[["polygon"]][["id"]]))
vegetation_type[which(domain[["polygon"]][["id"]] == 2)] = "agricultural_matrix"

cov_data[["e_data"]] = data.frame(vegetation_type = as.factor(vegetation_type))
cov_data[["t_data"]] = data.frame(air_temperature = scale(as.numeric(air_temperature_data[, 2] )))
cov_data[["t_data_times"]] = 0:(nrow(air_temperature_data) - 1)

plot(x = cov_data[["t_data_times"]], y = cov_data[["t_data"]][, 1], col="red", pch = 16, cex = 0.9 , xlab = "time (day)",
     ylab = expression(paste("Standarized air temperature") ), xaxt = "n")
grid(nx = NA, ny = NULL, lty = 2, col = "gray", lwd = 2)
lines(x = cov_data[["t_data_times"]], y = cov_data[["t_data"]][, 1], col = "red", lty = 1, )
axis(1, at = seq(min(cov_data[["t_data_times"]]), max(cov_data[["t_data_times"]]), by = 1) , las = 2, cex.axis = 0.8 )


#---------------------------------------------------------------------------
# Trait and phylogenetic data
#---------------------------------------------------------------------------

trait_data = read.csv(file = "data_moths/trait_data.csv")

tr_data = data.frame(wingspan_mm = scale(trait_data[, 4]), wing_shape = as.factor(trait_data[, 5]),
                     larval_feeding_guild = as.factor(trait_data[, 6]), larval_diet_breadth = as.factor(trait_data[, 7]),
                     adult_feeding = as.factor(trait_data[, 8]))

C = read.csv(file = "data_moths/phylogenetic_matrix_data.csv")

ns = nrow(C)

colnames(C) = trait_data[, 2]
rownames(C) = trait_data[, 2]

C = as.matrix(C)

ggcorrplot(C)


#---------------------------------------------------------------------------
# Set model formula
#---------------------------------------------------------------------------

model_formula = list()
model_formula[["diffusion"]] = ~ air_temperature
model_formula[["advection_1"]] = NULL
model_formula[["advection_2"]] = NULL
model_formula[["mortality"]] = ~ 1
model_formula[["habitat_preference"]] = ~ vegetation_type
model_formula[["traits"]] = ~ wingspan_mm
model_formula[["observation"]] = ~ 1


#---------------------------------------------------------------------------
# Set capture data
#---------------------------------------------------------------------------

releases_data = read.csv(file = "data_moths/releases_data.csv")
captures_data = read.csv(file = "data_moths/captures_data.csv")

releases = as.matrix(releases_data)

colnames(releases) = c("sp", "release_event")

captures = as.matrix(captures_data)
colnames(captures) = c("individual", "capture_event", "secondary_release" )

data = list()
data[["releases"]] = releases
data[["captures"]] = captures


#---------------------------------------------------------------------------
# Define model m
#---------------------------------------------------------------------------

m = Jsmm::jsmm(domain = domain, observation_effort = observation_effort,
         releases = data[["releases"]], captures = data[["captures"]],
         cov_data = cov_data,
         ns = ns, sp_names =  trait_data[, 2], tr_data = tr_data, C = C,
         model_formula = model_formula)

model_summary(m)

save(m, file = paste0("models/unfitted_model_moths.RData"))
```
