## Supplementary material for "R-package Jsmm: Joint species movement modelling of mark-recapture data": S2 Vignettes demonstrating the application of the R-package Jsmm: 2a_fit_model.R

```
library(here)
library(Jsmm)

thin = 5
samples = 250
transient = round(0.5*samples*thin)
nChains = 6

startFromEarlier = TRUE
excludeBadChains = TRUE
iii = 2

if(startFromEarlier){
  load("models/fitted_model_moths_samples_250_thin_5_nChains_6.RData")

  tmp = m$postList[[iii]]
  init_pars = tmp[[length(tmp)]]

  if(excludeBadChains){
    bad.chains = matrix(FALSE,nrow=m$ns,ncol=length(m$postList))
    for(sp in 1:m$ns){
      li = matrix(ncol=nChains,nrow = length(m$postList[[1]]))
      for(i in 1:length(m$postList)){
        for(j in 1:length(m$postList[[1]])){
          li[j, i] = m$postList[[i]][[j]]$li[sp]
        }
      }
      plot(NULL, xlim = c(0,length(m$postList[[1]])), ylim = c(min(li), max(li)))
      for(i in 1:length(m$postList)) lines(li[, i])
      me = colMeans(li)
      sd = sqrt(diag(var(li)))
      bad.chains[sp,me<max(me - 3*sd)] = TRUE
    }
    print(colMeans(bad.chains))
    for(sp in which(bad.chains[, iii])){
      iii2 = sample(which(!bad.chains[sp, ]), size = 1)
      tmp2 = m$postList[[iii2]]
      be = tmp2[[length(tmp2)]][["Beta"]]
      for(na in names(init_pars[["Beta"]])){
        init_pars[["Beta"]][[na]][sp, ] = be[[na]][sp, ]
      }
    }
  }
} else {
  init_pars = NULL
}

mname = "moths"
for(mname in c("moths")){
  load(paste0("models/unfitted_model_", mname, ".RData"))
  m = Jsmm::merge_rc_data(m)

  for(i in iii){
    set.seed(i)
    print(i)
    m = Jsmm::sampleMcmc(m = m, samples = samples, transient = transient, thin = thin,
                   nChains = 1,init_pars = init_pars)

    save(m, file = paste0(paste0("models/", i, "_fitted_model_", mname, "_samples_",
                                 samples, "_thin_", thin, "_nChains_", nChains, ".RData")))
  }

  if(FALSE){

    postList = list()

    for(i in 1:nChains){
      fname = paste0(paste0("models/", i, "_fitted_model_", mname, "_samples_",
                            samples, "_thin_", thin, "_nChains_", nChains,".RData"))
      load(file = fname)
      postList[[i]] = m[["postList"]][[1]]
      #file.remove(fname)
    }

    m[["postList"]] = postList

    save(m, file = paste0(paste0("models/fitted_model_", mname, "_samples_", samples, "_thin_", thin,
         "_nChains_", nChains, ".RData")))
  }
}
```
