## Supplementary material for "R-package Jsmm: Joint species movement modelling of mark-recapture data": S2 Vignettes demonstrating the application of the R-package Jsmm: 2b_remove_bad_chains.R

```
library(Jsmm)

thin = 5
samples = 250
transient = round(0.5*samples*thin)
nChains = 6

load("models/fitted_model_moths_samples_250_thin_5_nChains_6.RData")


bad.chains = matrix(FALSE, nrow = m$ns, ncol = length(m$postList))
for(sp in 1:m$ns){
  li = matrix(ncol = nChains, nrow = length(m$postList[[1]]))
  for(i in 1:length(m$postList)){
    for(j in 1:length(m$postList[[1]])){
      li[j, i] = m$postList[[i]][[j]]$li[sp]
    }
  }
  plot(NULL, xlim = c(0, length(m$postList[[1]])), ylim = c(min(li), max(li)))
  for(i in 1:length(m$postList)) lines(li[, i])
  me = colMeans(li)
  sd = sqrt(diag(var(li)))
  bad.chains[sp, me < max(me - 3*sd)] = TRUE
}
print(colMeans(bad.chains))

p = m$postList
m$postList = list(p[[1]], p[[3]], p[[4]], p[[5]], p[[6]])
save(m, file = "models/fitted_model_moths_samples_250_thin_5_nChains_5.RData")
```
