## Supplementary material for "R-package Jsmm: Joint species movement modelling of mark-recapture data": S2 Vignettes demonstrating the application of the R-package Jsmm: 2c_evaluate_MCMC_convergence.R

```
library(coda)

for(mname in c("moths")){
  thin = 5
  samples = 250
  nChains = 5
  load(paste0(paste0("models/fitted_model_", mname, "_samples_", samples, "_thin_", thin, "_nChains_", nChains, ".RData")))
  mpost = Jsmm::convert_to_coda_object(m)
#  plot(mpost$Beta)
#  plot(mpost$Gamma)
  pdf(paste0("results/MCMC_convergence_", mname, ".pdf"))
  plot(mpost$Beta)
  plot(mpost$Gamma)
  if(!is.null(m$C)) plot(mpost$rho)
  summary(coda::gelman.diag(mpost$Beta,multivariate = FALSE)$psrf)
  summary(coda::gelman.diag(mpost$Gamma,multivariate = FALSE)$psrf)
  summary(coda::gelman.diag(mpost$rho,multivariate = FALSE)$psrf)
  dev.off()
}

bad.chains = matrix(FALSE, nrow = m$ns, ncol = nChains)
for(sp in 1:m$ns){
  li = matrix(ncol = nChains, nrow = samples)
  for(i in 1:nChains){
    for(j in 1:samples){
      li[j, i] = m$postList[[i]][[j]]$li[sp]
    }
  }
  plot(NULL, xlim = c(0, samples), ylim = c(min(li), max(li)), main = sp)
  for(i in 1:nChains) lines(li[, i])
  me = colMeans(li)
  sd = sqrt(diag(var(li)))
  bad.chains[sp, me < max(me - 2*sd)] = TRUE
}
colMeans(bad.chains)
```
