## Supplementary material for "R-package Jsmm: Joint species movement modelling of mark-recapture data": S2 Vignettes demonstrating the application of the R-package Jsmm: 3a_show_posteriors_vs_priors.R

```
library(Jsmm)

showTrueValues = FALSE
for(mname in c("moths")){
  thin = 5
  samples = 250
  nChains = 5
  filename = paste0(paste0("models/fitted_model_", mname, "_samples_", samples, "_thin_", thin, "_nChains_", nChains, ".RData"))
  load(filename)
  sel = 1:m$ns
  if(mname=="moths"){sel = order(m$tr_data$wingspan_mm, decreasing = F)}
  post = poolMcmcChains(m[["postList"]])
  npost = length(post)
  vars = names(m[["nc"]][m[["nc"]] > 0])
  ns = m[["ns"]]
  nt = m[["nt"]]

  nprior = npost
  prior = list()
  for(repl in 1:nprior){
    prior[[repl]] = simulate_from_prior(m)
  }

  if(showTrueValues){
    if(mname == "simulated_ICP") load("parameters/true_parameters_ICP.RData")
    if(mname == "simulated_CCP") load("parameters/true_parameters_CCP.RData")
  }

 pdf(paste0("results/posteriors_vs_priors_", mname, ".pdf"))
  par(mfrow = c(2, 2))
  if(m[["ns"]] < 3) par(mfrow = c(2, 3))
  if(m[["ns"]] > 10) par(mfrow = c(2, 1))
  cc = 0
  for(va in vars){
    nv = m[["nc"]][[va]]
    for(i in 1:nv){
      cc = cc + 1

      re.pri = matrix(ncol = ns,nrow = nprior)
      re.pri.G = matrix(ncol = nt, nrow = nprior)
      for(repl in 1:nprior){
        re.pri[repl,] = prior[[repl]][["Beta"]][[va]][,i]
        re.pri.G[repl,] = prior[[repl]][["Gamma"]][,cc]
      }
      re.post = matrix(ncol = ns,nrow = npost)
      re.post.G = matrix(ncol = nt,nrow = nprior)
      for(repl in 1:npost){
        re.post[repl,] = post[[repl]][["Beta"]][[va]][,i]
        re.post.G[repl,] = post[[repl]][["Gamma"]][,cc]
      }
      ymin = min(quantile(re.pri,probs = 0.05),min(re.post))
      ymax = max(quantile(re.pri,probs = 0.95),max(re.post))
      if(showTrueValues){
        ymin = min(ymin,pars$Beta[[va]][,i])
        ymax = max(ymax,pars$Beta[[va]][,i])
      }
      colnames(re.pri) = m$sp_names
      re.pri = re.pri[,sel]
      re.post = re.post[,sel]
      boxplot(re.pri,outline = FALSE,at=3*(1:ns)-0.75,ylim=c(ymin,ymax),
              main=m[["parNames"]][cc],las=2,cex.axis=0.7,
              xlim = c(2,3*ns+2),
              ylab = "parameter value",col="grey",xaxt = "n")
      axis(1,at=3*(1:m$ns),label=colnames(re.pri),las=2,cex.axis=0.7)
      boxplot(re.post,outline = FALSE,at=3*(1:ns)+0.75,col="pink",add=TRUE,xaxt="n",yaxt="n")
      abline(h=0)
      if(showTrueValues){
        for(j in 1:m[["ns"]]){
          lines(x=c(3*j,3*j+1.5),y=c(pars$Beta[[va]][j,i],pars$Beta[[va]][j,i]),lwd=2,col="red")
        }
      }
      ymin = min(quantile(re.pri.G,probs = 0.05),min(re.post.G))
      ymax = max(quantile(re.pri.G,probs = 0.95),max(re.post.G))
      if(showTrueValues){
        ymin = min(ymin,pars$Gamma[,cc])
        ymax = max(ymax,pars$Gamma[,cc])
      }
      colnames(re.pri.G) = colnames(m$X[["trait"]])
      boxplot(re.pri.G,outline = FALSE,at=3*(1:nt),ylim=c(ymin,ymax),
              main=m[["parNames"]][cc],las=2,cex.axis=0.7,
              xlim = c(2,3*nt+2),
              ylab = "parameter value",col="grey")
      boxplot(re.post.G,outline = FALSE,at=3*(1:nt)+1.5,col="pink",add=TRUE,xaxt="n",yaxt="n")
      abline(h=0)
      if(showTrueValues){
        for(j in 1:m[["nt"]]){
          lines(x=c(3*j,3*j+1.5),y=c(pars$Gamma[j,cc],pars$Gamma[j,cc]),lwd=2,col="red")
        }
      }
    }
  }
  if(!is.null(post[[1]][["rho"]])){
    re.pri = rep(nprior)
    re.post = rep(npost)
    for(repl in 1:nprior){
      re.pri[repl] = prior[[repl]][["rho"]]
    }
    for(repl in 1:npost){
      re.post[repl] = post[[repl]][["rho"]]
    }
    boxplot(re.pri,ylim=c(0,1),xlim=c(0,2),at = 0.5, main = "rho", col = "grey",
            ylab = "parameter value",las=2,cex.axis=0.7)
    boxplot(re.post,add=TRUE,at = 1.5, col = "pink",yaxt='n')
    if(showTrueValues){
      lines(x=c(0.27,1.75),y=rep(pars$rho,2),lwd=2,col="red")
    }
  }

  dev.off()
}

np = length(post)
gamma.mean = post[[1]]$Gamma
gamma.pos = 1*(post[[1]]$Gamma>0)
for(i in 2:npost){
  gamma.mean = gamma.mean + post[[i]]$Gamma
  gamma.pos = gamma.pos + 1*(post[[i]]$Gamma>0)
}
gamma.mean = gamma.mean/npost
gamma.pos = gamma.pos/npost
colnames(gamma.mean) = m[["parNames"]]
rownames(gamma.mean) = colnames(m[["X"]][["trait"]])
colnames(gamma.pos) = m[["parNames"]]
rownames(gamma.pos) = colnames(m[["X"]][["trait"]])
gamma.mean
gamma.pos

rho.mean = post[[1]]$rho
rho.pos = 1*(post[[1]]$rho>0)
for(i in 2:npost){
  rho.mean = rho.mean + post[[i]]$rho
  rho.pos = rho.pos + 1*(post[[i]]$rho>0)
}
rho.mean = rho.mean/npost
rho.pos = rho.pos/npost
rho.mean
rho.pos
```
