## Supplementary material for "R-package Jsmm: Joint species movement modelling of mark-recapture data": S2 Vignettes demonstrating the application of the R-package Jsmm: 3b_show_moths_results.R

```
library(Jsmm)

mname = "moths"
thin = 5
samples = 250
nChains = 5
filename = paste0(paste0("models/fitted_model_", mname, "_samples_", samples,
                         "_thin_", thin, "_nChains_", nChains, ".RData"))
load(filename)
sel = order(m$tr_data$wingspan_mm, decreasing = F)
post = Jsmm::poolMcmcChains(m[["postList"]])
npost = length(post)
vars = names(m[["nc"]][m[["nc"]] > 0])
ns = m[["ns"]]
nt = m[["nt"]]
for(i in 1:npost){
  beta = post[[i]]$Beta
  beta = cbind(beta$diffusion, beta$mortality, beta$habitat_preference, beta$observation)
  rownames(beta) = m$sp_names
  beta = beta[sel, ]
  colnames(beta) = c("D_0", "D_AT", "M0", "H0", "C0")
  if(i==1){
    sbeta = beta
  } else {
    sbeta = sbeta + beta
  }
}
beta = sbeta/npost
plot(as.data.frame(beta))
rel = table(m$releases[, 1])
rec = table(m$releases[m$captures[, 1], 1])
names(rel) = m$sp_names
names(rec) = m$sp_names

plot(as.numeric(rel), as.numeric(rec), xlab = "number of released individuals", ylab = "number of recaptured individuals")
```
