## Supplementary material for "R-package Jsmm: Joint species movement modelling of mark-recapture data": S2 Vignettes demonstrating the application of the R-package Jsmm: data_source.txt

The empirical data used in the main example in this manuscript was originally published by:

Slade, E. M., Merckx, T., Riutta, T., Bebber, D. P., Redhead, D., & Riordan, P. (2013). Life-history traits and landscape characteristics predict macro-moth responses to forest fragmentation. Ecology, 94(7), 1519–1530. https://doi.org/10.1890/12-1366.1


The air temperature covariate data used in the main example in this manuscript was originally published by: 

Rennie, S., Adamson, J., Anderson, R., Andrews, C., Bater, J., Bayfield, N., Beaton, K., Beaumont, D., Benham, S., Bowmaker, V., Britt, C., Brooker, R., Brooks, D., Brunt, J., Common, G., Cooper, R., Corbett, S., Critchley, N., Dennis, P., … Wood, C. (2017). UK Environmental Change Network (ECN) meteorology data: 1991-2015. NERC Environmental Information Data Centre. https://doi.org/10.5285/fc9bcd1c-e3fc-4c5a-b569-2fe62d40f2f5
