## Supplementary material for "R-package Jsmm: Joint species movement modelling of mark-recapture data": S2 Vignettes demonstrating the application of the R-package Jsmm: MCMC_convergence_moths.pdf

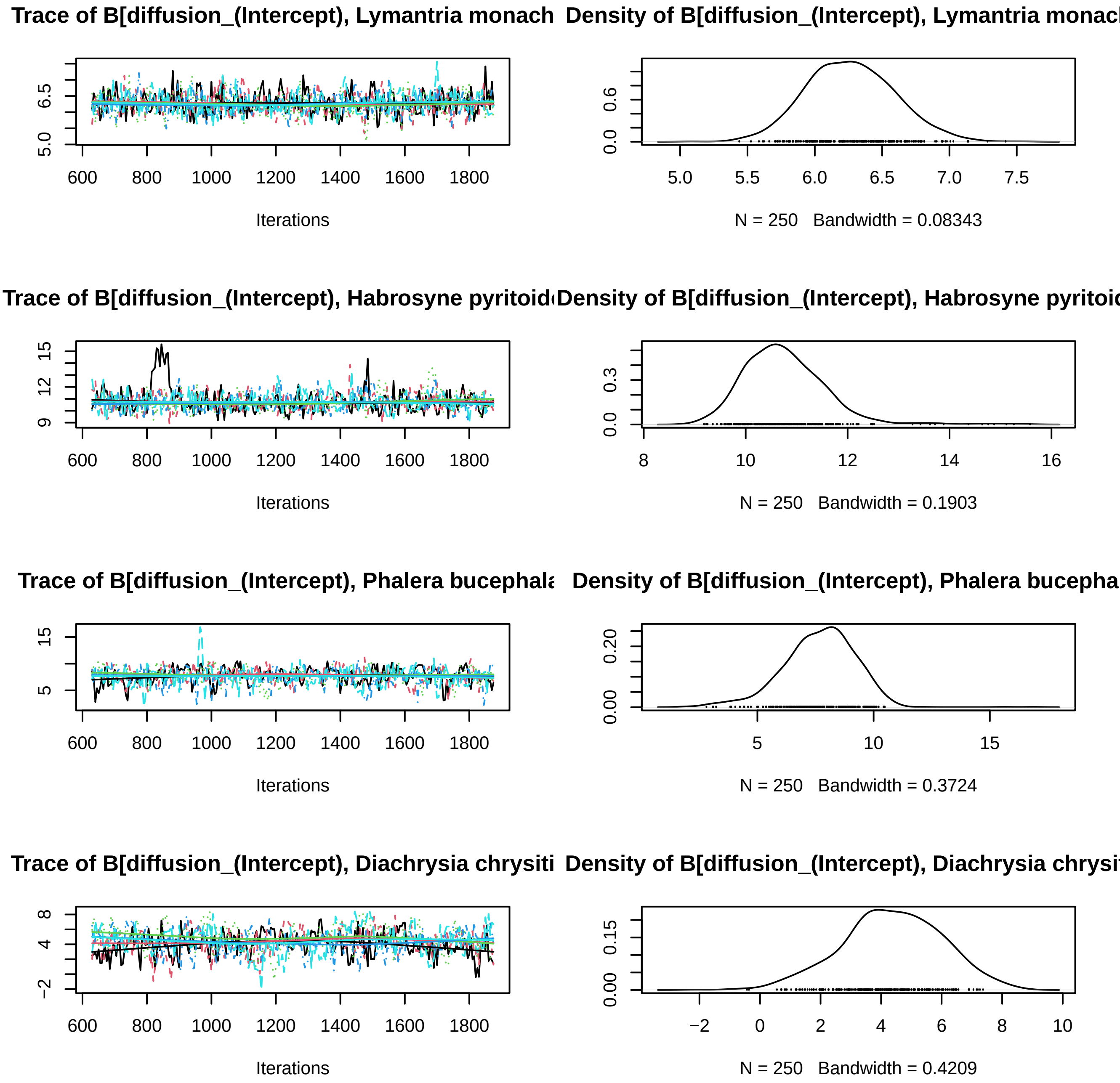

Trace of B[diffusion_(Intercept), Lymantria monach
Density of B[diffusion_(Intercept), Lymantria monac
Iterations
N = 250 Bandwidth = 0.08343
Trace of B[diffusion_(Intercept), Habrosyne pyritoid
Density of B[diffusion_(Intercept), Habrosyne pyritoi
Iterations
N = 250 Bandwidth = 0.1903
Trace of B[diffusion_(Intercept), Phalera bucephala
Density of B[diffusion_(Intercept), Phalera bucepha
Iterations
N = 250 Bandwidth = 0.3724
Trace of B[diffusion_(Intercept), Diachrysia chrysiti
Density of B[diffusion_(Intercept), Diachrysia chrysi
Iterations
N = 250 Bandwidth = 0.4209

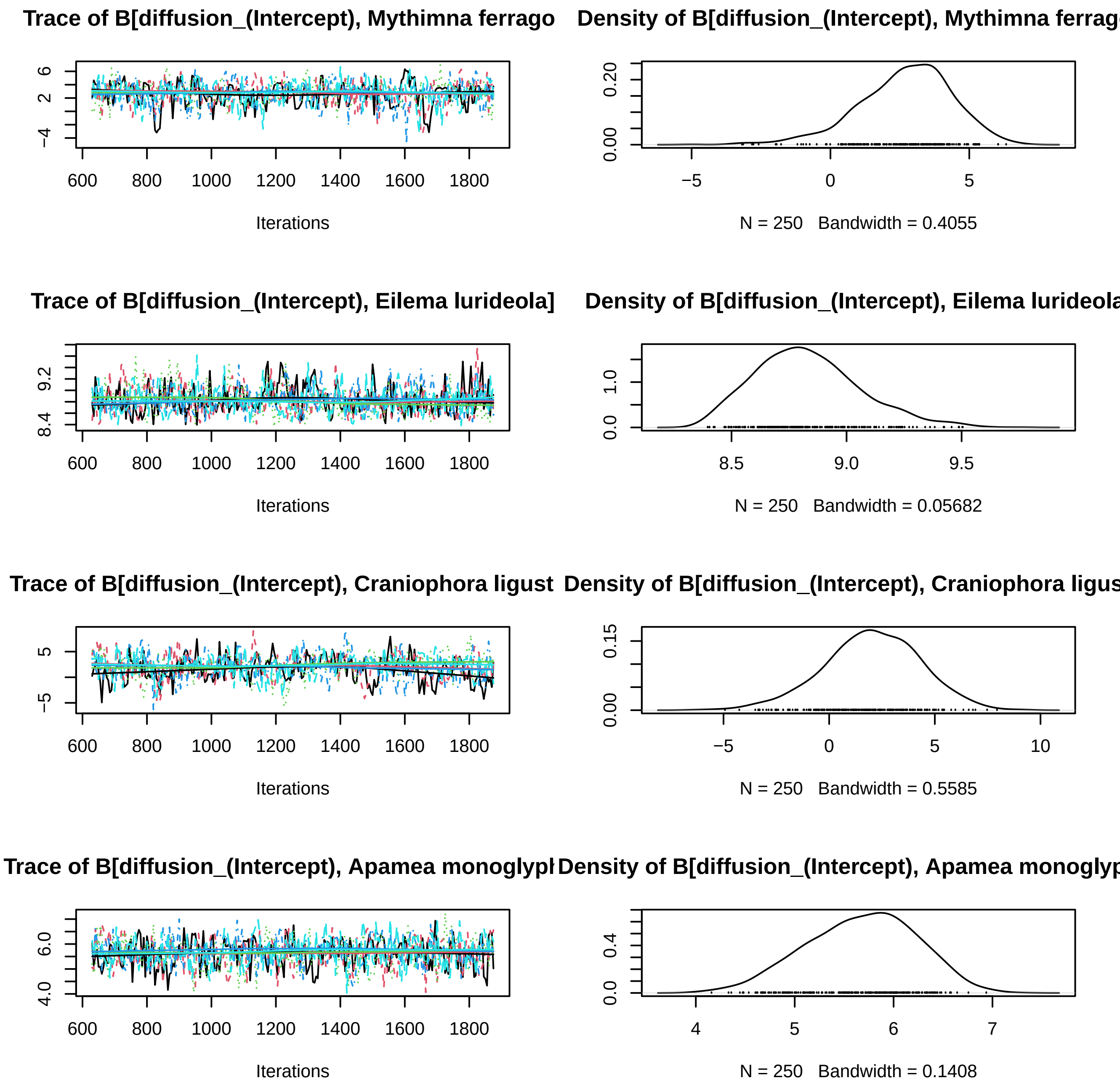

Trace of B[diffusion_(Intercept), Mythimna ferrago
Density of B[diffusion_(Intercept), Mythimna ferrag
Iterations
N = 250 Bandwidth = 0.4055
Trace of B[diffusion_(Intercept), Eilema lurideola]
Density of B[diffusion_(Intercept), Eilema lurideola
Iterations
N = 250 Bandwidth = 0.05682
Trace of B[diffusion_(Intercept), Craniophora ligust
Density of B[diffusion_(Intercept), Craniophora ligus
Iterations
N = 250 Bandwidth = 0.5585
Trace of B[diffusion_(Intercept), Apamea monoglypl
Density of B[diffusion_(Intercept), Apamea monoglyp
Iterations
N = 250 Bandwidth = 0.1408

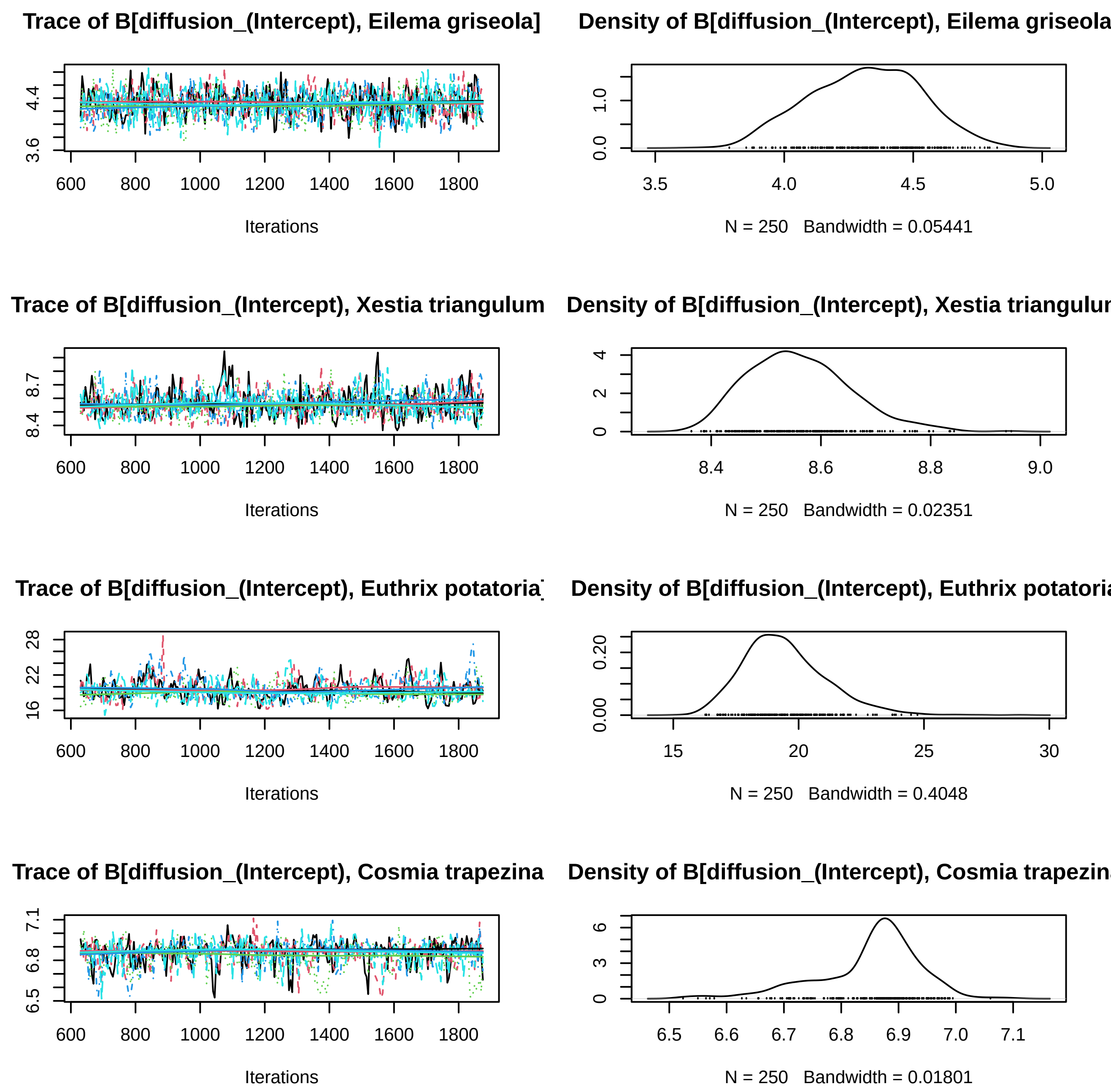
Trace of B[diffusion_(Intercept), Eilema griseola]
Density of B[diffusion_(Intercept), Eilema griseola
Iterations
N = 250 Bandwidth = 0.05441
Trace of B[diffusion_(Intercept), Xestia triangulum
Density of B[diffusion_(Intercept), Xestia triangulur
Iterations
N = 250 Bandwidth = 0.02351
Trace of B[diffusion_(Intercept), Euthrix potatoria]
Density of B[diffusion_(Intercept), Euthrix potatoria
Iterations
N = 250 Bandwidth = 0.4048
Trace of B[diffusion_(Intercept), Cosmia trapezina
Density of B[diffusion_(Intercept), Cosmia trapezin
Iterations
N = 250 Bandwidth = 0.01801

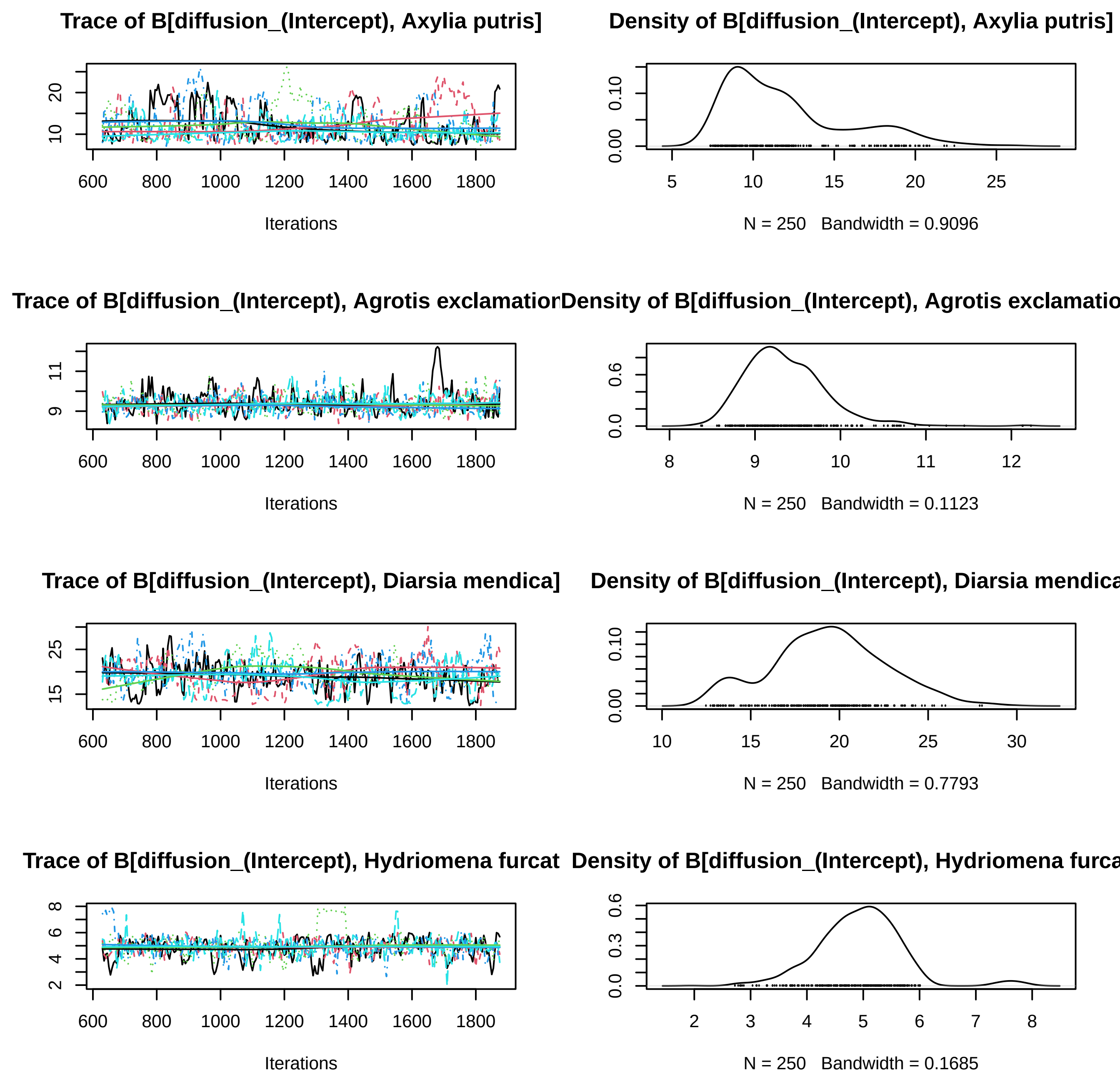
Trace of B[diffusion_(Intercept), Axylia putris]
Density of B[diffusion_(Intercept), Axylia putris]
Iterations
N = 250 Bandwidth = 0.9096
Trace of B[diffusion_(Intercept), Agrotis exclamatior
Density of B[diffusion_(Intercept), Agrotis exclamatio
Iterations
N = 250 Bandwidth = 0.1123
Trace of B[diffusion_(Intercept), Diarsia mendica]
Density of B[diffusion_(Intercept), Diarsia mendica
Iterations
N = 250 Bandwidth = 0.7793
Trace of B[diffusion_(Intercept), Hydriomena furcat
Density of B[diffusion_(Intercept), Hydriomena furca
Iterations
N = 250 Bandwidth = 0.1685

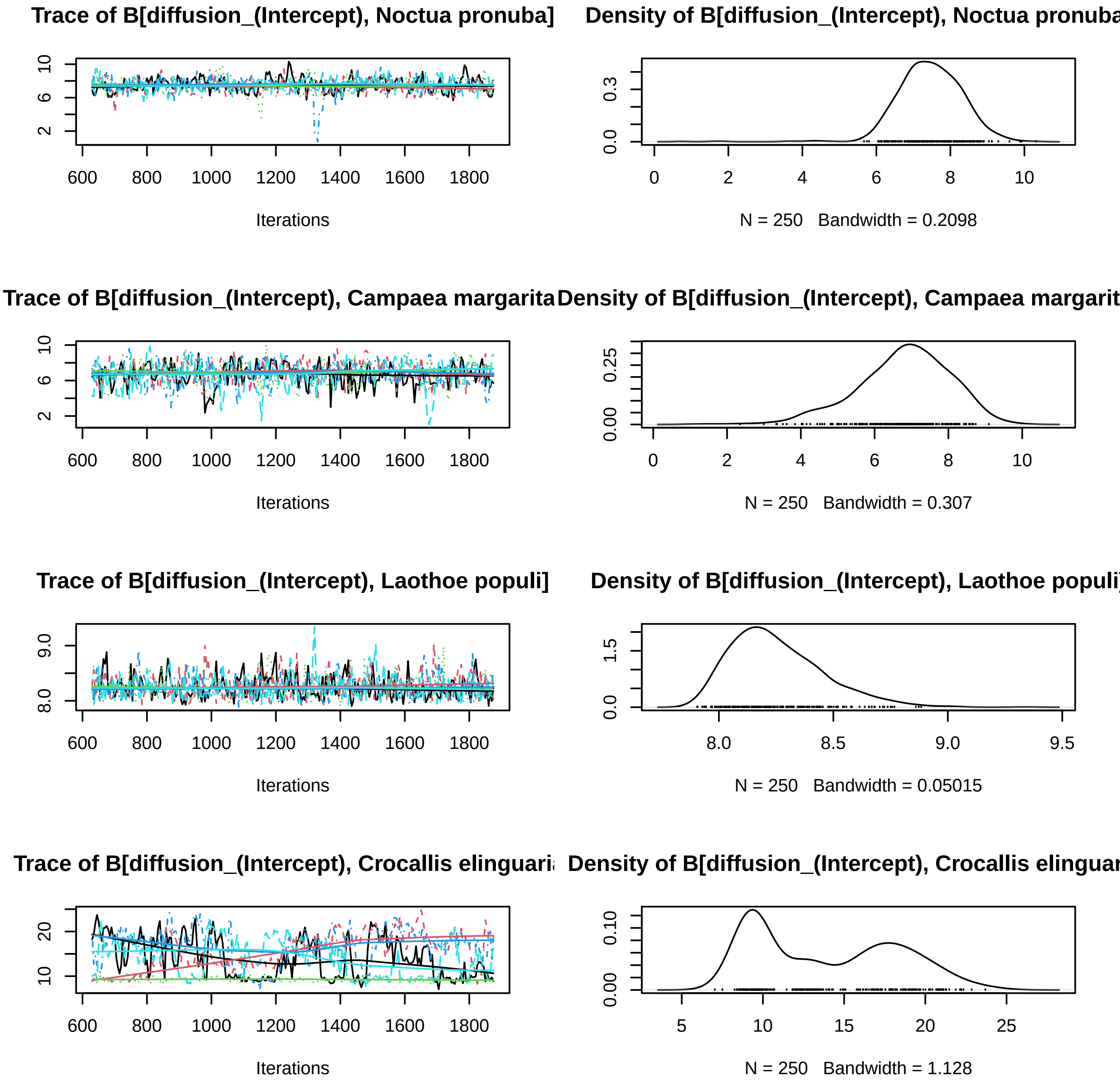
Trace of B[diffusion_(Intercept), Noctua pronuba]
Density of B[diffusion_(Intercept), Noctua pronuba
Iterations
N = 250 Bandwidth = 0.2098
Trace of B[diffusion_(Intercept), Campaea margarita
Density of B[diffusion_(Intercept), Campaea margarit
Iterations
N = 250 Bandwidth = 0.307
Trace of B[diffusion_(Intercept), Laothoe populi]
Density of B[diffusion_(Intercept), Laothoe populi
Iterations
N = 250 Bandwidth = 0.05015
Trace of B[diffusion_(Intercept), Crocallis elinguari
Density of B[diffusion_(Intercept), Crocallis elinguar
Iterations
N = 250 Bandwidth = 1.128

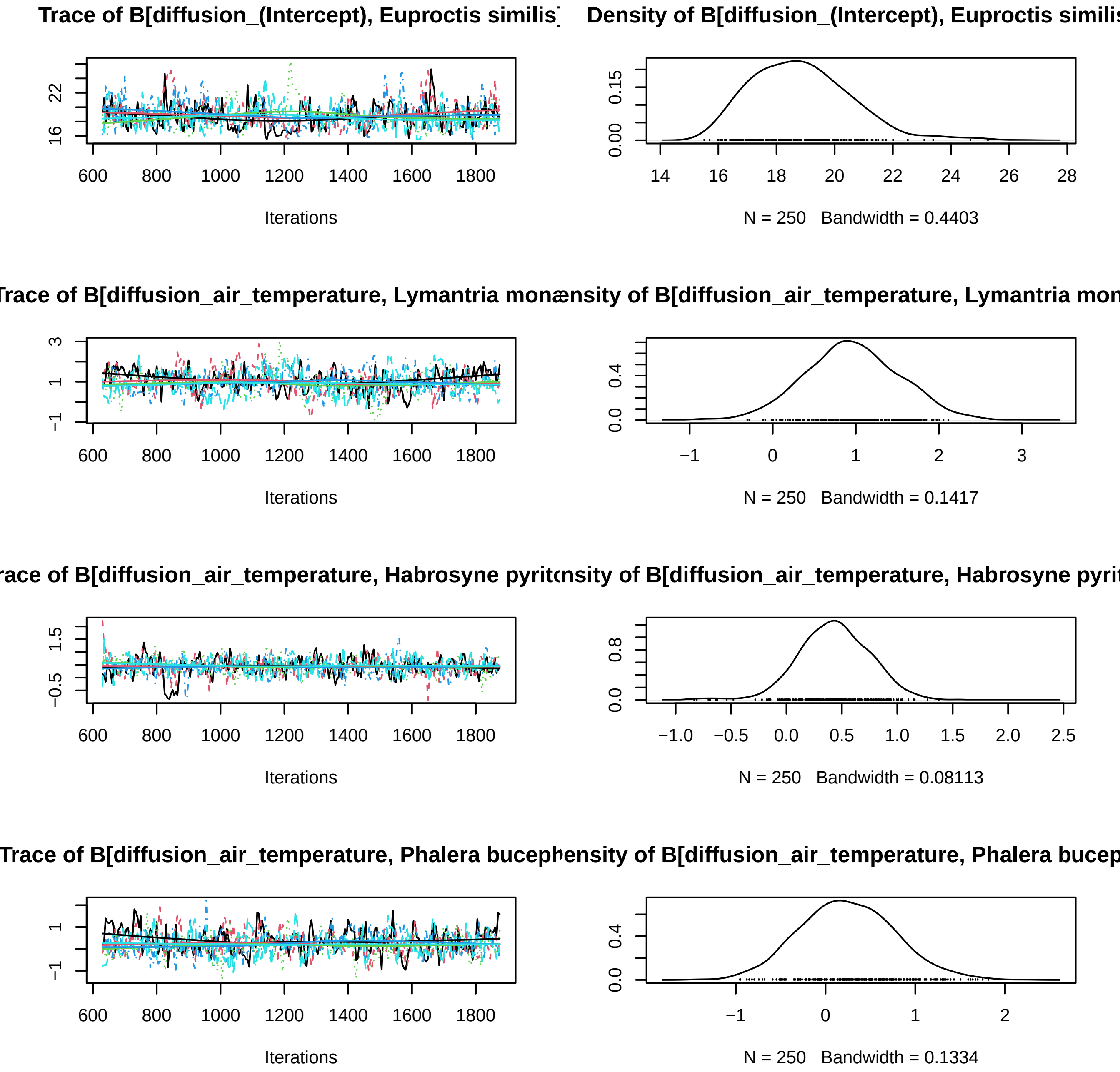
Trace of B[diffusion_(Intercept), Euproctis similis]
Density of B[diffusion_(Intercept), Euproctis similis
Iterations
N = 250 Bandwidth = 0.4403
Trace of B[diffusion_air_temperature, Lymantria mon
ensity of B[diffusion_air_temperature, Lymantria mon
Iterations
N = 250 Bandwidth = 0.1417
race of B[diffusion_air_temperature, Habrosyne pyrit
nsity of B[diffusion_air_temperature, Habrosyne pyrit
Iterations
N = 250 Bandwidth = 0.08113
Trace of B[diffusion_air_temperature, Phalera bucep
hensity of B[diffusion_air_temperature, Phalera bucep
Iterations
N = 250 Bandwidth = 0.1334

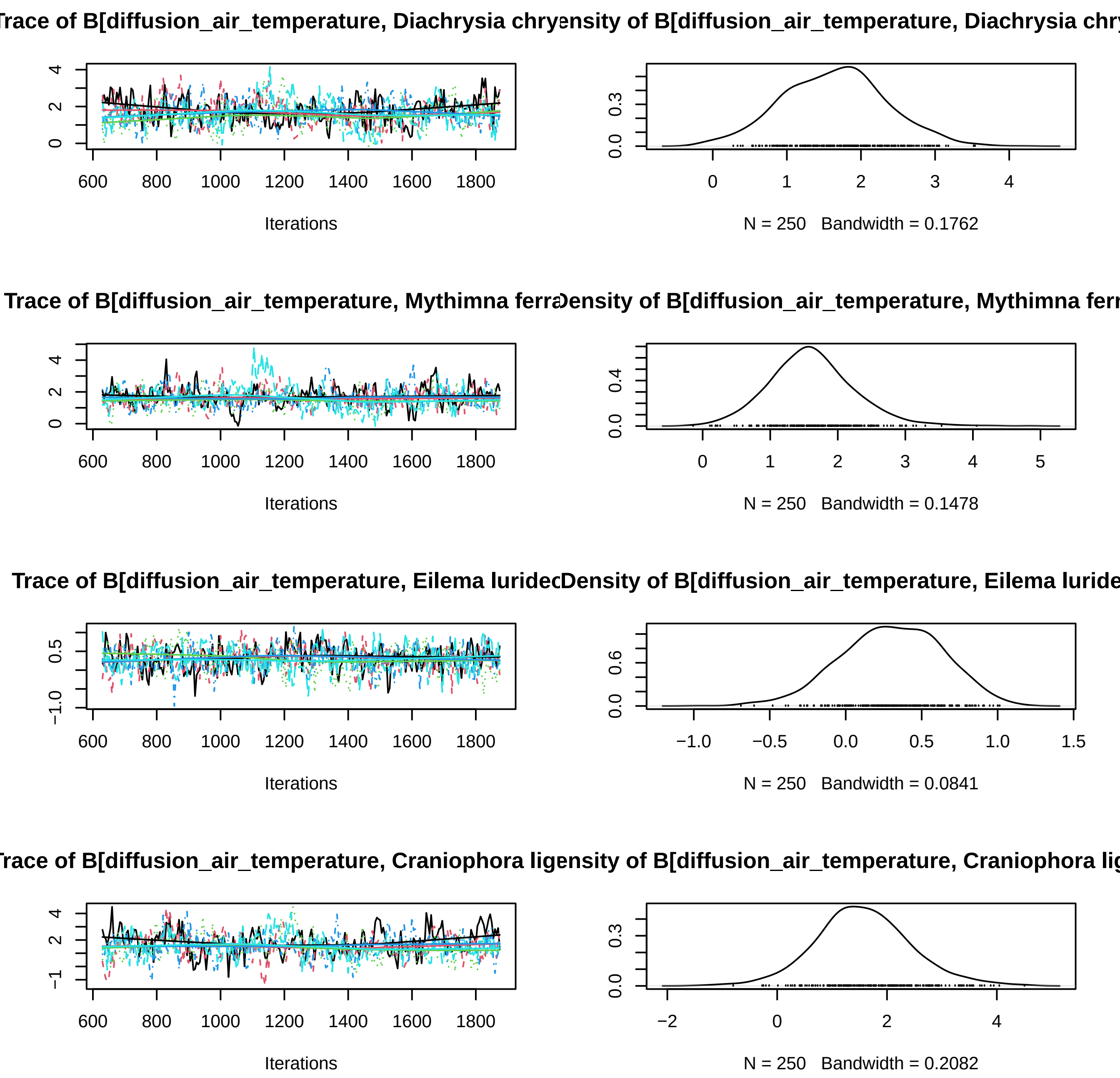
Trace of B[diffusion_air_temperature, Diachrysia chry
Density of B[diffusion_air_temperature, Diachrysia chry
Iterations
N = 250 Bandwidth = 0.1762
Trace of B[diffusion_air_temperature, Mythimna ferra
Density of B[diffusion_air_temperature, Mythimna ferr
Iterations
N = 250 Bandwidth = 0.1478
Trace of B[diffusion_air_temperature, Eilema luridec
Density of B[diffusion_air_temperature, Eilema luride
Iterations
N = 250 Bandwidth = 0.0841
Trace of B[diffusion_air_temperature, Craniophora lig
Density of B[diffusion_air_temperature, Craniophora lig
Iterations
N = 250 Bandwidth = 0.2082

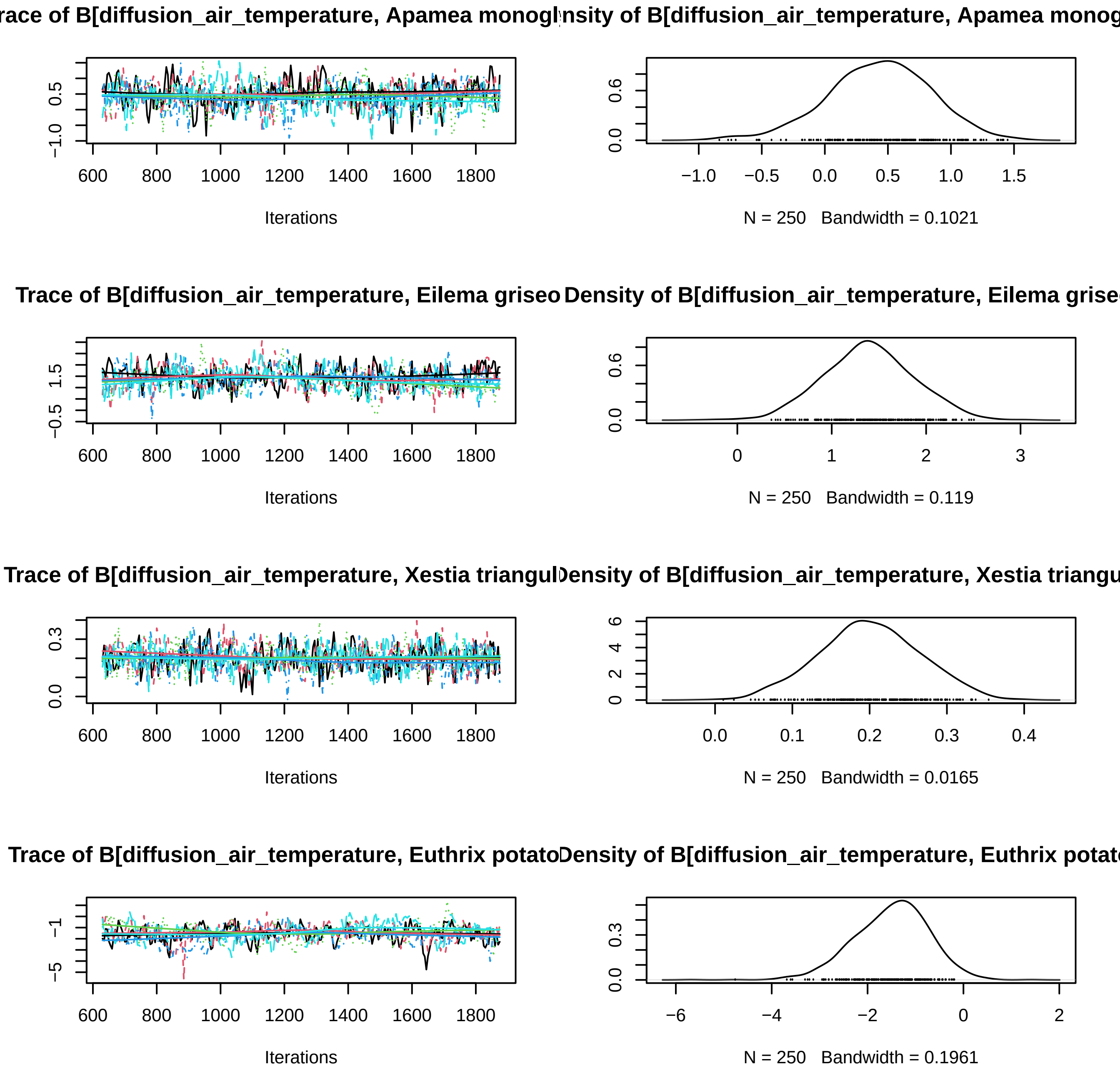
race of B[diffusion_air_temperature, Apamea monogl
nsity of B[diffusion_air_temperature, Apamea monog
Iterations
N = 250 Bandwidth = 0.1021
Trace of B[diffusion_air_temperature, Eilema griseo
Density of B[diffusion_air_temperature, Eilema grise
Iterations
N = 250 Bandwidth = 0.119
Trace of B[diffusion_air_temperature, Xestia triangul
ensity of B[diffusion_air_temperature, Xestia triangu
Iterations
N = 250 Bandwidth = 0.0165
Trace of B[diffusion_air_temperature, Euthrix potato
ensity of B[diffusion_air_temperature, Euthrix potat
Iterations
N = 250 Bandwidth = 0.1961

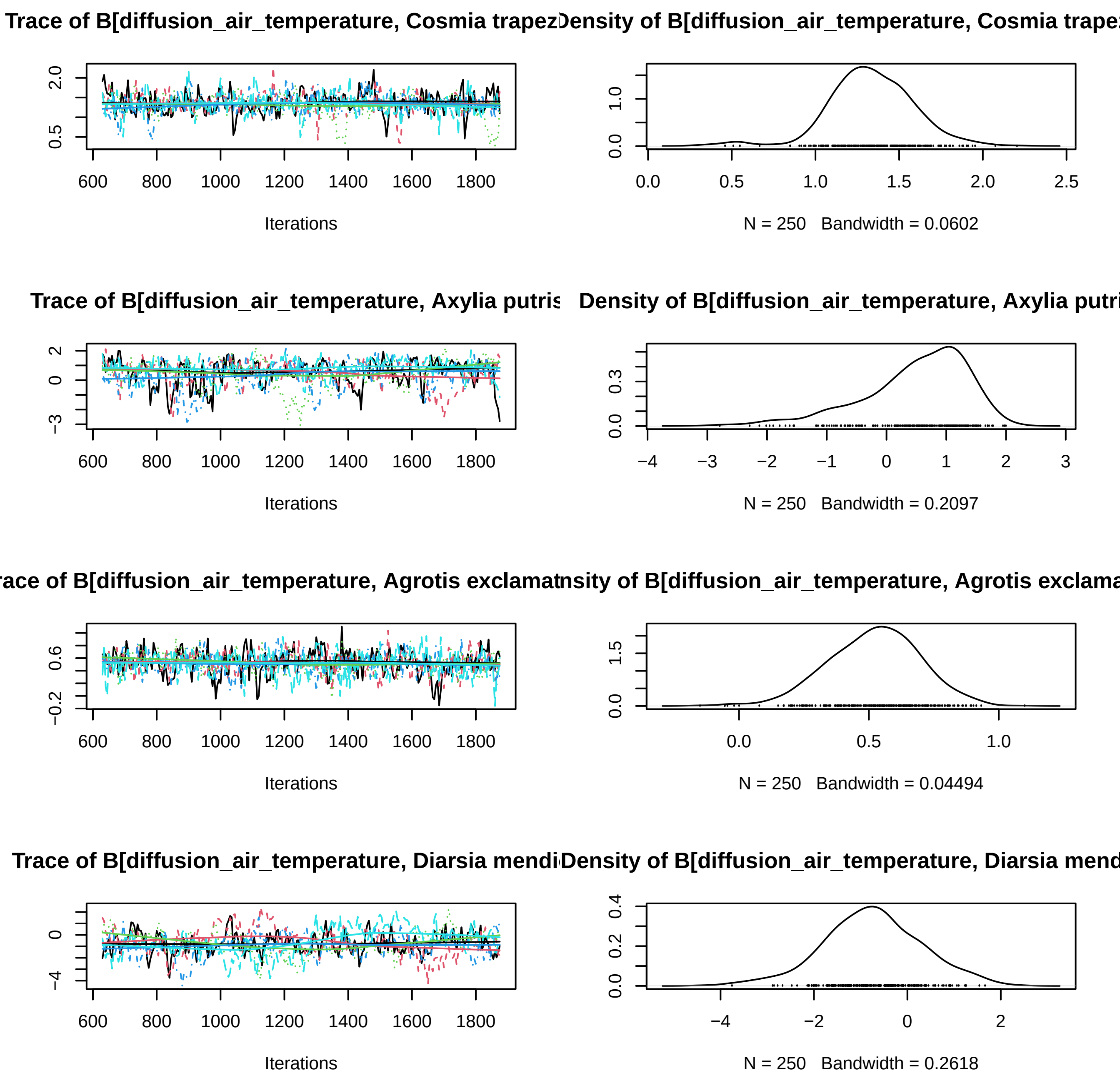

Trace of B[diffusion_air_temperature, Cosmia trapez
Density of B[diffusion_air_temperature, Cosmia trape
Iterations
N = 250 Bandwidth = 0.0602
Trace of B[diffusion_air_temperature, Axylia putris
Density of B[diffusion_air_temperature, Axylia putri
Iterations
N = 250 Bandwidth = 0.2097
race of B[diffusion_air_temperature, Agrotis exclamat
nsity of B[diffusion_air_temperature, Agrotis exclama
Iterations
N = 250 Bandwidth = 0.04494
Trace of B[diffusion_air_temperature, Diarsia mendi
Density of B[diffusion_air_temperature, Diarsia mend
Iterations
N = 250 Bandwidth = 0.2618

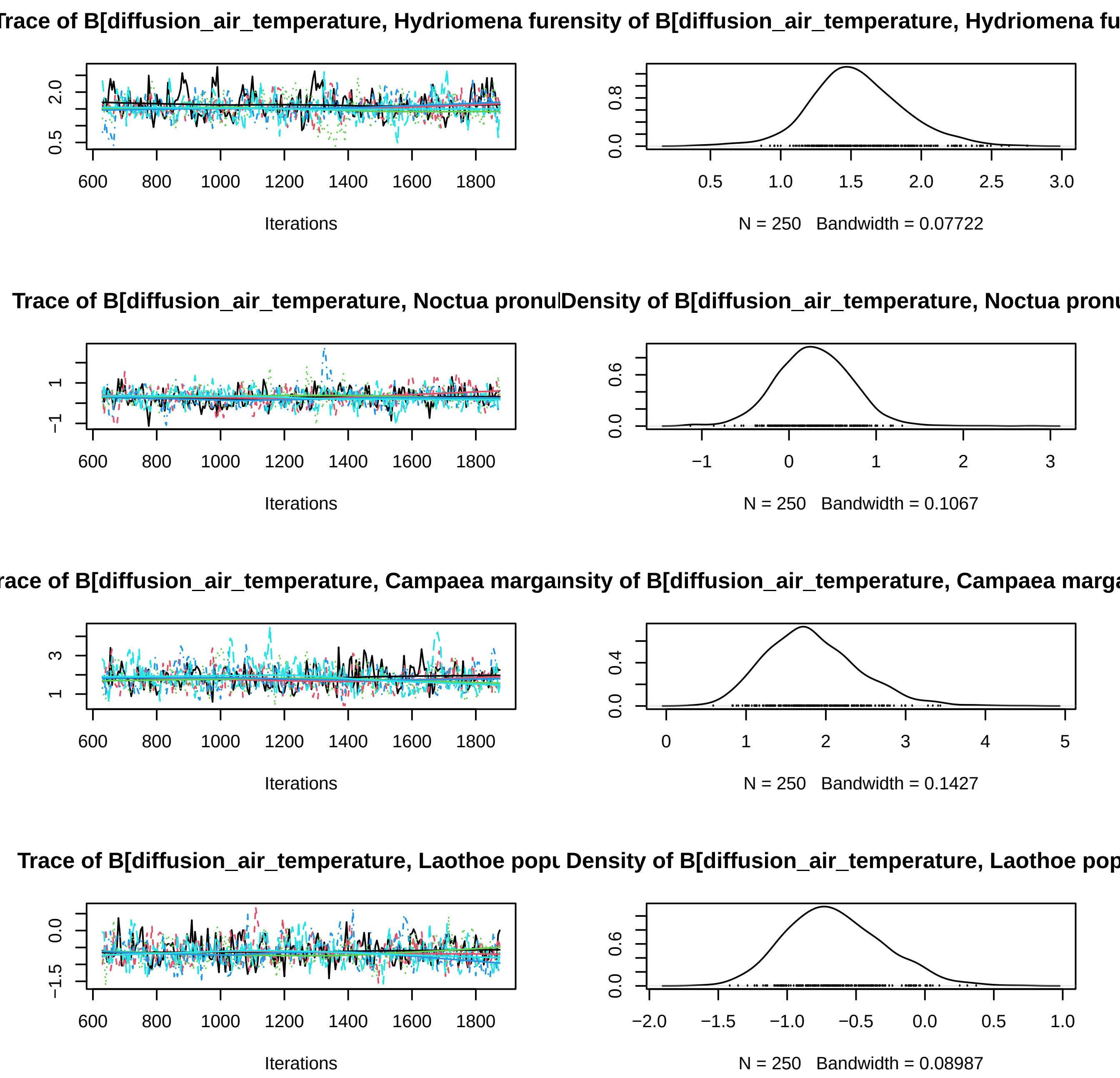
Trace of B[diffusion_air_temperature, Hydriomena furensity of B[diffusion_air_temperature, Hydriomena fu
Iterations
N = 250 Bandwidth = 0.07722
Trace of B[diffusion_air_temperature, Noctua pronuDensity of B[diffusion_air_temperature, Noctua pronu
Iterations
N = 250 Bandwidth = 0.1067
race of B[diffusion_air_temperature, Campaea margansity of B[diffusion_air_temperature, Campaea marga
Iterations
N = 250 Bandwidth = 0.1427
Trace of B[diffusion_air_temperature, Laothoe popu Density of B[diffusion_air_temperature, Laothoe pop
Iterations
N = 250 Bandwidth = 0.08987

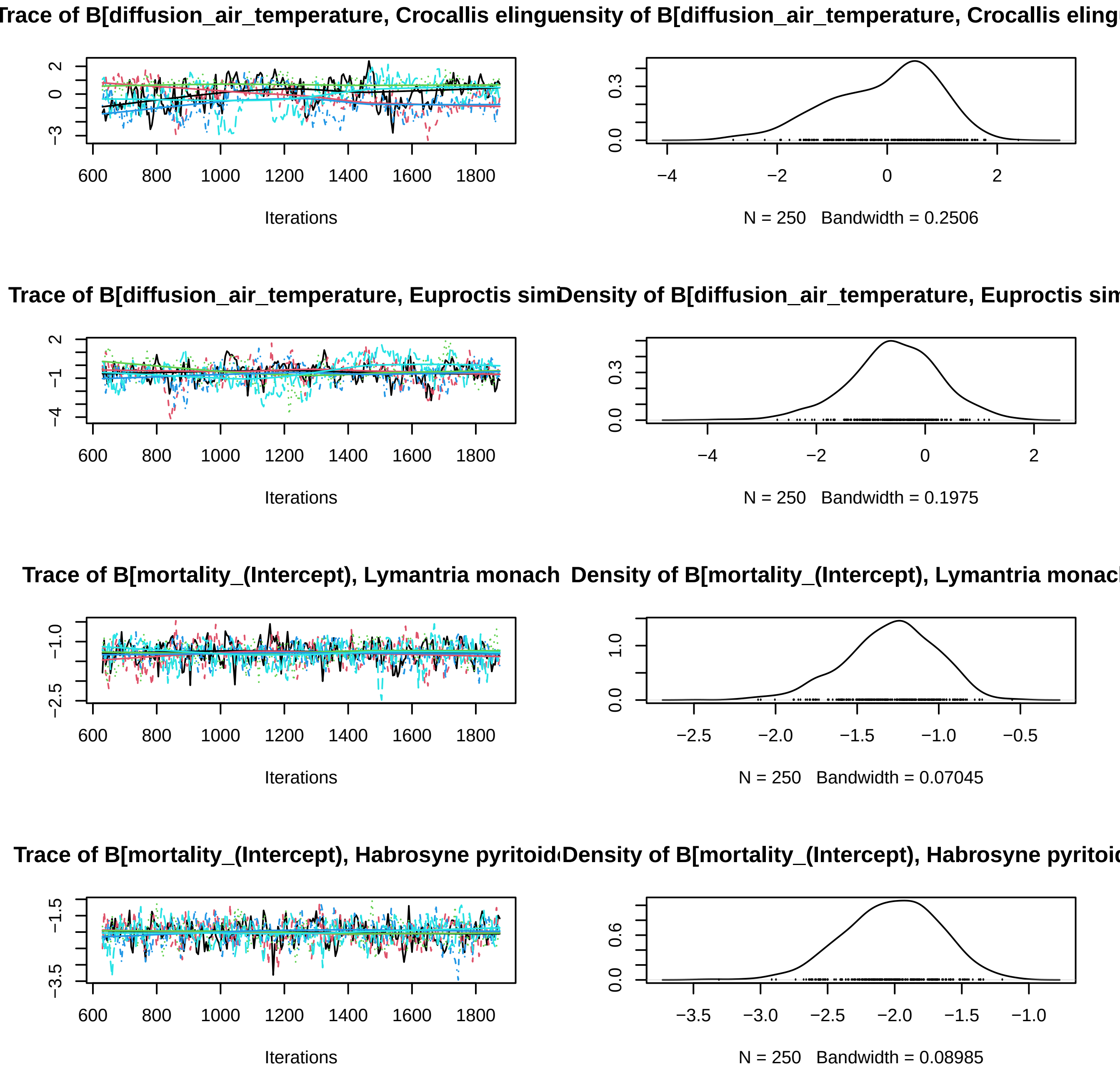

Trace of B[diffusion_air_temperature, Crocallis elingu
Density of B[diffusion_air_temperature, Crocallis eling
Iterations
N = 250 Bandwidth = 0.2506
Trace of B[diffusion_air_temperature, Euproctis sim
Density of B[diffusion_air_temperature, Euproctis sim
Iterations
N = 250 Bandwidth = 0.1975
Trace of B[mortality_(Intercept), Lymantria monach
Density of B[mortality_(Intercept), Lymantria monac
Iterations
N = 250 Bandwidth = 0.07045
Trace of B[mortality_(Intercept), Habrosyne pyritoid
Density of B[mortality_(Intercept), Habrosyne pyritoi
Iterations
N = 250 Bandwidth = 0.08985

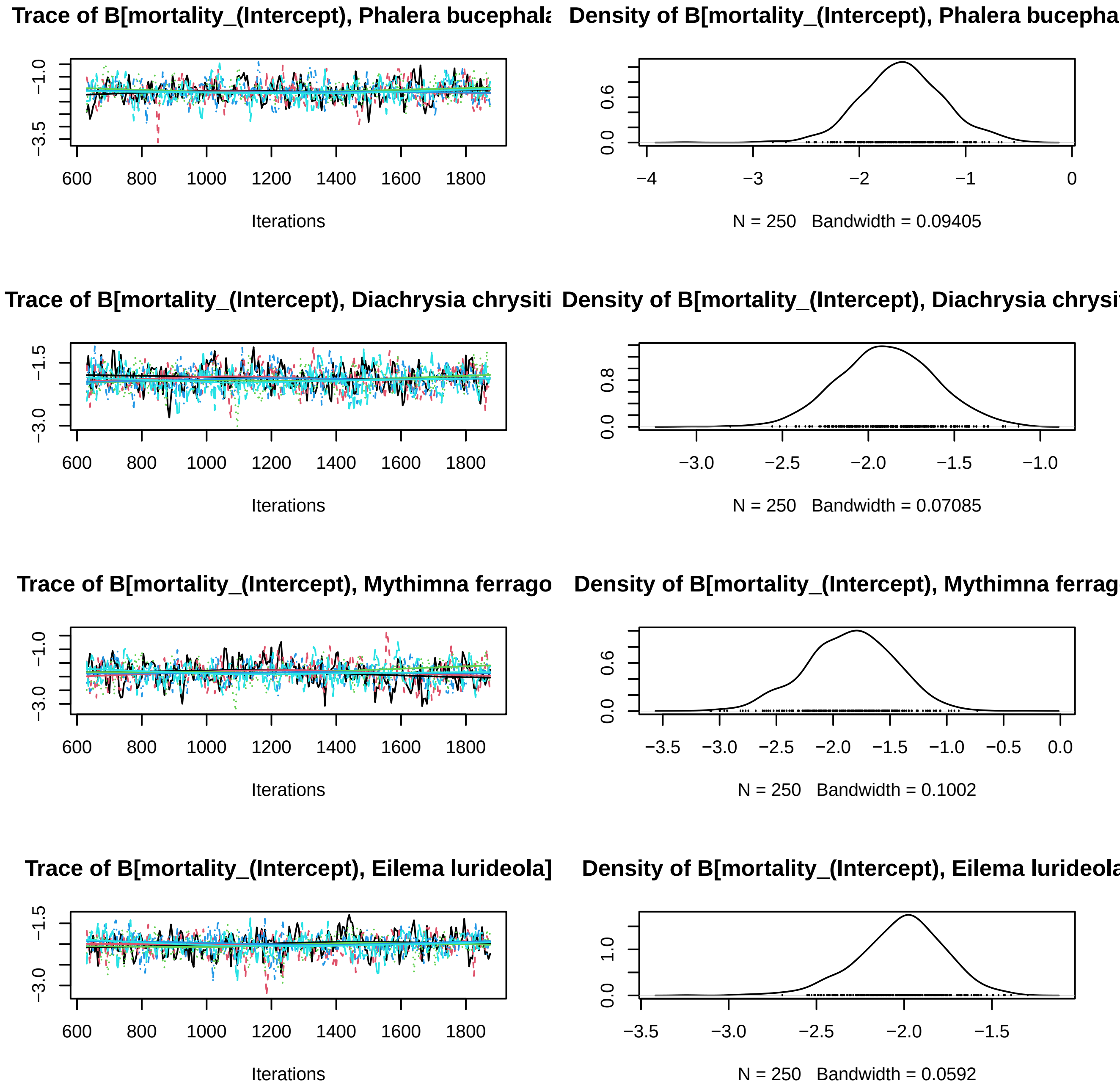

Trace of B[mortality_(Intercept), Phalera bucephala
Density of B[mortality_(Intercept), Phalera bucepha
−1.0
−3.5
600
800
1000
1200
1400
1600
1800
Iterations
0.0
0.6
−4
−3
−2
−1
0
N = 250 Bandwidth = 0.09405
Trace of B[mortality_(Intercept), Diachrysia chrysiti
Density of B[mortality_(Intercept), Diachrysia chrysi
−1.5
−3.0
600
800
1000
1200
1400
1600
1800
Iterations
0.0
0.8
−3.0
−2.5
−2.0
−1.5
−1.0
N = 250 Bandwidth = 0.07085
Trace of B[mortality_(Intercept), Mythimna ferrago
Density of B[mortality_(Intercept), Mythimna ferrag
−1.0
−3.0
600
800
1000
1200
1400
1600
1800
Iterations
0.0
0.6
−3.5
−3.0
−2.5
−2.0
−1.5
−1.0
−0.5
0.0
N = 250 Bandwidth = 0.1002
Trace of B[mortality_(Intercept), Eilema lurideola]
Density of B[mortality_(Intercept), Eilema lurideola
−1.5
−3.0
600
800
1000
1200
1400
1600
1800
Iterations
0.0
1.0
−3.5
−3.0
−2.5
−2.0
−1.5
N = 250 Bandwidth = 0.0592

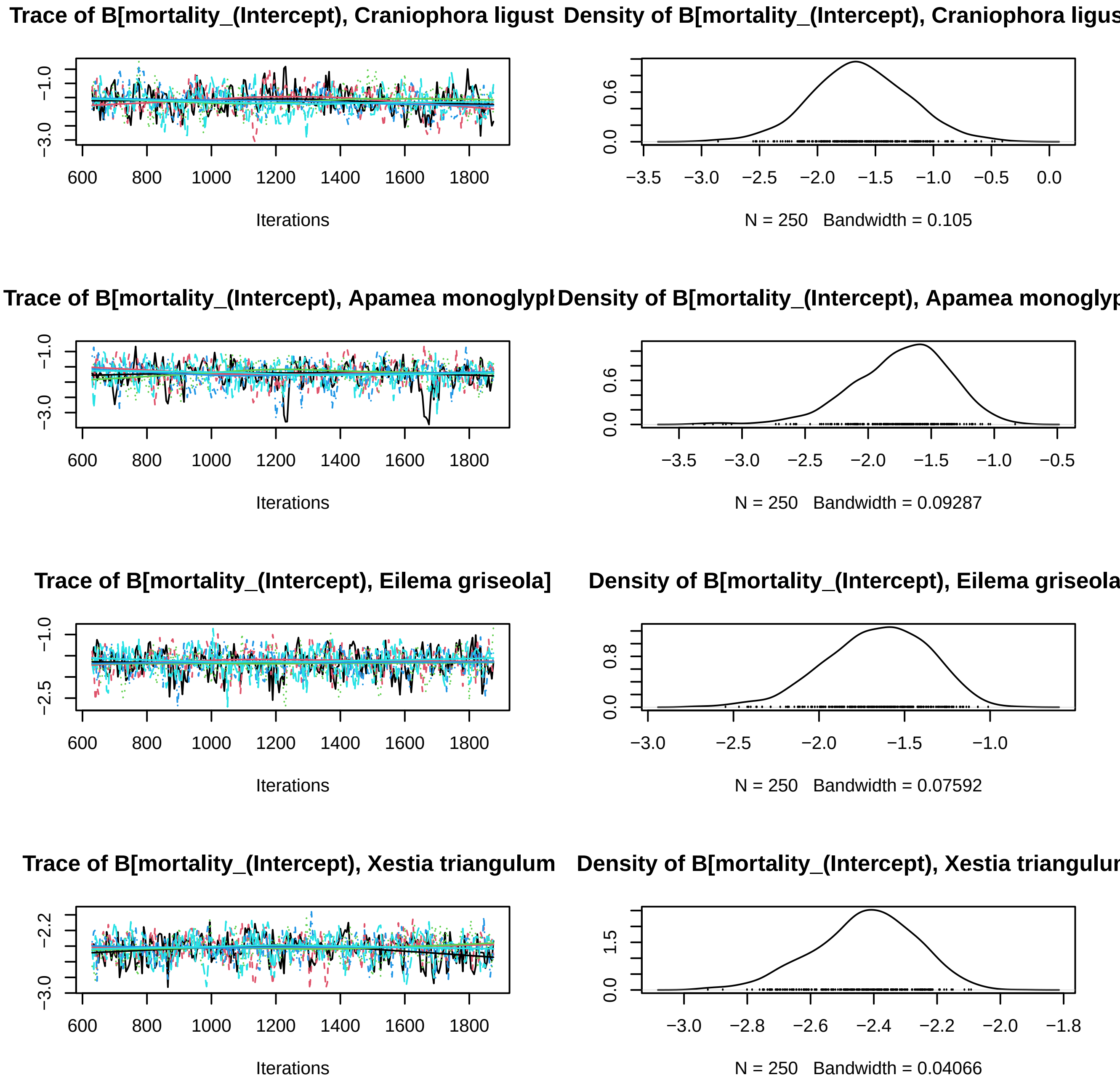

Trace of B[mortality_(Intercept), Craniophora ligust
Density of B[mortality_(Intercept), Craniophora ligus
Iterations
N = 250 Bandwidth = 0.105
Trace of B[mortality_(Intercept), Apamea monoglypl
Density of B[mortality_(Intercept), Apamea monoglyp
Iterations
N = 250 Bandwidth = 0.09287
Trace of B[mortality_(Intercept), Eilema griseola]
Density of B[mortality_(Intercept), Eilema griseola
Iterations
N = 250 Bandwidth = 0.07592
Trace of B[mortality_(Intercept), Xestia triangulum
Density of B[mortality_(Intercept), Xestia triangulur
Iterations
N = 250 Bandwidth = 0.04066

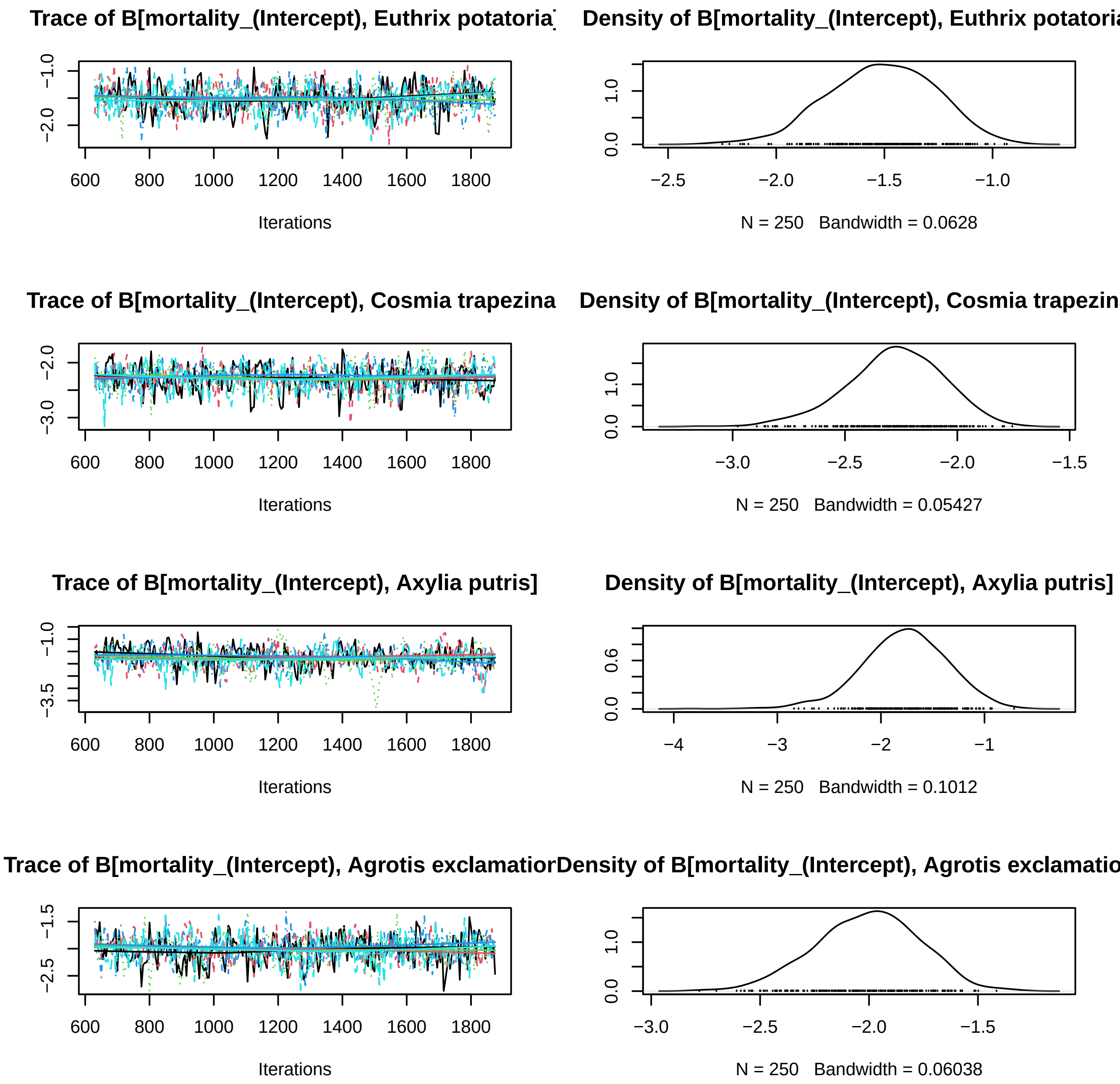
Trace of B[mortality_(Intercept), Euthrix potatoria]
Density of B[mortality_(Intercept), Euthrix potatoria
Iterations
N = 250 Bandwidth = 0.0628
Trace of B[mortality_(Intercept), Cosmia trapezina
Density of B[mortality_(Intercept), Cosmia trapezin
Iterations
N = 250 Bandwidth = 0.05427
Trace of B[mortality_(Intercept), Axylia putris]
Density of B[mortality_(Intercept), Axylia putris]
Iterations
N = 250 Bandwidth = 0.1012
Trace of B[mortality_(Intercept), Agrotis exclamatior
Density of B[mortality_(Intercept), Agrotis exclamatio
Iterations
N = 250 Bandwidth = 0.06038

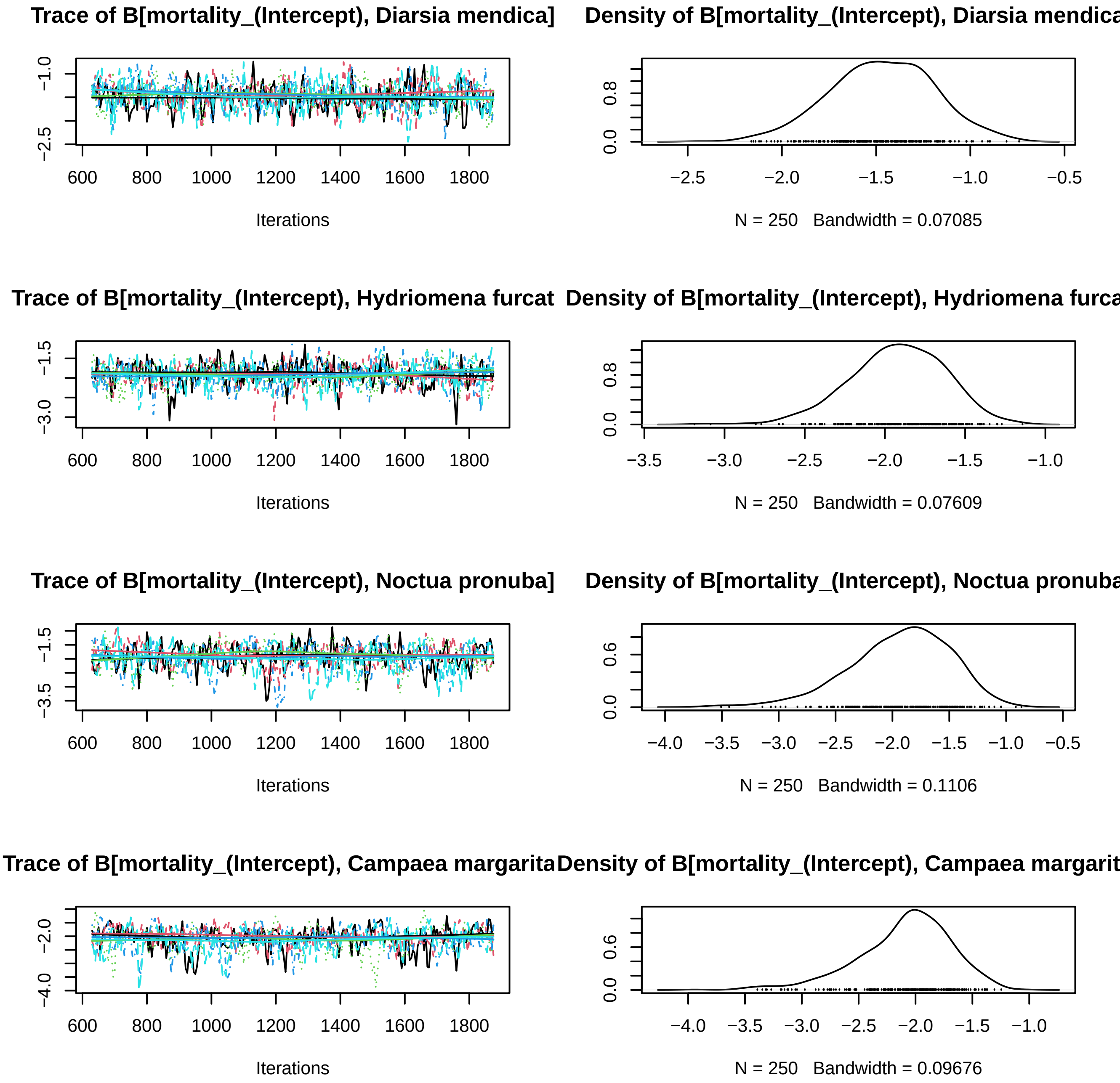
Trace of B[mortality_(Intercept), Diarsia mendica]
Density of B[mortality_(Intercept), Diarsia mendica
Iterations
N = 250 Bandwidth = 0.07085
Trace of B[mortality_(Intercept), Hydriomena furcat
Density of B[mortality_(Intercept), Hydriomena furca
Iterations
N = 250 Bandwidth = 0.07609
Trace of B[mortality_(Intercept), Noctua pronuba]
Density of B[mortality_(Intercept), Noctua pronuba
Iterations
N = 250 Bandwidth = 0.1106
Trace of B[mortality_(Intercept), Campaea margarita
Density of B[mortality_(Intercept), Campaea margarit
Iterations
N = 250 Bandwidth = 0.09676

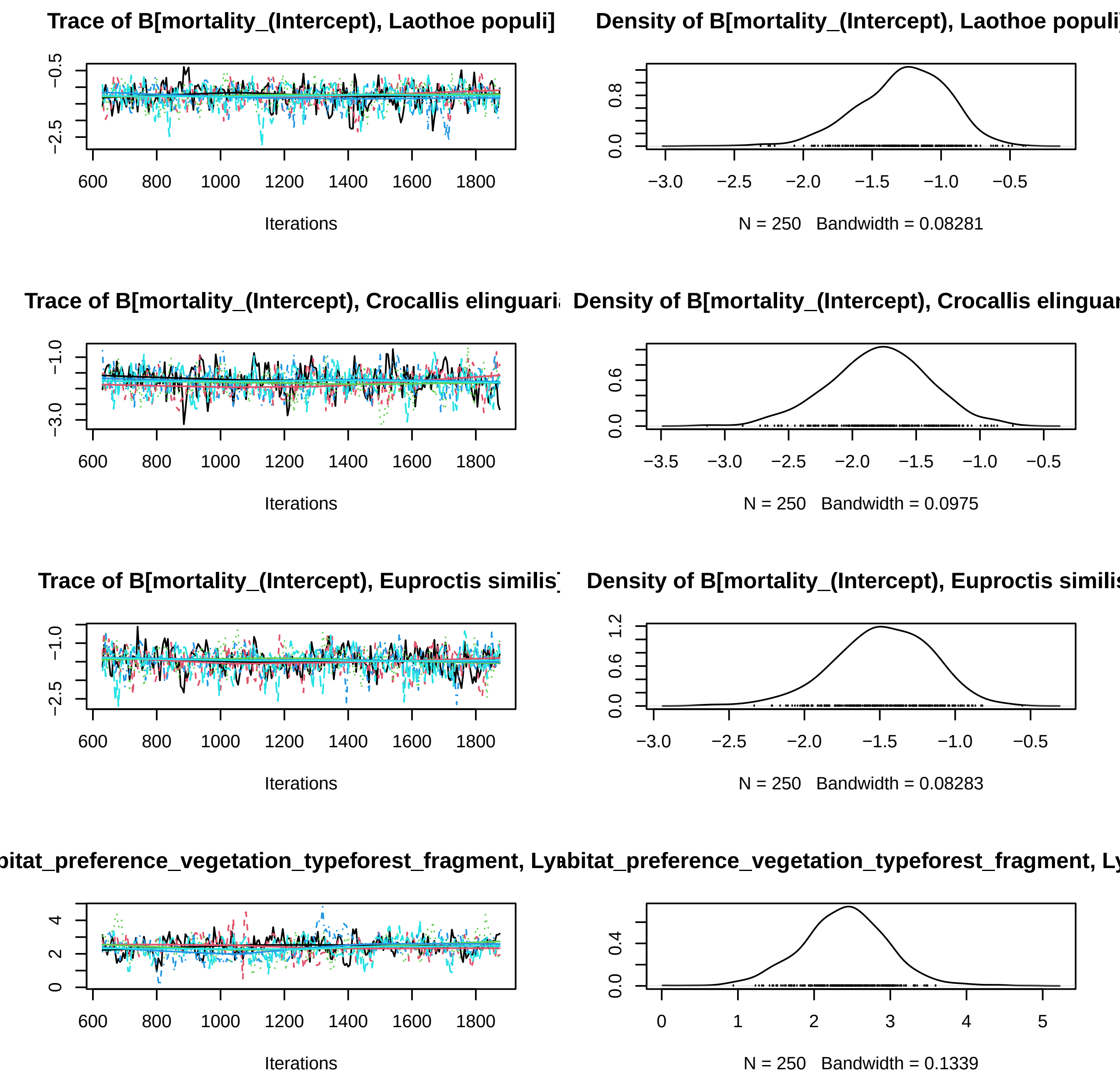
Trace of B[mortality_(Intercept), Laothoe populi]
Density of B[mortality_(Intercept), Laothoe populi
Iterations
N = 250 Bandwidth = 0.08281
Trace of B[mortality_(Intercept), Crocallis elinguari
Density of B[mortality_(Intercept), Crocallis elinguar
Iterations
N = 250 Bandwidth = 0.0975
Trace of B[mortality_(Intercept), Euproctis similis
Density of B[mortality_(Intercept), Euproctis similis
Iterations
N = 250 Bandwidth = 0.08283
bitat_preference_vegetation_typeforest_fragment, Ly
abitat_preference_vegetation_typeforest_fragment, Ly
Iterations
N = 250 Bandwidth = 0.1339

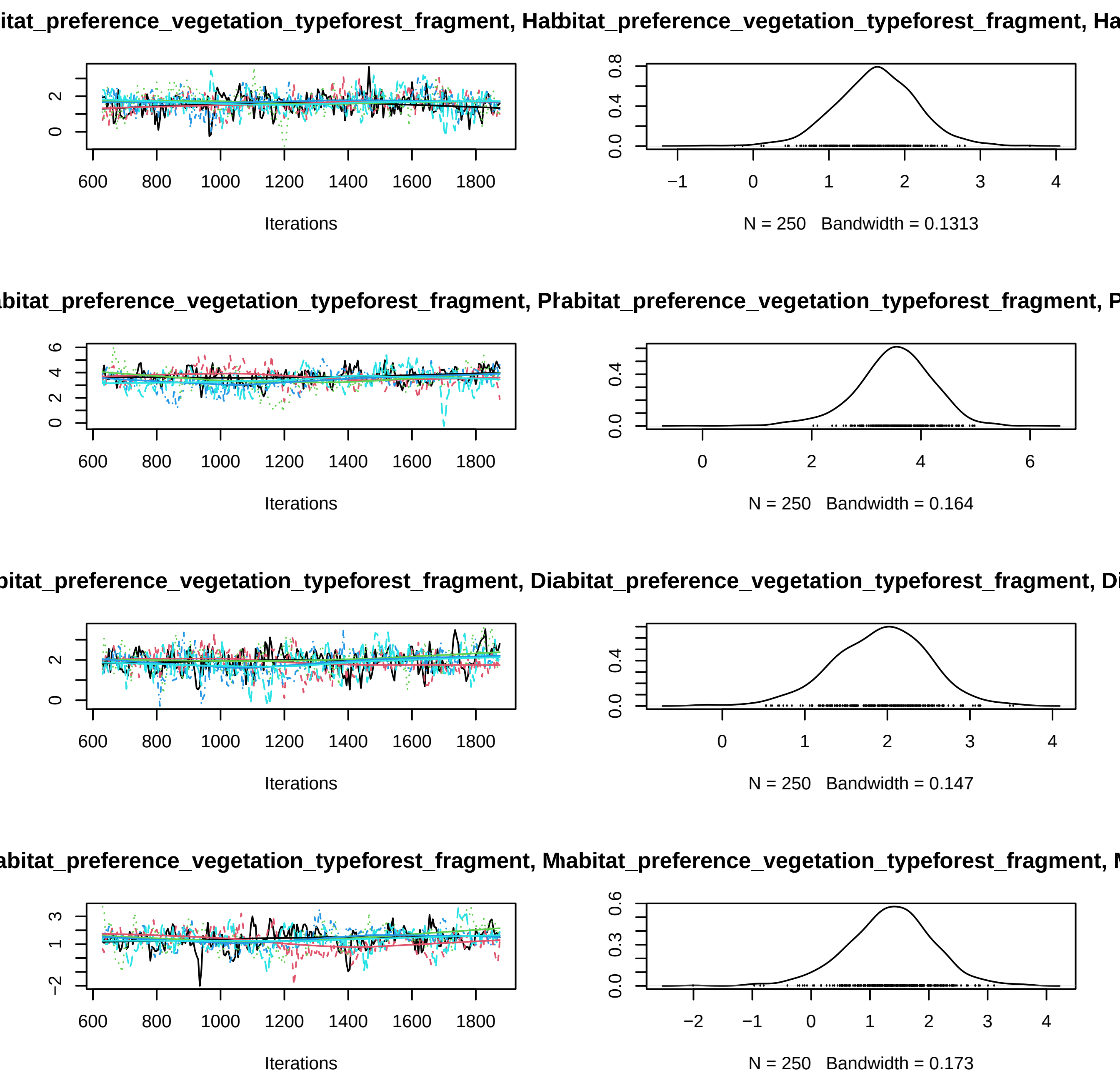
itat_preference_vegetation_typeforest_fragment, Habitat_preference_vegetation_typeforest_fragment, Ha
Iterations
N = 250 Bandwidth = 0.1313
abitat_preference_vegetation_typeforest_fragment, Phabitat_preference_vegetation_typeforest_fragment, P
Iterations
N = 250 Bandwidth = 0.164
bitat_preference_vegetation_typeforest_fragment, Diabitat_preference_vegetation_typeforest_fragment, Di
Iterations
N = 250 Bandwidth = 0.147
abitat_preference_vegetation_typeforest_fragment, Mhabitat_preference_vegetation_typeforest_fragment, M
Iterations
N = 250 Bandwidth = 0.173

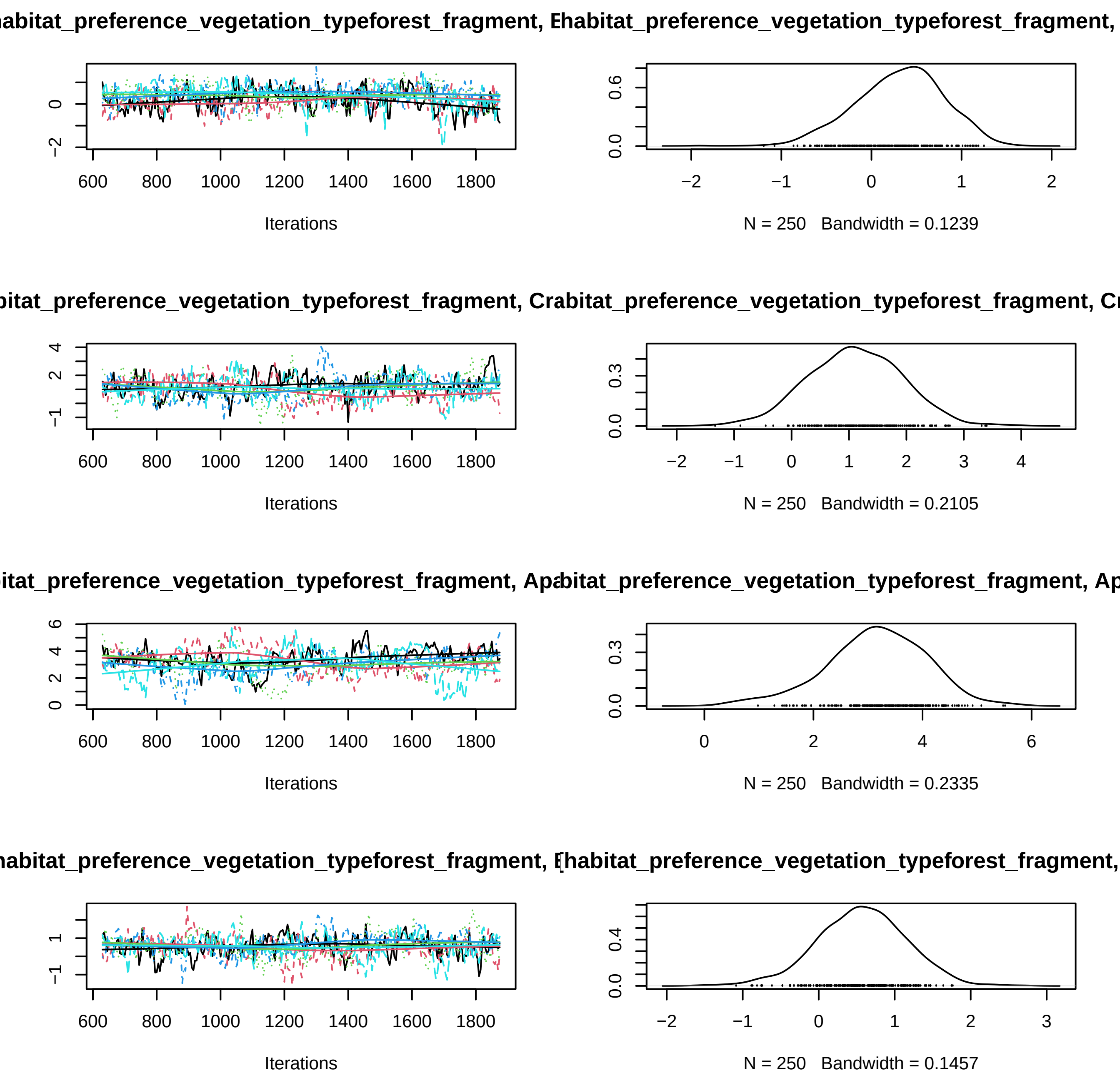

abitat_preference_vegetation_typeforest_fragment, E
habitat_preference_vegetation_typeforest_fragment,
Iterations
N = 250 Bandwidth = 0.1239
bitat_preference_vegetation_typeforest_fragment, Cr
abitat_preference_vegetation_typeforest_fragment, Cr
Iterations
N = 250 Bandwidth = 0.2105
bitat_preference_vegetation_typeforest_fragment, Ap
abitat_preference_vegetation_typeforest_fragment, Ap
Iterations
N = 250 Bandwidth = 0.2335
abitat_preference_vegetation_typeforest_fragment, E
habitat_preference_vegetation_typeforest_fragment,
Iterations
N = 250 Bandwidth = 0.1457

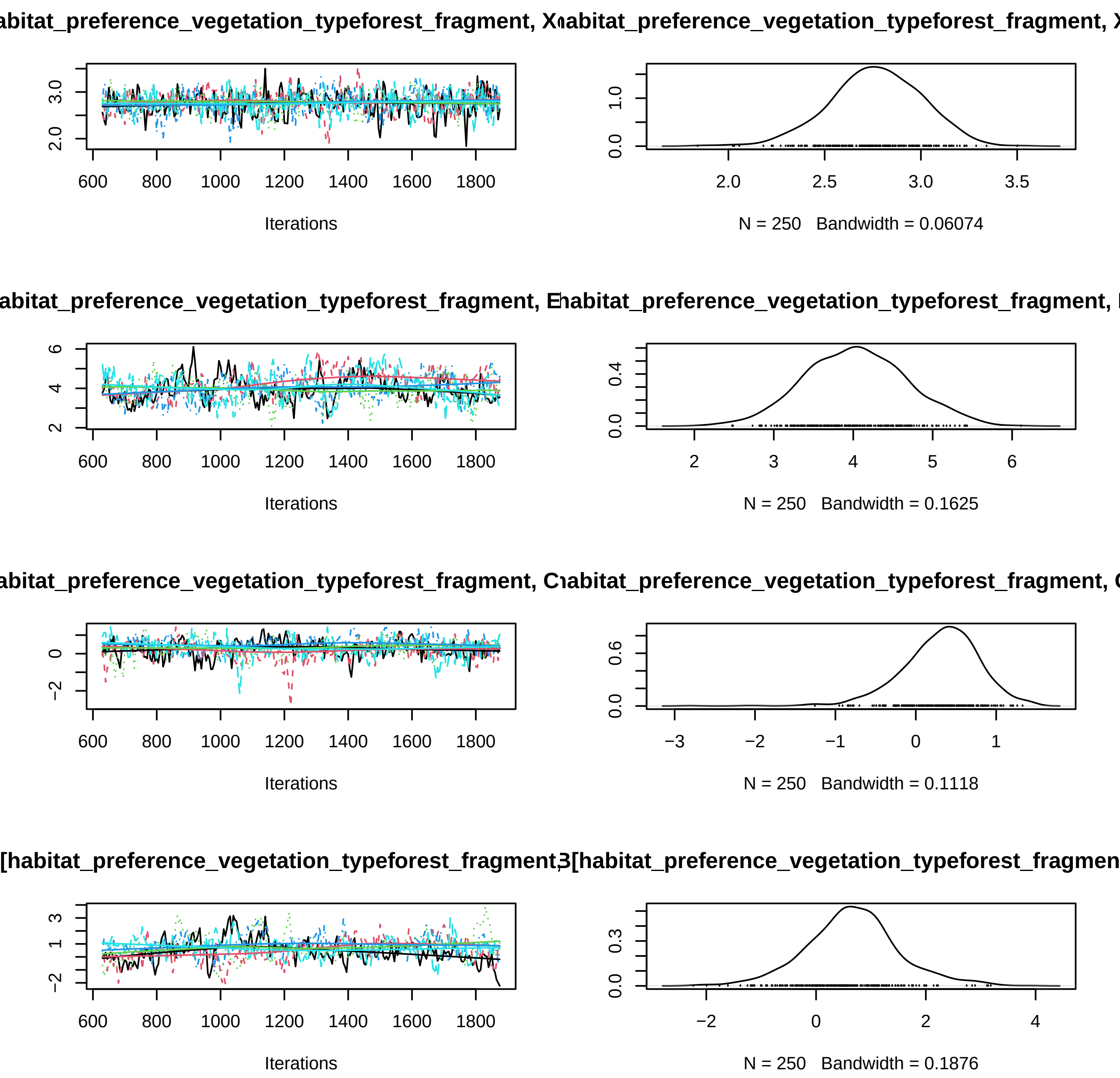
abitat_preference_vegetation_typeforest_fragment, X
abitat_preference_vegetation_typeforest_fragment, X
Iterations
N = 250 Bandwidth = 0.06074
abitat_preference_vegetation_typeforest_fragment, E
habitat_preference_vegetation_typeforest_fragment, I
Iterations
N = 250 Bandwidth = 0.1625
abitat_preference_vegetation_typeforest_fragment, C
abitat_preference_vegetation_typeforest_fragment, C
Iterations
N = 250 Bandwidth = 0.1118
[habitat_preference_vegetation_typeforest_fragment,
3[habitat_preference_vegetation_typeforest_fragmen
Iterations
N = 250 Bandwidth = 0.1876

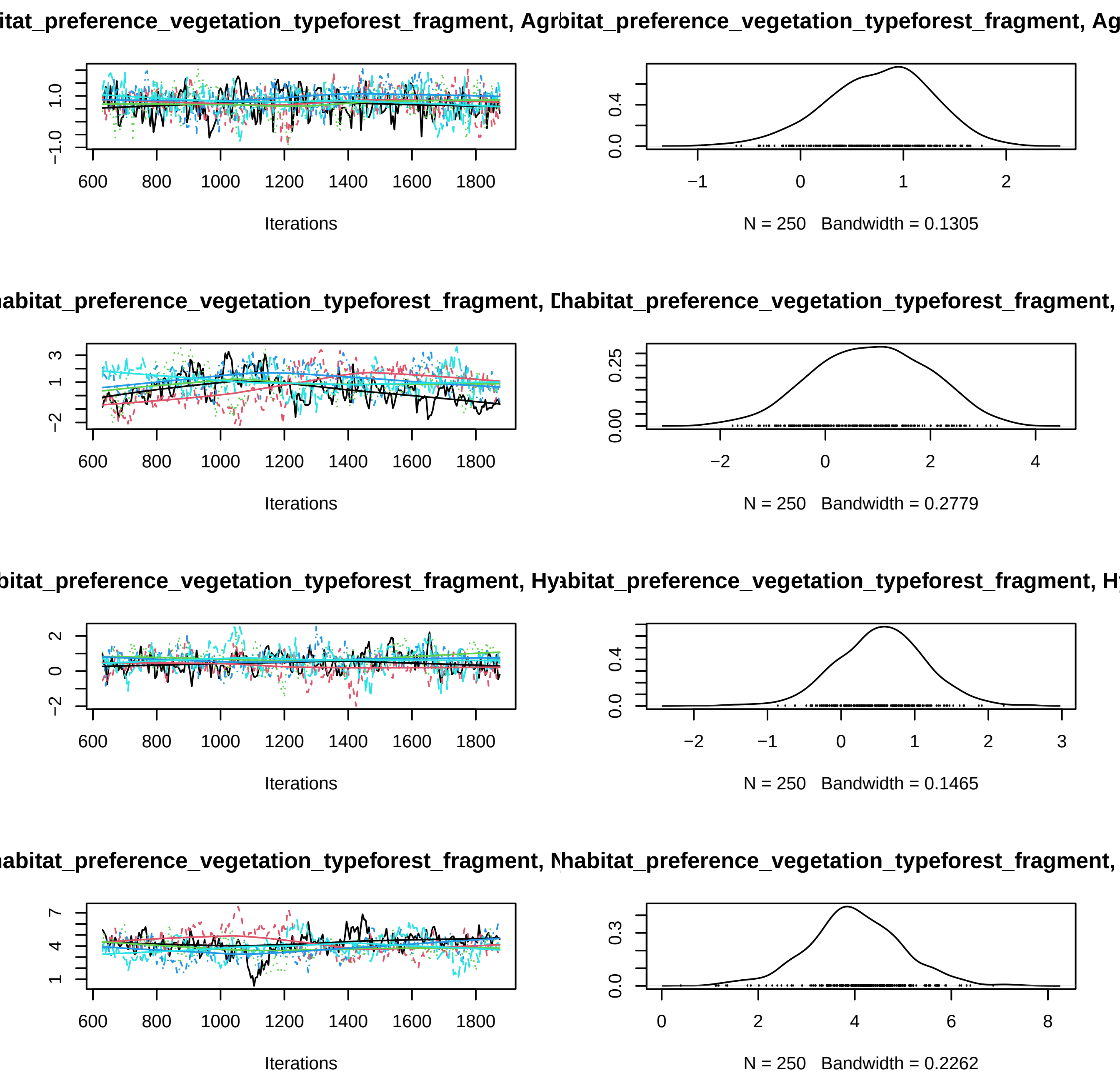
itat_preference_vegetation_typeforest_fragment, Agr
bitat_preference_vegetation_typeforest_fragment, Ag
Iterations
N = 250 Bandwidth = 0.1305
habitat_preference_vegetation_typeforest_fragment, D
habitat_preference_vegetation_typeforest_fragment,
Iterations
N = 250 Bandwidth = 0.2779
bitat_preference_vegetation_typeforest_fragment, Hy
abitat_preference_vegetation_typeforest_fragment, H
Iterations
N = 250 Bandwidth = 0.1465
habitat_preference_vegetation_typeforest_fragment, N
habitat_preference_vegetation_typeforest_fragment,
Iterations
N = 250 Bandwidth = 0.2262

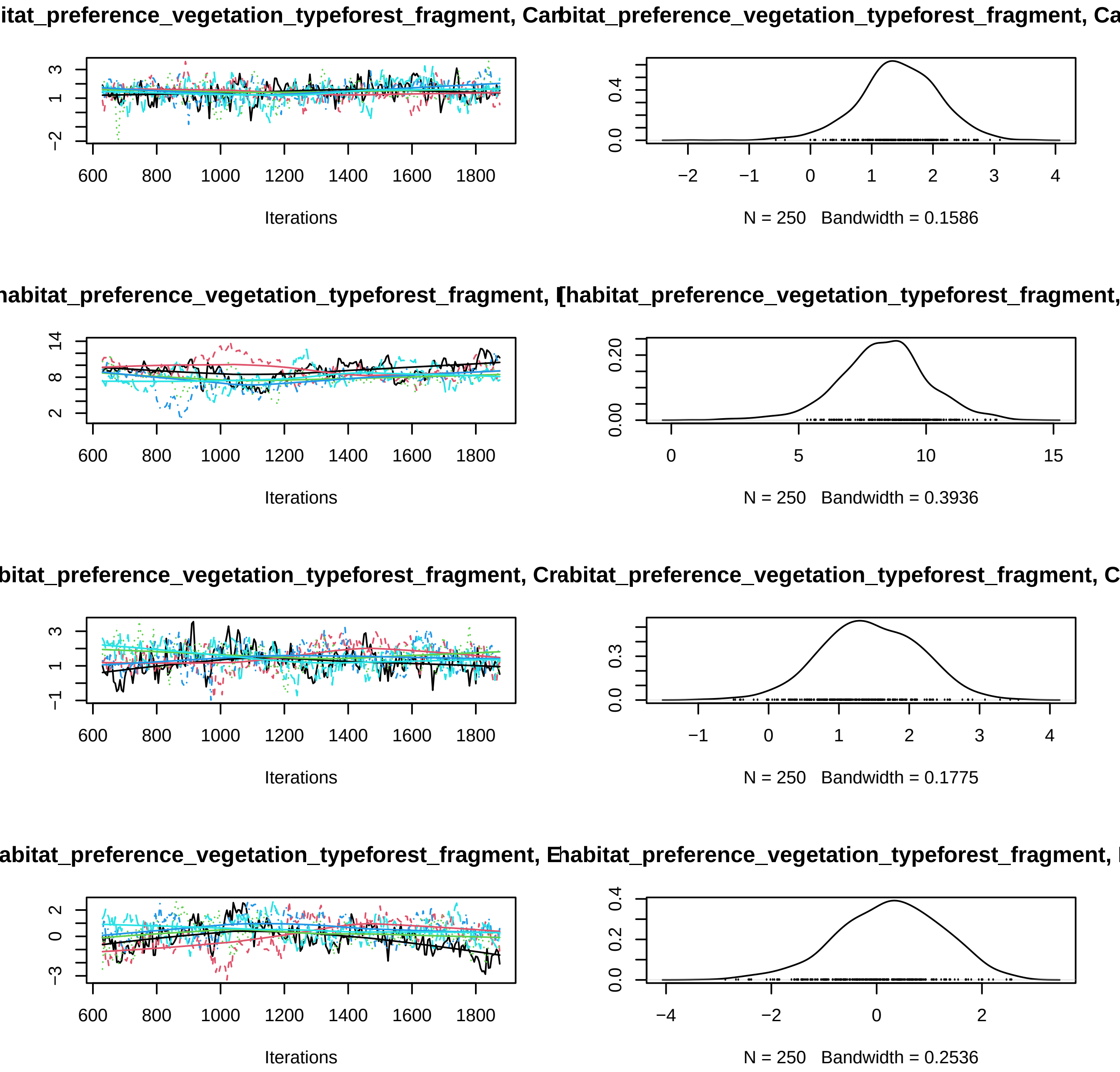
itat_preference_vegetation_typeforest_fragment, Car
bitat_preference_vegetation_typeforest_fragment, Ca
Iterations
N = 250 Bandwidth = 0.1586
habitat_preference_vegetation_typeforest_fragment, I
[habitat_preference_vegetation_typeforest_fragment,
Iterations
N = 250 Bandwidth = 0.3936
bitat_preference_vegetation_typeforest_fragment, Cr
abitat_preference_vegetation_typeforest_fragment, C
Iterations
N = 250 Bandwidth = 0.1775
abitat_preference_vegetation_typeforest_fragment, E
habitat_preference_vegetation_typeforest_fragment, I
Iterations
N = 250 Bandwidth = 0.2536

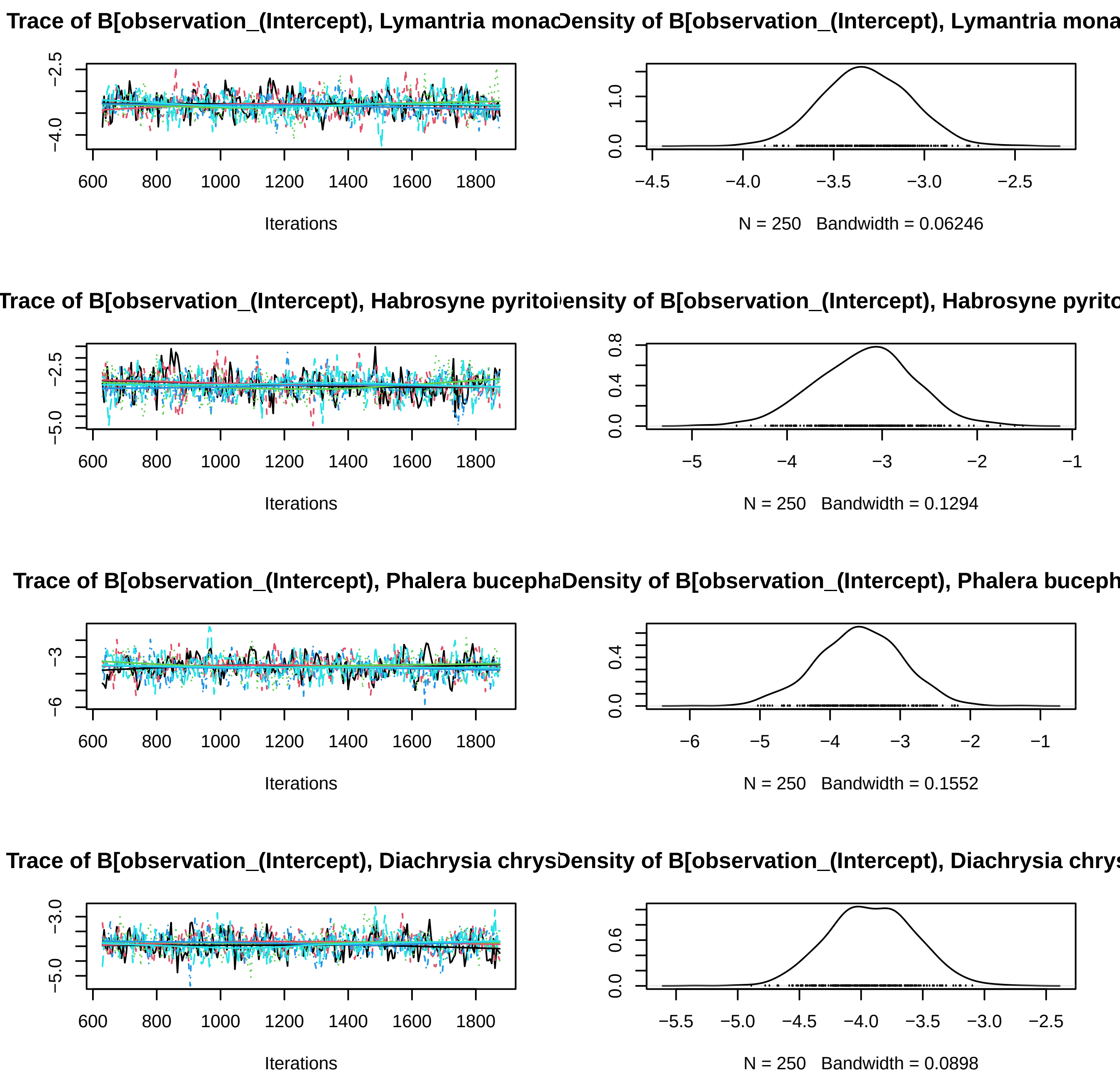
Trace of B[observation_(Intercept), Lymantria monac
Density of B[observation_(Intercept), Lymantria mona
Iterations
N = 250 Bandwidth = 0.06246
Trace of B[observation_(Intercept), Habrosyne pyritoi
Density of B[observation_(Intercept), Habrosyne pyrito
Iterations
N = 250 Bandwidth = 0.1294
Trace of B[observation_(Intercept), Phalera bucepha
Density of B[observation_(Intercept), Phalera buceph
Iterations
N = 250 Bandwidth = 0.1552
Trace of B[observation_(Intercept), Diachrysia chrys
Density of B[observation_(Intercept), Diachrysia chrys
Iterations
N = 250 Bandwidth = 0.0898

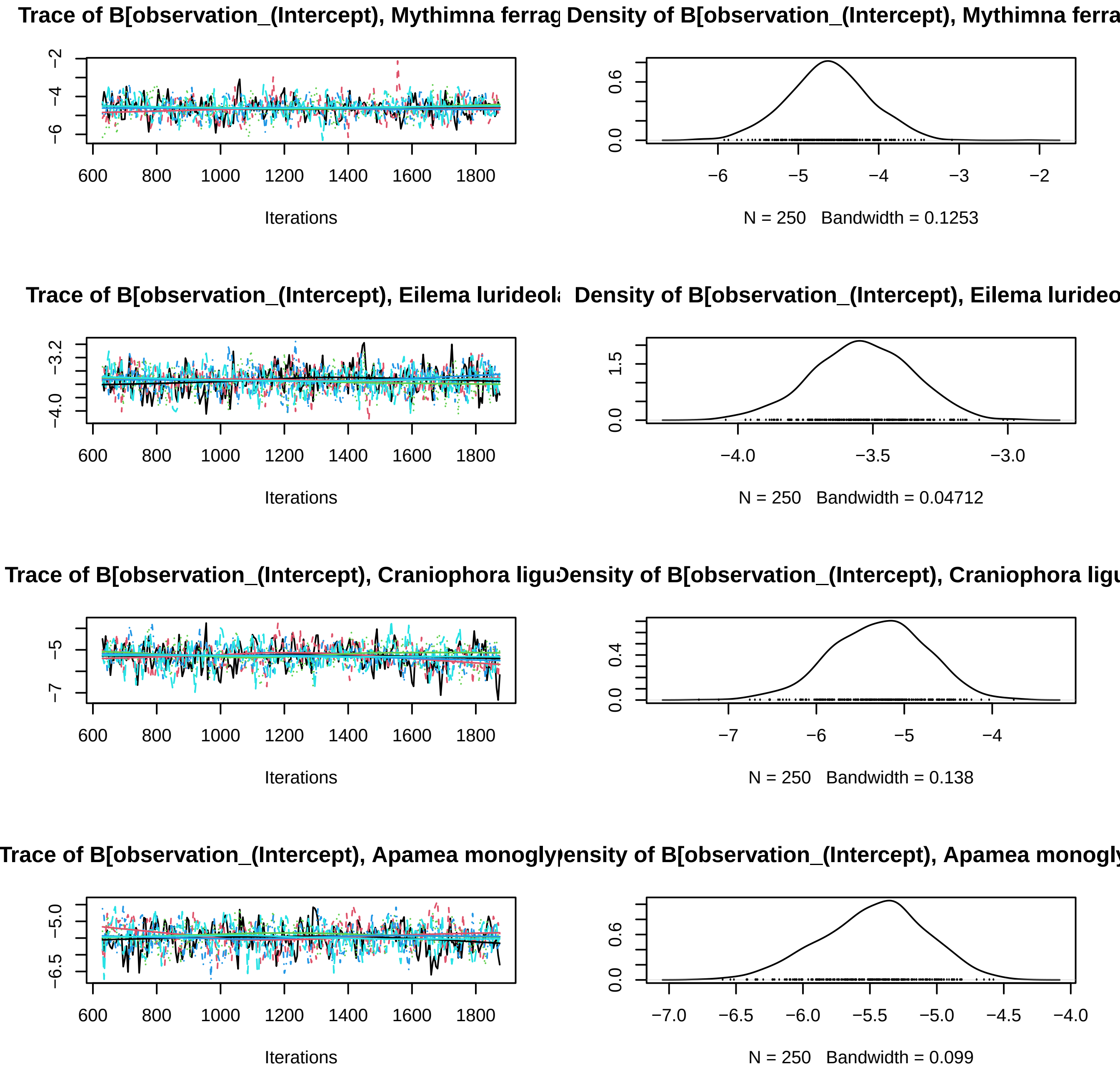
Trace of B[observation_(Intercept), Mythimna ferrag
Density of B[observation_(Intercept), Mythimna ferra
Iterations
N = 250 Bandwidth = 0.1253
Trace of B[observation_(Intercept), Eilema lurideola
Density of B[observation_(Intercept), Eilema lurideo
Iterations
N = 250 Bandwidth = 0.04712
Trace of B[observation_(Intercept), Craniophora ligu
Density of B[observation_(Intercept), Craniophora ligu
Iterations
N = 250 Bandwidth = 0.138
Trace of B[observation_(Intercept), Apamea monogly
Density of B[observation_(Intercept), Apamea monogly
Iterations
N = 250 Bandwidth = 0.099

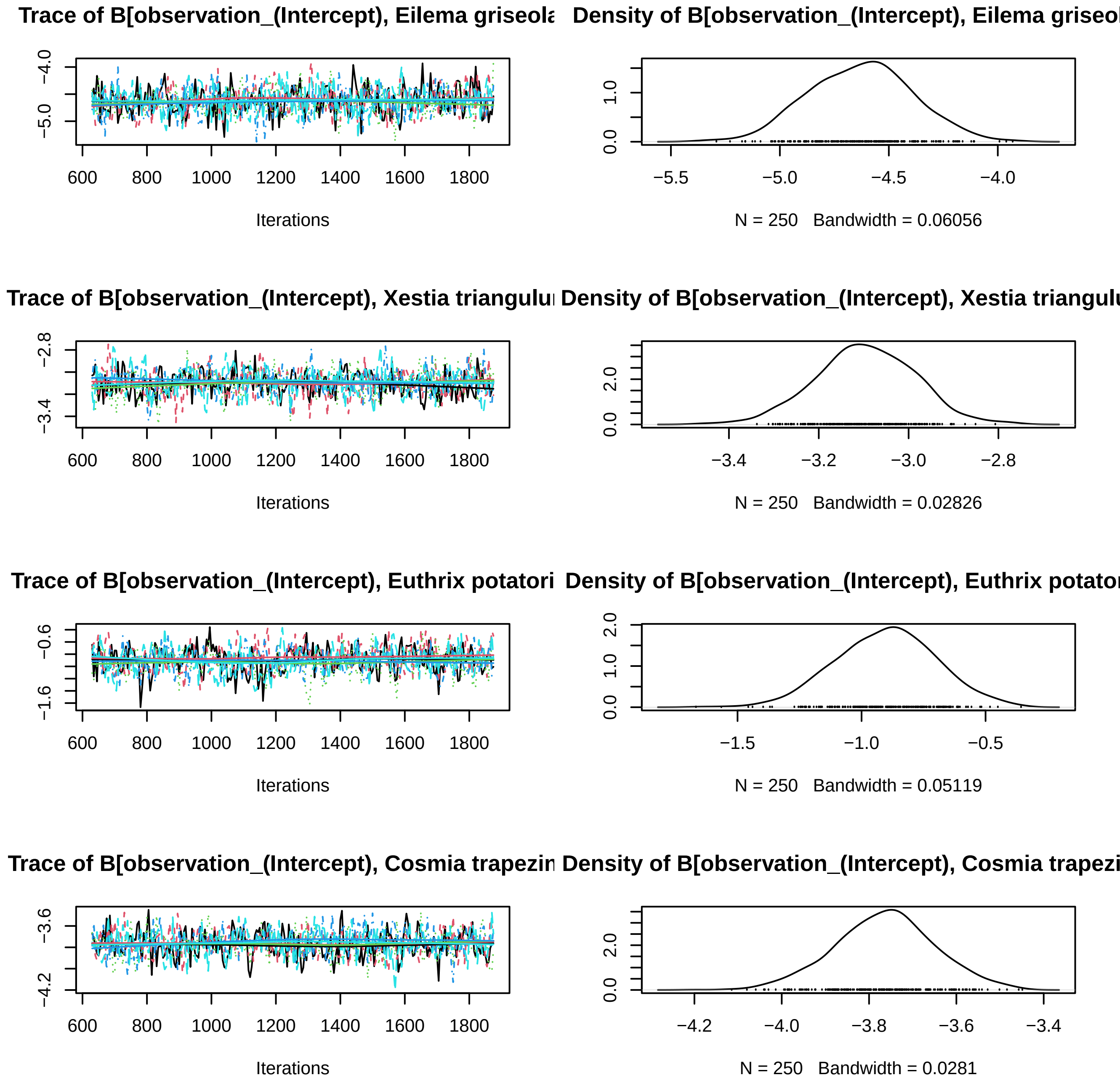

Trace of B[observation_(Intercept), Eilema griseola
Iterations
Density of B[observation_(Intercept), Eilema griseol
N = 250 Bandwidth = 0.06056
Trace of B[observation_(Intercept), Xestia triangulu
Iterations
Density of B[observation_(Intercept), Xestia triangulu
N = 250 Bandwidth = 0.02826
Trace of B[observation_(Intercept), Euthrix potatori
Iterations
Density of B[observation_(Intercept), Euthrix potator
N = 250 Bandwidth = 0.05119
Trace of B[observation_(Intercept), Cosmia trapezin
Iterations
Density of B[observation_(Intercept), Cosmia trapezi
N = 250 Bandwidth = 0.0281

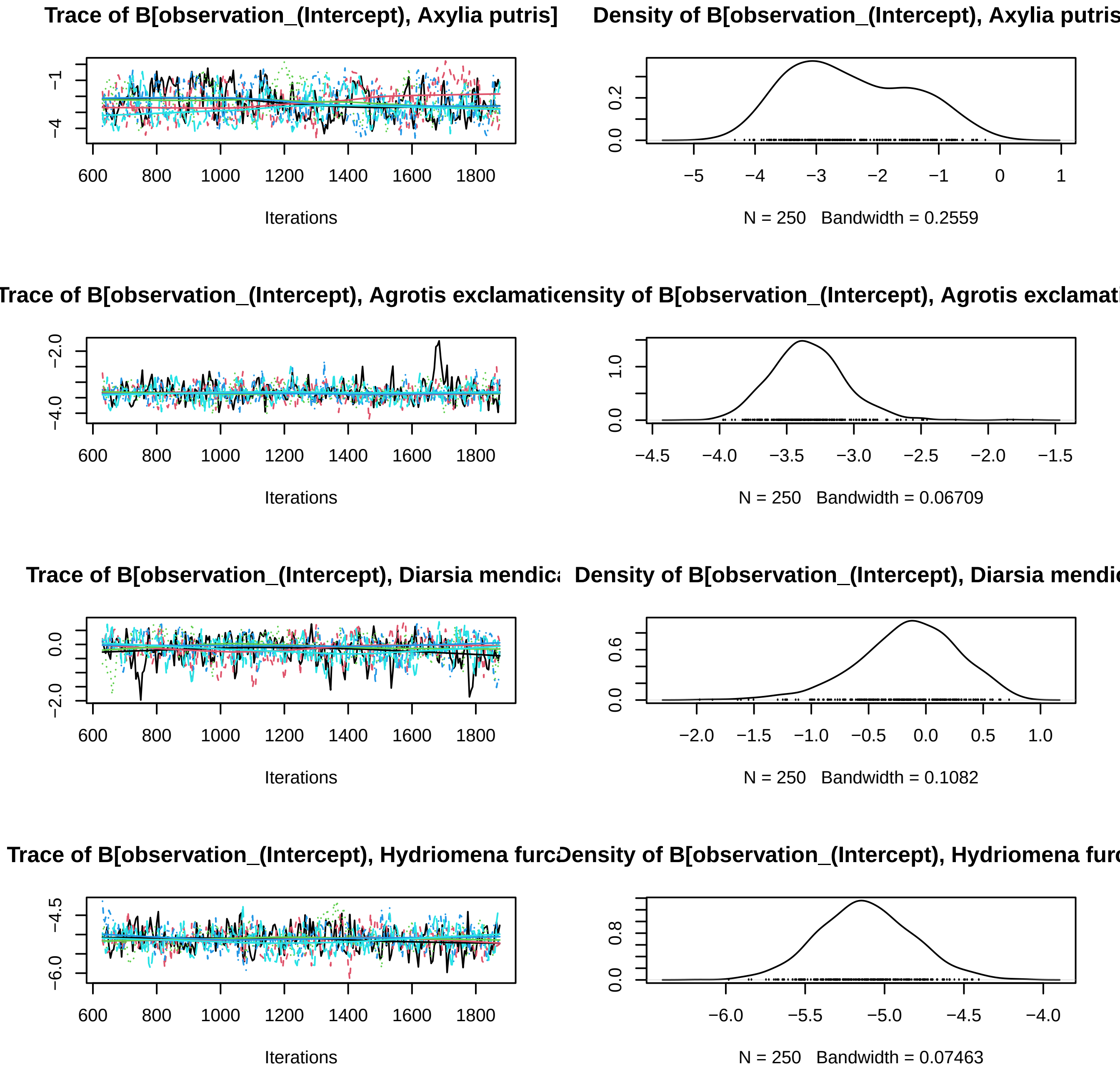

Trace of B[observation_(Intercept), Axylia putris]
Density of B[observation_(Intercept), Axylia putris
Iterations
N = 250 Bandwidth = 0.2559
Trace of B[observation_(Intercept), Agrotis exclamati
ensity of B[observation_(Intercept), Agrotis exclamat
Iterations
N = 250 Bandwidth = 0.06709
Trace of B[observation_(Intercept), Diarsia mendica
Density of B[observation_(Intercept), Diarsia mendic
Iterations
N = 250 Bandwidth = 0.1082
Trace of B[observation_(Intercept), Hydriomena furc
Density of B[observation_(Intercept), Hydriomena furc
Iterations
N = 250 Bandwidth = 0.07463

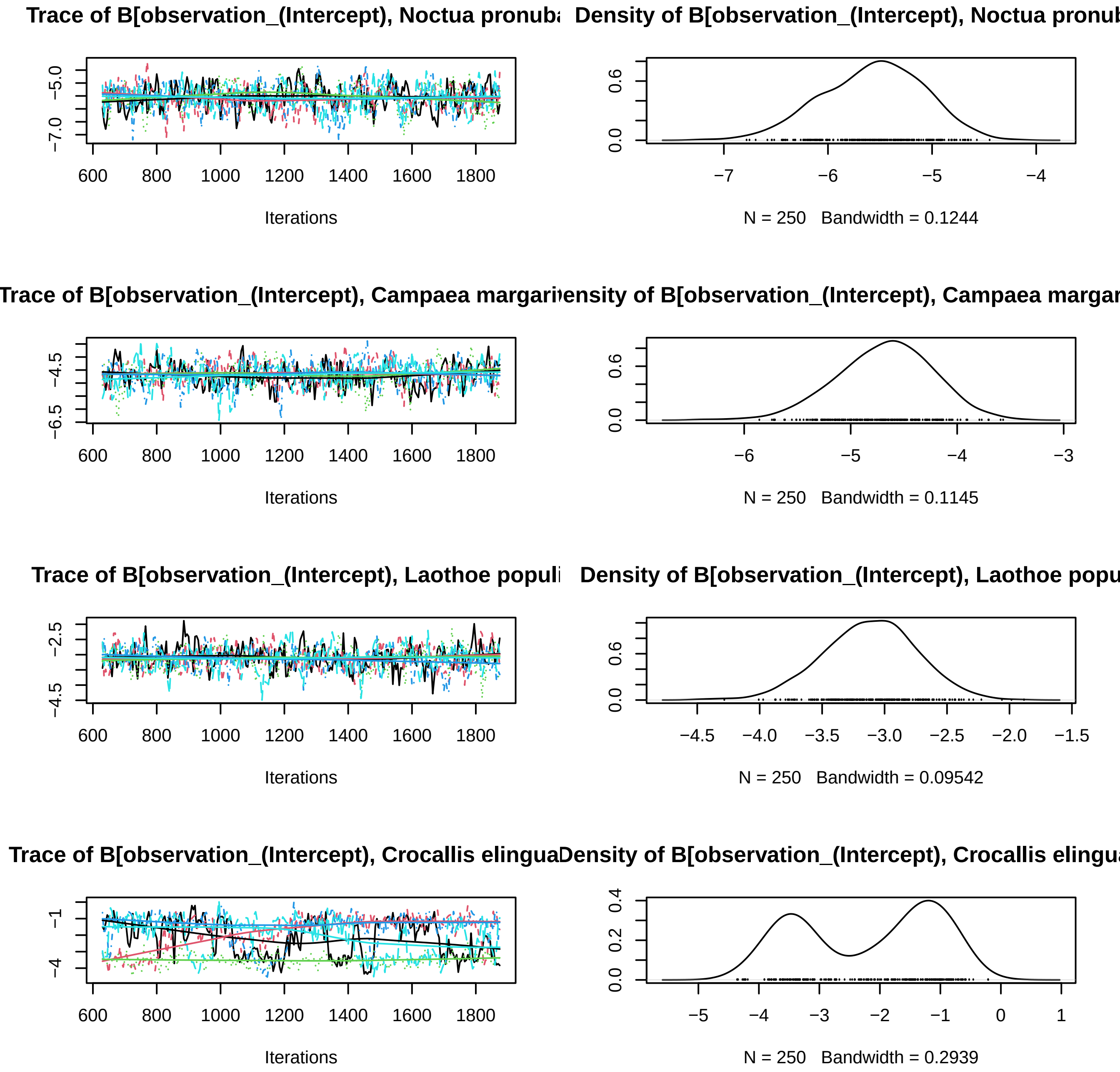
Trace of B[observation_(Intercept), Noctua pronuba
Density of B[observation_(Intercept), Noctua pronub
−5.0
−7.0
600
800
1000
1200
1400
1600
1800
Iterations
0.0
0.6
−7
−6
−5
−4
N = 250 Bandwidth = 0.1244
Trace of B[observation_(Intercept), Campaea margari
Density of B[observation_(Intercept), Campaea margar
−4.5
−6.5
600
800
1000
1200
1400
1600
1800
Iterations
0.0
0.6
−6
−5
−4
−3
N = 250 Bandwidth = 0.1145
Trace of B[observation_(Intercept), Laothoe populi
Density of B[observation_(Intercept), Laothoe popu
−2.5
−4.5
600
800
1000
1200
1400
1600
1800
Iterations
0.0
0.6
−4.5
−4.0
−3.5
−3.0
−2.5
−2.0
−1.5
N = 250 Bandwidth = 0.09542
Trace of B[observation_(Intercept), Crocallis elingua
Density of B[observation_(Intercept), Crocallis elingu
−1
−4
600
800
1000
1200
1400
1600
1800
Iterations
0.0
0.2
0.4
−5
−4
−3
−2
−1
0
1
N = 250 Bandwidth = 0.2939

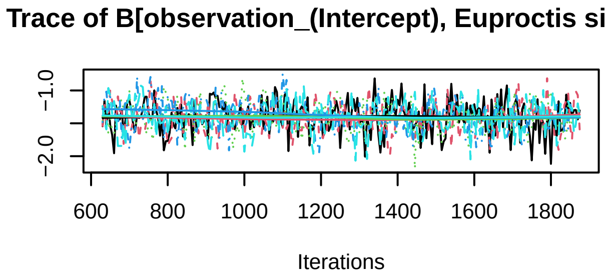
Trace of B[observation_(Intercept), Euproctis simili
−2.0
−1.0
600
800
1000
1200
1400
1600
1800
Iterations

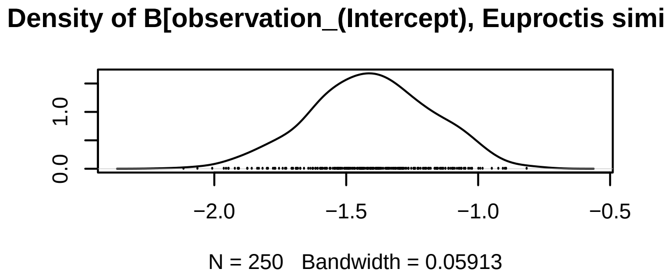
Density of B[observation_(Intercept), Euproctis simi
0.0
1.0
−2.0
−1.5
−1.0
−0.5
N = 250   Bandwidth = 0.05913

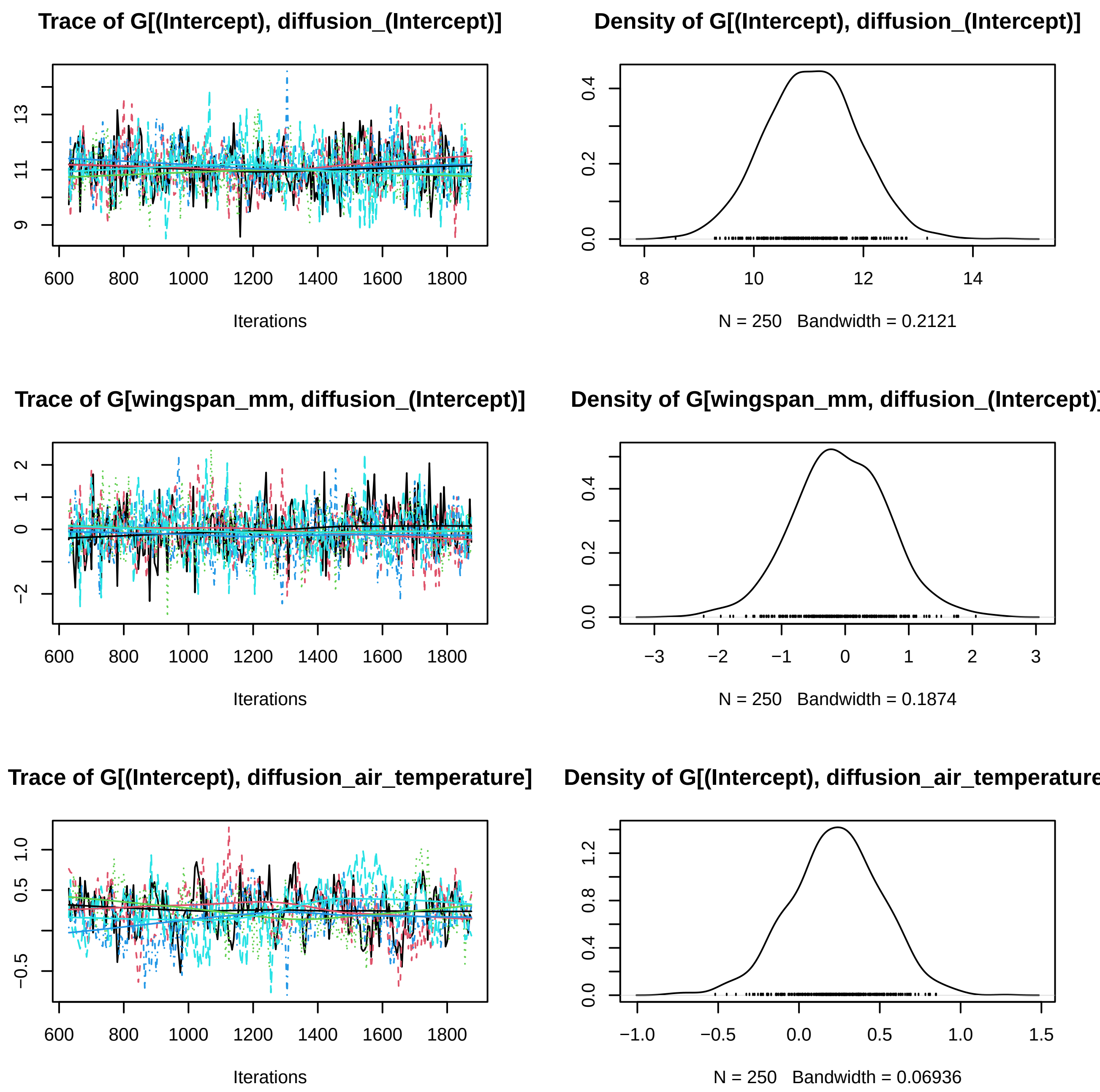
Trace of G[(Intercept), diffusion_(Intercept)]
Iterations
Density of G[(Intercept), diffusion_(Intercept)]
N = 250 Bandwidth = 0.2121
Trace of G[wingspan_mm, diffusion_(Intercept)]
Iterations
Density of G[wingspan_mm, diffusion_(Intercept)
N = 250 Bandwidth = 0.1874
Trace of G[(Intercept), diffusion_air_temperature]
Iterations
Density of G[(Intercept), diffusion_air_temperature
N = 250 Bandwidth = 0.06936

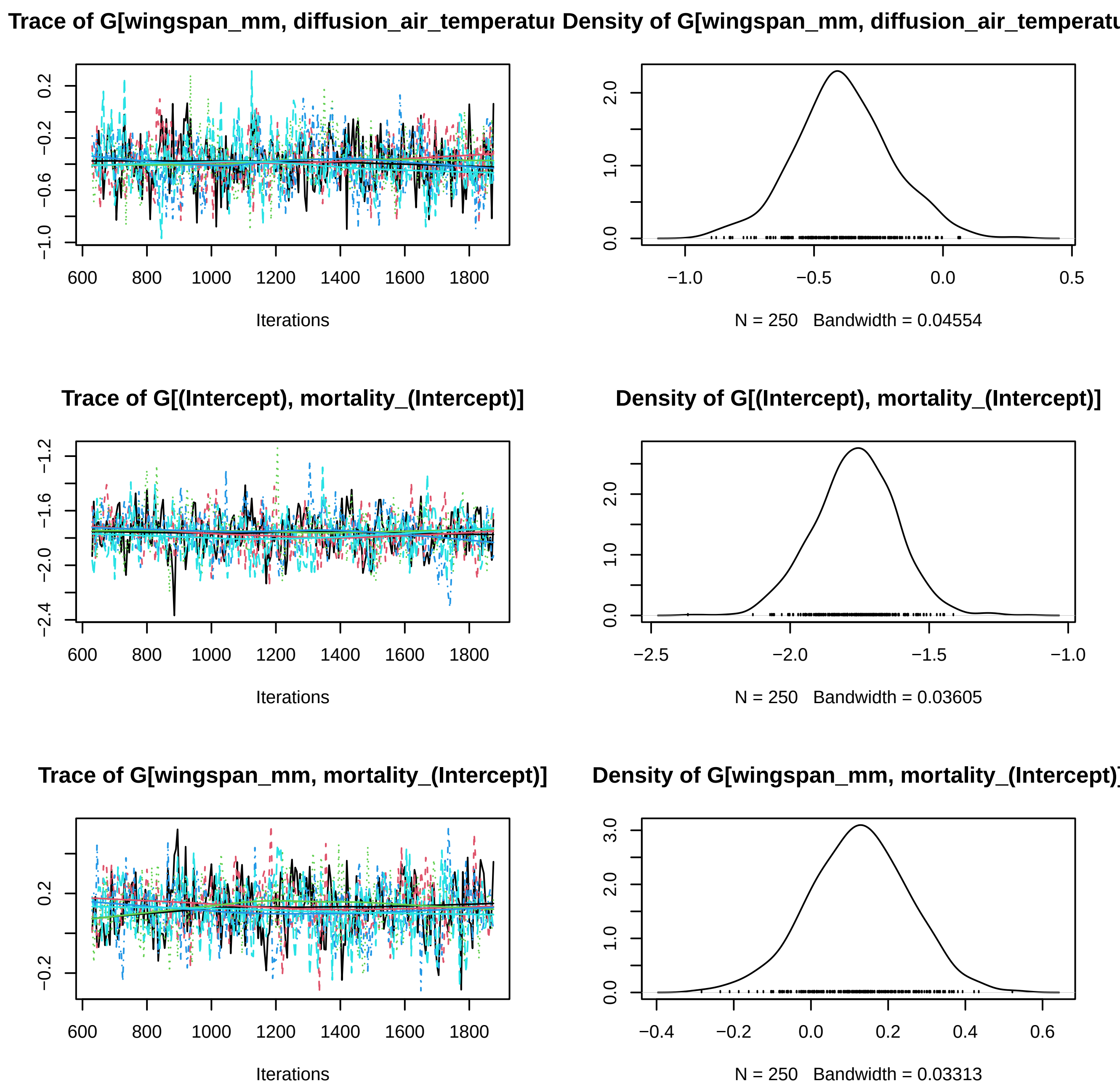
Trace of G[wingspan_mm, diffusion_air_temperatu
Density of G[wingspan_mm, diffusion_air_temperatu
Iterations
N = 250 Bandwidth = 0.04554
Trace of G[(Intercept), mortality_(Intercept)]
Density of G[(Intercept), mortality_(Intercept)]
Iterations
N = 250 Bandwidth = 0.03605
Trace of G[wingspan_mm, mortality_(Intercept)]
Density of G[wingspan_mm, mortality_(Intercept)
Iterations
N = 250 Bandwidth = 0.03313

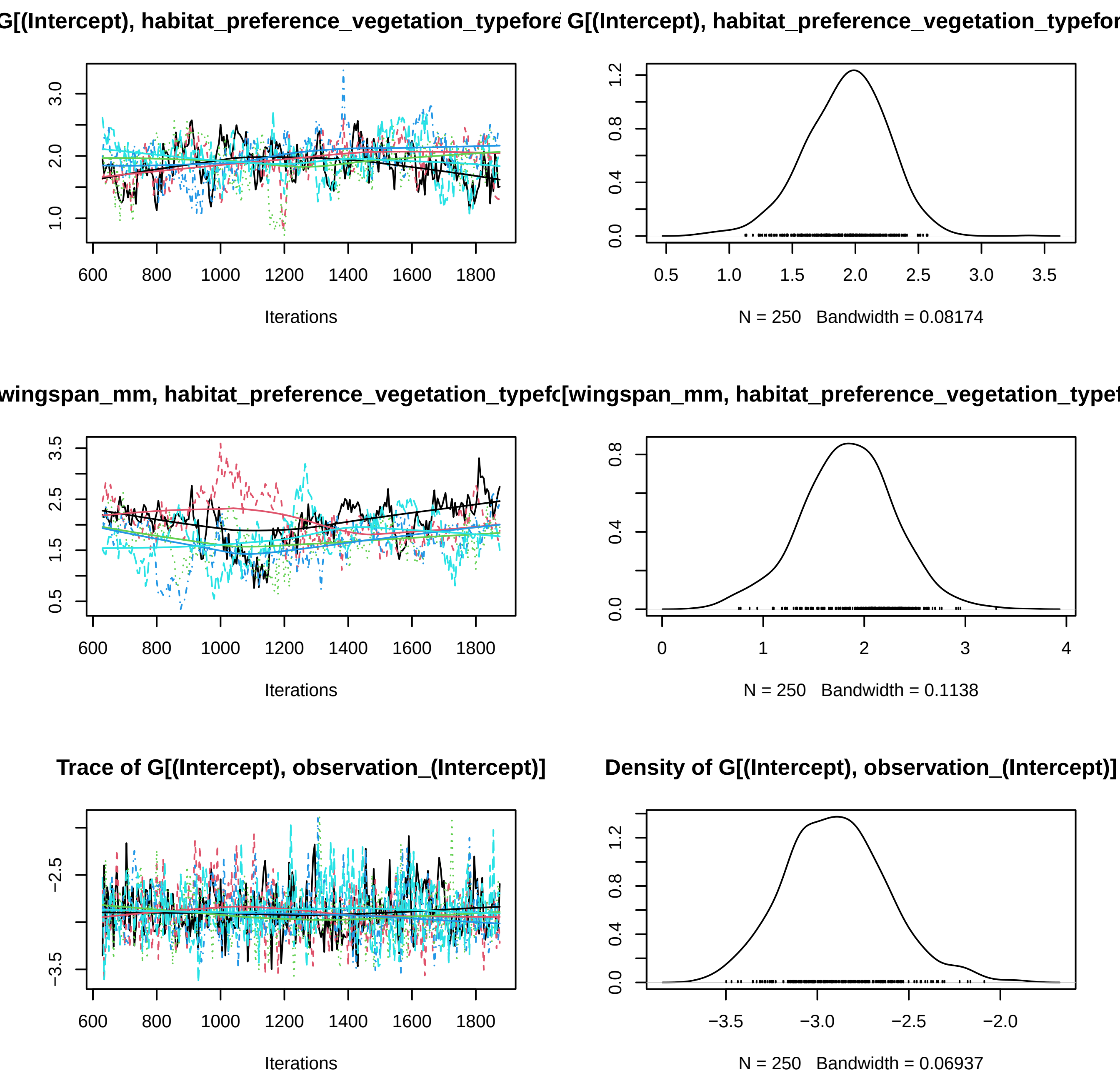

G[(Intercept), habitat_preference_vegetation_typefore
G[(Intercept), habitat_preference_vegetation_typefor
Iterations
N = 250 Bandwidth = 0.08174
wingspan_mm, habitat_preference_vegetation_typefc
[wingspan_mm, habitat_preference_vegetation_typef
Iterations
N = 250 Bandwidth = 0.1138
Trace of G[(Intercept), observation_(Intercept)]
Density of G[(Intercept), observation_(Intercept)]
Iterations
N = 250 Bandwidth = 0.06937

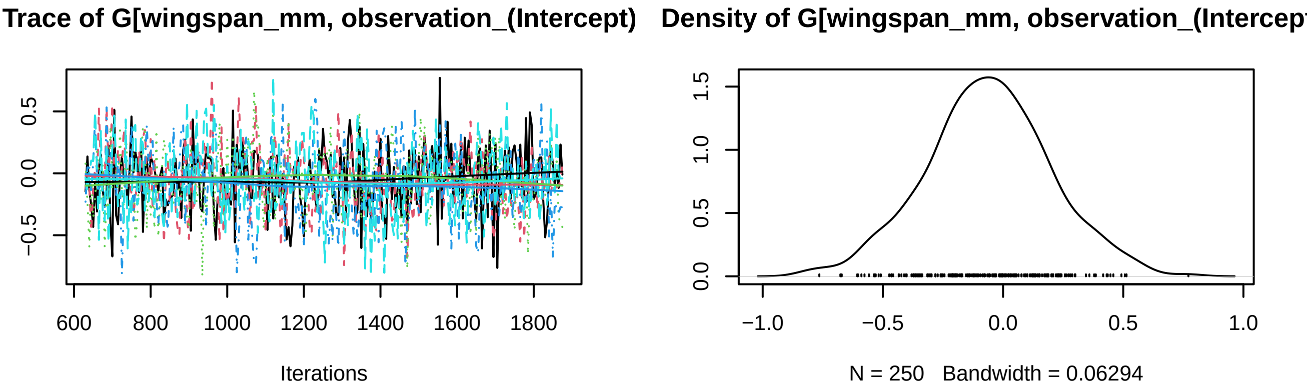
Trace of G[wingspan_mm, observation_(Intercept)
Density of G[wingspan_mm, observation_(Intercep
0.5
0.0
−0.5
600
800
1000
1200
1400
1600
1800
Iterations
1.5
1.0
0.5
0.0
−1.0
−0.5
0.0
0.5
1.0
N = 250   Bandwidth = 0.06294

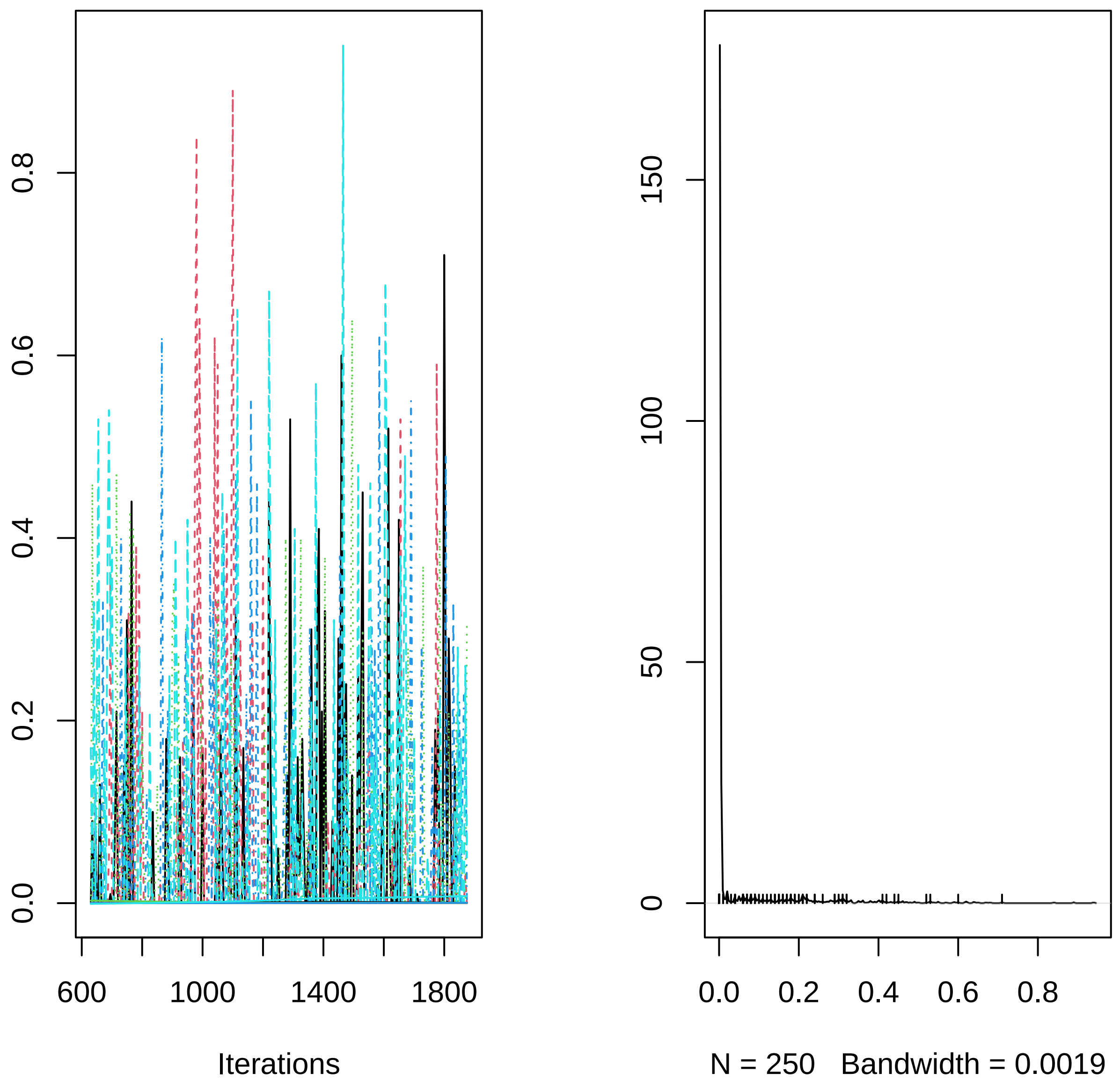

0.0
0.2
0.4
0.6
0.8
600
1000
1400
1800
Iterations
0
50
100
150
0.0
0.2
0.4
0.6
0.8
N = 250   Bandwidth = 0.0019
