## Supplementary material for "R-package Jsmm: Joint species movement modelling of mark-recapture data": S2 Vignettes demonstrating the application of the R-package Jsmm: 1a_define_model_simulated.R

```
library(Jsmm)
library(mapview)
library(ggcorrplot)

set.seed(3)

#---------------------------------------------------------------------------
# CREATE domain AND ITS TRIANGULATION
#---------------------------------------------------------------------------

p = c(2.5/15, 7.5/15, 12.5/15)

lc = as.matrix(expand.grid("x" = p, "y" = p)) # Positions capture and release sites
lc_label = 1:nrow(lc)

buffer_x = c(2.5/15, 2.5/15)
buffer_y = c(2.5/15, 2.5/15)

nnod_side_x = 16 # number of nodes per side x
nnod_side_y = 16 # number of nodes per side y

rad_capture_site     =  sqrt(2)/10 # site radius
n_sides_capture_site = 4           # number of site sides
angle_capture_site   = pi/4        # site rotation angle
int_buffer           = FALSE       # insert intermediate buffer OPTIONAL

domain_p = Jsmm::creating_domain_from_locations(lc = lc, lc_label = lc_label,
                                          buffer_x = buffer_x,
                                          buffer_y = buffer_y,
                                          nnod_side_x = nnod_side_x,
                                          nnod_side_y = nnod_side_y,
                                          rad_capture_site = rad_capture_site,
                                          n_sides_capture_site = n_sides_capture_site,
                                          angle_capture_site = angle_capture_site,
                                          int_buffer = int_buffer)

# max area and min angle for the triangulation (See R-package RTriangle)
max_t_area = NULL
min_t_angle = 30

domain = Jsmm::jsmm_add_triangulation(domain = domain_p, max_t_area = max_t_area,
                                min_t_angle = min_t_angle)

Jsmm::plot_domain(domain = domain)

mapview(domain$polygon, zcol = "id") # visualize triangulation with zoom

#---------------------------------------------------------------------------
# CREATE observation_effort
#---------------------------------------------------------------------------

for(method in c("ICP","CCP")){
  study_duration  = 10 # duration of the entire study
  recapture_sites = 1:9
  trap_checking_interval = 1

  nloc = length(recapture_sites)
  observation_effort = list()
  observation_effort[["method"]] = method

  locations = list()  # Characterizing the elements that belong to each capture/release site

  for(i in 1:nloc){
    locations[[i]] = which(domain[["polygon"]][["id_location"]] == recapture_sites[i])
  }

  observation_effort[["locations"]] = locations
  observation_effort[["releases"]][["location"]] = sample(recapture_sites,
                                                          size = study_duration,
                                                          replace = T)

  observation_effort[["releases"]][["time"]] = 0:(study_duration - 1)

  recapture_times = c()
  for(i in seq(trap_checking_interval, study_duration, by = trap_checking_interval)){
    recapture_times = c(recapture_times, rep(i, nloc))
  }

  if(method=="CCP"){
    recapture_times = cbind(recapture_times-trap_checking_interval,recapture_times)
    colnames(recapture_times) = c("t_0", "t_1")
  }

  recapture_locations = rep(recapture_sites, study_duration/trap_checking_interval)
  observation_effort[["captures"]][["location"]] = recapture_locations
  observation_effort[["captures"]][["time"]] = recapture_times

  Jsmm::plot_observation_effort(observation_effort , xlab = "Time", ylab = "Location",
                          by_x = 1, cex.axisx = 1, cex.axisy = 1, lasy = 2, tcky = 0)

  #---------------------------------------------------------------------------
  # Set covariate and phylogenetic relationships data
  #---------------------------------------------------------------------------

  cov_data = list()
  altitude = domain[["triangulation"]][["node"]][, 2]
  cov_data[["n_data"]] = data.frame(altitude)
  t_data_times = seq(0,study_duration)
  air_temperature = -1 + 2*(t_data_times>study_duration/2)

  cov_data[["t_data_times"]] = t_data_times
  cov_data[["t_data"]] = data.frame(air_temperature)

  nevents = length(observation_effort[["captures"]][["location"]])
  capture_intensity = factor(sample(x = c("low","high"), size = nevents,replace = TRUE), levels = c("low", "high"))
  cov_data[["obs_data"]] = data.frame(capture_intensity)

  ns = 20 #number of species
  sp_names = paste0("sp_", 1:ns)

  tr_data = data.frame(size = rnorm(ns))

  order = rep(c(1, 1, 2, 2), ceiling(ns/4))[1:ns]
  genus = rep(c(1, 2, 3, 4), ceiling(ns/4))[1:ns]

  C = matrix(0, nrow = ns, ncol = ns)
  colnames(C) = sp_names
  rownames(C) = sp_names

  for(i in 1:ns){
    for(j in 1:ns){
      if(order[i] == order[j]) C[i, j] = C[i, j] + 0.45
      if(genus[i] == genus[j]) C[i, j] = C[i, j] + 0.45
      if(i == j) C[i, j] = 1
    }
  }

  corrplot_data = ggcorrplot(C)
  corrplot_data + theme(axis.text.x = element_text(size = 8),
                        axis.text.y = element_text(size = 8))


  model_formula = list()
  model_formula[["diffusion"]] = ~ air_temperature
  model_formula[["advection_1"]] = NULL
  model_formula[["advection_2"]] = NULL
  model_formula[["mortality"]] = ~ altitude
  model_formula[["habitat_preference"]] = NULL
  model_formula[["traits"]] = ~size
  model_formula[["observation"]] = ~ capture_intensity

  data = list()
  data[["captures"]] = NULL
  data[["releases"]] = NULL

  m = Jsmm::jsmm(domain = domain, observation_effort = observation_effort,
           releases = data[["releases"]], captures = data[["captures"]],
           cov_data = cov_data,
           ns = ns, sp_names = sp_names, tr_data = tr_data, C = C,
           model_formula = model_formula)

  save(m, file = paste0("models/unfitted_model_simulated_", method, "_no_data.RData"))
}
```
