## Supplementary material for "R-package Jsmm: Joint species movement modelling of mark-recapture data": S2 Vignettes demonstrating the application of the R-package Jsmm: 1b_simulate_parameters.R

```
library(Jsmm)

for(method in c("ICP", "CCP")){
  set.seed(3)
  file = paste0("models/unfitted_model_simulated_", method, "_no_data.RData")
  load(file)
  muZ = m[["prior"]][["muZ"]]
  nc = m[["nc"]]
  np = sum(nc)
  nt = m[["nt"]]
  idx = 1

  for(j in 1:nt){
    for(i in 1:length(nc)){
      for(ii in seq_len(nc[i])){
        if(names(nc)[i]=="diffusion" & ii==1 & j==1) muZ[idx] = -4 # baseline diffusion
        if(names(nc)[i]=="diffusion" & ii==2 & j==1) muZ[idx] = 1  # diffusion increases with air temperature
        if(names(nc)[i]=="diffusion" & ii==1 & j==2) muZ[idx] = 1  # diffusion intercept increases with body size
        if(names(nc)[i]=="mortality" & ii==1 & j==1) muZ[idx] = -2 # baseline mortality
        if(names(nc)[i]=="mortality" & ii==2 & j==1) muZ[idx] = 1  # mortality increases with altitude
        if(method=="ICP"){
          if(names(nc)[i]=="observation" & ii==1 & j==1) muZ[idx] = -2 # baseline observation rate
        }
        if(method=="CCP"){
          if(names(nc)[i]=="observation" & ii==1 & j==1) muZ[idx] = -2 # baseline observation rate
        }

        if(names(nc)[i]=="observation" & ii==2 & j==1) muZ[idx] = 1    # observation rate higher with high intensity
        idx = idx + 1
      }
    }
  }
  m[["prior"]][["muZ"]] = muZ
  m[["prior"]][["VZ"]] = 0*m[["prior"]][["VZ"]]
  m[["prior"]][["rho"]] = matrix(c(0.75, 1), nrow = 1)
  pars = Jsmm::simulate_from_prior(m)

  if(m$ns>1 & !is.null(m$tr_data)) plot(pars$Beta$diffusion[, 1] ~ m$tr_data[, 1])
  save(pars, file = paste0("parameters/true_parameters_", method, ".RData"))
  print(pars$Beta)
}

i = 2

plot(pars$Beta$diffusion[, i], col = c("blue", "cyan", "red", "pink"), pch = 16)
plot(pars$Beta$mortality[, i], col = c("blue", "cyan", "red", "pink"), pch = 16)
plot(pars$Beta$observation[, i], col = c("blue", "cyan", "red", "pink"), pch = 16)
```
