## Supplementary material for "R-package Jsmm: Joint species movement modelling of mark-recapture data": S2 Vignettes demonstrating the application of the R-package Jsmm: 1c_simulate_capture_data.R

```
library(Jsmm)
library(colorspace)

set.seed(3)

for(method in c("ICP", "CCP")){
  load(paste0("parameters/true_parameters_", method, ".RData"))
  load(paste0("models/unfitted_model_simulated_", method, "_no_data.RData"))

  ni = round(10000/m[["ns"]]) # number of individuals per species
  releases = matrix(NA, ncol = 2, nrow = ni*m[["ns"]])
  releases[, 1] = rep(1:m[["ns"]], ni)
  releases[, 2] = rep(1, ni*m[["ns"]])
  releases[, 2] = sample(seq_len(length(m$observation_effort$releases$location)), size = ni*m[["ns"]], replace = TRUE)
  colnames(releases) = c("sp", "release_event")
  head(releases)

  secondary_release = FALSE
  max_dt = NULL

  m = Jsmm::simulate_captures(m = m, pars = pars, releases = releases,
                        max_dt = max_dt, secondary_release = secondary_release)

  save(m, file = paste0("models/unfitted_model_simulated_",method,".RData"))
  ta = table(m$times[m$observation_effort$captures$time[m$captures[,2]]])
  print("Numbers of recaptures per species:")
  print(table(m[["releases"]][m[["captures"]][,1],1]))
}

model_summary(m)
```
