## Supplementary material for "R-package Jsmm: Joint species movement modelling of mark-recapture data": S2 Vignettes demonstrating the application of the R-package Jsmm: 1d_plot_likelihood_profiles.R

```
library(Jsmm)

set.seed(1)
max_dt = NULL

for(method in c("ICP", "CCP")){
  mname = paste0("simulated_",method)
  pdf(paste0("results/likelihood_profiles_", mname, ".pdf"))
  load(paste0("models/unfitted_model_", mname, ".RData"))
  m = merge_rc_data(m)
  load(paste0("parameters/true_parameters_", method, ".RData"))
  Beta.TRUE = pars$Beta
  vars = names(Beta.TRUE)
  tmp = compute_likelihood(m, Beta = Beta.TRUE, max_dt = max_dt)
  li.TRUE = tmp$log.likelihood.sp
  nb.TRUE = tmp$numerical_problems
  for(va in vars){
    print(va)
    for(i in 1:ncol(Beta.TRUE[[va]])){
      adds = (-5:5)/5
      na = length(adds)
      re = matrix(nrow = na, ncol = m$ns)
      renb = matrix(nrow = na, ncol = m$ns)
      for(ai in 1:na){
        print(adds[ai])
        Beta =  Beta.TRUE
        Beta[[va]][, i] = Beta[[va]][, i] + adds[ai]
        tmp = compute_likelihood(m,Beta = Beta, max_dt = max_dt)
        li = tmp$log.likelihood.sp
        re[ai,] = li - li.TRUE
        renb[ai,] = tmp$numerical_problems
      }
      plot(adds,re[,1],type="l",col="red", main = paste(va, i), ylim = c(min(re), max(re)))
      abline(h = 0)
      abline(v = 0)
      if(m$ns>1) for(j in 2:m$ns) lines(adds, re[, j], col = "blue")
    #  plot(adds,renb[,1],type="l",col="red",main=paste(va,i),ylab="numerical problems")
    #  abline(v=0)
    }
  }
  dev.off()
}
```
