## Supplementary material for "R-package Jsmm: Joint species movement modelling of mark-recapture data": S2 Vignettes demonstrating the application of the R-package Jsmm: MCMC_convergence_simulated_CCP.pdf

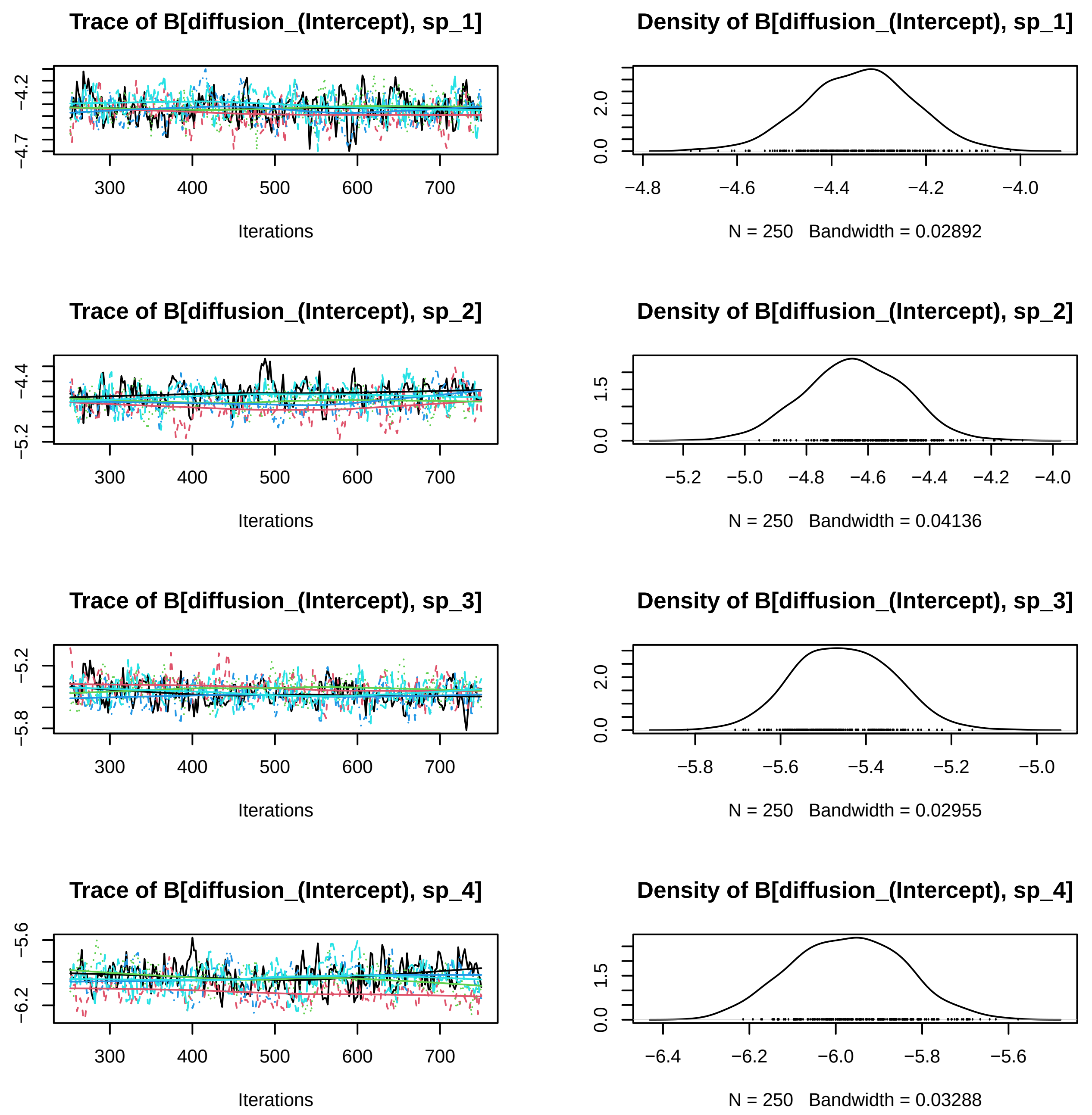
Trace of B[diffusion_(Intercept), sp_1]
-4.2
-4.7
300
400
500
600
700
Iterations
Density of B[diffusion_(Intercept), sp_1]
0.0
2.0
-4.8
-4.6
-4.4
-4.2
-4.0
N = 250 Bandwidth = 0.02892
Trace of B[diffusion_(Intercept), sp_2]
-4.4
-5.2
300
400
500
600
700
Iterations
Density of B[diffusion_(Intercept), sp_2]
0.0
1.5
-5.2
-5.0
-4.8
-4.6
-4.4
-4.2
-4.0
N = 250 Bandwidth = 0.04136
Trace of B[diffusion_(Intercept), sp_3]
-5.2
-5.8
300
400
500
600
700
Iterations
Density of B[diffusion_(Intercept), sp_3]
0.0
2.0
-5.8
-5.6
-5.4
-5.2
-5.0
N = 250 Bandwidth = 0.02955
Trace of B[diffusion_(Intercept), sp_4]
-5.6
-6.2
300
400
500
600
700
Iterations
Density of B[diffusion_(Intercept), sp_4]
0.0
1.5
-6.4
-6.2
-6.0
-5.8
-5.6
N = 250 Bandwidth = 0.03288

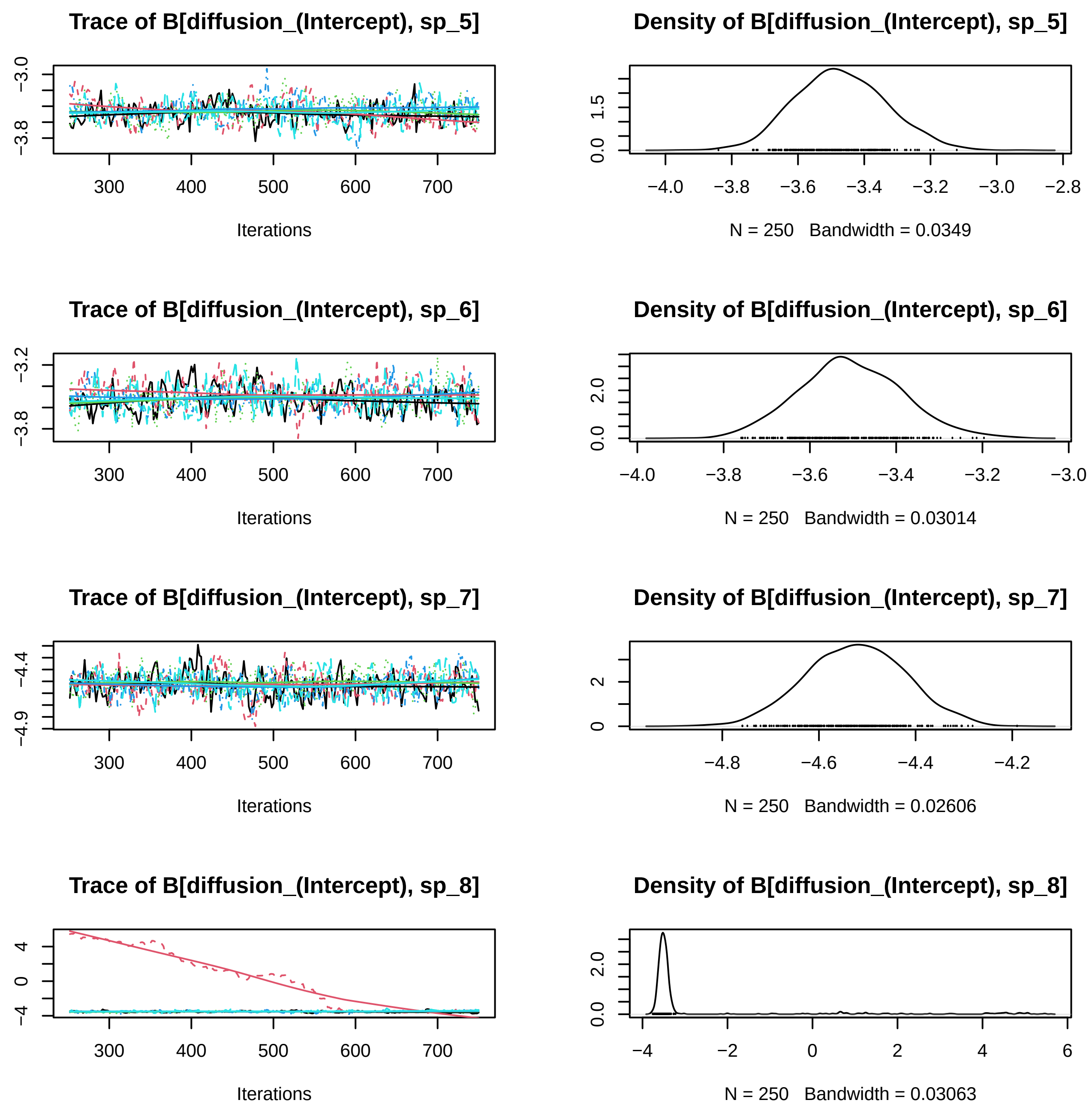
Trace of B[diffusion_(Intercept), sp_5]
Density of B[diffusion_(Intercept), sp_5]
Iterations
N = 250 Bandwidth = 0.0349
Trace of B[diffusion_(Intercept), sp_6]
Density of B[diffusion_(Intercept), sp_6]
Iterations
N = 250 Bandwidth = 0.03014
Trace of B[diffusion_(Intercept), sp_7]
Density of B[diffusion_(Intercept), sp_7]
Iterations
N = 250 Bandwidth = 0.02606
Trace of B[diffusion_(Intercept), sp_8]
Density of B[diffusion_(Intercept), sp_8]
Iterations
N = 250 Bandwidth = 0.03063

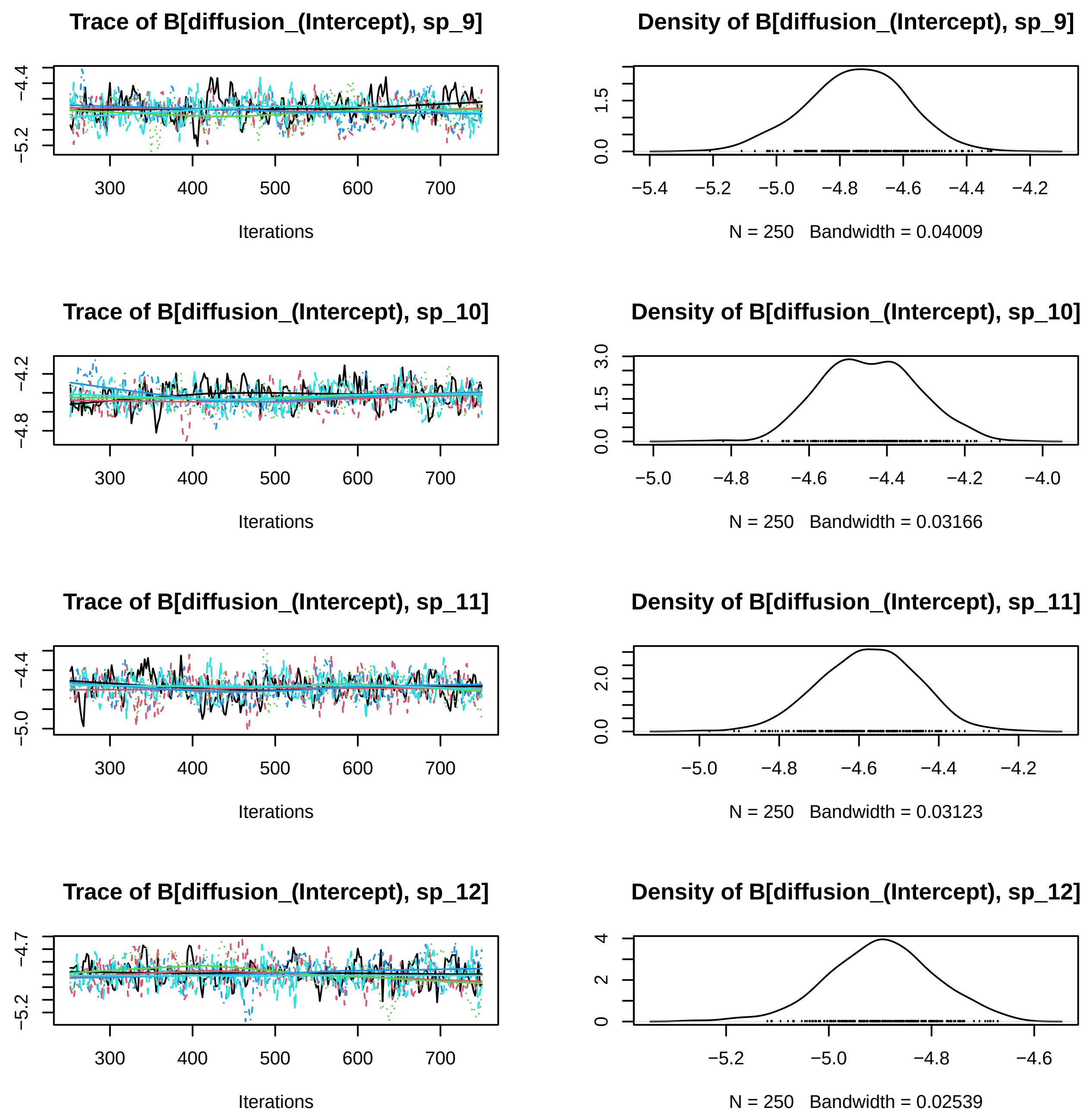
Trace of B[diffusion_(Intercept), sp_9]
Iterations
Density of B[diffusion_(Intercept), sp_9]
N = 250 Bandwidth = 0.04009
Trace of B[diffusion_(Intercept), sp_10]
Iterations
Density of B[diffusion_(Intercept), sp_10]
N = 250 Bandwidth = 0.03166
Trace of B[diffusion_(Intercept), sp_11]
Iterations
Density of B[diffusion_(Intercept), sp_11]
N = 250 Bandwidth = 0.03123
Trace of B[diffusion_(Intercept), sp_12]
Iterations
Density of B[diffusion_(Intercept), sp_12]
N = 250 Bandwidth = 0.02539

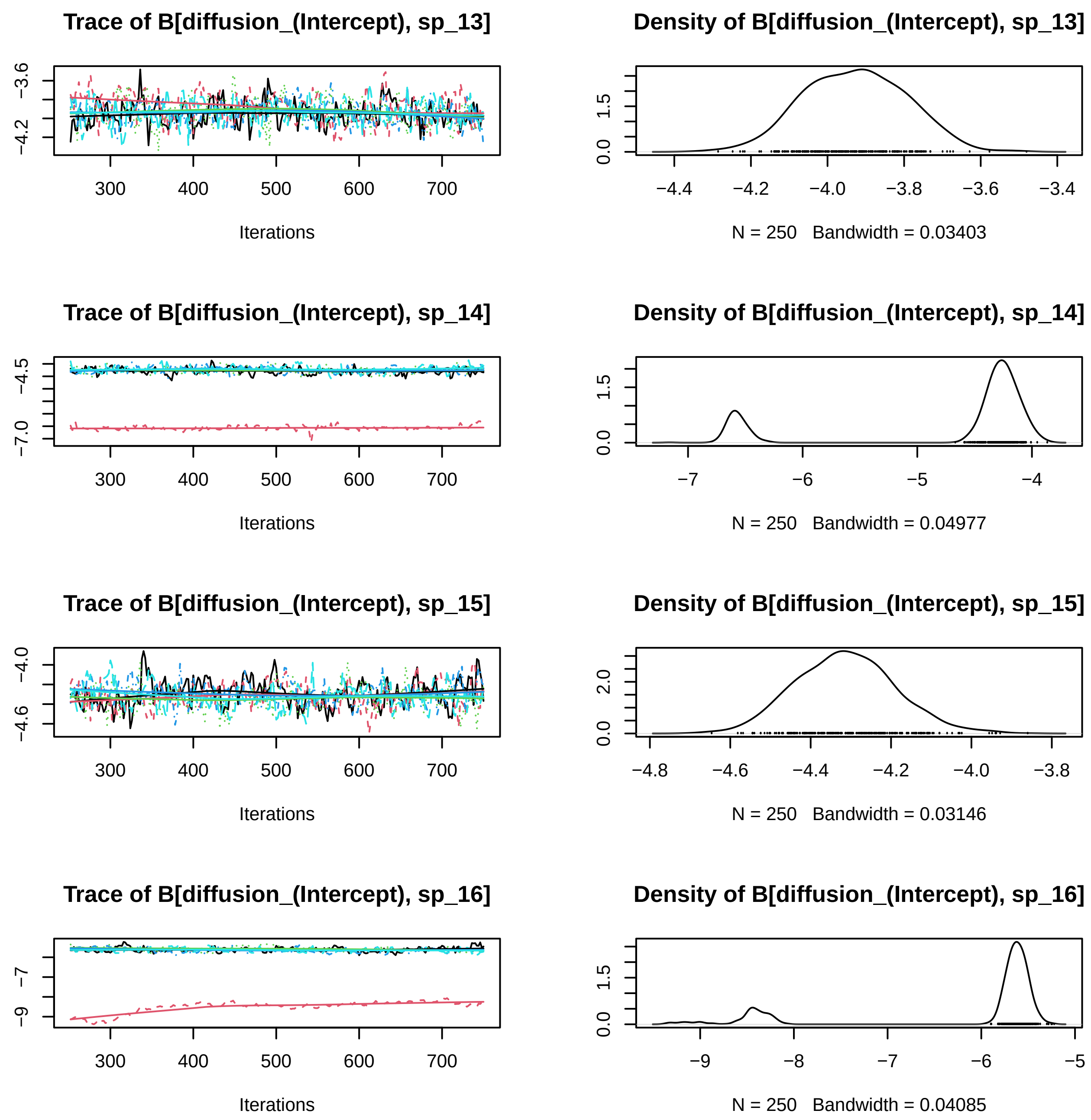

Trace of B[diffusion_(Intercept), sp_13]
Iterations
Density of B[diffusion_(Intercept), sp_13]
N = 250 Bandwidth = 0.03403
Trace of B[diffusion_(Intercept), sp_14]
Iterations
Density of B[diffusion_(Intercept), sp_14]
N = 250 Bandwidth = 0.04977
Trace of B[diffusion_(Intercept), sp_15]
Iterations
Density of B[diffusion_(Intercept), sp_15]
N = 250 Bandwidth = 0.03146
Trace of B[diffusion_(Intercept), sp_16]
Iterations
Density of B[diffusion_(Intercept), sp_16]
N = 250 Bandwidth = 0.04085

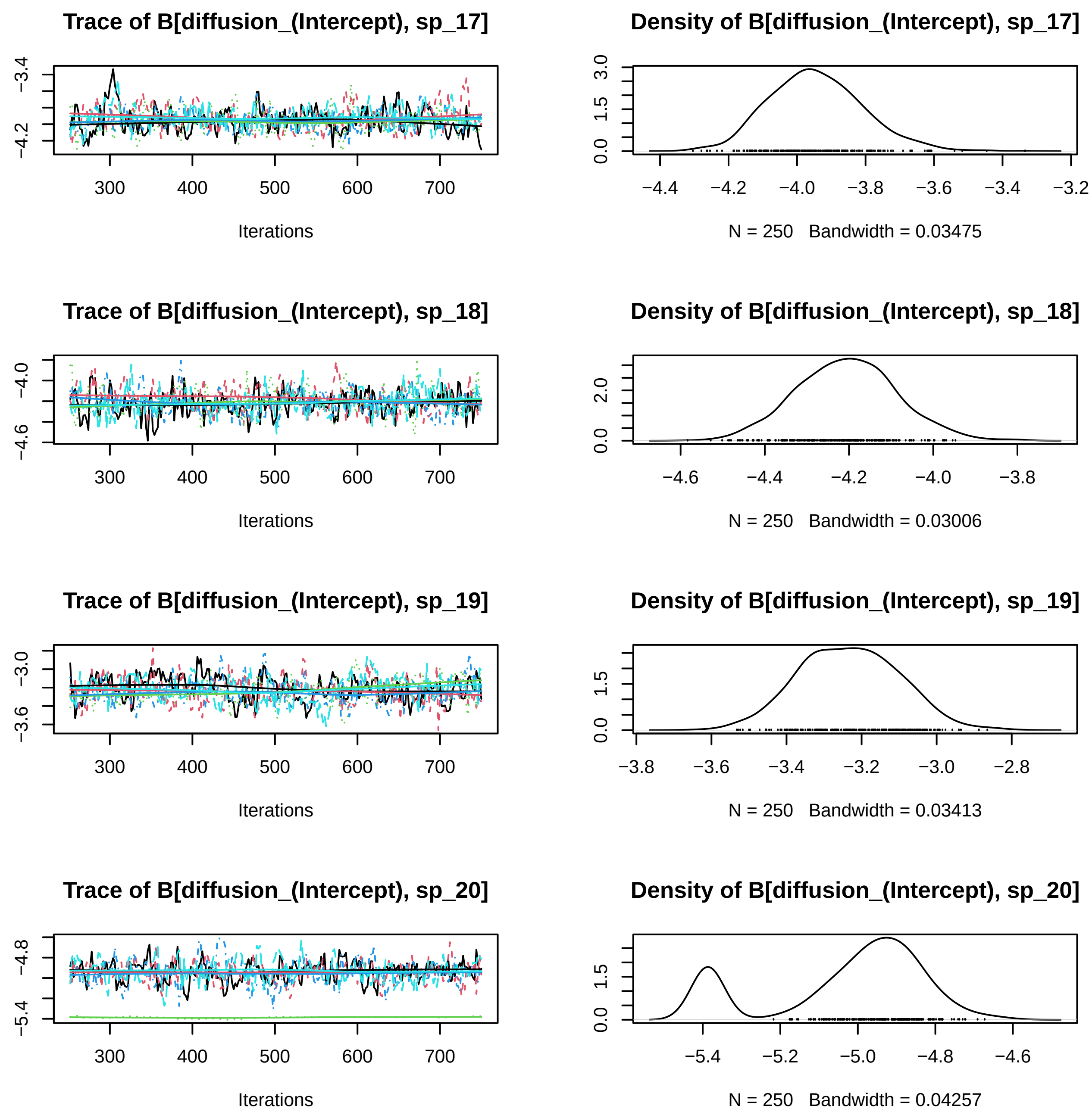
Trace of B[diffusion_(Intercept), sp_17]
−3.4
−4.2
300
400
500
600
700
Iterations
Density of B[diffusion_(Intercept), sp_17]
0.0
1.5
3.0
−4.4
−4.2
−4.0
−3.8
−3.6
−3.4
−3.2
N = 250 Bandwidth = 0.03475
Trace of B[diffusion_(Intercept), sp_18]
−4.0
−4.6
300
400
500
600
700
Iterations
Density of B[diffusion_(Intercept), sp_18]
0.0
2.0
−4.6
−4.4
−4.2
−4.0
−3.8
N = 250 Bandwidth = 0.03006
Trace of B[diffusion_(Intercept), sp_19]
−3.0
−3.6
300
400
500
600
700
Iterations
Density of B[diffusion_(Intercept), sp_19]
0.0
1.5
−3.8
−3.6
−3.4
−3.2
−3.0
−2.8
N = 250 Bandwidth = 0.03413
Trace of B[diffusion_(Intercept), sp_20]
−4.8
−5.4
300
400
500
600
700
Iterations
Density of B[diffusion_(Intercept), sp_20]
0.0
1.5
−5.4
−5.2
−5.0
−4.8
−4.6
N = 250 Bandwidth = 0.04257

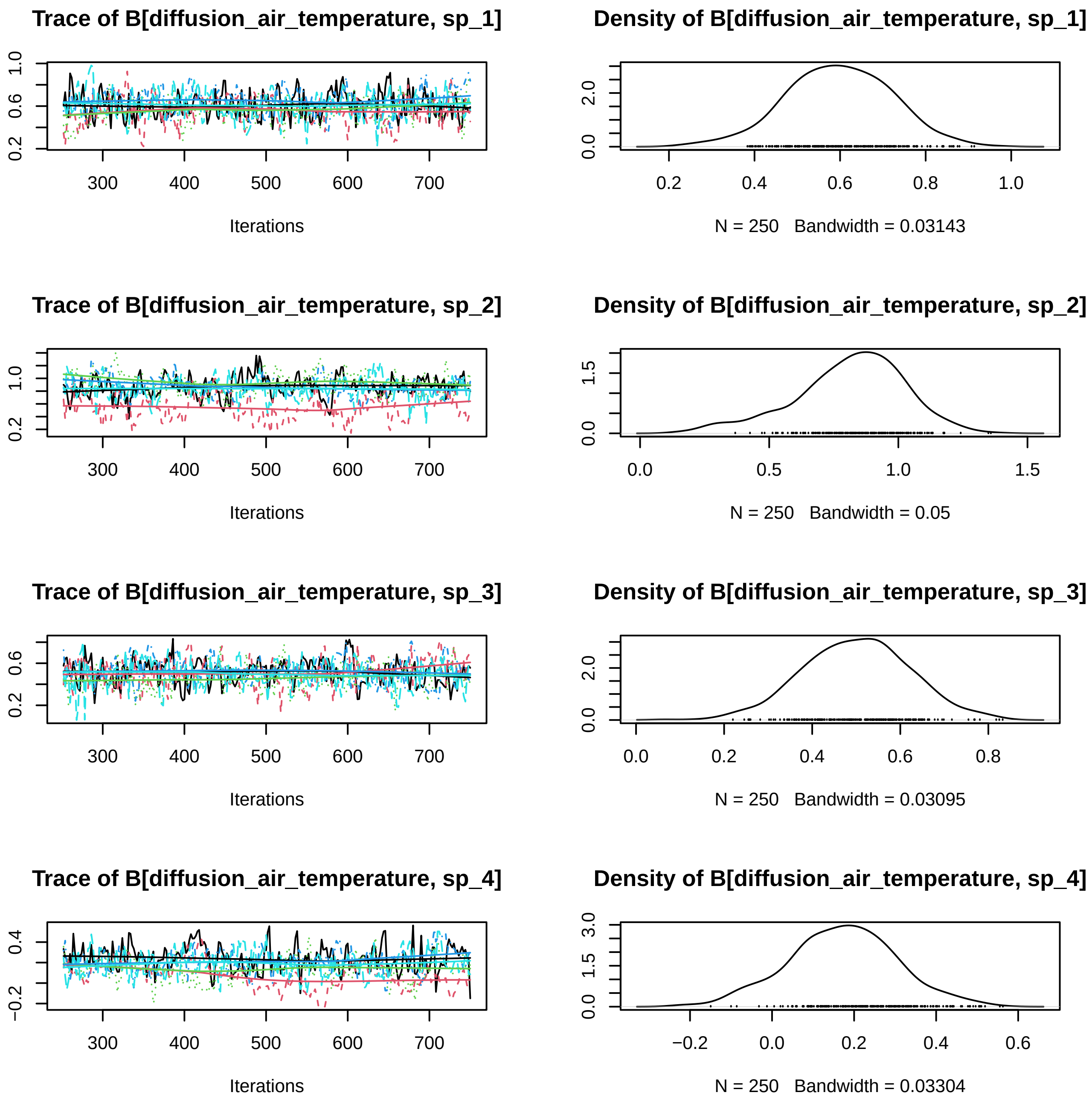
Trace of B[diffusion_air_temperature, sp_1]
Iterations
Density of B[diffusion_air_temperature, sp_1]
N = 250 Bandwidth = 0.03143
Trace of B[diffusion_air_temperature, sp_2]
Iterations
Density of B[diffusion_air_temperature, sp_2]
N = 250 Bandwidth = 0.05
Trace of B[diffusion_air_temperature, sp_3]
Iterations
Density of B[diffusion_air_temperature, sp_3]
N = 250 Bandwidth = 0.03095
Trace of B[diffusion_air_temperature, sp_4]
Iterations
Density of B[diffusion_air_temperature, sp_4]
N = 250 Bandwidth = 0.03304

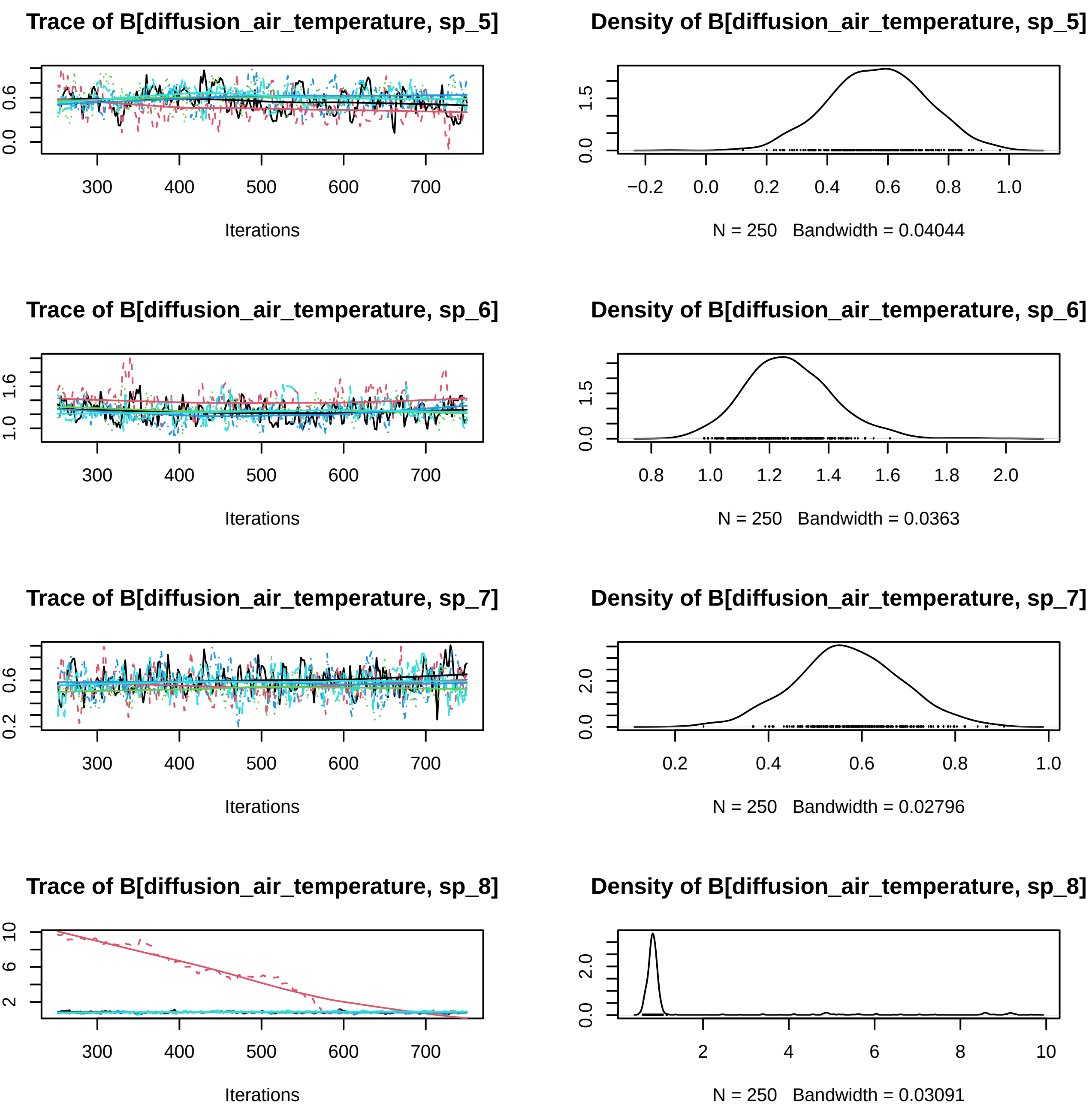
Trace of B[diffusion_air_temperature, sp_5]
Iterations
Density of B[diffusion_air_temperature, sp_5]
N = 250 Bandwidth = 0.04044
Trace of B[diffusion_air_temperature, sp_6]
Iterations
Density of B[diffusion_air_temperature, sp_6]
N = 250 Bandwidth = 0.0363
Trace of B[diffusion_air_temperature, sp_7]
Iterations
Density of B[diffusion_air_temperature, sp_7]
N = 250 Bandwidth = 0.02796
Trace of B[diffusion_air_temperature, sp_8]
Iterations
Density of B[diffusion_air_temperature, sp_8]
N = 250 Bandwidth = 0.03091

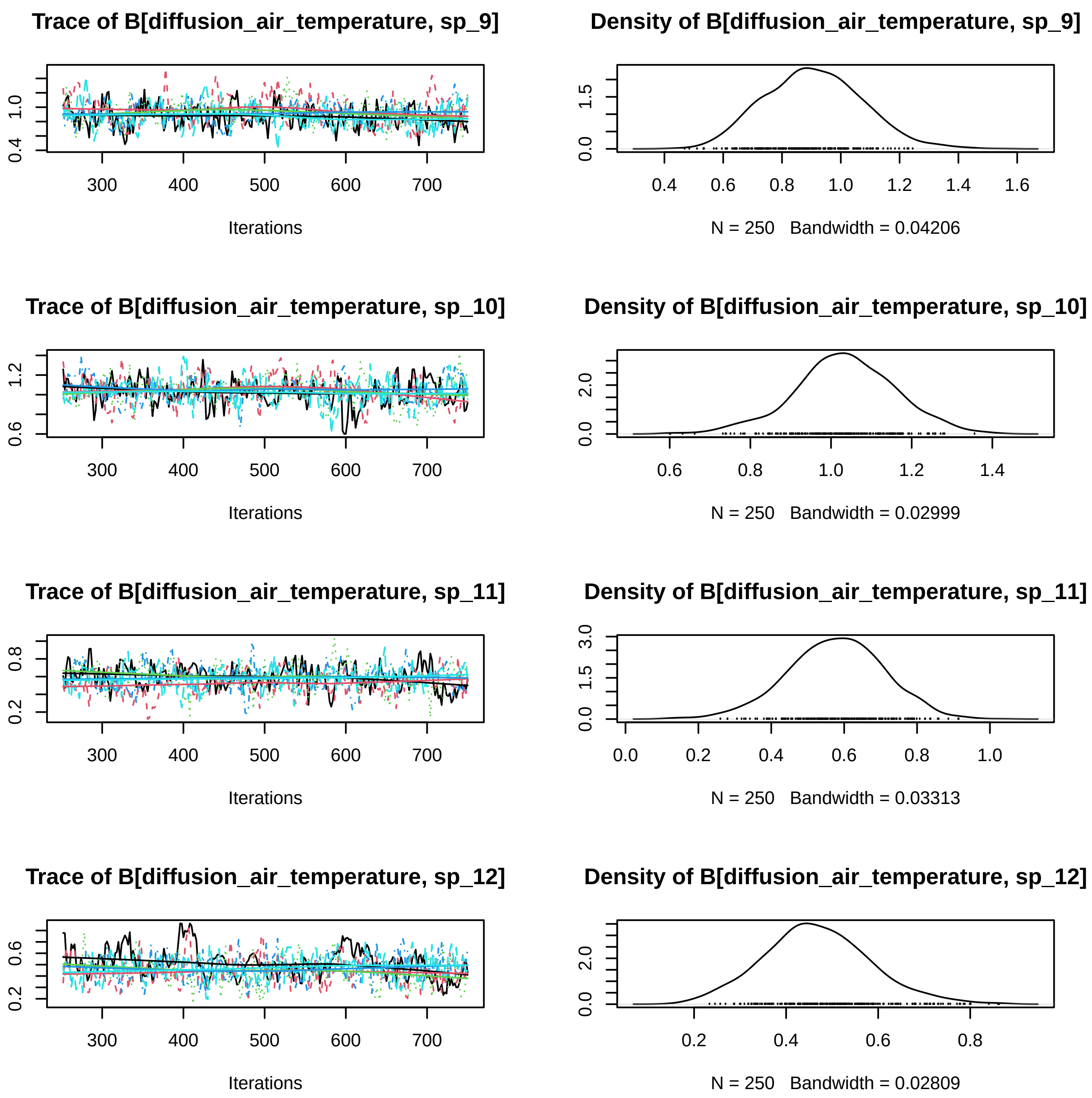
Trace of B[diffusion_air_temperature, sp_9]
Density of B[diffusion_air_temperature, sp_9]
0.4
1.0
300
400
500
600
700
Iterations
0.0
1.5
0.4
0.6
0.8
1.0
1.2
1.4
1.6
N = 250   Bandwidth = 0.04206
Trace of B[diffusion_air_temperature, sp_10]
Density of B[diffusion_air_temperature, sp_10]
0.6
1.2
300
400
500
600
700
Iterations
0.0
2.0
0.6
0.8
1.0
1.2
1.4
N = 250   Bandwidth = 0.02999
Trace of B[diffusion_air_temperature, sp_11]
Density of B[diffusion_air_temperature, sp_11]
0.2
0.8
300
400
500
600
700
Iterations
0.0
1.5
3.0
0.0
0.2
0.4
0.6
0.8
1.0
N = 250   Bandwidth = 0.03313
Trace of B[diffusion_air_temperature, sp_12]
Density of B[diffusion_air_temperature, sp_12]
0.2
0.6
300
400
500
600
700
Iterations
0.0
2.0
0.2
0.4
0.6
0.8
N = 250   Bandwidth = 0.02809

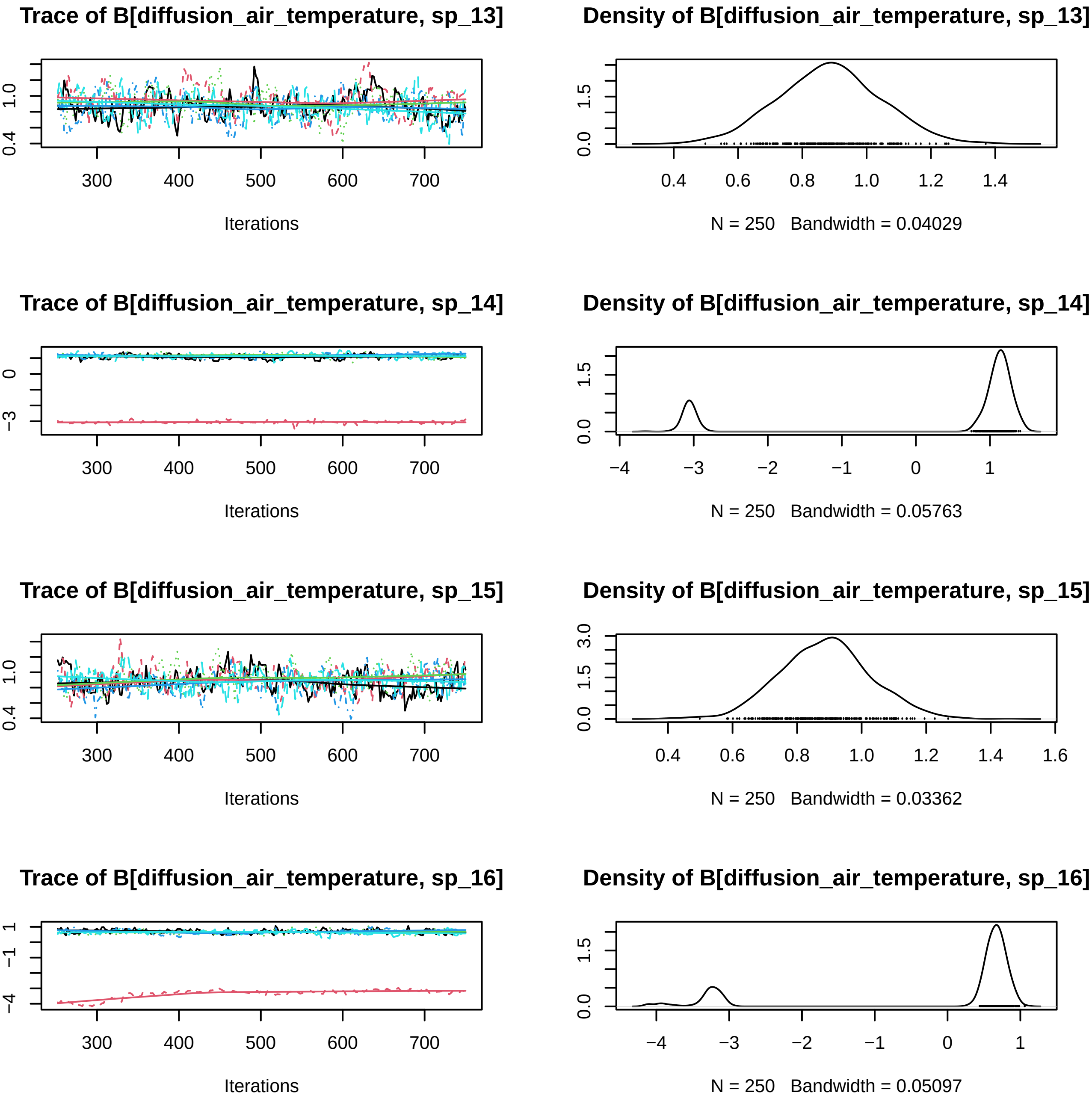
Trace of B[diffusion_air_temperature, sp_13]
Iterations
Density of B[diffusion_air_temperature, sp_13]
N = 250 Bandwidth = 0.04029
Trace of B[diffusion_air_temperature, sp_14]
Iterations
Density of B[diffusion_air_temperature, sp_14]
N = 250 Bandwidth = 0.05763
Trace of B[diffusion_air_temperature, sp_15]
Iterations
Density of B[diffusion_air_temperature, sp_15]
N = 250 Bandwidth = 0.03362
Trace of B[diffusion_air_temperature, sp_16]
Iterations
Density of B[diffusion_air_temperature, sp_16]
N = 250 Bandwidth = 0.05097

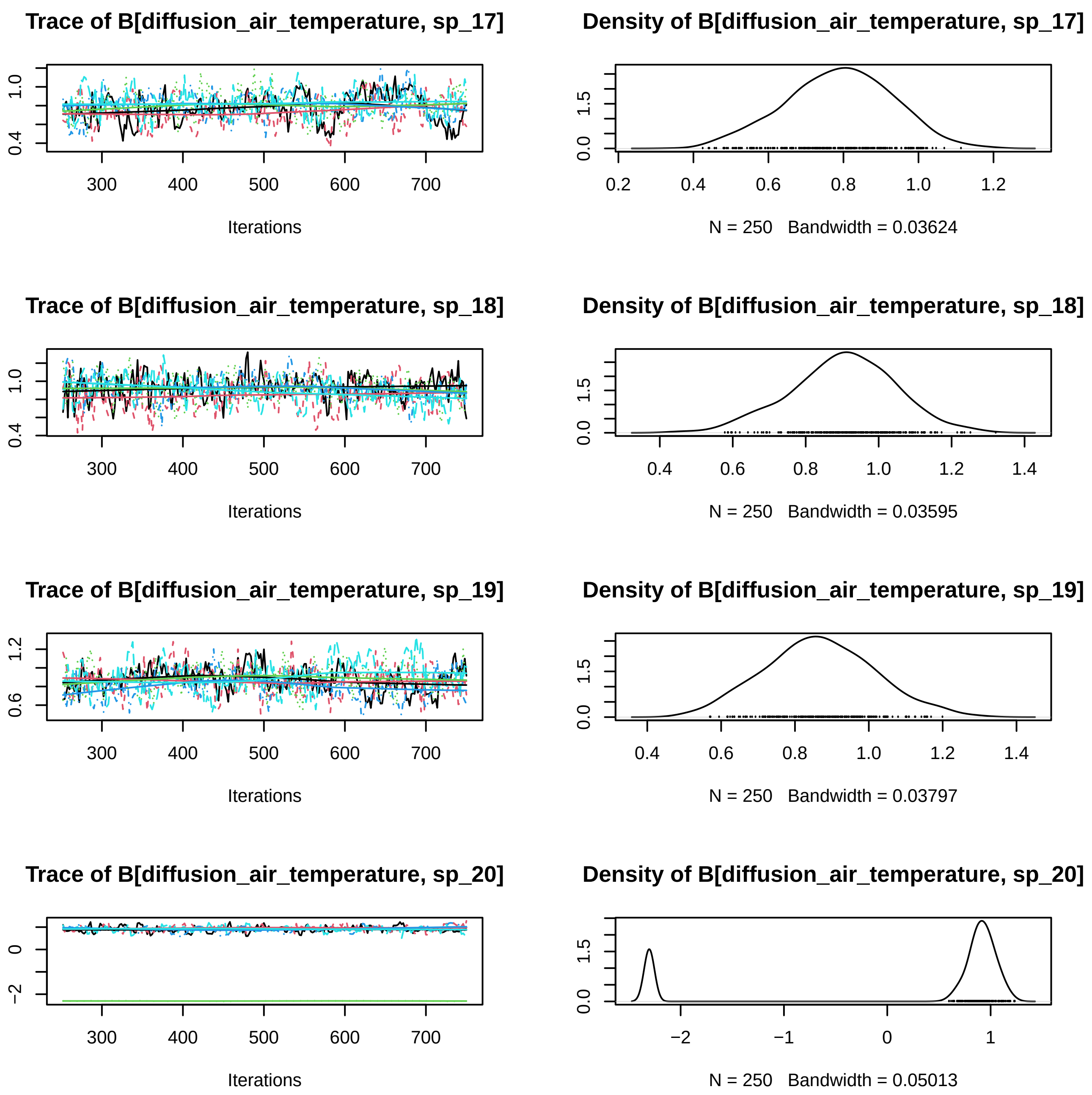
Trace of B[diffusion_air_temperature, sp_17]
Iterations
Density of B[diffusion_air_temperature, sp_17]
N = 250 Bandwidth = 0.03624
Trace of B[diffusion_air_temperature, sp_18]
Iterations
Density of B[diffusion_air_temperature, sp_18]
N = 250 Bandwidth = 0.03595
Trace of B[diffusion_air_temperature, sp_19]
Iterations
Density of B[diffusion_air_temperature, sp_19]
N = 250 Bandwidth = 0.03797
Trace of B[diffusion_air_temperature, sp_20]
Iterations
Density of B[diffusion_air_temperature, sp_20]
N = 250 Bandwidth = 0.05013

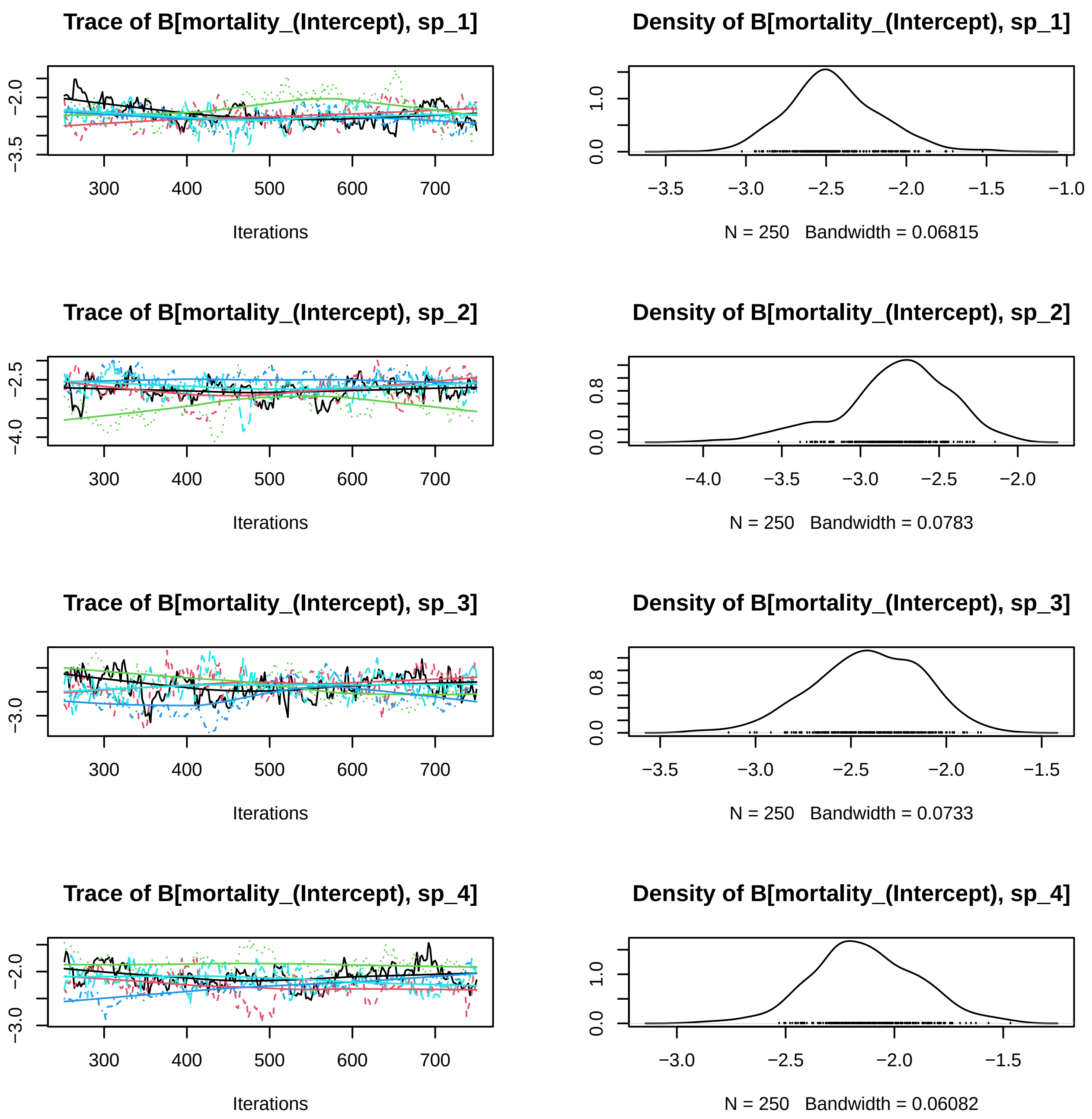
Trace of B[mortality_(Intercept), sp_1]
−3.5
−2.0
300
400
500
600
700
Iterations
Density of B[mortality_(Intercept), sp_1]
0.0
1.0
−3.5
−3.0
−2.5
−2.0
−1.5
−1.0
N = 250 Bandwidth = 0.06815
Trace of B[mortality_(Intercept), sp_2]
−4.0
−2.5
300
400
500
600
700
Iterations
Density of B[mortality_(Intercept), sp_2]
0.0
0.8
−4.0
−3.5
−3.0
−2.5
−2.0
N = 250 Bandwidth = 0.0783
Trace of B[mortality_(Intercept), sp_3]
−3.0
300
400
500
600
700
Iterations
Density of B[mortality_(Intercept), sp_3]
0.0
0.8
−3.5
−3.0
−2.5
−2.0
−1.5
N = 250 Bandwidth = 0.0733
Trace of B[mortality_(Intercept), sp_4]
−3.0
−2.0
300
400
500
600
700
Iterations
Density of B[mortality_(Intercept), sp_4]
0.0
1.0
−3.0
−2.5
−2.0
−1.5
N = 250 Bandwidth = 0.06082

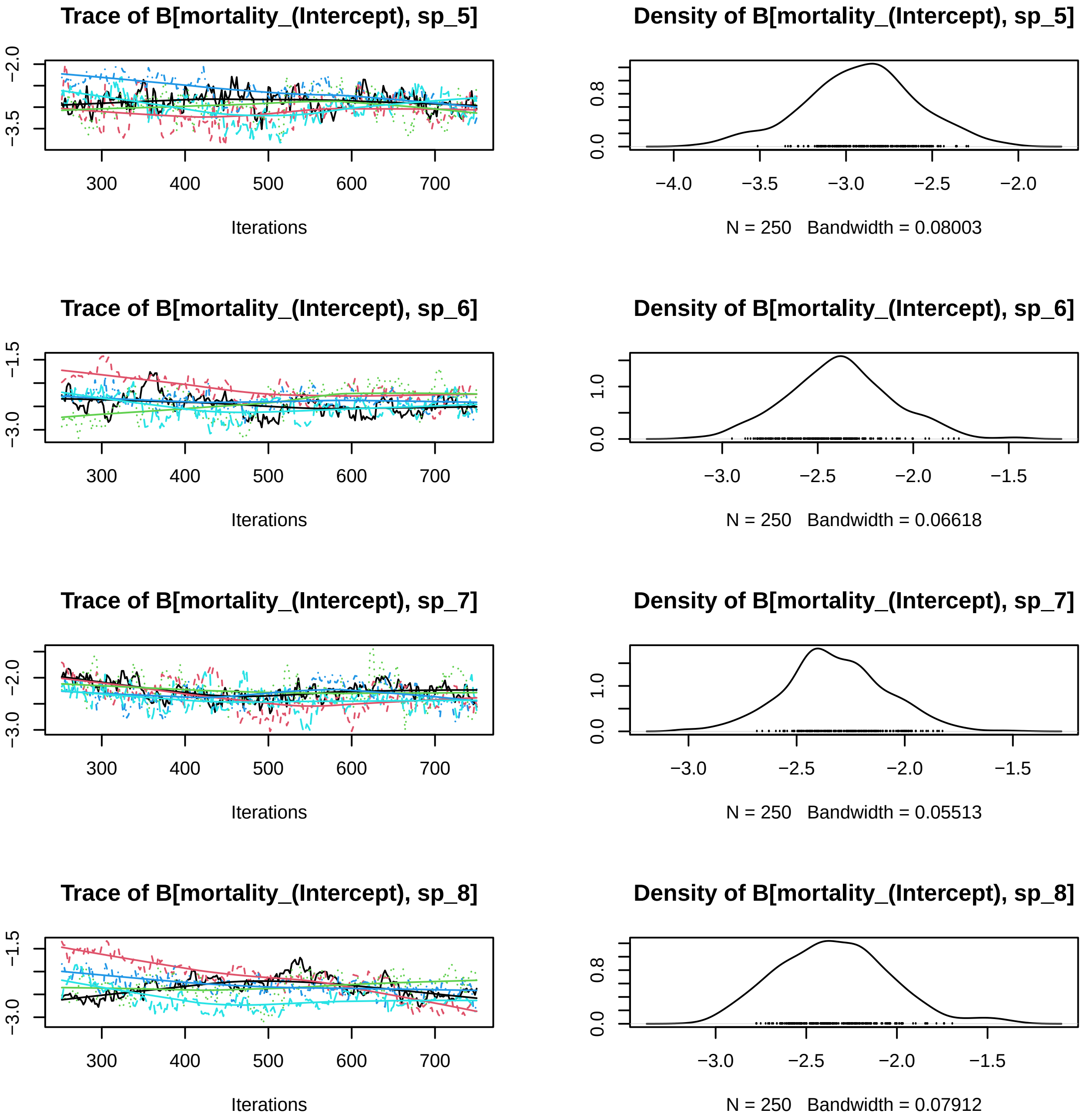
Trace of B[mortality_(Intercept), sp_5]
Density of B[mortality_(Intercept), sp_5]
Iterations
N = 250 Bandwidth = 0.08003
Trace of B[mortality_(Intercept), sp_6]
Density of B[mortality_(Intercept), sp_6]
Iterations
N = 250 Bandwidth = 0.06618
Trace of B[mortality_(Intercept), sp_7]
Density of B[mortality_(Intercept), sp_7]
Iterations
N = 250 Bandwidth = 0.05513
Trace of B[mortality_(Intercept), sp_8]
Density of B[mortality_(Intercept), sp_8]
Iterations
N = 250 Bandwidth = 0.07912

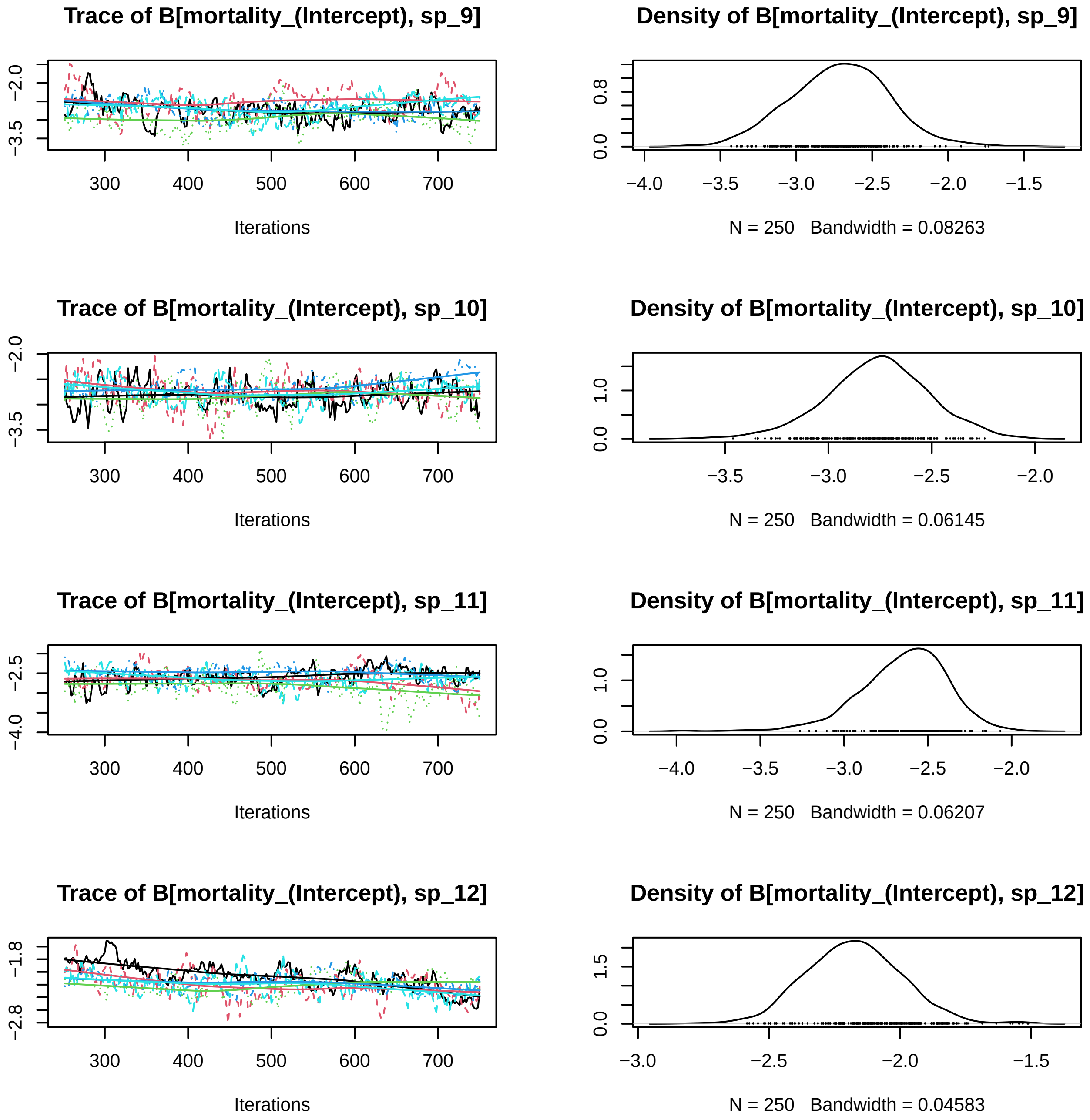
Trace of B[mortality_(Intercept), sp_9]
Density of B[mortality_(Intercept), sp_9]
Iterations
N = 250 Bandwidth = 0.08263
Trace of B[mortality_(Intercept), sp_10]
Density of B[mortality_(Intercept), sp_10]
Iterations
N = 250 Bandwidth = 0.06145
Trace of B[mortality_(Intercept), sp_11]
Density of B[mortality_(Intercept), sp_11]
Iterations
N = 250 Bandwidth = 0.06207
Trace of B[mortality_(Intercept), sp_12]
Density of B[mortality_(Intercept), sp_12]
Iterations
N = 250 Bandwidth = 0.04583

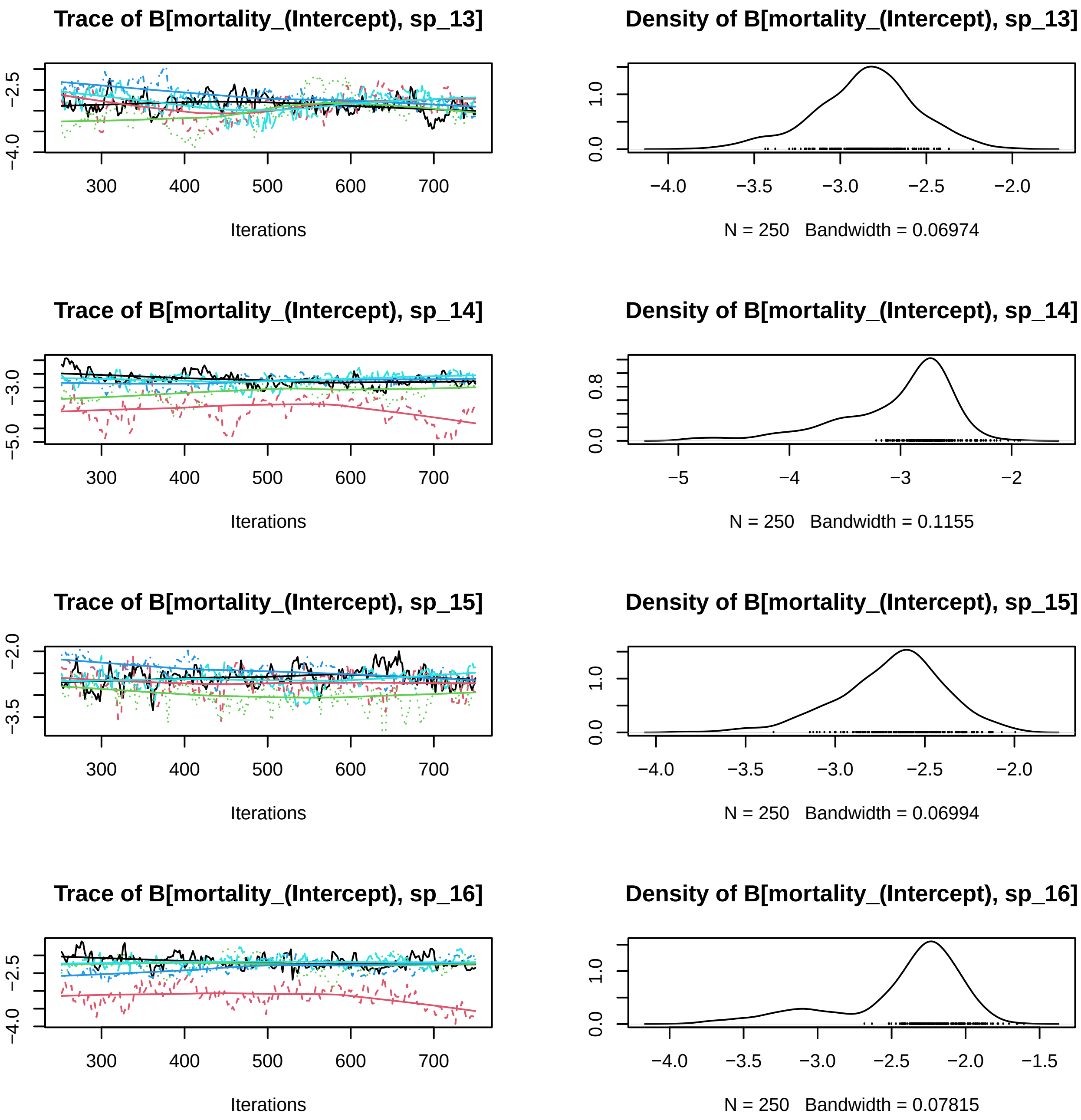
Trace of B[mortality_(Intercept), sp_13]
−4.0
−2.5
300
400
500
600
700
Iterations
Density of B[mortality_(Intercept), sp_13]
0.0
1.0
−4.0
−3.5
−3.0
−2.5
−2.0
N = 250 Bandwidth = 0.06974
Trace of B[mortality_(Intercept), sp_14]
−5.0
−3.0
300
400
500
600
700
Iterations
Density of B[mortality_(Intercept), sp_14]
0.0
0.8
−5
−4
−3
−2
N = 250 Bandwidth = 0.1155
Trace of B[mortality_(Intercept), sp_15]
−3.5
−2.0
300
400
500
600
700
Iterations
Density of B[mortality_(Intercept), sp_15]
0.0
1.0
−4.0
−3.5
−3.0
−2.5
−2.0
N = 250 Bandwidth = 0.06994
Trace of B[mortality_(Intercept), sp_16]
−4.0
−2.5
300
400
500
600
700
Iterations
Density of B[mortality_(Intercept), sp_16]
0.0
1.0
−4.0
−3.5
−3.0
−2.5
−2.0
−1.5
N = 250 Bandwidth = 0.07815

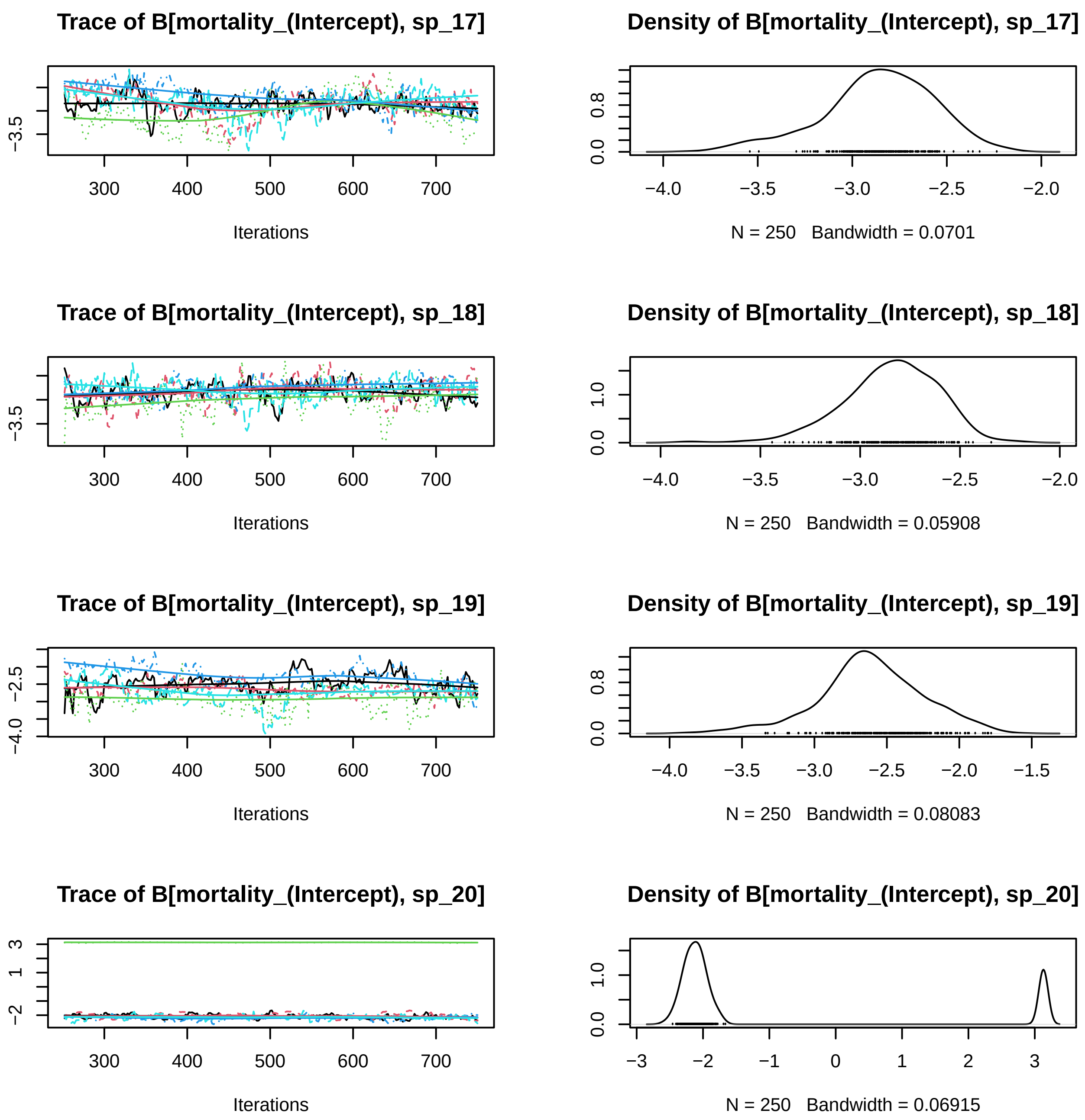
Trace of B[mortality_(Intercept), sp_17]
−3.5
300
400
500
600
700
Iterations
Density of B[mortality_(Intercept), sp_17]
0.0
0.8
−4.0
−3.5
−3.0
−2.5
−2.0
N = 250 Bandwidth = 0.0701
Trace of B[mortality_(Intercept), sp_18]
−3.5
300
400
500
600
700
Iterations
Density of B[mortality_(Intercept), sp_18]
0.0
1.0
−4.0
−3.5
−3.0
−2.5
−2.0
N = 250 Bandwidth = 0.05908
Trace of B[mortality_(Intercept), sp_19]
−4.0
−2.5
300
400
500
600
700
Iterations
Density of B[mortality_(Intercept), sp_19]
0.0
0.8
−4.0
−3.5
−3.0
−2.5
−2.0
−1.5
N = 250 Bandwidth = 0.08083
Trace of B[mortality_(Intercept), sp_20]
−2
1
3
300
400
500
600
700
Iterations
Density of B[mortality_(Intercept), sp_20]
0.0
1.0
−3
−2
−1
0
1
2
3
N = 250 Bandwidth = 0.06915

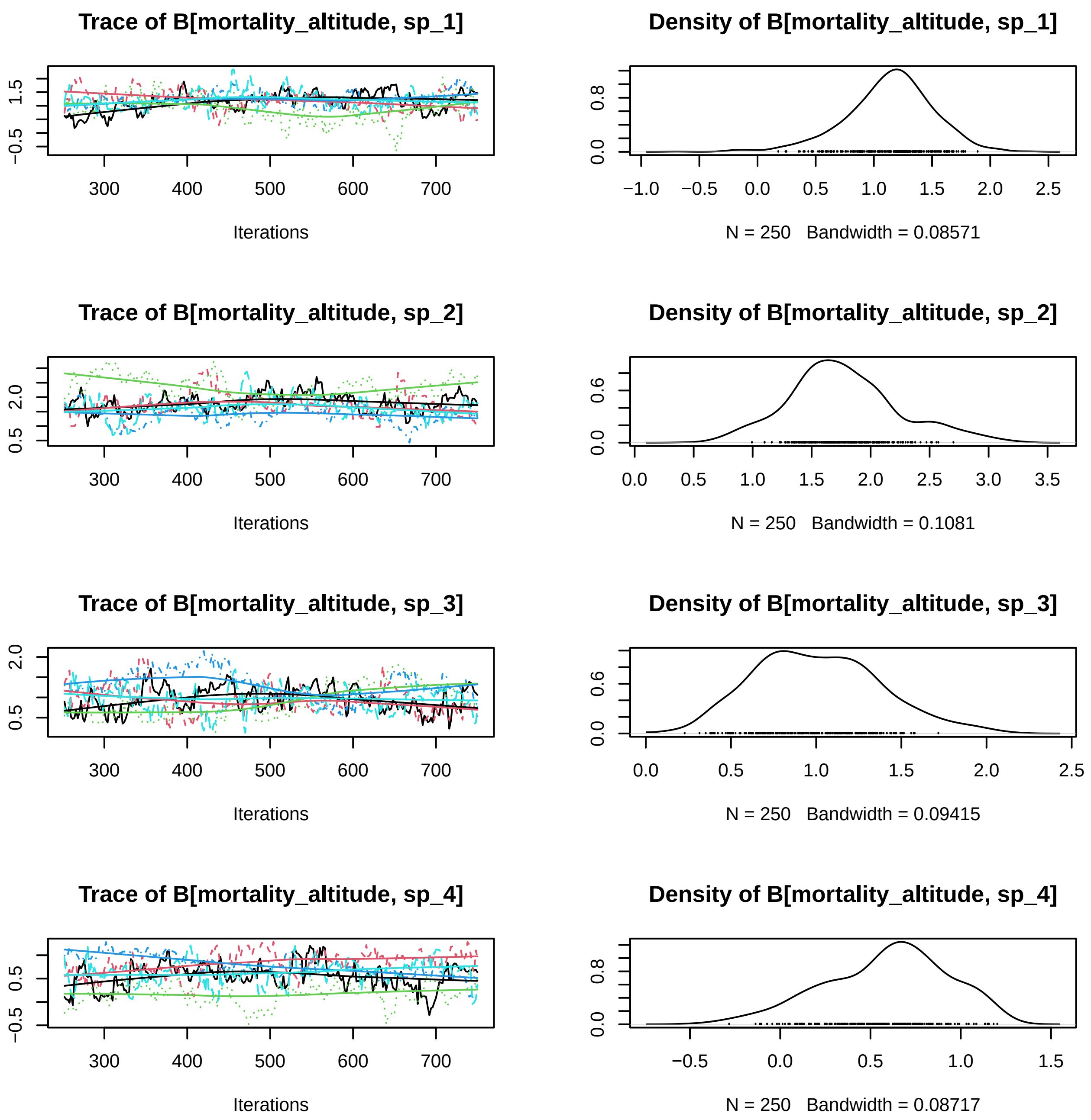
Trace of B[mortality_altitude, sp_1]
Iterations
Density of B[mortality_altitude, sp_1]
N = 250 Bandwidth = 0.08571
Trace of B[mortality_altitude, sp_2]
Iterations
Density of B[mortality_altitude, sp_2]
N = 250 Bandwidth = 0.1081
Trace of B[mortality_altitude, sp_3]
Iterations
Density of B[mortality_altitude, sp_3]
N = 250 Bandwidth = 0.09415
Trace of B[mortality_altitude, sp_4]
Iterations
Density of B[mortality_altitude, sp_4]
N = 250 Bandwidth = 0.08717

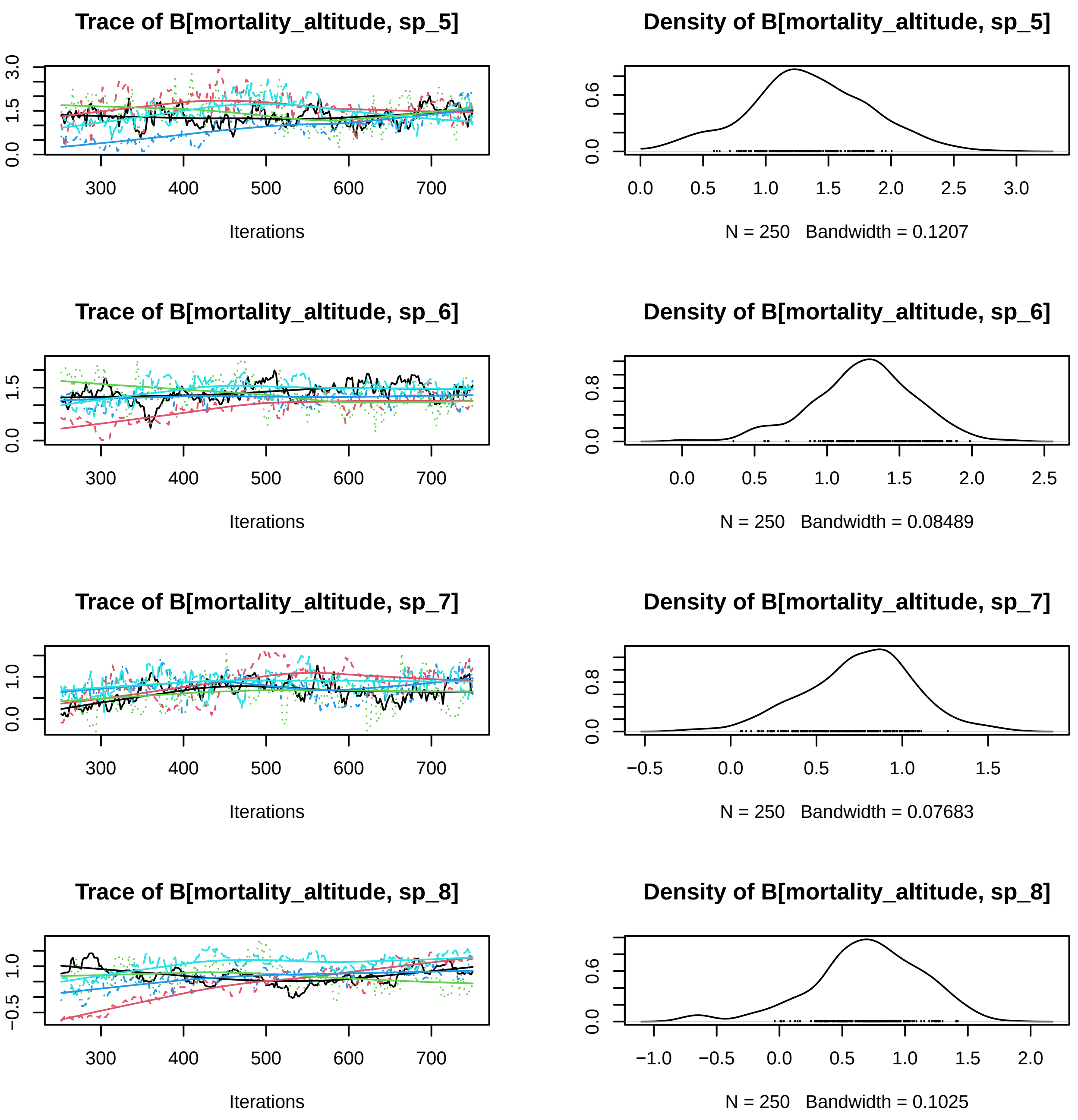
Trace of B[mortality_altitude, sp_5]
Iterations
Density of B[mortality_altitude, sp_5]
N = 250 Bandwidth = 0.1207
Trace of B[mortality_altitude, sp_6]
Iterations
Density of B[mortality_altitude, sp_6]
N = 250 Bandwidth = 0.08489
Trace of B[mortality_altitude, sp_7]
Iterations
Density of B[mortality_altitude, sp_7]
N = 250 Bandwidth = 0.07683
Trace of B[mortality_altitude, sp_8]
Iterations
Density of B[mortality_altitude, sp_8]
N = 250 Bandwidth = 0.1025

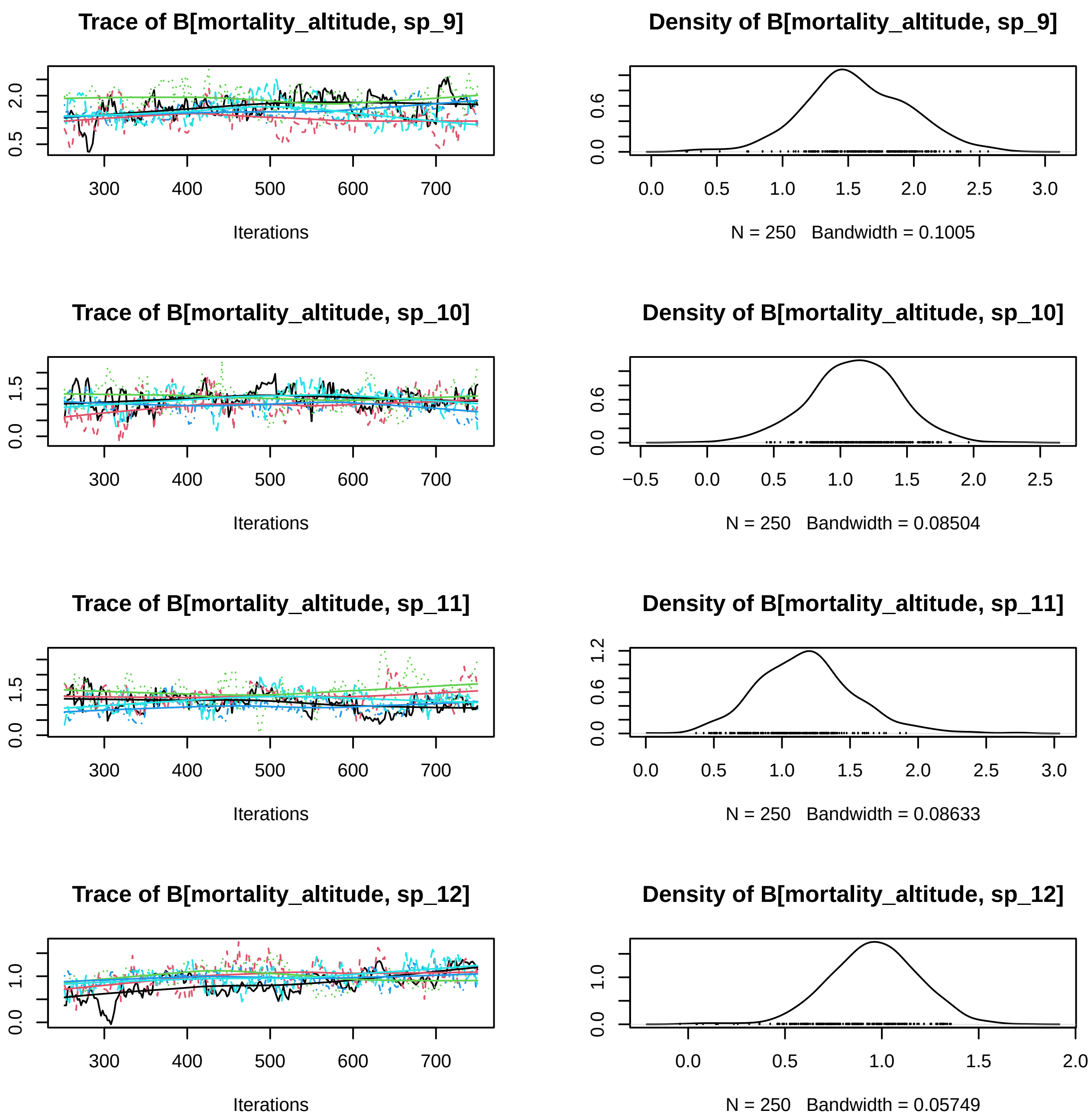

Trace of B[mortality_altitude, sp_9]
Iterations
Density of B[mortality_altitude, sp_9]
N = 250 Bandwidth = 0.1005
Trace of B[mortality_altitude, sp_10]
Iterations
Density of B[mortality_altitude, sp_10]
N = 250 Bandwidth = 0.08504
Trace of B[mortality_altitude, sp_11]
Iterations
Density of B[mortality_altitude, sp_11]
N = 250 Bandwidth = 0.08633
Trace of B[mortality_altitude, sp_12]
Iterations
Density of B[mortality_altitude, sp_12]
N = 250 Bandwidth = 0.05749

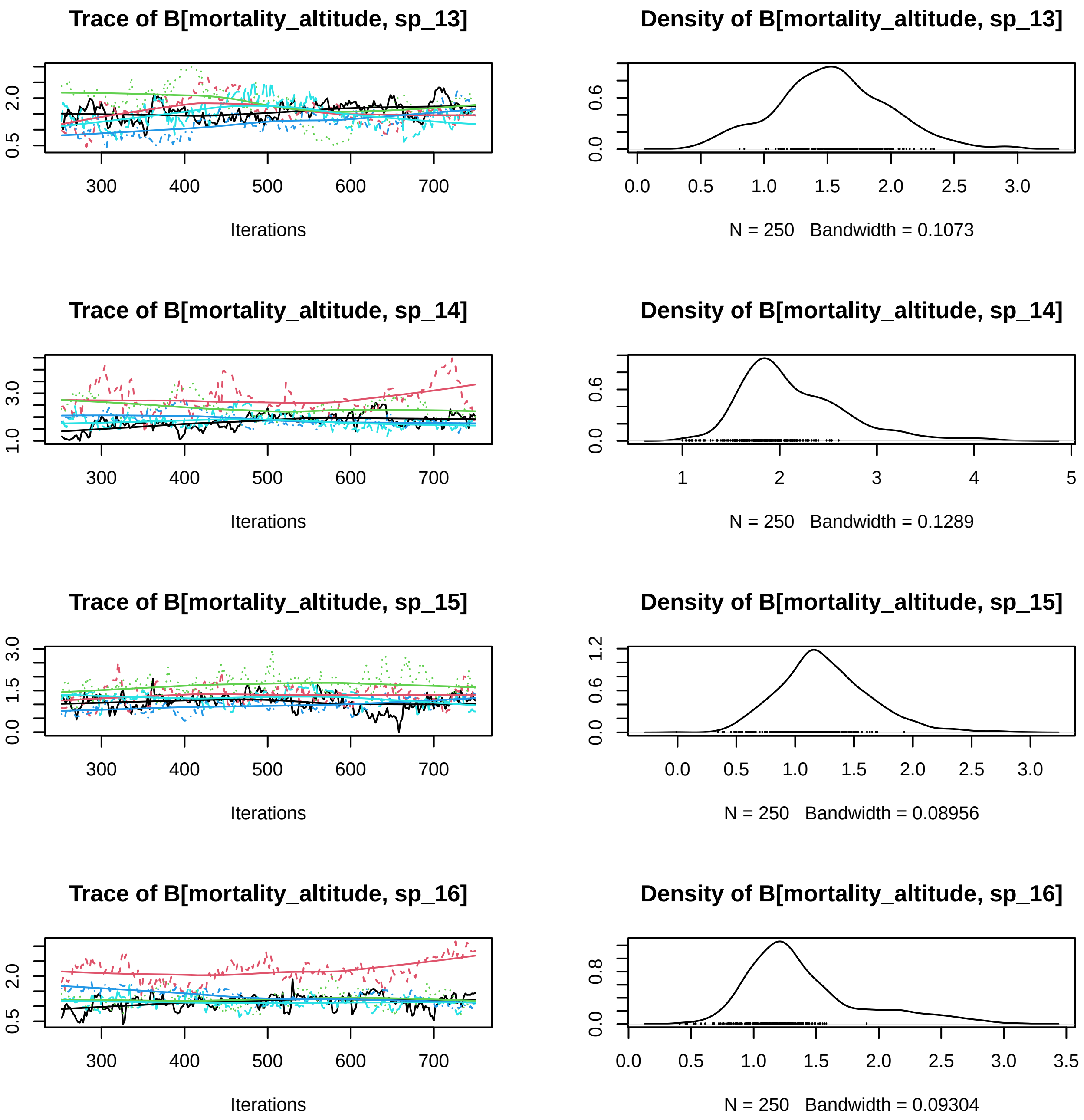

Trace of B[mortality_altitude, sp_13]
Iterations
Density of B[mortality_altitude, sp_13]
N = 250 Bandwidth = 0.1073
Trace of B[mortality_altitude, sp_14]
Iterations
Density of B[mortality_altitude, sp_14]
N = 250 Bandwidth = 0.1289
Trace of B[mortality_altitude, sp_15]
Iterations
Density of B[mortality_altitude, sp_15]
N = 250 Bandwidth = 0.08956
Trace of B[mortality_altitude, sp_16]
Iterations
Density of B[mortality_altitude, sp_16]
N = 250 Bandwidth = 0.09304

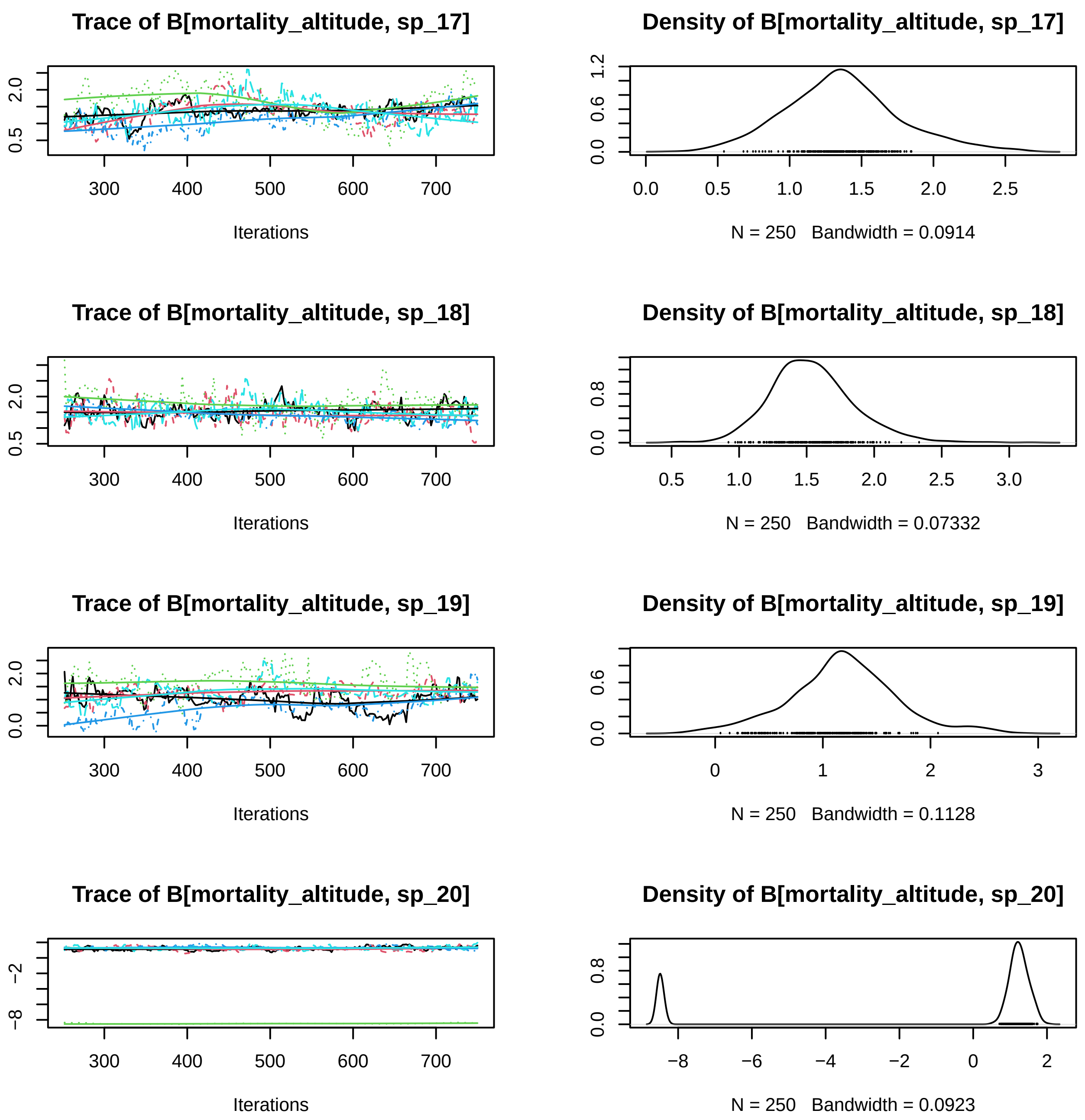
Trace of B[mortality_altitude, sp_17]
Density of B[mortality_altitude, sp_17]
0.5
2.0
300
400
500
600
700
Iterations
0.0
0.6
1.2
0.0
0.5
1.0
1.5
2.0
2.5
N = 250 Bandwidth = 0.0914
Trace of B[mortality_altitude, sp_18]
Density of B[mortality_altitude, sp_18]
0.5
2.0
300
400
500
600
700
Iterations
0.0
0.8
0.5
1.0
1.5
2.0
2.5
3.0
N = 250 Bandwidth = 0.07332
Trace of B[mortality_altitude, sp_19]
Density of B[mortality_altitude, sp_19]
0.0
2.0
300
400
500
600
700
Iterations
0.0
0.6
0
1
2
3
N = 250 Bandwidth = 0.1128
Trace of B[mortality_altitude, sp_20]
Density of B[mortality_altitude, sp_20]
−8
−2
300
400
500
600
700
Iterations
0.0
0.8
−8
−6
−4
−2
0
2
N = 250 Bandwidth = 0.0923

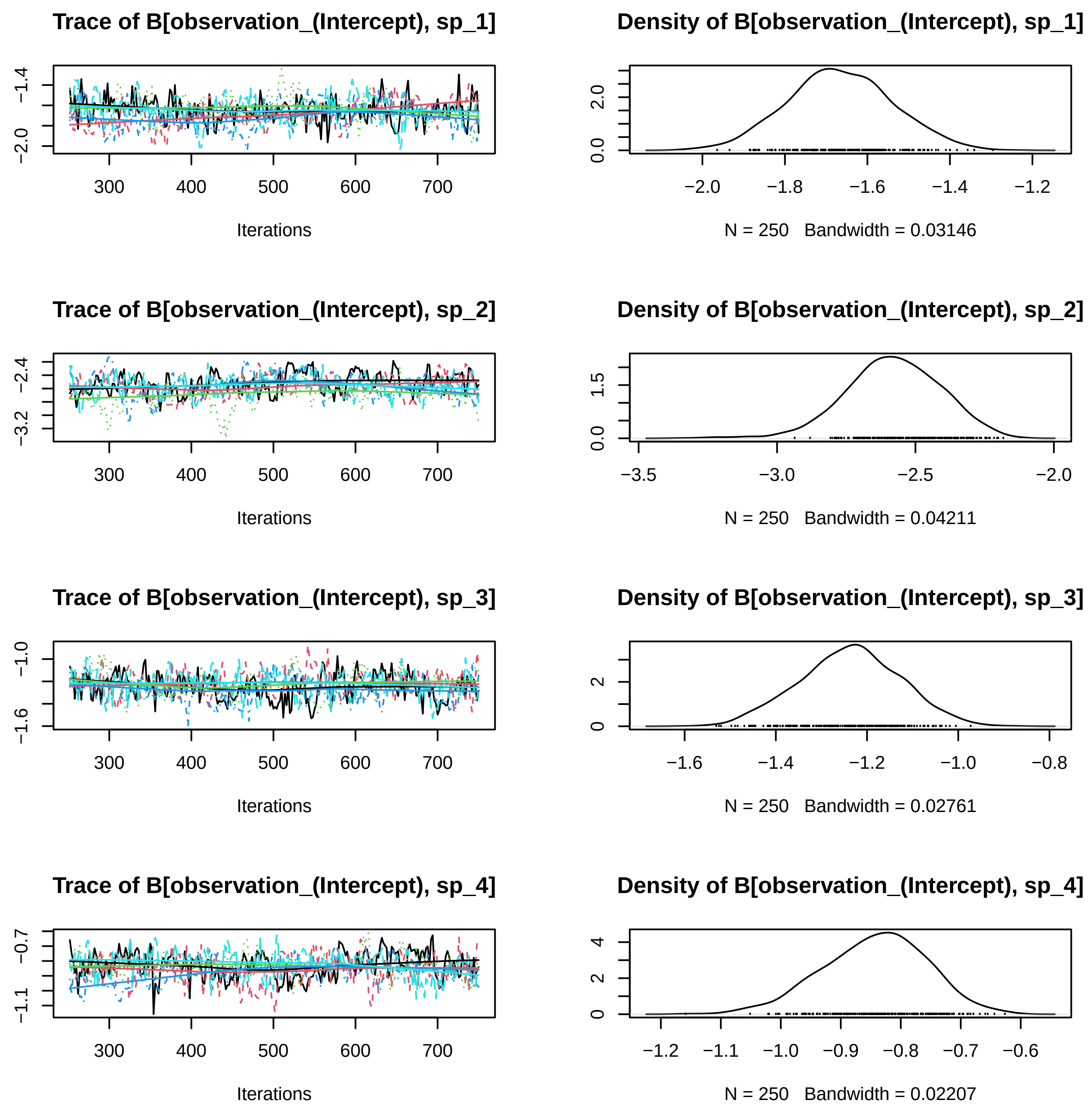

Trace of B[observation_(Intercept), sp_1]
−2.0
−1.4
300
400
500
600
700
Iterations
Density of B[observation_(Intercept), sp_1]
0.0
2.0
−2.0
−1.8
−1.6
−1.4
−1.2
N = 250 Bandwidth = 0.03146
Trace of B[observation_(Intercept), sp_2]
−3.2
−2.4
300
400
500
600
700
Iterations
Density of B[observation_(Intercept), sp_2]
0.0
1.5
−3.5
−3.0
−2.5
−2.0
N = 250 Bandwidth = 0.04211
Trace of B[observation_(Intercept), sp_3]
−1.6
−1.0
300
400
500
600
700
Iterations
Density of B[observation_(Intercept), sp_3]
0
2
−1.6
−1.4
−1.2
−1.0
−0.8
N = 250 Bandwidth = 0.02761
Trace of B[observation_(Intercept), sp_4]
−1.1
−0.7
300
400
500
600
700
Iterations
Density of B[observation_(Intercept), sp_4]
0
2
4
−1.2
−1.1
−1.0
−0.9
−0.8
−0.7
−0.6
N = 250 Bandwidth = 0.02207

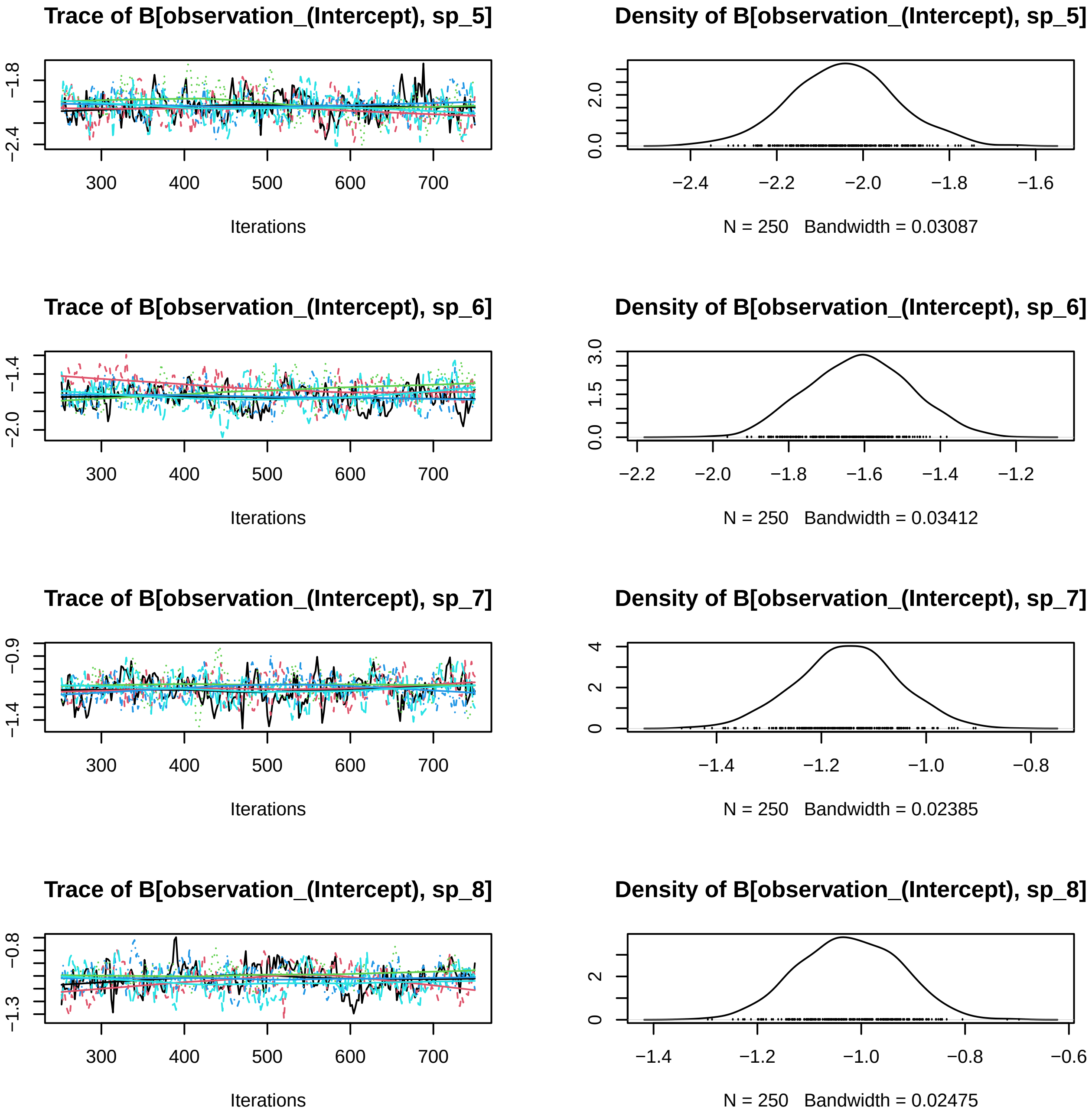

Trace of B[observation_(Intercept), sp_5]
−1.8
−2.4
300
400
500
600
700
Iterations
Density of B[observation_(Intercept), sp_5]
0.0
2.0
−2.4
−2.2
−2.0
−1.8
−1.6
N = 250 Bandwidth = 0.03087
Trace of B[observation_(Intercept), sp_6]
−1.4
−2.0
300
400
500
600
700
Iterations
Density of B[observation_(Intercept), sp_6]
0.0
1.5
3.0
−2.2
−2.0
−1.8
−1.6
−1.4
−1.2
N = 250 Bandwidth = 0.03412
Trace of B[observation_(Intercept), sp_7]
−0.9
−1.4
300
400
500
600
700
Iterations
Density of B[observation_(Intercept), sp_7]
0
2
4
−1.4
−1.2
−1.0
−0.8
N = 250 Bandwidth = 0.02385
Trace of B[observation_(Intercept), sp_8]
−0.8
−1.3
300
400
500
600
700
Iterations
Density of B[observation_(Intercept), sp_8]
0
2
−1.4
−1.2
−1.0
−0.8
−0.6
N = 250 Bandwidth = 0.02475

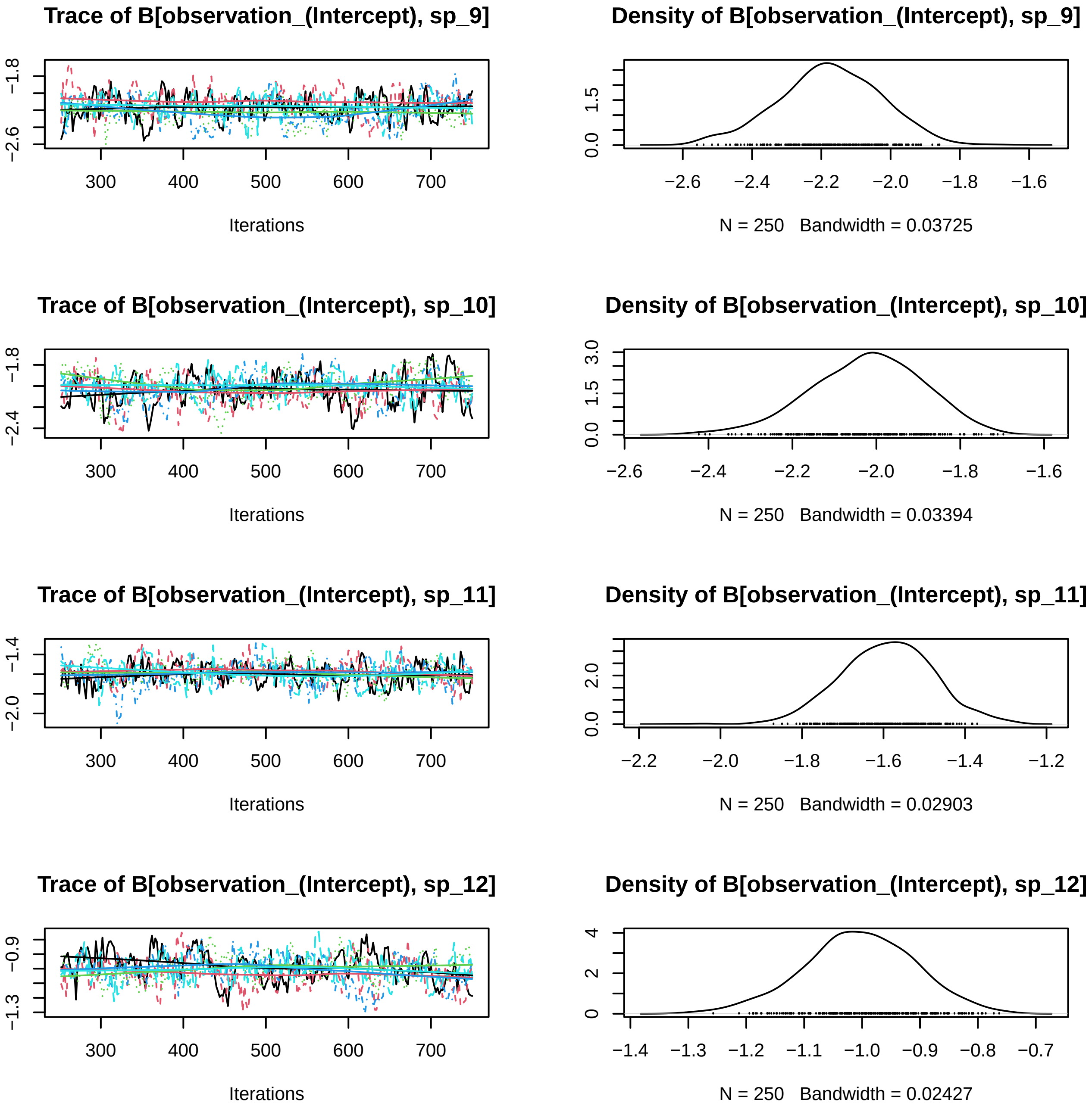

Trace of B[observation_(Intercept), sp_9]
−2.6
−1.8
300
400
500
600
700
Iterations
Density of B[observation_(Intercept), sp_9]
0.0
1.5
−2.6
−2.4
−2.2
−2.0
−1.8
−1.6
N = 250 Bandwidth = 0.03725
Trace of B[observation_(Intercept), sp_10]
−2.4
−1.8
300
400
500
600
700
Iterations
Density of B[observation_(Intercept), sp_10]
0.0
1.5
3.0
−2.6
−2.4
−2.2
−2.0
−1.8
−1.6
N = 250 Bandwidth = 0.03394
Trace of B[observation_(Intercept), sp_11]
−2.0
−1.4
300
400
500
600
700
Iterations
Density of B[observation_(Intercept), sp_11]
0.0
2.0
−2.2
−2.0
−1.8
−1.6
−1.4
−1.2
N = 250 Bandwidth = 0.02903
Trace of B[observation_(Intercept), sp_12]
−1.3
−0.9
300
400
500
600
700
Iterations
Density of B[observation_(Intercept), sp_12]
0
2
4
−1.4
−1.3
−1.2
−1.1
−1.0
−0.9
−0.8
−0.7
N = 250 Bandwidth = 0.02427

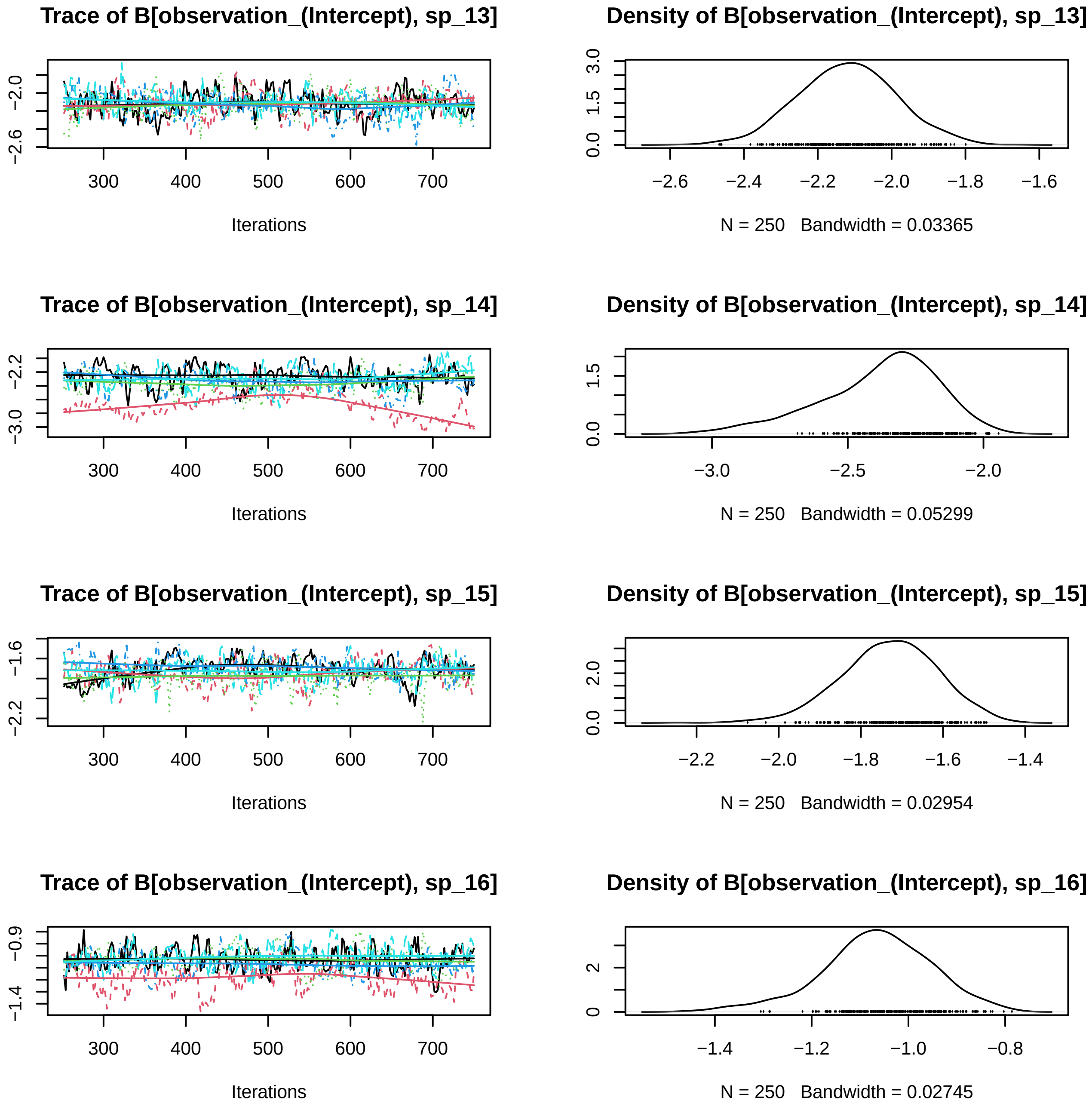
Trace of B[observation_(Intercept), sp_13]
Iterations
Density of B[observation_(Intercept), sp_13]
N = 250   Bandwidth = 0.03365
Trace of B[observation_(Intercept), sp_14]
Iterations
Density of B[observation_(Intercept), sp_14]
N = 250   Bandwidth = 0.05299
Trace of B[observation_(Intercept), sp_15]
Iterations
Density of B[observation_(Intercept), sp_15]
N = 250   Bandwidth = 0.02954
Trace of B[observation_(Intercept), sp_16]
Iterations
Density of B[observation_(Intercept), sp_16]
N = 250   Bandwidth = 0.02745

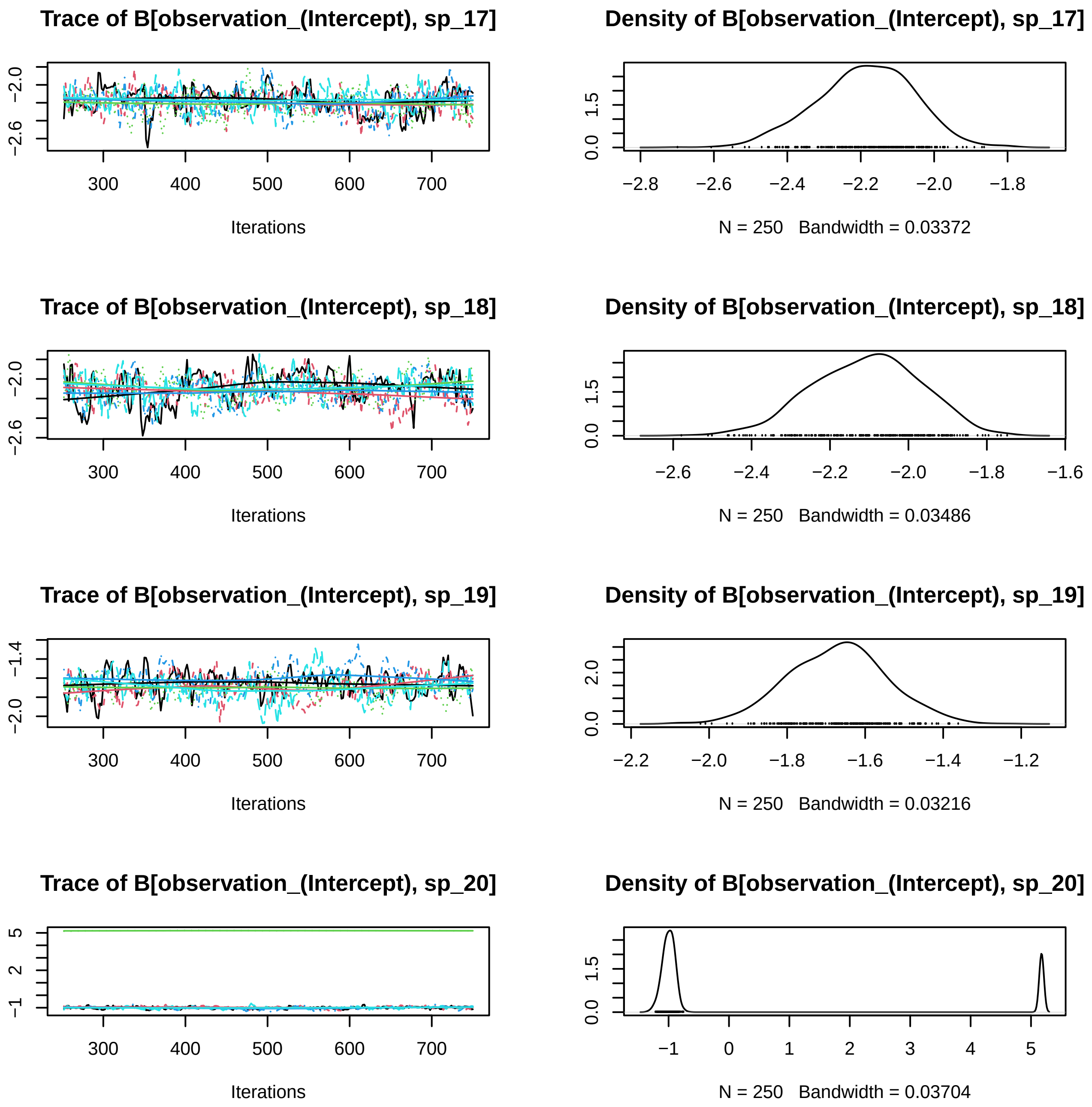

Trace of B[observation_(Intercept), sp_17]
Iterations
Density of B[observation_(Intercept), sp_17]
N = 250 Bandwidth = 0.03372
Trace of B[observation_(Intercept), sp_18]
Iterations
Density of B[observation_(Intercept), sp_18]
N = 250 Bandwidth = 0.03486
Trace of B[observation_(Intercept), sp_19]
Iterations
Density of B[observation_(Intercept), sp_19]
N = 250 Bandwidth = 0.03216
Trace of B[observation_(Intercept), sp_20]
Iterations
Density of B[observation_(Intercept), sp_20]
N = 250 Bandwidth = 0.03704

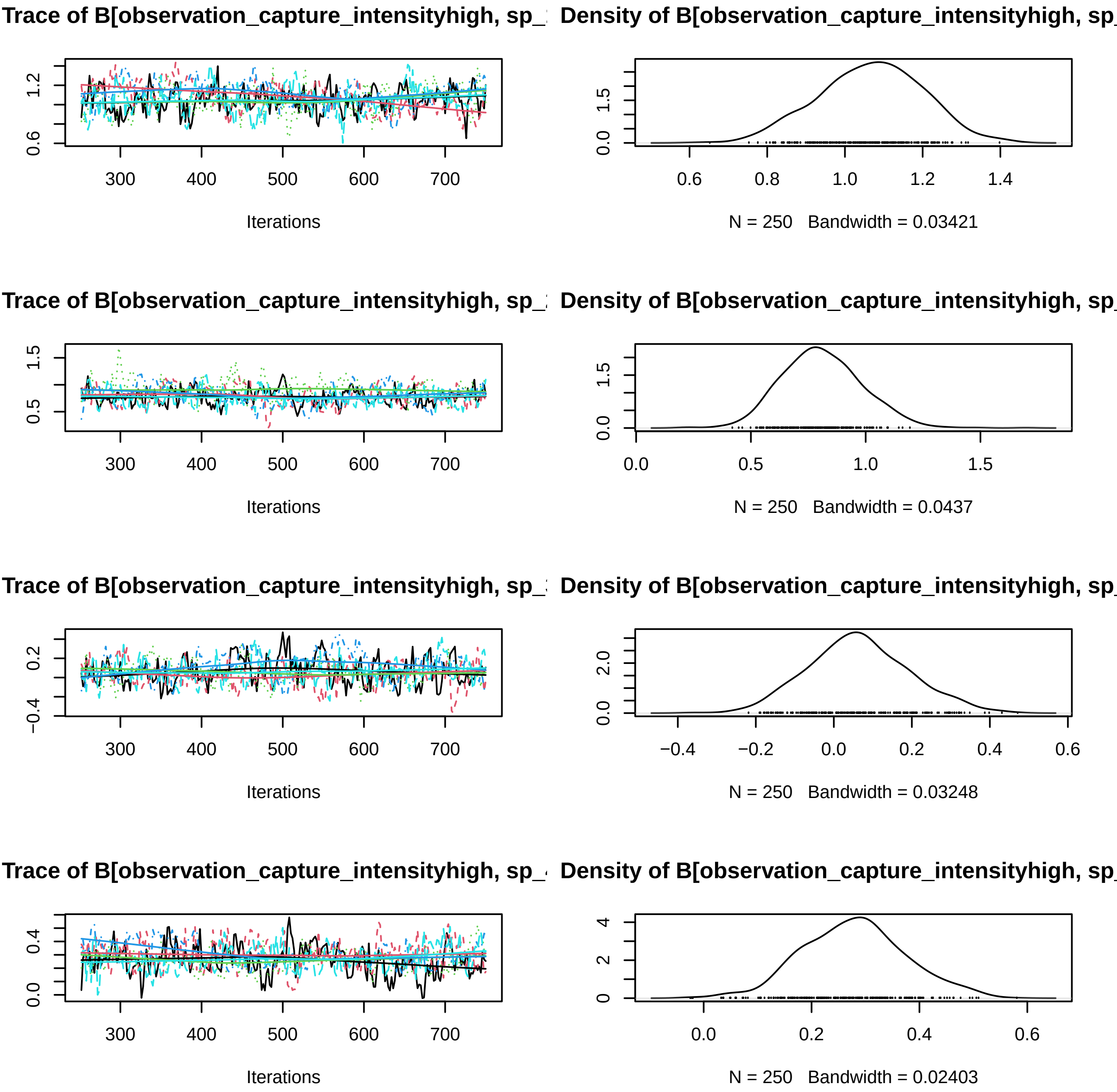
Trace of B[observation_capture_intensityhigh, sp_
Density of B[observation_capture_intensityhigh, sp_
Iterations
N = 250 Bandwidth = 0.03421
Trace of B[observation_capture_intensityhigh, sp_
Density of B[observation_capture_intensityhigh, sp_
Iterations
N = 250 Bandwidth = 0.0437
Trace of B[observation_capture_intensityhigh, sp_
Density of B[observation_capture_intensityhigh, sp_
Iterations
N = 250 Bandwidth = 0.03248
Trace of B[observation_capture_intensityhigh, sp_
Density of B[observation_capture_intensityhigh, sp_
Iterations
N = 250 Bandwidth = 0.02403

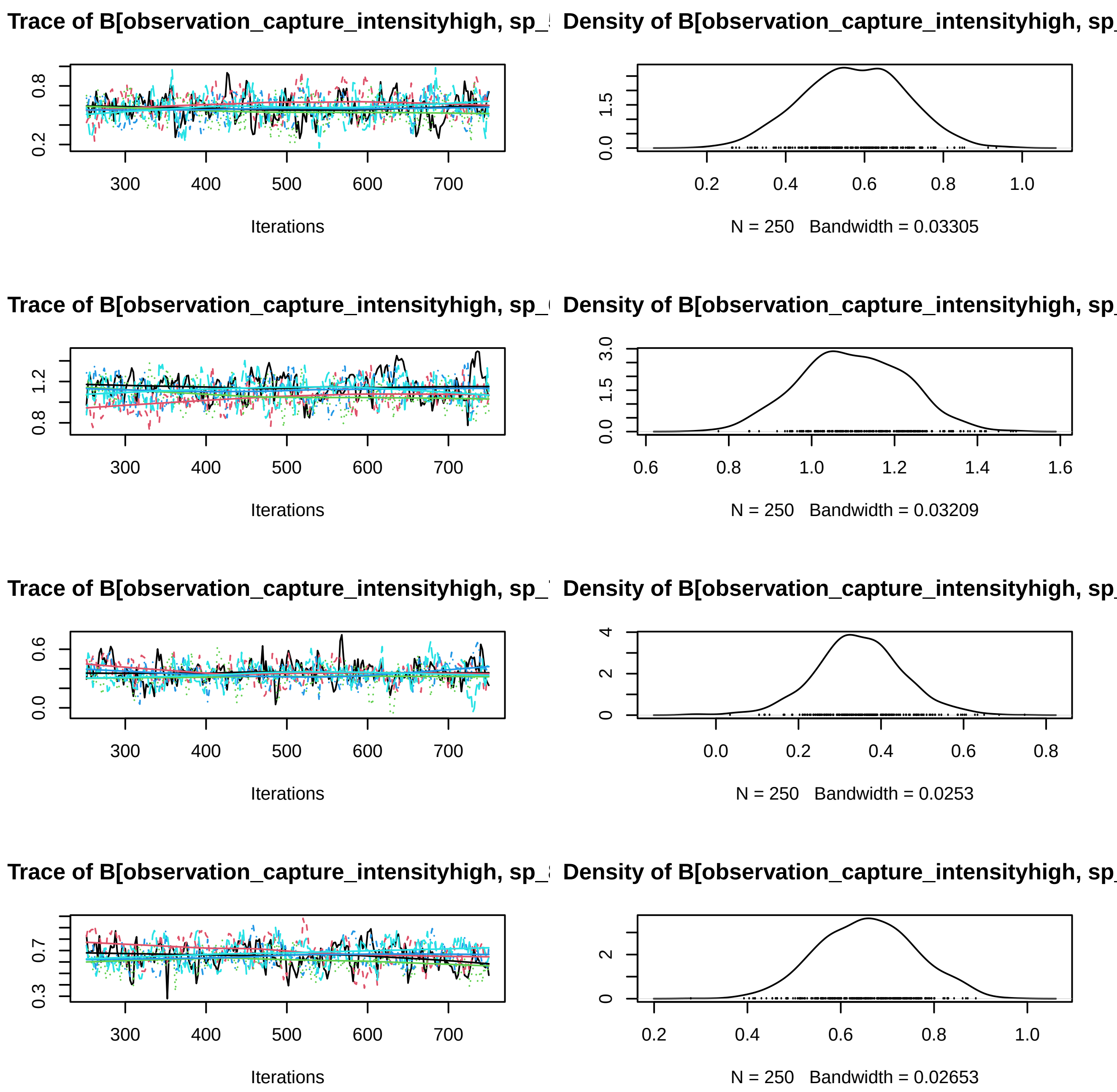
Trace of B[observation_capture_intensityhigh, sp_
Density of B[observation_capture_intensityhigh, sp_
Iterations
N = 250 Bandwidth = 0.03305
Trace of B[observation_capture_intensityhigh, sp_
Density of B[observation_capture_intensityhigh, sp_
Iterations
N = 250 Bandwidth = 0.03209
Trace of B[observation_capture_intensityhigh, sp_
Density of B[observation_capture_intensityhigh, sp_
Iterations
N = 250 Bandwidth = 0.0253
Trace of B[observation_capture_intensityhigh, sp_
Density of B[observation_capture_intensityhigh, sp_
Iterations
N = 250 Bandwidth = 0.02653

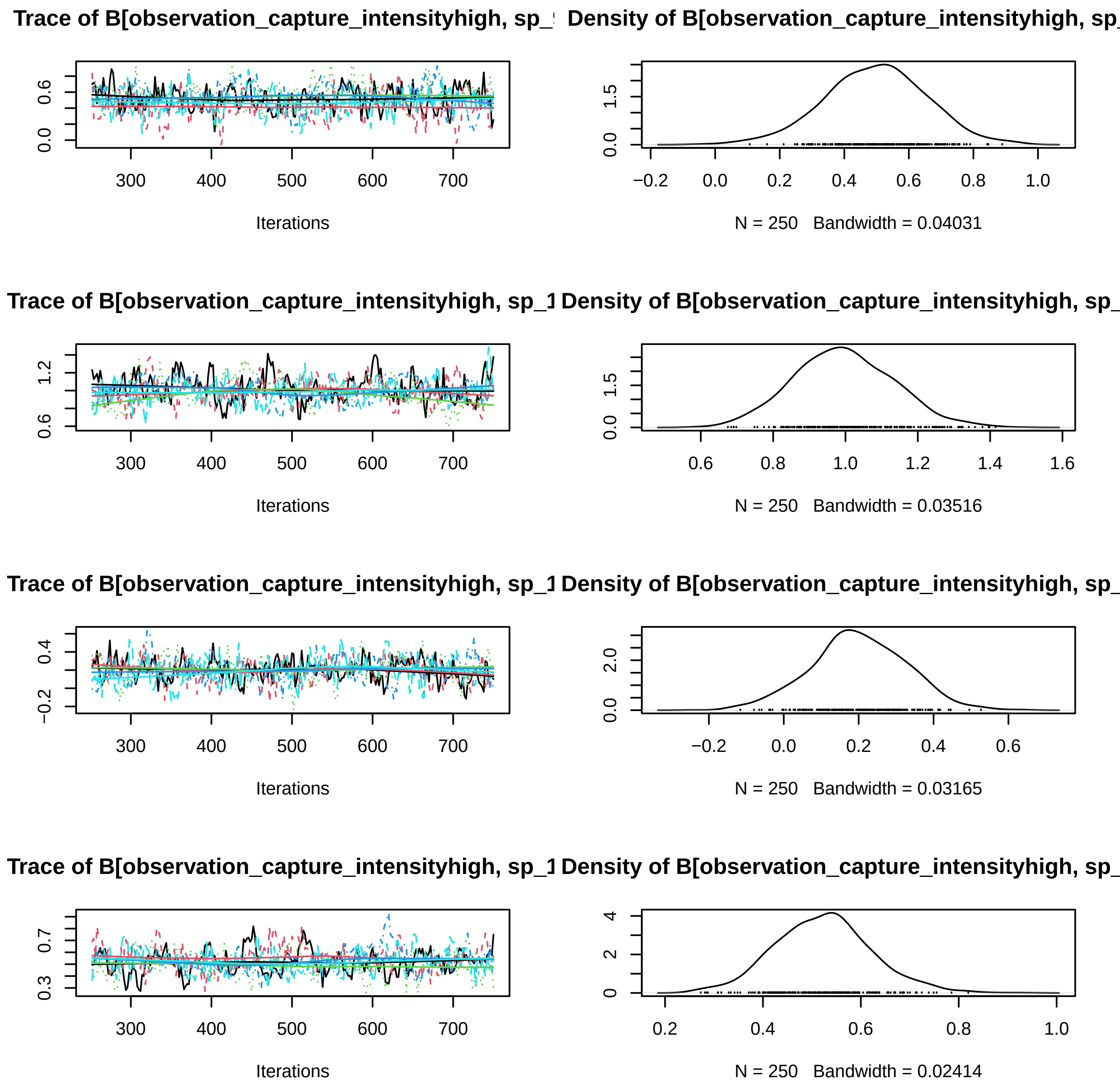
Trace of B[observation_capture_intensityhigh, sp_
Density of B[observation_capture_intensityhigh, sp_
Iterations
N = 250 Bandwidth = 0.04031
Trace of B[observation_capture_intensityhigh, sp_1
Density of B[observation_capture_intensityhigh, sp_
Iterations
N = 250 Bandwidth = 0.03516
Trace of B[observation_capture_intensityhigh, sp_1
Density of B[observation_capture_intensityhigh, sp_
Iterations
N = 250 Bandwidth = 0.03165
Trace of B[observation_capture_intensityhigh, sp_1
Density of B[observation_capture_intensityhigh, sp_
Iterations
N = 250 Bandwidth = 0.02414

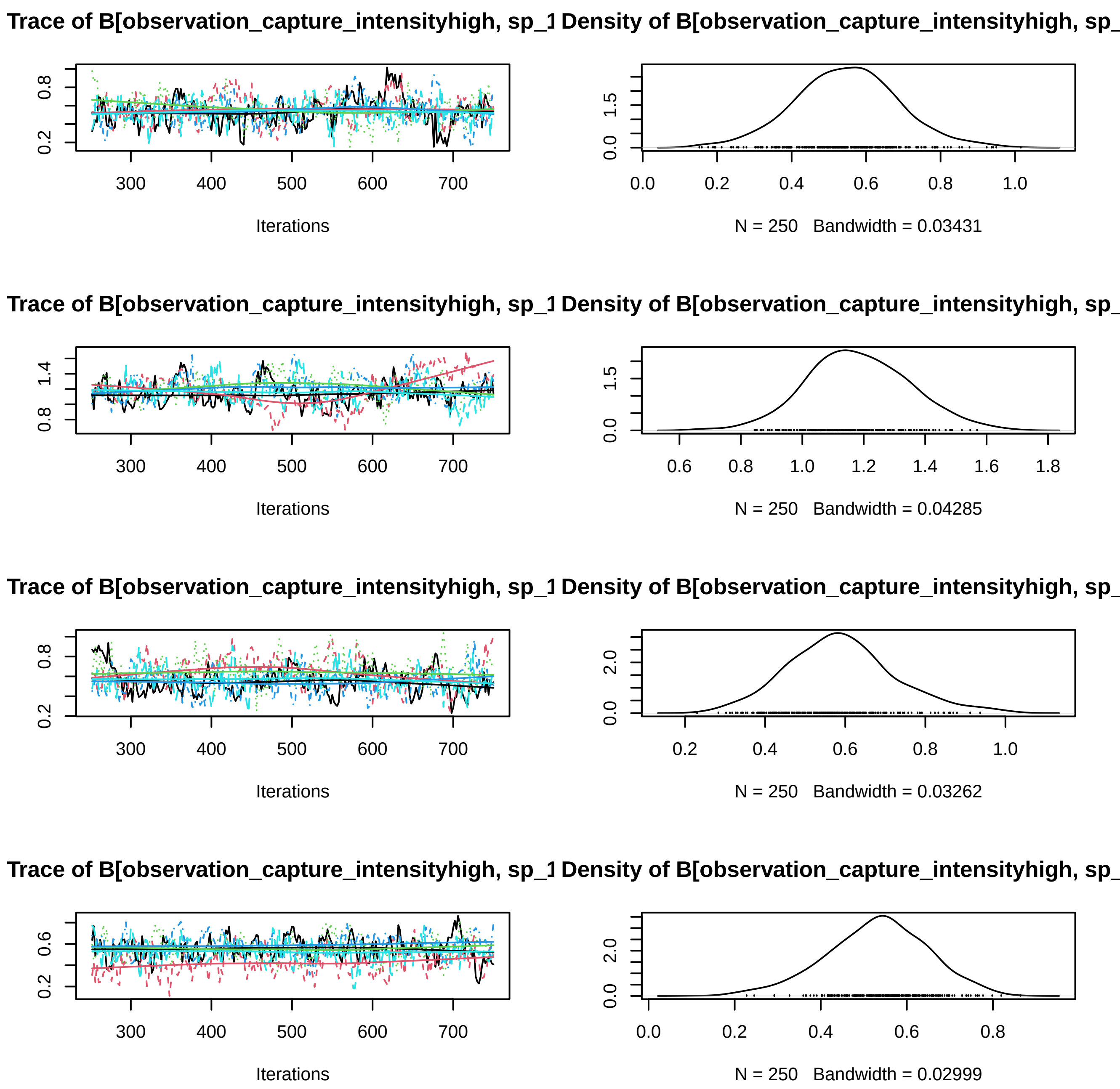
Trace of B[observation_capture_intensityhigh, sp_1
Density of B[observation_capture_intensityhigh, sp_
Iterations
N = 250 Bandwidth = 0.03431
Trace of B[observation_capture_intensityhigh, sp_1
Density of B[observation_capture_intensityhigh, sp_
Iterations
N = 250 Bandwidth = 0.04285
Trace of B[observation_capture_intensityhigh, sp_1
Density of B[observation_capture_intensityhigh, sp_
Iterations
N = 250 Bandwidth = 0.03262
Trace of B[observation_capture_intensityhigh, sp_1
Density of B[observation_capture_intensityhigh, sp_
Iterations
N = 250 Bandwidth = 0.02999

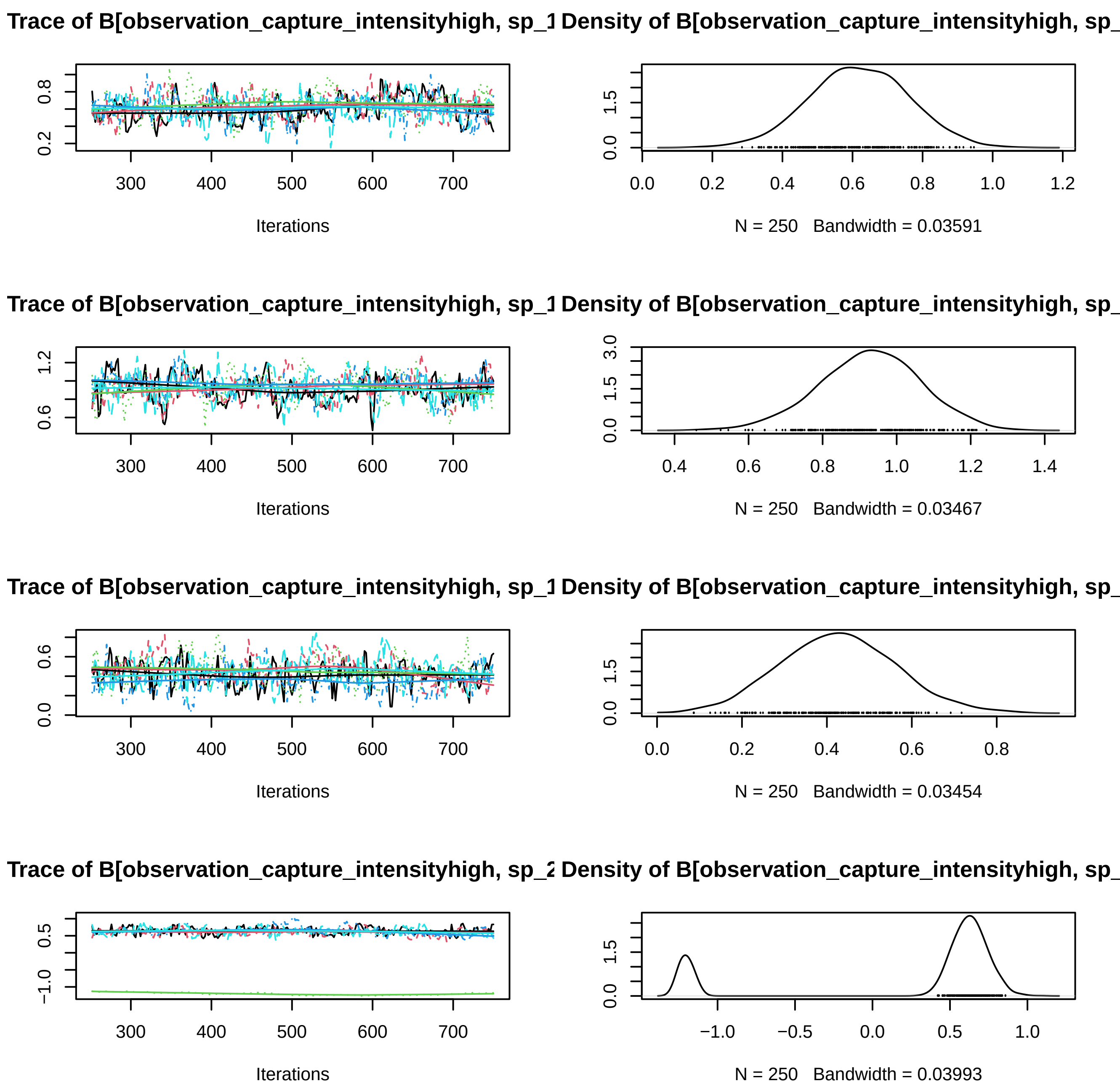
Trace of B[observation_capture_intensityhigh, sp_1
Density of B[observation_capture_intensityhigh, sp_
Iterations
N = 250 Bandwidth = 0.03591
Trace of B[observation_capture_intensityhigh, sp_1
Density of B[observation_capture_intensityhigh, sp_
Iterations
N = 250 Bandwidth = 0.03467
Trace of B[observation_capture_intensityhigh, sp_1
Density of B[observation_capture_intensityhigh, sp_
Iterations
N = 250 Bandwidth = 0.03454
Trace of B[observation_capture_intensityhigh, sp_2
Density of B[observation_capture_intensityhigh, sp_
Iterations
N = 250 Bandwidth = 0.03993

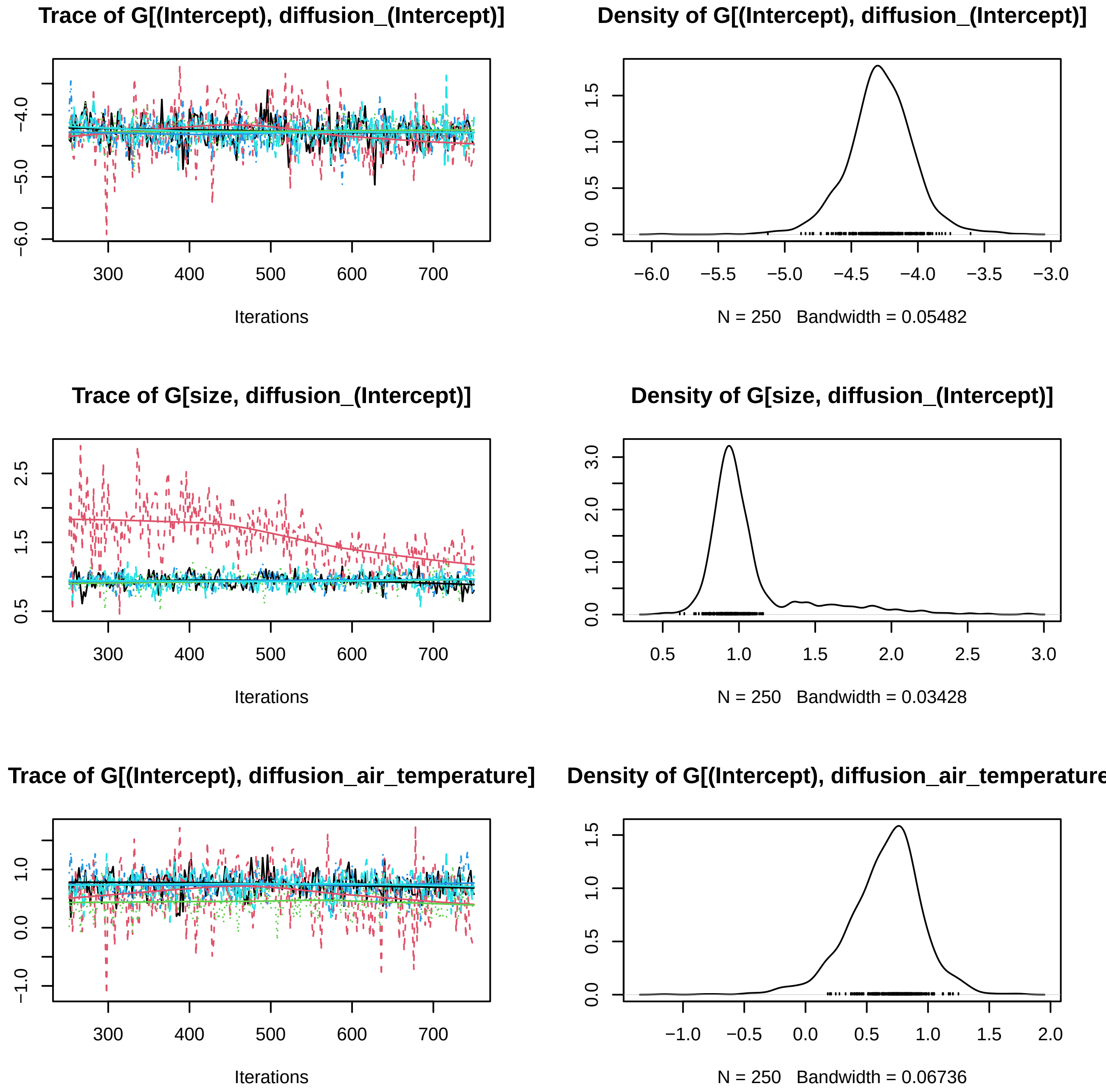
Trace of G[(Intercept), diffusion_(Intercept)]
Iterations
Density of G[(Intercept), diffusion_(Intercept)]
N = 250 Bandwidth = 0.05482
Trace of G[size, diffusion_(Intercept)]
Iterations
Density of G[size, diffusion_(Intercept)]
N = 250 Bandwidth = 0.03428
Trace of G[(Intercept), diffusion_air_temperature]
Iterations
Density of G[(Intercept), diffusion_air_temperature
N = 250 Bandwidth = 0.06736

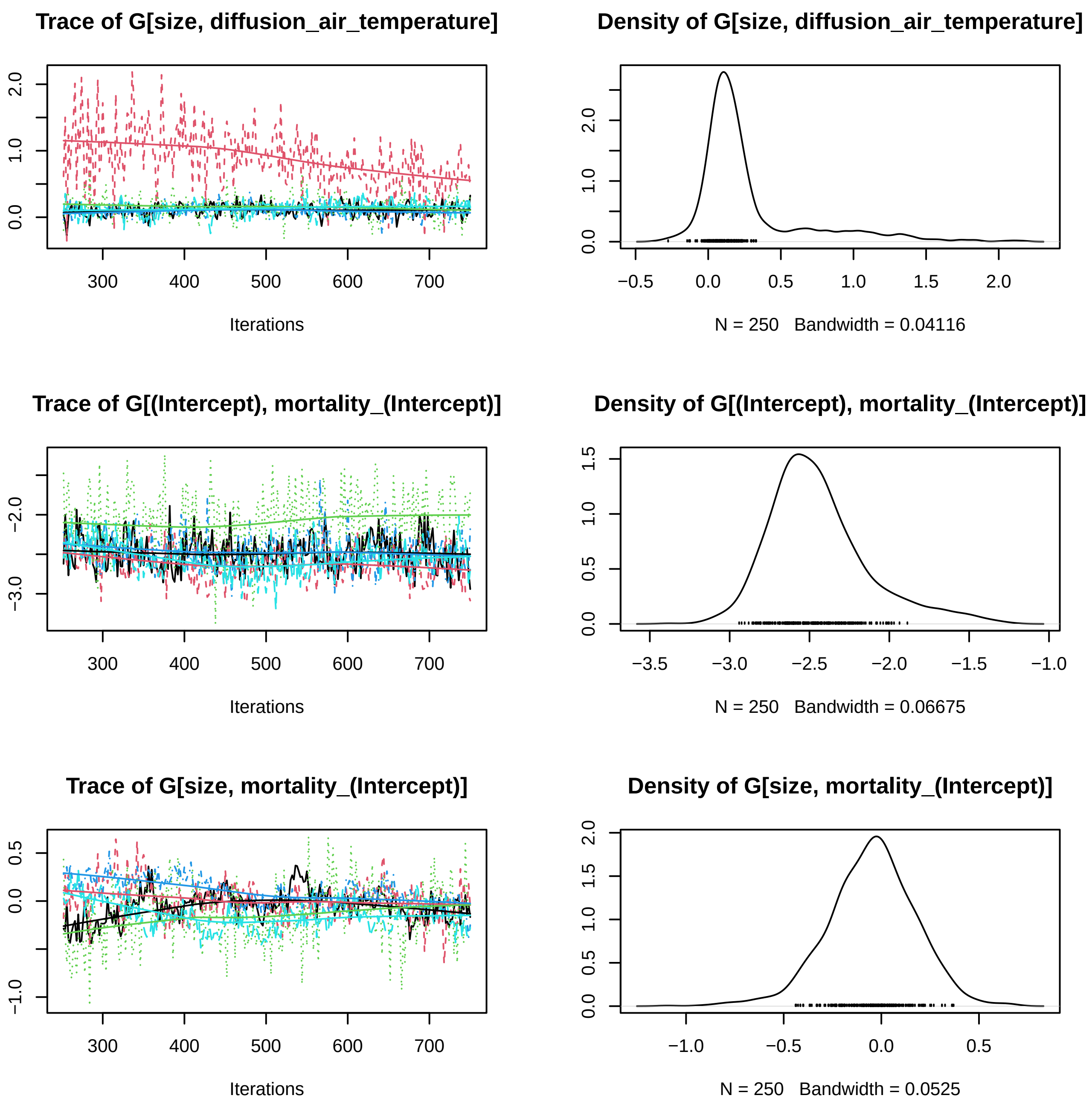

Trace of G[size, diffusion_air_temperature]
0.0
1.0
2.0
300
400
500
600
700
Iterations
Density of G[size, diffusion_air_temperature]
0.0
1.0
2.0
−0.5
0.0
0.5
1.0
1.5
2.0
N = 250 Bandwidth = 0.04116
Trace of G[(Intercept), mortality_(Intercept)]
−3.0
−2.0
300
400
500
600
700
Iterations
Density of G[(Intercept), mortality_(Intercept)]
0.0
0.5
1.0
1.5
−3.5
−3.0
−2.5
−2.0
−1.5
−1.0
N = 250 Bandwidth = 0.06675
Trace of G[size, mortality_(Intercept)]
−1.0
0.0
0.5
300
400
500
600
700
Iterations
Density of G[size, mortality_(Intercept)]
0.0
0.5
1.0
1.5
2.0
−1.0
−0.5
0.0
0.5
N = 250 Bandwidth = 0.0525

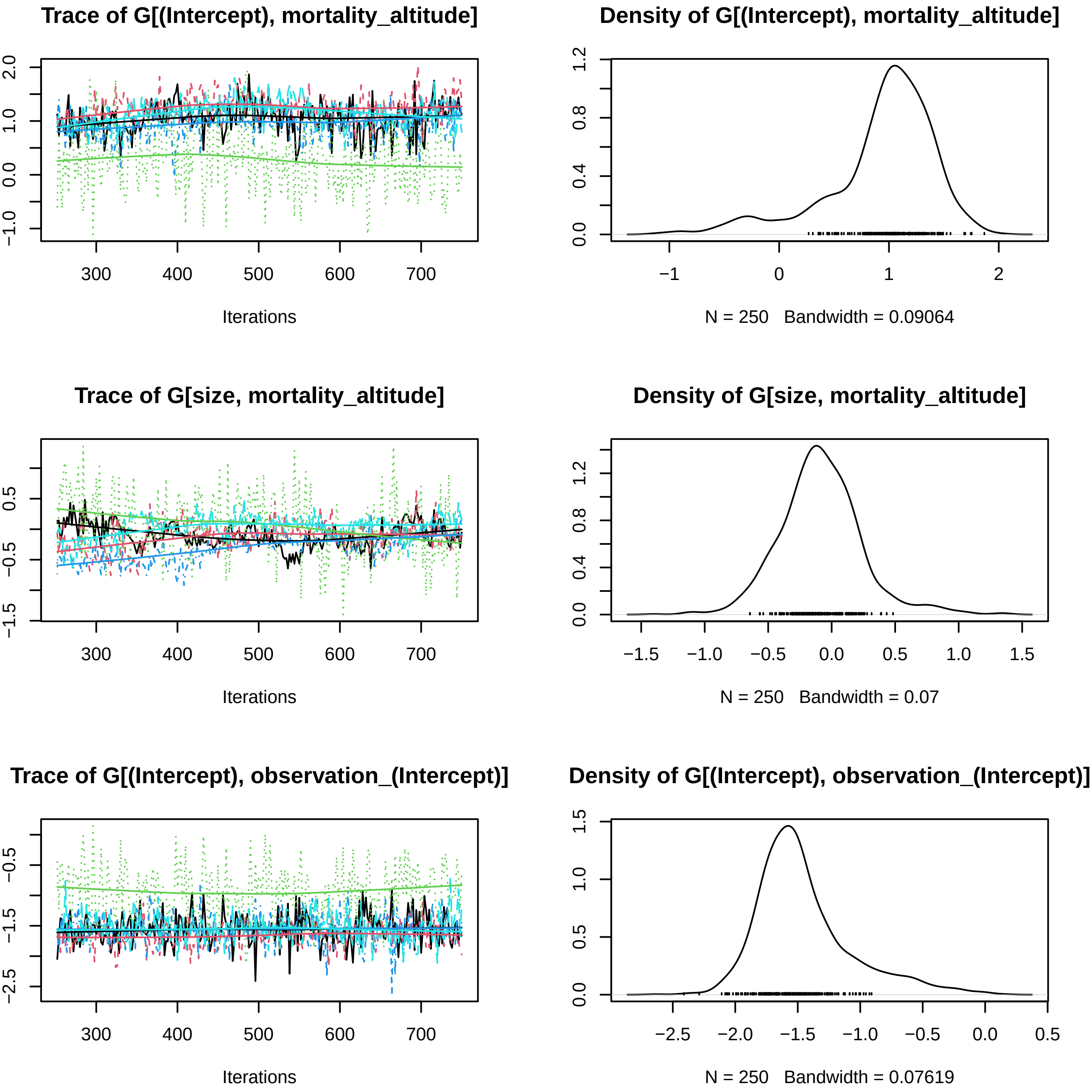

Trace of G[(Intercept), mortality_altitude]
Iterations
Density of G[(Intercept), mortality_altitude]
N = 250 Bandwidth = 0.09064
Trace of G[size, mortality_altitude]
Iterations
Density of G[size, mortality_altitude]
N = 250 Bandwidth = 0.07
Trace of G[(Intercept), observation_(Intercept)]
Iterations
Density of G[(Intercept), observation_(Intercept)]
N = 250 Bandwidth = 0.07619

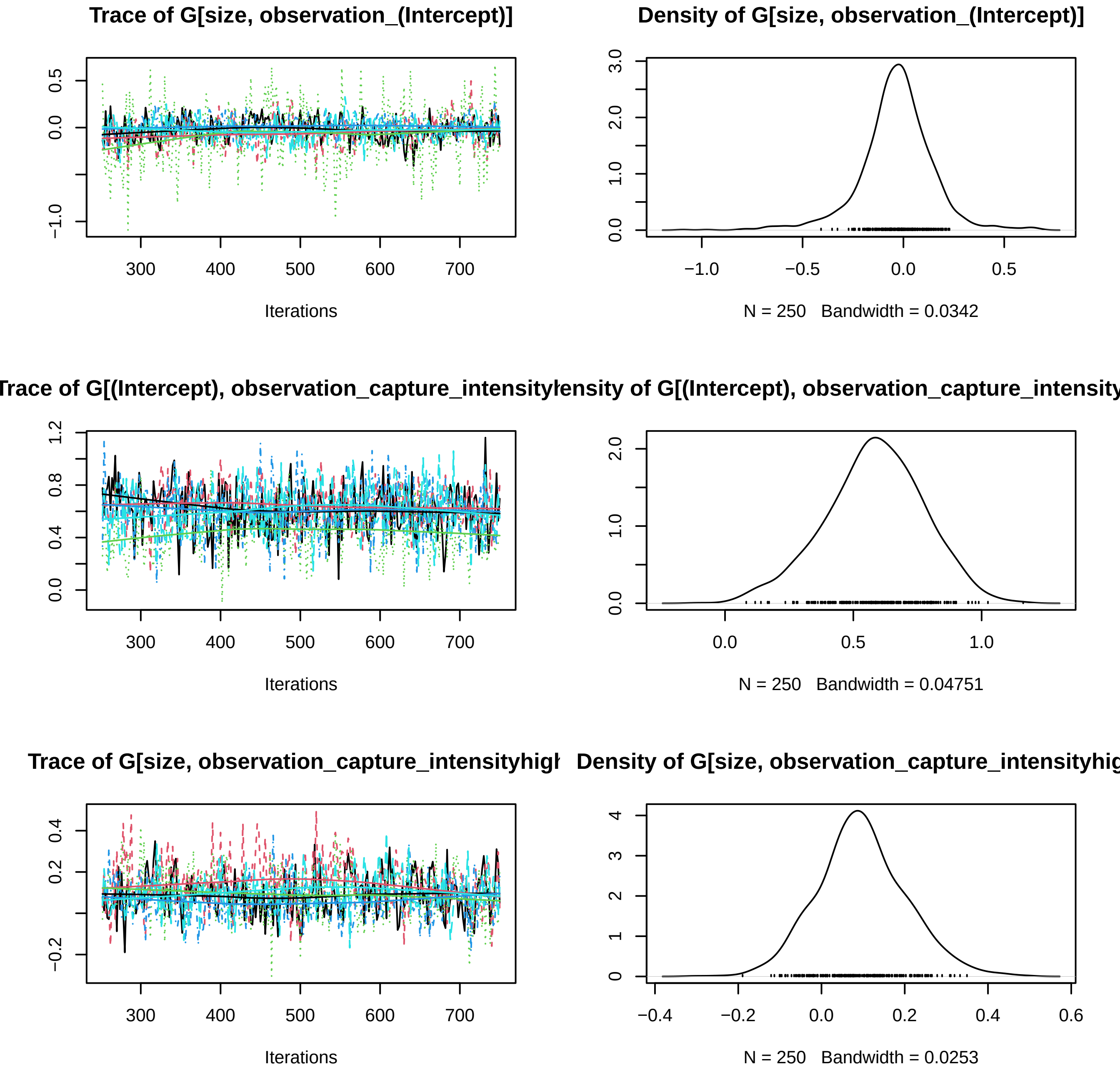
Trace of G[size, observation_(Intercept)]
0.5
0.0
−1.0
300
400
500
600
700
Iterations
Density of G[size, observation_(Intercept)]
3.0
2.0
1.0
0.0
−1.0
−0.5
0.0
0.5
N = 250 Bandwidth = 0.0342
Trace of G[(Intercept), observation_capture_intensity
1.2
0.8
0.4
0.0
300
400
500
600
700
Iterations
ensity of G[(Intercept), observation_capture_intensity
2.0
1.0
0.0
0.0
0.5
1.0
N = 250 Bandwidth = 0.04751
Trace of G[size, observation_capture_intensityhigh
0.4
0.2
−0.2
300
400
500
600
700
Iterations
Density of G[size, observation_capture_intensityhig
4
3
2
1
0
−0.4
−0.2
0.0
0.2
0.4
0.6
N = 250 Bandwidth = 0.0253

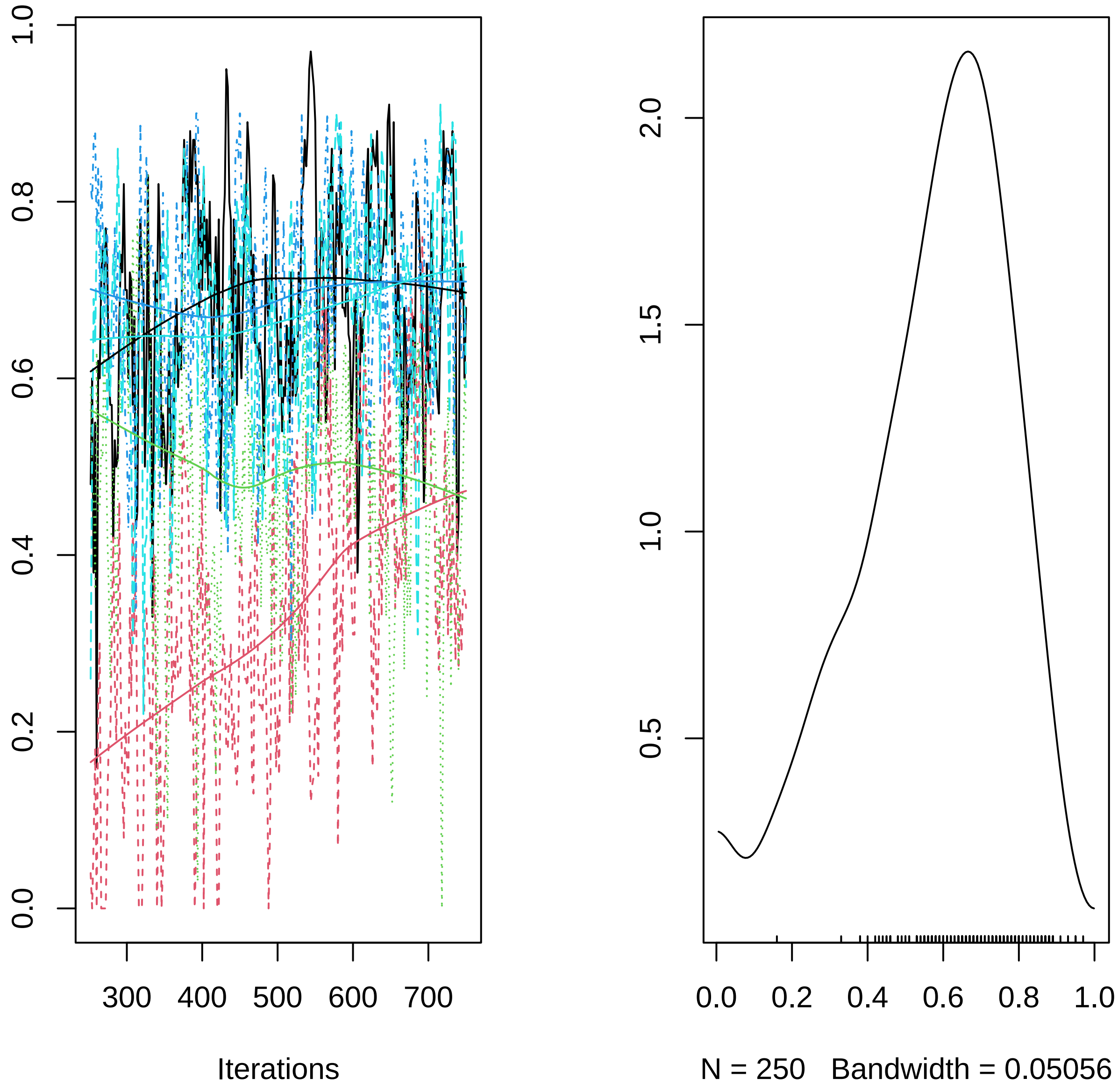

1.0
0.8
0.6
0.4
0.2
0.0
300
400
500
600
700
Iterations
2.0
1.5
1.0
0.5
0.0
0.2
0.4
0.6
0.8
1.0
N = 250   Bandwidth = 0.05056
