## Supplementary material for "R-package Jsmm: Joint species movement modelling of mark-recapture data": S2 Vignettes demonstrating the application of the R-package Jsmm: MCMC_convergence_simulated_ICP.pdf

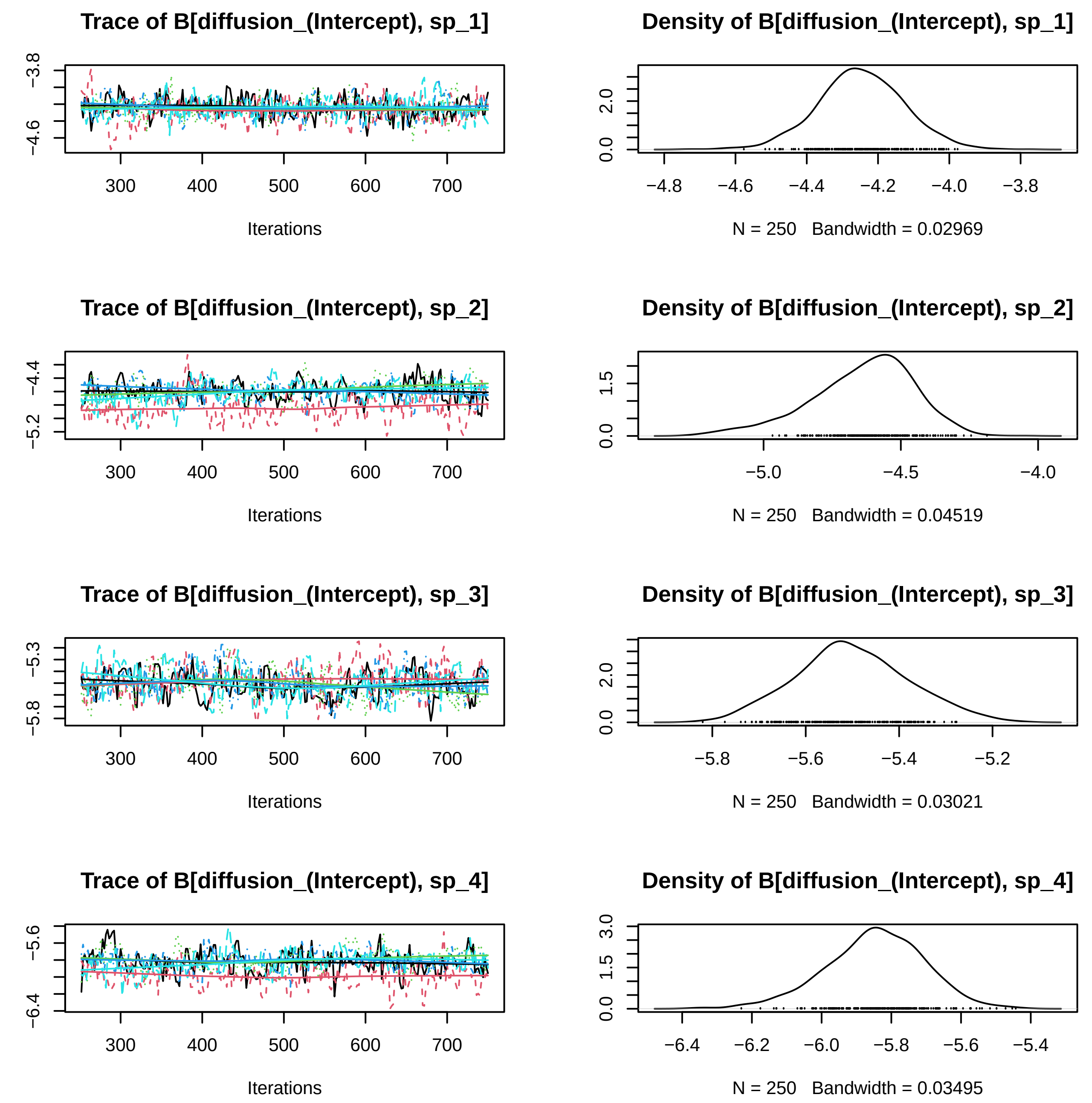
Trace of B[diffusion_(Intercept), sp_1]
Iterations
Density of B[diffusion_(Intercept), sp_1]
N = 250 Bandwidth = 0.02969
Trace of B[diffusion_(Intercept), sp_2]
Iterations
Density of B[diffusion_(Intercept), sp_2]
N = 250 Bandwidth = 0.04519
Trace of B[diffusion_(Intercept), sp_3]
Iterations
Density of B[diffusion_(Intercept), sp_3]
N = 250 Bandwidth = 0.03021
Trace of B[diffusion_(Intercept), sp_4]
Iterations
Density of B[diffusion_(Intercept), sp_4]
N = 250 Bandwidth = 0.03495

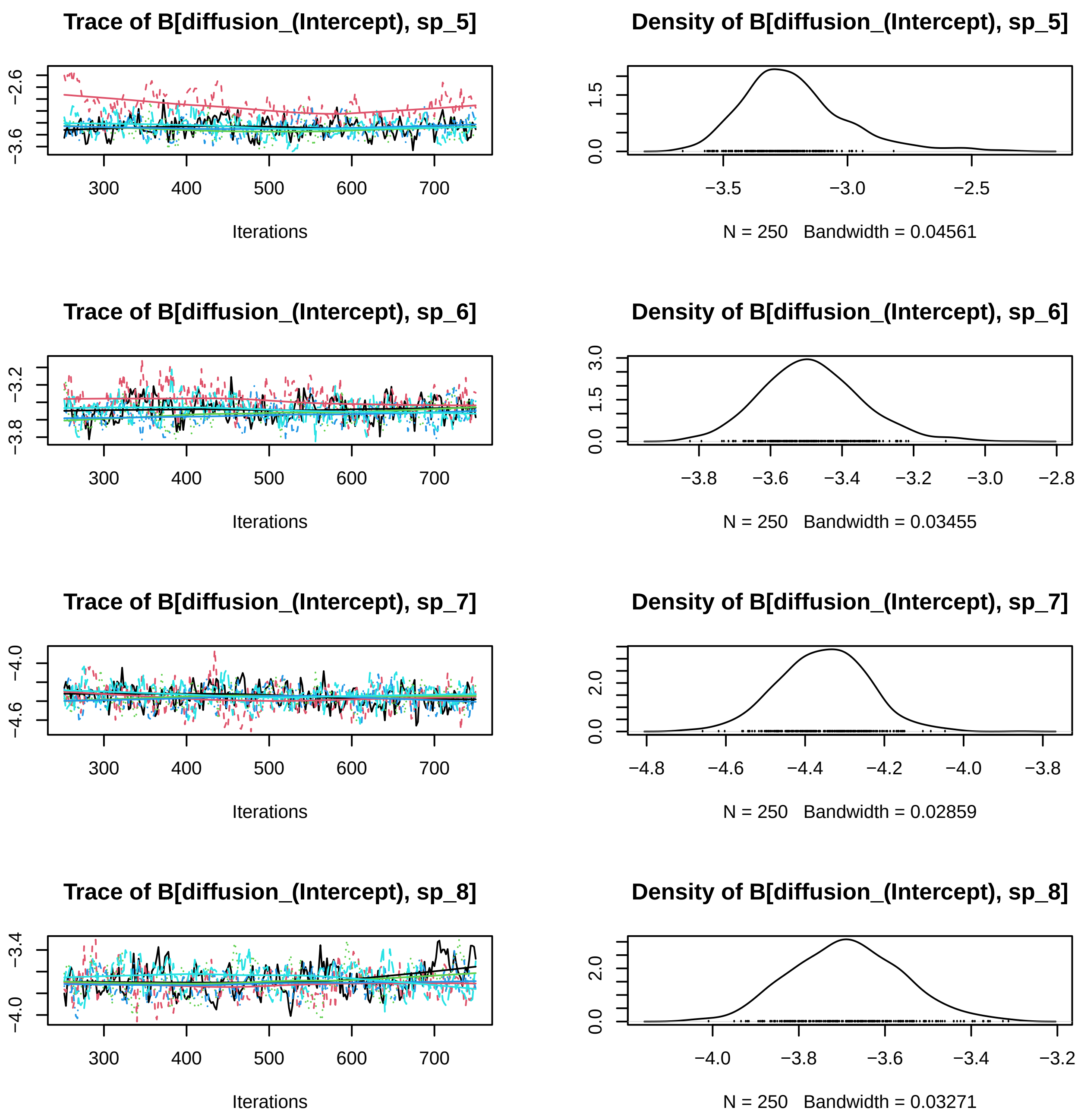
Trace of B[diffusion_(Intercept), sp_5]
Iterations
Density of B[diffusion_(Intercept), sp_5]
N = 250 Bandwidth = 0.04561
Trace of B[diffusion_(Intercept), sp_6]
Iterations
Density of B[diffusion_(Intercept), sp_6]
N = 250 Bandwidth = 0.03455
Trace of B[diffusion_(Intercept), sp_7]
Iterations
Density of B[diffusion_(Intercept), sp_7]
N = 250 Bandwidth = 0.02859
Trace of B[diffusion_(Intercept), sp_8]
Iterations
Density of B[diffusion_(Intercept), sp_8]
N = 250 Bandwidth = 0.03271

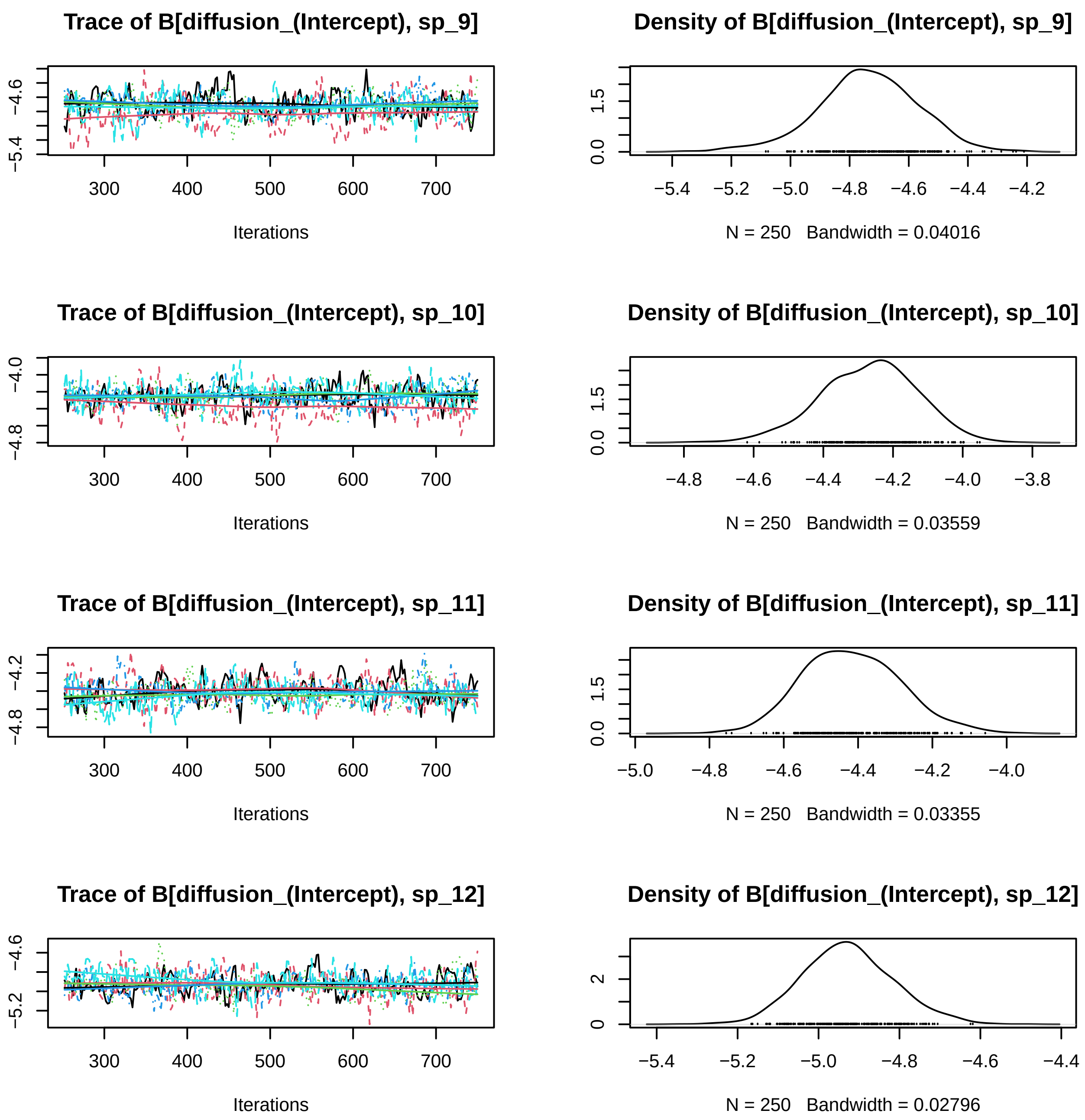
Trace of B[diffusion_(Intercept), sp_9]
Iterations
Density of B[diffusion_(Intercept), sp_9]
N = 250 Bandwidth = 0.04016
Trace of B[diffusion_(Intercept), sp_10]
Iterations
Density of B[diffusion_(Intercept), sp_10]
N = 250 Bandwidth = 0.03559
Trace of B[diffusion_(Intercept), sp_11]
Iterations
Density of B[diffusion_(Intercept), sp_11]
N = 250 Bandwidth = 0.03355
Trace of B[diffusion_(Intercept), sp_12]
Iterations
Density of B[diffusion_(Intercept), sp_12]
N = 250 Bandwidth = 0.02796

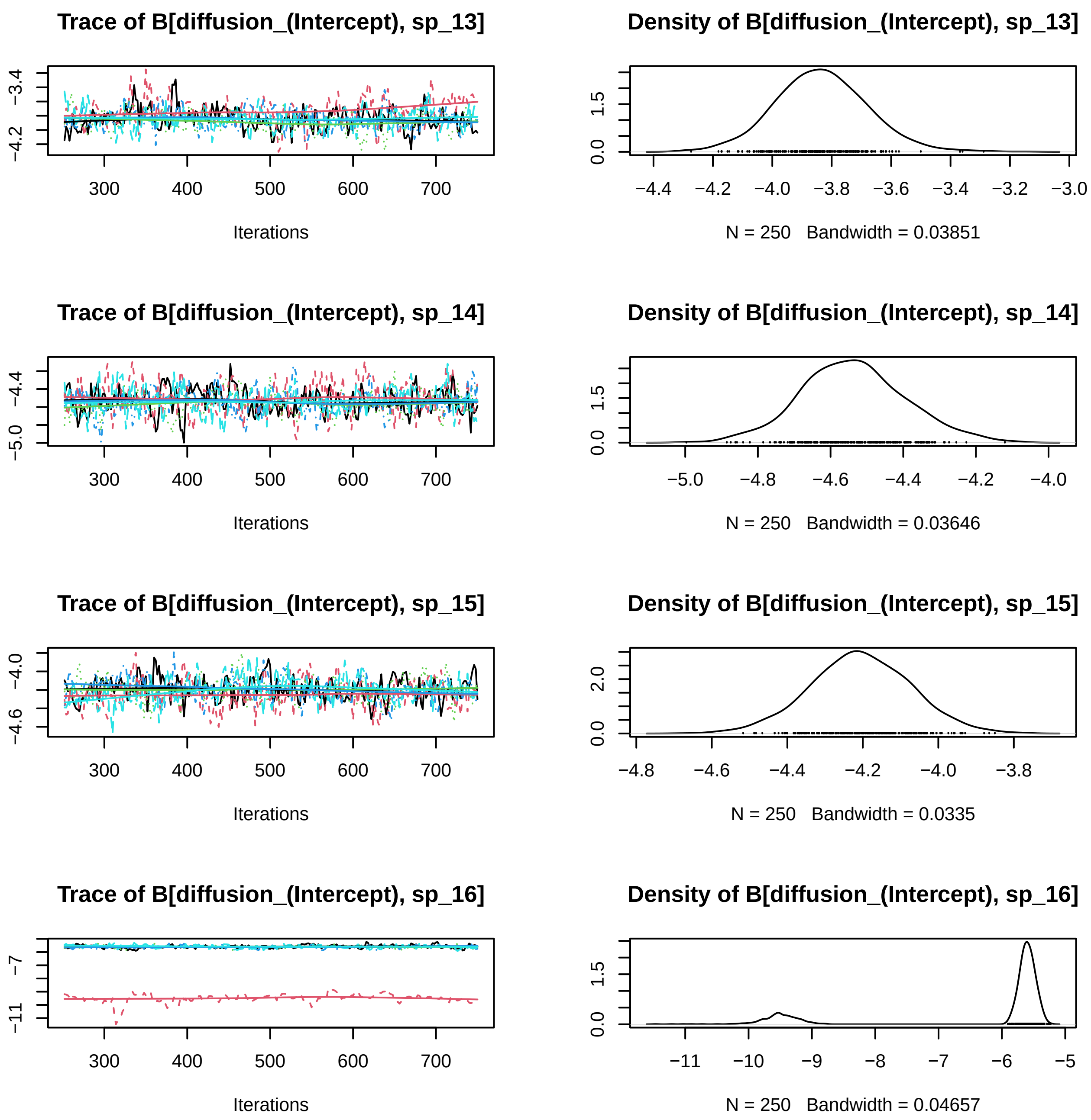
Trace of B[diffusion_(Intercept), sp_13]
Iterations
Density of B[diffusion_(Intercept), sp_13]
N = 250   Bandwidth = 0.03851
Trace of B[diffusion_(Intercept), sp_14]
Iterations
Density of B[diffusion_(Intercept), sp_14]
N = 250   Bandwidth = 0.03646
Trace of B[diffusion_(Intercept), sp_15]
Iterations
Density of B[diffusion_(Intercept), sp_15]
N = 250   Bandwidth = 0.0335
Trace of B[diffusion_(Intercept), sp_16]
Iterations
Density of B[diffusion_(Intercept), sp_16]
N = 250   Bandwidth = 0.04657

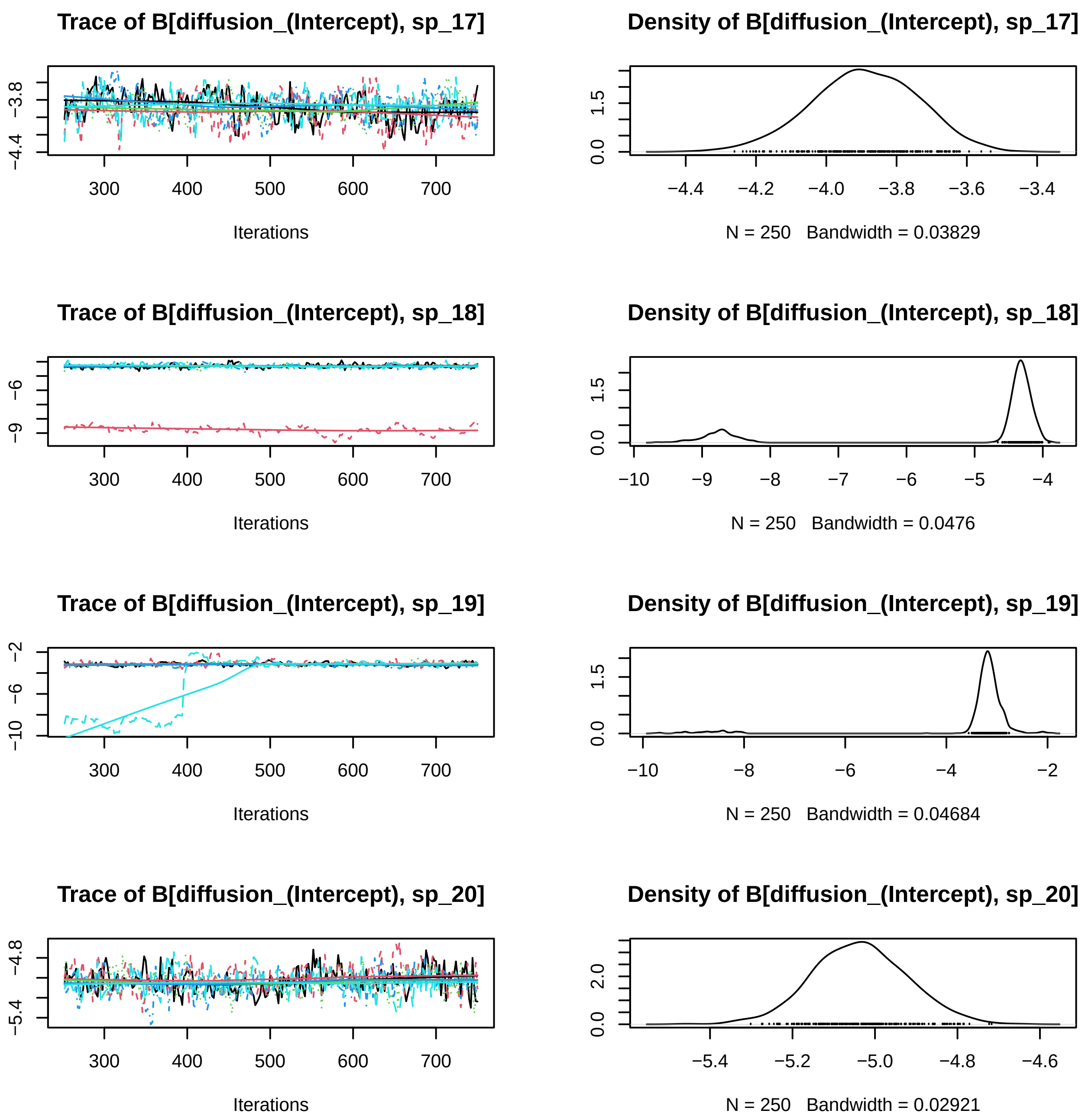
Trace of B[diffusion_(Intercept), sp_17]
Iterations
Density of B[diffusion_(Intercept), sp_17]
N = 250 Bandwidth = 0.03829
Trace of B[diffusion_(Intercept), sp_18]
Iterations
Density of B[diffusion_(Intercept), sp_18]
N = 250 Bandwidth = 0.0476
Trace of B[diffusion_(Intercept), sp_19]
Iterations
Density of B[diffusion_(Intercept), sp_19]
N = 250 Bandwidth = 0.04684
Trace of B[diffusion_(Intercept), sp_20]
Iterations
Density of B[diffusion_(Intercept), sp_20]
N = 250 Bandwidth = 0.02921

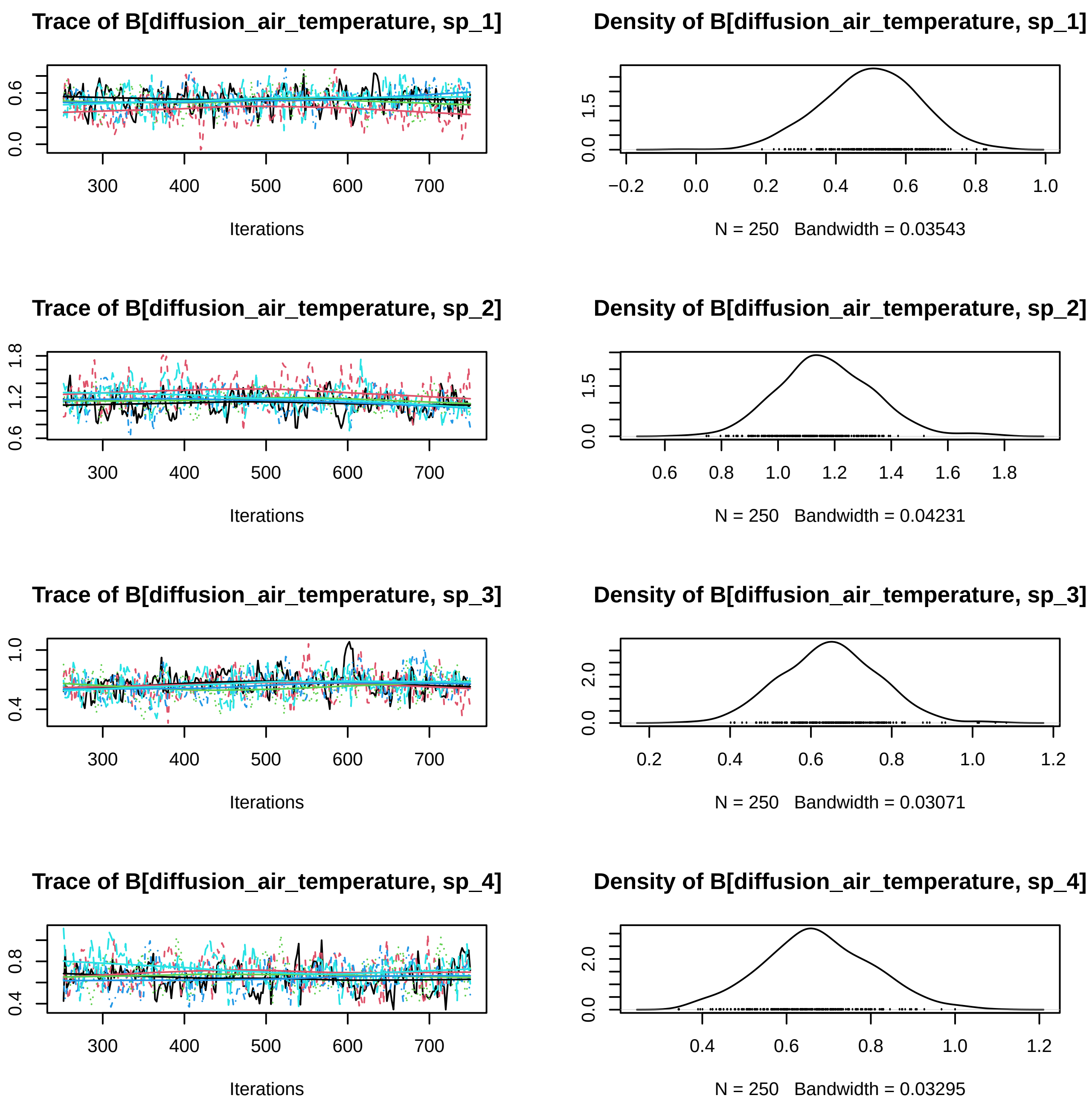
Trace of B[diffusion_air_temperature, sp_1]
Density of B[diffusion_air_temperature, sp_1]
Iterations
N = 250 Bandwidth = 0.03543
Trace of B[diffusion_air_temperature, sp_2]
Density of B[diffusion_air_temperature, sp_2]
Iterations
N = 250 Bandwidth = 0.04231
Trace of B[diffusion_air_temperature, sp_3]
Density of B[diffusion_air_temperature, sp_3]
Iterations
N = 250 Bandwidth = 0.03071
Trace of B[diffusion_air_temperature, sp_4]
Density of B[diffusion_air_temperature, sp_4]
Iterations
N = 250 Bandwidth = 0.03295

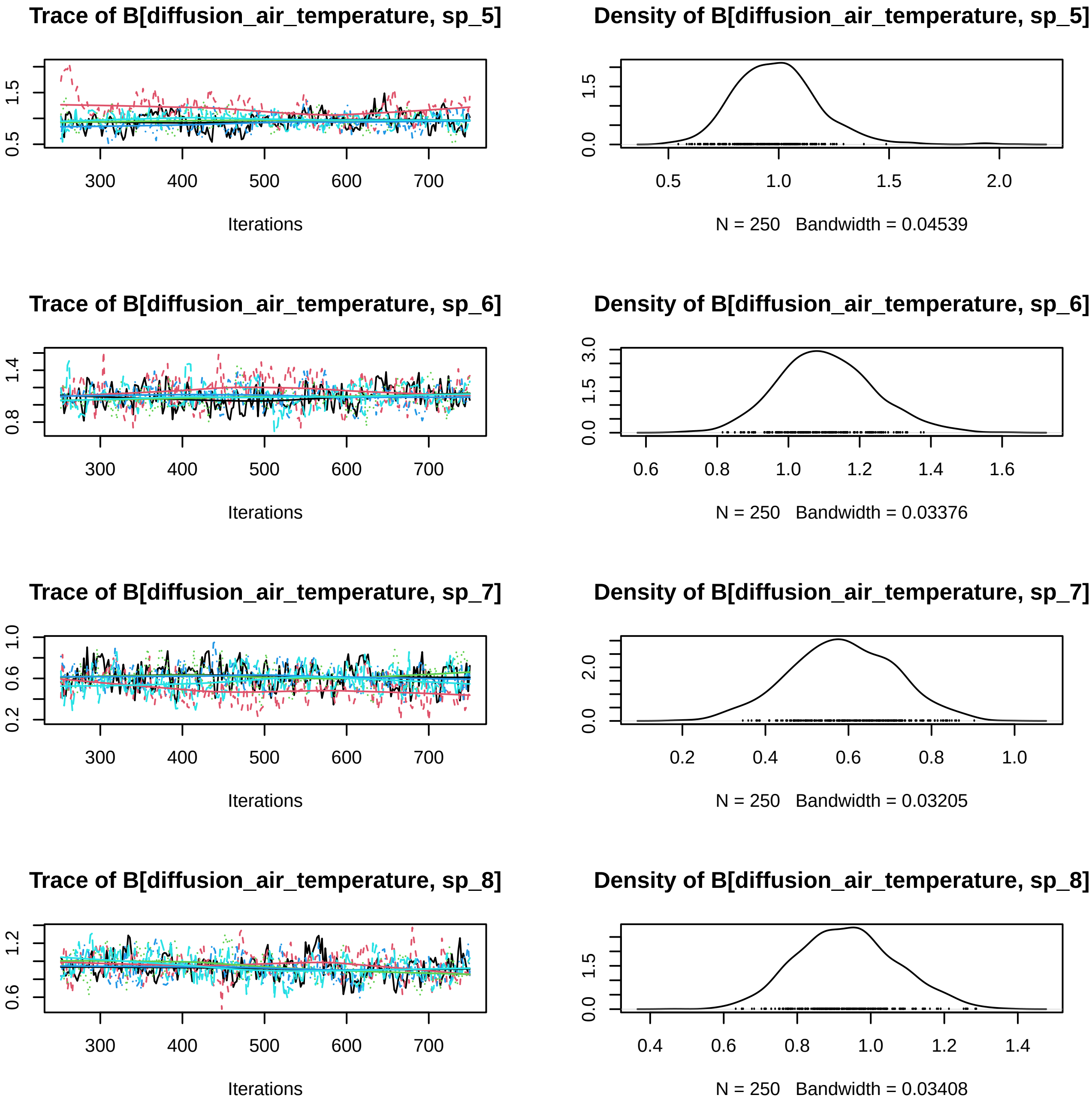
Trace of B[diffusion_air_temperature, sp_5]
Iterations
Density of B[diffusion_air_temperature, sp_5]
N = 250   Bandwidth = 0.04539
Trace of B[diffusion_air_temperature, sp_6]
Iterations
Density of B[diffusion_air_temperature, sp_6]
N = 250   Bandwidth = 0.03376
Trace of B[diffusion_air_temperature, sp_7]
Iterations
Density of B[diffusion_air_temperature, sp_7]
N = 250   Bandwidth = 0.03205
Trace of B[diffusion_air_temperature, sp_8]
Iterations
Density of B[diffusion_air_temperature, sp_8]
N = 250   Bandwidth = 0.03408

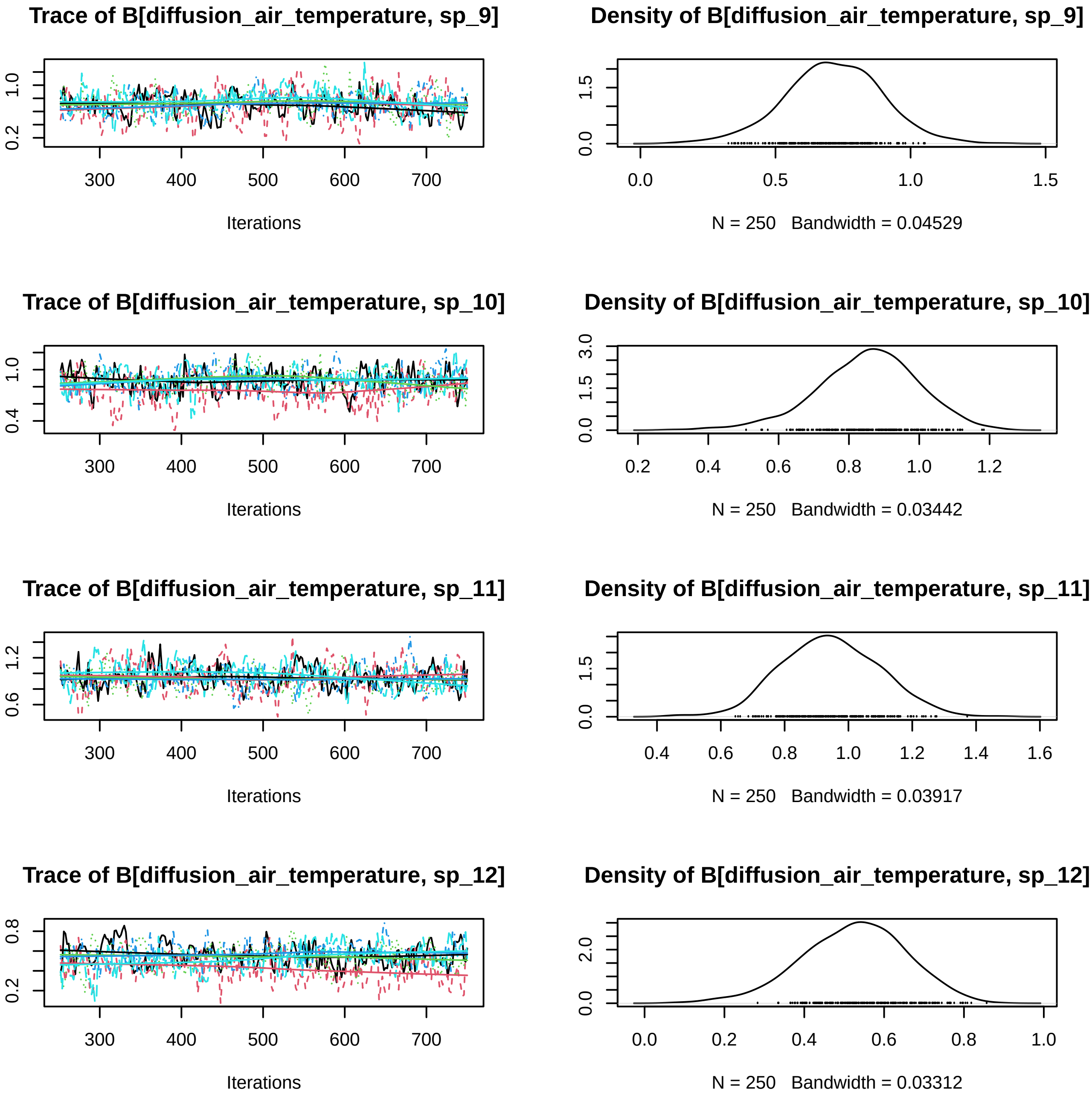
Trace of B[diffusion_air_temperature, sp_9]
Density of B[diffusion_air_temperature, sp_9]
Iterations
N = 250 Bandwidth = 0.04529
Trace of B[diffusion_air_temperature, sp_10]
Density of B[diffusion_air_temperature, sp_10]
Iterations
N = 250 Bandwidth = 0.03442
Trace of B[diffusion_air_temperature, sp_11]
Density of B[diffusion_air_temperature, sp_11]
Iterations
N = 250 Bandwidth = 0.03917
Trace of B[diffusion_air_temperature, sp_12]
Density of B[diffusion_air_temperature, sp_12]
Iterations
N = 250 Bandwidth = 0.03312

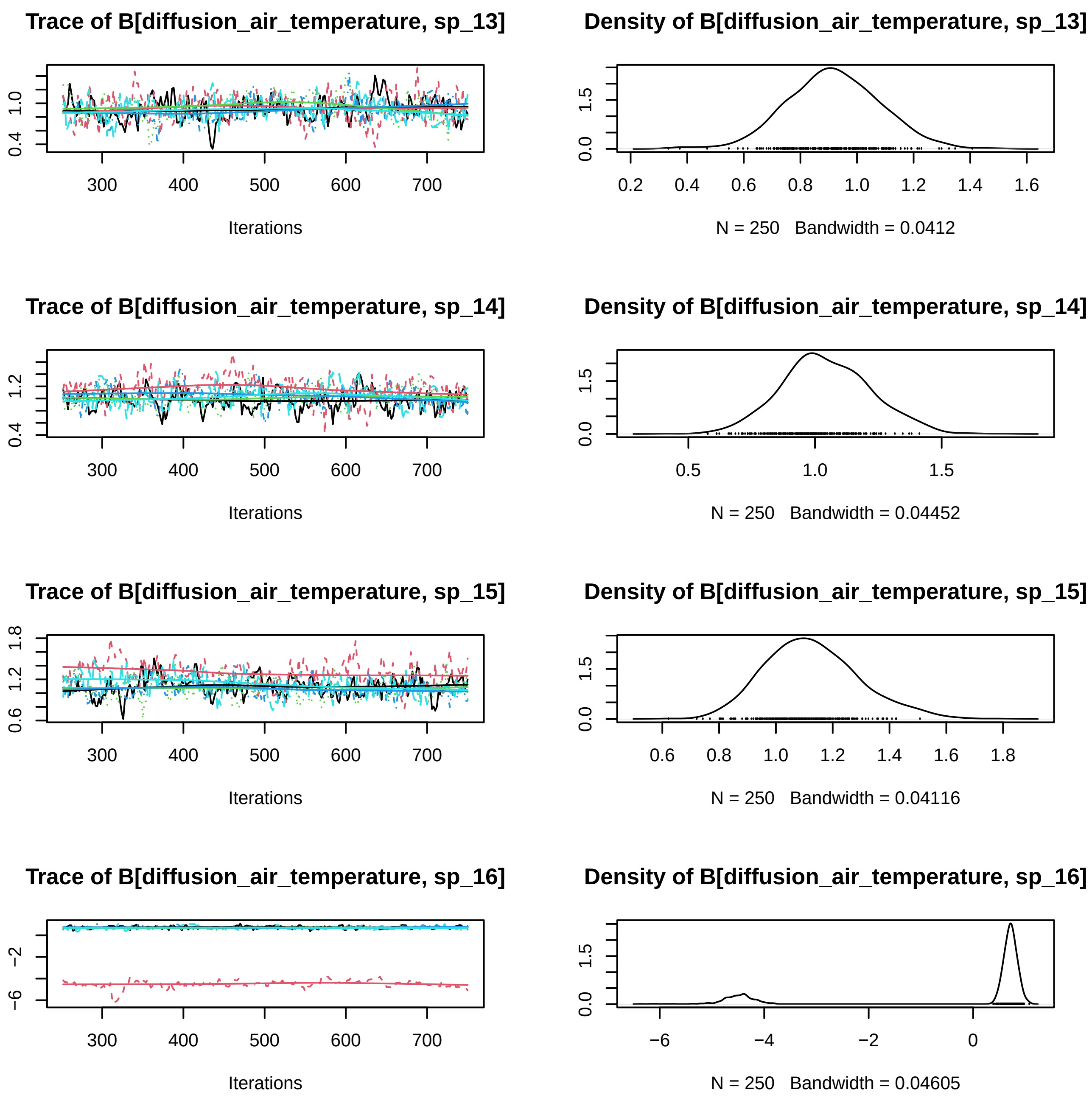

Trace of B[diffusion_air_temperature, sp_13]
0.4
1.0
300
400
500
600
700
Iterations
Density of B[diffusion_air_temperature, sp_13]
0.0
1.5
0.2
0.4
0.6
0.8
1.0
1.2
1.4
1.6
N = 250 Bandwidth = 0.0412
Trace of B[diffusion_air_temperature, sp_14]
0.4
1.2
300
400
500
600
700
Iterations
Density of B[diffusion_air_temperature, sp_14]
0.0
1.5
0.5
1.0
1.5
N = 250 Bandwidth = 0.04452
Trace of B[diffusion_air_temperature, sp_15]
0.6
1.2
1.8
300
400
500
600
700
Iterations
Density of B[diffusion_air_temperature, sp_15]
0.0
1.5
0.6
0.8
1.0
1.2
1.4
1.6
1.8
N = 250 Bandwidth = 0.04116
Trace of B[diffusion_air_temperature, sp_16]
−6
−2
300
400
500
600
700
Iterations
Density of B[diffusion_air_temperature, sp_16]
0.0
1.5
−6
−4
−2
0
N = 250 Bandwidth = 0.04605

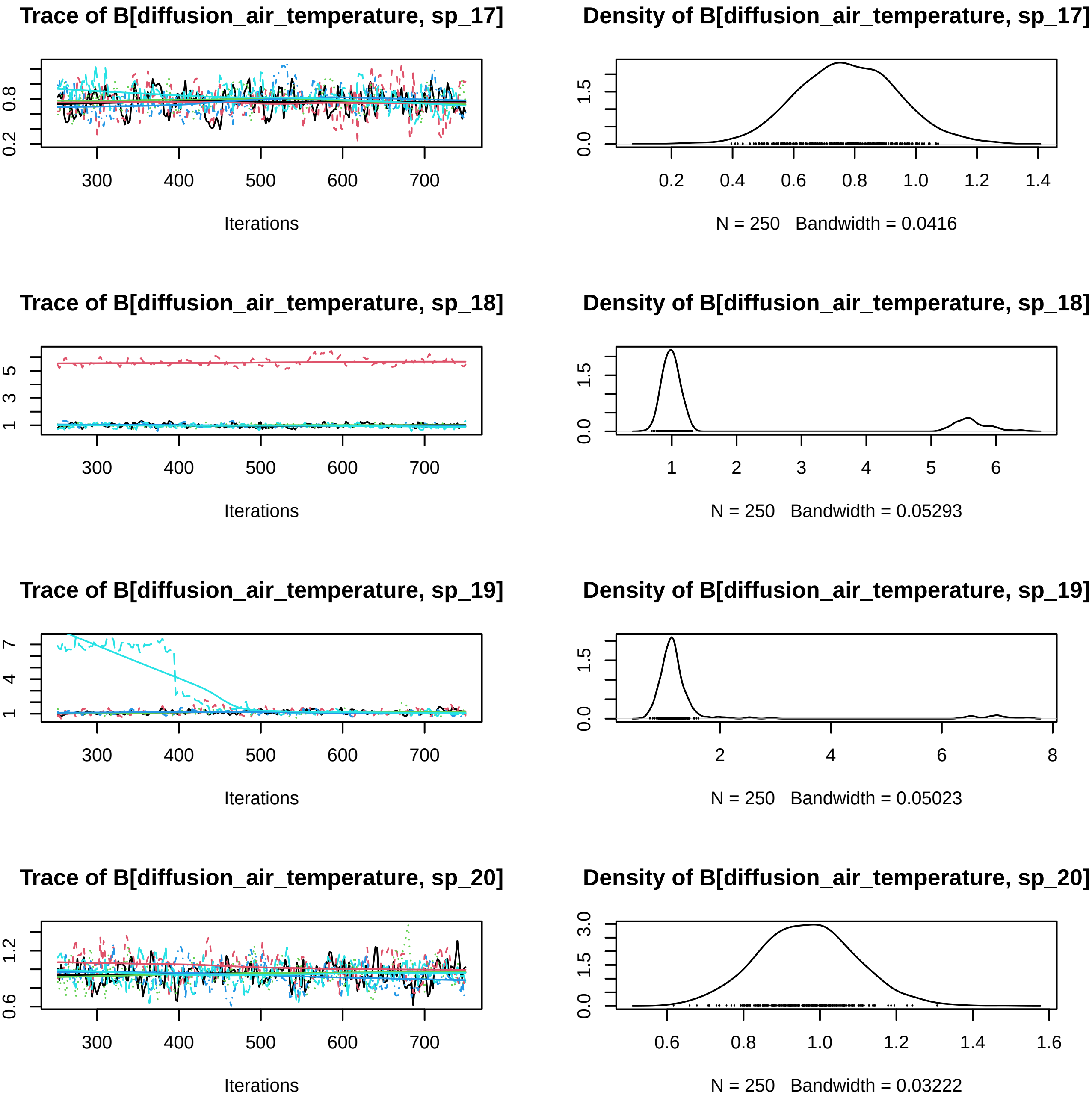
Trace of B[diffusion_air_temperature, sp_17]
Density of B[diffusion_air_temperature, sp_17]
Iterations
N = 250 Bandwidth = 0.0416
Trace of B[diffusion_air_temperature, sp_18]
Density of B[diffusion_air_temperature, sp_18]
Iterations
N = 250 Bandwidth = 0.05293
Trace of B[diffusion_air_temperature, sp_19]
Density of B[diffusion_air_temperature, sp_19]
Iterations
N = 250 Bandwidth = 0.05023
Trace of B[diffusion_air_temperature, sp_20]
Density of B[diffusion_air_temperature, sp_20]
Iterations
N = 250 Bandwidth = 0.03222

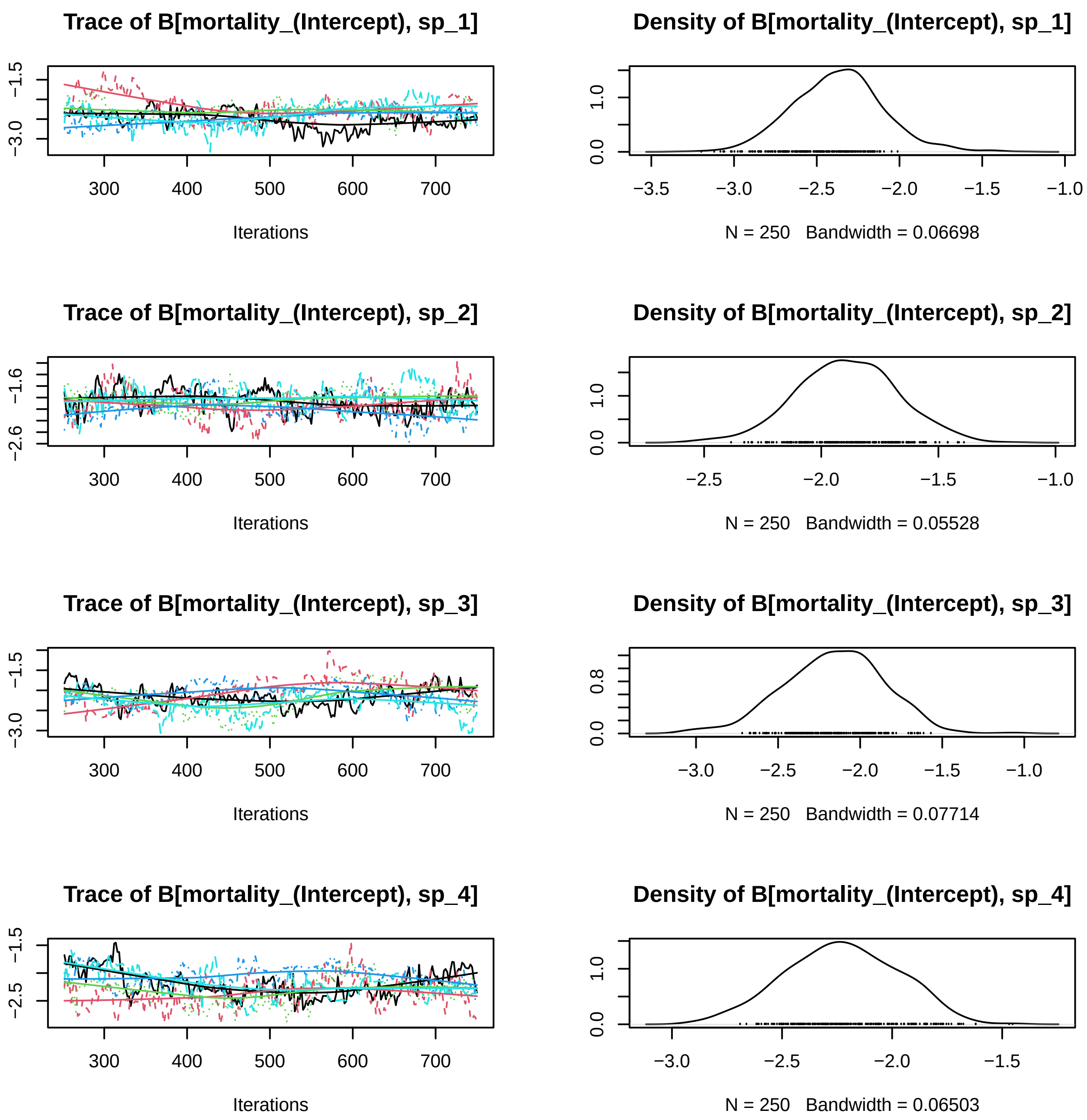

Trace of B[mortality_(Intercept), sp_1]
−1.5
−3.0
300
400
500
600
700
Iterations
Density of B[mortality_(Intercept), sp_1]
0.0
1.0
−3.5
−3.0
−2.5
−2.0
−1.5
−1.0
N = 250 Bandwidth = 0.06698
Trace of B[mortality_(Intercept), sp_2]
−1.6
−2.6
300
400
500
600
700
Iterations
Density of B[mortality_(Intercept), sp_2]
0.0
1.0
−2.5
−2.0
−1.5
−1.0
N = 250 Bandwidth = 0.05528
Trace of B[mortality_(Intercept), sp_3]
−1.5
−3.0
300
400
500
600
700
Iterations
Density of B[mortality_(Intercept), sp_3]
0.0
0.8
−3.0
−2.5
−2.0
−1.5
−1.0
N = 250 Bandwidth = 0.07714
Trace of B[mortality_(Intercept), sp_4]
−1.5
−2.5
300
400
500
600
700
Iterations
Density of B[mortality_(Intercept), sp_4]
0.0
1.0
−3.0
−2.5
−2.0
−1.5
N = 250 Bandwidth = 0.06503

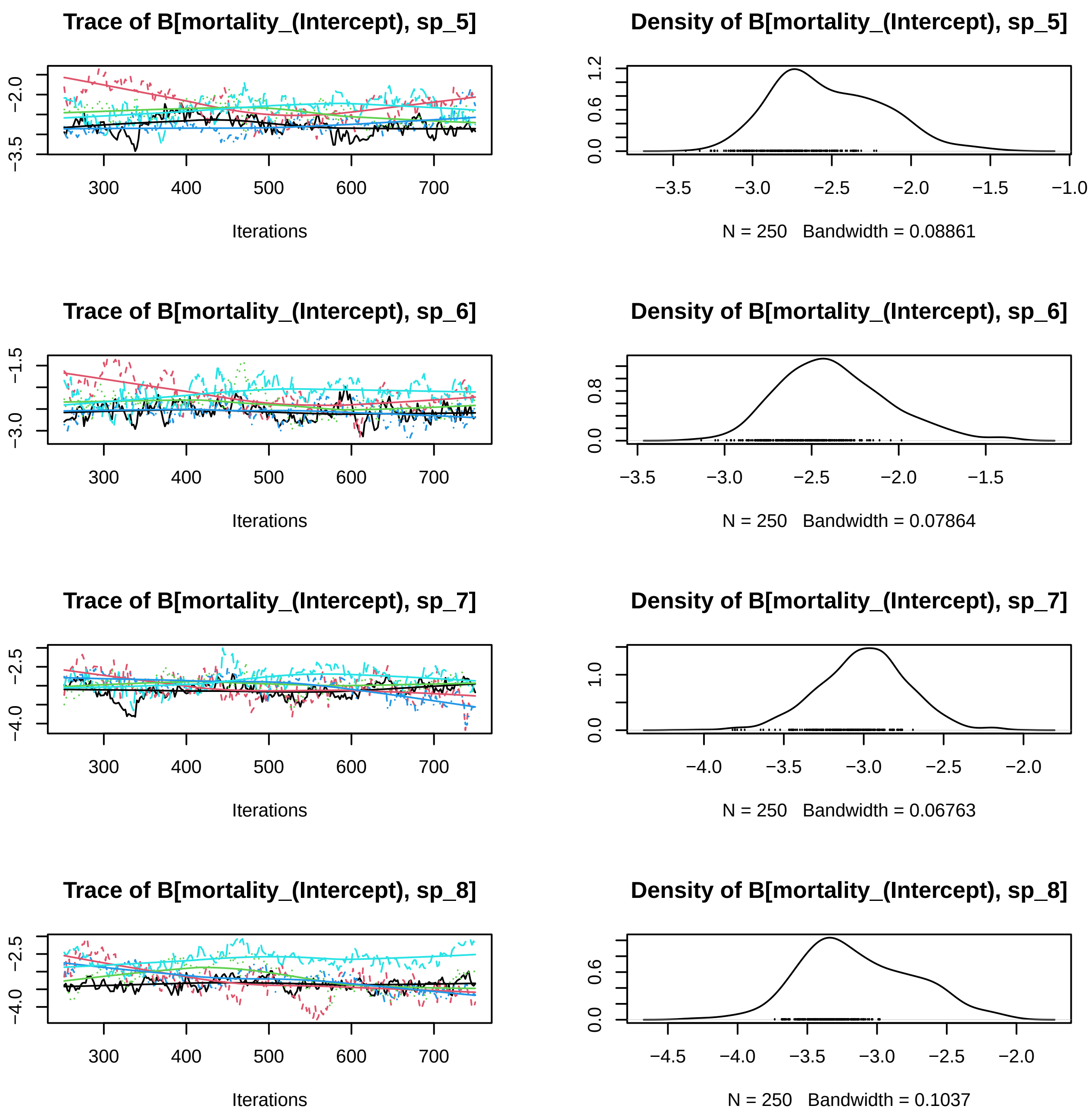

Trace of B[mortality_(Intercept), sp_5]
Iterations
Density of B[mortality_(Intercept), sp_5]
N = 250 Bandwidth = 0.08861
Trace of B[mortality_(Intercept), sp_6]
Iterations
Density of B[mortality_(Intercept), sp_6]
N = 250 Bandwidth = 0.07864
Trace of B[mortality_(Intercept), sp_7]
Iterations
Density of B[mortality_(Intercept), sp_7]
N = 250 Bandwidth = 0.06763
Trace of B[mortality_(Intercept), sp_8]
Iterations
Density of B[mortality_(Intercept), sp_8]
N = 250 Bandwidth = 0.1037

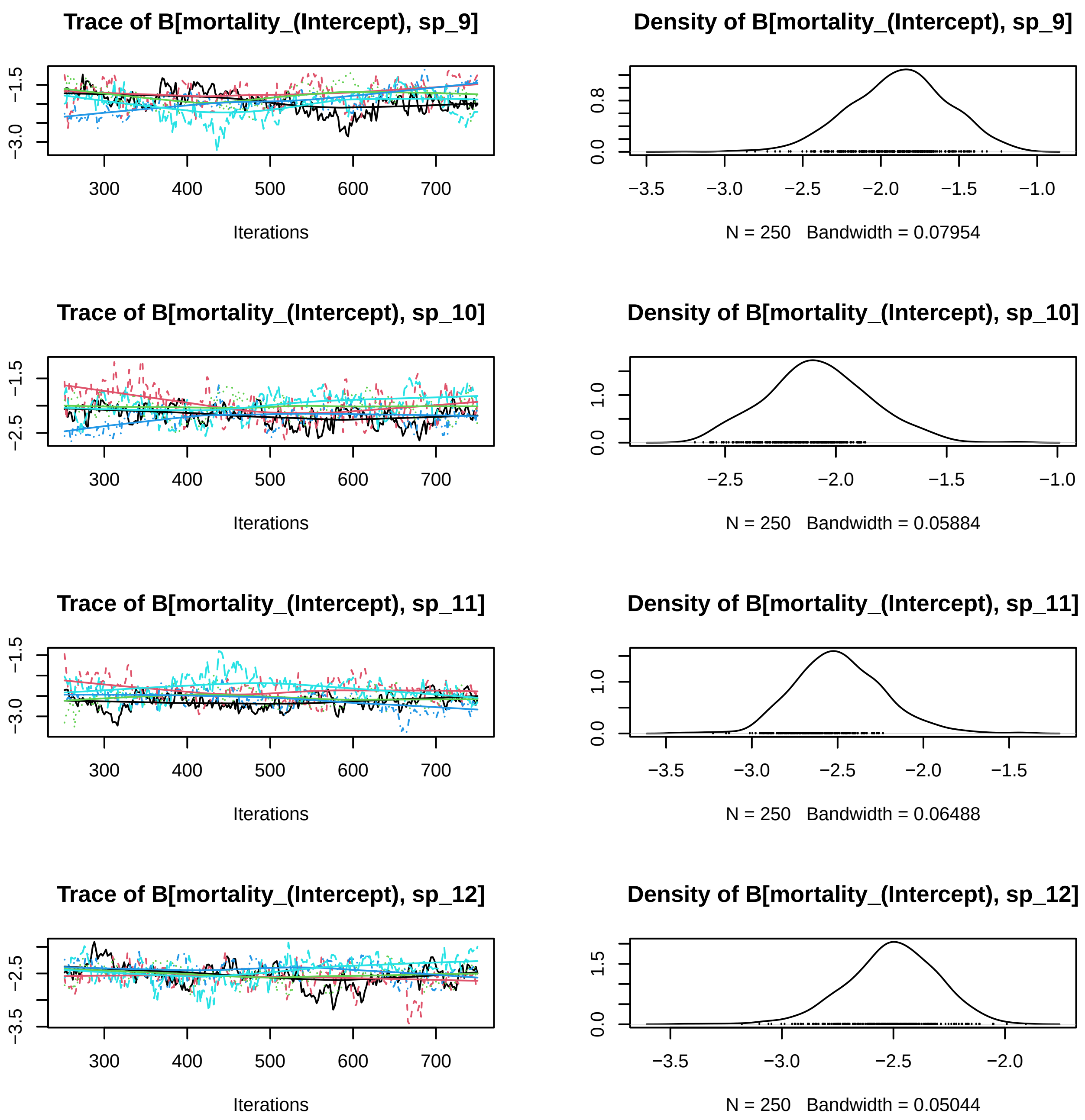
Trace of B[mortality_(Intercept), sp_9]
−3.0
−1.5
300
400
500
600
700
Iterations
Density of B[mortality_(Intercept), sp_9]
0.0
0.8
−3.5
−3.0
−2.5
−2.0
−1.5
−1.0
N = 250 Bandwidth = 0.07954
Trace of B[mortality_(Intercept), sp_10]
−2.5
−1.5
300
400
500
600
700
Iterations
Density of B[mortality_(Intercept), sp_10]
0.0
1.0
−2.5
−2.0
−1.5
−1.0
N = 250 Bandwidth = 0.05884
Trace of B[mortality_(Intercept), sp_11]
−3.0
−1.5
300
400
500
600
700
Iterations
Density of B[mortality_(Intercept), sp_11]
0.0
1.0
−3.5
−3.0
−2.5
−2.0
−1.5
N = 250 Bandwidth = 0.06488
Trace of B[mortality_(Intercept), sp_12]
−3.5
−2.5
300
400
500
600
700
Iterations
Density of B[mortality_(Intercept), sp_12]
0.0
1.5
−3.5
−3.0
−2.5
−2.0
N = 250 Bandwidth = 0.05044

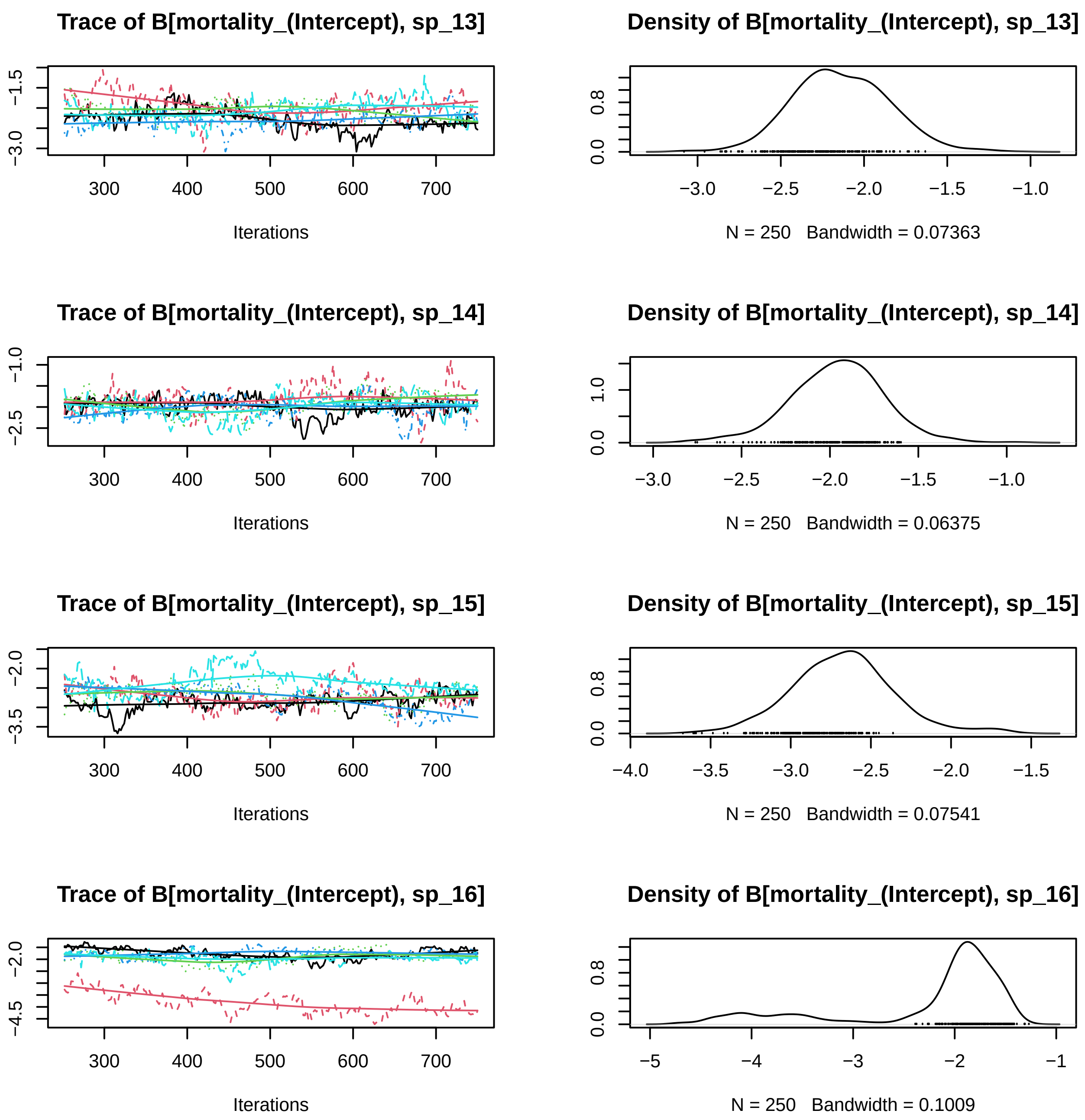
Trace of B[mortality_(Intercept), sp_13]
Density of B[mortality_(Intercept), sp_13]
−3.0
−1.5
300
400
500
600
700
Iterations
0.0
0.8
−3.0
−2.5
−2.0
−1.5
−1.0
N = 250 Bandwidth = 0.07363
Trace of B[mortality_(Intercept), sp_14]
Density of B[mortality_(Intercept), sp_14]
−2.5
−1.0
300
400
500
600
700
Iterations
0.0
1.0
−3.0
−2.5
−2.0
−1.5
−1.0
N = 250 Bandwidth = 0.06375
Trace of B[mortality_(Intercept), sp_15]
Density of B[mortality_(Intercept), sp_15]
−3.5
−2.0
300
400
500
600
700
Iterations
0.0
0.8
−4.0
−3.5
−3.0
−2.5
−2.0
−1.5
N = 250 Bandwidth = 0.07541
Trace of B[mortality_(Intercept), sp_16]
Density of B[mortality_(Intercept), sp_16]
−4.5
−2.0
300
400
500
600
700
Iterations
0.0
0.8
−5
−4
−3
−2
−1
N = 250 Bandwidth = 0.1009

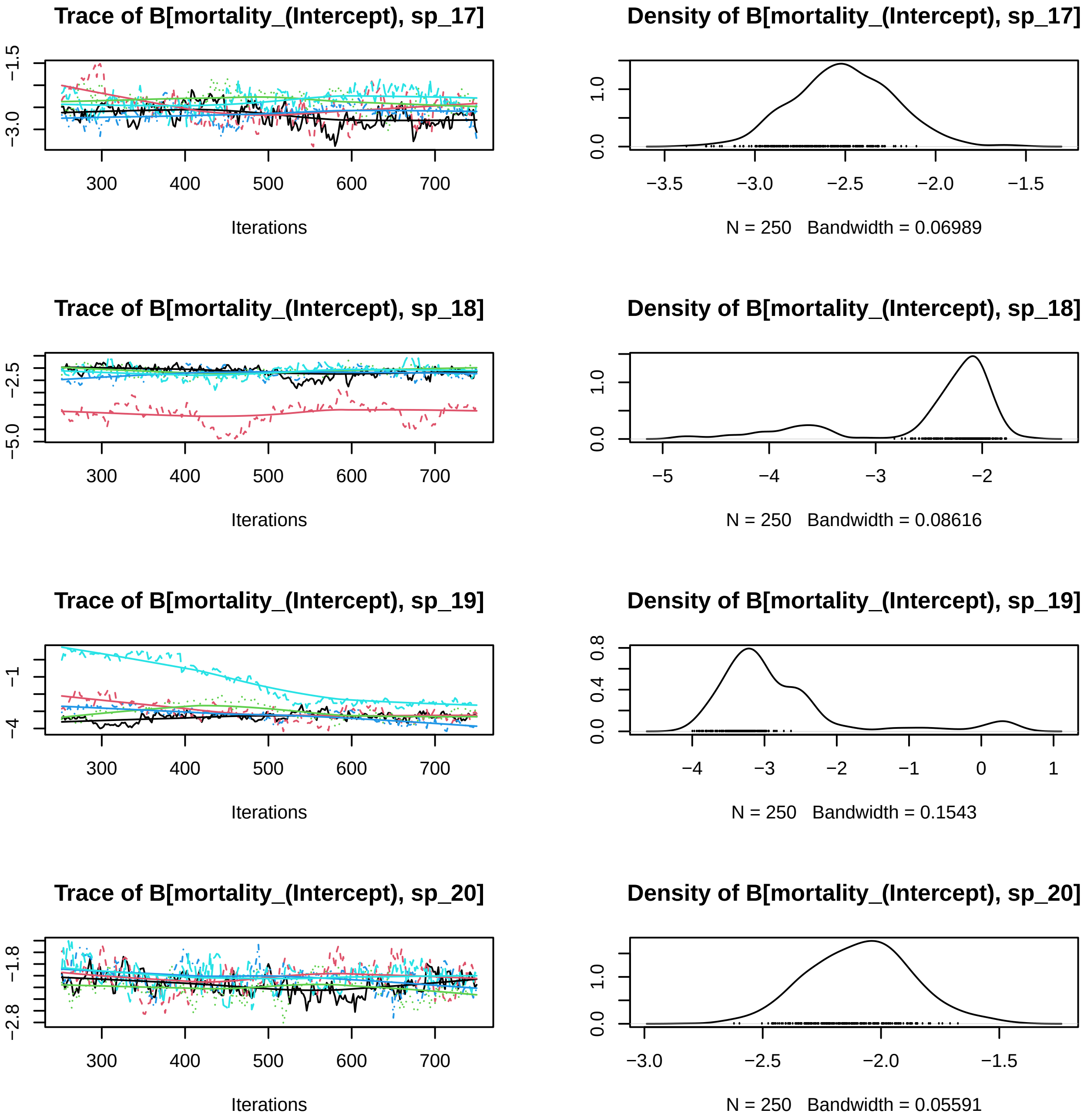

Trace of B[mortality_(Intercept), sp_17]
−1.5
−3.0
300
400
500
600
700
Iterations
Density of B[mortality_(Intercept), sp_17]
0.0
1.0
−3.5
−3.0
−2.5
−2.0
−1.5
N = 250 Bandwidth = 0.06989
Trace of B[mortality_(Intercept), sp_18]
−2.5
−5.0
300
400
500
600
700
Iterations
Density of B[mortality_(Intercept), sp_18]
0.0
1.0
−5
−4
−3
−2
N = 250 Bandwidth = 0.08616
Trace of B[mortality_(Intercept), sp_19]
−1
−4
300
400
500
600
700
Iterations
Density of B[mortality_(Intercept), sp_19]
0.0
0.4
0.8
−4
−3
−2
−1
0
1
N = 250 Bandwidth = 0.1543
Trace of B[mortality_(Intercept), sp_20]
−1.8
−2.8
300
400
500
600
700
Iterations
Density of B[mortality_(Intercept), sp_20]
0.0
1.0
−3.0
−2.5
−2.0
−1.5
N = 250 Bandwidth = 0.05591

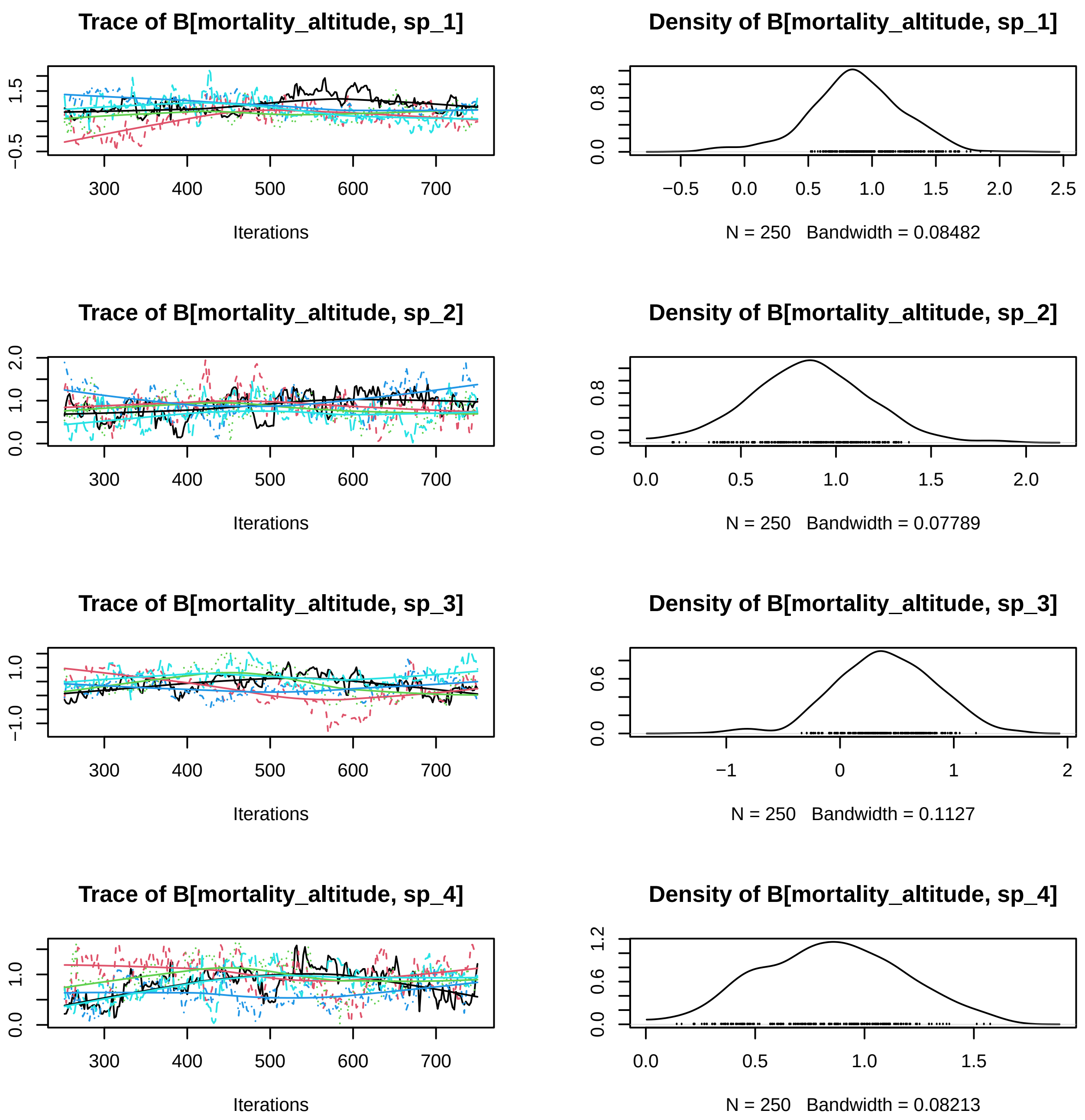
Trace of B[mortality_altitude, sp_1]
Density of B[mortality_altitude, sp_1]
Iterations
N = 250 Bandwidth = 0.08482
Trace of B[mortality_altitude, sp_2]
Density of B[mortality_altitude, sp_2]
Iterations
N = 250 Bandwidth = 0.07789
Trace of B[mortality_altitude, sp_3]
Density of B[mortality_altitude, sp_3]
Iterations
N = 250 Bandwidth = 0.1127
Trace of B[mortality_altitude, sp_4]
Density of B[mortality_altitude, sp_4]
Iterations
N = 250 Bandwidth = 0.08213

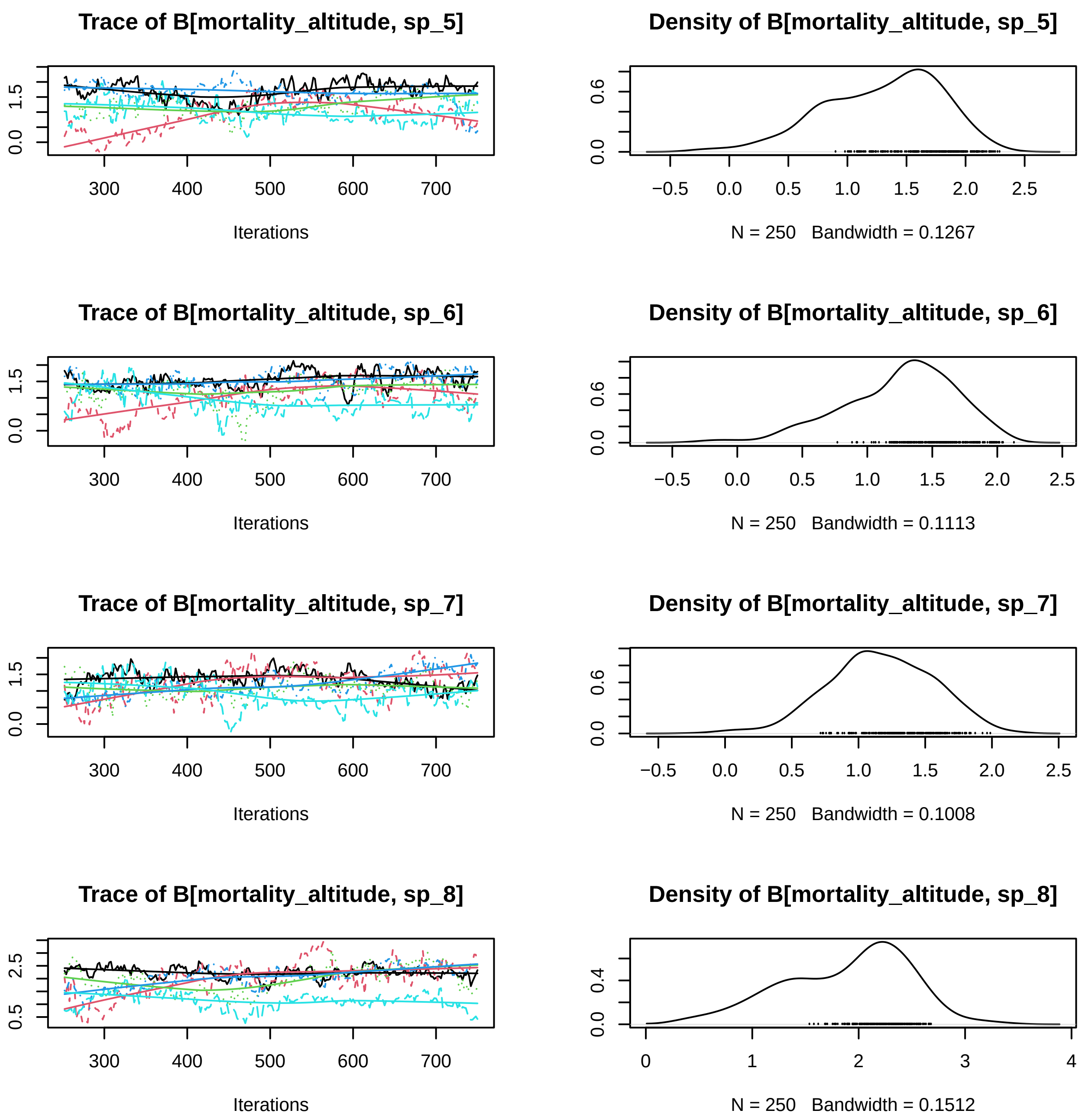
Trace of B[mortality_altitude, sp_5]
Iterations
Density of B[mortality_altitude, sp_5]
N = 250 Bandwidth = 0.1267
Trace of B[mortality_altitude, sp_6]
Iterations
Density of B[mortality_altitude, sp_6]
N = 250 Bandwidth = 0.1113
Trace of B[mortality_altitude, sp_7]
Iterations
Density of B[mortality_altitude, sp_7]
N = 250 Bandwidth = 0.1008
Trace of B[mortality_altitude, sp_8]
Iterations
Density of B[mortality_altitude, sp_8]
N = 250 Bandwidth = 0.1512

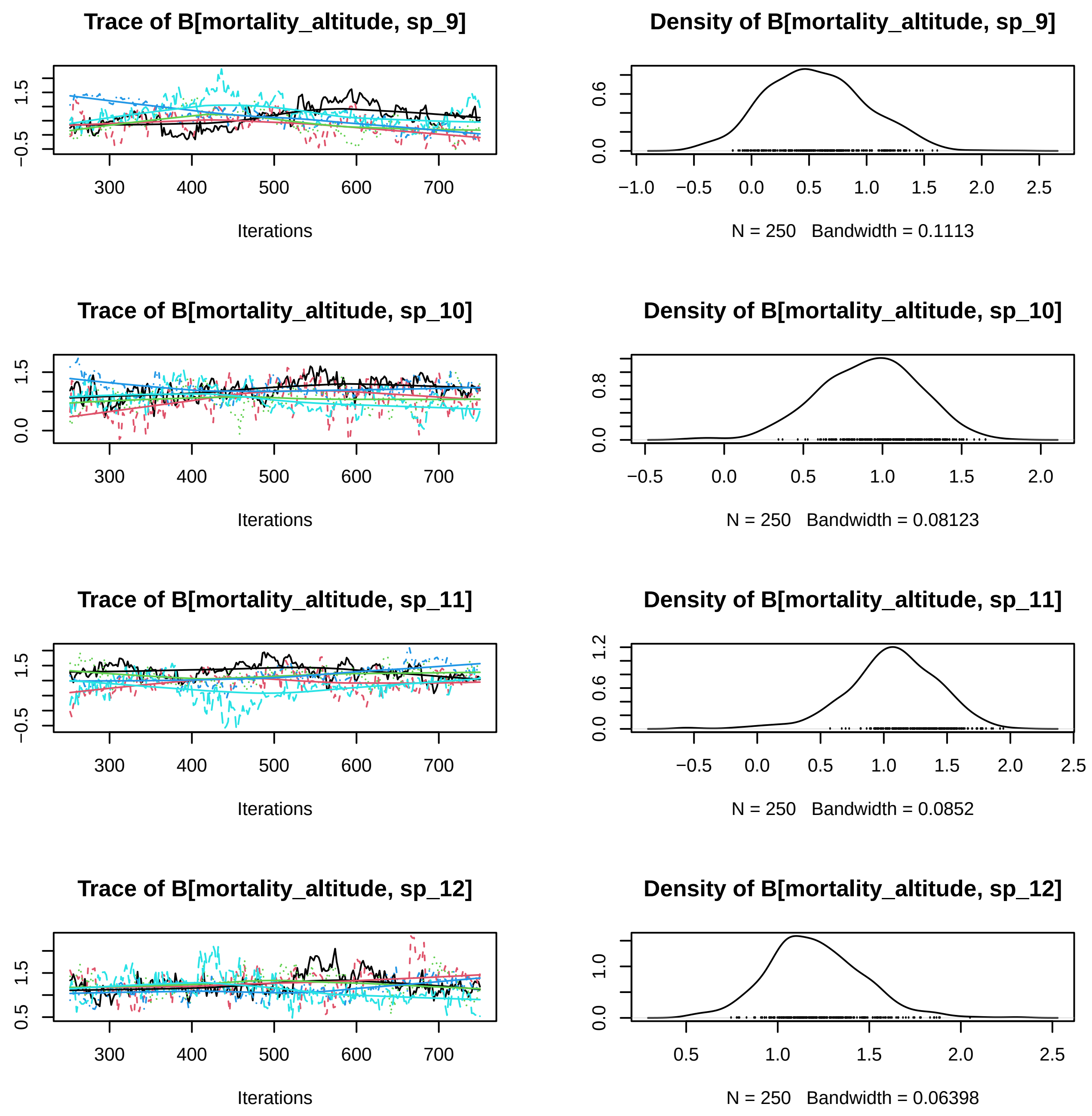

Trace of B[mortality_altitude, sp_9]
Iterations
Density of B[mortality_altitude, sp_9]
N = 250 Bandwidth = 0.1113
Trace of B[mortality_altitude, sp_10]
Iterations
Density of B[mortality_altitude, sp_10]
N = 250 Bandwidth = 0.08123
Trace of B[mortality_altitude, sp_11]
Iterations
Density of B[mortality_altitude, sp_11]
N = 250 Bandwidth = 0.0852
Trace of B[mortality_altitude, sp_12]
Iterations
Density of B[mortality_altitude, sp_12]
N = 250 Bandwidth = 0.06398

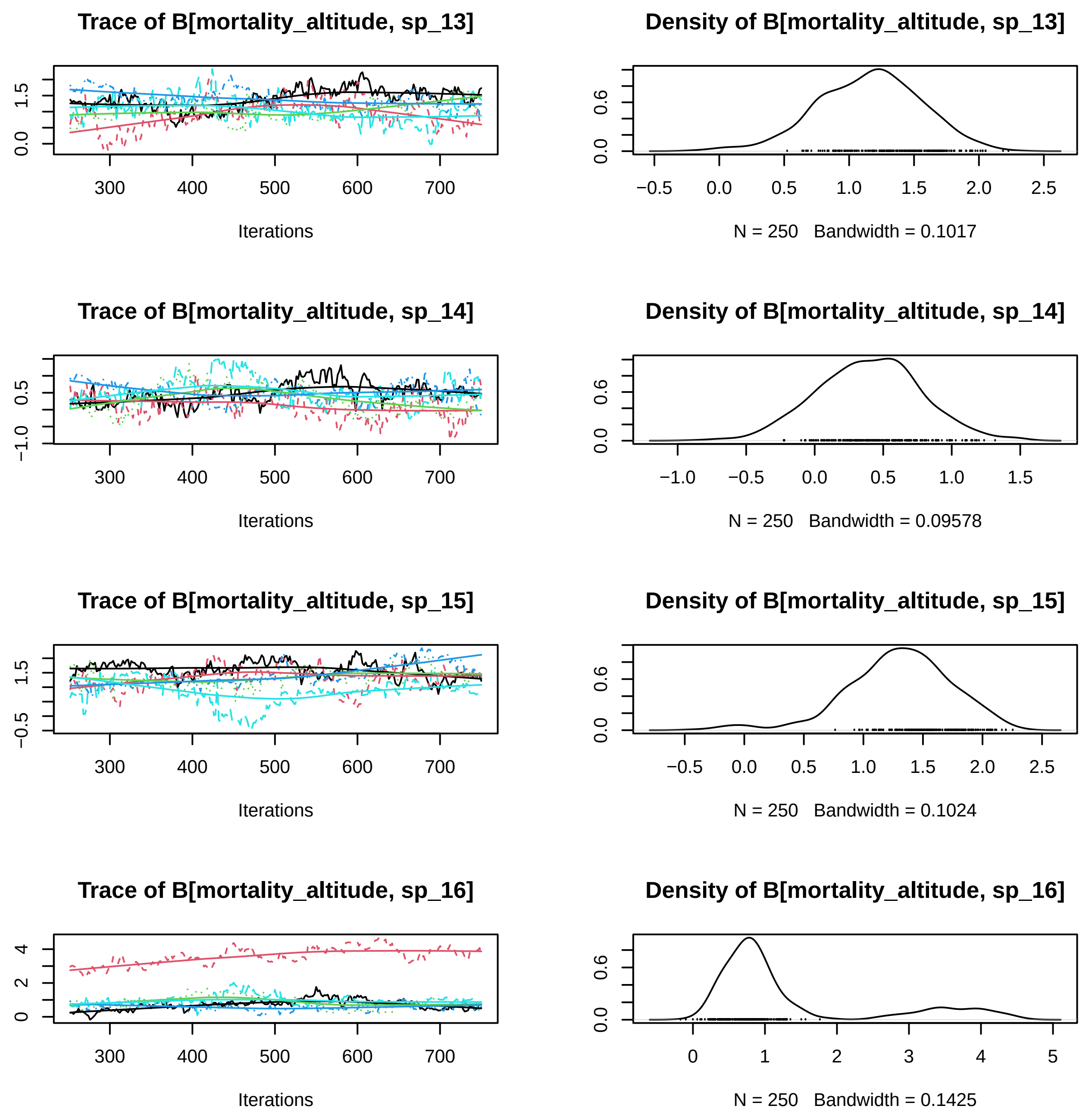
Trace of B[mortality_altitude, sp_13]
Iterations
Density of B[mortality_altitude, sp_13]
N = 250 Bandwidth = 0.1017
Trace of B[mortality_altitude, sp_14]
Iterations
Density of B[mortality_altitude, sp_14]
N = 250 Bandwidth = 0.09578
Trace of B[mortality_altitude, sp_15]
Iterations
Density of B[mortality_altitude, sp_15]
N = 250 Bandwidth = 0.1024
Trace of B[mortality_altitude, sp_16]
Iterations
Density of B[mortality_altitude, sp_16]
N = 250 Bandwidth = 0.1425

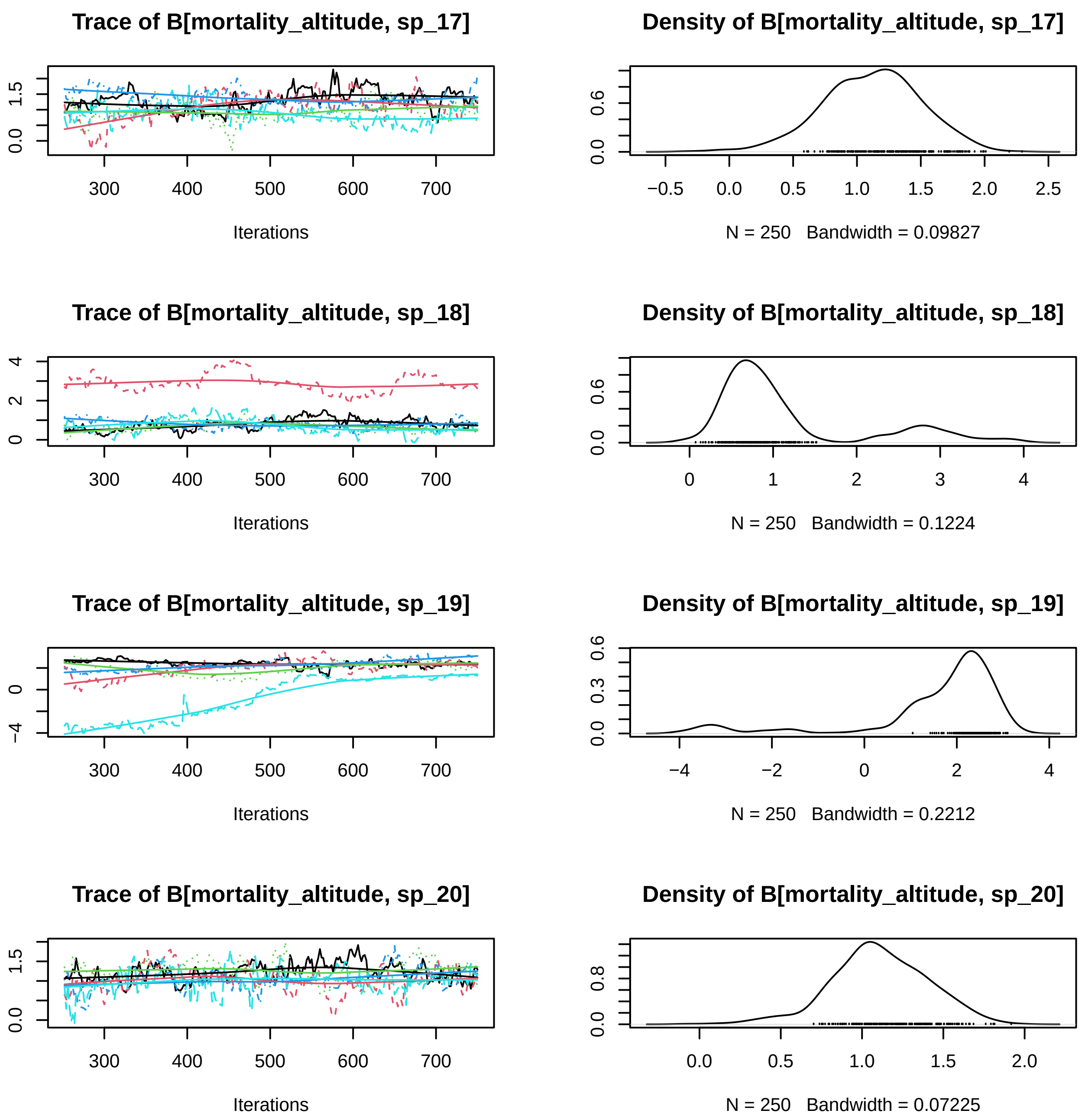
Trace of B[mortality_altitude, sp_17]
Iterations
Density of B[mortality_altitude, sp_17]
N = 250 Bandwidth = 0.09827
Trace of B[mortality_altitude, sp_18]
Iterations
Density of B[mortality_altitude, sp_18]
N = 250 Bandwidth = 0.1224
Trace of B[mortality_altitude, sp_19]
Iterations
Density of B[mortality_altitude, sp_19]
N = 250 Bandwidth = 0.2212
Trace of B[mortality_altitude, sp_20]
Iterations
Density of B[mortality_altitude, sp_20]
N = 250 Bandwidth = 0.07225

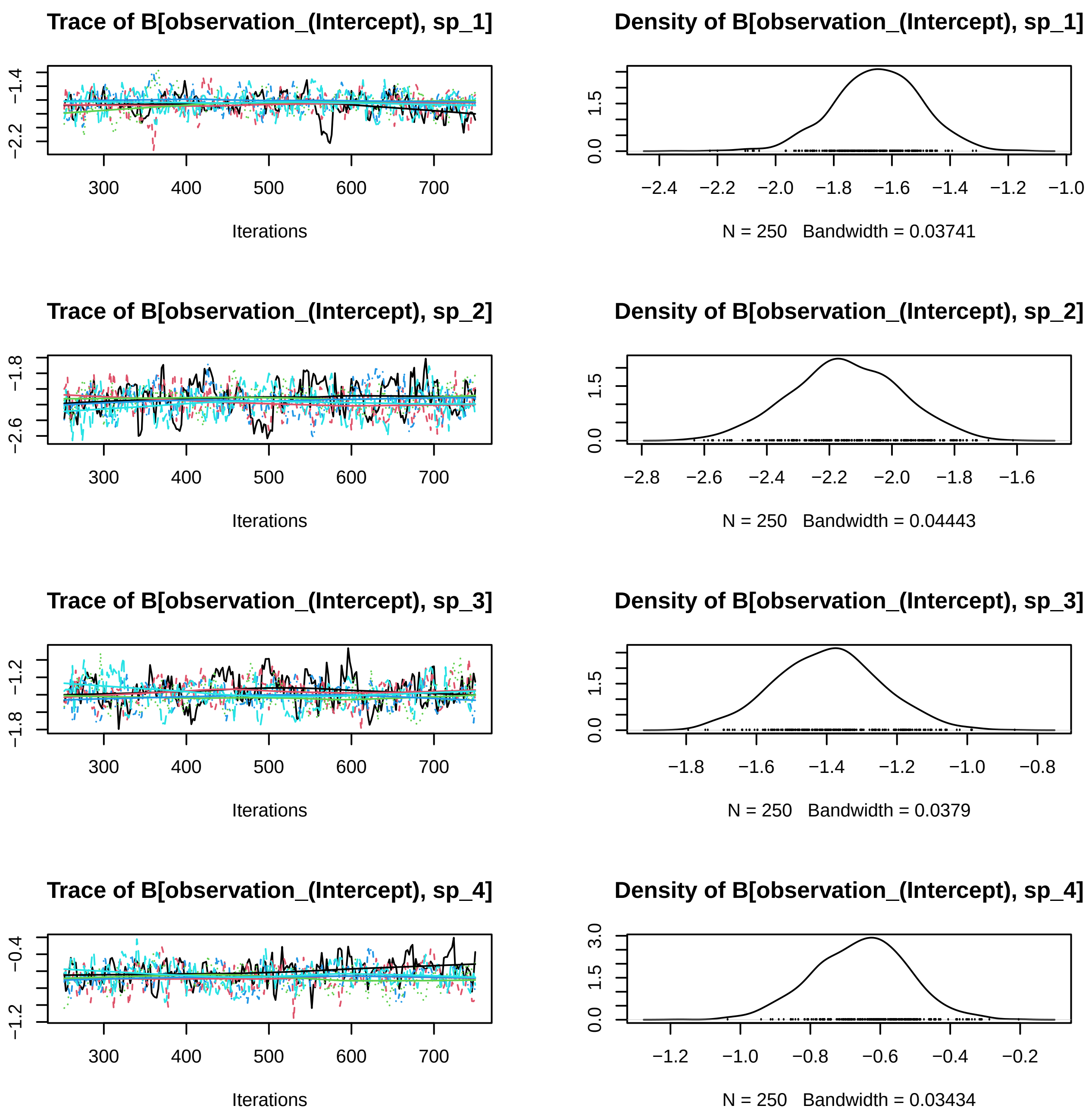
Trace of B[observation_(Intercept), sp_1]
Iterations
Density of B[observation_(Intercept), sp_1]
N = 250   Bandwidth = 0.03741
Trace of B[observation_(Intercept), sp_2]
Iterations
Density of B[observation_(Intercept), sp_2]
N = 250   Bandwidth = 0.04443
Trace of B[observation_(Intercept), sp_3]
Iterations
Density of B[observation_(Intercept), sp_3]
N = 250   Bandwidth = 0.0379
Trace of B[observation_(Intercept), sp_4]
Iterations
Density of B[observation_(Intercept), sp_4]
N = 250   Bandwidth = 0.03434

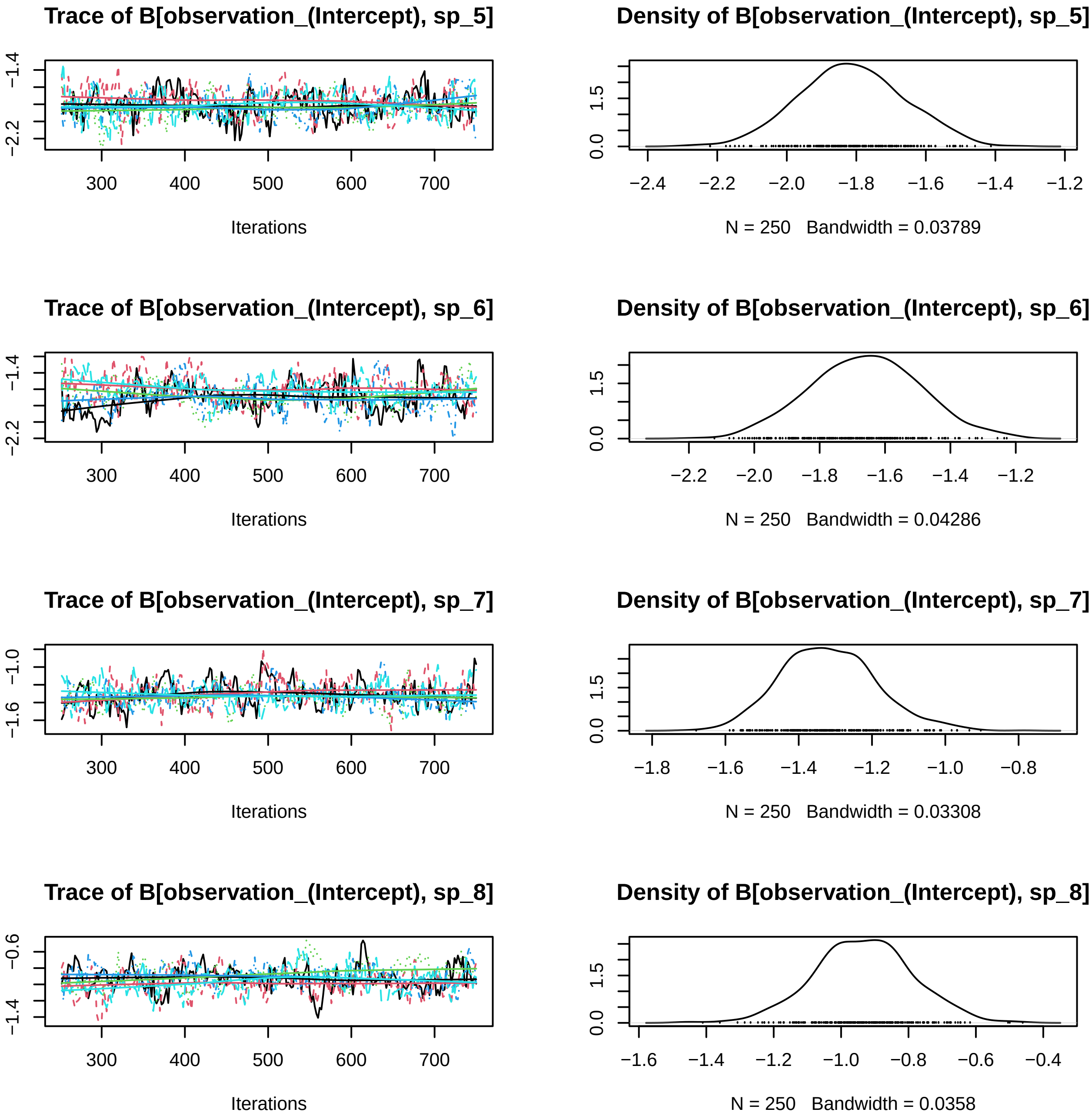
Trace of B[observation_(Intercept), sp_5]
Iterations
Density of B[observation_(Intercept), sp_5]
N = 250 Bandwidth = 0.03789
Trace of B[observation_(Intercept), sp_6]
Iterations
Density of B[observation_(Intercept), sp_6]
N = 250 Bandwidth = 0.04286
Trace of B[observation_(Intercept), sp_7]
Iterations
Density of B[observation_(Intercept), sp_7]
N = 250 Bandwidth = 0.03308
Trace of B[observation_(Intercept), sp_8]
Iterations
Density of B[observation_(Intercept), sp_8]
N = 250 Bandwidth = 0.0358

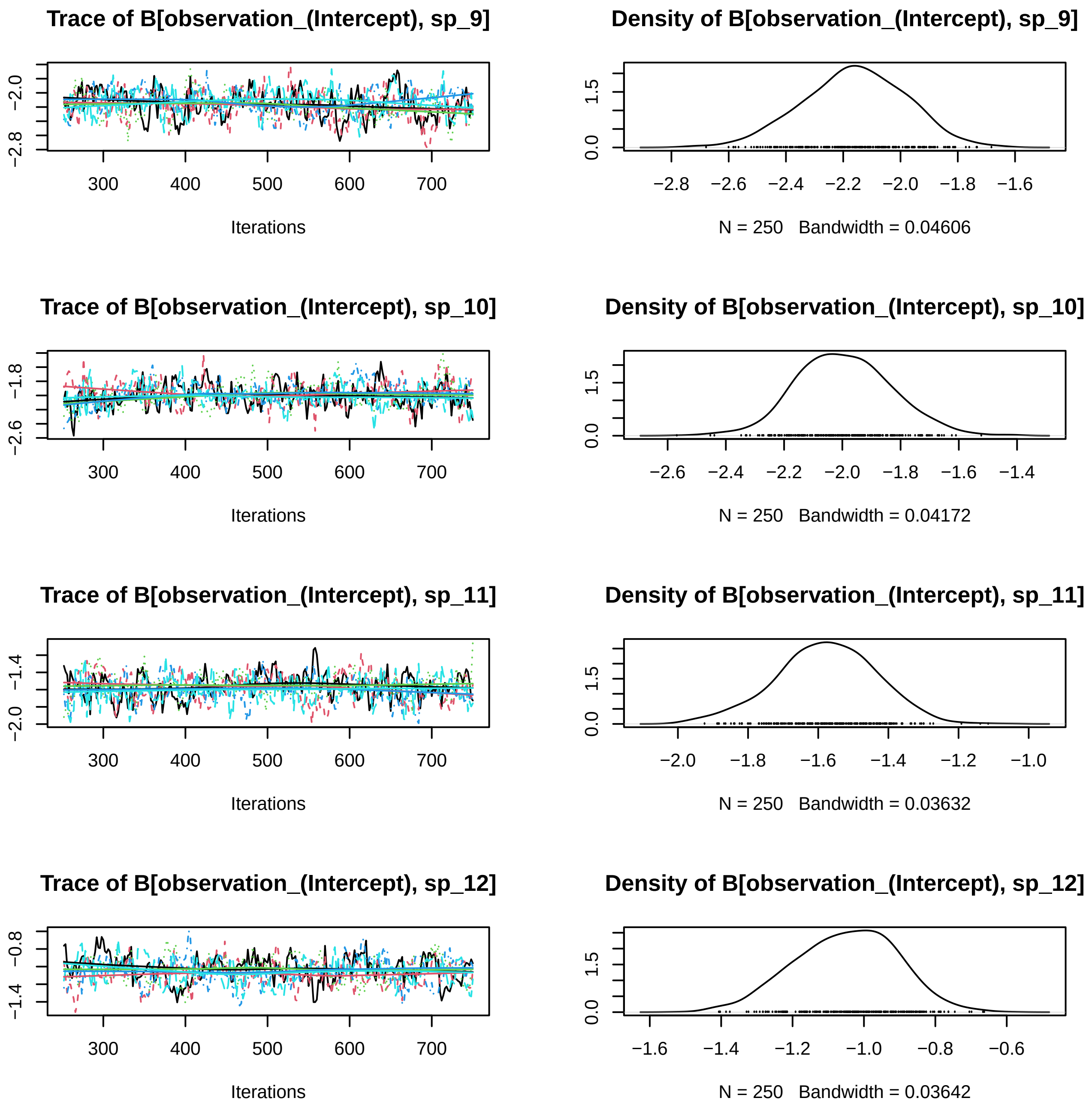
Trace of B[observation_(Intercept), sp_9]
Iterations
Density of B[observation_(Intercept), sp_9]
N = 250   Bandwidth = 0.04606
Trace of B[observation_(Intercept), sp_10]
Iterations
Density of B[observation_(Intercept), sp_10]
N = 250   Bandwidth = 0.04172
Trace of B[observation_(Intercept), sp_11]
Iterations
Density of B[observation_(Intercept), sp_11]
N = 250   Bandwidth = 0.03632
Trace of B[observation_(Intercept), sp_12]
Iterations
Density of B[observation_(Intercept), sp_12]
N = 250   Bandwidth = 0.03642

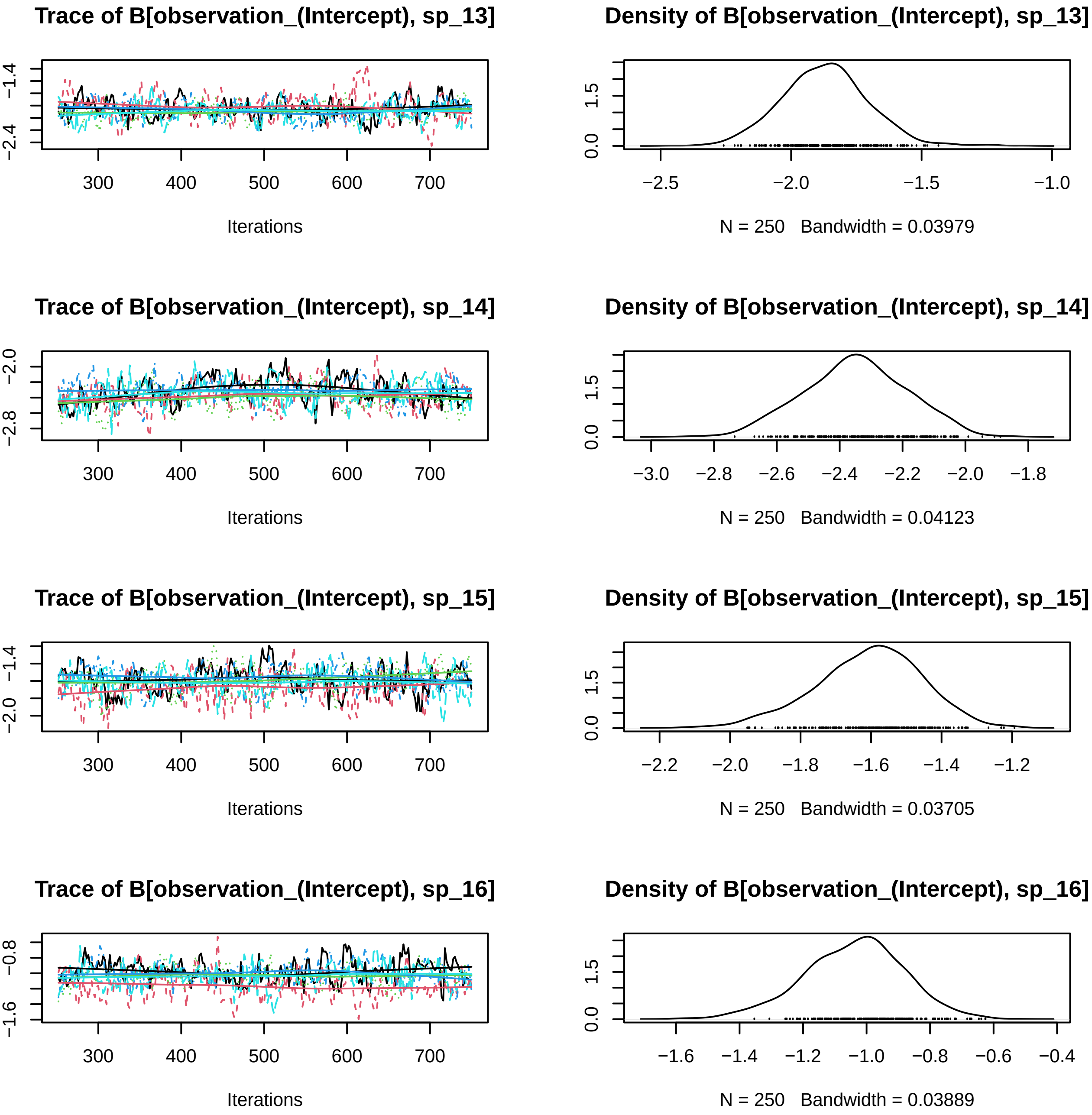
Trace of B[observation_(Intercept), sp_13]
−2.4
−1.4
300
400
500
600
700
Iterations
Density of B[observation_(Intercept), sp_13]
0.0
1.5
−2.5
−2.0
−1.5
−1.0
N = 250 Bandwidth = 0.03979
Trace of B[observation_(Intercept), sp_14]
−2.8
−2.0
300
400
500
600
700
Iterations
Density of B[observation_(Intercept), sp_14]
0.0
1.5
−3.0
−2.8
−2.6
−2.4
−2.2
−2.0
−1.8
N = 250 Bandwidth = 0.04123
Trace of B[observation_(Intercept), sp_15]
−2.0
−1.4
300
400
500
600
700
Iterations
Density of B[observation_(Intercept), sp_15]
0.0
1.5
−2.2
−2.0
−1.8
−1.6
−1.4
−1.2
N = 250 Bandwidth = 0.03705
Trace of B[observation_(Intercept), sp_16]
−1.6
−0.8
300
400
500
600
700
Iterations
Density of B[observation_(Intercept), sp_16]
0.0
1.5
−1.6
−1.4
−1.2
−1.0
−0.8
−0.6
−0.4
N = 250 Bandwidth = 0.03889

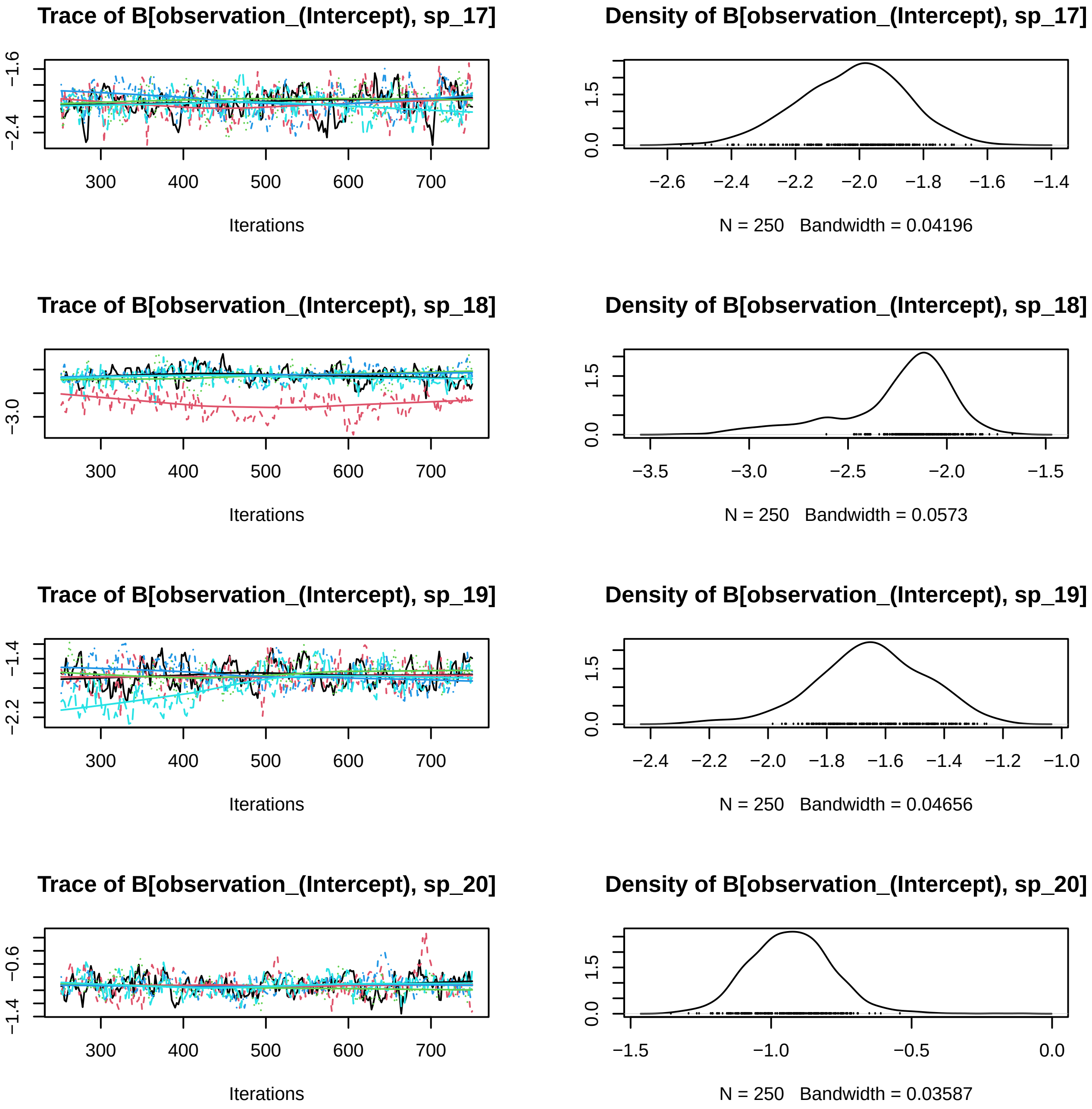

Trace of B[observation_(Intercept), sp_17]
−1.6
−2.4
300
400
500
600
700
Iterations
Density of B[observation_(Intercept), sp_17]
0.0
1.5
−2.6
−2.4
−2.2
−2.0
−1.8
−1.6
−1.4
N = 250 Bandwidth = 0.04196
Trace of B[observation_(Intercept), sp_18]
−3.0
300
400
500
600
700
Iterations
Density of B[observation_(Intercept), sp_18]
0.0
1.5
−3.5
−3.0
−2.5
−2.0
−1.5
N = 250 Bandwidth = 0.0573
Trace of B[observation_(Intercept), sp_19]
−1.4
−2.2
300
400
500
600
700
Iterations
Density of B[observation_(Intercept), sp_19]
0.0
1.5
−2.4
−2.2
−2.0
−1.8
−1.6
−1.4
−1.2
−1.0
N = 250 Bandwidth = 0.04656
Trace of B[observation_(Intercept), sp_20]
−0.6
−1.4
300
400
500
600
700
Iterations
Density of B[observation_(Intercept), sp_20]
0.0
1.5
−1.5
−1.0
−0.5
0.0
N = 250 Bandwidth = 0.03587

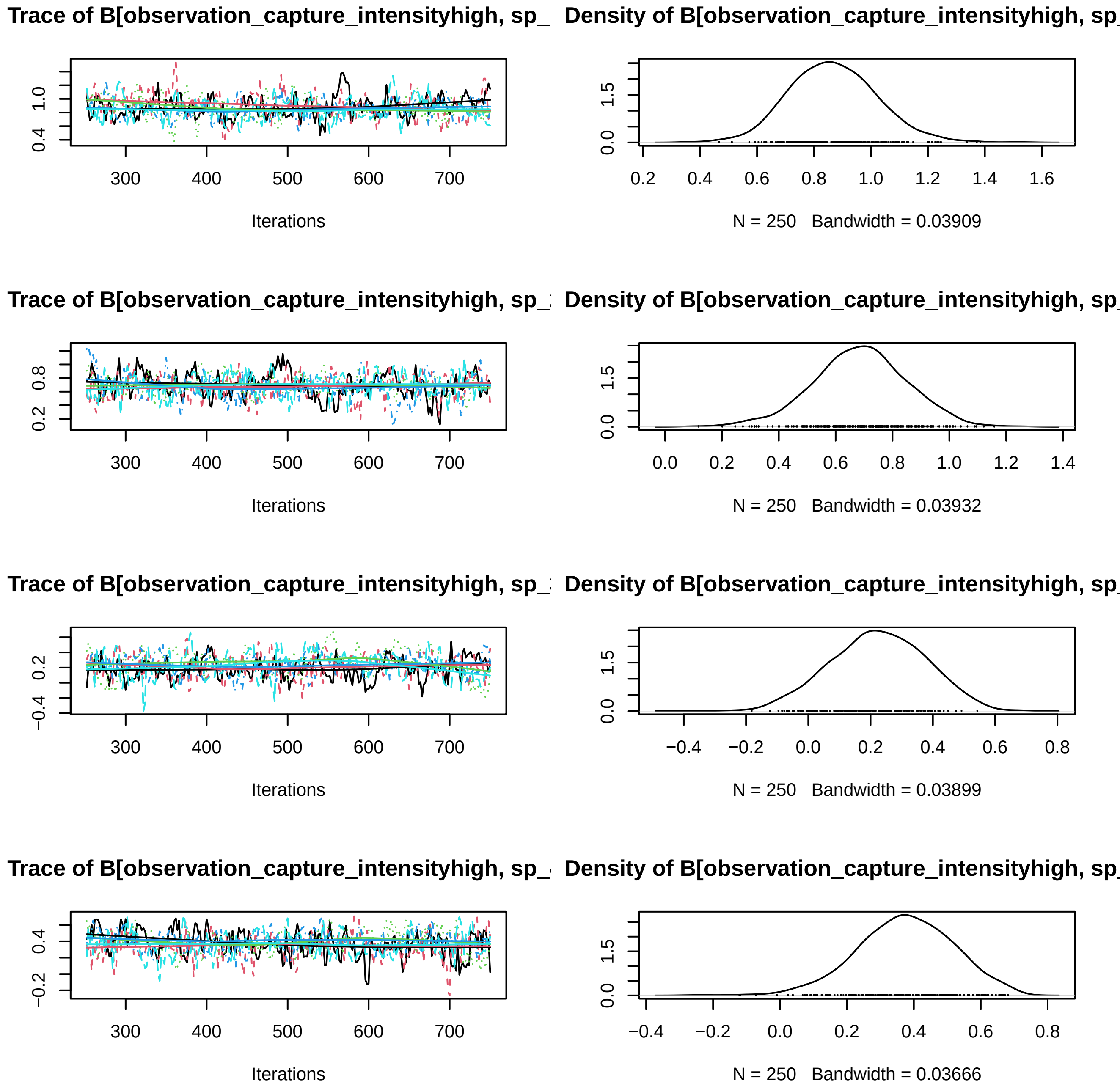

Trace of B[observation_capture_intensityhigh, sp_
Density of B[observation_capture_intensityhigh, sp_
Iterations
N = 250 Bandwidth = 0.03909
Trace of B[observation_capture_intensityhigh, sp_
Density of B[observation_capture_intensityhigh, sp_
Iterations
N = 250 Bandwidth = 0.03932
Trace of B[observation_capture_intensityhigh, sp_
Density of B[observation_capture_intensityhigh, sp_
Iterations
N = 250 Bandwidth = 0.03899
Trace of B[observation_capture_intensityhigh, sp_
Density of B[observation_capture_intensityhigh, sp_
Iterations
N = 250 Bandwidth = 0.03666

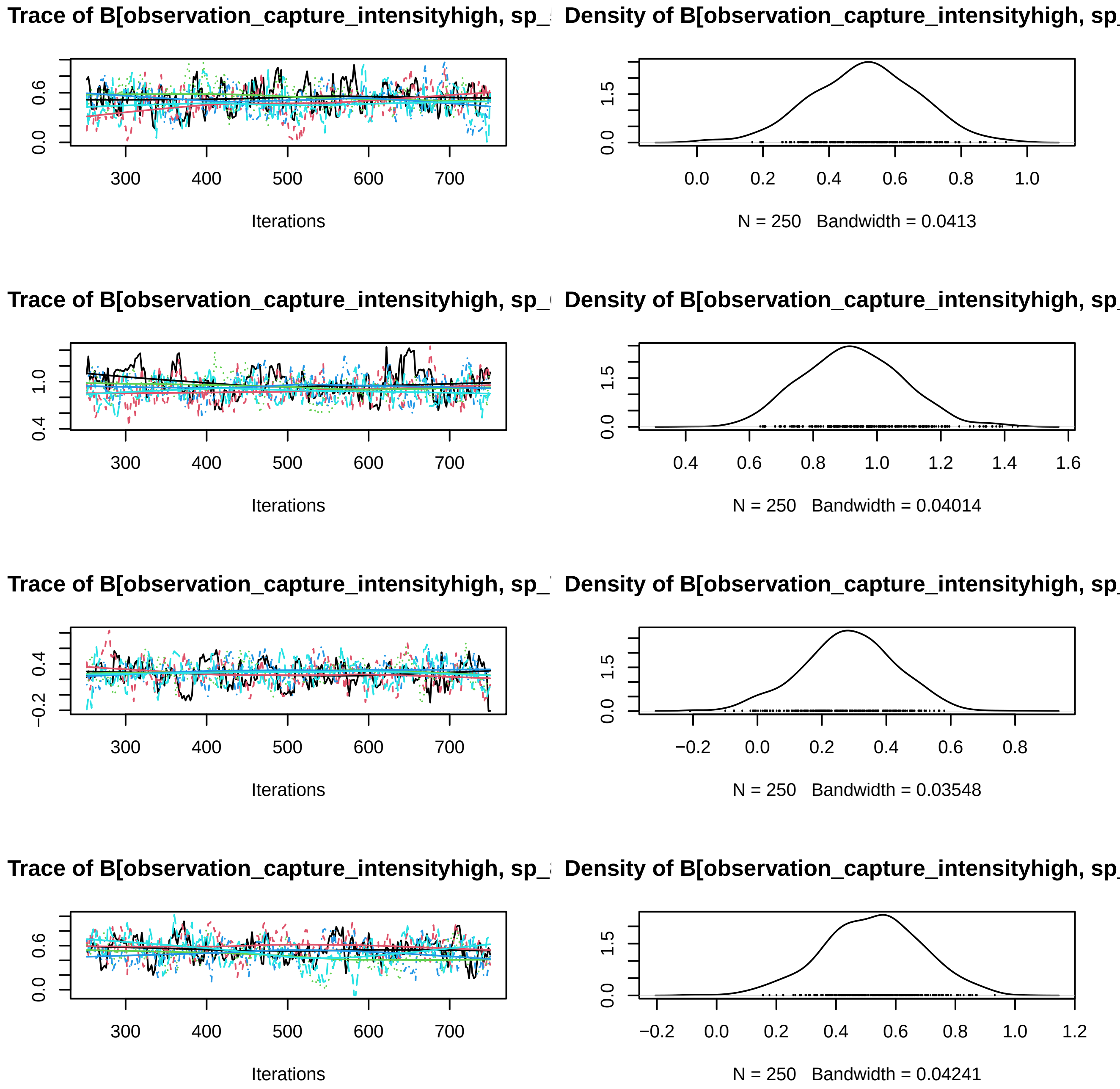
Trace of B[observation_capture_intensityhigh, sp_
Density of B[observation_capture_intensityhigh, sp_
Iterations
N = 250 Bandwidth = 0.0413
Trace of B[observation_capture_intensityhigh, sp_
Density of B[observation_capture_intensityhigh, sp_
Iterations
N = 250 Bandwidth = 0.04014
Trace of B[observation_capture_intensityhigh, sp_
Density of B[observation_capture_intensityhigh, sp_
Iterations
N = 250 Bandwidth = 0.03548
Trace of B[observation_capture_intensityhigh, sp_
Density of B[observation_capture_intensityhigh, sp_
Iterations
N = 250 Bandwidth = 0.04241

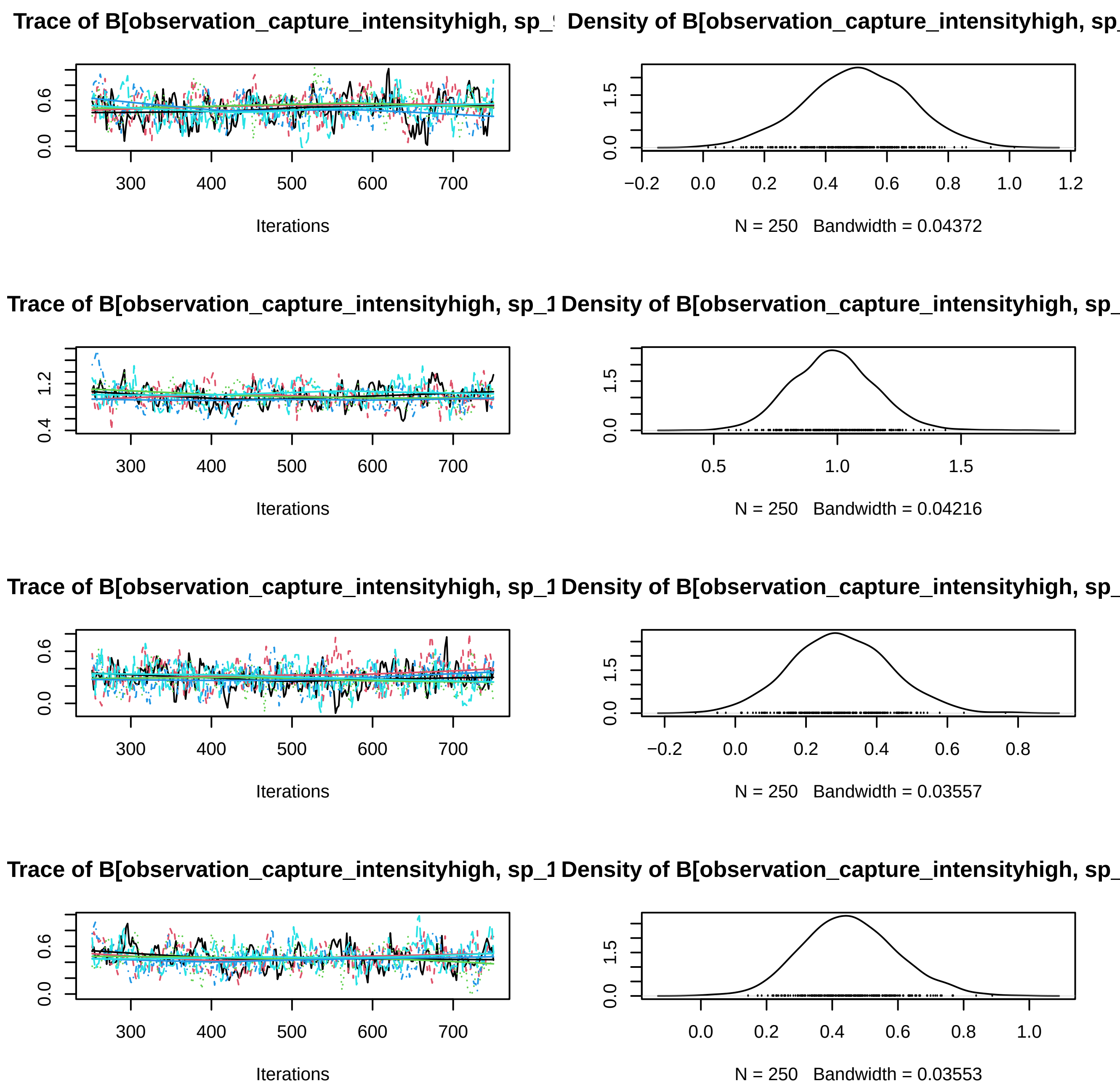
Trace of B[observation_capture_intensityhigh, sp_
Density of B[observation_capture_intensityhigh, sp_
Iterations
N = 250 Bandwidth = 0.04372
Trace of B[observation_capture_intensityhigh, sp_1
Density of B[observation_capture_intensityhigh, sp_
Iterations
N = 250 Bandwidth = 0.04216
Trace of B[observation_capture_intensityhigh, sp_1
Density of B[observation_capture_intensityhigh, sp_
Iterations
N = 250 Bandwidth = 0.03557
Trace of B[observation_capture_intensityhigh, sp_1
Density of B[observation_capture_intensityhigh, sp_
Iterations
N = 250 Bandwidth = 0.03553

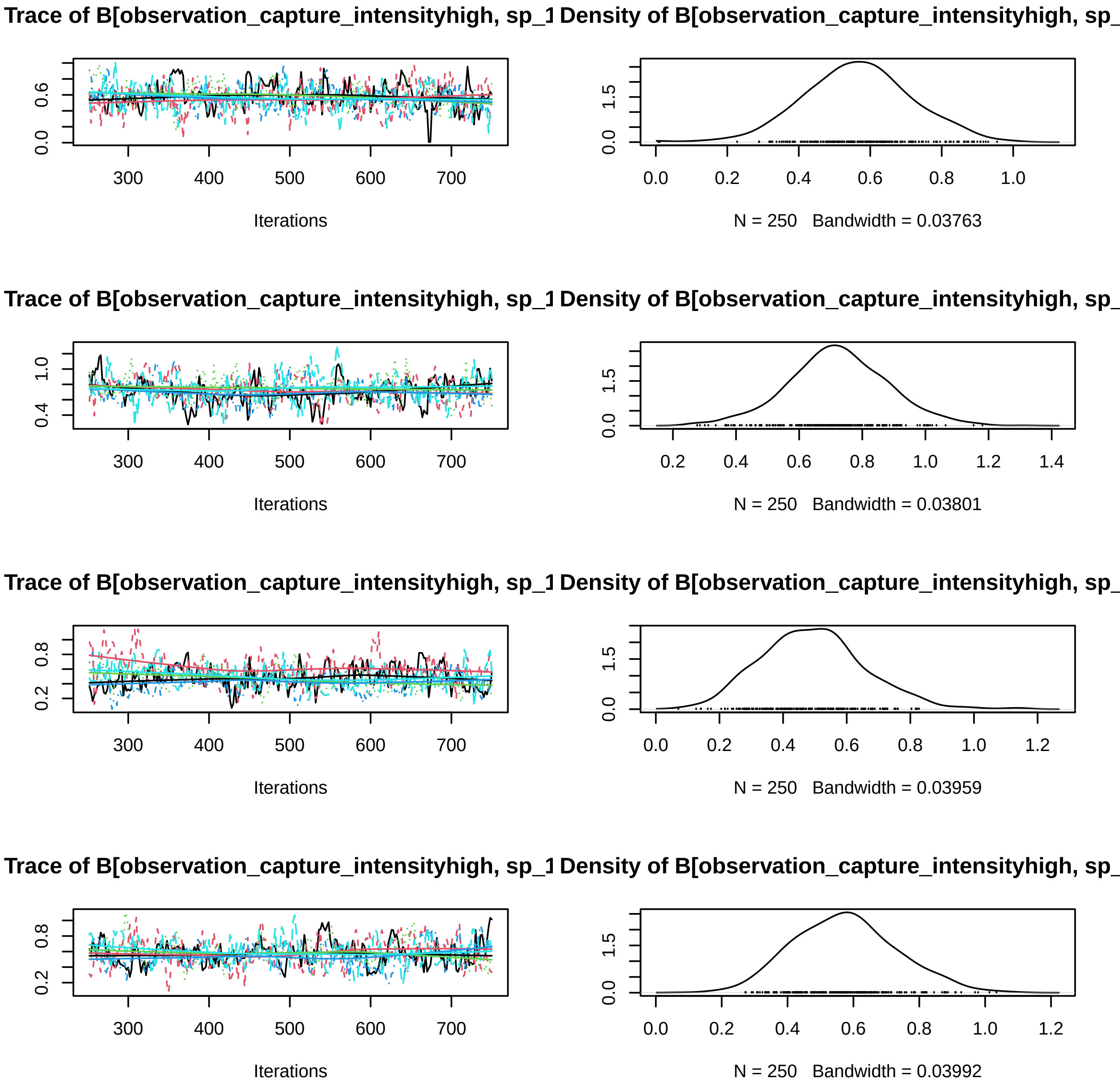

Trace of B[observation_capture_intensityhigh, sp_1
Density of B[observation_capture_intensityhigh, sp_
Iterations
N = 250 Bandwidth = 0.03763
Trace of B[observation_capture_intensityhigh, sp_1
Density of B[observation_capture_intensityhigh, sp_
Iterations
N = 250 Bandwidth = 0.03801
Trace of B[observation_capture_intensityhigh, sp_1
Density of B[observation_capture_intensityhigh, sp_
Iterations
N = 250 Bandwidth = 0.03959
Trace of B[observation_capture_intensityhigh, sp_1
Density of B[observation_capture_intensityhigh, sp_
Iterations
N = 250 Bandwidth = 0.03992

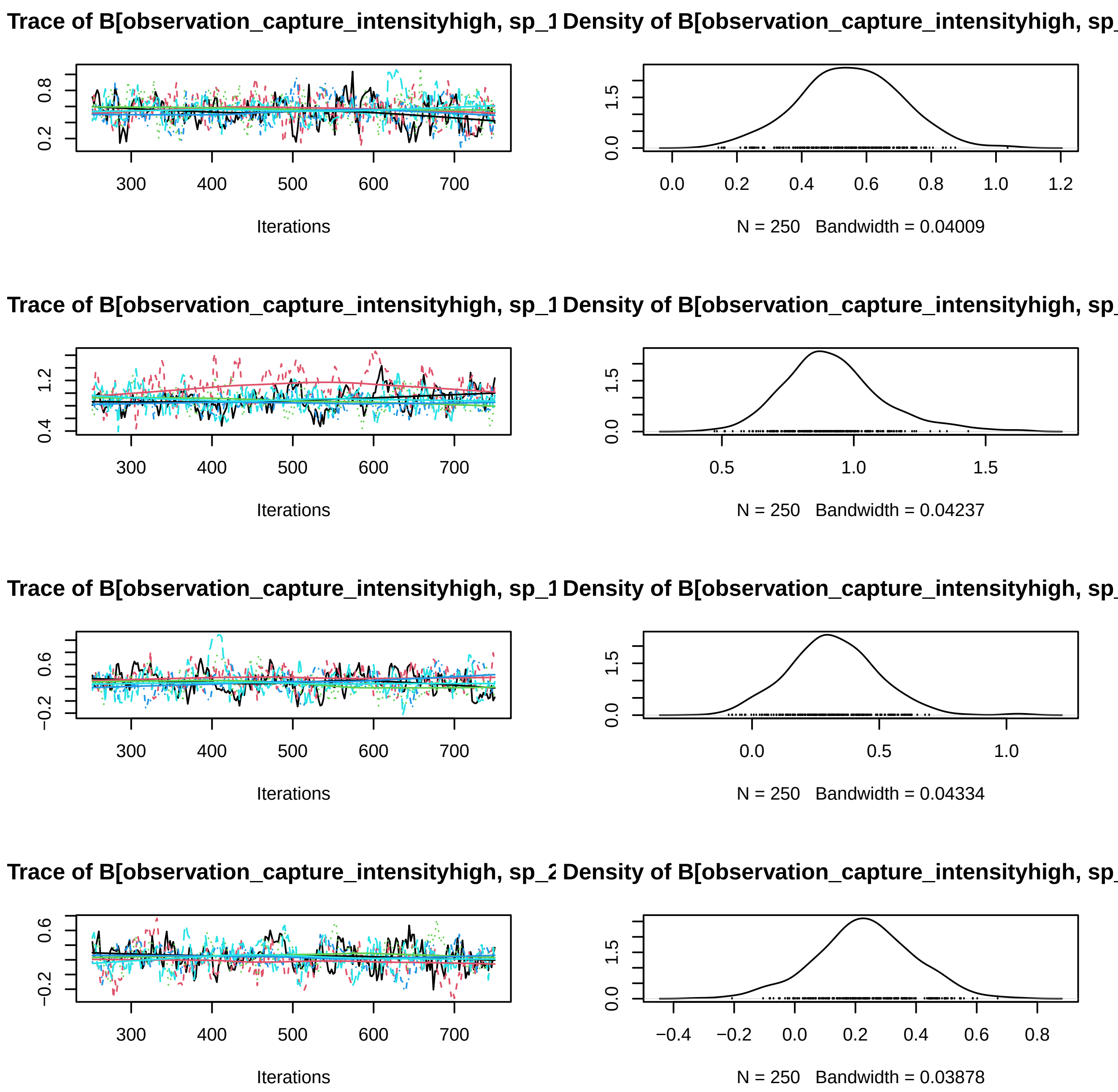
Trace of B[observation_capture_intensityhigh, sp_1
Density of B[observation_capture_intensityhigh, sp_
Iterations
N = 250 Bandwidth = 0.04009
Trace of B[observation_capture_intensityhigh, sp_1
Density of B[observation_capture_intensityhigh, sp_
Iterations
N = 250 Bandwidth = 0.04237
Trace of B[observation_capture_intensityhigh, sp_1
Density of B[observation_capture_intensityhigh, sp_
Iterations
N = 250 Bandwidth = 0.04334
Trace of B[observation_capture_intensityhigh, sp_2
Density of B[observation_capture_intensityhigh, sp_
Iterations
N = 250 Bandwidth = 0.03878

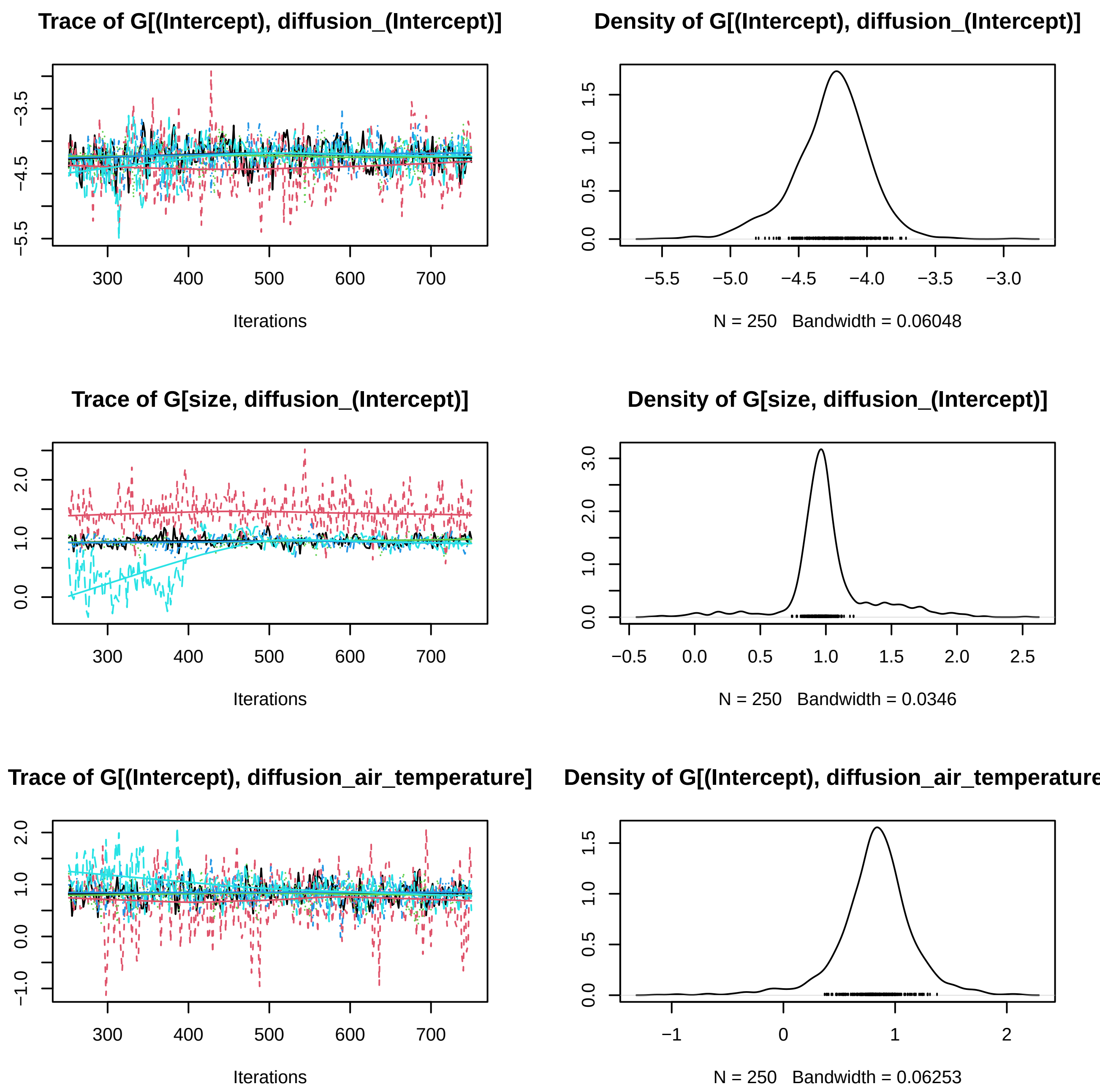
Trace of G[(Intercept), diffusion_(Intercept)]
Iterations
Density of G[(Intercept), diffusion_(Intercept)]
N = 250 Bandwidth = 0.06048
Trace of G[size, diffusion_(Intercept)]
Iterations
Density of G[size, diffusion_(Intercept)]
N = 250 Bandwidth = 0.0346
Trace of G[(Intercept), diffusion_air_temperature]
Iterations
Density of G[(Intercept), diffusion_air_temperature
N = 250 Bandwidth = 0.06253

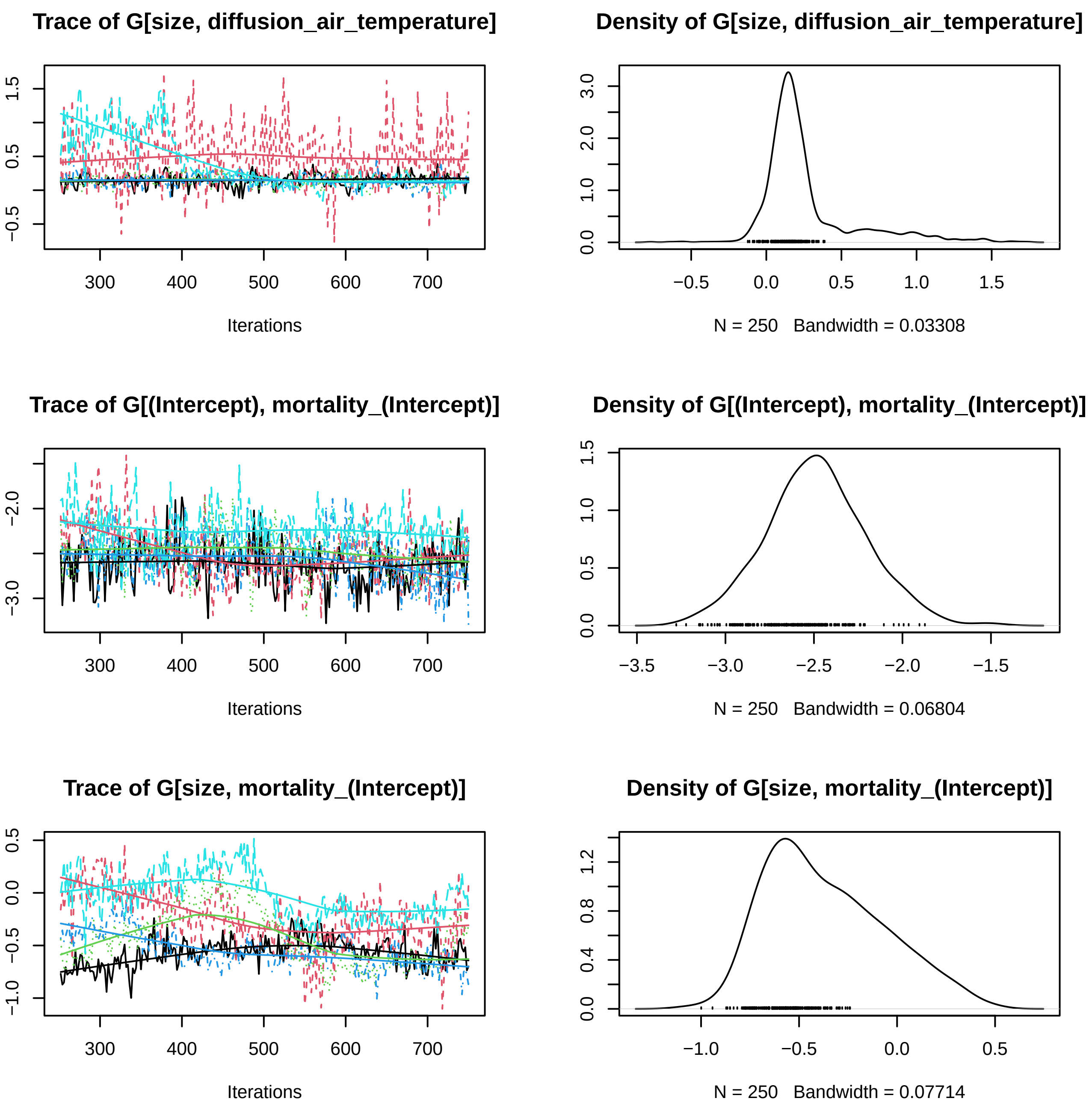
Trace of G[size, diffusion_air_temperature]
−0.5
0.5
1.5
300
400
500
600
700
Iterations
Density of G[size, diffusion_air_temperature]
0.0
1.0
2.0
3.0
−0.5
0.0
0.5
1.0
1.5
N = 250 Bandwidth = 0.03308
Trace of G[(Intercept), mortality_(Intercept)]
−3.0
−2.0
300
400
500
600
700
Iterations
Density of G[(Intercept), mortality_(Intercept)]
0.0
0.5
1.0
1.5
−3.5
−3.0
−2.5
−2.0
−1.5
N = 250 Bandwidth = 0.06804
Trace of G[size, mortality_(Intercept)]
−1.0
−0.5
0.0
0.5
300
400
500
600
700
Iterations
Density of G[size, mortality_(Intercept)]
0.0
0.4
0.8
1.2
−1.0
−0.5
0.0
0.5
N = 250 Bandwidth = 0.07714

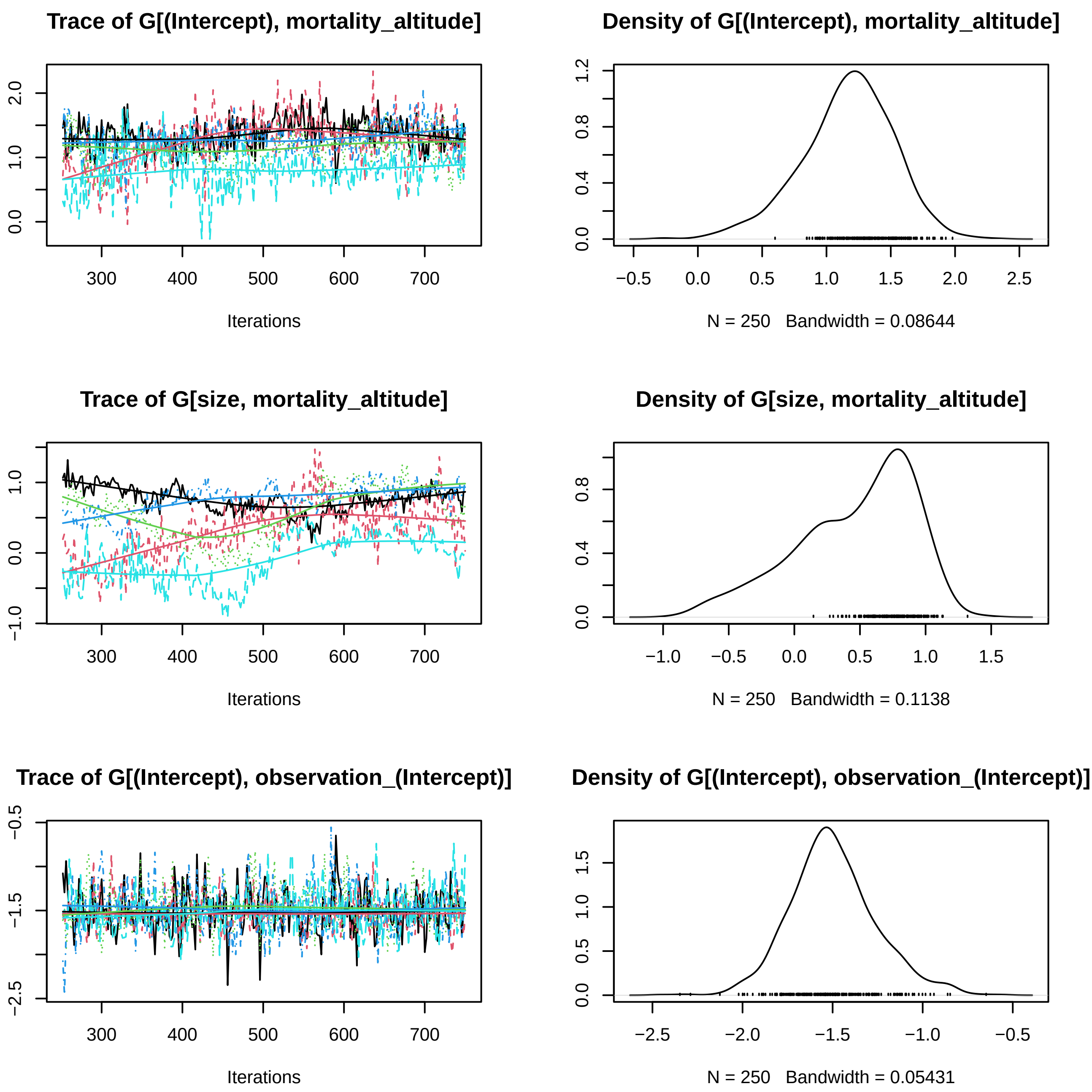
Trace of G[(Intercept), mortality_altitude]
Iterations
N = 250 Bandwidth = 0.08644
Density of G[(Intercept), mortality_altitude]
Trace of G[size, mortality_altitude]
Iterations
Density of G[size, mortality_altitude]
N = 250 Bandwidth = 0.1138
Trace of G[(Intercept), observation_(Intercept)]
Iterations
Density of G[(Intercept), observation_(Intercept)]
N = 250 Bandwidth = 0.05431

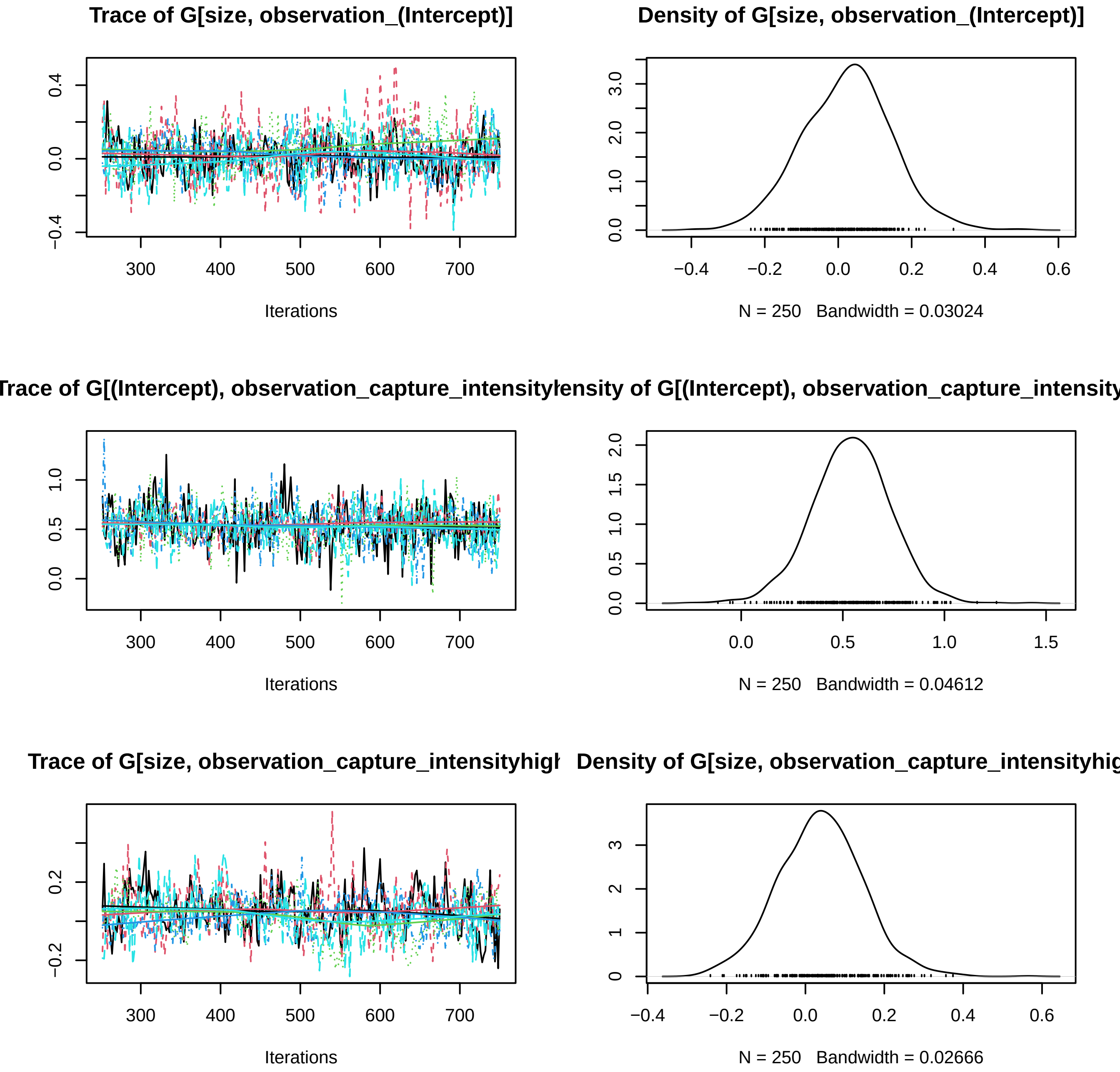
Trace of G[size, observation_(Intercept)]
Iterations
Density of G[size, observation_(Intercept)]
N = 250   Bandwidth = 0.03024
Trace of G[(Intercept), observation_capture_intensity
Iterations
ensity of G[(Intercept), observation_capture_intensity
N = 250   Bandwidth = 0.04612
Trace of G[size, observation_capture_intensityhigl
Iterations
Density of G[size, observation_capture_intensityhig
N = 250   Bandwidth = 0.02666

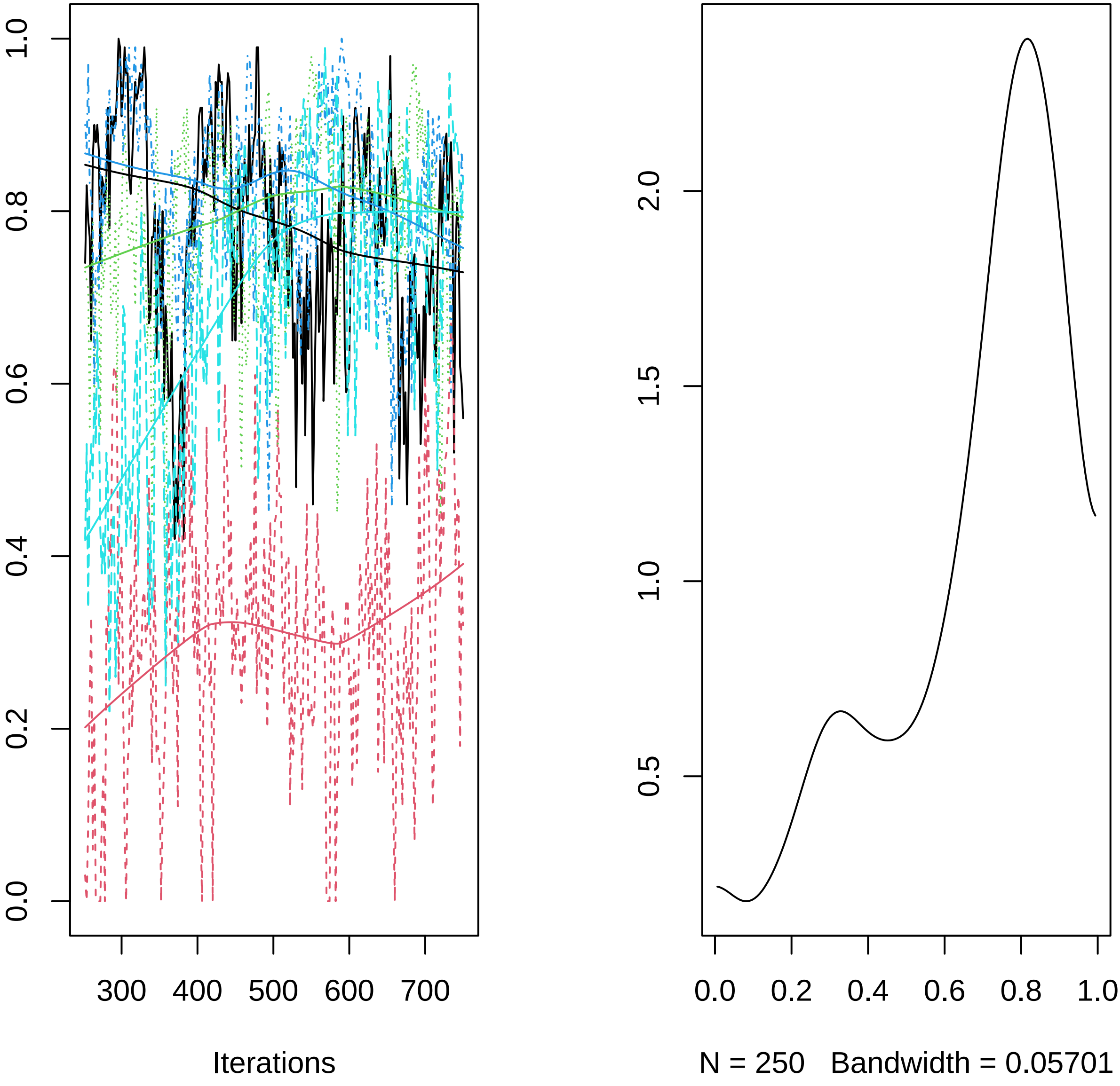

0.0
0.2
0.4
0.6
0.8
1.0
300
400
500
600
700
Iterations
0.5
1.0
1.5
2.0
0.0
0.2
0.4
0.6
0.8
1.0
N = 250   Bandwidth = 0.05701
