## Supplementary material for "R-package Jsmm: Joint species movement modelling of mark-recapture data": S2 Vignettes demonstrating the application of the R-package Jsmm: Supporting_information_S2.html

#### 2026-05-27

### Preface

Rodriguez, L. F., & Ovaskainen, O. (2026). R-package Jsmm: Joint species movement modelling of mark-recapture data. bioRxiv. doi:10.64898/2026.02.24.707702

This book demonstrates the use of the Jsmm R-package. The Jsmm implementation and the demo scripts for running analyses can be found in GitHub.

The folder scripts\_simulated contains all the steps to reproduce the simulated examples (Demos simulated example ICP and CCP (covariates)).

For running Jsmm analyses, the main project folder should have subfolders called models, parameters, and results. The models folder saves the data corresponding to the model structure and all its posterior modifications, the optional folder parameters saves the simulated model parameters data, and the results folder stores the main results as .PDF format.

License CC BY-NC-ND 4.0. Rodriguez and Ovaskainen. All rights reserved, including those for text and data mining, AI training, and similar technologies.

### Demo simulated example ICP (no covariates)

#### Example Description

Let’s suppose that a researcher is interested in exploring the movement activity and vital dynamics for *two* focal species. To do this, an experiment is conducted over a landscape by setting nine trapping devices for eleven days, as shown in Fig. 1. Initially, 500 animals of each species are marked and released from different locations at different times. All capture sites are visited daily, starting from the second day until the end of the experiment. While the capture sites are visited, animals are marked, and captures are recorded. Secondary releases are possible for this example.

To run the analyses within the Jsmm implementation, we will follow the following steps:

- **Step 1a: Define model**

  - Creating the domain
  - Creating the observation effort
  - Setting data model
  - Information about the focal species
  - Defining model m
- **Step 1b: Simulate model parameters from priors**
- **Step 1c: Simulate capture data**
- **Step 1d: Visualize likelihood profiles**
- **Step 2a: Fit the model**
- **Step 2b: Evaluate MCMC convergence**
- **Step 2c: Simulate predictive data**
- **Step 2d: Evaluate model fit**
- **Step 3a: Show posteriors vs priors**
- **Step 3b: Visualize movement trajectories**

For detailed information about the functions and their corresponding outcomes involved in our analyses, please refer to the Jsmm documentation and Supplementary material. Main paper.

#### Step 0: Loading the package

We start by loading the Jsmm package.

```
library(Jsmm)
```

#### Step 1a: Define model

##### Creating a domain

To create the simulated domain, we use the function **`creating_domain_from_locations(.)`**. This function allows the creation of a customized rectangular domain and capture and release sites. The output of this function is a polygonal object from the class sf (Pebesma and Bivand (2023), Pebesma (2018)).

For this particular example, we created a square domain with 16 nodes or 15 units per side and 9 square capture/release sites (See Fig. 1.).

```
p = c(2.5/15, 7.5/15, 12.5/15) 

lc = as.matrix(expand.grid("x" = p, "y" = p)) # Positions capture and release sites 
lc_label = 1:nrow(lc) 

buffer_x = c(2.5/15, 2.5/15) 
buffer_y = c(2.5/15, 2.5/15) 

nnod_side_x = 16 # number of nodes per side x
nnod_side_y = 16 # number of nodes per side y

rad_capture_site     =  sqrt(2)/10 # site radius
n_sides_capture_site = 4           # number of site sides
angle_capture_site   = pi/4        # site rotation angle
int_buffer           = FALSE       # insert intermediate buffer OPTIONAL

domain_p = Jsmm::creating_domain_from_locations(lc = lc, lc_label = lc_label, 
                                          buffer_x = buffer_x, 
                                          buffer_y = buffer_y, 
                                          nnod_side_x = nnod_side_x, 
                                          nnod_side_y = nnod_side_y, 
                                          rad_capture_site = rad_capture_site,
                                          n_sides_capture_site = n_sides_capture_site, 
                                          angle_capture_site = angle_capture_site, 
                                          int_buffer = int_buffer)
```

The numerical method on which the software is based requires a triangulation of the domain. To do this, we utilize the function **`jsmm_add_triangulation(.)`**. This function returns a named list called **`domain`** that contains the corresponding triangulation **`domain$triangulation`**, and its polygonal representation **`domain$polygon`**. The function parameters **`min_t_angle`** and **`max_t_area`** set the triangles’ minimum angles and maximum areas, respectively (Shewchuk (1996)).

```
max_t_area = NULL
min_t_angle = 30

domain = Jsmm::jsmm_add_triangulation(domain = domain_p, max_t_area = max_t_area, 
                                min_t_angle = min_t_angle)
```

The **`domain$triangulation`** list contains spatial information about the domain triangulation. For this particular case, we have a triangulation composed of 258 elements and 160 nodes.

```
head(domain$triangulation$ele)
```

```
##      node_1 node_2 node_3
## [1,]     62     98     97
## [2,]     15     16     62
## [3,]     14     15     62
## [4,]     14     62     97
## [5,]     14     97     13
## [6,]     62     16     17
```

```
nrow(domain$triangulation$ele)
```

```
## [1] 258
```

```
head(domain$triangulation$node)
```

```
##      node_x    node_y
## [1,]      0 1.0000000
## [2,]      0 0.9333333
## [3,]      0 0.8666667
## [4,]      0 0.8000000
## [5,]      0 0.7333333
## [6,]      0 0.6666667
```

```
nrow(domain$triangulation$node)
```

```
## [1] 160
```

Within **`domain$polygon`**, it is possible to set column indicators for the habitat classification (**`id`**), and for the release/capture locations (**`id_location`**). Each row contains information about each triangle or element.

```
head(domain$polygon)
```

```
## Simple feature collection with 6 features and 2 fields
## Geometry type: POLYGON
## Dimension:     XY
## Bounding box:  xmin: 0 ymin: 0 xmax: 0.1916667 ymax: 0.2
## CRS:           NA
##   id id_location                       geometry
## 1  2           1 POLYGON ((0.06666667 0.0666...
## 2  2           0 POLYGON ((0 0.06666667, 0 0...
## 3  2           0 POLYGON ((0 0.1333333, 0 0....
## 4  2           0 POLYGON ((0 0.1333333, 0.06...
## 5  2           0 POLYGON ((0 0.1333333, 0.06...
## 6  2           0 POLYGON ((0.06666667 0.0666...
```

The function **`plot_domain(.)`** shows the triangulated domain. The **`customize_plot`** list shown below provides control of the main aspects of the visualization.

```
  customize_plot = list()
  customize_plot[["color_map"]]   = "#f0f0f0"
  customize_plot[["labs_axis"]]   = c("X", "Y")
  customize_plot[["lab_legend"]]  = "Matrix"
  customize_plot[["name_legend"]] = "Habitat"
  customize_plot[["color_polyb"]] = "#d6dbdf"
  customize_plot[["lwd_poly"]]    = 0.001
  customize_plot[["site_col"]]    = "red"
  customize_plot[["siteb_col"]]   = "darkred"
  customize_plot[["site_text_pos"]]   = c(-0.05, -0.05) 
  customize_plot[["site_text_size"]]  = 4.2
  customize_plot[["panelb_color"]]    = "black" 
  customize_plot[["panelb_lwd"]]  = 1
```

```
 Jsmm::plot_domain(domain = domain, customize_plot = customize_plot)
```

Figure 1: Study region for the simulated domain with 9 capture/release sites. The triangulation is composed of 258 elements and 160 nodes.

Note: To explore in detail the generated triangulation, is possible to utilize the function **`mapview()`** from the R-package mapview:

```
mapview(domain$polygon, zcol = "id") # visualize triangulation with zoom
```

##### Creating the observation effort

The **`observation_effort`** is a named list that contains all information about how releases and captures occur in the experiment. The list is composed of a string called **`method`**, named lists called **`locations`**, **`releases`**, and **`captures`**. Fig. 2. shows information about the release and capture events in the experiment. The list **`locations`** contains spatial information about the sites in which releases or captures occurred. For this example, we set the capture process within **`method`** as ICP. Below is the script to set the **`observation_effort`**.

```
method          = "ICP"
study_duration  = 10 # duration of the entire study
recapture_sites = 1:9
trap_checking_interval = 1
  
nloc = length(recapture_sites)
observation_effort = list()
observation_effort[["method"]] = method

locations = list()  # Characterizing the elements that belong to each capture/release site
  
for(i in 1:nloc){
  locations[[i]] = which(domain[["polygon"]][["id_location"]] == recapture_sites[i])
}
  
observation_effort[["locations"]] = locations
```

```
observation_effort$method
```

```
## [1] "ICP"
```

```
length(observation_effort$locations)
```

```
## [1] 9
```

Information about the elements that belong to location 5 can be retrieved by

```
observation_effort$locations[[5]]
```

```
## [1]  43  46 108 114 116 217
```

The list **`releases`**, has the vectors **`time`**, and **`location`**, which provide information about when and where releases are possible. Each possible relationship is called a release event. For this simple example, we have set ten release events.

```
observation_effort[["releases"]][["location"]] = sample(recapture_sites,
                                                        size = study_duration, 
                                                        replace = T)

observation_effort[["releases"]][["time"]] = 0:(study_duration - 1)
```

The release event 1 was held on location 5 at time \(t = 0\). This information can be retrieved as

```
observation_effort$releases$time[1]
```

```
## [1] 0
```

```
observation_effort$releases$location[1]
```

```
## [1] 5
```

The list **`captures`** is composed of the vector **`location`** and the vector **`time`**. Below, we set this list for the ICP capture process.

```
recapture_times = c()

for(i in seq(trap_checking_interval, study_duration, by = trap_checking_interval)){
  recapture_times = c(recapture_times, rep(i, nloc)) 
}

recapture_locations = rep(recapture_sites, study_duration/trap_checking_interval) 
observation_effort[["captures"]][["location"]] = recapture_locations
observation_effort[["captures"]][["time"]]     = recapture_times
```

The number of capture events is

```
length(observation_effort$captures$time)
```

```
## [1] 90
```

Information corresponding to the capture event 12 is retrieved as

```
observation_effort$captures$time[12]
```

```
## [1] 2
```

```
observation_effort$captures$location[12]
```

```
## [1] 3
```

Read as: the twelfth capture event occurred at capture site number 3 at time \(t = 2\) (third day).

Figure 2: Observation effort scheme for ICP capture process. Times in which releases and captures are possible are represented by blue and red colors, respectively.

##### Set covariate data

The Jsmm framework allows the inclusion of spatial, temporal, spatio-temporal covariates, and trait covariates. To do this, first, we are required to provide the data in a suitable format. The covariates are set within the named lists **`cov_data`** and **`tr_data`**.

For this example, we do not have any covariates; for illustration, we declare this empty list as:

```
cov_data = NULL
ns       = 2
sp_names = paste0("sp_", 1:ns)
tr_data  = NULL
C        = NULL
```

##### Setting the data model

In this section, we are going to set the named list **`model_formula`** to declare explicitly which parameters are going to be included in the model. See Appendix S1. Main paper for a detailed description.

The **`model_formula`** list is composed of (some combination of) the objects **`diffusion`**, **`advection_1`** (advection in x), **`advection_2`** (advection in y), **`mortality`**, **`habitat_preference`**, **`observation`**, and **`traits`**.

For declaring the formulas, the ~ symbol is used. If the model does not include a given parameter, this should be declared as **`NULL`**. If the parameter is included but no covariates are declared, this should be set as **`~1`**, this applies to all parameters except for the habitat preference parameter **`habitat_preference`**. This parameter is declared in the model formula only when covariates are included.

For our example, the model includes diffusion and mortality movement parameters, and the observation parameter. Table 1. shows the formulas declarations for this model. This model formula is set as:

```
model_formula = list()
model_formula[["diffusion"]]          = ~ 1
model_formula[["advection_1"]]        = NULL
model_formula[["advection_2"]]        = NULL
model_formula[["mortality"]]          = ~ 1
model_formula[["habitat_preference"]] = NULL
model_formula[["traits"]]             = NULL
model_formula[["observation"]]        = ~ 1
```

Model Formulas declarations for the simulated example.

| Model parameter | Model formula |
| --- | --- |
| **`diffusion`** | ~ 1 |
| **`advection_1`** | - |
| **`advection_2`** | - |
| **`habitat_preference`** | - |
| **`mortality`** | ~ 1 |
| **`observation`** | ~ 1 |
| **`traits`** | - |

At this stage, for our simulated example, we do not have mark-recapture data. For models for which this information is available, the data should be included on the named list **`data`** with sublists **`data$releases`** and **`data$captures`**. In the next section, more explanation will be provided about this. For illustration, we declare the empty list **`data`** here as:

```
  data = list()
  data[["captures"]] = NULL
  data[["releases"]] = NULL
```

##### Define the model object **`m`**

The model object **`m`** is a named list composed of all the elements created in previous steps. To define the model **`m`**, some actions occur within the function **`jsmm(.)`**. See Appendix S1. of the main manuscript for detailed information.

The model **`m`** for our example is set as

```
m = Jsmm::jsmm(domain = domain, observation_effort = observation_effort,
          releases = data[["releases"]], captures = data[["captures"]], 
          cov_data = cov_data, 
          ns = ns, sp_names = sp_names, tr_data = tr_data, C = C,
          model_formula = model_formula)
```

The function **`model_summary()`** retrieves a summary of our model specifications:

```
Jsmm::model_summary(m)
```

```
## Domain: nele = 258 | nnod = 160 
## Capture method: ICP | Number release sites: 4 | Number capture sites: 9 
## Number of time partitions: 11 
## ------------------------------------- 
## Model formulas: 
##     Diffusion ~ 1 | nc = 1 
##     Mortality ~ 1 | nc = 1 
##     Observation ~ 1 | nc = 1 
## ------------------------------------- 
## Number of species: 2
```

#### Step 1b: Simulate model parameters from priors

The function **`simulate_from_prior(m)`** allows generating the model parameters from predefined priors (See Appendix S1. and Fig. 1. main manuscript). The script below sets the diffusion, mortality, and observation parameters.

```
muZ = m[["prior"]][["muZ"]]
nc  = m[["nc"]]
np  = sum(nc)
nt  = m[["nt"]]
idx = 1

for(j in 1:nt){
  for(i in 1:length(nc)){
    for(ii in seq_len(nc[i])){
      # baseline diffusion
      if(names(nc)[i] == "diffusion" & ii == 1 & j == 1) muZ[idx] = -4
      
      # baseline mortality
      if(names(nc)[i] == "mortality" & ii == 1 & j == 1) muZ[idx] = -2 
      
      # baseline observation rate
      if(names(nc)[i] == "observation" & ii == 1 & j == 1) muZ[idx] = -2 
      
      idx = idx + 1
    }
  }
}

m[["prior"]][["muZ"]] = muZ
m[["prior"]][["VZ"]]  = 0*m[["prior"]][["VZ"]]
m[["prior"]][["rho"]] = matrix(c(0.75, 1), nrow = 1)

pars = Jsmm::simulate_from_prior(m)
```

Table 1: Model coeficients generated from priors for diffusion, mortality and observation parameters for the two species.

| species | betaa0 | betam0 | betao0 |
| --- | --- | --- | --- |
| 1 | -3.331 | -2.099 | -2.973 |
| 2 | -3.815 | -2.285 | -3.581 |

The model parameters are set up within the named list **`pars$Beta`**. Within each element of this list, there are matrices for each of the movement and observation parameters, that is, **`pars$Beta$diffusion`**, **`pars$Beta$mortality`**, and **`pars$Beta$observation`**. For example, for the diffusion parameter, a matrix with two rows and one column is created. Rows correspond to the species, the column corresponds to the coefficient \(\beta^a\_0\).

#### Step 1c: Simulate capture data

To start the simulation, we need to provide release information about the individuals. This is done by defining a matrix called **`releases`**, which is composed of two columns. The first and second columns indicate the species and release event, respectively. Each row provides information about each unique individual. For this example, we simulated 500 individuals for each species, allowing secondary releases.

```
ni = round(1000/m[["ns"]]) #number of individuals per species
releases = matrix(NA, ncol = 2, nrow = ni*m[["ns"]])
releases[, 1] = rep(1:m[["ns"]], ni)
releases[, 2] = rep(1, ni*m[["ns"]])
releases[, 2] = sample(seq_len(length(m$observation_effort$releases$location)),
                       size = ni*m[["ns"]], replace = TRUE)

colnames(releases) = c("sp", "release_event")
  
secondary_release = TRUE
max_dt = NULL

head(releases)
```

```
##      sp release_event
## [1,]  1             5
## [2,]  2            10
## [3,]  1             7
## [4,]  2             4
## [5,]  1            10
## [6,]  2             8
```

For example, the second row of the **`releases`** matrix indicates that one individual from species 2 was released at the release site number four at \(t = 9\). (release event was declared on the **`observation_effort`** list).

Now we can proceed to simulate capture-recapture data by using the function **`simulate_captures(.)`**.

```
  m = Jsmm::simulate_captures(m = m, pars = pars, releases = releases,
                        max_dt = max_dt, secondary_release = secondary_release)
```

After running **`simulate_captures(.)`**, the releases data created in the previous step is stored as **`m$releases`**, and the simulated data is stored as a matrix in **`m$captures`**. The first column indicates the individual to whom this register corresponds (row number from releases matrix), the second column shows the capture event, and the third column provides information about secondary releases.

```
head(m$captures)
```

```
##      individual capture_event secondary_release
## [1,]        235            14                 1
## [2,]        330            12                 1
## [3,]        349            11                 1
## [4,]        731            16                 1
## [5,]        837            16                 1
## [6,]        931            15                 1
```

For example, the first capture is related to the releases matrix as

```
m$releases[235, ]
```

```
##            sp release_event 
##             1             2
```

Summary: a unique individual from species one with release event 2 was recaptured at the location and time shown by the capture event 14.

Table 2: Numbers of recaptures per species.

| \(sp\_1\) | \(sp\_2\) |
| --- | --- |
| 55 | 42 |

#### Step 1d: Visualize likelihood profiles

This is an optional step for verifying that the log likelihood function is returning accurate results. Fig 3. shows the likelihood profile for our example.

```
max_dt = NULL
method = "ICP"
mname  = paste0("simulated_", method)

load(paste0("models/unfitted_model_", mname, "_0", ".RData"))
load(paste0("parameters/true_parameters_", method, "_0", ".RData"))

m         = Jsmm::merge_rc_data(m)
Beta.TRUE = pars$Beta
vars      = names(Beta.TRUE)
tmp       = Jsmm::compute_likelihood(m, Beta = Beta.TRUE, max_dt = max_dt)
li.TRUE   = tmp$log.likelihood.sp

par(mfrow = c(1, 3))

for(va in vars){
  for(i in 1:ncol(Beta.TRUE[[va]])){
    adds = (-5:5)/5
    na   = length(adds)
    re   = matrix(nrow = na, ncol = m$ns)
    renb = matrix(nrow = na, ncol = m$ns)
    
    for(ai in 1:na){
      Beta =  Beta.TRUE
      Beta[[va]][, i] = Beta[[va]][, i] + adds[ai]
      tmp = Jsmm::compute_likelihood(m, Beta = Beta, max_dt = max_dt)
      li  = tmp$log.likelihood.sp
      re[ai, ] = li - li.TRUE
    }
    
    plot(adds, re[, 1],type = "l", col = "red", main = paste(va, i), ylim = c(min(re), max(re)))
    
    abline(h = 0)
    abline(v = 0)
    
    if(m$ns > 1) for(j in 2:m$ns) lines(adds, re[, j], col = "blue")
  }
}
```

Figure 3: Log likelihood profiles for each of the model parameters.

#### Step 2a: Fit the model

The parameter estimation within the Jsmm is made via Markov Chain Monte Carlo (MCMC) through the function **`sampleMCMC(.)`**. The user should choose reasonable parameters for the sampler to ensure that the MCMC has good mixing, low autocorrelation, and reliable convergence. For this simple example, we will pretend that we do not know the model parameters, and we will proceed to estimate those. The parameters used in our sampler are set in Table 4.

Declaration of the model formulas for the simulated example.

| MCMC sampler parameter | Value |
| --- | --- |
| thin | 2 |
| samples | 500 |
| transcient | 500 |
| number of chains | 5 |

```
mname = "simulated_ICP_0"

load(paste0("models/unfitted_model_", mname, ".RData"))
  
load("parameters/true_parameters_ICP_0.RData")
  
m         = merge_rc_data(m)
thin      = 2
samples   = 500
transient = round(0.5*samples*thin)
nChains   = 5
true_pars = pars

for(i in 1:nChains){
  set.seed(i)
  m = Jsmm::sampleMcmc(m = m, samples = samples, transient = transient, thin = thin, nChains = 1,
                 init_pars = true_pars)
  
  save(m, file = paste0(paste0("models/", i, "_fitted_model_", mname, "_samples_", samples, 
                               "_thin_", thin, "_nChains_", nChains, ".RData")))
}

postList = list()
  
for(i in 1:nChains){
  fname = paste0(paste0("models/", i, "_fitted_model_", mname, "_samples_", samples, 
                        "_thin_", thin, "_nChains_", nChains, ".RData"))
  load(file = fname)
  postList[[i]] = m[["postList"]][[1]]
  file.remove(fname)
}
  
m[["postList"]] = postList
  
save(m, file = paste0(paste0("models/fitted_model_", mname, "_samples_", samples, 
                             "_thin_", thin, "_nChains_", nChains, ".RData")))
```

#### Step 2b: Evaluating MCMC convergence

After fitting the model, we will assess the MCMC convergence. First, we convert the generated chains to objects of the MCMC class. Within our framework, this is done through the function **`convert_to_coda_object(.)`**, which is based on the R-package Coda (Plummer et al. (2006)). Note that the Gelman-Rubin statistic (R-hat) diagnosis of convergence shows that the chains converge satisfactorily. Figs. 4-5 show the trace plots for our model.

Note: The script below generates the corresponding trace plots and saves results in PDF format within a folder called results.

```
mname = "simulated_ICP_0"

thin    = 2
samples = 500
nChains = 5

load(paste0(paste0("models/fitted_model_", mname, 
                   "_samples_", samples, "_thin_", thin, "_nChains_", nChains, ".RData")))

mpost = Jsmm::convert_to_coda_object(m)

summary(coda::gelman.diag(mpost$Beta, multivariate = FALSE)$psrf)
```

```
##    Point est.      Upper C.I.   
##  Min.   :1.005   Min.   :1.014  
##  1st Qu.:1.008   1st Qu.:1.022  
##  Median :1.013   Median :1.036  
##  Mean   :1.014   Mean   :1.036  
##  3rd Qu.:1.016   3rd Qu.:1.044  
##  Max.   :1.028   Max.   :1.068
```

Figure 4: Trace plots for the five MCMC chains.

Figure 5: Trace plots for the five MCMC chains.

#### Step 2c: Simulate prior and posterior predictive data

The main purpose of generating predictive data is to explore how the fitted model behaves in time and space compared to the original data. This method helps to infer if the fitted parameters provide a good approximation of the real data. The next script generates 10, 100, and 1000 datasets by utilizing the exact model.

```
mname = "simulated_ICP_0"

for(nrepl in c(10, 100, 1000)){
  showTrueValues = TRUE
  
  thin    = 2
  samples = 500
  nChains = 5
  
  load(paste0(paste0("models/fitted_model_", mname, "_samples_", samples, "_thin_", thin, 
                     "_nChains_", nChains, ".RData")))
  
  post = m[["postList"]]
  
  if(showTrueValues){
    types = c("true", "prior", "posterior")
    load("parameters/true_parameters_ICP_0.RData")
    true_pars = pars
  }else{
    types =  c("prior","posterior")
  }

  for(type in types){
    PPcaptures = list()

    for(repl in 1:nrepl){
      #print(paste0(type,"_repl_",repl))
      
      if(type == "true"){
        pars = true_pars
      }
      
      if(type == "posterior"){
        chain = sample(1:length(post), size = 1)
        sa    = sample(1:length(post[[chain]]), size = 1)
        pars  = post[[chain]][[sa]]
      }
      
      if(type == "prior"){
        pars = Jsmm::simulate_from_prior(m)
      }
      
      max_dt = NULL
      secondary_release = TRUE
      releases = m[["releases"]]
      
      m2 = Jsmm::simulate_captures(m = m, pars = pars, releases = releases,
                               max_dt = max_dt, secondary_release = secondary_release)
        
      captures.simulated = m2[["captures"]]
        
      if(nrow(captures.simulated) > 0){
        PPcaptures[[repl]] = captures.simulated      
      }else{
        PPcaptures[[repl]] = -1
      }
    }
    save(PPcaptures, file = paste0("models/predicted_captures_", mname, "_", 
                                   type, "_nrepl_", nrepl, ".RData"))
  }
}
```

#### Step 2d: Evaluating model fit

The script below shows the posterior predictive data generated in the previous step, the original simulated data, and the corresponding priors. Note that in space and time, the posterior predictive data (red color) represents the original data (black color) accurately.

```
showTrueValues = TRUE
showPrior      = TRUE
showPosterior  = TRUE

mname   = "simulated_ICP_0"
thin    = 2
samples = 500
nChains = 5
nrepl   = 1000

load(paste0(paste0("models/fitted_model_", mname, "_samples_", samples, "_thin_", thin,
                   "_nChains_", nChains, ".RData")))

sel = 1:m$ns

captures = m[["captures"]]

if(showPrior){
  load(file = paste0("models/predicted_captures_", mname, "_prior_nrepl_", nrepl, ".RData"))
  prior.captures = PPcaptures
}

if(showPosterior){
  load(file = paste0("models/predicted_captures_", mname, "_posterior_nrepl_", nrepl, ".RData"))
  posterior.captures = PPcaptures
}

if(showTrueValues){
  load(file = paste0("models/predicted_captures_", mname, "_true_nrepl_", nrepl, ".RData"))
  true.captures = PPcaptures
}

#pdf(paste0("results/model_fit_", mname, ".pdf"))

par(mfrow = c(2, 2))

for(k in 1:4){
  if(k == 1){
    fun  = function(m,captures) Jsmm::number_of_captures(m, captures)
    xlab = ""
    ylab = "total number of captures"
  }
  
  if(k == 2){
    fun  = function(m,captures) Jsmm::mean_captures(m, captures)
    xlab = ""
    ylab = "mean number of captures per individual"
  }
  
  if(k == 3){
    fun  = function(m, captures) Jsmm::mean_time_to_capture(m, captures)
    xlab = ""
    ylab = "mean time to capture"
  }
  
  if(k == 4){
    fun = function(m, captures) Jsmm::mean_distance_between_captures(m, captures)
    xlab = ""
    ylab = "mean distance between captures"
  }
    
  obs = fun(m, captures)
  ma = max(obs, na.rm = T)
  
  if(showPrior){
    prior.pred = matrix(ncol = length(obs), nrow = nrepl)
    for(i in 1:nrepl) prior.pred[i, ] = fun(m,prior.captures[[i]])
    ma = max(ma, max(prior.pred, na.rm = TRUE))
  }
  
  if(showPosterior){
    posterior.pred = matrix(ncol = length(obs), nrow = nrepl)
    for(i in 1:nrepl) posterior.pred[i, ] = fun(m,posterior.captures[[i]])
    ma = max(ma, max(posterior.pred, na.rm = TRUE), na.rm = TRUE)
  }
  
  if(showTrueValues){
    true.pred = matrix(ncol = length(obs), nrow = nrepl)
    for(i in 1:nrepl) true.pred[i, ] = fun(m, true.captures[[i]])
    ma = max(ma, max(true.pred, na.rm = TRUE))
  }
  
  plot(obs[sel], ylim = c(0, ma), xlim = c(0.5, m$ns + 0.5), xlab = xlab, ylab = ylab, 
       pch = 16, col = "black", cex = 1,
        xaxt = "n")
  axis(1, at = 1:m$ns, label = m$sp_names[sel], las = 2, cex.axis = 0.7)
  
  if(showTrueValues){
    qu = apply(true.pred, MARGIN = 2, FUN = quantile, prob = c(0.025, 0.975), na.rm = TRUE)
    for(i in 1:length(obs)) lines(x = c(i, i), y = qu[, sel[i]], col = "black")
  }
  
  if(showPrior){
    qu = apply(prior.pred, MARGIN = 2, FUN = quantile, prob = c(0.025, 0.975), na.rm = TRUE)
    me = apply(prior.pred, MARGIN = 2, FUN = median, na.rm = TRUE)
    points(x = (1:length(me)) - 0.25, y = me[sel], pch = 16, col = "grey")
    
    for(i in 1:length(obs)) lines(x = c(i, i) - 0.25, y = qu[, sel[i]], col = "grey")
  }
  
  if(showPosterior){
    qu = apply(posterior.pred, MARGIN = 2, FUN = quantile, prob = c(0.025, 0.975), na.rm = TRUE)
    me = apply(posterior.pred, MARGIN = 2, FUN = median, na.rm = TRUE)
    points(x = (1:length(me)) + 0.25, y = me[sel], pch = 16, col = "red")
    
    for(i in 1:length(obs)) lines(x = c(i, i) + 0.25, y = qu[, sel[i]], col = "red")
  }
}
```

Figure 6: Posterior predictive data generated with 1000 datasets. Black, red, and grey colors represent the datasets generated with true, posterior, and prior parameters values.

```
#dev.off()
```

#### Step 3a: Show Posteriors vs priors

The script below provides a visualization of the posterior credible intervals for each model parameter (pink color) and its corresponding prior (grey color). See Fig. 6. As before, results are saved in PDF format.

```
showTrueValues = TRUE
mname = "simulated_ICP_0"

thin     = 2
samples  = 500
nChains  = 5
filename = paste0(paste0("models/fitted_model_", mname, "_samples_", samples, "_thin_", thin,
                         "_nChains_", nChains, ".RData"))
load(filename)

sel   = 1:m$ns
post  = Jsmm::poolMcmcChains(m[["postList"]])
npost = length(post)
vars  = names(m[["nc"]][m[["nc"]] > 0])
ns    = m[["ns"]]
nt    = m[["nt"]]
  
nprior = npost
prior  = list()

for(repl in 1:nprior){
  prior[[repl]] = Jsmm::simulate_from_prior(m)
}
  
load("parameters/true_parameters_ICP_0.RData")
  
#pdf(paste0("results/posteriors_vs_priors_", mname, ".pdf"))

par(mfrow = c(3, 2))
cc = 0
for(va in vars){
  nv = m[["nc"]][[va]]
  for(i in 1:nv){
    cc = cc + 1
      
    re.pri = matrix(ncol = ns, nrow = nprior)
    re.pri.G = matrix(ncol = nt, nrow = nprior)
    
    for(repl in 1:nprior){
      re.pri[repl, ]   = prior[[repl]][["Beta"]][[va]][, i]
      re.pri.G[repl, ] = prior[[repl]][["Gamma"]][, cc]
    }
    
    re.post   = matrix(ncol = ns, nrow = npost)
    re.post.G = matrix(ncol = nt, nrow = nprior)
    
    for(repl in 1:npost){
      re.post[repl, ]   = post[[repl]][["Beta"]][[va]][, i]
      re.post.G[repl, ] = post[[repl]][["Gamma"]][, cc]
    }
    
    ymin = min(quantile(re.pri, probs = 0.05), min(re.post))
    ymax = max(quantile(re.pri, probs = 0.95), max(re.post))
    
    if(showTrueValues){
      ymin = min(ymin, pars$Beta[[va]][, i])
      ymax = max(ymax, pars$Beta[[va]][, i])
    }
    
    colnames(re.pri) = m$sp_names
    re.pri  = re.pri[, sel]
    re.post = re.post[, sel]
    
    boxplot(re.pri, outline = FALSE, at = 3*(1:ns)-0.75, ylim = c(ymin, ymax),
            main = m[["parNames"]][cc], las = 2, cex.axis = 0.7,
            xlim = c(2, 3*ns + 2),
            ylab = "parameter value", col = "grey", xaxt = "n")
    
    axis(1, at = 3*(1:m$ns), label = colnames(re.pri), las = 2, cex.axis = 0.7)
    
    boxplot(re.post, outline = FALSE, at = 3*(1:ns) + 0.75, col = "pink", add = TRUE, 
            xaxt = "n", yaxt = "n")
    
    abline(h = 0)
    
    if(showTrueValues){
      for(j in 1:m[["ns"]]){
        lines(x = c(3*j, 3*j + 1.5), y = c(pars$Beta[[va]][j, i], pars$Beta[[va]][j, i]), 
              lwd = 2, col = "red")
      }
    }
    
    ymin = min(quantile(re.pri.G, probs = 0.05), min(re.post.G))
    ymax = max(quantile(re.pri.G, probs = 0.95), max(re.post.G))
    
    if(showTrueValues){
      ymin = min(ymin, pars$Gamma[, cc])
      ymax = max(ymax, pars$Gamma[, cc])
    }
    
    colnames(re.pri.G) = colnames(m$X[["trait"]])
    
    boxplot(re.pri.G, outline = FALSE, at = 3*(1:nt), ylim = c(ymin, ymax),
            main = m[["parNames"]][cc], las = 2, cex.axis = 0.7,
            xlim = c(2, 3*nt + 2),
            ylab = "parameter value", col = "grey")
    
    boxplot(re.post.G, outline = FALSE, at = 3*(1:nt) + 1.5, col = "pink", add = TRUE, 
            xaxt = "n", yaxt = "n")
    
    abline(h = 0)
    
    if(showTrueValues){
      for(j in 1:m[["nt"]]){
        lines(x = c(3*j, 3*j + 1.5), y = c(pars$Gamma[j, cc], pars$Gamma[j, cc]), 
              lwd = 2, col = "red")
      }
    }
  }
}
```

Figure 7: Posterior credible intervals for the model parameters. Left panels show the posteriors for each of the model parameters (pink color), their corresponding priors (Grey), and the parameter true value (Red). The right panels show the corresponding values for the Gamma associated parameter.

```
#dev.off()
```

#### Step 3b: Simulate movement trajectories

The function **`simulate_trajectories()`** simulates the movement trajectories and adds them in matrix form to the named list **`m`**. The matrix **`m$trajectories`** has a number of rows equivalent to the time partitions, and each column represents a unique individual.

```
method = "ICP"

load(paste0("parameters/true_parameters_", method,"_0", ".RData"))
load(paste0("models/unfitted_model_simulated_", method,"_0", "_no_data.RData"))

ni = round(6/m[["ns"]]) #number of individuals per species
releases = matrix(NA, ncol = 2, nrow = ni*m[["ns"]])
releases[, 1] = rep(1:m[["ns"]], ni)
releases[, 2] = rep(1, ni*m[["ns"]])
releases[, 2] = sample(seq_len(length(m$observation_effort$releases$location)),
                       size = ni*m[["ns"]], replace = TRUE)

colnames(releases) = c("sp", "release_event")
  
secondary_release = TRUE
max_dt = NULL

head(releases)
```

```
##      sp release_event
## [1,]  1             5
## [2,]  2            10
## [3,]  1             7
## [4,]  2             4
## [5,]  1            10
## [6,]  2             8
```

```
secondary_release = TRUE
max_dt = NULL

m = Jsmm::simulate_trajectories(m = m, pars = pars, releases = releases,
                          max_dt = max_dt)

m$trajectories
```

```
##       [,1] [,2] [,3] [,4] [,5] [,6]
##  [1,]   NA   NA   NA   NA   NA   NA
##  [2,]   NA   NA   NA   NA   NA   NA
##  [3,]   NA   NA   NA   NA   NA   NA
##  [4,]   NA   NA   NA  136   NA   NA
##  [5,]   87   NA   NA   89   NA   NA
##  [6,]   73   NA   NA  121   NA   NA
##  [7,]   64   NA  121  133   NA   NA
##  [8,]  119   NA   85  133   NA  104
##  [9,]  109   NA   85  124   NA   53
## [10,]  144   92   86  112   77   92
## [11,]   NA   NA   NA   NA   NA   NA
```

```
dim(m$trajectories)
```

```
## [1] 11  6
```

The simulated trajectories can be visualized by the function **`plot_trajectories()`**.

```
# optional parameters for customizing plot:
  customize_plot = list()
  customize_plot$color_map = "#EEEEEE"
  customize_plot$labs_axis = c("X", "Y")
  customize_plot$lab_legend =  "Matrix"
  customize_plot$name_legend = "Habitat"
  customize_plot$color_polyb = "darkgrey"
  customize_plot$lwd_poly = 0.01
  customize_plot$site_col = "red"
  customize_plot$siteb_col = "darkred"
  customize_plot$sites_text_pos = c(-0.0, -0.0)
  customize_plot$site_text_size = 5
  customize_plot$panelb_color = "black"
  customize_plot$panelb_lwd = 1
  customize_plot$trajectory_color = c("black","blue") #rep("black", m[["ns"]])
  customize_plot$trajectory_lwd = 1
  plot_sp = 1:2

  Jsmm::plot_trajectories(m = m, customize_plot = customize_plot, plot_sp = plot_sp  )
```

Figure 8: Simulated movement trajectories for three animals from each species. Black and blue colors correspond to the species 1 and 2, respectively.

### Demo simulated example ICP (covariates)

#### Example Description

Let’s suppose that a researcher is interested in exploring whether the air temperature, altitude, and the species’ body size influence the movement activity and vital dynamics for twenty focal species. In order to do this, an experiment is conducted over a landscape by setting nine trapping devices for eleven days, as shown in Fig. 1. Initially, 500 animals of each species are marked and released from different locations at different times. All capture sites are visited every day, starting from the second day until the end of the experiment. While the capture sites are visited, animals are marked, and captures are recorded. Secondary releases are not possible for this experiment.

For detailed information about the functions and their corresponding outcomes involved in our analyses, please refer to Appendix S1. Main paper.

#### Step 0: Loading the package

We start by loading the Jsmm package.

```
library(Jsmm)
```

#### Step 1a: Define model

For this example, we will use the same domain and observation effort lists as in the previous example.

##### Set covariate and phylogenetic relationships data

The Jsmm framework allows the inclusion of spatial, temporal, spatio-temporal covariates, and trait covariates. To do this, first, we are required to provide the data in a suitable format. The covariates are set within the named lists **`cov_data`** and **`tr_data`**.

For this example, we will include the spatial covariate altitude, the temporal covariate air temperature, a covariate for the observation process, and the body size as a species trait. Tables 5-7. show the model covariates characterizations.

```
cov_data = list()
altitude = domain[["triangulation"]][["node"]][, 2]
cov_data = list()
cov_data[["n_data"]] = data.frame(altitude)

head(cov_data[["n_data"]])
```

```
##    altitude
## 1 1.0000000
## 2 0.9333333
## 3 0.8666667
## 4 0.8000000
## 5 0.7333333
## 6 0.6666667
```

Figure 9: Altitude covariate. Measurements recorded at each node of the domain.

Covariates for the movement parameters.

| Covariate name | Spatial | Temporal | Spatio-temporal | Type |
| --- | --- | --- | --- | --- |
| altitude | node | No | No | Numeric |
| air\_temperature | No | Yes | No | Numeric |

Covariates for the observation parameter.

| Covariate name | Event | Time variation within events | Type |
| --- | --- | --- | --- |
| capture\_intensity | Yes | No | Factor |

Covariates for the species traits

| Covariate name | Type |
| --- | --- |
| size | Numeric |

The spatial data could be expressed as element or node resolution within the data frames **`e_data`** and **`n_data`**, respectively. The altitude covariate shown in Fig. 8., with values at each node, is set within the list **`cov_data$n_data`**.

The temporal data is set within the data frame **`t_data`**, and the times at which the observations were held are saved within the vector **`t_data_time`**. The covariate for the air temperature contains data for each day of the experiment. It is assumed that during the first six days of the experiment, cold temperatures occur, and during the rest of the days, the temperature increases. (See Fig. 9.).

```
t_data_times = seq(0, study_duration)
air_temperature = -1 + 2*(t_data_times > study_duration/2)

cov_data[["t_data"]] = data.frame(air_temperature)
cov_data[["t_data_times"]] = t_data_times 
  
cov_data[["t_data"]]
```

```
##    air_temperature
## 1               -1
## 2               -1
## 3               -1
## 4               -1
## 5               -1
## 6               -1
## 7                1
## 8                1
## 9                1
## 10               1
## 11               1
```

```
cov_data[["t_data_times"]]
```

```
##  [1]  0  1  2  3  4  5  6  7  8  9 10
```

Figure 10: Air temperature covariate. Measurements recorded each day of the experiment (red dots).

Within the Jsmm, the covariates for the capture process are included in the data frame **`obs_data`** when there are no time variations within events, the rows correspond to the capture events, and the number of columns to the number of covariates. The covariate that characterizes capture events as high or low intensity is set as:

```
nevents = length(observation_effort[["captures"]][["location"]])

capture_intensity = factor(sample(x = c("low","high"), size = nevents,
                                  replace = TRUE), levels = c("low", "high"))

cov_data[["obs_data"]] = data.frame(capture_intensity)

head(cov_data[["obs_data"]])
```

```
##   capture_intensity
## 1               low
## 2              high
## 3               low
## 4              high
## 5               low
## 6               low
```

The covariate information for the capture event c = 10 can be retrieved as

```
c = 10
cov_data$obs_data[c, ]
```

```
## [1] low
## Levels: low high
```

In our framework, the trait information and phylogenetic relationships are stored in the data frame called **`tr_data`**, and matrix **`C`** (See Fig. 10.). For **`tr_data`**, the number of rows and columns corresponds to the number of species and the number of traits, respectively. The phylogenetic relationships matrix has dimensions number of species \(\times\) number of species. If **`C`** and **`tr_data`** are not provided, by default the model sets the identity matrix \(I\_{ns}\), and intercept, respectively. For our example, we want to include information about the body size trait for the focal species.

```
ns = 20

sp_names = paste0("sp_", 1:ns)

tr_data = data.frame(size = rnorm(ns))
head(tr_data)
```

```
##         size
## 1 -0.2488135
## 2 -0.4886277
## 3 -1.4161455
## 4 -1.0260879
## 5  0.5308912
## 6  0.4276941
```

```
order = rep(c(1, 1, 2, 2), ceiling(ns/4))[1:ns]
genus = rep(c(1, 2, 3, 4), ceiling(ns/4))[1:ns]

C = matrix(0, nrow = ns, ncol = ns)
colnames(C) = sp_names
rownames(C) = sp_names

for(i in 1:ns){
  for(j in 1:ns){
    if(order[i] == order[j]) C[i, j] = C[i, j] + 0.45
    if(genus[i] == genus[j]) C[i, j] = C[i, j] + 0.45
    if(i == j) C[i, j] = 1
  }
}
```

Figure 11: Phylogenetic correlation matrix.

##### Setting the data model

Before, we explored how to set the different covariate data within the software. In this section, we are going to set the named list **`model_formula`** to declare explicitly the relationships between the covariates, and the movement and observation parameters within the diffusion-advection-reaction model on which our framework is based. Additionally, we will set the traits and possible relationships with trait data. See Appendix S1. Main paper for a detailed description.

The **`model_formula`** list is composed of (some combination of) the objects **`diffusion`**, **`advection_1`** (advection in x), **`advection_2`** (advection in y), **`mortality`**, **`habitat_preference`**, **`observation`**, and **`traits`**.

For declaring the formulas, the ~ symbol is used. If the model does not include a given parameter, this should be declared as **`NULL`**. If the parameter is included but no covariates are declared, this should be set as **`~1`**, this applies to all parameters except for the habitat preference parameter **`habitat_preference`**. This parameter is declared in the model formula only when covariates are included. By default, the intercept is included in all the formulas except in the **`habitat_preference`**. It is a requirement that for all formulas that include covariates, the name declarations in the model formula match exactly with the covariate data, and that the covariates for the observation process are declared separately. This is because the temporal resolution within the observation corresponds to capture events or time variations within capture events.

For our example, the diffusion is expressed as a function of the air temperature, and the mortality is expressed as a function of the altitude. We also include the species body size trait. Table 8. shows the formulas declarations for this model. This model formula is set as:

```
model_formula = list()
model_formula[["diffusion"]]          = ~ air_temperature
model_formula[["advection_1"]]        = NULL
model_formula[["advection_2"]]        = NULL
model_formula[["mortality"]]          = ~ altitude
model_formula[["habitat_preference"]] = NULL
model_formula[["traits"]]             = ~ size
model_formula[["observation"]]        = ~ capture_intensity
```

Model Formulas declarations for the simulated example.

| Model parameter | Model formula |
| --- | --- |
| **`diffusion`** | ~ air\_temperature |
| **`advection_1`** | - |
| **`advection_2`** | - |
| **`habitat_preference`** | - |
| **`mortality`** | ~ altitude |
| **`observation`** | ~ capture\_intensity |
| **`traits`** | ~ size |

##### Setting captures and releases data (optional)

At this stage, for our simulated example, we do not have capture and release data. For models for which this information is available, the data should be included on the named list **`data`** with sublists **`data$releases`** and **`data$captures`**. In the next section, more explanation will be provided about the releases and the capture data structure. For illustration, we declare the empty list **`data`** here as:

```
  data = list()
  data[["captures"]] = NULL
  data[["releases"]] = NULL
```

##### Define the model object **`m`**

The model object **`m`** is a named list that synthesizes all the necessary information for running the analyses by utilizing all the elements created in previous steps. To define the model **`m`**, some actions take place within the function **`jsmm(.)`**. First, a partition of time is set, taking into account all the times in which the data and experiment occur. Second, if present, the covariate data is combined into two objects, one related to covariates for the movement model, and the other one related to covariates used for the capture process. Third, design matrices are constructed by using the model (**`model_formula`**) declaration, movement, observation, and trait data. Fourth, some elements used for the underlying numerical method are created, indicators for the presence of covariates, and the relationship between the nodes and the capture/release locations. See Appendix S1. of the main manuscript for detailed information.

The model **`m`** for the Instantaneous Capture Process (ICP) is set as

```
  m = Jsmm::jsmm(domain = domain, observation_effort = observation_effort,
           releases = data[["releases"]], captures = data[["captures"]], 
           cov_data = cov_data, 
           ns = ns, sp_names = sp_names, tr_data = tr_data, C = C,
           model_formula = model_formula)
```

```
# Here we are saving the model **`m`** just created.

save(m, file = paste0("models/unfitted_model_simulated_", 
                      method, "_no_data.RData"))
```

#### Step 1b: Simulate model parameters from priors

This optional step allows generating the model parameters from predefined priors through the function **`simulate_from_prior(m)`** (See Appendix S1. and Fig. 1. main manuscript). The script below sets the diffusion parameter coefficients \(\beta^a\_0\) and \(\beta^a\_1\) increasing with the body size and air temperature, respectively; the mortality increases with altitude; and the observation rate is higher with high intensity.

First, we load the model created in the previous step.

```
method = "ICP"

file = paste0("models/unfitted_model_simulated_", method, "_no_data.RData")
  
load(file)
```

```
muZ = m[["prior"]][["muZ"]]
nc = m[["nc"]]
np = sum(nc)
nt = m[["nt"]]
idx = 1

for(j in 1:nt){
  for(i in 1:length(nc)){
    for(ii in seq_len(nc[i])){
      
      # baseline diffusion
      if(names(nc)[i] == "diffusion" & ii == 1 & j == 1) muZ[idx] = -4
      
      # diffusion increases with air temperature
      if(names(nc)[i] == "diffusion" & ii == 2 & j == 1) muZ[idx] = 1 
        
      # diffusion intercept increases with body size
      if(names(nc)[i] == "diffusion" & ii == 1 & j == 2) muZ[idx] = 1 
      
      # baseline mortality
      if(names(nc)[i] == "mortality" & ii == 1 & j == 1) muZ[idx] = -2 
      
      # mortality increases with altitude
      if(names(nc)[i] == "mortality" & ii == 2 & j == 1) muZ[idx] = 1 
      
      # baseline observation rate
      if(names(nc)[i] == "observation" & ii == 1 & j == 1) muZ[idx] = -2 
        
      # observation rate higher with high intensity
      if(names(nc)[i] == "observation" & ii == 2 & j == 1) muZ[idx] = 1 
      
      idx = idx + 1
    }
  }
}

m[["prior"]][["muZ"]] = muZ
m[["prior"]][["VZ"]] = 0*m[["prior"]][["VZ"]]
m[["prior"]][["rho"]] = matrix(c(0.75, 1), nrow = 1)

pars = Jsmm::simulate_from_prior(m)
```

```
save(pars, file = paste0("parameters/true_parameters_", method,".RData"))
```

The model parameters are set up within the named list **`pars$Beta`**. Within each element of this list, there are matrices for each of the movement and observation parameters, that is, **`pars$Beta$diffusion`**, **`pars$Beta$mortality`**, and **`pars$Beta$observation`**. For example, for diffusion, a matrix with twenty rows and two columns is created. Rows correspond to the species, and the first and second columns correspond to the coefficients \(\beta^a\_0\), and \(\beta^a\_1\).

```
head(pars$Beta$diffusion)
```

```
##           [,1]     [,2]
## [1,] -3.871749 1.313660
## [2,] -3.846895 1.211122
## [3,] -5.295459 1.000561
## [4,] -5.261582 1.289043
## [5,] -3.047360 1.597476
## [6,] -3.325745 1.548127
```

Figure 12: Diffusion intercept is influenced by the species body size.

Table 3: Model coeficients for diffusion, mortality and observation parameters for the twenty species.

| species | betaa0 | betaa1 | betam0 | betam1 | betao0 | betao1 |
| --- | --- | --- | --- | --- | --- | --- |
| 1 | -3.872 | 1.314 | -2.378 | 1.124 | -2.205 | 0.846 |
| 2 | -3.847 | 1.211 | -2.645 | 1.297 | -2.359 | 0.532 |
| 3 | -5.295 | 1.001 | -3.033 | 1.234 | -1.480 | 0.496 |
| 4 | -5.262 | 1.289 | -2.162 | 1.308 | -1.480 | 0.542 |
| 5 | -3.047 | 1.597 | -2.725 | 1.195 | -2.682 | 0.963 |
| 6 | -3.326 | 1.548 | -2.127 | 1.051 | -2.497 | 1.423 |
| 7 | -4.307 | 1.056 | -3.017 | 1.658 | -1.004 | 0.467 |
| 8 | -3.211 | 1.526 | -2.442 | 1.177 | -1.645 | 0.307 |
| 9 | -4.567 | 1.721 | -2.571 | 1.330 | -2.281 | 0.769 |
| 10 | -3.665 | 1.511 | -2.104 | 1.141 | -2.805 | 1.319 |
| 11 | -4.068 | 0.994 | -3.108 | 1.331 | -1.498 | 0.861 |
| 12 | -4.515 | 1.491 | -2.439 | 1.616 | -1.470 | 0.543 |
| 13 | -3.727 | 1.119 | -2.335 | 0.963 | -2.239 | 0.507 |
| 14 | -4.497 | 1.451 | -2.114 | 0.799 | -2.385 | 1.390 |
| 15 | -4.225 | 1.272 | -2.776 | 1.322 | -1.353 | 0.264 |
| 16 | -5.276 | 1.986 | -2.446 | 1.210 | -1.595 | 0.724 |
| 17 | -3.598 | 0.989 | -2.540 | 0.866 | -2.154 | 0.774 |
| 18 | -4.367 | 1.880 | -2.201 | 0.711 | -2.641 | 1.144 |
| 19 | -2.879 | 0.995 | -2.664 | 1.410 | -1.736 | 0.379 |
| 20 | -4.596 | 1.729 | -2.399 | 1.355 | -1.727 | 0.743 |

Figure 13: Model parameters for each of the twenty species. Upper to lower figures show diffusion, mortality, and observation dependence on the predictors; air temperature, altitude, and capture intensity, respectivelly.

#### Step 1c: Simulate capture data

For this example, we simulated 500 individuals for each species. We also declared that secondary releases are not possible.

```
ni = round(10000/m[["ns"]]) #number of individuals per species

releases = matrix(NA, ncol = 2, nrow = ni*m[["ns"]])

releases[, 1] = rep(1:m[["ns"]], ni)
releases[, 2] = rep(1, ni*m[["ns"]])
releases[, 2] = sample(seq_len(length(m$observation_effort$releases$location)),
                       size = ni*m[["ns"]], replace = TRUE)

colnames(releases) = c("sp", "release_event")
  
secondary_release = FALSE
max_dt = NULL

head(releases)
```

```
##      sp release_event
## [1,]  1             5
## [2,]  2            10
## [3,]  3             7
## [4,]  4             4
## [5,]  5            10
## [6,]  6             8
```

For example, the second row of the releases matrix indicates that one individual from species 2 was released at the release site number four at \(t = 9\). (release event was declared on the **`observation_effort`** list).

As before, we can proceed to simulate capture-recapture data by using the function **`simulate_captures(.)`**.

```
m = Jsmm::simulate_captures(m = m, pars = pars, releases = releases,
                      max_dt = max_dt, secondary_release = secondary_release)
```

```
save(m, file = paste0("models/unfitted_model_simulated_", method, ".RData"))
```

#### Step 2a: Fit the model

The parameter estimation within the Jsmm is made via Markov Chain Monte Carlo (MCMC) through the function **`sampleMCMC(.)`**. The user should choose reasonable parameters for the sampler to ensure that the MCMC has good mixing, low autocorrelation, and reliable convergence. For this simple example, we will pretend that we do not know the model parameters, and we will proceed to estimate those. The parameters used in our sampler are set in Table 11.

Model Formulas declarations for the simulated example.

| MCMC sampler parameter | Value |
| --- | --- |
| thin | 2 |
| samples | 250 |
| transcient | 250 |
| number of chains | 5 |

```
mname = "simulated_ICP"
load(paste0("models/unfitted_model_", mname, ".RData"))
  
m         = merge_rc_data(m)
thin      = 2
samples   = 250
transient = round(0.5*samples*thin)
nChains   = 5
init_pars = NULL

for(i in 1:nChains){
  set.seed(i)
  m = Jsmm::sampleMcmc(m = m, samples = samples, transient = transient, thin = thin, 
                 nChains = 1, init_pars = init_pars)
  
  save(m, file = paste0(paste0("models/", i, "_fitted_model_", mname, "_samples_", samples, 
                               "_thin_", thin, "_nChains_", nChains, ".RData")))
}

postList = list()
  
for(i in 1:nChains){
    fname = paste0(paste0("models/", i, "_fitted_model_", mname, "_samples_", samples, 
                          "_thin_", thin, "_nChains_", nChains, ".RData"))
    load(file = fname)
    postList[[i]] = m[["postList"]][[1]]
    file.remove(fname)
}
  
m[["postList"]] = postList
  
save(m, file = paste0(paste0("models/fitted_model_", mname, "_samples_", samples, 
                             "_thin_", thin, "_nChains_", nChains, ".RData")))
```

#### Step 2b: Evaluating MCMC convergence

After fitting the model, we will assess the MCMC convergence. First, we convert the generated chains to objects of the MCMC class. Here this is done through the function **`convert_to_coda_object(.)`**, this function is based on the R-package Coda (Plummer et al. (2006)). Note that the Gelman-Rubin statistic (R-hat) diagnosis of convergence shows that the chains converge satisfactorily. The Figures 14-20. Show the trace plots for our model for the first species.

```
mname = "simulated_ICP"

thin    = 2
samples = 250
nChains = 5

load(paste0(paste0("models/fitted_model_", mname, 
                   "_samples_", samples, "_thin_", thin, "_nChains_", nChains, ".RData")))

mpost = Jsmm::convert_to_coda_object(m)

summary(coda::gelman.diag(mpost$Beta, multivariate = FALSE)$psrf)
```

```
##    Point est.       Upper C.I.    
##  Min.   : 1.009   Min.   : 1.020  
##  1st Qu.: 1.028   1st Qu.: 1.070  
##  Median : 1.097   Median : 1.242  
##  Mean   : 1.773   Mean   : 2.726  
##  3rd Qu.: 1.263   3rd Qu.: 1.627  
##  Max.   :18.410   Max.   :39.043
```

Figure 14: First species trace plots model parameters Beta., five MCMC chains.

Figure 15: First species trace plots model parameters Beta., five MCMC chains.

Figure 16: First species trace plots, five MCMC chains Gamma.

Figure 17: First species trace plots, five MCMC chains Gamma.

Figure 18: First species trace plots, five MCMC chains Gamma.

Figure 19: First species trace plots, five MCMC chains Gamma.

Figure 20: First species trace plots, five MCMC chains Rho.

#### Step 2c: Simulate prior and posterior predictive data

The script below generates 1000 datasets. The main purpose of generating predictive data is to explore how the fitted model behaves in time and space compared to the original data. This method helps to infer if the fitted parameters provide a good approximation of the real data.

```
mname = "simulated_ICP"

for(nrepl in c(1000)){
  showTrueValues = TRUE
  
  thin    = 2
  samples = 250
  nChains = 5
  
  load(paste0(paste0("models/fitted_model_", mname, "_samples_", samples, "_thin_", thin, 
                     "_nChains_", nChains, ".RData")))
  
  post = m[["postList"]]
  
  if(showTrueValues){
    types = c("true", "prior", "posterior")
    load("parameters/true_parameters_ICP.RData")
    true_pars = pars
  }else{
    types =  c("prior", "posterior")
  }

  for(type in types){
    PPcaptures = list()

    for(repl in 1:nrepl){
      if(type == "true"){
        pars = true_pars
      }
      
      if(type == "posterior"){
        chain = sample(1:length(post), size = 1)
        sa    = sample(1:length(post[[chain]]), size = 1)
        pars  = post[[chain]][[sa]]
      }
      
      if(type == "prior"){
        pars = Jsmm::simulate_from_prior(m)
      }
      
      max_dt = NULL
      secondary_release = FALSE
      releases = m[["releases"]]
      
      m2 = Jsmm::simulate_captures(m = m, pars = pars, releases = releases,
                               max_dt = max_dt, secondary_release = secondary_release)
        
      captures.simulated = m2[["captures"]]
        
      if(nrow(captures.simulated) > 0){
        PPcaptures[[repl]] = captures.simulated      
      }else{
        PPcaptures[[repl]] = -1
      }
    }
    save(PPcaptures, file = paste0("models/predicted_captures_", mname, "_", 
                                   type, "_nrepl_", nrepl, ".RData"))
  }
}
```

#### Step 2d: Evaluating model fit

The script below provides a visualization of the posterior predictive data generated on the previous step compared with the original simulated dataset and priors.

```
showTrueValues = TRUE
showPrior      = TRUE
showPosterior  = TRUE

mname   = "simulated_ICP"
thin    = 2
samples = 250
nChains = 5
nrepl   = 1000

load(paste0(paste0("models/fitted_model_", mname, "_samples_", samples, "_thin_", thin,
                   "_nChains_", nChains, ".RData")))

sel = 1:m$ns

captures = m[["captures"]]

if(showPrior){
  load(file = paste0("models/predicted_captures_", mname, "_prior_nrepl_", nrepl, ".RData"))
  prior.captures = PPcaptures
}

if(showPosterior){
  load(file = paste0("models/predicted_captures_", mname, "_posterior_nrepl_", nrepl, ".RData"))
  posterior.captures = PPcaptures
}

if(showTrueValues){
  load(file = paste0("models/predicted_captures_", mname, "_true_nrepl_", nrepl, ".RData"))
  true.captures = PPcaptures
}

#pdf(paste0("results/model_fit_", mname, ".pdf"))

par(mfrow = c(2, 2))

for(k in 1:4){
  if(k == 1){
    fun  = function(m,captures) Jsmm::number_of_captures(m, captures)
    xlab = ""
    ylab = "total number of captures"
  }
  
  if(k == 2){
    fun  = function(m,captures) Jsmm::mean_captures(m, captures)
    xlab = ""
    ylab = "mean number of captures per individual"
  }
  
  if(k == 3){
    fun  = function(m, captures) Jsmm::mean_time_to_capture(m, captures)
    xlab = ""
    ylab = "mean time to capture"
  }
  
  if(k == 4){
    fun = function(m, captures) Jsmm::mean_distance_between_captures(m, captures)
    xlab = ""
    ylab = "mean distance between captures"
  }
    
  obs = fun(m, captures)
  ma = max(obs, na.rm = T)
  
  if(showPrior){
    prior.pred = matrix(ncol = length(obs), nrow = nrepl)
    for(i in 1:nrepl) prior.pred[i, ] = fun(m,prior.captures[[i]])
    ma = max(ma, max(prior.pred, na.rm = TRUE))
  }
  
  if(showPosterior){
    posterior.pred = matrix(ncol = length(obs), nrow = nrepl)
    for(i in 1:nrepl) posterior.pred[i, ] = fun(m,posterior.captures[[i]])
    ma = max(ma, max(posterior.pred, na.rm = TRUE), na.rm = TRUE)
  }
  
  if(showTrueValues){
    true.pred = matrix(ncol = length(obs), nrow = nrepl)
    for(i in 1:nrepl) true.pred[i, ] = fun(m, true.captures[[i]])
    ma = max(ma, max(true.pred, na.rm = TRUE))
  }
  
  plot(obs[sel], ylim = c(0, ma), xlim = c(0.5, m$ns + 0.5), xlab = xlab, ylab = ylab, 
       pch = 16, col = "black", cex = 1,
        xaxt = "n")
  axis(1, at = 1:m$ns, label = m$sp_names[sel], las = 2, cex.axis = 0.7)
  
  if(showTrueValues){
    qu = apply(true.pred, MARGIN = 2, FUN = quantile, prob = c(0.025, 0.975), na.rm = TRUE)
    for(i in 1:length(obs)) lines(x = c(i, i), y = qu[, sel[i]], col = "black")
  }
  
  if(showPrior){
    qu = apply(prior.pred, MARGIN = 2, FUN = quantile, prob = c(0.025, 0.975), na.rm = TRUE)
    me = apply(prior.pred, MARGIN = 2, FUN = median, na.rm = TRUE)
    points(x = (1:length(me)) - 0.25, y = me[sel], pch = 16, col = "grey")
    
    for(i in 1:length(obs)) lines(x = c(i, i) - 0.25, y = qu[, sel[i]], col = "grey")
  }
  
  if(showPosterior){
    qu = apply(posterior.pred, MARGIN = 2, FUN = quantile, prob = c(0.025, 0.975), na.rm = TRUE)
    me = apply(posterior.pred, MARGIN = 2, FUN = median, na.rm = TRUE)
    points(x = (1:length(me)) + 0.25, y = me[sel], pch = 16, col = "red")
    
    for(i in 1:length(obs)) lines(x = c(i, i) + 0.25, y = qu[, sel[i]], col = "red")
  }
}
```

Figure 21: Posterior predictive data generated with 1000 datasets. Black, red, and grey colors represent the datasets generated with true, posterior, and prior parameters values.

```
#dev.off()
```

#### Step 3a: Show Posteriors vs priors

The script below provides a visualization of the posterior credible intervals for each model parameter (pink color) and its corresponding prior (grey color). See Fig. 21-22.

```
showTrueValues = TRUE
mname = "simulated_ICP"

thin     = 2
samples  = 250
nChains  = 5
filename = paste0(paste0("models/fitted_model_", mname, "_samples_", samples, "_thin_", thin,
                         "_nChains_", nChains, ".RData"))
load(filename)

sel   = 1:m$ns
post  = poolMcmcChains(m[["postList"]])
npost = length(post)
vars  = names(m[["nc"]][m[["nc"]] > 0])
ns    = m[["ns"]]
nt    = m[["nt"]]
  
nprior = npost
prior  = list()

for(repl in 1:nprior){
  prior[[repl]] = simulate_from_prior(m)
}
  
load("parameters/true_parameters_ICP.RData")

par(mfrow=c(3,2))

cc = 0
for(va in vars){
  nv = m[["nc"]][[va]]
  for(i in 1:nv){
    cc = cc + 1
      
    re.pri = matrix(ncol = ns, nrow = nprior)
    re.pri.G = matrix(ncol = nt, nrow = nprior)
    
    for(repl in 1:nprior){
      re.pri[repl, ]   = prior[[repl]][["Beta"]][[va]][, i]
      re.pri.G[repl, ] = prior[[repl]][["Gamma"]][, cc]
    }
    
    re.post   = matrix(ncol = ns, nrow = npost)
    re.post.G = matrix(ncol = nt, nrow = nprior)
    
    for(repl in 1:npost){
      re.post[repl, ]   = post[[repl]][["Beta"]][[va]][, i]
      re.post.G[repl, ] = post[[repl]][["Gamma"]][, cc]
    }
    
    ymin = min(quantile(re.pri, probs = 0.05), min(re.post))
    ymax = max(quantile(re.pri, probs = 0.95), max(re.post))
    
    if(showTrueValues){
      ymin = min(ymin, pars$Beta[[va]][, i])
      ymax = max(ymax, pars$Beta[[va]][, i])
    }
    
    colnames(re.pri) = m$sp_names
    re.pri  = re.pri[, sel]
    re.post = re.post[, sel]
    
    boxplot(re.pri, outline = FALSE, at = 3*(1:ns)-0.75, ylim = c(ymin, ymax),
            main = m[["parNames"]][cc], las = 2, cex.axis = 0.7,
            xlim = c(2, 3*ns + 2),
            ylab = "parameter value", col = "grey", xaxt = "n")
    
    axis(1, at = 3*(1:m$ns), label = colnames(re.pri), las = 2, cex.axis = 0.7)
    
    boxplot(re.post, outline = FALSE, at = 3*(1:ns) + 0.75, col = "pink", add = TRUE, 
            xaxt = "n", yaxt = "n")
    
    abline(h = 0)
    
    if(showTrueValues){
      for(j in 1:m[["ns"]]){
        lines(x = c(3*j, 3*j + 1.5), y = c(pars$Beta[[va]][j, i], pars$Beta[[va]][j, i]), 
              lwd = 2, col = "red")
      }
    }
    
    ymin = min(quantile(re.pri.G, probs = 0.05), min(re.post.G))
    ymax = max(quantile(re.pri.G, probs = 0.95), max(re.post.G))
    
    if(showTrueValues){
      ymin = min(ymin, pars$Gamma[, cc])
      ymax = max(ymax, pars$Gamma[, cc])
    }
    
    colnames(re.pri.G) = colnames(m$X[["trait"]])
    
    boxplot(re.pri.G, outline = FALSE, at = 3*(1:nt), ylim = c(ymin, ymax),
            main = m[["parNames"]][cc], las = 2, cex.axis = 0.7,
            xlim = c(2, 3*nt + 2),
            ylab = "parameter value", col = "grey")
    
    boxplot(re.post.G, outline = FALSE, at = 3*(1:nt) + 1.5, col = "pink", add = TRUE, 
            xaxt = "n", yaxt = "n")
    
    abline(h = 0)
    
    if(showTrueValues){
      for(j in 1:m[["nt"]]){
        lines(x = c(3*j, 3*j + 1.5), y = c(pars$Gamma[j, cc], pars$Gamma[j, cc]), 
              lwd = 2, col = "red")
      }
    }
  }
}
```

Figure 22: Posterior credible intervals for the model parameters. Left panels show the posteriors for each of the model parameters (pink color), their corresponding priors (Grey), and the parameter true value (Red). Right panels show the corresponding values for the Gamma associated parameter.

Figure 23: Posterior credible intervals for the model parameters. Left panels show the posteriors for each of the model parameters (pink color), their corresponding priors (Grey), and the parameter true value (Red). Right panels show the corresponding values for the Gamma associated parameter.

Figure 24: Posterior credible intervals for the associated rho parameter. Posterior, prior, and true value are represented by pink, grey, and red colors, respectively.

### Demo simulated example CCP (covariates)

#### Example Description

Let’s suppose that a researcher is interested in exploring whether the air temperature, altitude, and the species’ body size influence the movement activity and vital dynamics for twenty focal species. In order to do this, an experiment is conducted over a landscape by setting nine trapping devices for eleven days, as shown in Fig. 1. Initially, 500 animals of each species are marked and released from different locations at different times. By simplicity, let’s suppose that all the traps are kept open, allowing captures every day, visited by the end of each day, and captures are recorded. Note that within our framework, the user can customize the time intervals in which the traps allow captures. Secondary releases are not possible for this experiment.

For detailed information about the functions and their corresponding outcomes involved in our analyses, please refer to Appendix S1. Main paper.

#### Step 0: Loading the package

We start by loading the Jsmm package.

```
library(Jsmm)
```

#### Step 1a: Define model

We set the same elements as before, except the **`observation_effort`** list. For this third example, the capture process has been set as continuous in the list **`observation_effort$method`**. In addition, **`observation_effort$captures$time`**, which indicates the times in which captures occur, is now set as a matrix with rows representing the capture events, and the first and second columns indicating the time interval \([t\_0, t\_1]\) in which the capture site starts and ends capturing individuals. The observation effort can be visualized in Fig. 25.

The script for setting the **`observation_effort`** object is:

Figure 25: Observation effort scheme for CCP capture process. Times in which releases and captures are possible are represented by blue and red colors, respectively.

For example, capture events 10 to 15 occur at times

```
observation_effort$captures$time[10:15, ]
```

```
##      t_0 t_1
## [1,]   1   2
## [2,]   1   2
## [3,]   1   2
## [4,]   1   2
## [5,]   1   2
## [6,]   1   2
```

Information corresponding to the capture event \(c = 12\), is retrieved as

```
observation_effort$captures$location[12]
```

```
## [1] 3
```

```
observation_effort$captures$time[12, ]
```

```
## t_0 t_1 
##   1   2
```

Read this as, the third capture site allows captures during the time interval \([1, 2]\).

Setting model formulas:

```
model_formula = list()
model_formula[["diffusion"]]          = ~ air_temperature
model_formula[["advection_1"]]        = NULL
model_formula[["advection_2"]]        = NULL
model_formula[["mortality"]]          = ~ altitude
model_formula[["habitat_preference"]] = NULL
model_formula[["traits"]]             = ~ size
model_formula[["observation"]]        = ~ capture_intensity
```

Model Formulas declarations for the simulated example.

| Model parameter | Model formula |
| --- | --- |
| **`diffusion`** | ~ air\_temperature |
| **`advection_1`** | - |
| **`advection_2`** | - |
| **`habitat_preference`** | - |
| **`mortality`** | ~ altitude |
| **`observation`** | ~ capture\_intensity |
| **`traits`** | ~ size |

The model **`m`** for the Continuous Capture Process (CCP) is set as

```
  m = Jsmm::jsmm(domain = domain, observation_effort = observation_effort,
           releases = data[["releases"]], captures = data[["captures"]], 
           cov_data = cov_data, 
           ns = ns, sp_names = sp_names, tr_data = tr_data, C = C,
           model_formula = model_formula)


model_summary(m)
```

```
## Domain: nele = 258 | nnod = 160 
## Capture method: CCP | Number release sites: 4 | Number capture sites: 9 
## Number of time partitions: 11 
## ------------------------------------- 
## Model formulas: 
##     Diffusion ~ air_temperature | nc = 2 
##     Mortality ~ altitude | nc = 2 
##     Traits ~ size 
##     Observation ~ capture_intensity | nc = 2 
## ------------------------------------- 
## Number of species: 20
```

```
#Here we are saving the model **`m`** just created.

save(m, file = paste0("models/unfitted_model_simulated_", 
                      method, "_no_data.RData"))
```

#### Step 1b: Simulate model parameters from priors

This optional step allows generating the model parameters from predefined priors through the function **`simulate_from_prior(m)`** (See Appendix S1. and Fig. 1. main manuscript). The script below sets the diffusion parameter coefficients \(\beta^a\_0\) and \(\beta^a\_1\) increasing with the body size and air temperature, respectively; the mortality increases with altitude; and the observation rate is higher with high intensity.

First, we load the model created in the previous step.

```
method = "CCP"

file = paste0("models/unfitted_model_simulated_", method, "_no_data.RData")
  
load(file)
```

```
muZ = m[["prior"]][["muZ"]]
nc = m[["nc"]]
np = sum(nc)
nt = m[["nt"]]
idx = 1

for(j in 1:nt){
  for(i in 1:length(nc)){
    for(ii in seq_len(nc[i])){
      
      # baseline diffusion
      if(names(nc)[i] == "diffusion" & ii == 1 & j == 1) muZ[idx] = -4
      
      # diffusion increases with air temperature
      if(names(nc)[i] == "diffusion" & ii == 2 & j == 1) muZ[idx] = 1 
        
      # diffusion intercept increases with body size
      if(names(nc)[i] == "diffusion" & ii == 1 & j == 2) muZ[idx] = 1 
      
      # baseline mortality
      if(names(nc)[i] == "mortality" & ii == 1 & j == 1) muZ[idx] = -2 
      
      # mortality increases with altitude
      if(names(nc)[i] == "mortality" & ii == 2 & j == 1) muZ[idx] = 1 
      
      # baseline observation rate
      if(names(nc)[i] == "observation" & ii == 1 & j == 1) muZ[idx] = -2 
        
      # observation rate higher with high intensity
      if(names(nc)[i] == "observation" & ii == 2 & j == 1) muZ[idx] = 1 
      
      idx = idx + 1
    }
  }
}

m[["prior"]][["muZ"]] = muZ
m[["prior"]][["VZ"]] = 0*m[["prior"]][["VZ"]]
m[["prior"]][["rho"]] = matrix(c(0.75, 1), nrow = 1)

pars = Jsmm::simulate_from_prior(m)
```

```
save(pars, file = paste0("parameters/true_parameters_", method,".RData"))
```

The model parameters are set up within the named list **`pars$Beta`**. Within each element of this list, there are matrices for each of the movement and observation parameters, that is, **`pars$Beta$diffusion`**, **`pars$Beta$mortality`**, and **`pars$Beta$observation`**. For example, for diffusion, a matrix with twenty rows and two columns is created. Rows correspond to the species, and the first and second columns correspond to the coefficients \(\beta^a\_0\), and \(\beta^a\_1\).

```
head(pars$Beta$diffusion)
```

```
##           [,1]     [,2]
## [1,] -3.871749 1.313660
## [2,] -3.846895 1.211122
## [3,] -5.295459 1.000561
## [4,] -5.261582 1.289043
## [5,] -3.047360 1.597476
## [6,] -3.325745 1.548127
```

Figure 26: Diffusion intercept is influenced by the species body size.

Table 4: Model coeficients for diffusion, mortality and observation parameters for the twenty species.

| species | betaa0 | betaa1 | betam0 | betam1 | betao0 | betao1 |
| --- | --- | --- | --- | --- | --- | --- |
| 1 | -3.872 | 1.314 | -2.378 | 1.124 | -2.205 | 0.846 |
| 2 | -3.847 | 1.211 | -2.645 | 1.297 | -2.359 | 0.532 |
| 3 | -5.295 | 1.001 | -3.033 | 1.234 | -1.480 | 0.496 |
| 4 | -5.262 | 1.289 | -2.162 | 1.308 | -1.480 | 0.542 |
| 5 | -3.047 | 1.597 | -2.725 | 1.195 | -2.682 | 0.963 |
| 6 | -3.326 | 1.548 | -2.127 | 1.051 | -2.497 | 1.423 |
| 7 | -4.307 | 1.056 | -3.017 | 1.658 | -1.004 | 0.467 |
| 8 | -3.211 | 1.526 | -2.442 | 1.177 | -1.645 | 0.307 |
| 9 | -4.567 | 1.721 | -2.571 | 1.330 | -2.281 | 0.769 |
| 10 | -3.665 | 1.511 | -2.104 | 1.141 | -2.805 | 1.319 |
| 11 | -4.068 | 0.994 | -3.108 | 1.331 | -1.498 | 0.861 |
| 12 | -4.515 | 1.491 | -2.439 | 1.616 | -1.470 | 0.543 |
| 13 | -3.727 | 1.119 | -2.335 | 0.963 | -2.239 | 0.507 |
| 14 | -4.497 | 1.451 | -2.114 | 0.799 | -2.385 | 1.390 |
| 15 | -4.225 | 1.272 | -2.776 | 1.322 | -1.353 | 0.264 |
| 16 | -5.276 | 1.986 | -2.446 | 1.210 | -1.595 | 0.724 |
| 17 | -3.598 | 0.989 | -2.540 | 0.866 | -2.154 | 0.774 |
| 18 | -4.367 | 1.880 | -2.201 | 0.711 | -2.641 | 1.144 |
| 19 | -2.879 | 0.995 | -2.664 | 1.410 | -1.736 | 0.379 |
| 20 | -4.596 | 1.729 | -2.399 | 1.355 | -1.727 | 0.743 |

Figure 27: Model parameters for each of the twenty species. Upper to lower figures show diffusion, mortality, and observation dependence on the predictors; air temperature, altitude, and capture intensity, respectively.

#### Step 1c: Simulate capture data

For this example, we simulated 500 individuals for each species. We also declared that secondary releases are not possible.

```
method = "CCP"

load(paste0("parameters/true_parameters_", method, ".RData"))
load(paste0("models/unfitted_model_simulated_", method, "_no_data.RData"))
```

```
ni = round(10000/m[["ns"]]) #number of individuals per species

releases = matrix(NA, ncol = 2, nrow = ni*m[["ns"]])

releases[, 1] = rep(1:m[["ns"]], ni)
releases[, 2] = rep(1, ni*m[["ns"]])
releases[, 2] = sample(seq_len(length(m$observation_effort$releases$location)),
                       size = ni*m[["ns"]], replace = TRUE)

colnames(releases) = c("sp", "release_event")
  
secondary_release = FALSE
max_dt = NULL

head(releases)
```

```
##      sp release_event
## [1,]  1             5
## [2,]  2            10
## [3,]  3             7
## [4,]  4             4
## [5,]  5            10
## [6,]  6             8
```

For example, the second row of the releases matrix indicates that one individual from species 2 was released at the release site number four at \(t = 9\). (release event was declared on the **`observation_effort`** list).

As before, we can proceed to simulate capture-recapture data by using the function **`simulate_captures(.)`**.

```
m = Jsmm::simulate_captures(m = m, pars = pars, releases = releases,
                      max_dt = max_dt, secondary_release = secondary_release)
```

```
save(m, file = paste0("models/unfitted_model_simulated_", method, ".RData"))
```

```
Jsmm::model_summary(m)
```

```
## Domain: nele = 258 | nnod = 160 
## Capture method: CCP | Number release sites: 4 | Number capture sites: 9 
## Number of time partitions: 11 
## ------------------------------------- 
## Model formulas: 
##     Diffusion ~ air_temperature | nc = 2 
##     Mortality ~ altitude | nc = 2 
##     Traits ~ size 
##     Observation ~ capture_intensity | nc = 2 
## ------------------------------------- 
## Number of species: 20 
## Releases Data:
##      10000 unique releases 
## Captures Data:
##      4866 captures
```

#### Step 2a: Fit the model

The parameter estimation within the Jsmm is made via Markov Chain Monte Carlo (MCMC) through the function **`sampleMCMC(.)`**. The user should choose reasonable parameters for the sampler to ensure that the MCMC has good mixing, low autocorrelation, and reliable convergence. For this simple example, we will pretend that we do not know the model parameters, and we will proceed to estimate those. The parameters used in our sampler are set in Table 13.

Model Formulas declarations for the simulated example.

| MCMC sampler parameter | Value |
| --- | --- |
| thin | 2 |
| samples | 250 |
| transcient | 250 |
| number of chains | 5 |

```
mname = "simulated_CCP"
load(paste0("models/unfitted_model_", mname, ".RData"))
  
m         = merge_rc_data(m)
thin      = 2
samples   = 250
transient = round(0.5*samples*thin)
nChains   = 5
init_pars = NULL

for(i in 1:nChains){
  set.seed(i)
  m = Jsmm::sampleMcmc(m = m, samples = samples, transient = transient, thin = thin, 
                 nChains = 1, init_pars = init_pars)
  
  save(m, file = paste0(paste0("models/", i, "_fitted_model_", mname, "_samples_", samples, 
                               "_thin_", thin, "_nChains_", nChains, ".RData")))
}

postList = list()
  
for(i in 1:nChains){
    fname = paste0(paste0("models/", i, "_fitted_model_", mname, "_samples_", samples, 
                          "_thin_", thin, "_nChains_", nChains, ".RData"))
    load(file = fname)
    postList[[i]] = m[["postList"]][[1]]
    file.remove(fname)
}
  
m[["postList"]] = postList
  
save(m, file = paste0(paste0("models/fitted_model_", mname, "_samples_", samples, 
                             "_thin_", thin, "_nChains_", nChains, ".RData")))
```

#### Step 2b: Evaluating MCMC convergence

After fitting the model, we will assess the MCMC convergence. First, we convert the generated chains to objects of the MCMC class. Here, this is done through the function **`convert_to_coda_object(.)`**, which is based on the R-package Coda (Plummer et al. (2006)). Note that the Gelman-Rubin statistic (R-hat) diagnosis of convergence shows that the chains converge satisfactorily. The Figs. 28-34. Show the trace plots for our model for the first species.

```
mname = "simulated_CCP"

thin    = 2
samples = 250
nChains = 5

load(paste0(paste0("models/fitted_model_", mname, 
                   "_samples_", samples, "_thin_", thin, "_nChains_", nChains, ".RData")))

mpost = Jsmm::convert_to_coda_object(m)

summary(coda::gelman.diag(mpost$Beta, multivariate = FALSE)$psrf)
```

```
##    Point est.       Upper C.I.    
##  Min.   : 1.003   Min.   : 1.013  
##  1st Qu.: 1.032   1st Qu.: 1.084  
##  Median : 1.090   Median : 1.227  
##  Mean   : 2.566   Mean   : 4.029  
##  3rd Qu.: 1.286   3rd Qu.: 1.690  
##  Max.   :39.149   Max.   :73.189
```

Figure 28: First species trace plots model parameters Beta., five MCMC chains.

Figure 29: First species trace plots model parameters Beta., five MCMC chains.

Figure 30: First species trace plots, five MCMC chains Gamma.

Figure 31: First species trace plots, five MCMC chains Gamma.

Figure 32: First species trace plots, five MCMC chains Gamma.

Figure 33: First species trace plots, five MCMC chains Gamma.

Figure 34: First species trace plots, five MCMC chains Rho.

#### Step 2c: Simulate prior and posterior predictive data

The script below generates 1000 datasets for the CCP simulated example. The main purpose of generating predictive data is exploring how the fitted model behaves in time and space compared to the original data. This method helps to infer if the fitted parameters provide a good approximation of the real data.

```
mname = "simulated_CCP"

for(nrepl in c(1000)){
  showTrueValues = TRUE
  
  thin    = 2
  samples = 250
  nChains = 5
  
  load(paste0(paste0("models/fitted_model_", mname, "_samples_", samples, "_thin_", thin, 
                     "_nChains_", nChains, ".RData")))
  
  post = m[["postList"]]
  
  if(showTrueValues){
    types = c("true", "prior", "posterior")
    load("parameters/true_parameters_CCP.RData")
    true_pars = pars
  }else{
    types =  c("prior", "posterior")
  }

  for(type in types){
    PPcaptures = list()

    for(repl in 1:nrepl){
      if(type == "true"){
        pars = true_pars
      }
      
      if(type == "posterior"){
        chain = sample(1:length(post), size = 1)
        sa    = sample(1:length(post[[chain]]), size = 1)
        pars  = post[[chain]][[sa]]
      }
      
      if(type == "prior"){
        pars = Jsmm::simulate_from_prior(m)
      }
      
      max_dt = NULL
      secondary_release = FALSE
      releases = m[["releases"]]
      
      m2 = Jsmm::simulate_captures(m = m, pars = pars, releases = releases,
                               max_dt = max_dt, secondary_release = secondary_release)
        
      captures.simulated = m2[["captures"]]
        
      if(nrow(captures.simulated) > 0){
        PPcaptures[[repl]] = captures.simulated      
      }else{
        PPcaptures[[repl]] = -1
      }
    }
    save(PPcaptures, file = paste0("models/predicted_captures_", mname, "_", 
                                   type, "_nrepl_", nrepl, ".RData"))
  }
}
```

#### Step 2d: Evaluating model fit

The script below provides a visualization of the posterior predictive data generated on the previous step compared with the original simulated dataset and priors. See Fig. 35.

```
showTrueValues = TRUE
showPrior      = TRUE
showPosterior  = TRUE

mname   = "simulated_CCP"
thin    = 2
samples = 250
nChains = 5
nrepl   = 1000

load(paste0(paste0("models/fitted_model_", mname, "_samples_", samples, "_thin_", thin,
                   "_nChains_", nChains, ".RData")))

sel = 1:m$ns

captures = m[["captures"]]

if(showPrior){
  load(file = paste0("models/predicted_captures_", mname, "_prior_nrepl_", nrepl, ".RData"))
  prior.captures = PPcaptures
}

if(showPosterior){
  load(file = paste0("models/predicted_captures_", mname, "_posterior_nrepl_", nrepl, ".RData"))
  posterior.captures = PPcaptures
}

if(showTrueValues){
  load(file = paste0("models/predicted_captures_", mname, "_true_nrepl_", nrepl, ".RData"))
  true.captures = PPcaptures
}

#pdf(paste0("results/model_fit_", mname, ".pdf"))

par(mfrow = c(2, 2))

for(k in 1:4){
  if(k == 1){
    fun  = function(m,captures) Jsmm::number_of_captures(m, captures)
    xlab = ""
    ylab = "total number of captures"
  }
  
  if(k == 2){
    fun  = function(m,captures) Jsmm::mean_captures(m, captures)
    xlab = ""
    ylab = "mean number of captures per individual"
  }
  
  if(k == 3){
    fun  = function(m, captures) Jsmm::mean_time_to_capture(m, captures)
    xlab = ""
    ylab = "mean time to capture"
  }
  
  if(k == 4){
    fun = function(m, captures) Jsmm::mean_distance_between_captures(m, captures)
    xlab = ""
    ylab = "mean distance between captures"
  }
    
  obs = fun(m, captures)
  ma = max(obs, na.rm = T)
  
  if(showPrior){
    prior.pred = matrix(ncol = length(obs), nrow = nrepl)
    for(i in 1:nrepl) prior.pred[i, ] = fun(m,prior.captures[[i]])
    ma = max(ma, max(prior.pred, na.rm = TRUE))
  }
  
  if(showPosterior){
    posterior.pred = matrix(ncol = length(obs), nrow = nrepl)
    for(i in 1:nrepl) posterior.pred[i, ] = fun(m,posterior.captures[[i]])
    ma = max(ma, max(posterior.pred, na.rm = TRUE), na.rm = TRUE)
  }
  
  if(showTrueValues){
    true.pred = matrix(ncol = length(obs), nrow = nrepl)
    for(i in 1:nrepl) true.pred[i, ] = fun(m, true.captures[[i]])
    ma = max(ma, max(true.pred, na.rm = TRUE))
  }
  
  plot(obs[sel], ylim = c(0, ma), xlim = c(0.5, m$ns + 0.5), xlab = xlab, ylab = ylab, 
       pch = 16, col = "black", cex = 1,
        xaxt = "n")
  axis(1, at = 1:m$ns, label = m$sp_names[sel], las = 2, cex.axis = 0.7)
  
  if(showTrueValues){
    qu = apply(true.pred, MARGIN = 2, FUN = quantile, prob = c(0.025, 0.975), na.rm = TRUE)
    for(i in 1:length(obs)) lines(x = c(i, i), y = qu[, sel[i]], col = "black")
  }
  
  if(showPrior){
    qu = apply(prior.pred, MARGIN = 2, FUN = quantile, prob = c(0.025, 0.975), na.rm = TRUE)
    me = apply(prior.pred, MARGIN = 2, FUN = median, na.rm = TRUE)
    points(x = (1:length(me)) - 0.25, y = me[sel], pch = 16, col = "grey")
    
    for(i in 1:length(obs)) lines(x = c(i, i) - 0.25, y = qu[, sel[i]], col = "grey")
  }
  
  if(showPosterior){
    qu = apply(posterior.pred, MARGIN = 2, FUN = quantile, prob = c(0.025, 0.975), na.rm = TRUE)
    me = apply(posterior.pred, MARGIN = 2, FUN = median, na.rm = TRUE)
    points(x = (1:length(me)) + 0.25, y = me[sel], pch = 16, col = "red")
    
    for(i in 1:length(obs)) lines(x = c(i, i) + 0.25, y = qu[, sel[i]], col = "red")
  }
}
```

Figure 35: Posterior predictive data generated with 1000 datasets. Black, red, and grey colors represent the datasets generated with true, posterior, and prior parameters values.

```
#dev.off()
```

#### Step 3a: Show Posteriors vs priors

The script below provides a visualization of the posterior credible intervals for each model parameter (pink color) and its corresponding prior (grey color). See Fig. 36-38.

```
showTrueValues = TRUE
mname = "simulated_CCP"

thin     = 2
samples  = 250
nChains  = 5
filename = paste0(paste0("models/fitted_model_", mname, "_samples_", samples, "_thin_", thin,
                         "_nChains_", nChains, ".RData"))
load(filename)

sel   = 1:m$ns
post  = poolMcmcChains(m[["postList"]])
npost = length(post)
vars  = names(m[["nc"]][m[["nc"]] > 0])
ns    = m[["ns"]]
nt    = m[["nt"]]
  
nprior = npost
prior  = list()

for(repl in 1:nprior){
  prior[[repl]] = Jsmm::simulate_from_prior(m)
}
  
load("parameters/true_parameters_CCP.RData")

par(mfrow=c(3,2))

cc = 0
for(va in vars){
  nv = m[["nc"]][[va]]
  for(i in 1:nv){
    cc = cc + 1
      
    re.pri = matrix(ncol = ns, nrow = nprior)
    re.pri.G = matrix(ncol = nt, nrow = nprior)
    
    for(repl in 1:nprior){
      re.pri[repl, ]   = prior[[repl]][["Beta"]][[va]][, i]
      re.pri.G[repl, ] = prior[[repl]][["Gamma"]][, cc]
    }
    
    re.post   = matrix(ncol = ns, nrow = npost)
    re.post.G = matrix(ncol = nt, nrow = nprior)
    
    for(repl in 1:npost){
      re.post[repl, ]   = post[[repl]][["Beta"]][[va]][, i]
      re.post.G[repl, ] = post[[repl]][["Gamma"]][, cc]
    }
    
    ymin = min(quantile(re.pri, probs = 0.05), min(re.post))
    ymax = max(quantile(re.pri, probs = 0.95), max(re.post))
    
    if(showTrueValues){
      ymin = min(ymin, pars$Beta[[va]][, i])
      ymax = max(ymax, pars$Beta[[va]][, i])
    }
    
    colnames(re.pri) = m$sp_names
    re.pri  = re.pri[, sel]
    re.post = re.post[, sel]
    
    boxplot(re.pri, outline = FALSE, at = 3*(1:ns)-0.75, ylim = c(ymin, ymax),
            main = m[["parNames"]][cc], las = 2, cex.axis = 0.7,
            xlim = c(2, 3*ns + 2),
            ylab = "parameter value", col = "grey", xaxt = "n")
    
    axis(1, at = 3*(1:m$ns), label = colnames(re.pri), las = 2, cex.axis = 0.7)
    
    boxplot(re.post, outline = FALSE, at = 3*(1:ns) + 0.75, col = "pink", add = TRUE, 
            xaxt = "n", yaxt = "n")
    
    abline(h = 0)
    
    if(showTrueValues){
      for(j in 1:m[["ns"]]){
        lines(x = c(3*j, 3*j + 1.5), y = c(pars$Beta[[va]][j, i], pars$Beta[[va]][j, i]), 
              lwd = 2, col = "red")
      }
    }
    
    ymin = min(quantile(re.pri.G, probs = 0.05), min(re.post.G))
    ymax = max(quantile(re.pri.G, probs = 0.95), max(re.post.G))
    
    if(showTrueValues){
      ymin = min(ymin, pars$Gamma[, cc])
      ymax = max(ymax, pars$Gamma[, cc])
    }
    
    colnames(re.pri.G) = colnames(m$X[["trait"]])
    
    boxplot(re.pri.G, outline = FALSE, at = 3*(1:nt), ylim = c(ymin, ymax),
            main = m[["parNames"]][cc], las = 2, cex.axis = 0.7,
            xlim = c(2, 3*nt + 2),
            ylab = "parameter value", col = "grey")
    
    boxplot(re.post.G, outline = FALSE, at = 3*(1:nt) + 1.5, col = "pink", add = TRUE, 
            xaxt = "n", yaxt = "n")
    
    abline(h = 0)
    
    if(showTrueValues){
      for(j in 1:m[["nt"]]){
        lines(x = c(3*j, 3*j + 1.5), y = c(pars$Gamma[j, cc], pars$Gamma[j, cc]), 
              lwd = 2, col = "red")
      }
    }
  }
}
```

Figure 36: Posterior credible intervals for the model parameters. Left panels show the posteriors for each of the model parameters (pink color), their corresponding priors (Grey), and the parameter true value (Red). Right panels show the corresponding values for the Gamma associated parameter.

Figure 37: Posterior credible intervals for the model parameters. Left panels show the posteriors for each of the model parameters (pink color), their corresponding priors (Grey), and the parameter true value (Red). Right panels show the corresponding values for the Gamma associated parameter.

Figure 38: Posterior credible intervals for the associated rho parameter. Posterior, prior, and true value are represented by pink, grey, and red colors, respectively.

### How to set temporally varying capture event-specific covariates for CCP capture process

The list **`obs_t_data`** allows temporal variations within capture events. This list has a length equal to the number of capture events. Within each index \(c\), there is a data frame (**`obs_t_data[[c]]$cov`**), and a vector (**`obs_t_data[[c]]$times`**) with covariate information and time instants, respectively, for the capture event \(c\).

Below we set **`obs_t_data`** for an example in which the domain and observation\_effort are as in the Supplementary Material (S 02). We included numeric variables for microclimate and humidity, and factor variables representing the different possible light settings (See Table 1.).

Covariates for the observation parameter

| Covariate name | Event | Time variation within events | Type |
| --- | --- | --- | --- |
| microclimate | No | Yes | Numeric |
| humidity | No | Yes | Numeric |
| light\_settings | No | Yes | Factor |

```
nc = length(observation_effort[["captures"]][["location"]])

obs_t_data = list(list())
                      
for(c in 1:nc){
  t = observation_effort[["captures"]][["time"]][c, ]
  h = sample(1:8, 1)
  ts = seq(from = t[1], to = t[2], by = 1/h)
  li = list()
                    
  li[["covs"]] = data.frame(microclimate = rnorm(length(ts)), 
                            humidity = rnorm(length(ts)),
                            light_settings = factor(sample(c("a", "b", "c"), length(ts), 
                                                          replace = TRUE) , 
                                                          levels = c("a", "b", "c")))
                                              
  li[["times"]] = ts
  obs_t_data[[c]] = li
}
```

```
cov_data[["obs_t_data"]] = obs_t_data
```

The information for the time-dependent covariate for the capture event \(c =\) 35 can be retrieved as

```
cov_data$obs_t_data[[c]]$covs
```

```
##   microclimate   humidity light_settings
## 1   -1.7872898  0.4376164              b
## 2    0.3413938  0.2702748              a
## 3    2.7339644 -1.6601651              a
## 4    1.1542943  0.1134256              b
## 5   -1.9944199  0.7366299              c
```

```
cov_data$obs_t_data[[c]]$times
```

```
## [1] 3.00 3.25 3.50 3.75 4.00
```

Note from **`obs_effort$captures`** that the capture event \(c =\) 35 occurs on location 8 within the time interval \(t\_{35,1} =\) 3 and \(t\_{35,2} =\) 4.

Below is the declaration of the corresponding model formula for the observation parameter:

```
model_formula[["observation"]] = ~ humidity*microclimate + light_settings

model_formula$observation
```

```
## ~humidity * microclimate + light_settings
```

Note: It is also possible to define formulas as a combination of constant covariates within events and temporally varying within events covariates.

### Summary input data Jsmm software

The function **`jsmm(.)`** that define the model **`m`** object takes as input the following parameters: **`domain`**, **`observation_effort`**, **`releases`**, **`captures`**, **`cov_data`**, **`ns`**,**`sp_names`**, **`C`**, **`tr_data`**, and **`model_formula`**.

#### Domain

The input parameter **`domain`** initially is a polygonal object from the class **`sf`**. Each sub polygon within **`domain`** has a label **`id`** that provides information on whether it belongs to a certain habitat and a label **`id_location`** that indicates if it is a release or capture site.

On a second step, **`domain`** is transformed via the function **`domain = jsmm_add_triangulation(.)`** in a named list that consists of two elements. The first element is **`polygon`**, which is a modified polygonal object from the **`sf`** class with triangular polygons defined over \(\Omega\) that inherits the labels **`id`**, and **`id_location`** from before.

The second element from the modified **`domain`** is the named list **`triangulation`**, this list consists of the elements **`ele`** and **`node`**. The object **`ele`** is a matrix where the rows correspond to the triangular elements, and the three columns list the nodes that define each element. The object **`node`** is a matrix where the rows correspond to the nodes, and the two columns give the x- and y-coordinates of the nodes.

The function **`jsmm_add_triangulation(domain, max_t_area, min_t_angle)`** has as input the parameters **`max_t_area`**, and **`min_t_angle`**, which provide specifications about area and triangle internal angles constrains for the triangulation. The triangulation within **`jsmm_add_triangulation(.)`** is built based on the triangulation function **`triangulate(.)`** from the R-package RTriangle (Shewchuk (1996)).

#### Observation effort

The input parameter **`observation_effort`** object is a named list that describes where and when the releases and captures occur within the experiment. It consists of the objects **`method`**, **`locations`**, **`releases`**, and **`captures`**.

The object **`method`** is a string that describes the nature of the method used for the capture process. Within our framework, the capture process can be defined as instantaneous (**`method = "ICP"`**) or continuous (**`method = "CCP"`**).

The object \*\*`locations} is a list that describes the spatial locations of the capture and release sites. Each element of the list corresponds to one site, and it is a vector of integers that shows which triangular elements belong to that particular site.

The object **`releases`** is a named list that contains the objects **`releases$location`** and **`releases$time`**. This object describes the release events, i.e., the locations and times at which individuals have been released. Both **`location`** and **`time`** are vectors, where each element corresponds to a particular release event. The value of **`location[i]`** describes the release location by giving the relevant index to the list **`observation_effort$locations`**. The value of **`releases$time[i]`** gives the time at which releases took place.

The object **`captures`** is a named list that contains the objects **`captures$location`** and **`captures$time`**. This object describes the capture events, i.e., the locations and times where captures have been attempted. As with **`releases`**, **`location`** is a vector, where each element corresponds to a particular capture event, and its values describe the capture locations by giving the relevant index to the list **`observation_effort$locations`**. In the case of the ICP, **`time`** is a vector, with values indicating the times when the captures were attempted. In the case of the CCP, **`time`** is a matrix, where the rows correspond to the capture events and the two columns show the times when the capture process was started and finished.

#### Releases

The object **`releases`** is a matrix that describes the species, locations, and times of the released individuals. The rows of this matrix correspond to the individuals, the first column describes the species, and the second column the release event (which codes the information for location and time).

#### Captures

The object **`captures`** contains the capture histories of the individuals (except the release of the individual, which is described in the object **`releases`**).
The object **`captures`** is a matrix with three columns, which describe the identity of the individual (index to the row of the matrix **`releases`**), the capture event in which the individual was captured, and a boolean variable describing if the individual was re-released after the capture or not.

#### Covariate data

The input parameter **`cov_data`** is a named list that consists of (some combination of) the objects described below.

- *Spatial covariates defined through nodes can be used for diffusion, advection, and mortality*. These covariates are incorporated through a dataframe named **`n_data`**, where the columns correspond to covariates, and the rows correspond to the nodes.
- *Spatial covariates defined through elements can be used for habitat preference, diffusion, advection, and mortality*. These covariates are incorporated through a dataframe named **`e_data`**, where the columns correspond to the covariates, and the rows correspond to the elements.
- *Spatially invariant temporal covariates can be used for diffusion, advection, and mortality.* The values \(t\_j\) are given in the numeric vector **`t_data_times`**. The values of the covariates are given in the dataframe **`t_data`**, where the columns correspond to the covariates and the rows to the discretization over time.
- *Spatiotemporal covariates defined through nodes can be used for diffusion, advection, and mortality*. The values \(t\_j\) are given in the numeric vector **`n_t_data_times`**. The values of the covariates are given in the list **`n_t_data`**, where the list element **`n_t_data[[j]]`** corresponds to the time point \(t\_j\). The object **`n_t_data[[j]]`** is a dataframe, where the columns correspond to the covariates and the rows to the nodes.
- *Spatiotemporal covariates defined through elements can be used for habitat preference, diffusion, advection, and mortality.* The values \(t\_j\) are given in the numeric vector **`e_t_data_times`**. The values of the covariates are given in the list **`e_t_data`**, where the list element **`e_t_data[[j]]`** corresponds to the time point \(t\_j\). The object **`e_t_data[[j]]`** is a dataframe, where the columns correspond to the covariates and the rows to the elements.
- *Capture event-specific covariates can be used for ICP and CCP.* These covariates are incorporated through a dataframe named **`obs_data`**, where the columns correspond to the covariates, and the rows correspond to the capture events.
- *Temporally varying capture event-specific covariates can be used for CCP.* These covariates are incorporated through a list named **`obs_t_data`**, where each list element **`c`** corresponds to a particular capture event. For event \(c\), the values \(t\_j\) are given in the numeric vector **`obs_t_data[[c]]$times`**. The values of the covariates are given in the dataframe **`obs_t_data[[c]]$covs`**, where the columns correspond to the covariates and the rows to the discretization over time.

#### Model formula

The input parameter **`model_formula`** is a named list that consists of (some combination of) the objects **`diffusion`**, **`advection`**, **`mortality`**, **`habitat_preference`**, **`observation`**, and **`traits`**. These follow the R-notation for model formulae, e.g., **`model_formula$diffusion =`** \(\sim\) **`habitat*temperature`**, if diffusion is assumed to depend on habitat type, temperature, and their interaction. The intercept is included in all other model formulas except for **`habitat_preference`**, as adding an intercept to habitat preference would overparameterize the model. The user does not need to include \(\sim -1\) in the model of habitat preference as the intercept is removed automatically.

- The model formulae for **`diffusion`**, **`advection`**, **`mortality`** and **`habitat_preference`** can involve any variables that are included in the objects **`n_data`**, **`e_data`**, **`t_data`**, **`n_t_data`** or **`e_t_data`**.
- The model formula for **`observation`** can involve any variables that are included in the objects **`obs_data`** (for both ICP and CCP) or **`obs_t_data`** (for CCP only).
- The model formula for **`traits`** can involve any variables that are included in the objects **`tr_data`**.

#### Number of species

The integer **`ns`** is the number of species included in the model.

#### Species names

The string vector **`sp_names`** is an optional parameter assigning the species names.

#### Trait and phylogenetic data

The trait data is characterized within the **`tr_data`** dataframe component. The columns correspond to the covariates, and the rows to the number of species.

The matrix **`C`** includes the phylogenetic relationships among the species.

### References

Pebesma, Edzer. 2018. “Simple Features for R: Standardized Support for Spatial Vector Data.” *The R Journal* 10 (1): 439–46. https://doi.org/10.32614/RJ-2018-009.

Pebesma, Edzer, and Roger Bivand. 2023. *Spatial Data Science: With applications in R*. Chapman and Hall/CRC. https://doi.org/10.1201/9780429459016.

Plummer, Martyn, Nicky Best, Kate Cowles, and Karen Vines. 2006. “CODA: Convergence Diagnosis and Output Analysis for MCMC.” *R News* 6 (1): 7–11. https://journal.r-project.org/archive/.

Shewchuk, Jonathan Richard. 1996. “Triangle: Engineering a 2D Quality Mesh Generator and Delaunay Triangulator.” In *Applied Computational Geometry: Towards Geometric Engineering*, edited by Ming C. Lin and Dinesh Manocha, 1148:203–22. Lecture Notes in Computer Science. Springer-Verlag.
