## Supplementary material for "R-package Jsmm: Joint species movement modelling of mark-recapture data": S2 Vignettes demonstrating the application of the R-package Jsmm: Supporting_information_S2.pdf

```
head(domain$triangulation$ele)
```

```
##      node_1 node_2 node_3
## [1,]     62     98     97
## [2,]     15     16     62
## [3,]     14     15     62
## [4,]     14     62     97
## [5,]     14     97     13
## [6,]     62     16     17
```

```r
nrow(domain$triangulation$ele)
```

```
## [1] 258
```

```r
head(domain$triangulation$node)
```

```
##      node_x    node_y
## [1,]      0 1.0000000
## [2,]      0 0.9333333
## [3,]      0 0.8666667
## [4,]      0 0.8000000
## [5,]      0 0.7333333
## [6,]      0 0.6666667
```

```r
nrow(domain$triangulation$node)
```

```
## [1] 160
```

Within **domain$polygon**, it is possible to set column indicators for the habitat classification (**id**), and for the release/capture locations (**id_location**). Each row contains information about each triangle or element.

```r
head(domain$polygon)
```

```
## Simple feature collection with 6 features and 2 fields
## Geometry type: POLYGON
## Dimension:     XY
## Bounding box:  xmin: 0 ymin: 0 xmax: 0.1916667 ymax: 0.2
## CRS:           NA
##   id id_location                       geometry
## 1  2           1 POLYGON ((0.06666667 0.0666...
## 2  2           0 POLYGON ((0 0.06666667, 0 0...
## 3  2           0 POLYGON ((0 0.1333333, 0 0....
## 4  2           0 POLYGON ((0 0.1333333, 0.06...
## 5  2           0 POLYGON ((0 0.1333333, 0.06...
## 6  2           0 POLYGON ((0.06666667 0.0666...
```

The function **plot_domain(.)** shows the triangulated domain. The **customize_plot** list shown below provides control of the main aspects of the visualization.

```r
  customize_plot = list()
  customize_plot[["color_map"]]   = "#f0f0f0"
  customize_plot[["labs_axis"]]   = c("X", "Y")
  customize_plot[["lab_legend"]]  = "Matrix"
  customize_plot[["name_legend"]] = "Habitat"
  customize_plot[["color_polyb"]] = "#d6dbdf"
  customize_plot[["lwd_poly"]]    = 0.001
  customize_plot[["site_col"]]    = "red"
  customize_plot[["siteb_col"]]   = "darkred"
  customize_plot[["site_text_pos"]]   = c(-0.05, -0.05)
  customize_plot[["site_text_size"]]  = 4.2
  customize_plot[["panelb_color"]]    = "black"
  customize_plot[["panelb_lwd"]]  = 1
```

```r
 Jsmm::plot_domain(domain = domain, customize_plot = customize_plot)
```

Note: To explore in detail the generated triangulation, is possible to utilize the function **mapview()** from the R-package mapview:

```r
mapview(domain$polygon, zcol = "id") # visualize triangulation with zoom
```

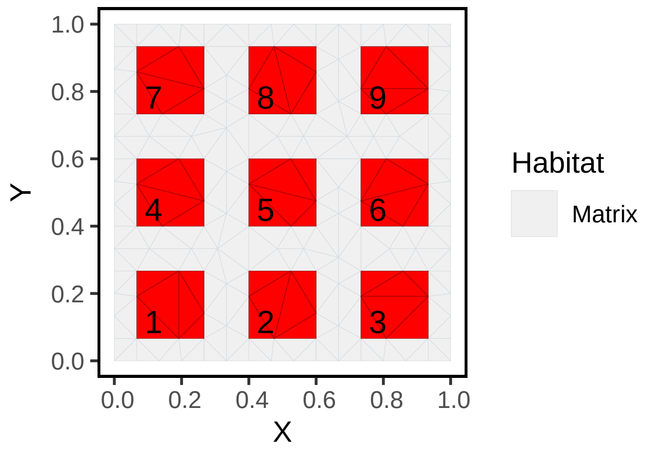

Figure 1: Study region for the simulated domain with 9 capture/release sites. The triangulation is composed of 258 elements and 160 nodes.

#### Creating the observation effort

The **observation_effort** is a named list that contains all information about how releases and captures occur in the experiment. The list is composed of a string called **method**, named lists called **locations**, **releases**, and **captures**. Fig. 2. shows information about the release and capture events in the experiment. The list **locations** contains spatial information about the sites in which releases or captures occurred. For this example, we set the capture process within **method** as ICP. Below is the script to set the **observation_effort**.

```
method           = "ICP"
study_duration  = 10 # duration of the entire study
recapture_sites = 1:9
trap_checking_interval = 1

nloc = length(recapture_sites)
observation_effort = list()
observation_effort[["method"]] = method

locations = list()  # Characterizing the elements that belong to each capture/release site

for(i in 1:nloc){
  locations[[i]] = which(domain[["polygon"]][["id_location"]] == recapture_sites[i])
}

observation_effort[["locations"]] = locations
```

```
observation_effort$method
```

```
## [1] "ICP"
```

```
length(observation_effort$locations)
```

```
## [1] 9
```

Information about the elements that belong to location 5 can be retrieved by

```
observation_effort$locations[[5]]
```

```
## [1]  43  46 108 114 116 217
```

The list **releases**, has the vectors **time**, and **location**, which provide information about when and where releases are possible. Each possible relationship is called a release event. For this simple example, we have set ten release events.

```r
observation_effort[["releases"]][["location"]] = sample(recapture_sites,
                                                         size = study_duration,
                                                         replace = T)

observation_effort[["releases"]][["time"]] = 0:(study_duration - 1)
```

The release event 1 was held on location 5 at time $t = 0$. This information can be retrieved as

```r
observation_effort$releases$time[1]
```

```
## [1] 0
```

```r
observation_effort$releases$location[1]
```

```
## [1] 5
```

The list **captures** is composed of the vector **location** and the vector **time**. Below, we set this list for the ICP capture process.

```r
recapture_times = c()

for(i in seq(trap_checking_interval, study_duration, by = trap_checking_interval)){
  recapture_times = c(recapture_times, rep(i, nloc))
}

recapture_locations = rep(recapture_sites, study_duration/trap_checking_interval)
observation_effort[["captures"]][["location"]] = recapture_locations
observation_effort[["captures"]][["time"]]     = recapture_times
```

The number of capture events is

```r
length(observation_effort$captures$time)
```

```
## [1] 90
```

Information corresponding to the capture event 12 is retrieved as

```r
observation_effort$captures$time[12]
```

```
## [1] 2
```

```r
observation_effort$captures$location[12]
```

```
## [1] 3
```

Read as: the twelfth capture event occurred at capture site number 3 at time $t = 2$ (third day).

#### Set covariate data

The Jsmm framework allows the inclusion of spatial, temporal, spatio-temporal covariates, and trait covariates. To do this, first, we are required to provide the data in a suitable format. The covariates are set within the named lists **cov_data** and **tr_data**.

For this example, we do not have any covariates; for illustration, we declare this empty list as:

```r
cov_data = NULL
ns       = 2
sp_names = paste0("sp_", 1:ns)
tr_data  = NULL
C        = NULL
```

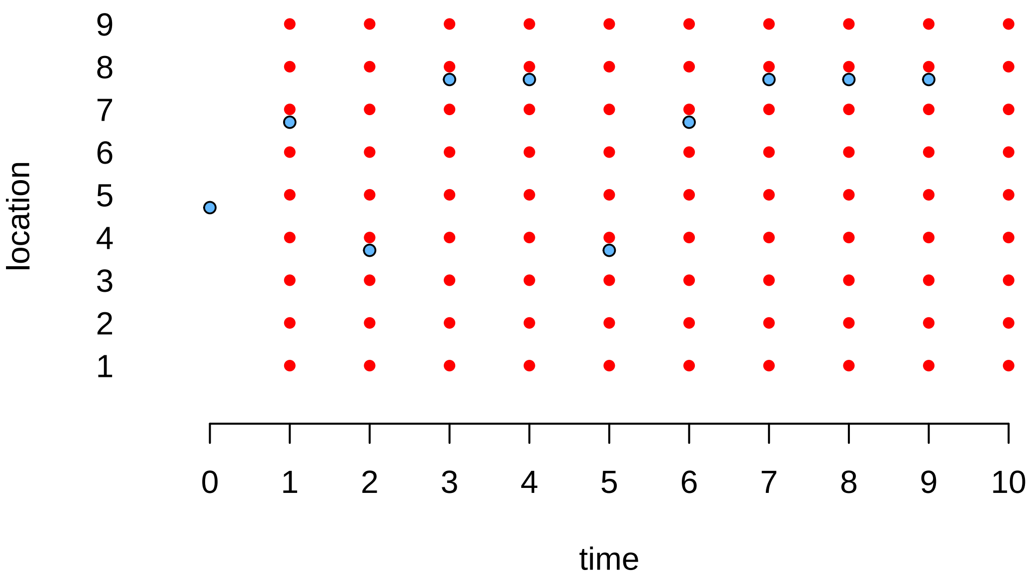

Figure 2: Observation effort scheme for ICP capture process. Times in which releases and captures are possible are represented by blue and red colors, respectively.

Now we can proceed to simulate capture-recapture data by using the function **simulate_captures(.)**.

Table 3: Numbers of recaptures per species.

| $sp_1$ | $sp_2$ |
|---|---|
| 55 | 42 |

```
m = Jsmm::simulate_captures(m = m, pars = pars, releases = releases,
                           max_dt = max_dt, secondary_release = secondary_release)
```

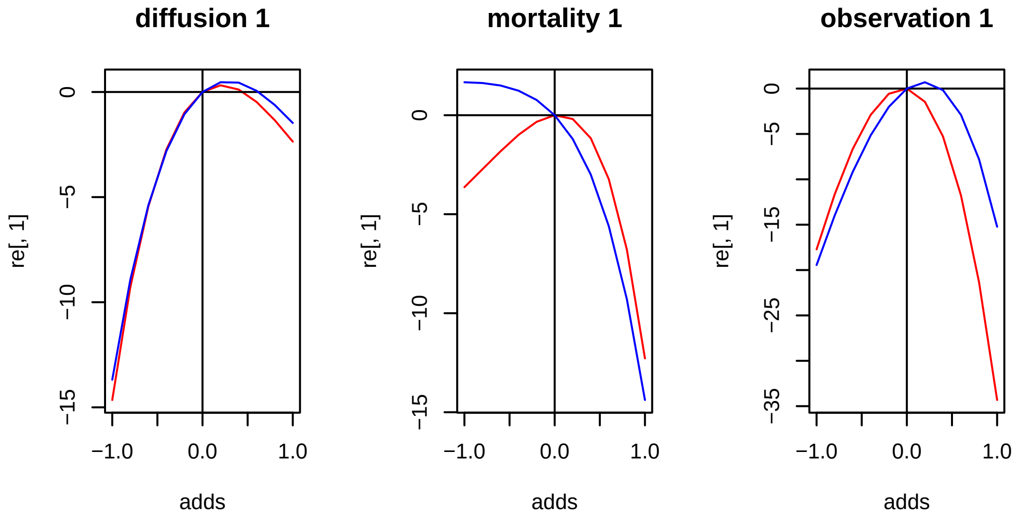

Figure 3: Log likelihood profiles for each of the model parameters.

#### Step 2a: Fit the model

The parameter estimation within the Jsmm is made via Markov Chain Monte Carlo (MCMC) through the function **`sampleMCMC(.)`**. The user should choose reasonable parameters for the sampler to ensure that the MCMC has good mixing, low autocorrelation, and reliable convergence. For this simple example, we will pretend that we do not know the model parameters, and we will proceed to estimate those. The parameters used in our sampler are set in Table 4.

Table 4: Declaration of the model formulas for the simulated example.

| MCMC sampler parameter | Value |
|---|---|
| thin | 2 |
| samples | 500 |
| transcient | 500 |
| number of chains | 5 |

```r
mname = "simulated_ICP_0"

load(paste0("models/unfitted_model_", mname, ".RData"))

load("parameters/true_parameters_ICP_0.RData")

m         = merge_rc_data(m)
thin      = 2
samples   = 500
transient = round(0.5*samples*thin)
nChains   = 5
true_pars = pars

for(i in 1:nChains){
  set.seed(i)
  m = Jsmm::sampleMcmc(m = m, samples = samples, transient = transient, thin = thin, nChains = 1,
                init_pars = true_pars)

  save(m, file = paste0(paste0("models/", i, "_fitted_model_", mname, "_samples_", samples,
                               "_thin_", thin, "_nChains_", nChains, ".RData")))
}

postList = list()

for(i in 1:nChains){
  fname = paste0(paste0("models/", i, "_fitted_model_", mname, "_samples_", samples,
                        "_thin_", thin, "_nChains_", nChains, ".RData"))
  load(file = fname)
  postList[[i]] = m[["postList"]][[1]]
  file.remove(fname)
}

m[["postList"]] = postList

save(m, file = paste0(paste0("models/fitted_model_", mname, "_samples_", samples,
                             "_thin_", thin, "_nChains_", nChains, ".RData")))
```

#### Step 2b: Evaluating MCMC convergence

After fitting the model, we will assess the MCMC convergence. First, we convert the generated chains to objects of the MCMC class. Within our framework, this is done through the function **`convert_to_coda_object(.)`**, which is based on the R-package Coda (Plummer et al. (2006)). Note that the Gelman-Rubin statistic (R-hat) diagnosis of convergence shows that the chains converge satisfactorily. Figs. 4-5 show the trace plots for our model.

Note: The script below generates the corresponding trace plots and saves results in PDF format within a folder called results.

```r
mname = "simulated_ICP_0"

thin    = 2
samples = 500
nChains = 5

load(paste0(paste0("models/fitted_model_", mname,
                   "_samples_", samples, "_thin_", thin, "_nChains_", nChains, ".RData")))
```

```r
mpost = Jsmm::convert_to_coda_object(m)

summary(coda::gelman.diag(mpost$Beta, multivariate = FALSE)$psrf)
```

```
##    Point est.      Upper C.I.
##  Min.   :1.005   Min.   :1.014
##  1st Qu.:1.008   1st Qu.:1.022
##  Median :1.013   Median :1.036
##  Mean   :1.014   Mean   :1.036
##  3rd Qu.:1.016   3rd Qu.:1.044
##  Max.   :1.028   Max.   :1.068
```

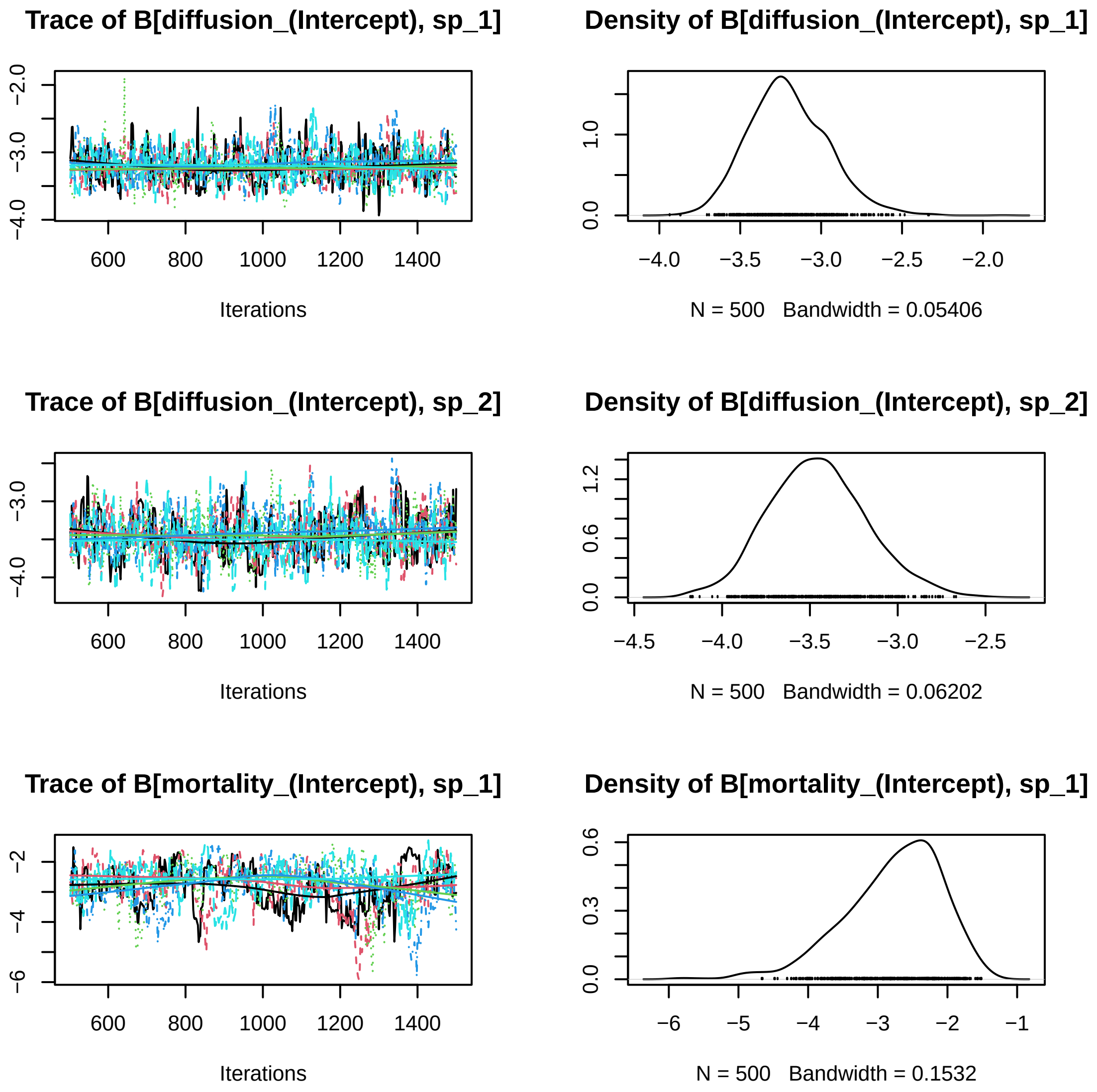

Figure 4: Trace plots for the five MCMC chains.

#### Step 2c: Simulate prior and posterior predictive data

The main purpose of generating predictive data is to explore how the fitted model behaves in time and space compared to the original data. This method helps to infer if the fitted parameters provide a good approximation of the real data. The next script generates 10, 100, and 1000 datasets by utilizing the exact model.

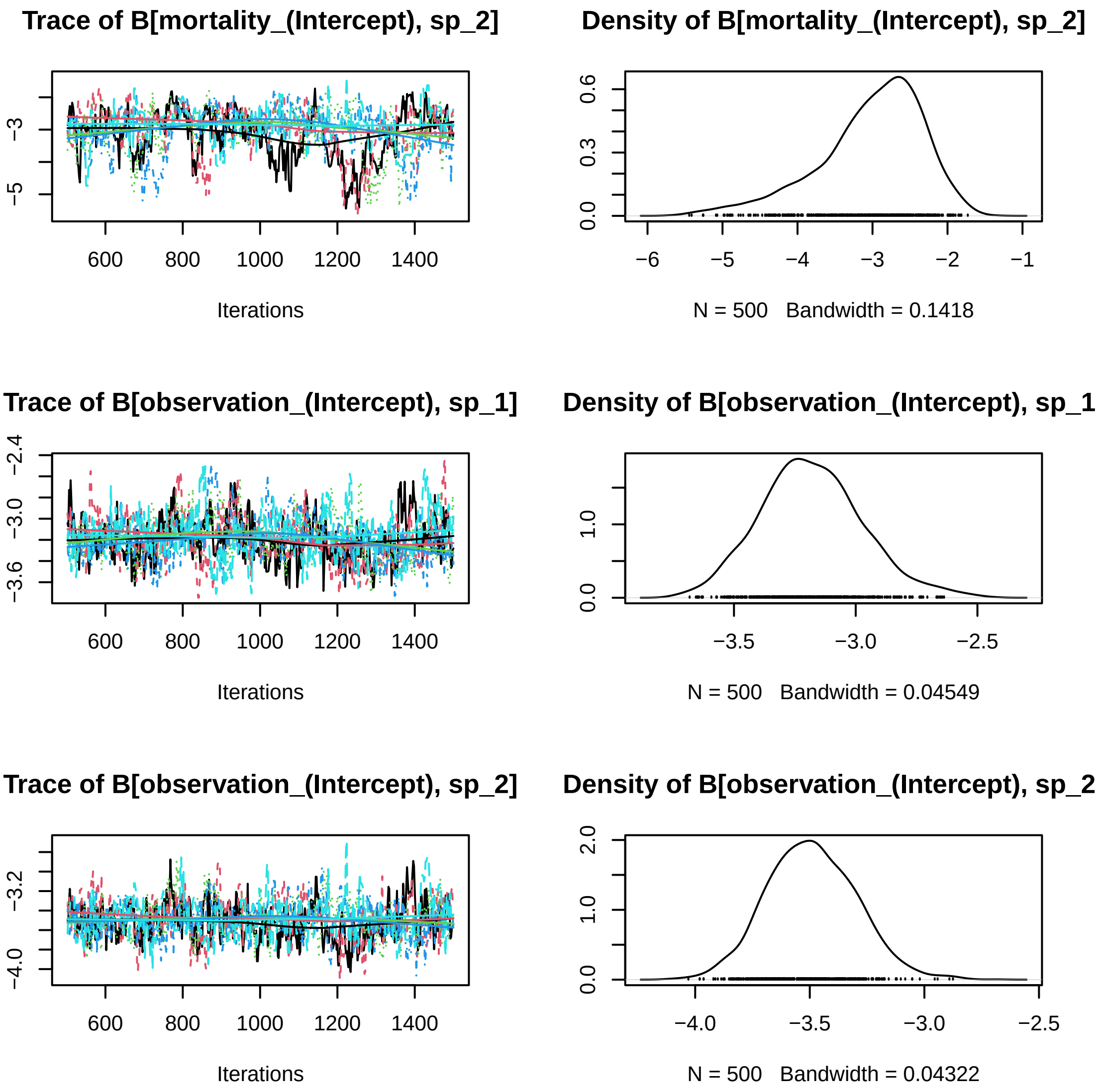

Figure 5: Trace plots for the five MCMC chains.

```
mname = "simulated_ICP_0"

for(nrepl in c(10, 100, 1000)){
  showTrueValues = TRUE

  thin    = 2
  samples = 500
  nChains = 5

  load(paste0(paste0("models/fitted_model_", mname, "_samples_", samples, "_thin_", thin,
                     "_nChains_", nChains, ".RData")))

  post = m[["postList"]]

  if(showTrueValues){
    types = c("true", "prior", "posterior")
    load("parameters/true_parameters_ICP_0.RData")
    true_pars = pars
  }else{
    types =  c("prior","posterior")
  }

  for(type in types){
    PPcaptures = list()

    for(repl in 1:nrepl){
      #print(paste0(type,"_repl_",repl))

      if(type == "true"){
        pars = true_pars
      }

      if(type == "posterior"){
        chain = sample(1:length(post), size = 1)
        sa    = sample(1:length(post[[chain]]), size = 1)
        pars  = post[[chain]][[sa]]
      }

      if(type == "prior"){
        pars = Jsmm::simulate_from_prior(m)
      }

      max_dt = NULL
      secondary_release = TRUE
      releases = m[["releases"]]

      m2 = Jsmm::simulate_captures(m = m, pars = pars, releases = releases,
                                   max_dt = max_dt, secondary_release = secondary_release)

      captures.simulated = m2[["captures"]]

      if(nrow(captures.simulated) > 0){
        PPcaptures[[repl]] = captures.simulated
      }else{
        PPcaptures[[repl]] = -1
      }
```

```
    }
    save(PPcaptures, file = paste0("models/predicted_captures_", mname, "_",
                                   type, "_nrepl_", nrepl, ".RData"))
  }
}
```

#### Step 2d: Evaluating model fit

The script below shows the posterior predictive data generated in the previous step, the original simulated data, and the corresponding priors. Note that in space and time, the posterior predictive data (red color) represents the original data (black color) accurately.

```
showTrueValues = TRUE
showPrior      = TRUE
showPosterior  = TRUE

mname   = "simulated_ICP_0"
thin    = 2
samples = 500
nChains = 5
nrepl   = 1000

load(paste0(paste0("models/fitted_model_", mname, "_samples_", samples, "_thin_", thin,
                   "_nChains_", nChains, ".RData")))

sel = 1:m$ns

captures = m[["captures"]]

if(showPrior){
  load(file = paste0("models/predicted_captures_", mname, "_prior_nrepl_", nrepl, ".RData"))
  prior.captures = PPcaptures
}

if(showPosterior){
  load(file = paste0("models/predicted_captures_", mname, "_posterior_nrepl_", nrepl, ".RData"))
  posterior.captures = PPcaptures
}

if(showTrueValues){
  load(file = paste0("models/predicted_captures_", mname, "_true_nrepl_", nrepl, ".RData"))
  true.captures = PPcaptures
}

#pdf(paste0("results/model_fit_", mname, ".pdf"))

par(mfrow = c(2, 2))

for(k in 1:4){
  if(k == 1){
    fun  = function(m,captures) Jsmm::number_of_captures(m, captures)
    xlab = ""
    ylab = "total number of captures"
  }

  if(k == 2){
```

```r
  fun  = function(m,captures) Jsmm::mean_captures(m, captures)
  xlab = ""
  ylab = "mean number of captures per individual"
}

if(k == 3){
  fun  = function(m, captures) Jsmm::mean_time_to_capture(m, captures)
  xlab = ""
  ylab = "mean time to capture"
}

if(k == 4){
  fun = function(m, captures) Jsmm::mean_distance_between_captures(m, captures)
  xlab = ""
  ylab = "mean distance between captures"
}

obs = fun(m, captures)
ma = max(obs, na.rm = T)

if(showPrior){
  prior.pred = matrix(ncol = length(obs), nrow = nrepl)
  for(i in 1:nrepl) prior.pred[i, ] = fun(m,prior.captures[[i]])
  ma = max(ma, max(prior.pred, na.rm = TRUE))
}

if(showPosterior){
  posterior.pred = matrix(ncol = length(obs), nrow = nrepl)
  for(i in 1:nrepl) posterior.pred[i, ] = fun(m,posterior.captures[[i]])
  ma = max(ma, max(posterior.pred, na.rm = TRUE), na.rm = TRUE)
}

if(showTrueValues){
  true.pred = matrix(ncol = length(obs), nrow = nrepl)
  for(i in 1:nrepl) true.pred[i, ] = fun(m, true.captures[[i]])
  ma = max(ma, max(true.pred, na.rm = TRUE))
}

plot(obs[sel], ylim = c(0, ma), xlim = c(0.5, m$ns + 0.5), xlab = xlab, ylab = ylab,
     pch = 16, col = "black", cex = 1,
      xaxt = "n")
axis(1, at = 1:m$ns, label = m$sp_names[sel], las = 2, cex.axis = 0.7)

if(showTrueValues){
  qu = apply(true.pred, MARGIN = 2, FUN = quantile, prob = c(0.025, 0.975), na.rm = TRUE)
  for(i in 1:length(obs)) lines(x = c(i, i), y = qu[, sel[i]], col = "black")
}

if(showPrior){
  qu = apply(prior.pred, MARGIN = 2, FUN = quantile, prob = c(0.025, 0.975), na.rm = TRUE)
  me = apply(prior.pred, MARGIN = 2, FUN = median, na.rm = TRUE)
  points(x = (1:length(me)) - 0.25, y = me[sel], pch = 16, col = "grey")

  for(i in 1:length(obs)) lines(x = c(i, i) - 0.25, y = qu[, sel[i]], col = "grey")
}
```

```
  if(showPosterior){
    qu = apply(posterior.pred, MARGIN = 2, FUN = quantile, prob = c(0.025, 0.975), na.rm = TRUE)
    me = apply(posterior.pred, MARGIN = 2, FUN = median, na.rm = TRUE)
    points(x = (1:length(me)) + 0.25, y = me[sel], pch = 16, col = "red")

    for(i in 1:length(obs)) lines(x = c(i, i) + 0.25, y = qu[, sel[i]], col = "red")
  }
}
```

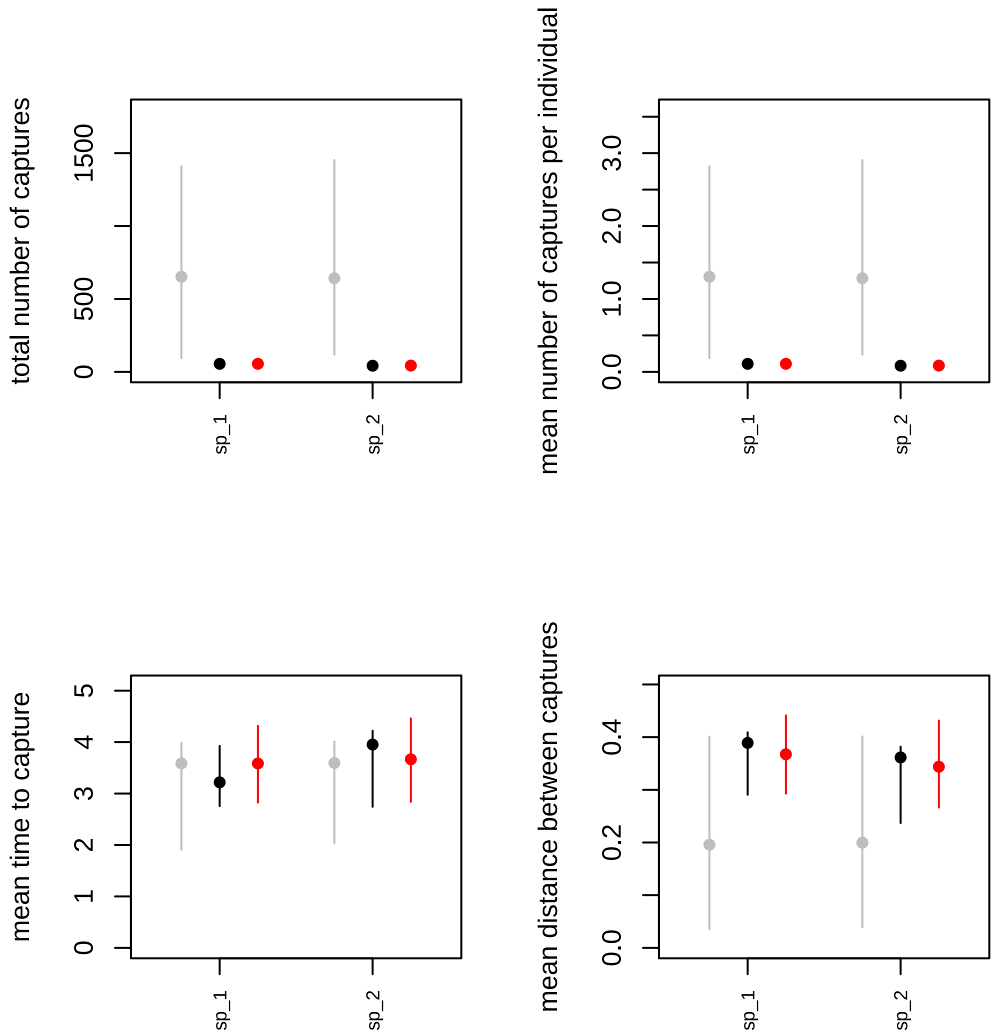

Figure 6: Posterior predictive data generated with 1000 datasets. Black, red, and grey colors represent the datasets generated with true, posterior, and prior parameters values.

```
#dev.off()
```

#### Step 3a: Show Posteriors vs priors

The script below provides a visualization of the posterior credible intervals for each model parameter (pink color) and its corresponding prior (grey color). See Fig. 6. As before, results are saved in PDF format.

```
showTrueValues = TRUE
mname = "simulated_ICP_0"

thin     = 2
```

```r
samples  = 500
nChains  = 5
filename = paste0(paste0("models/fitted_model_", mname, "_samples_", samples, "_thin_", thin,
                         "_nChains_", nChains, ".RData"))
load(filename)

sel   = 1:m$ns
post  = Jsmm::poolMcmcChains(m[["postList"]])
npost = length(post)
vars  = names(m[["nc"]][m[["nc"]] > 0])
ns    = m[["ns"]]
nt    = m[["nt"]]

nprior = npost
prior  = list()

for(repl in 1:nprior){
  prior[[repl]] = Jsmm::simulate_from_prior(m)
}

load("parameters/true_parameters_ICP_0.RData")

#pdf(paste0("results/posteriors_vs_priors_", mname, ".pdf"))

par(mfrow = c(3, 2))
cc = 0
for(va in vars){
  nv = m[["nc"]][[va]]
  for(i in 1:nv){
    cc = cc + 1

    re.pri = matrix(ncol = ns, nrow = nprior)
    re.pri.G = matrix(ncol = nt, nrow = nprior)

    for(repl in 1:nprior){
      re.pri[repl, ]   = prior[[repl]][["Beta"]][[va]][, i]
      re.pri.G[repl, ] = prior[[repl]][["Gamma"]][, cc]
    }

    re.post   = matrix(ncol = ns, nrow = npost)
    re.post.G = matrix(ncol = nt, nrow = nprior)

    for(repl in 1:npost){
      re.post[repl, ]   = post[[repl]][["Beta"]][[va]][, i]
      re.post.G[repl, ] = post[[repl]][["Gamma"]][, cc]
    }

    ymin = min(quantile(re.pri, probs = 0.05), min(re.post))
    ymax = max(quantile(re.pri, probs = 0.95), max(re.post))

    if(showTrueValues){
      ymin = min(ymin, pars$Beta[[va]][, i])
      ymax = max(ymax, pars$Beta[[va]][, i])
    }

    colnames(re.pri) = m$sp_names
```

```
    re.pri  = re.pri[, sel]
    re.post = re.post[, sel]

    boxplot(re.pri, outline = FALSE, at = 3*(1:ns)-0.75, ylim = c(ymin, ymax),
            main = m[["parNames"]][cc], las = 2, cex.axis = 0.7,
            xlim = c(2, 3*ns + 2),
            ylab = "parameter value", col = "grey", xaxt = "n")

    axis(1, at = 3*(1:m$ns), label = colnames(re.pri), las = 2, cex.axis = 0.7)

    boxplot(re.post, outline = FALSE, at = 3*(1:ns) + 0.75, col = "pink", add = TRUE,
            xaxt = "n", yaxt = "n")

    abline(h = 0)

    if(showTrueValues){
      for(j in 1:m[["ns"]]){
        lines(x = c(3*j, 3*j + 1.5), y = c(pars$Beta[[va]][j, i], pars$Beta[[va]][j, i]),
              lwd = 2, col = "red")
      }
    }

    ymin = min(quantile(re.pri.G, probs = 0.05), min(re.post.G))
    ymax = max(quantile(re.pri.G, probs = 0.95), max(re.post.G))

    if(showTrueValues){
      ymin = min(ymin, pars$Gamma[, cc])
      ymax = max(ymax, pars$Gamma[, cc])
    }

    colnames(re.pri.G) = colnames(m$X[["trait"]])

    boxplot(re.pri.G, outline = FALSE, at = 3*(1:nt), ylim = c(ymin, ymax),
            main = m[["parNames"]][cc], las = 2, cex.axis = 0.7,
            xlim = c(2, 3*nt + 2),
            ylab = "parameter value", col = "grey")

    boxplot(re.post.G, outline = FALSE, at = 3*(1:nt) + 1.5, col = "pink", add = TRUE,
            xaxt = "n", yaxt = "n")

    abline(h = 0)

    if(showTrueValues){
      for(j in 1:m[["nt"]]){
        lines(x = c(3*j, 3*j + 1.5), y = c(pars$Gamma[j, cc], pars$Gamma[j, cc]),
              lwd = 2, col = "red")
      }
    }
  }
}
```

```
#dev.off()
```

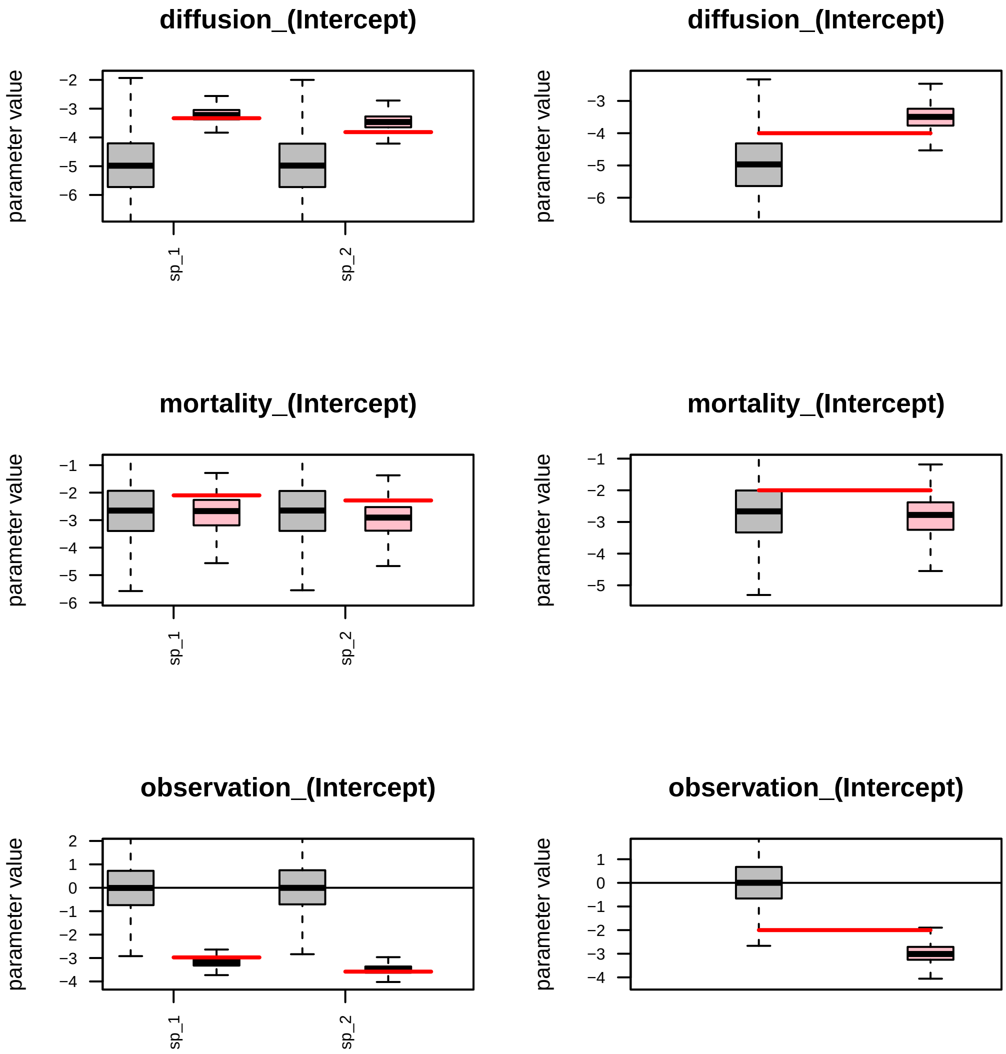

Figure 7: Posterior credible intervals for the model parameters. Left panels show the posteriors for each of the model parameters (pink color), their corresponding priors (Grey), and the parameter true value (Red). The right panels show the corresponding values for the Gamma associated parameter.

#### Step 3b: Simulate movement trajectories

The function **simulate_trajectories()** simulates the movement trajectories and adds them in matrix form to the named list **m**. The matrix **m$trajectories** has a number of rows equivalent to the time partitions, and each column represents a unique individual.

```
method = "ICP"

load(paste0("parameters/true_parameters_", method,"_0", ".RData"))
load(paste0("models/unfitted_model_simulated_", method,"_0", "_no_data.RData"))

ni = round(6/m[["ns"]]) #number of individuals per species
releases = matrix(NA, ncol = 2, nrow = ni*m[["ns"]])
releases[, 1] = rep(1:m[["ns"]], ni)
releases[, 2] = rep(1, ni*m[["ns"]])
releases[, 2] = sample(seq_len(length(m$observation_effort$releases$location)),
                       size = ni*m[["ns"]], replace = TRUE)

colnames(releases) = c("sp", "release_event")

secondary_release = TRUE
max_dt = NULL

head(releases)
```

```
##      sp release_event
## [1,]  1             5
## [2,]  2            10
## [3,]  1             7
## [4,]  2             4
## [5,]  1            10
## [6,]  2             8
```

```
secondary_release = TRUE
max_dt = NULL

m = Jsmm::simulate_trajectories(m = m, pars = pars, releases = releases,
                                max_dt = max_dt)

m$trajectories
```

```
##       [,1] [,2] [,3] [,4] [,5] [,6]
##  [1,]   NA   NA   NA   NA   NA   NA
##  [2,]   NA   NA   NA   NA   NA   NA
##  [3,]   NA   NA   NA   NA   NA   NA
##  [4,]   NA   NA   NA  136   NA   NA
##  [5,]   87   NA   NA   89   NA   NA
##  [6,]   73   NA   NA  121   NA   NA
##  [7,]   64   NA  121  133   NA   NA
##  [8,]  119   NA   85  133   NA  104
##  [9,]  109   NA   85  124   NA   53
## [10,]  144   92   86  112   77   92
## [11,]   NA   NA   NA   NA   NA   NA
```

```
dim(m$trajectories)
```

```
## [1] 11  6
```

The simulated trajectories can be visualized by the function **plot_trajectories()**.

```r
# optional parameters for customizing plot:
  customize_plot = list()
  customize_plot$color_map = "#EEEEEE"
  customize_plot$labs_axis = c("X", "Y")
  customize_plot$lab_legend =  "Matrix"
  customize_plot$name_legend = "Habitat"
  customize_plot$color_polyb = "darkgrey"
  customize_plot$lwd_poly = 0.01
  customize_plot$site_col = "red"
  customize_plot$siteb_col = "darkred"
  customize_plot$sites_text_pos = c(-0.0, -0.0)
  customize_plot$site_text_size = 5
  customize_plot$panelb_color = "black"
  customize_plot$panelb_lwd = 1
  customize_plot$trajectory_color = c("black","blue") #rep("black", m[["ns"]])
  customize_plot$trajectory_lwd = 1
  plot_sp = 1:2

  Jsmm::plot_trajectories(m = m, customize_plot = customize_plot, plot_sp = plot_sp  )
```

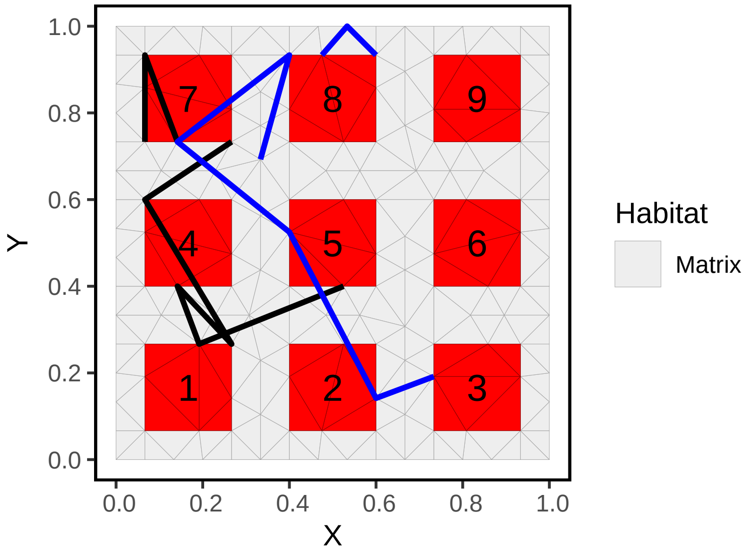

Figure 8: Simulated movement trajectories for three animals from each species. Black and blue colors correspond to the species 1 and 2, respectively.

### Demo simulated example ICP (covariates)

#### Example Description

Let's suppose that a researcher is interested in exploring whether the air temperature, altitude, and the species' body size influence the movement activity and vital dynamics for twenty focal species. In order to do this, an experiment is conducted over a landscape by setting nine trapping devices for eleven days, as shown in Fig. 1. Initially, 500 animals of each species are marked and released from different locations at different times. All capture sites are visited every day, starting from the second day until the end of the experiment. While the capture sites are visited, animals are marked, and captures are recorded. Secondary releases are not possible for this experiment.

For detailed information about the functions and their corresponding outcomes involved in our analyses, please refer to Appendix S1. Main paper.

#### Step 0: Loading the package

We start by loading the Jsmm package.

```
library(Jsmm)
```

#### Step 1a: Define model

For this example, we will use the same domain and observation effort lists as in the previous example.

##### Set covariate and phylogenetic relationships data

The Jsmm framework allows the inclusion of spatial, temporal, spatio-temporal covariates, and trait covariates. To do this, first, we are required to provide the data in a suitable format. The covariates are set within the named lists **cov_data** and **tr_data**.

For this example, we will include the spatial covariate altitude, the temporal covariate air temperature, a covariate for the observation process, and the body size as a species trait. Tables 5-7. show the model covariates characterizations.

```
cov_data = list()
altitude = domain[["triangulation"]][["node"]][, 2]
cov_data = list()
cov_data[["n_data"]] = data.frame(altitude)

head(cov_data[["n_data"]])
```

```
##    altitude
## 1 1.0000000
## 2 0.9333333
## 3 0.8666667
## 4 0.8000000
## 5 0.7333333
## 6 0.6666667
```

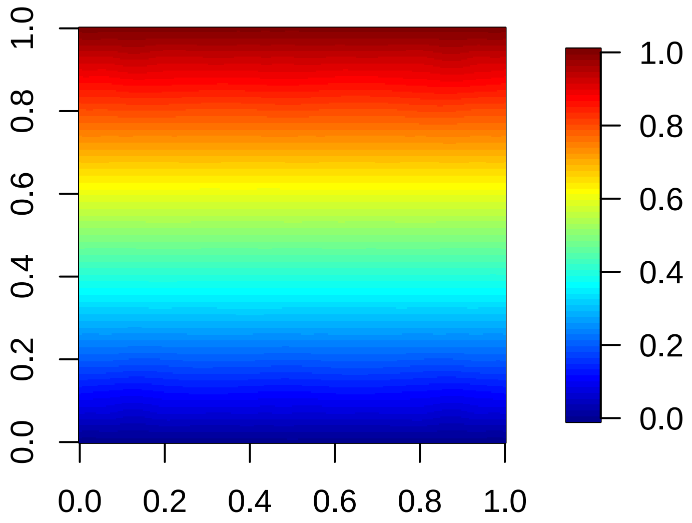

Figure 9: Altitude covariate. Measurements recorded at each node of the domain.

Table 5: Covariates for the movement parameters.

| Covariate name | Spatial | Temporal | Spatio-temporal | Type |
|---|---|---|---|---|
| altitude | node | No | No | Numeric |
| air_temperature | No | Yes | No | Numeric |

Table 6: Covariates for the observation parameter.

| Covariate name | Event | Time variation within events | Type |
|---|---|---|---|
| capture_intensity | Yes | No | Factor |

Table 7: Covariates for the species traits

| Covariate name | Type |
|---|---|
| size | Numeric |

The spatial data could be expressed as element or node resolution within the data frames **e_data** and **n_data**, respectively. The altitude covariate shown in Fig. 8., with values at each node, is set within the list **cov_data$n_data**.

The temporal data is set within the data frame **t_data**, and the times at which the observations were held are saved within the vector **t_data_time**. The covariate for the air temperature contains data for each day of the experiment. It is assumed that during the first six days of the experiment, cold temperatures occur, and during the rest of the days, the temperature increases. (See Fig. 9.).

```
t_data_times = seq(0, study_duration)
air_temperature = -1 + 2*(t_data_times > study_duration/2)

cov_data[["t_data"]] = data.frame(air_temperature)
cov_data[["t_data_times"]] = t_data_times

cov_data[["t_data"]]
```

```
##    air_temperature
## 1               -1
```

```
## 2                -1
## 3                -1
## 4                -1
## 5                -1
## 6                -1
## 7                 1
## 8                 1
## 9                 1
## 10                1
## 11                1
```

```
cov_data[["t_data_times"]]
```

```
##  [1]  0  1  2  3  4  5  6  7  8  9 10
```

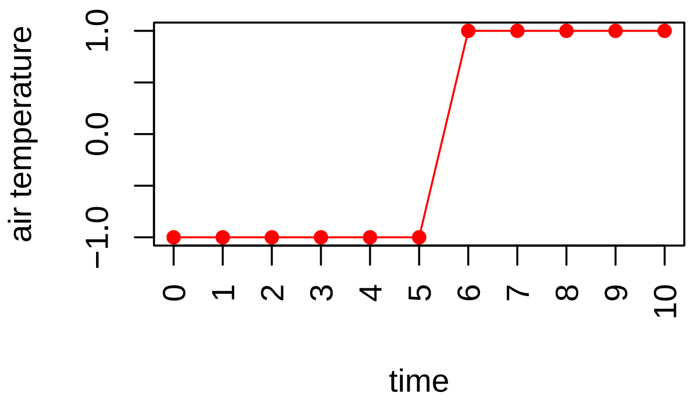

Figure 10: Air temperature covariate. Measurements recorded each day of the experiment (red dots).

Within the Jsmm, the covariates for the capture process are included in the data frame **obs_data** when there are no time variations within events, the rows correspond to the capture events, and the number of columns to the number of covariates. The covariate that characterizes capture events as high or low intensity is set as:

```
model_formula = list()
model_formula[["diffusion"]]          = ~ air_temperature
```

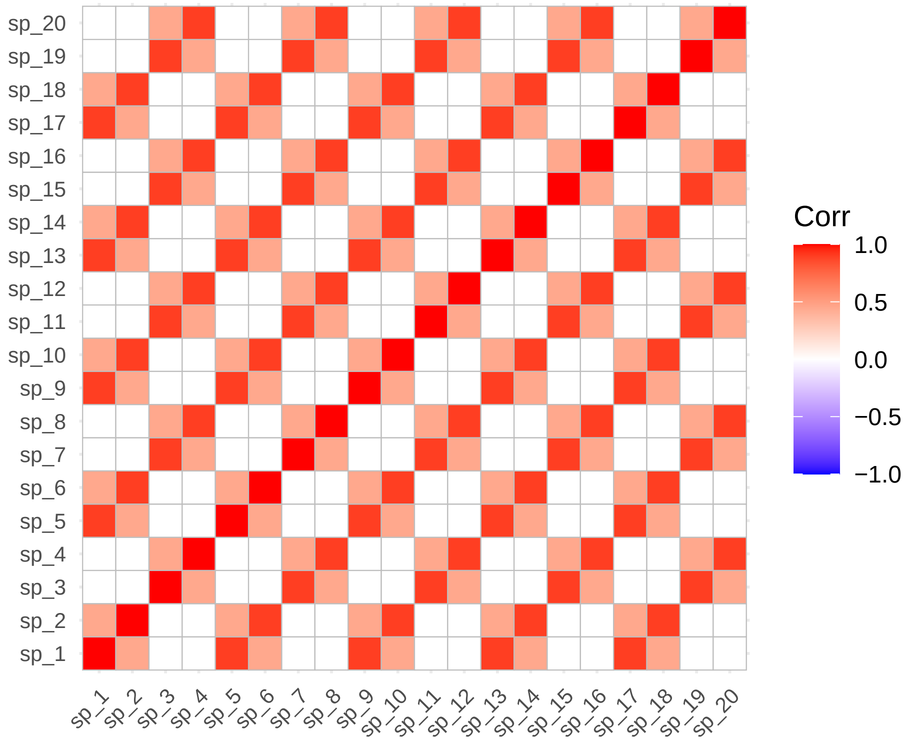

Figure 11: Phylogenetic correlation matrix.

```
model_formula[["advection_1"]]        = NULL
model_formula[["advection_2"]]        = NULL
model_formula[["mortality"]]          = ~ altitude
model_formula[["habitat_preference"]] = NULL
model_formula[["traits"]]             = ~ size
model_formula[["observation"]]        = ~ capture_intensity
```

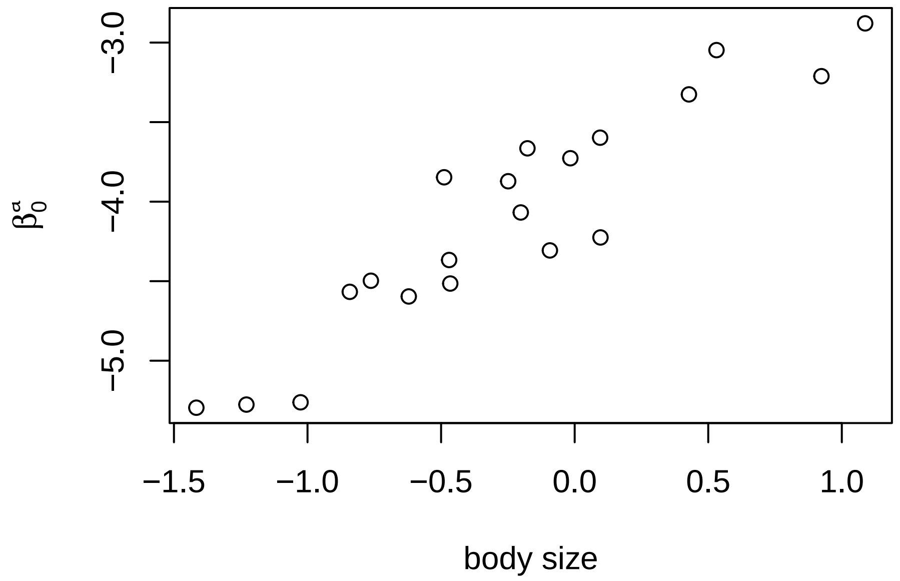

Figure 12: Diffusion intercept is influenced by the species body size.

Table 9: Model coeficients for diffusion, mortality and observation parameters for the twenty species.

| species | betaa0 | betaa1 | betam0 | betam1 | betao0 | betao1 |
|---|---|---|---|---|---|---|
| 1 | -3.872 | 1.314 | -2.378 | 1.124 | -2.205 | 0.846 |
| 2 | -3.847 | 1.211 | -2.645 | 1.297 | -2.359 | 0.532 |
| 3 | -5.295 | 1.001 | -3.033 | 1.234 | -1.480 | 0.496 |
| 4 | -5.262 | 1.289 | -2.162 | 1.308 | -1.480 | 0.542 |
| 5 | -3.047 | 1.597 | -2.725 | 1.195 | -2.682 | 0.963 |
| 6 | -3.326 | 1.548 | -2.127 | 1.051 | -2.497 | 1.423 |
| 7 | -4.307 | 1.056 | -3.017 | 1.658 | -1.004 | 0.467 |
| 8 | -3.211 | 1.526 | -2.442 | 1.177 | -1.645 | 0.307 |
| 9 | -4.567 | 1.721 | -2.571 | 1.330 | -2.281 | 0.769 |
| 10 | -3.665 | 1.511 | -2.104 | 1.141 | -2.805 | 1.319 |
| 11 | -4.068 | 0.994 | -3.108 | 1.331 | -1.498 | 0.861 |
| 12 | -4.515 | 1.491 | -2.439 | 1.616 | -1.470 | 0.543 |
| 13 | -3.727 | 1.119 | -2.335 | 0.963 | -2.239 | 0.507 |
| 14 | -4.497 | 1.451 | -2.114 | 0.799 | -2.385 | 1.390 |
| 15 | -4.225 | 1.272 | -2.776 | 1.322 | -1.353 | 0.264 |
| 16 | -5.276 | 1.986 | -2.446 | 1.210 | -1.595 | 0.724 |
| 17 | -3.598 | 0.989 | -2.540 | 0.866 | -2.154 | 0.774 |
| 18 | -4.367 | 1.880 | -2.201 | 0.711 | -2.641 | 1.144 |
| 19 | -2.879 | 0.995 | -2.664 | 1.410 | -1.736 | 0.379 |
| 20 | -4.596 | 1.729 | -2.399 | 1.355 | -1.727 | 0.743 |

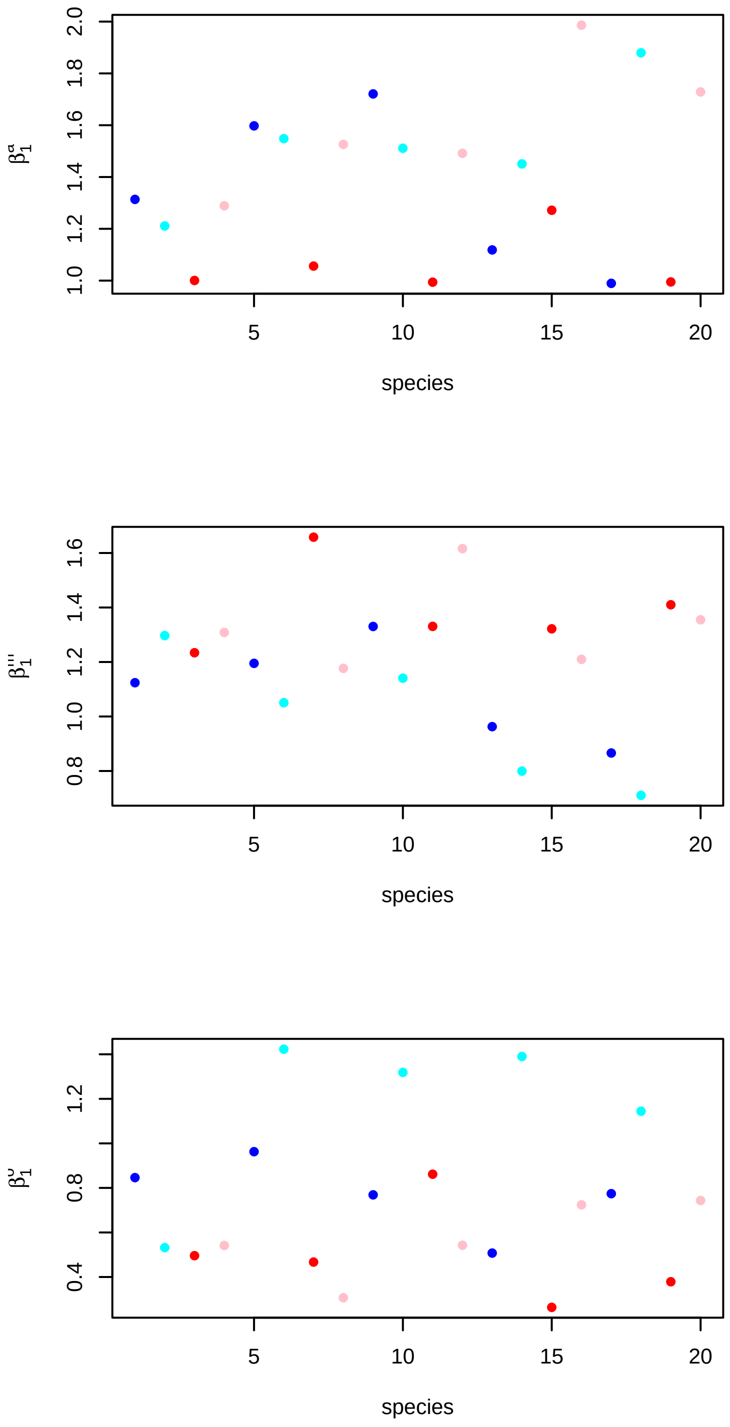

Figure 13: Model parameters for each of the twenty species. Upper to lower figures show diffusion, mortality, and observation dependence on the predictors; air temperature, altitude, and capture intensity, respectivelly.

```r
head(releases)
```

```
##      sp release_event
## [1,]  1             5
## [2,]  2            10
## [3,]  3             7
## [4,]  4             4
## [5,]  5            10
## [6,]  6             8
```

For example, the second row of the releases matrix indicates that one individual from species 2 was released at the release site number four at $t = 9$. (release event was declared on the **observation_effort** list).

As before, we can proceed to simulate capture-recapture data by using the function **simulate_captures(.)**.

```r
m = Jsmm::simulate_captures(m = m, pars = pars, releases = releases,
                      max_dt = max_dt, secondary_release = secondary_release)
```

```r
save(m, file = paste0("models/unfitted_model_simulated_", method, ".RData"))
```

Table 10: Numbers of recaptures per species.

| $sp_1$ | $sp_2$ | $sp_3$ | $sp_4$ | $sp_5$ | $sp_6$ | $sp_7$ | $sp_8$ | $sp_9$ | $sp_{10}$ | $sp_{11}$ | $sp_{12}$ | $sp_{13}$ | $sp_{14}$ | $sp_{15}$ | $sp_{16}$ | $sp_{17}$ | $sp_{18}$ | $sp_{19}$ | $sp_{20}$ |
|---|---|---|---|---|---|---|---|---|---|---|---|---|---|---|---|---|---|---|---|
| 185 | 94 | 180 | 237 | 139 | 166 | 210 | 236 | 91 | 133 | 148 | 211 | 118 | 104 | 161 | 182 | 122 | 130 | 141 | 173 |

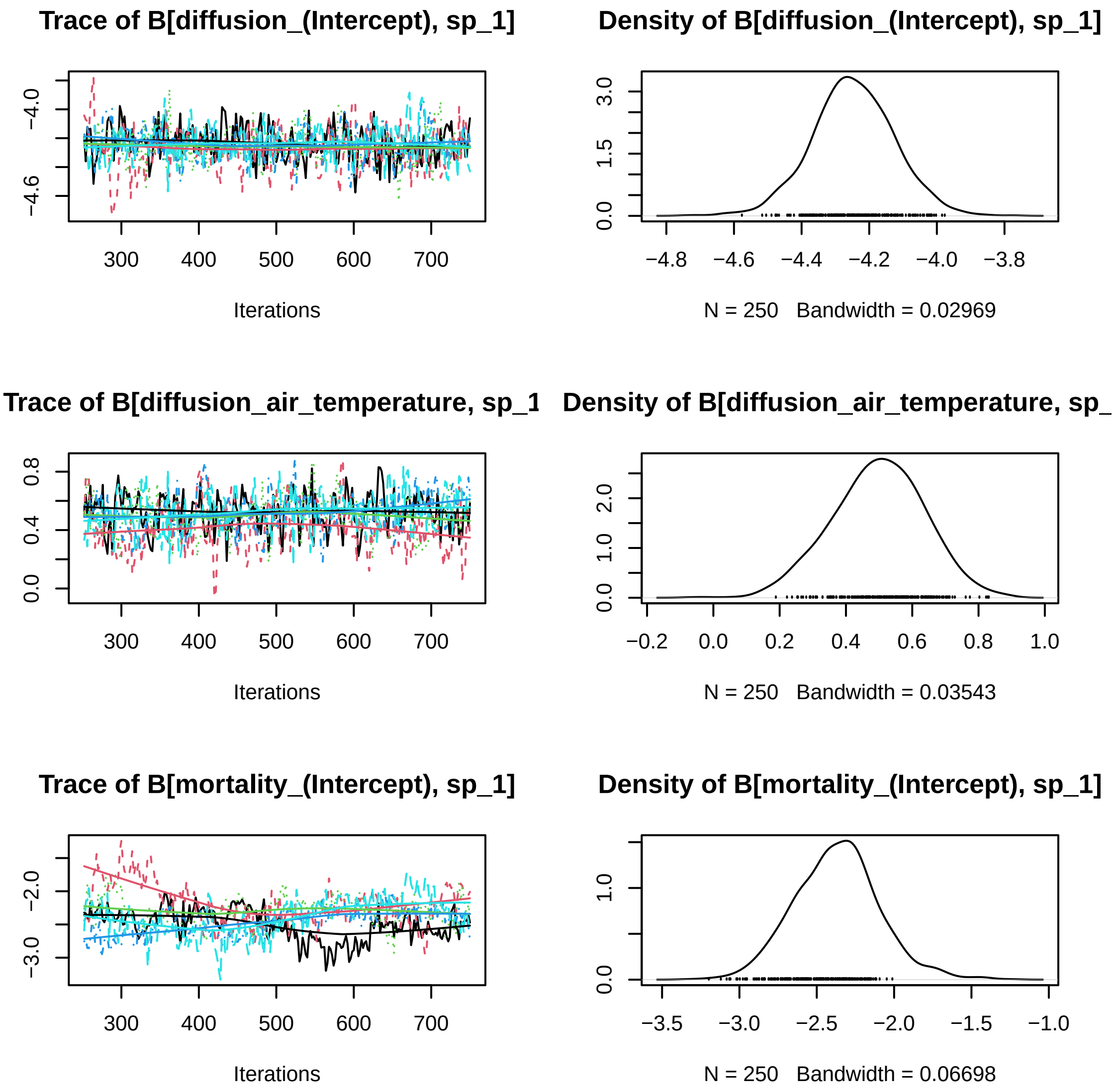

Figure 14: First species trace plots model parameters Beta., five MCMC chains.

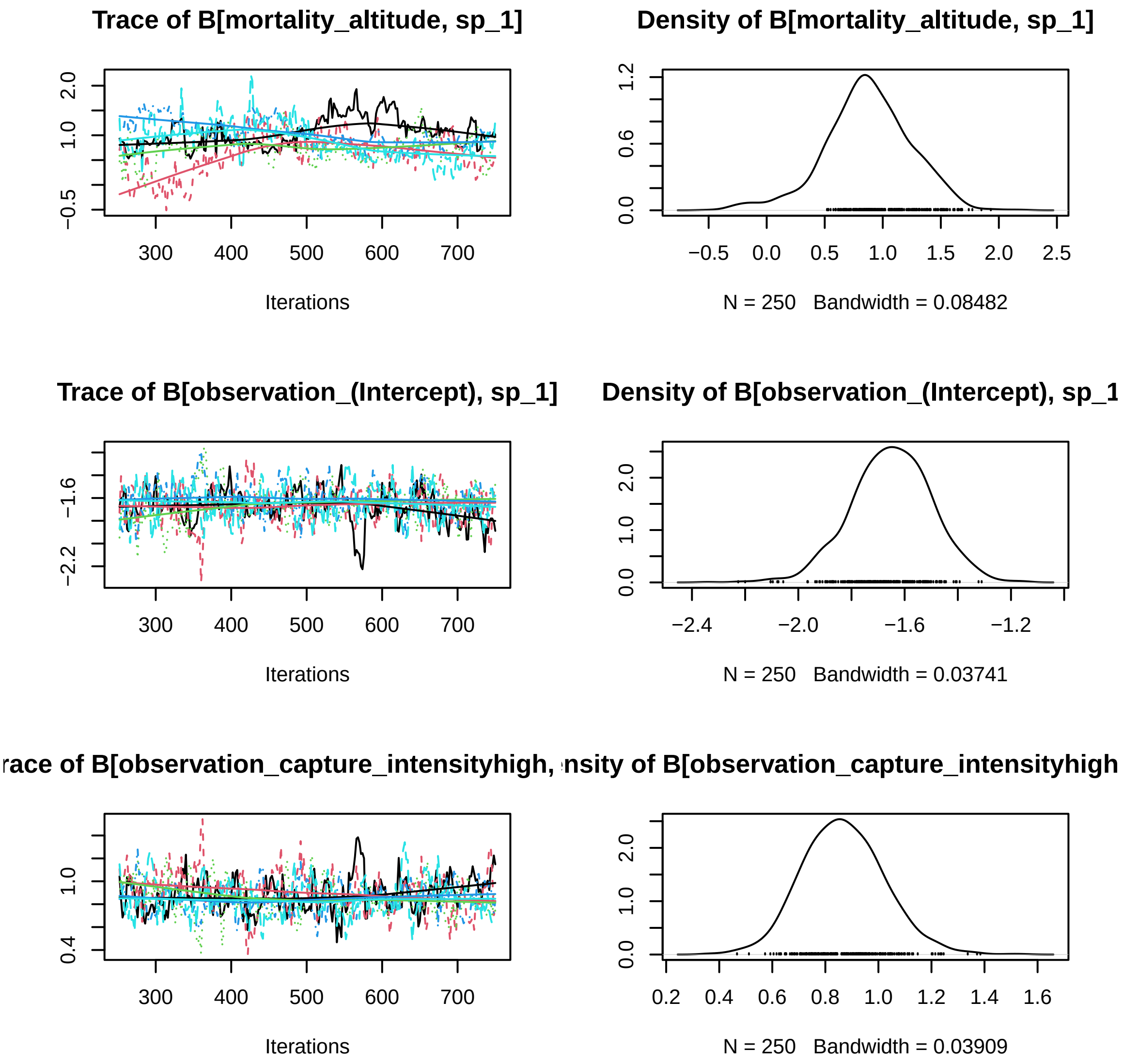

Figure 15: First species trace plots model parameters Beta., five MCMC chains.

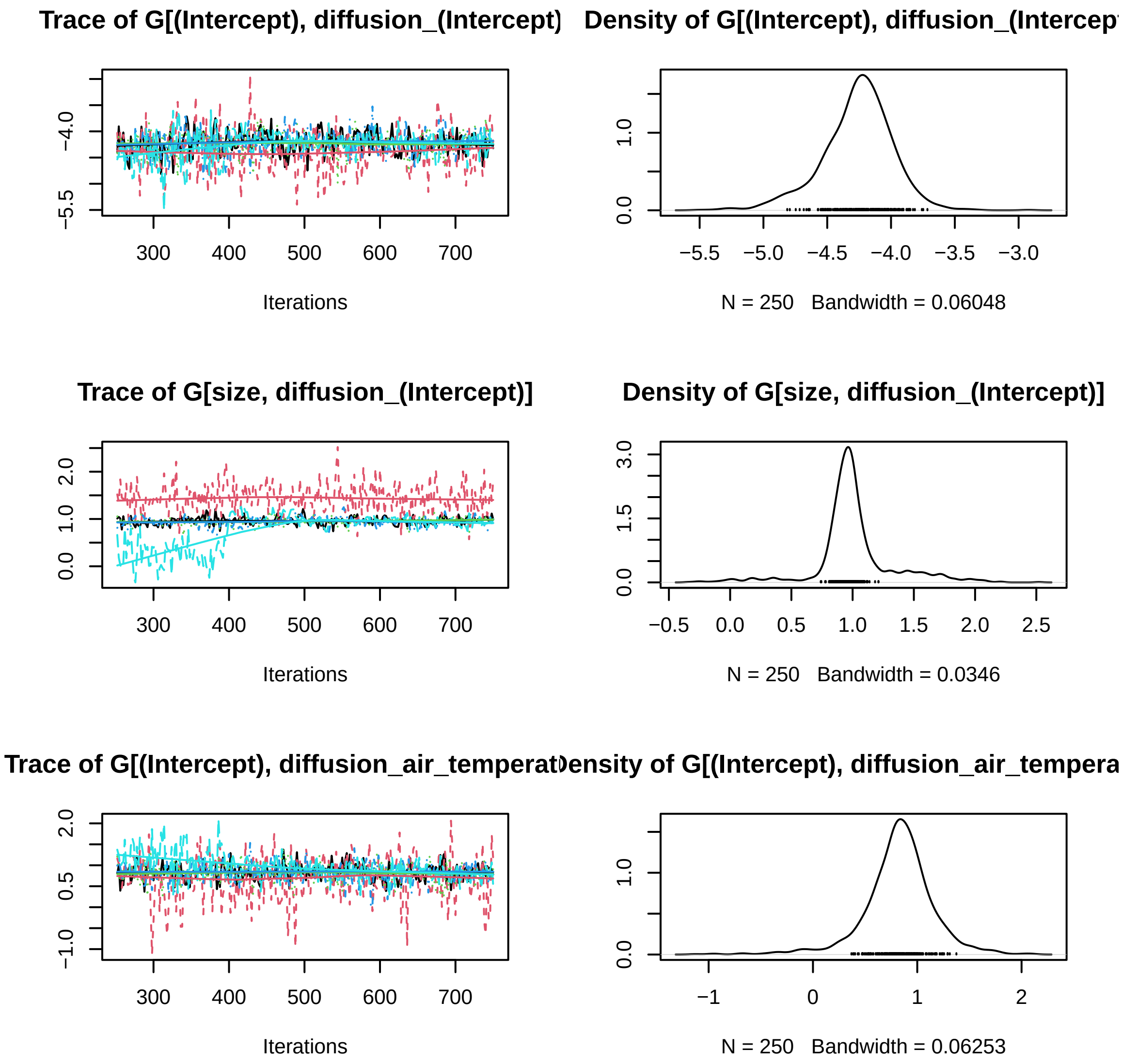

Figure 16: First species trace plots, five MCMC chains Gamma.

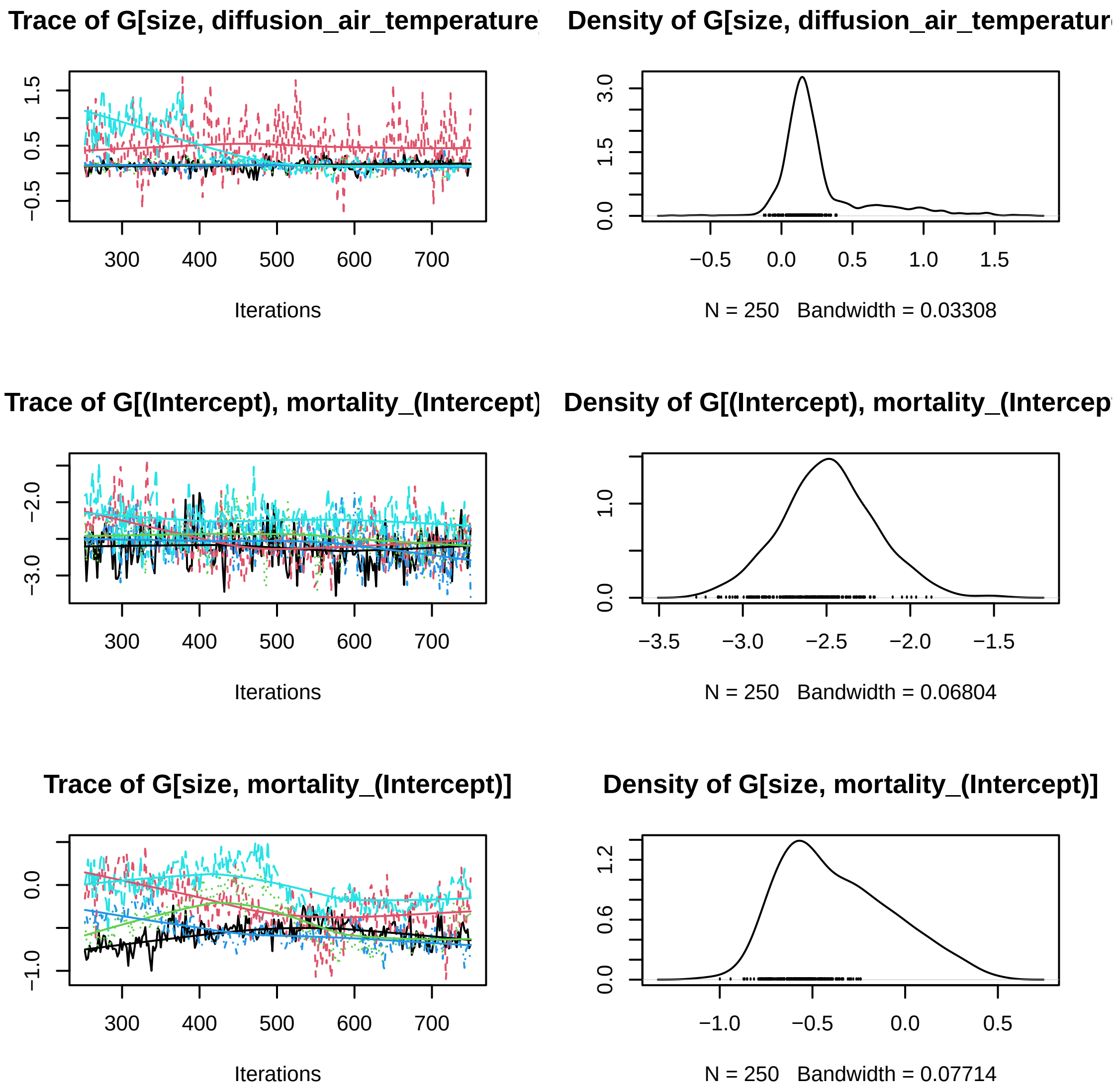

Figure 17: First species trace plots, five MCMC chains Gamma.

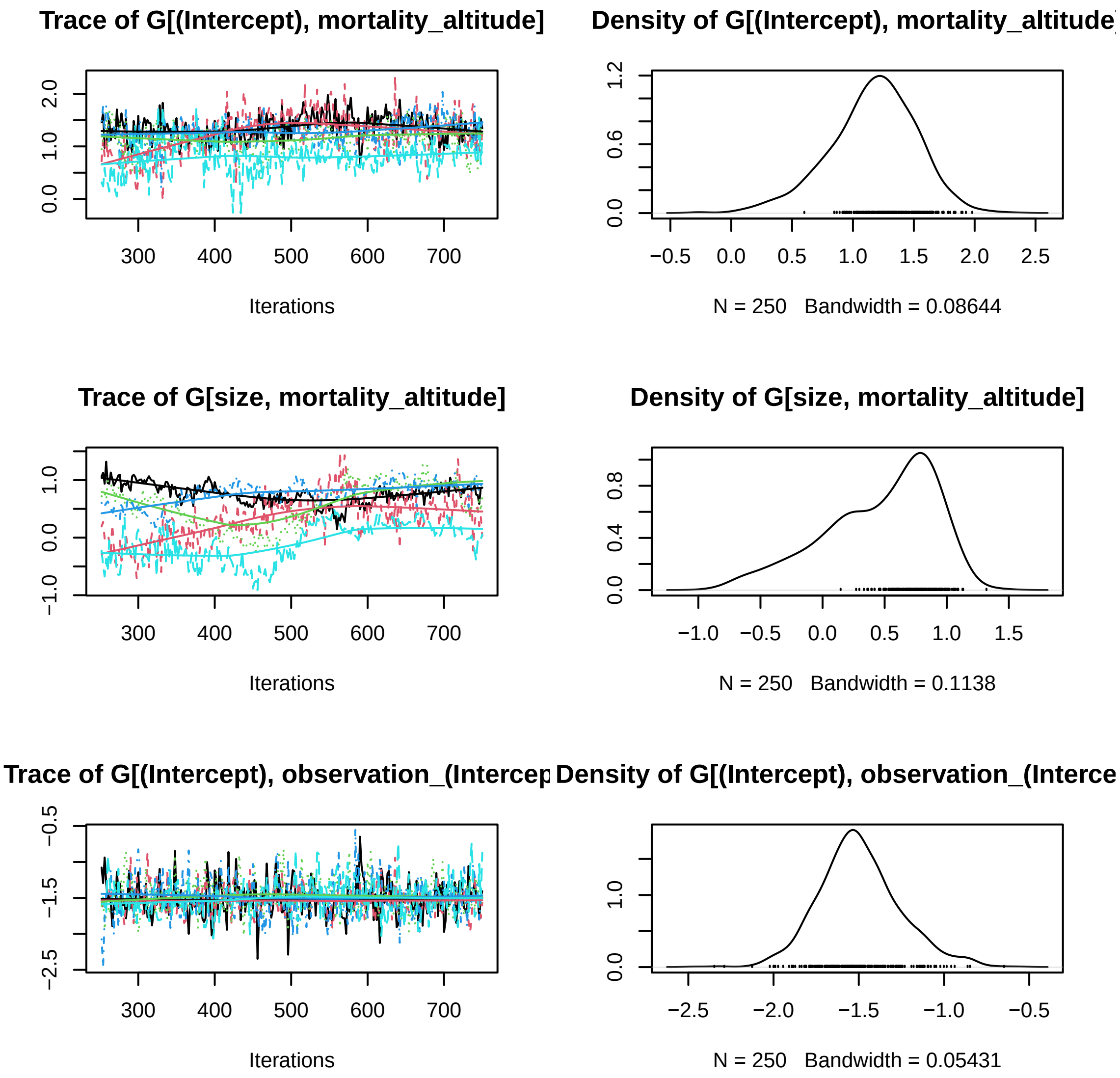

Figure 18: First species trace plots, five MCMC chains Gamma.

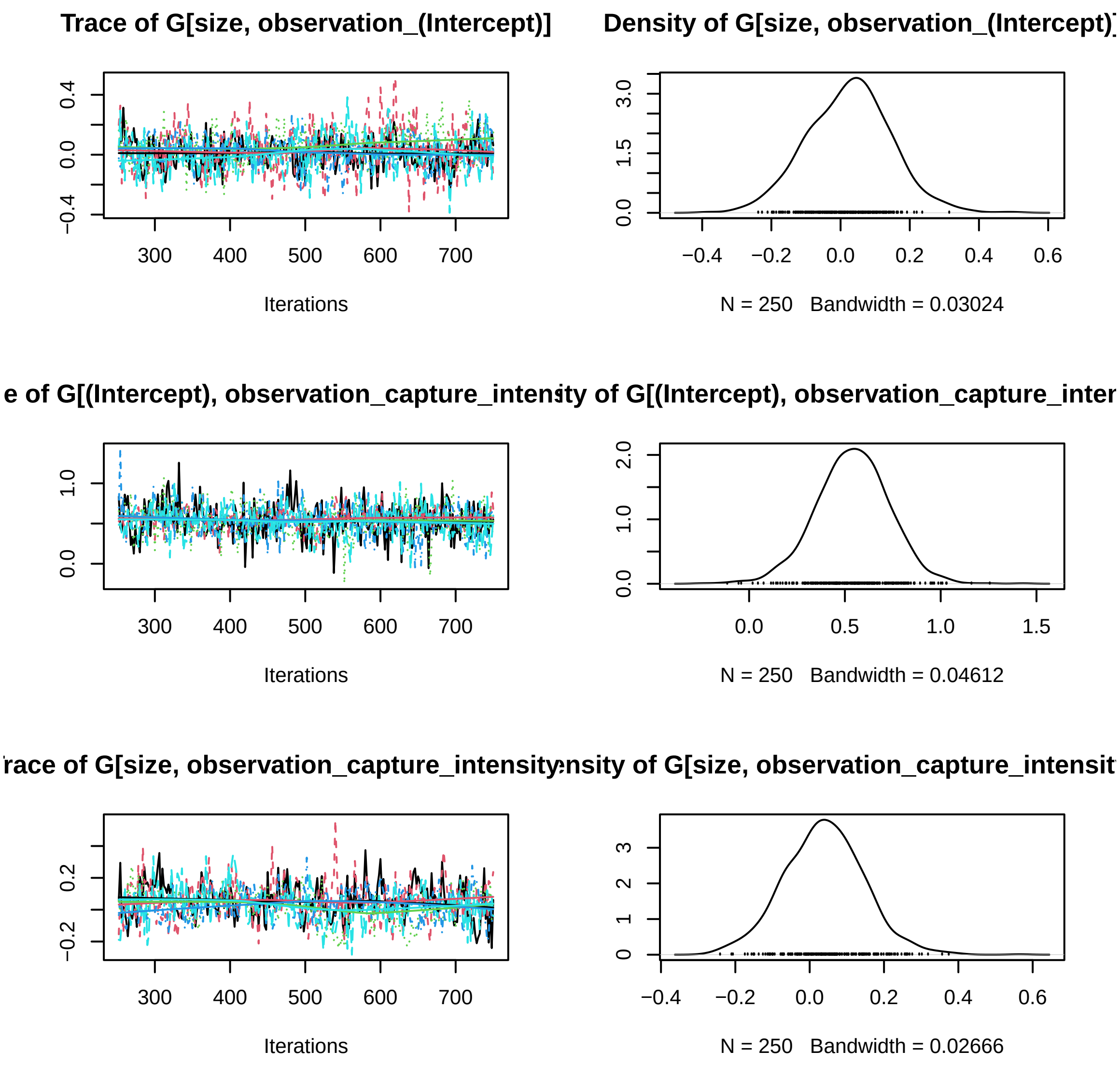

Figure 19: First species trace plots, five MCMC chains Gamma.

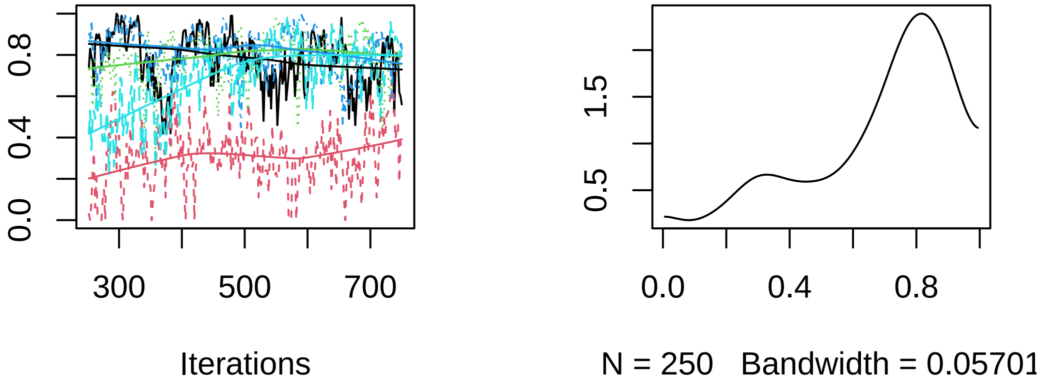

Figure 20: First species trace plots, five MCMC chains Rho.

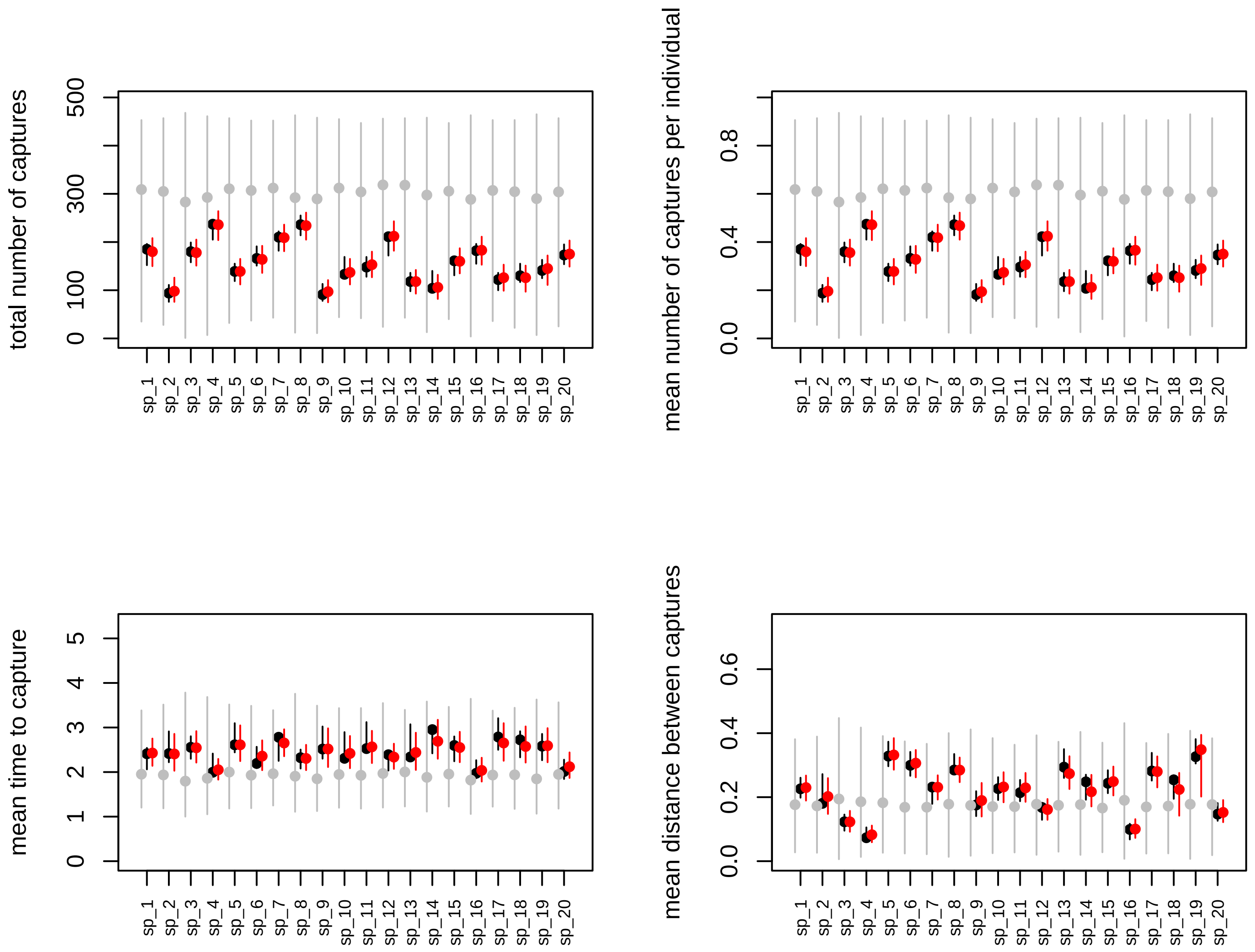

Figure 21: Posterior predictive data generated with 1000 datasets. Black, red, and grey colors represent the datasets generated with true, posterior, and prior parameters values.

```
ns     = m[["ns"]]
nt     = m[["nt"]]

nprior = npost
prior  = list()

for(repl in 1:nprior){
  prior[[repl]] = simulate_from_prior(m)
}

load("parameters/true_parameters_ICP.RData")

par(mfrow=c(3,2))

cc = 0
for(va in vars){
  nv = m[["nc"]][[va]]
  for(i in 1:nv){
    cc = cc + 1

    re.pri = matrix(ncol = ns, nrow = nprior)
    re.pri.G = matrix(ncol = nt, nrow = nprior)

    for(repl in 1:nprior){
      re.pri[repl, ]   = prior[[repl]][["Beta"]][[va]][, i]
      re.pri.G[repl, ] = prior[[repl]][["Gamma"]][, cc]
    }

    re.post   = matrix(ncol = ns, nrow = npost)
    re.post.G = matrix(ncol = nt, nrow = nprior)

    for(repl in 1:npost){
      re.post[repl, ]   = post[[repl]][["Beta"]][[va]][, i]
      re.post.G[repl, ] = post[[repl]][["Gamma"]][, cc]
    }

    ymin = min(quantile(re.pri, probs = 0.05), min(re.post))
    ymax = max(quantile(re.pri, probs = 0.95), max(re.post))

    if(showTrueValues){
      ymin = min(ymin, pars$Beta[[va]][, i])
      ymax = max(ymax, pars$Beta[[va]][, i])
    }

    colnames(re.pri) = m$sp_names
    re.pri  = re.pri[, sel]
    re.post = re.post[, sel]

    boxplot(re.pri, outline = FALSE, at = 3*(1:ns)-0.75, ylim = c(ymin, ymax),
            main = m[["parNames"]][cc], las = 2, cex.axis = 0.7,
            xlim = c(2, 3*ns + 2),
            ylab = "parameter value", col = "grey", xaxt = "n")

    axis(1, at = 3*(1:m$ns), label = colnames(re.pri), las = 2, cex.axis = 0.7)

    boxplot(re.post, outline = FALSE, at = 3*(1:ns) + 0.75, col = "pink", add = TRUE,
```

```
            xaxt = "n", yaxt = "n")

    abline(h = 0)

    if(showTrueValues){
      for(j in 1:m[["ns"]]){
        lines(x = c(3*j, 3*j + 1.5), y = c(pars$Beta[[va]][j, i], pars$Beta[[va]][j, i]),
              lwd = 2, col = "red")
      }
    }

    ymin = min(quantile(re.pri.G, probs = 0.05), min(re.post.G))
    ymax = max(quantile(re.pri.G, probs = 0.95), max(re.post.G))

    if(showTrueValues){
      ymin = min(ymin, pars$Gamma[, cc])
      ymax = max(ymax, pars$Gamma[, cc])
    }

    colnames(re.pri.G) = colnames(m$X[["trait"]])

    boxplot(re.pri.G, outline = FALSE, at = 3*(1:nt), ylim = c(ymin, ymax),
            main = m[["parNames"]][cc], las = 2, cex.axis = 0.7,
            xlim = c(2, 3*nt + 2),
            ylab = "parameter value", col = "grey")

    boxplot(re.post.G, outline = FALSE, at = 3*(1:nt) + 1.5, col = "pink", add = TRUE,
            xaxt = "n", yaxt = "n")

    abline(h = 0)

    if(showTrueValues){
      for(j in 1:m[["nt"]]){
        lines(x = c(3*j, 3*j + 1.5), y = c(pars$Gamma[j, cc], pars$Gamma[j, cc]),
              lwd = 2, col = "red")
      }
    }
  }
}
```

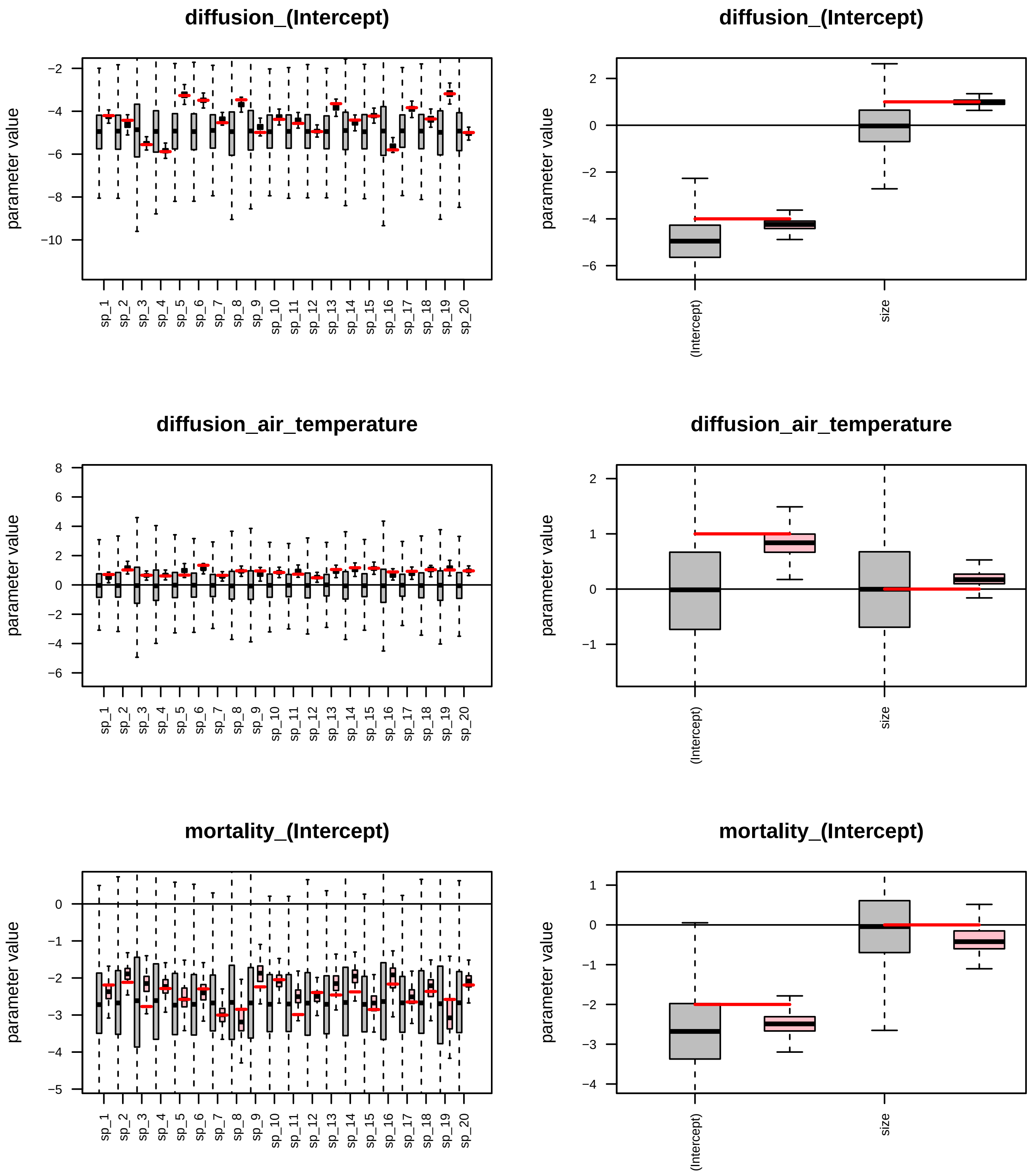

Figure 22: Posterior credible intervals for the model parameters. Left panels show the posteriors for each of the model parameters (pink color), their corresponding priors (Grey), and the parameter true value (Red). Right panels show the corresponding values for the Gamma associated parameter.

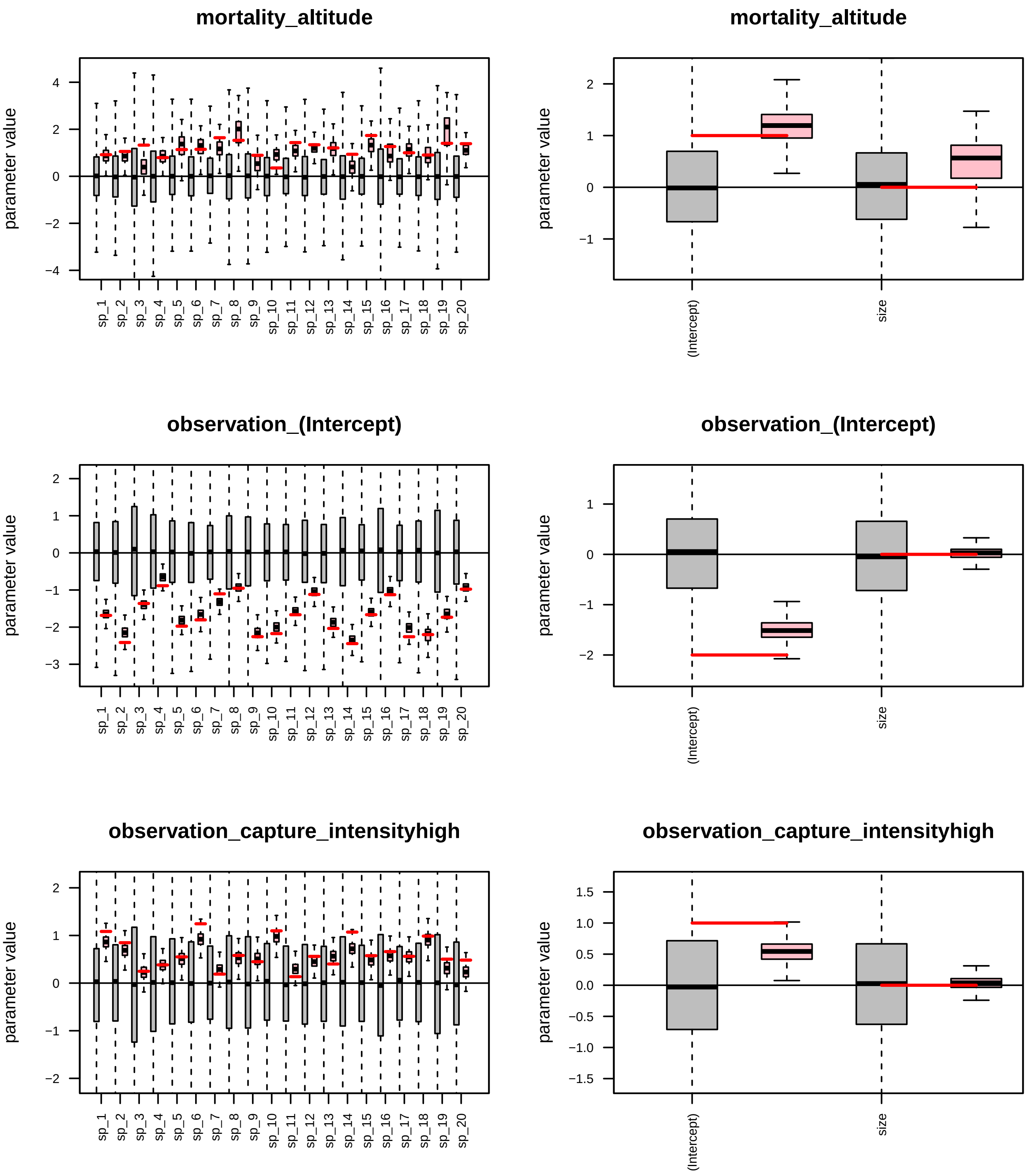

Figure 23: Posterior credible intervals for the model parameters. Left panels show the posteriors for each of the model parameters (pink color), their corresponding priors (Grey), and the parameter true value (Red). Right panels show the corresponding values for the Gamma associated parameter.

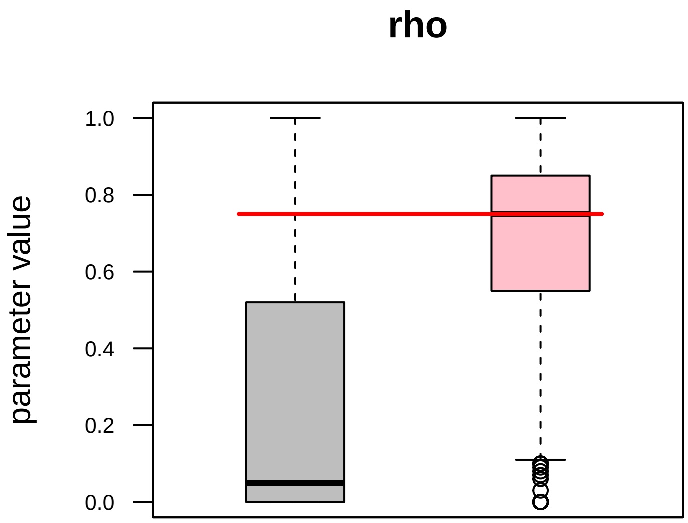

Figure 24: Posterior credible intervals for the associated rho parameter. Posterior, prior, and true value are represented by pink, grey, and red colors, respectively.

### Demo simulated example CCP (covariates)

#### Example Description

Let's suppose that a researcher is interested in exploring whether the air temperature, altitude, and the species' body size influence the movement activity and vital dynamics for twenty focal species. In order to do this, an experiment is conducted over a landscape by setting nine trapping devices for eleven days, as shown in Fig. 1. Initially, 500 animals of each species are marked and released from different locations at different times. By simplicity, let's suppose that all the traps are kept open, allowing captures every day, visited by the end of each day, and captures are recorded. Note that within our framework, the user can customize the time intervals in which the traps allow captures. Secondary releases are not possible for this experiment.

For detailed information about the functions and their corresponding outcomes involved in our analyses, please refer to Appendix S1. Main paper.

#### Step 0: Loading the package

We start by loading the Jsmm package.

```
library(Jsmm)
```

#### Step 1a: Define model

We set the same elements as before, except the **observation_effort** list. For this third example, the capture process has been set as continuous in the list **observation_effort$method**. In addition, **observation_effort$captures$time**, which indicates the times in which captures occur, is now set as a matrix with rows representing the capture events, and the first and second columns indicating the time interval $[t_0, t_1]$ in which the capture site starts and ends capturing individuals. The observation effort can be visualized in Fig. 25.

The script for setting the **observation_effort** object is:

For example, capture events 10 to 15 occur at times

```
observation_effort$captures$time[10:15, ]
```

```
##      t_0 t_1
## [1,]   1   2
## [2,]   1   2
## [3,]   1   2
## [4,]   1   2
## [5,]   1   2
## [6,]   1   2
```

Information corresponding to the capture event $c = 12$, is retrieved as

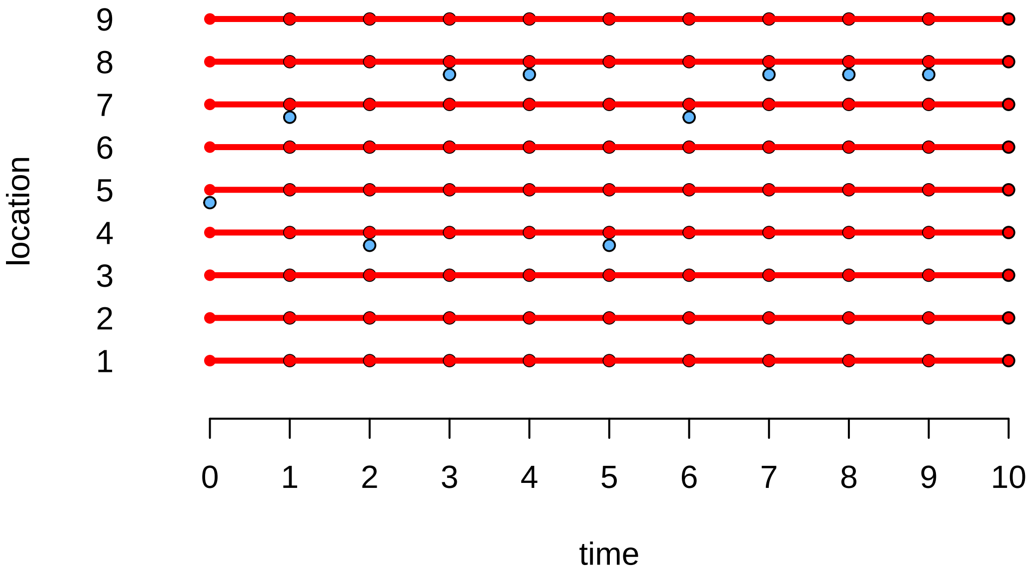

Figure 25: Observation effort scheme for CCP capture process. Times in which releases and captures are possible are represented by blue and red colors, respectively.

```
observation_effort$captures$location[12]
```

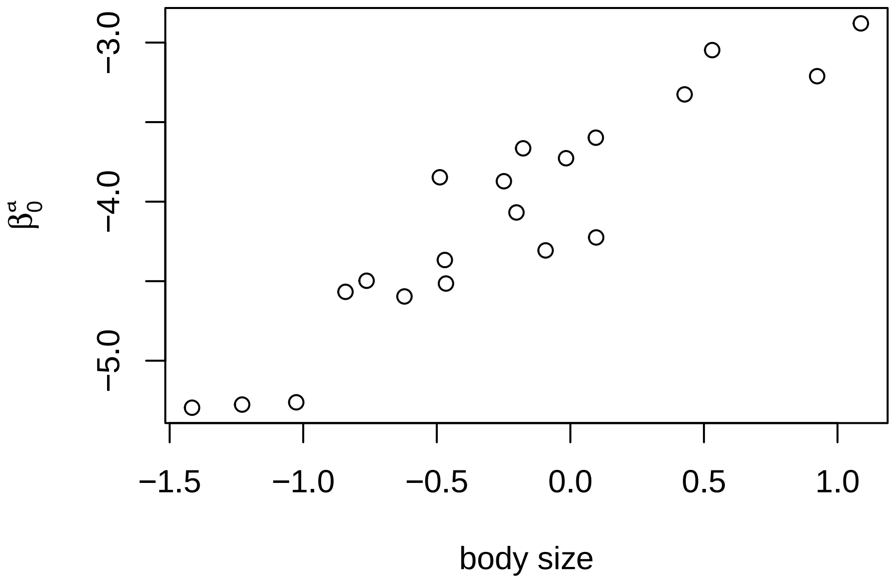

Figure 26: Diffusion intercept is influenced by the species body size.

Table 13: Model coeficients for diffusion, mortality and observation parameters for the twenty species.

| species | betaa0 | betaa1 | betam0 | betam1 | betao0 | betao1 |
|---|---|---|---|---|---|---|
| 1 | -3.872 | 1.314 | -2.378 | 1.124 | -2.205 | 0.846 |
| 2 | -3.847 | 1.211 | -2.645 | 1.297 | -2.359 | 0.532 |
| 3 | -5.295 | 1.001 | -3.033 | 1.234 | -1.480 | 0.496 |
| 4 | -5.262 | 1.289 | -2.162 | 1.308 | -1.480 | 0.542 |
| 5 | -3.047 | 1.597 | -2.725 | 1.195 | -2.682 | 0.963 |
| 6 | -3.326 | 1.548 | -2.127 | 1.051 | -2.497 | 1.423 |
| 7 | -4.307 | 1.056 | -3.017 | 1.658 | -1.004 | 0.467 |
| 8 | -3.211 | 1.526 | -2.442 | 1.177 | -1.645 | 0.307 |
| 9 | -4.567 | 1.721 | -2.571 | 1.330 | -2.281 | 0.769 |
| 10 | -3.665 | 1.511 | -2.104 | 1.141 | -2.805 | 1.319 |
| 11 | -4.068 | 0.994 | -3.108 | 1.331 | -1.498 | 0.861 |
| 12 | -4.515 | 1.491 | -2.439 | 1.616 | -1.470 | 0.543 |
| 13 | -3.727 | 1.119 | -2.335 | 0.963 | -2.239 | 0.507 |
| 14 | -4.497 | 1.451 | -2.114 | 0.799 | -2.385 | 1.390 |
| 15 | -4.225 | 1.272 | -2.776 | 1.322 | -1.353 | 0.264 |
| 16 | -5.276 | 1.986 | -2.446 | 1.210 | -1.595 | 0.724 |
| 17 | -3.598 | 0.989 | -2.540 | 0.866 | -2.154 | 0.774 |
| 18 | -4.367 | 1.880 | -2.201 | 0.711 | -2.641 | 1.144 |
| 19 | -2.879 | 0.995 | -2.664 | 1.410 | -1.736 | 0.379 |
| 20 | -4.596 | 1.729 | -2.399 | 1.355 | -1.727 | 0.743 |

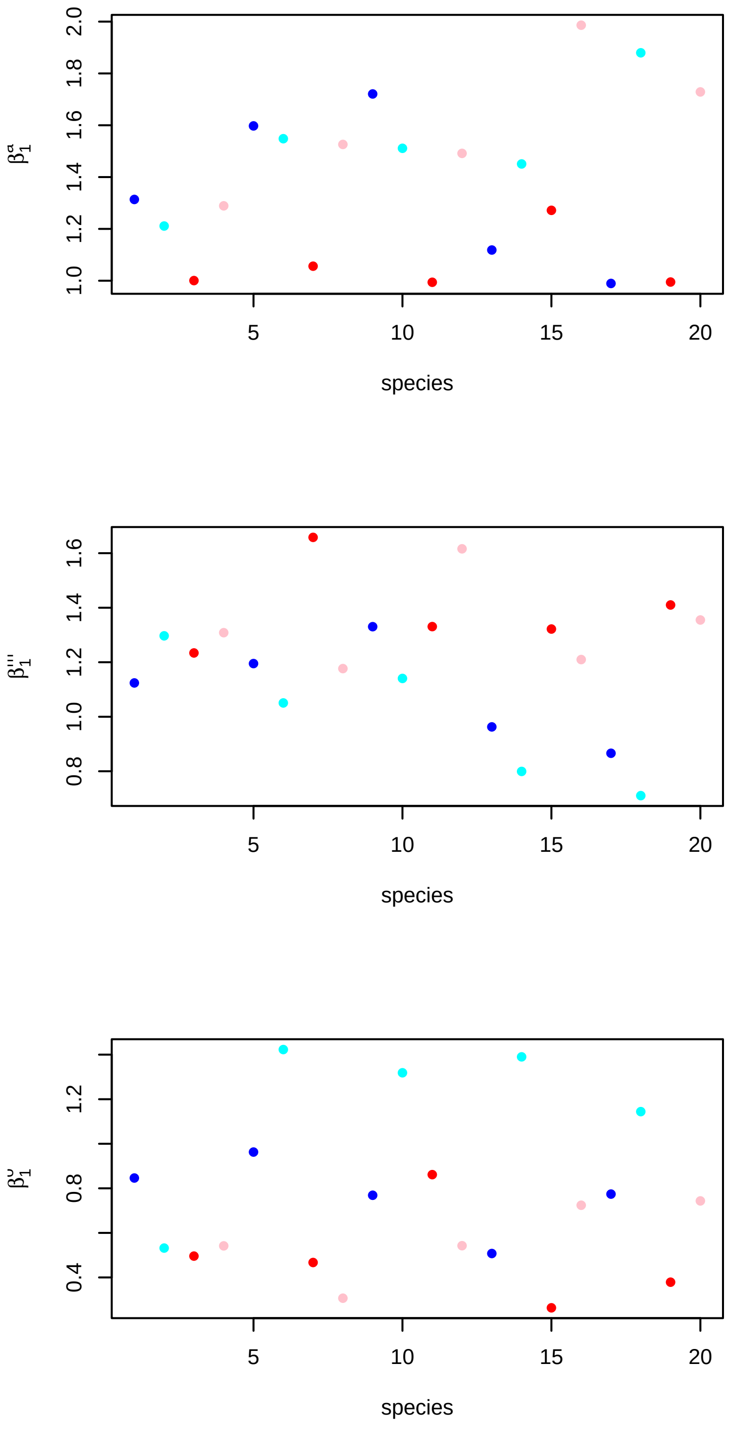

Figure 27: Model parameters for each of the twenty species. Upper to lower figures show diffusion, mortality, and observation dependence on the predictors; air temperature, altitude, and capture intensity, respectively.

```r
releases[, 1] = rep(1:m[["ns"]], ni)
releases[, 2] = rep(1, ni*m[["ns"]])
releases[, 2] = sample(seq_len(length(m$observation_effort$releases$location)),
                       size = ni*m[["ns"]], replace = TRUE)

colnames(releases) = c("sp", "release_event")

secondary_release = FALSE
max_dt = NULL

head(releases)
```

```
##      sp release_event
## [1,]  1             5
## [2,]  2            10
## [3,]  3             7
## [4,]  4             4
## [5,]  5            10
## [6,]  6             8
```

For example, the second row of the releases matrix indicates that one individual from species 2 was released at the release site number four at $t = 9$. (release event was declared on the **observation_effort** list).

As before, we can proceed to simulate capture-recapture data by using the function **simulate_captures(.)**.

```r
m = Jsmm::simulate_captures(m = m, pars = pars, releases = releases,
                     max_dt = max_dt, secondary_release = secondary_release)
```

```r
save(m, file = paste0("models/unfitted_model_simulated_", method, ".RData"))
```

```r
Jsmm::model_summary(m)
```

```
## Domain: nele = 258 | nnod = 160
## Capture method: CCP | Number release sites: 4 | Number capture sites: 9
## Number of time partitions: 11
## ----------------------------------------
## Model formulas:
##     Diffusion ~ air_temperature | nc = 2
##     Mortality ~ altitude | nc = 2
##     Traits ~ size
##     Observation ~ capture_intensity | nc = 2
## ----------------------------------------
## Number of species: 20
## Releases Data:
##      10000 unique releases
## Captures Data:
##      4866 captures
```

Table 14: Numbers of recaptures per species.

| $sp_1$ | $sp_2$ | $sp_3$ | $sp_4$ | $sp_5$ | $sp_6$ | $sp_7$ | $sp_8$ | $sp_9$ | $sp_{10}$ | $sp_{11}$ | $sp_{12}$ | $sp_{13}$ | $sp_{14}$ | $sp_{15}$ | $sp_{16}$ | $sp_{17}$ | $sp_{18}$ | $sp_{19}$ | $sp_{20}$ |
|---|---|---|---|---|---|---|---|---|---|---|---|---|---|---|---|---|---|---|---|
| 289 | 111 | 255 | 366 | 184 | 288 | 308 | 375 | 135 | 245 | 207 | 339 | 164 | 174 | 217 | 323 | 162 | 201 | 211 | 312 |

```r
mname = "simulated_CCP"

thin    = 2
samples = 250
nChains = 5

load(paste0(paste0("models/fitted_model_", mname,
                   "_samples_", samples, "_thin_", thin, "_nChains_", nChains, ".RData")))

mpost = Jsmm::convert_to_coda_object(m)

summary(coda::gelman.diag(mpost$Beta, multivariate = FALSE)$psrf)
```

```
##    Point est.       Upper C.I.
##  Min.   : 1.003   Min.   : 1.013
##  1st Qu.: 1.032   1st Qu.: 1.084
##  Median : 1.090   Median : 1.227
##  Mean   : 2.566   Mean   : 4.029
##  3rd Qu.: 1.286   3rd Qu.: 1.690
##  Max.   :39.149   Max.   :73.189
```

#### Step 2c: Simulate prior and posterior predictive data

The script below generates 1000 datasets for the CCP simulated example. The main purpose of generating predictive data is exploring how the fitted model behaves in time and space compared to the original data. This method helps to infer if the fitted parameters provide a good approximation of the real data.

```r
mname = "simulated_CCP"

for(nrepl in c(1000)){
  showTrueValues = TRUE

  thin    = 2
  samples = 250
  nChains = 5

  load(paste0(paste0("models/fitted_model_", mname, "_samples_", samples, "_thin_", thin,
                     "_nChains_", nChains, ".RData")))

  post = m[["postList"]]

  if(showTrueValues){
    types = c("true", "prior", "posterior")
    load("parameters/true_parameters_CCP.RData")
    true_pars = pars
  }else{
    types =  c("prior", "posterior")
  }

  for(type in types){
    PPcaptures = list()

    for(repl in 1:nrepl){
      if(type == "true"){
        pars = true_pars
      }
```

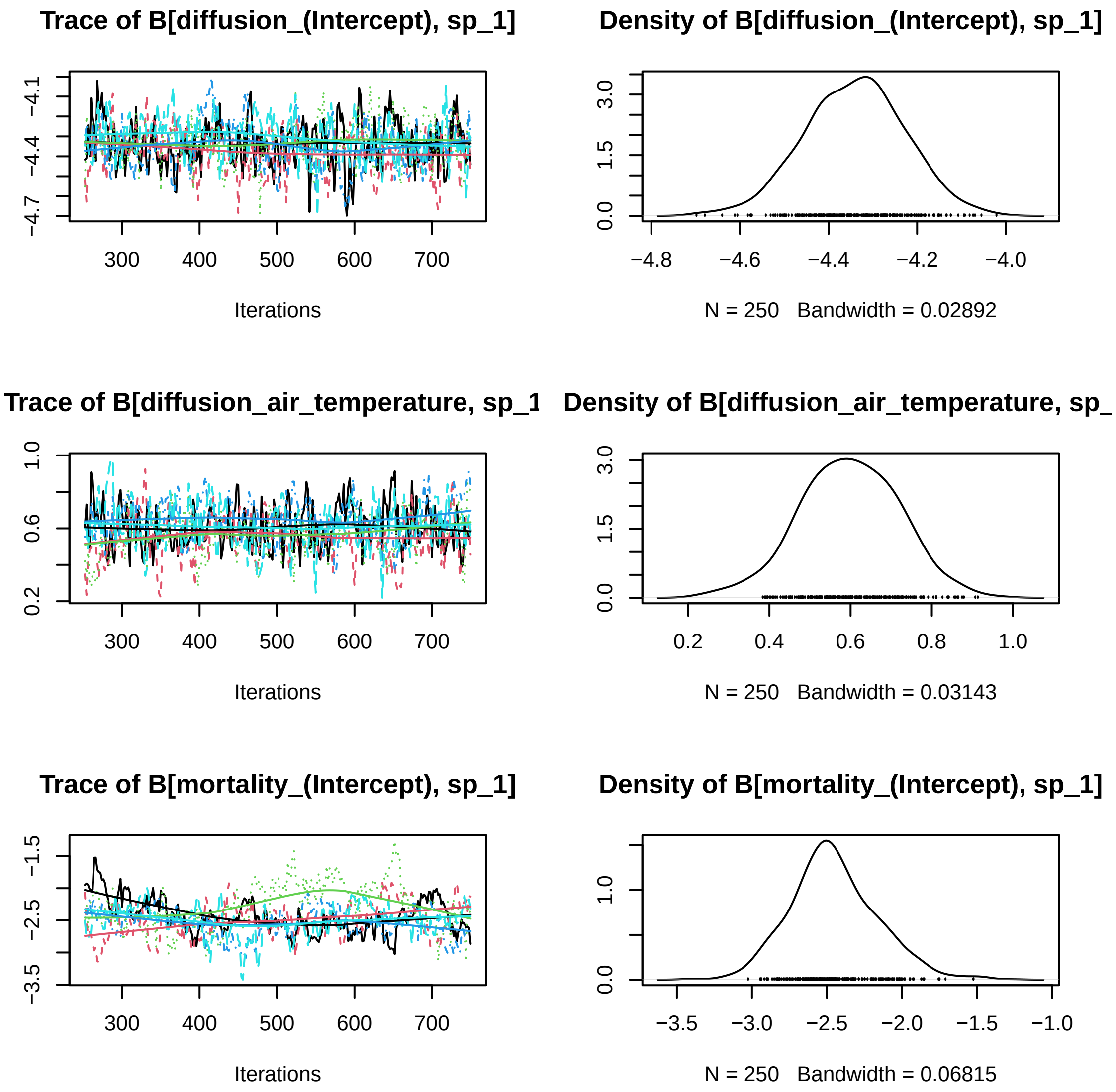

Figure 28: First species trace plots model parameters Beta., five MCMC chains.

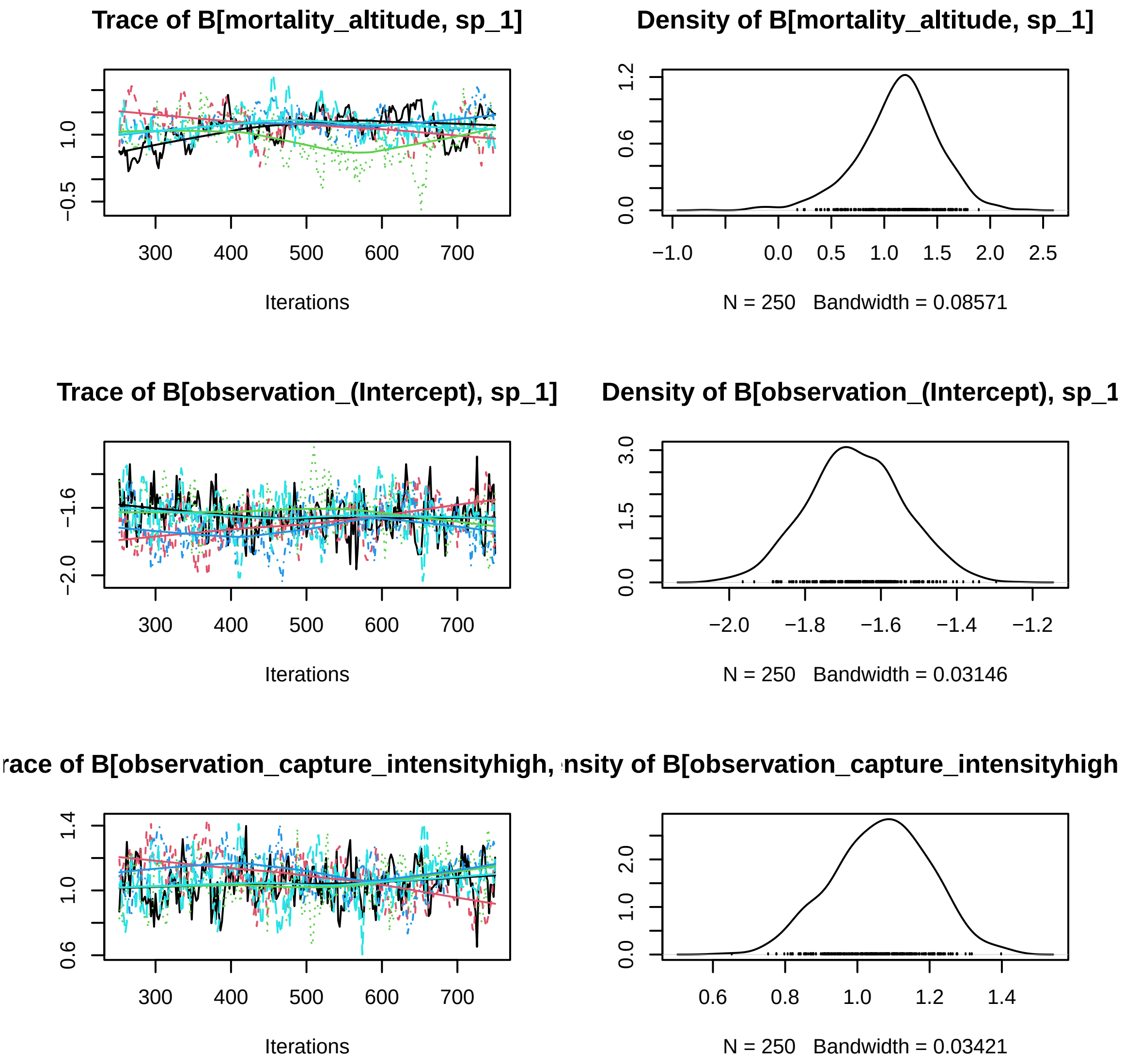

Figure 29: First species trace plots model parameters Beta., five MCMC chains.

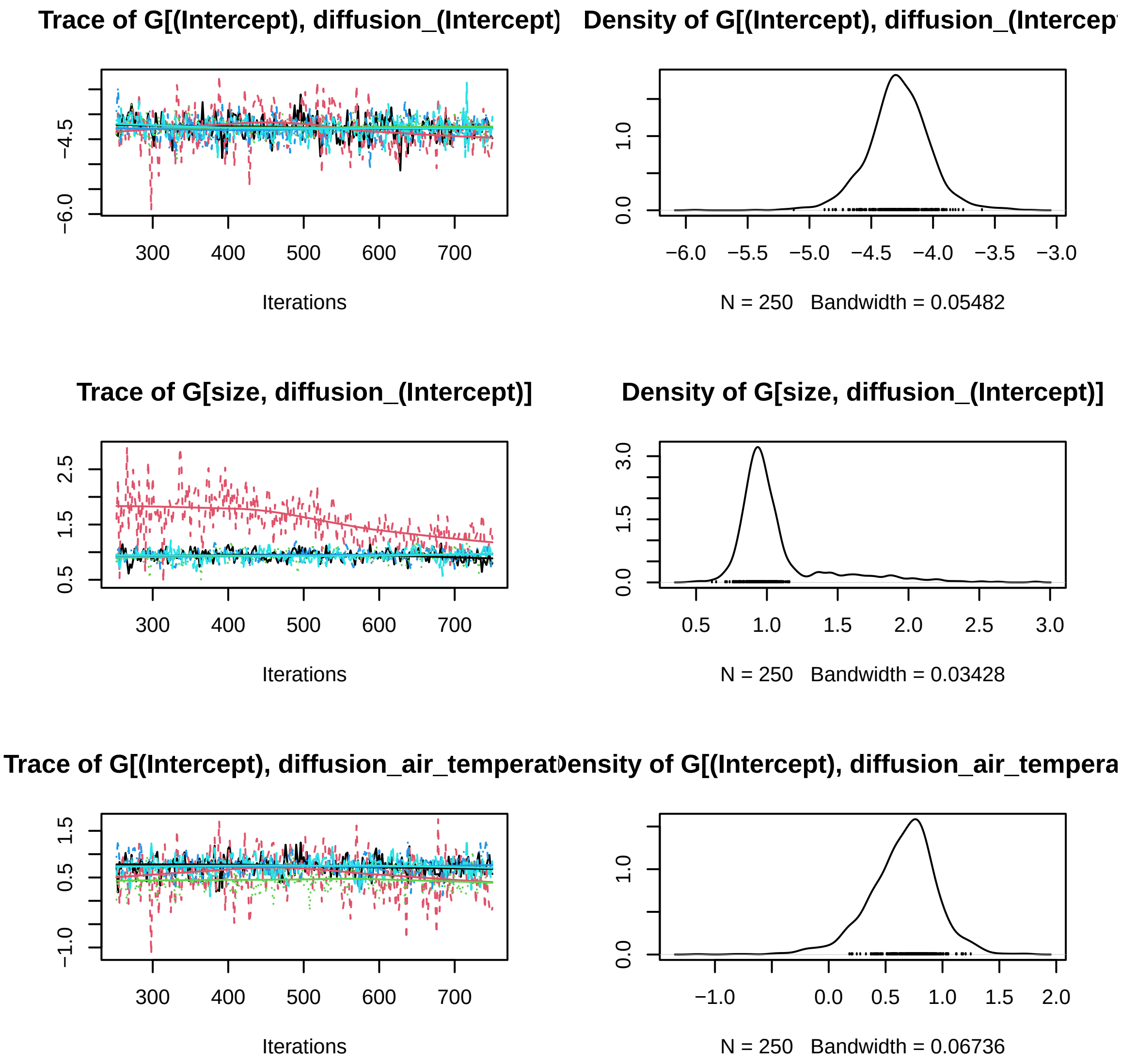

Figure 30: First species trace plots, five MCMC chains Gamma.

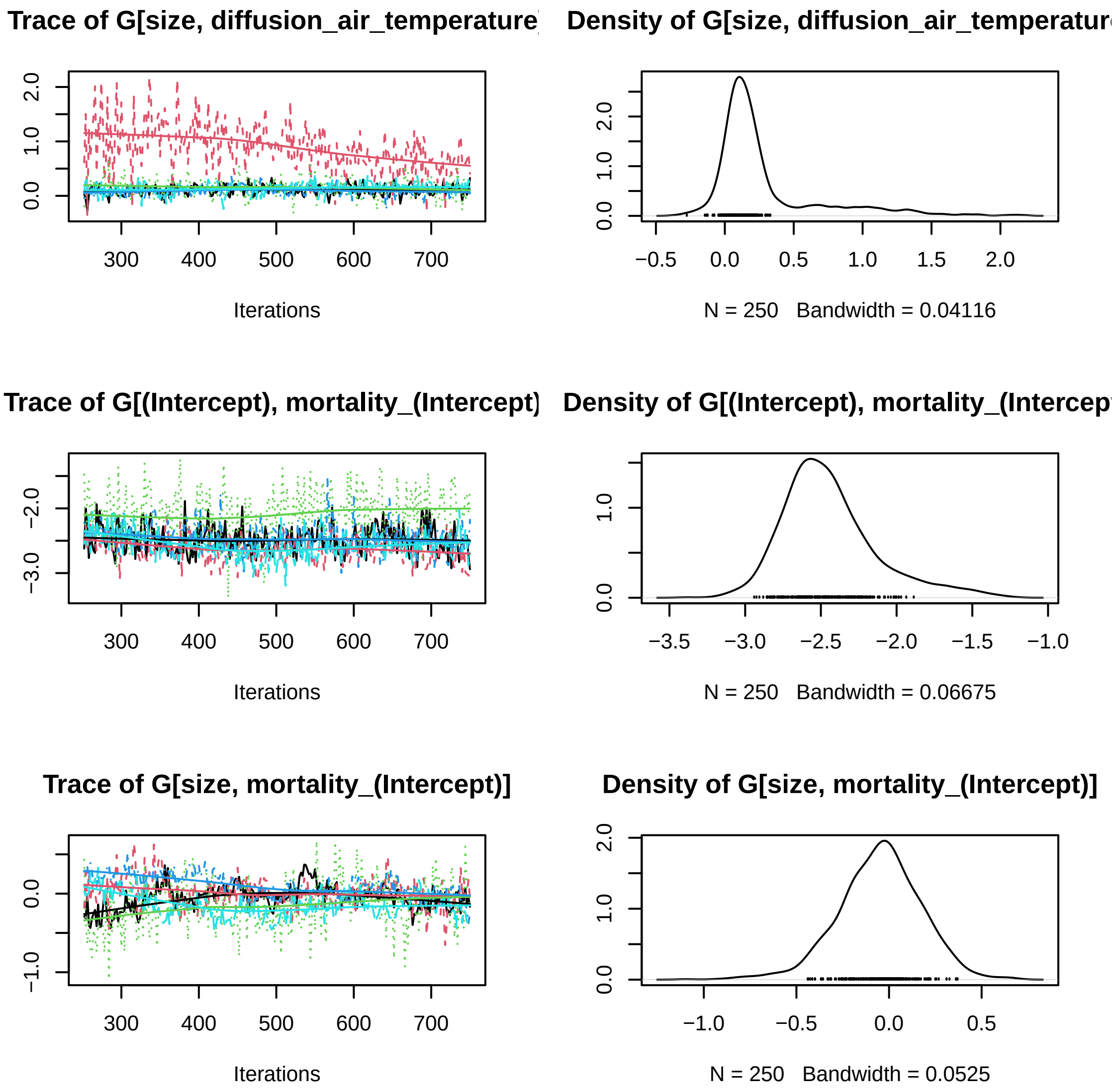

Figure 31: First species trace plots, five MCMC chains Gamma.

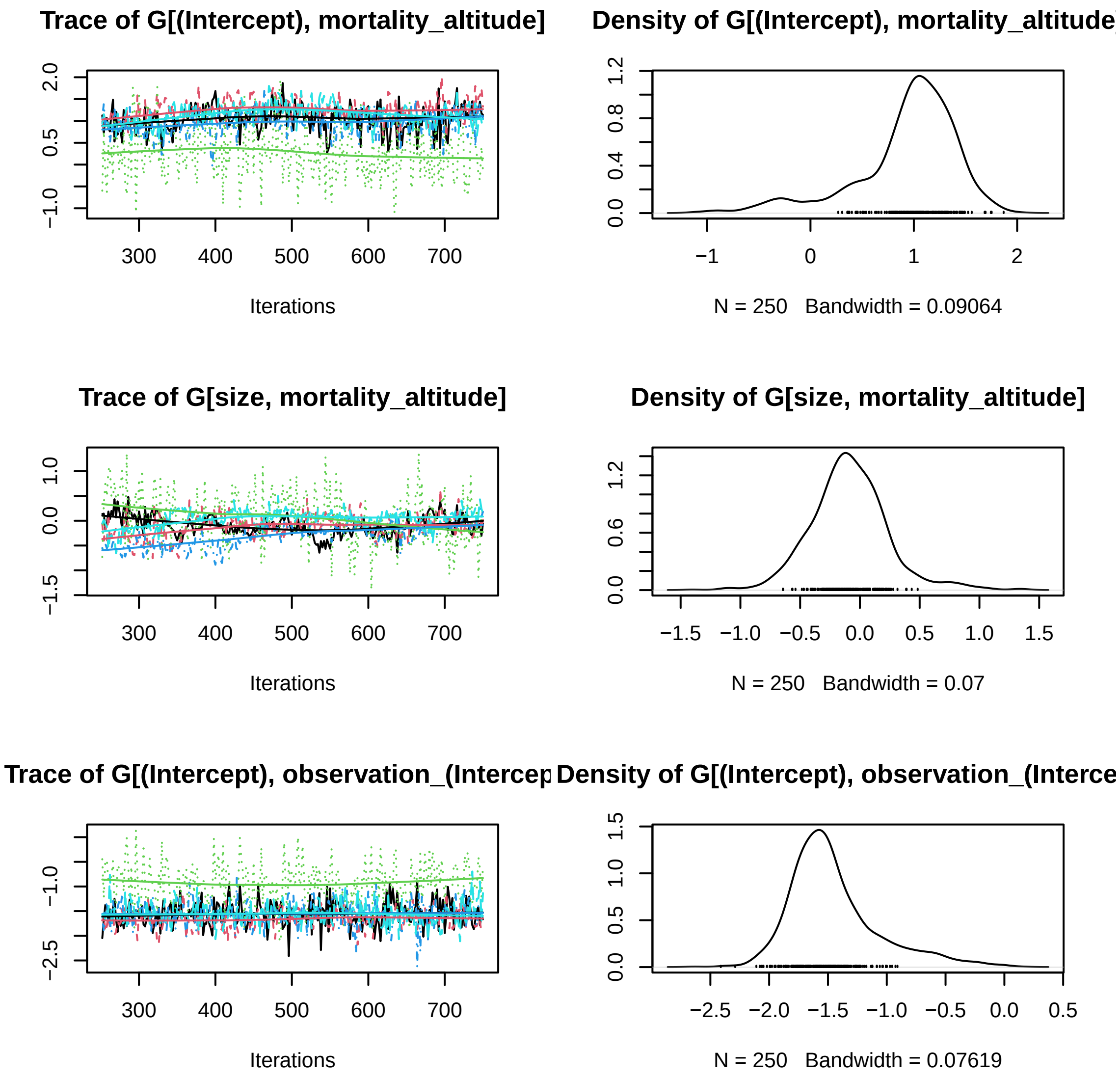

Figure 32: First species trace plots, five MCMC chains Gamma.

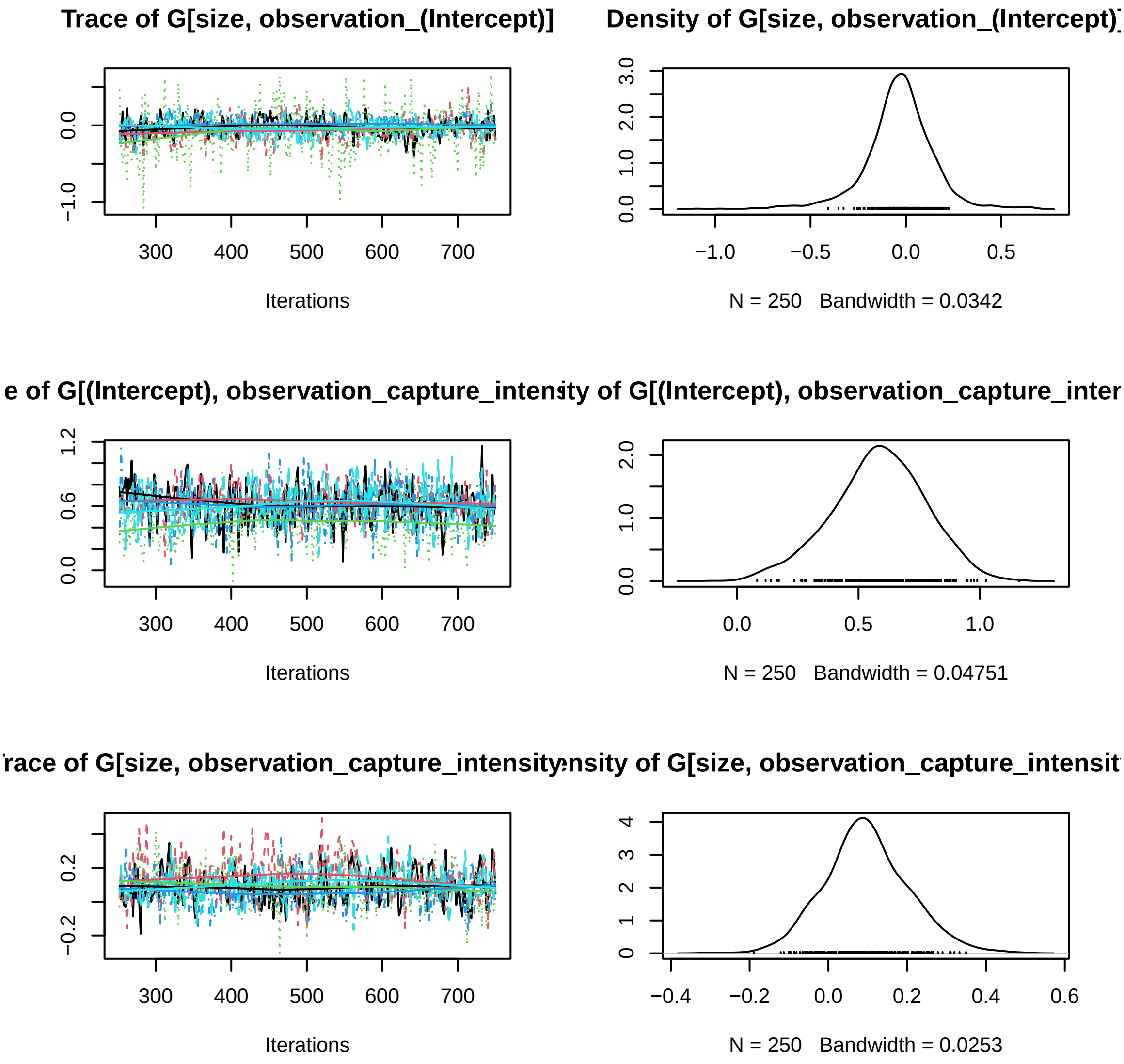

Figure 33: First species trace plots, five MCMC chains Gamma.

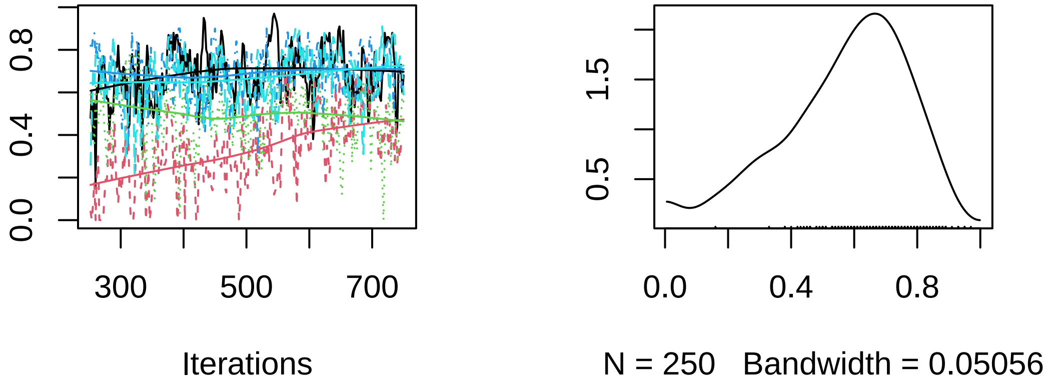

Figure 34: First species trace plots, five MCMC chains Rho.

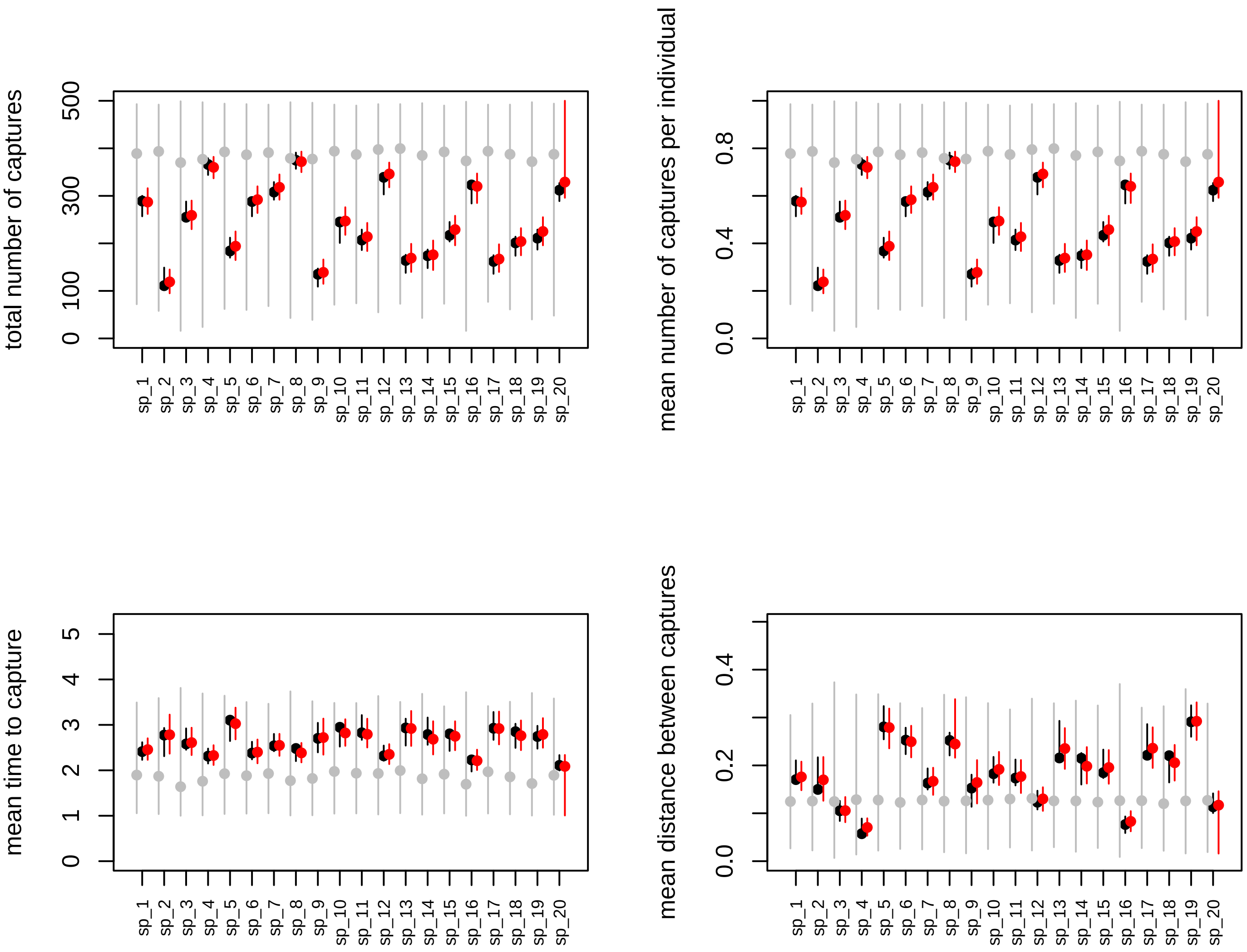

Figure 35: Posterior predictive data generated with 1000 datasets. Black, red, and grey colors represent the datasets generated with true, posterior, and prior parameters values.

```r
nprior = npost
prior  = list()

for(repl in 1:nprior){
  prior[[repl]] = Jsmm::simulate_from_prior(m)
}

load("parameters/true_parameters_CCP.RData")

par(mfrow=c(3,2))

cc = 0
for(va in vars){
  nv = m[["nc"]][[va]]
  for(i in 1:nv){
    cc = cc + 1

    re.pri = matrix(ncol = ns, nrow = nprior)
    re.pri.G = matrix(ncol = nt, nrow = nprior)

    for(repl in 1:nprior){
      re.pri[repl, ]   = prior[[repl]][["Beta"]][[va]][, i]
      re.pri.G[repl, ] = prior[[repl]][["Gamma"]][, cc]
    }

    re.post   = matrix(ncol = ns, nrow = npost)
    re.post.G = matrix(ncol = nt, nrow = nprior)

    for(repl in 1:npost){
      re.post[repl, ]   = post[[repl]][["Beta"]][[va]][, i]
      re.post.G[repl, ] = post[[repl]][["Gamma"]][, cc]
    }

    ymin = min(quantile(re.pri, probs = 0.05), min(re.post))
    ymax = max(quantile(re.pri, probs = 0.95), max(re.post))

    if(showTrueValues){
      ymin = min(ymin, pars$Beta[[va]][, i])
      ymax = max(ymax, pars$Beta[[va]][, i])
    }

    colnames(re.pri) = m$sp_names
    re.pri  = re.pri[, sel]
    re.post = re.post[, sel]

    boxplot(re.pri, outline = FALSE, at = 3*(1:ns)-0.75, ylim = c(ymin, ymax),
            main = m[["parNames"]][cc], las = 2, cex.axis = 0.7,
            xlim = c(2, 3*ns + 2),
            ylab = "parameter value", col = "grey", xaxt = "n")

    axis(1, at = 3*(1:m$ns), label = colnames(re.pri), las = 2, cex.axis = 0.7)

    boxplot(re.post, outline = FALSE, at = 3*(1:ns) + 0.75, col = "pink", add = TRUE,
            xaxt = "n", yaxt = "n")
```

```
    abline(h = 0)

    if(showTrueValues){
      for(j in 1:m[["ns"]]){
        lines(x = c(3*j, 3*j + 1.5), y = c(pars$Beta[[va]][j, i], pars$Beta[[va]][j, i]),
              lwd = 2, col = "red")
      }
    }

    ymin = min(quantile(re.pri.G, probs = 0.05), min(re.post.G))
    ymax = max(quantile(re.pri.G, probs = 0.95), max(re.post.G))

    if(showTrueValues){
      ymin = min(ymin, pars$Gamma[, cc])
      ymax = max(ymax, pars$Gamma[, cc])
    }

    colnames(re.pri.G) = colnames(m$X[["trait"]])

    boxplot(re.pri.G, outline = FALSE, at = 3*(1:nt), ylim = c(ymin, ymax),
            main = m[["parNames"]][cc], las = 2, cex.axis = 0.7,
            xlim = c(2, 3*nt + 2),
            ylab = "parameter value", col = "grey")

    boxplot(re.post.G, outline = FALSE, at = 3*(1:nt) + 1.5, col = "pink", add = TRUE,
            xaxt = "n", yaxt = "n")

    abline(h = 0)

    if(showTrueValues){
      for(j in 1:m[["nt"]]){
        lines(x = c(3*j, 3*j + 1.5), y = c(pars$Gamma[j, cc], pars$Gamma[j, cc]),
              lwd = 2, col = "red")
      }
    }
  }
}
```

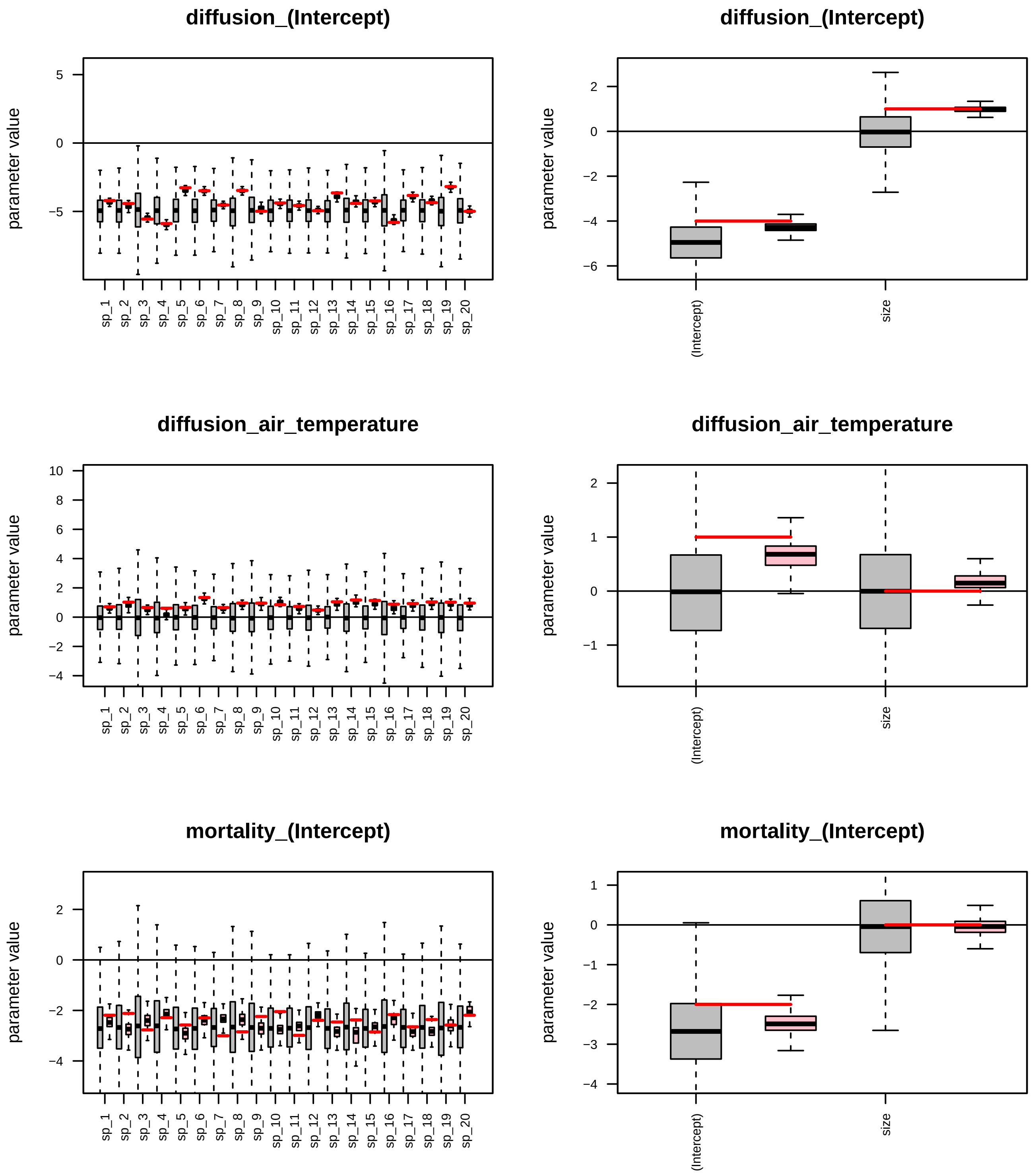

Figure 36: Posterior credible intervals for the model parameters. Left panels show the posteriors for each of the model parameters (pink color), their corresponding priors (Grey), and the parameter true value (Red). Right panels show the corresponding values for the Gamma associated parameter.

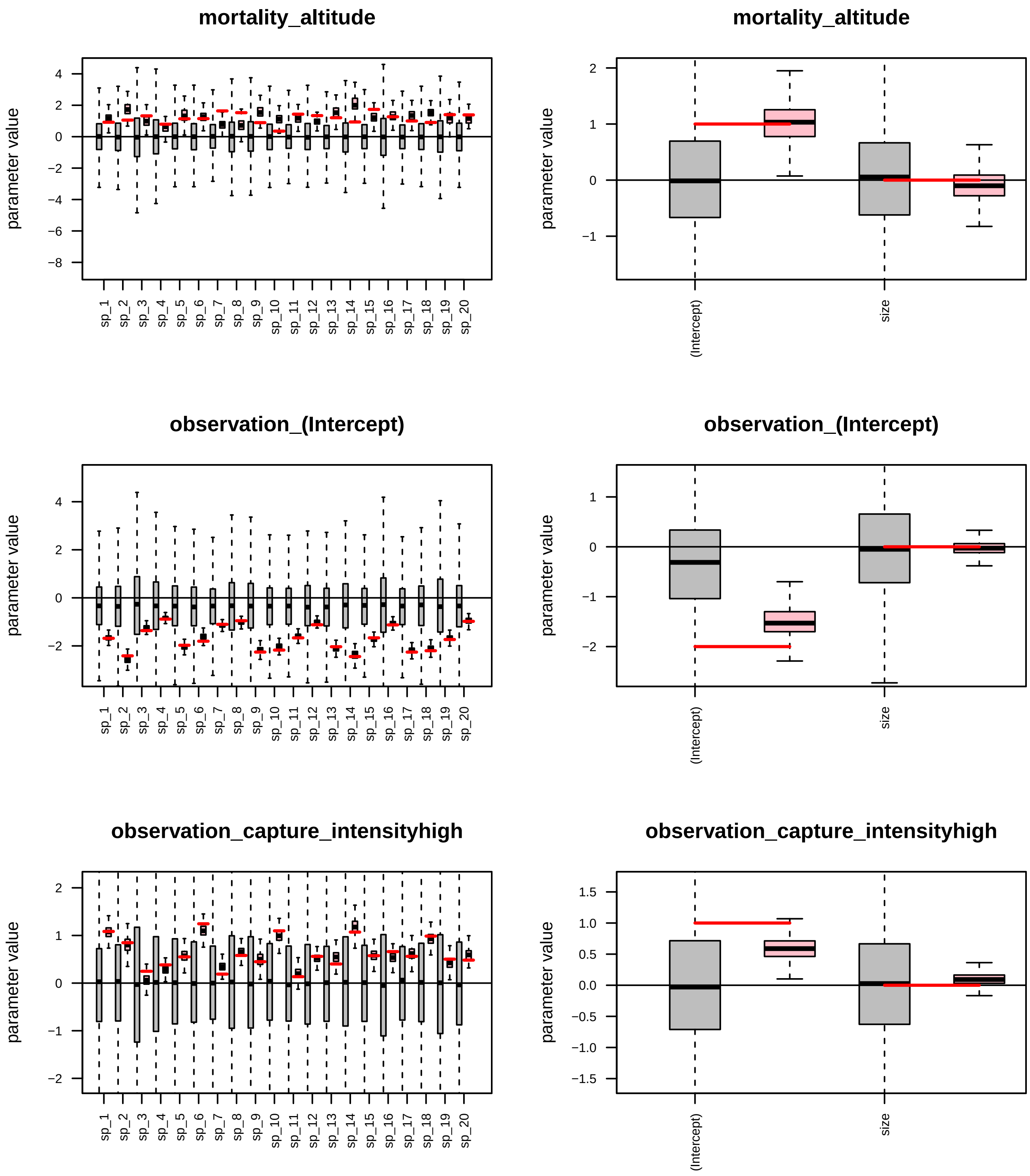

Figure 37: Posterior credible intervals for the model parameters. Left panels show the posteriors for each of the model parameters (pink color), their corresponding priors (Grey), and the parameter true value (Red). Right panels show the corresponding values for the Gamma associated parameter.

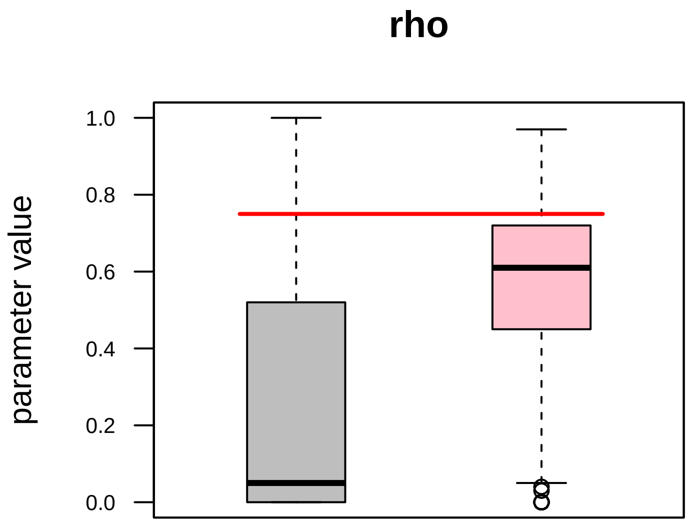

Figure 38: Posterior credible intervals for the associated rho parameter. Posterior, prior, and true value are represented by pink, grey, and red colors, respectively.

### How to set temporally varying capture event-specific covariates for CCP capture process
