## Supplementary figures and images for "R-package Jsmm: Joint species movement modelling of mark-recapture data"

### likelihood_profiles_simulated_ICP.pdf

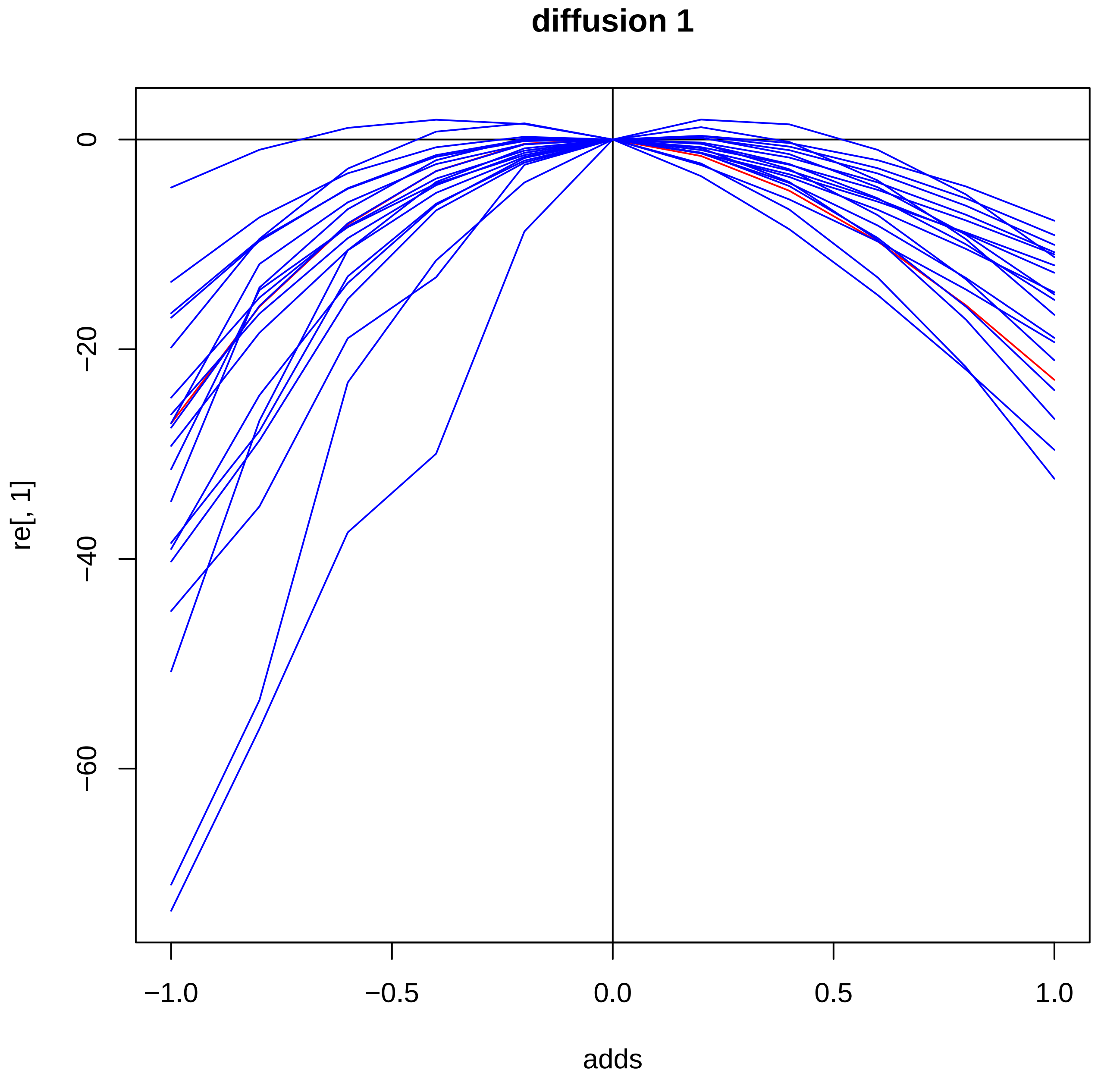

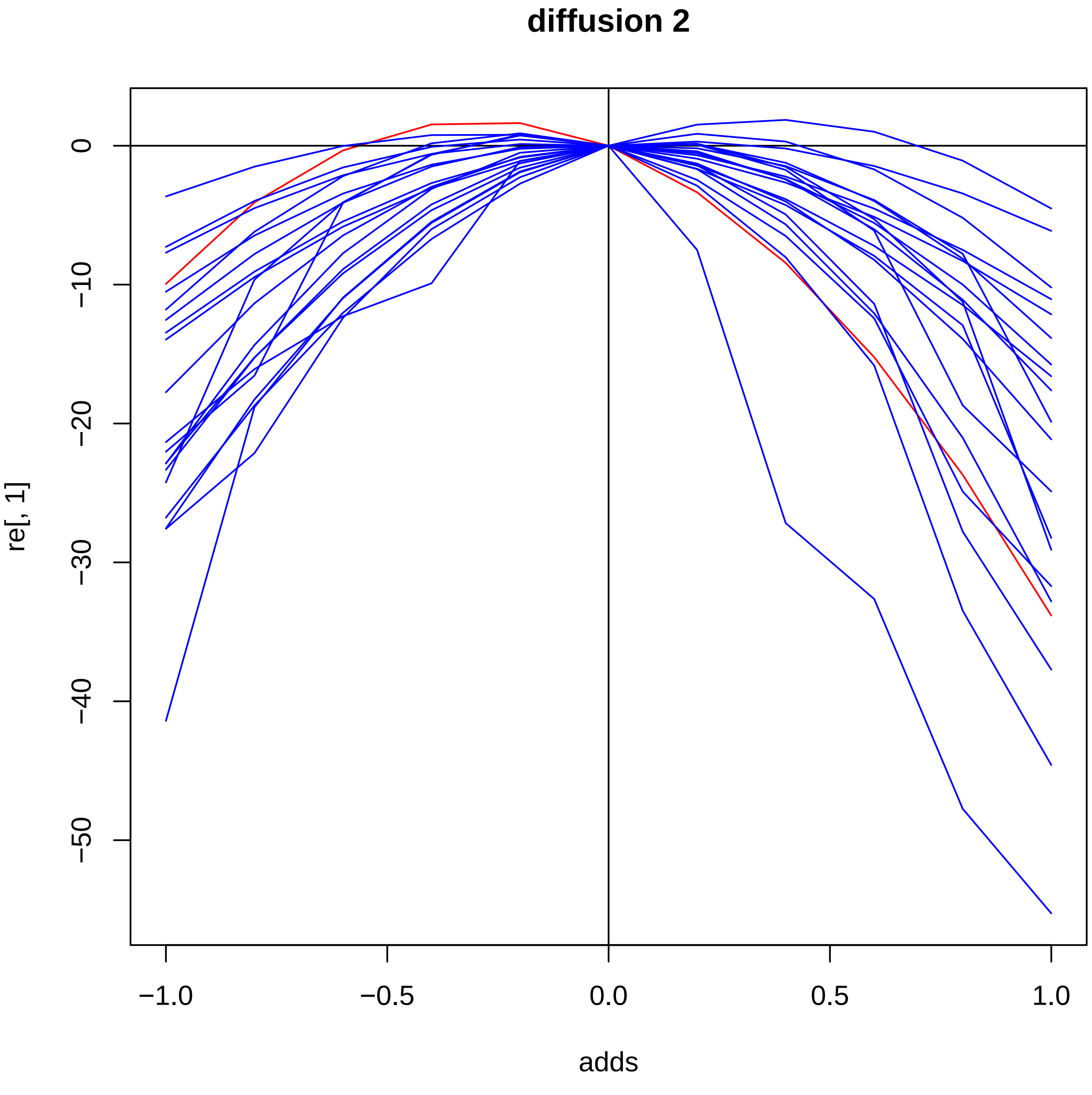

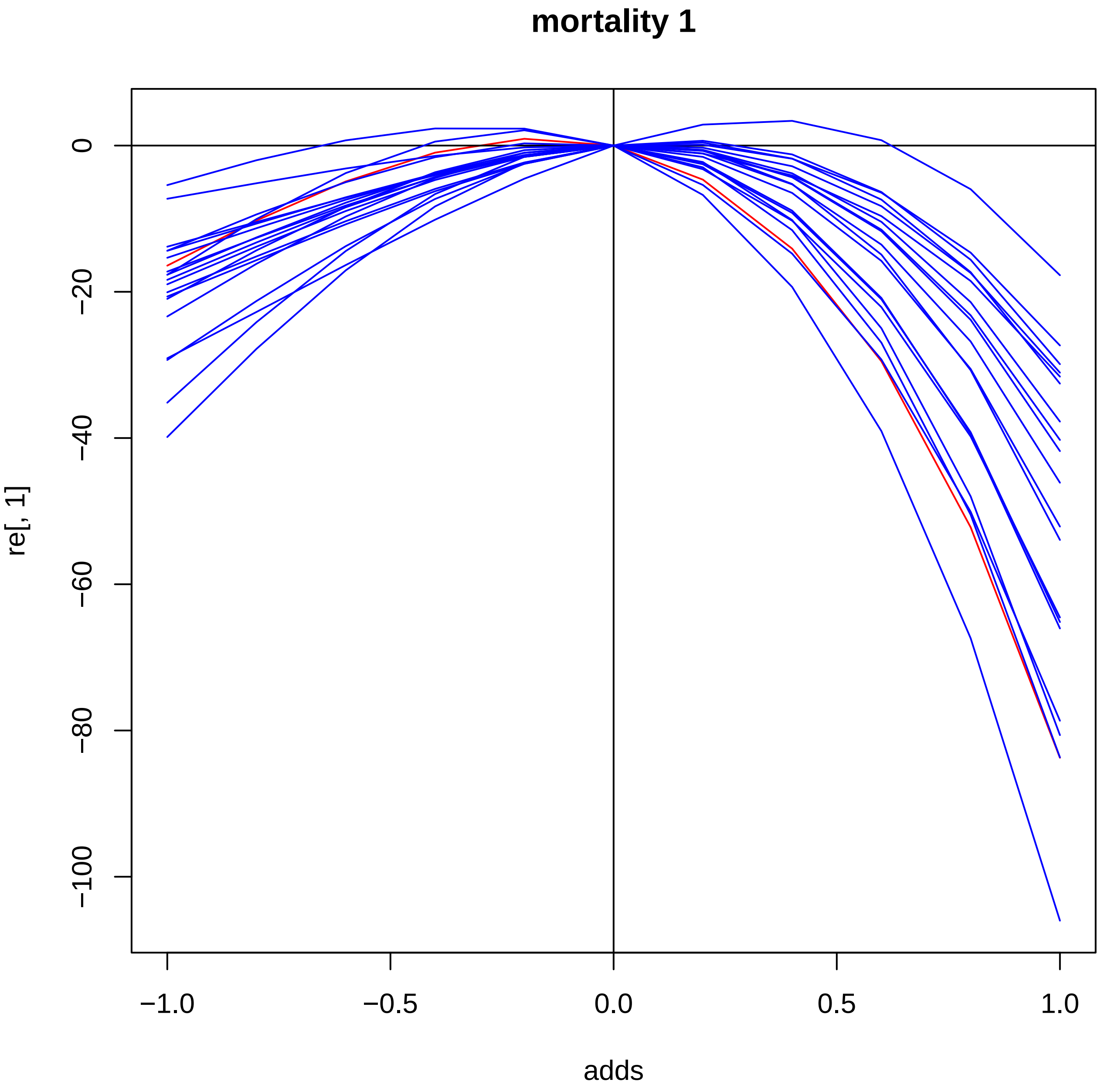

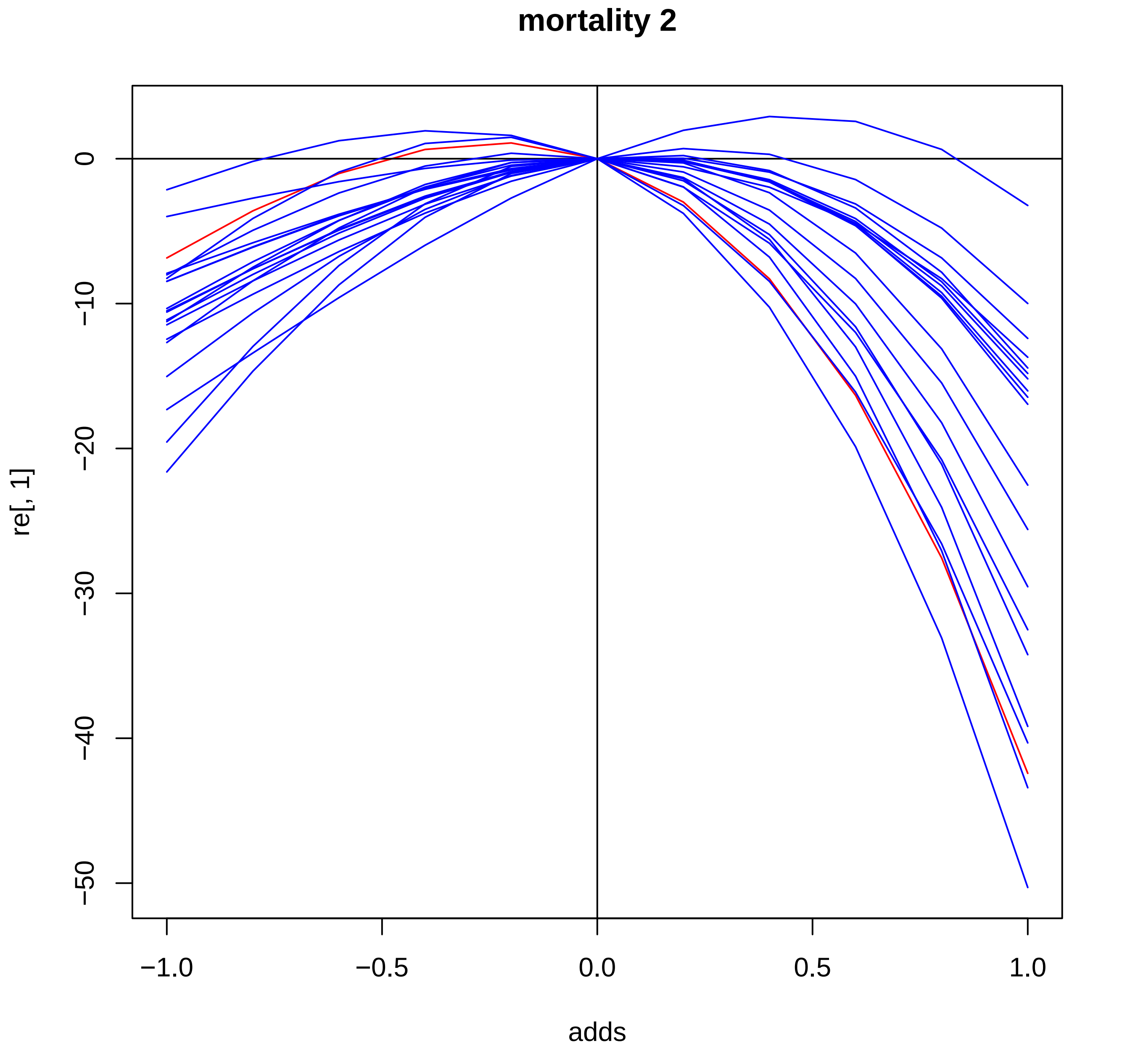

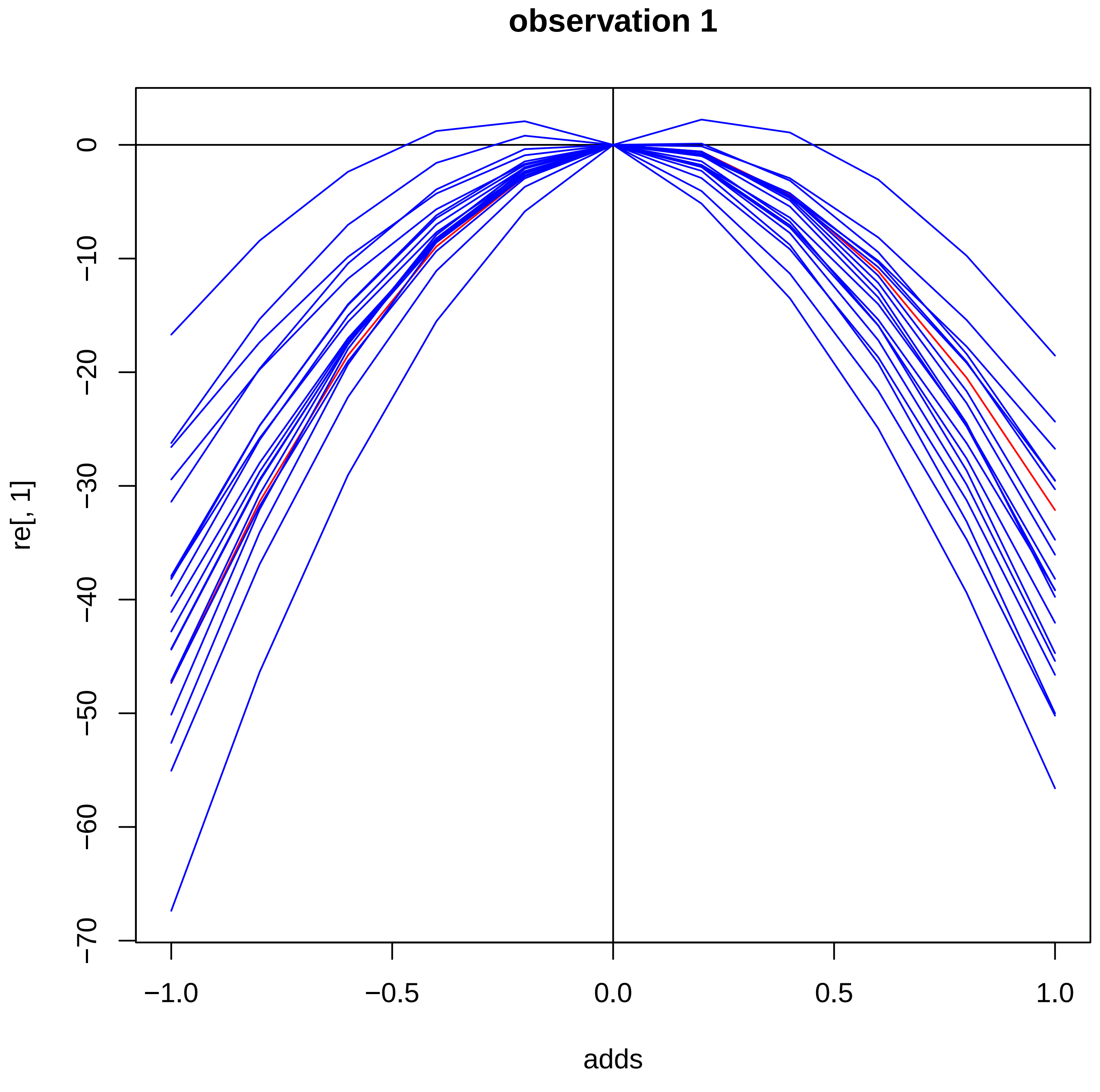

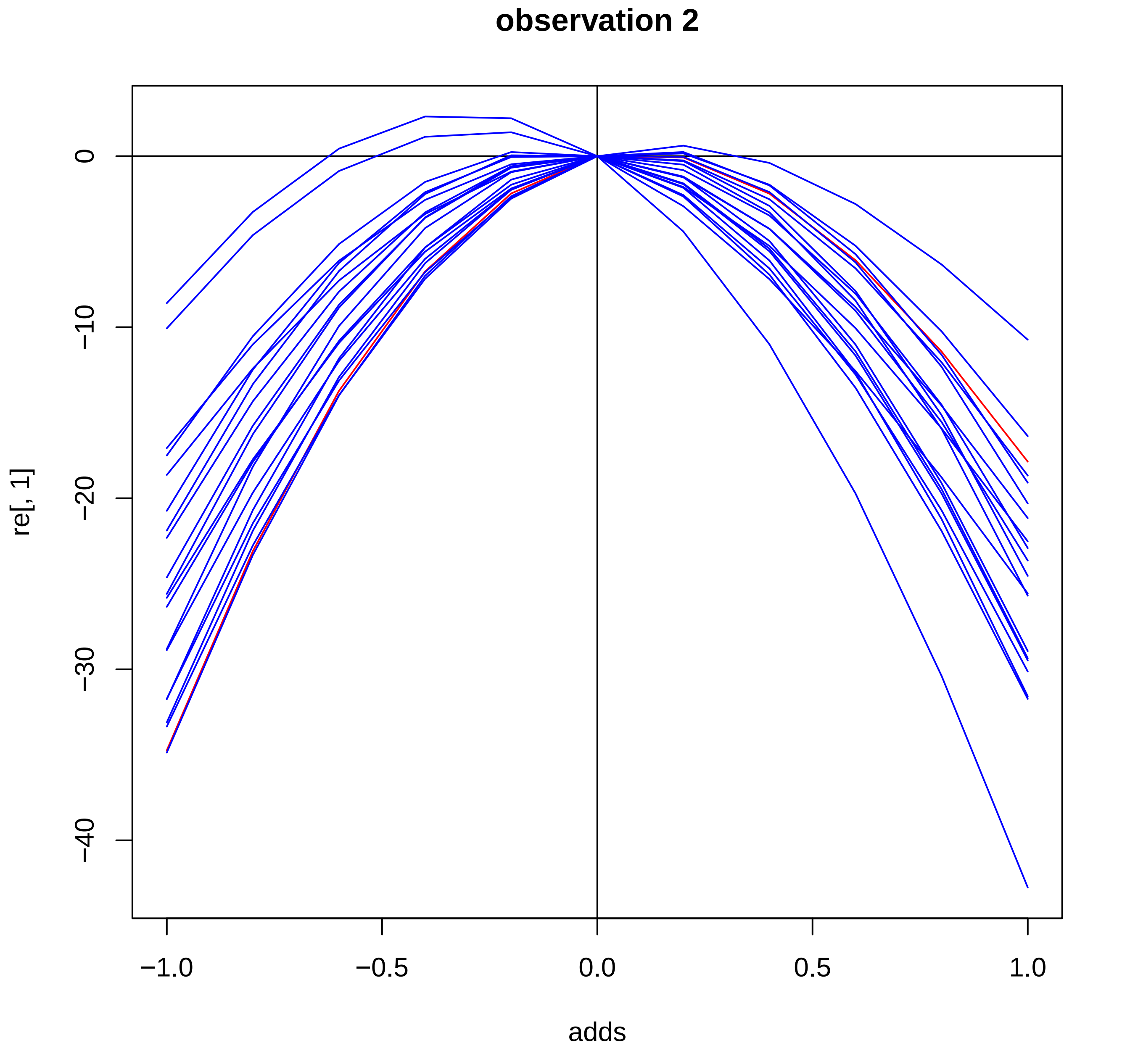

### model_fit_moths.pdf

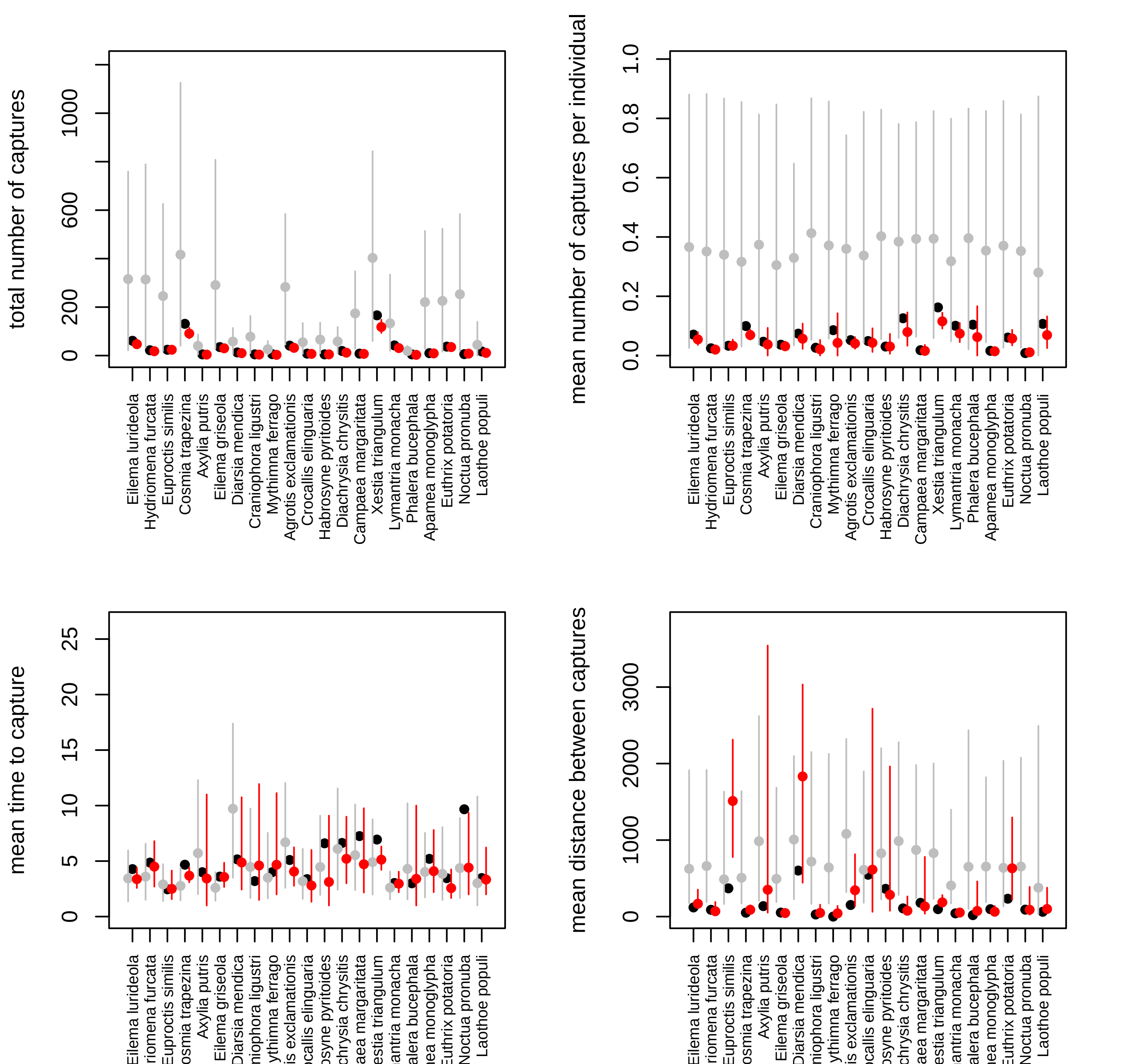

### model_fit_simulated_CCP.pdf

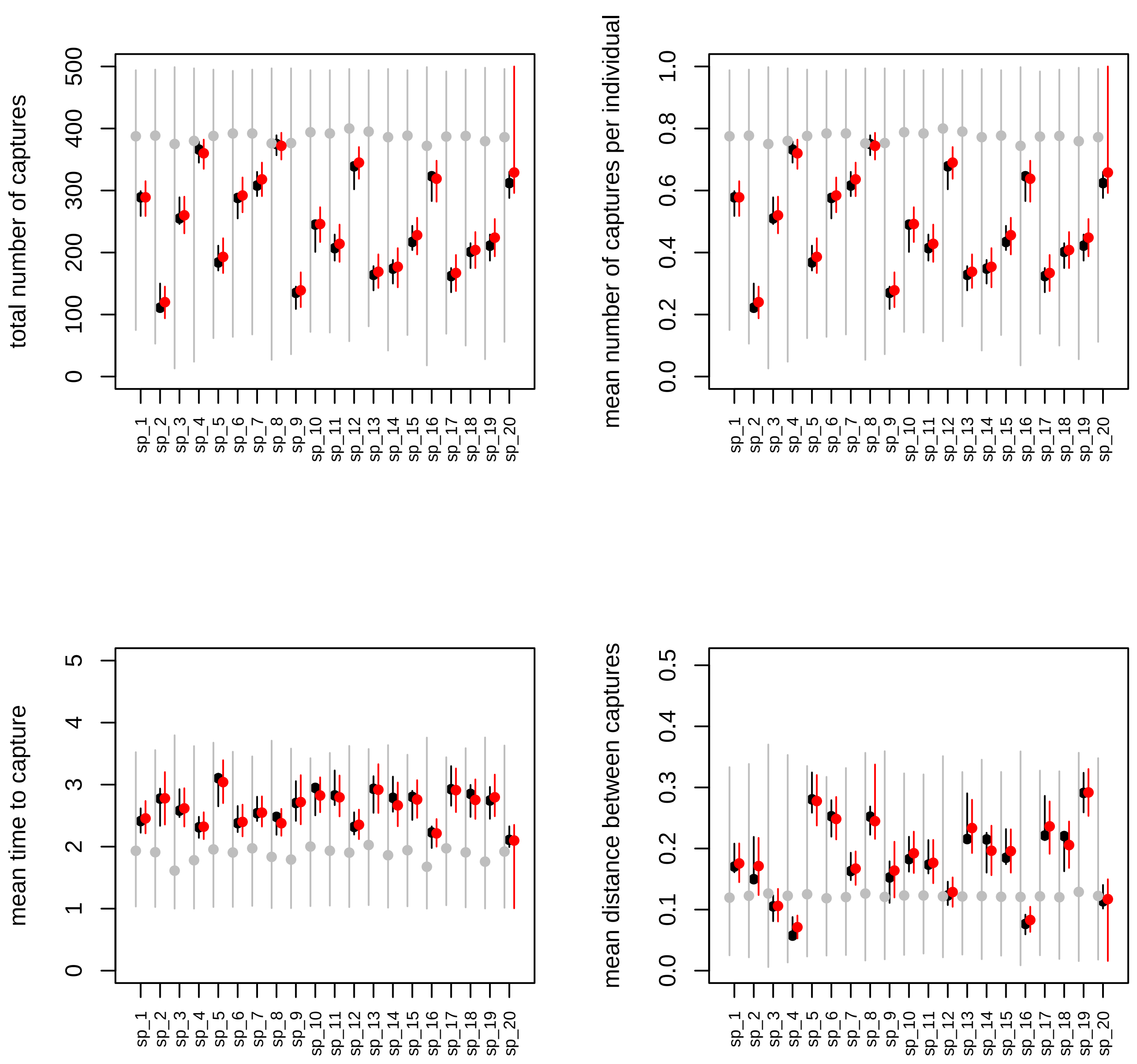

### model_fit_simulated_ICP.pdf

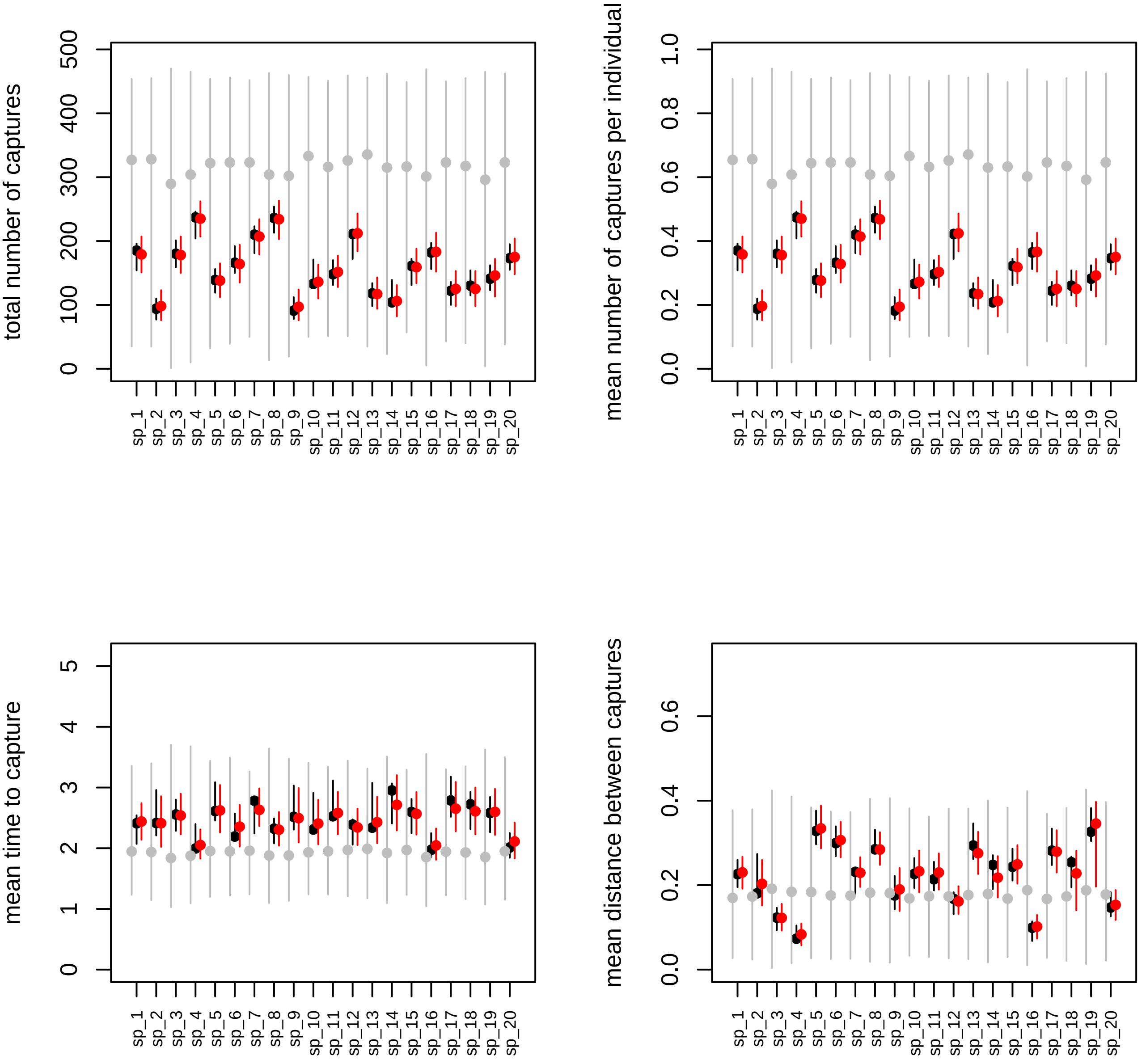

### posteriors_vs_priors_moths.pdf

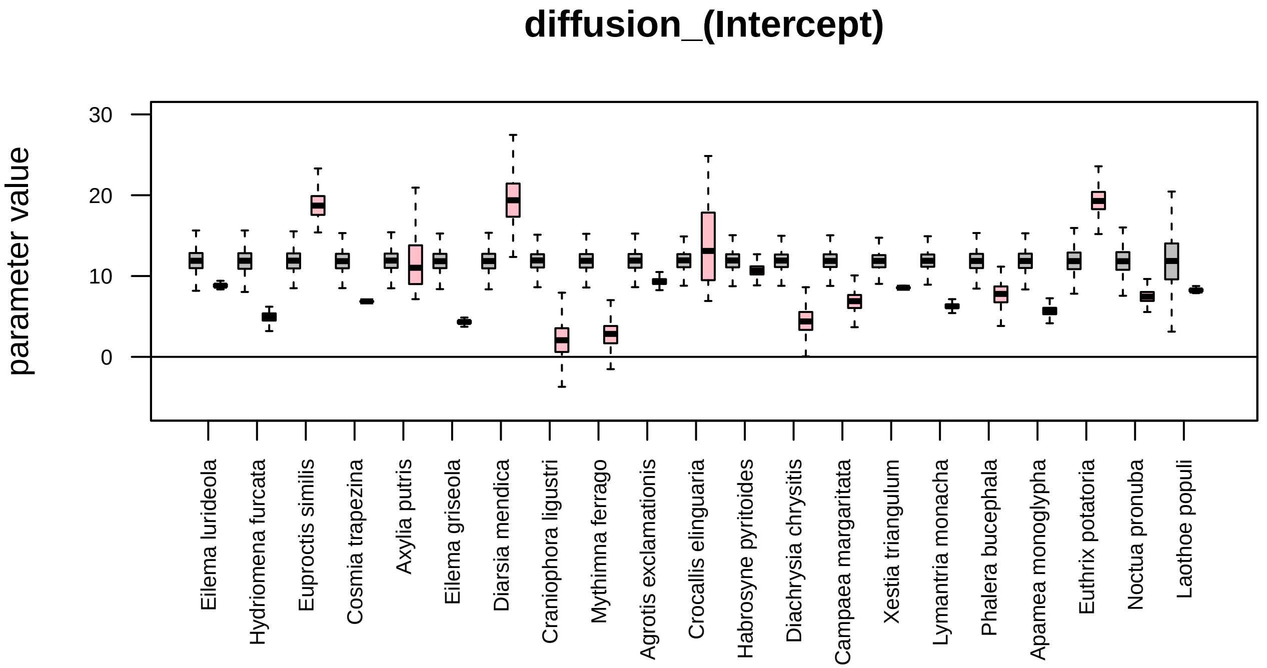

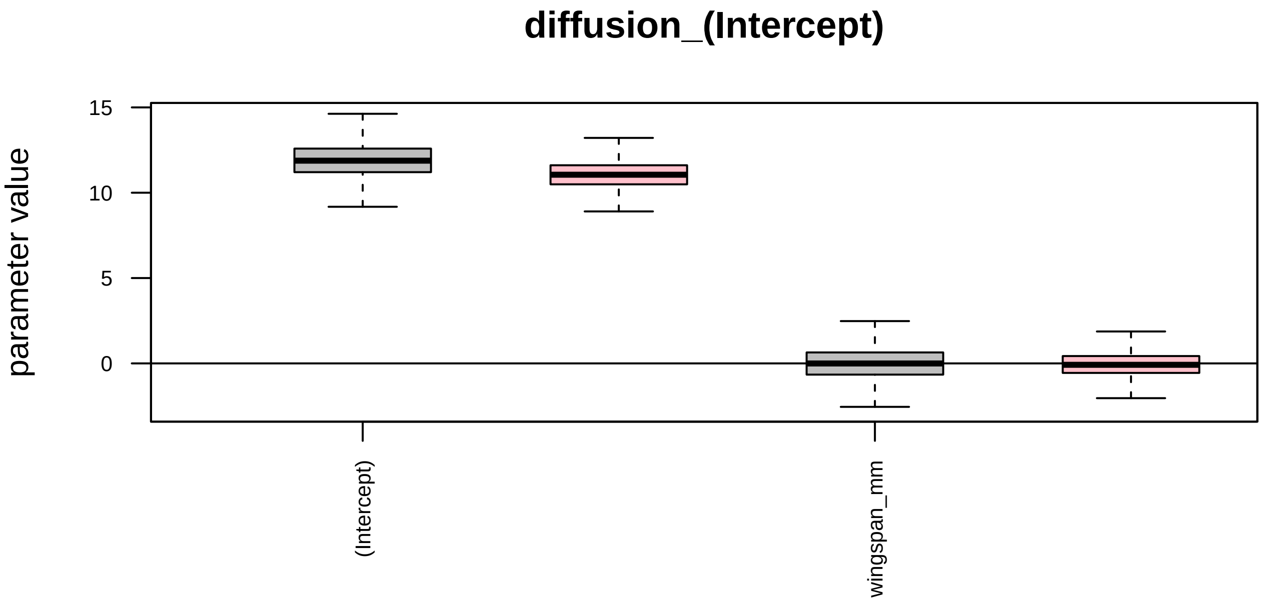

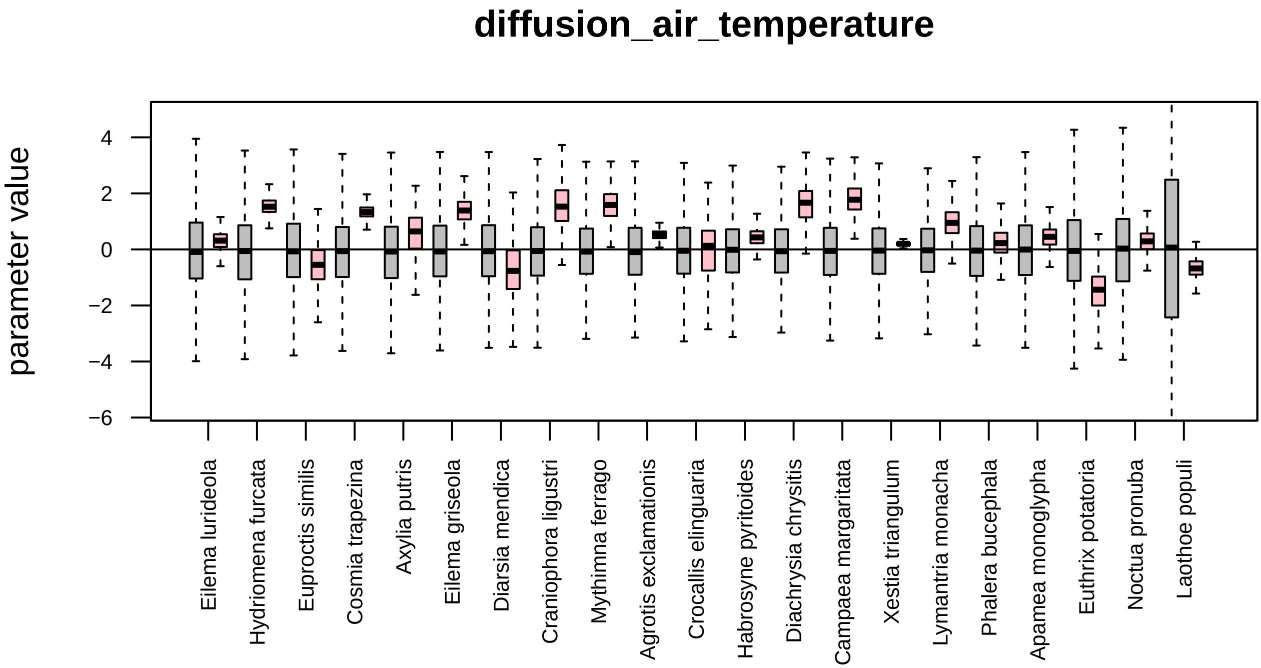

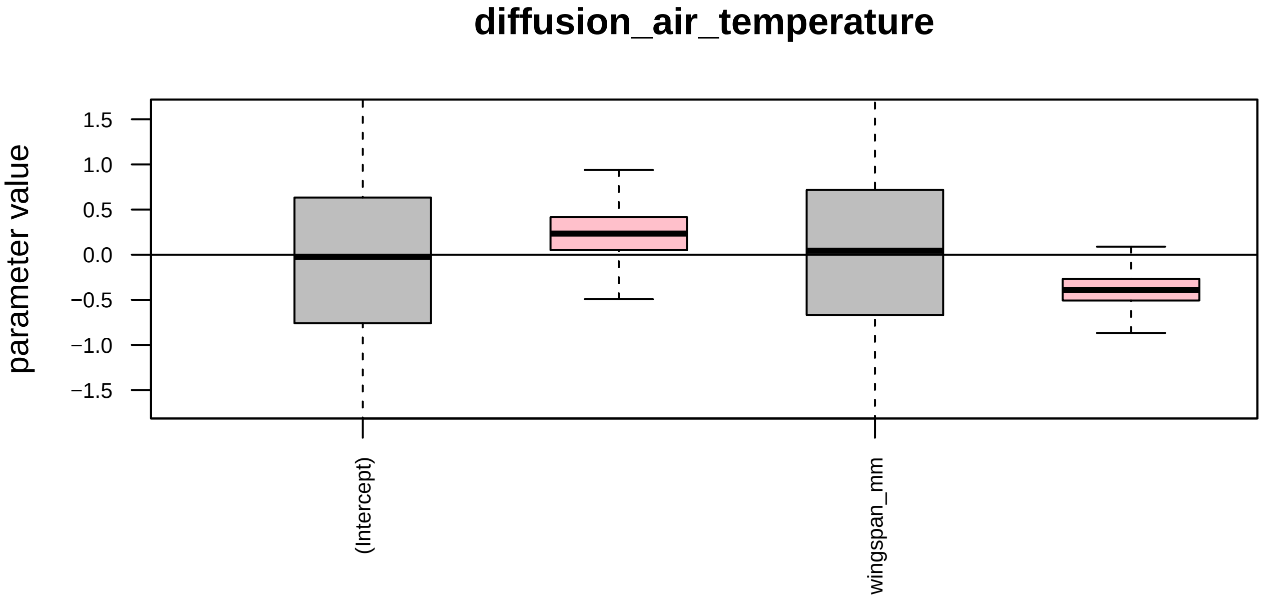

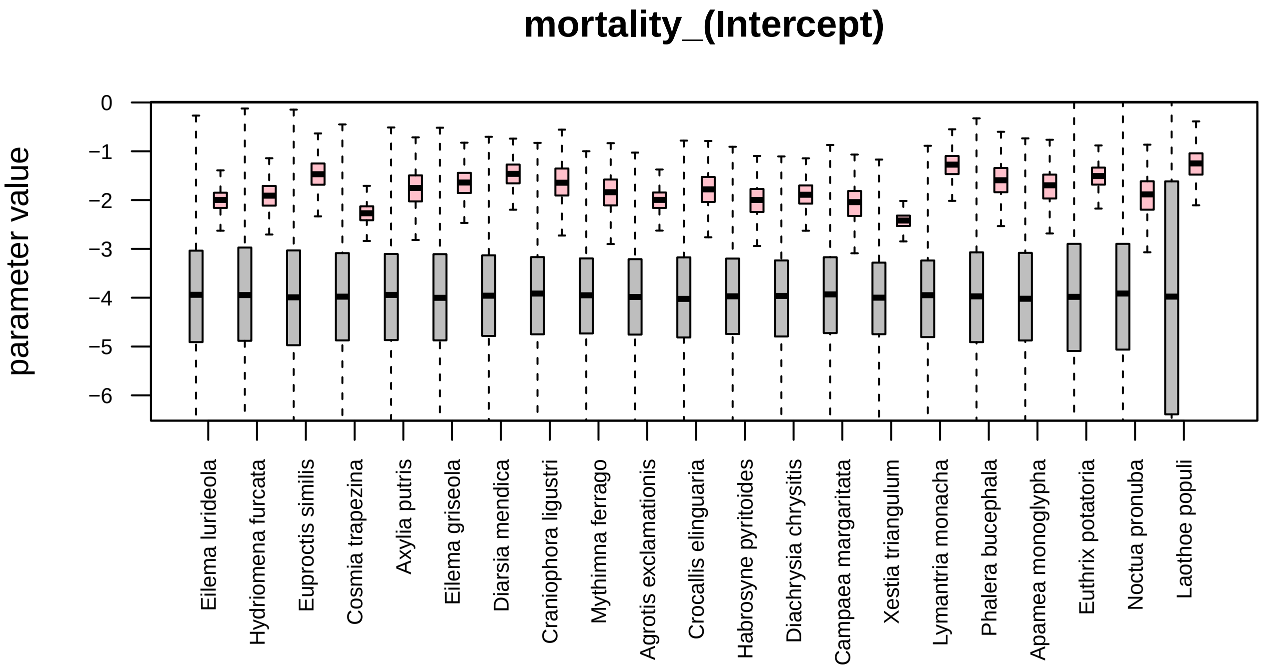

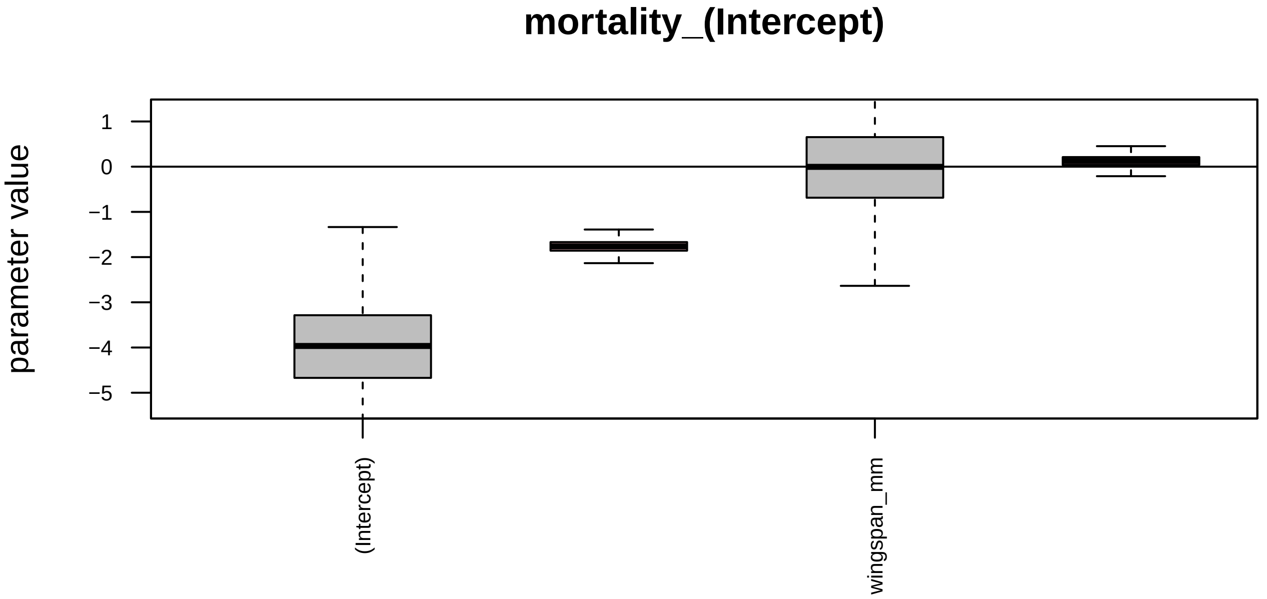

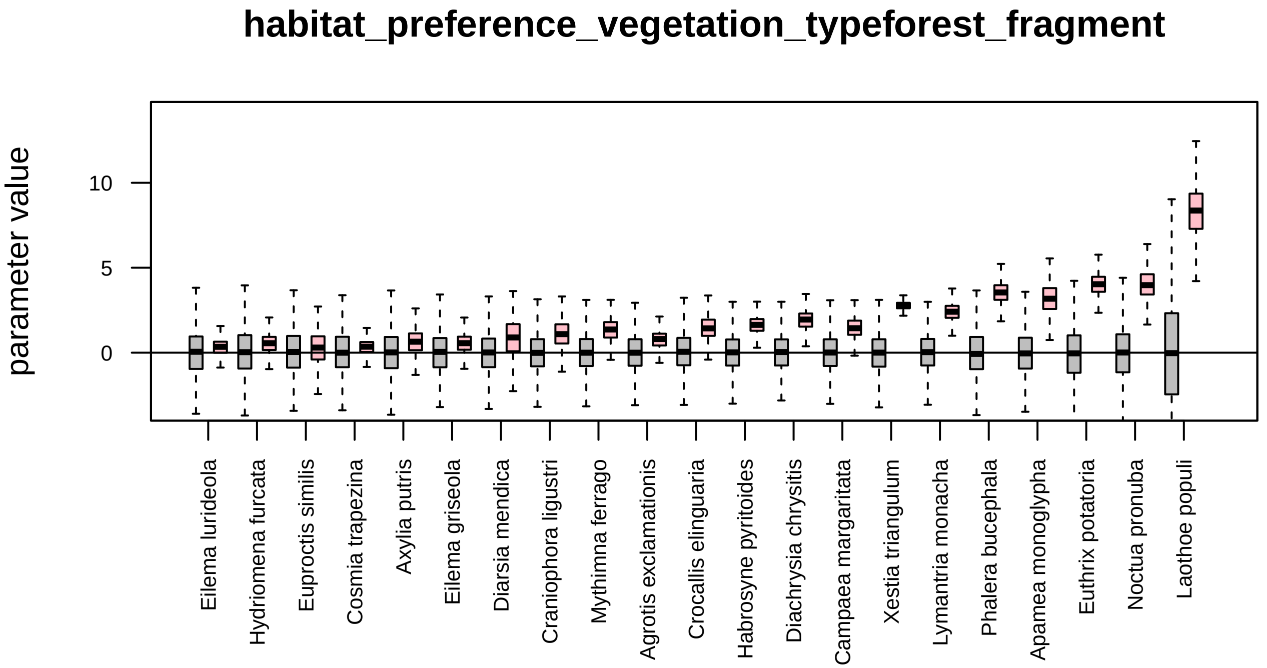

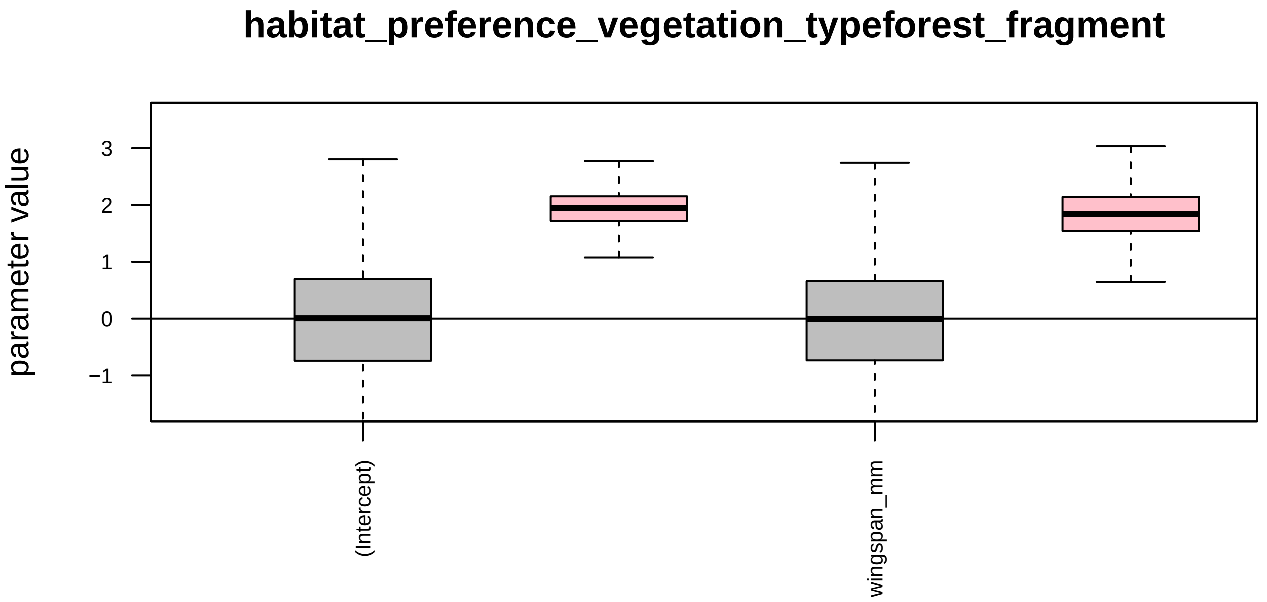

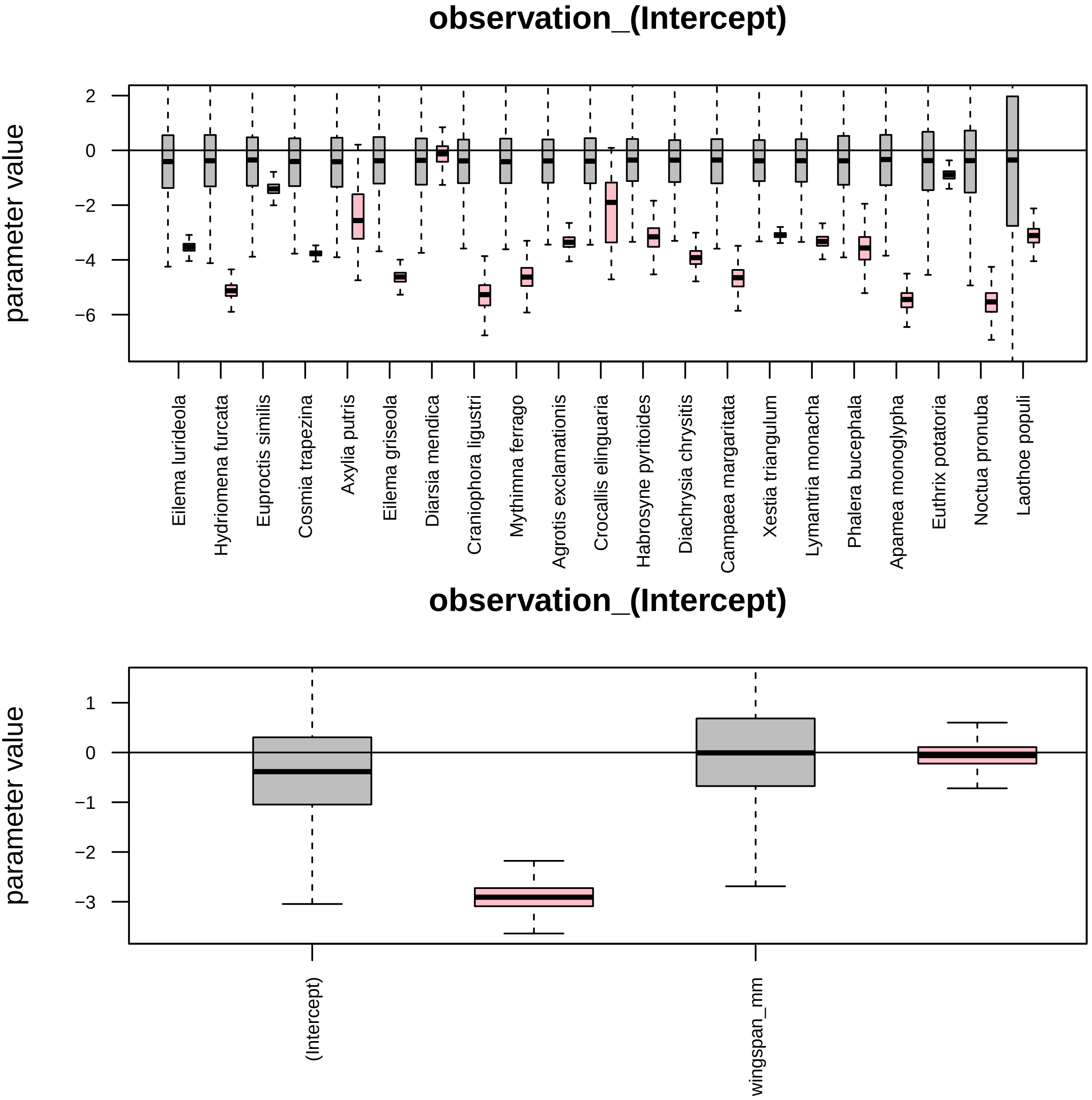

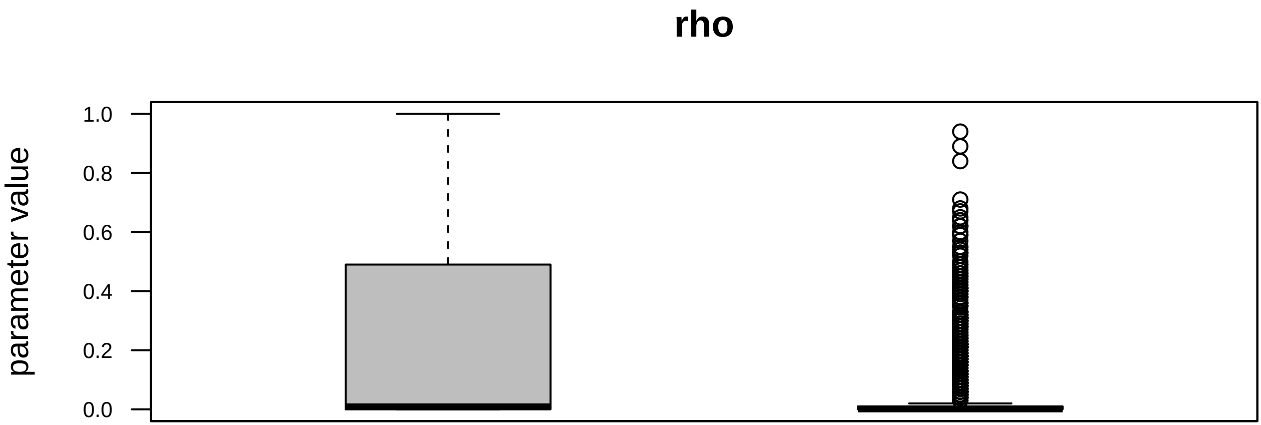

### posteriors_vs_priors_simulated_CCP.pdf

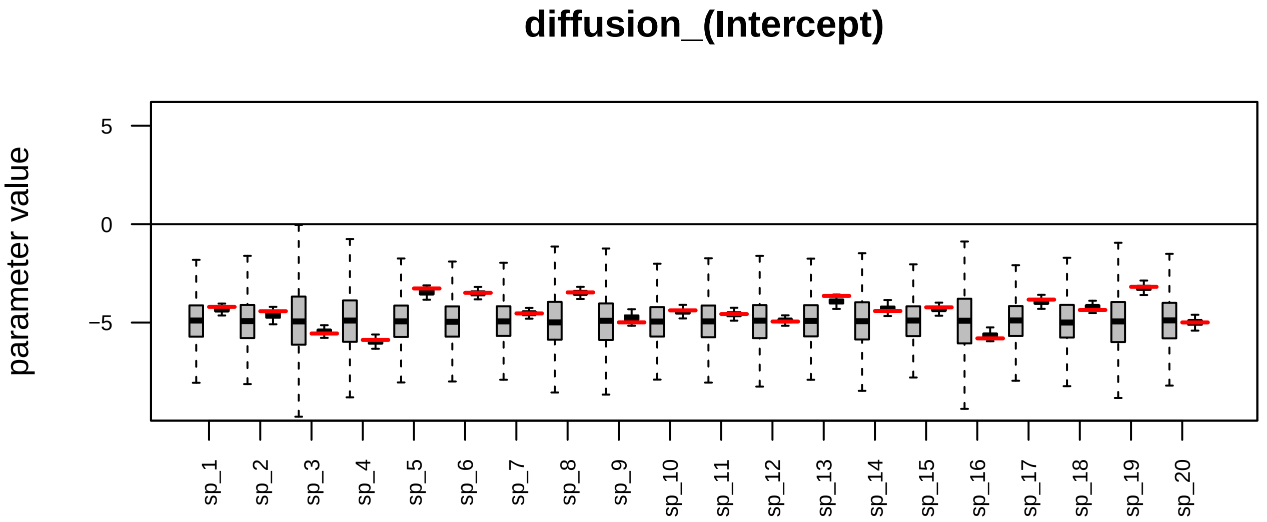

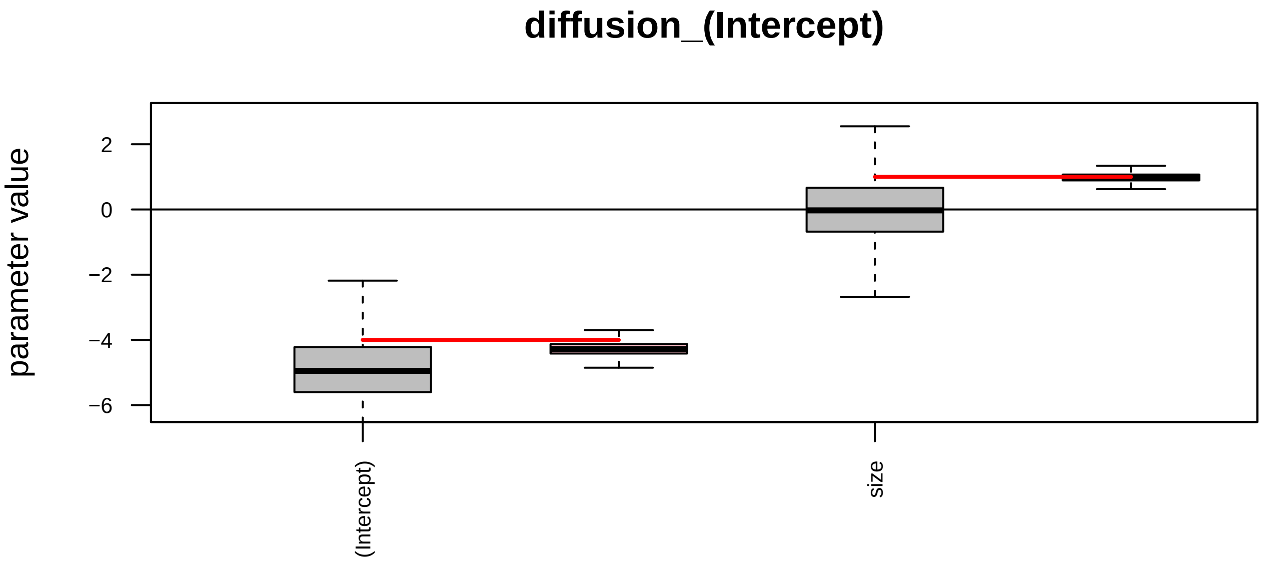

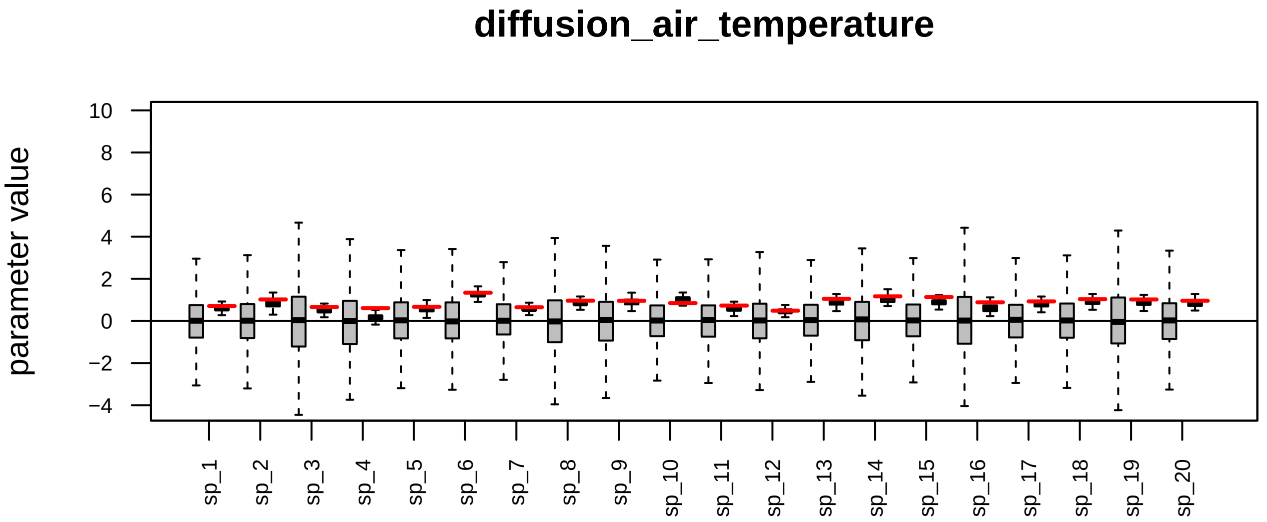

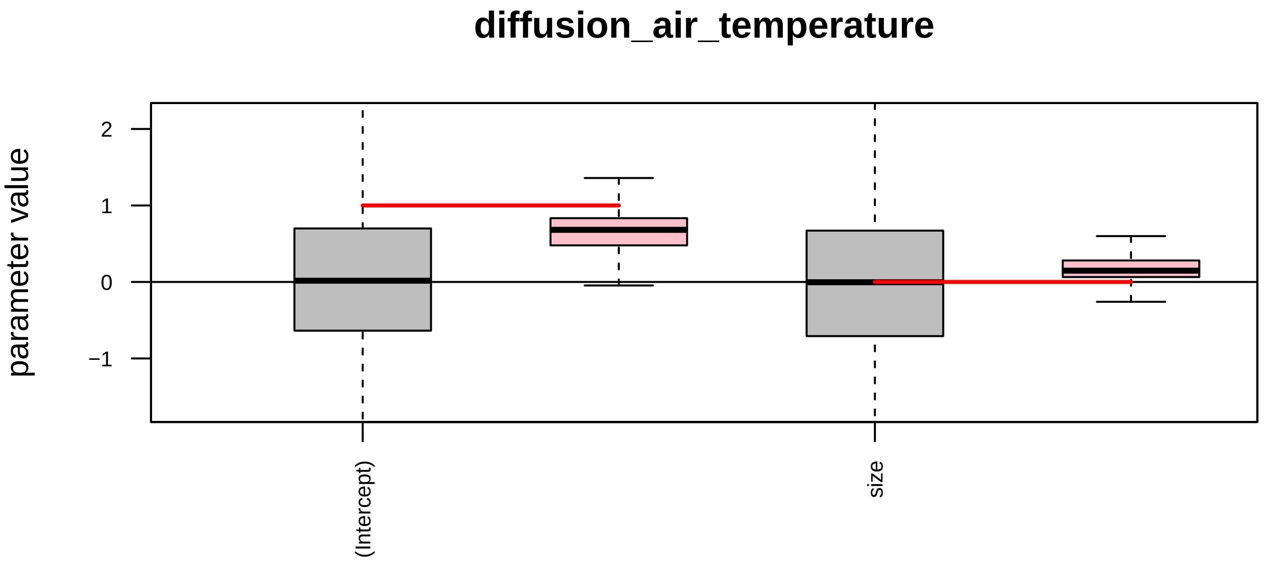

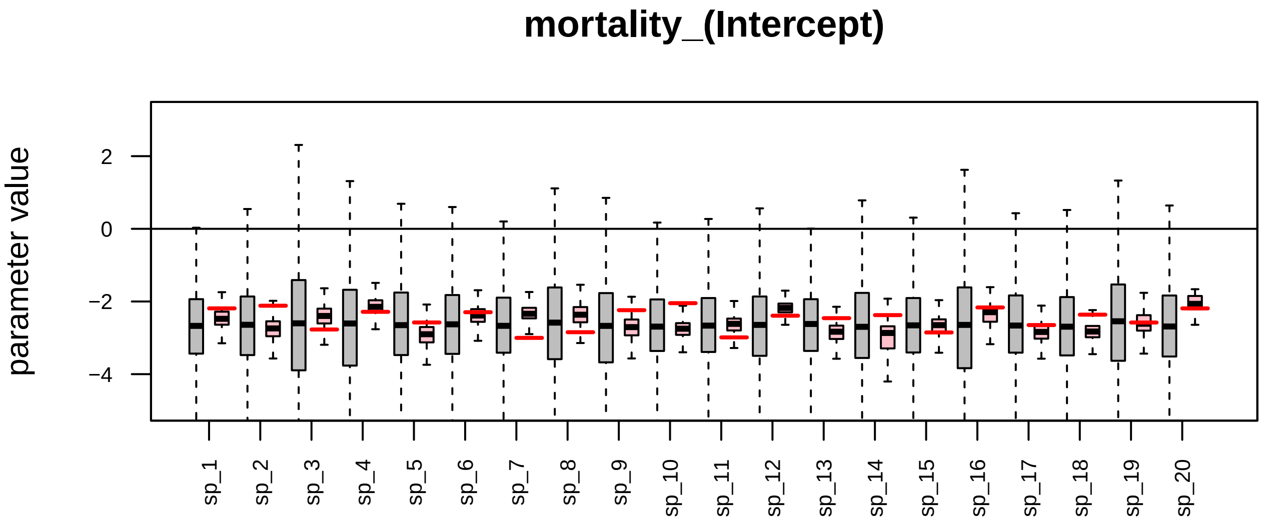

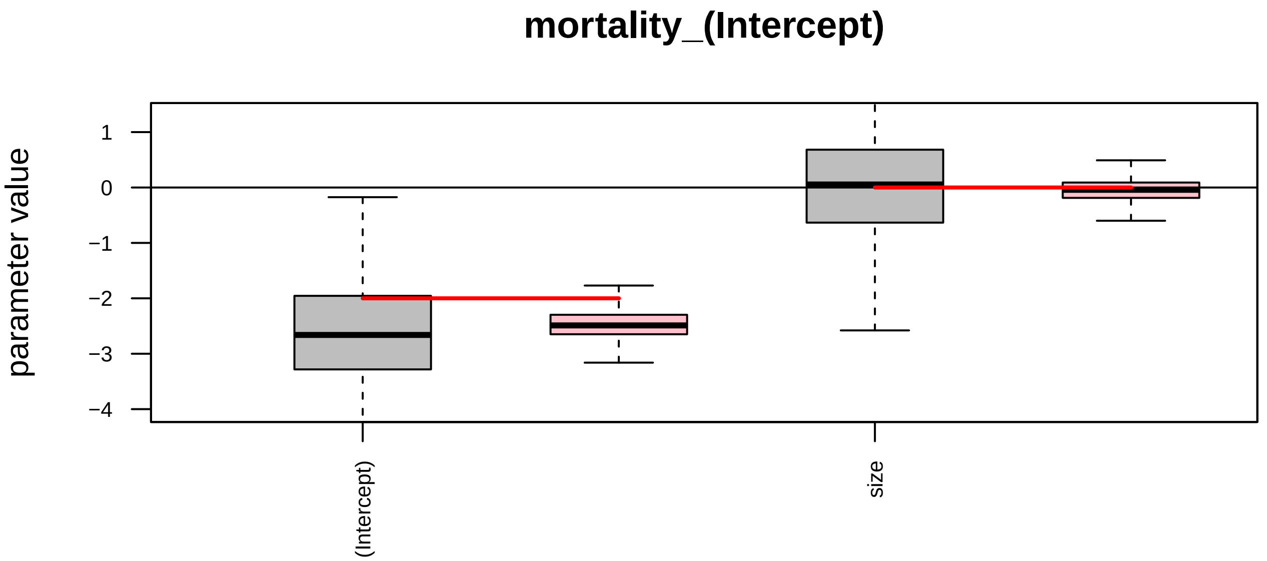

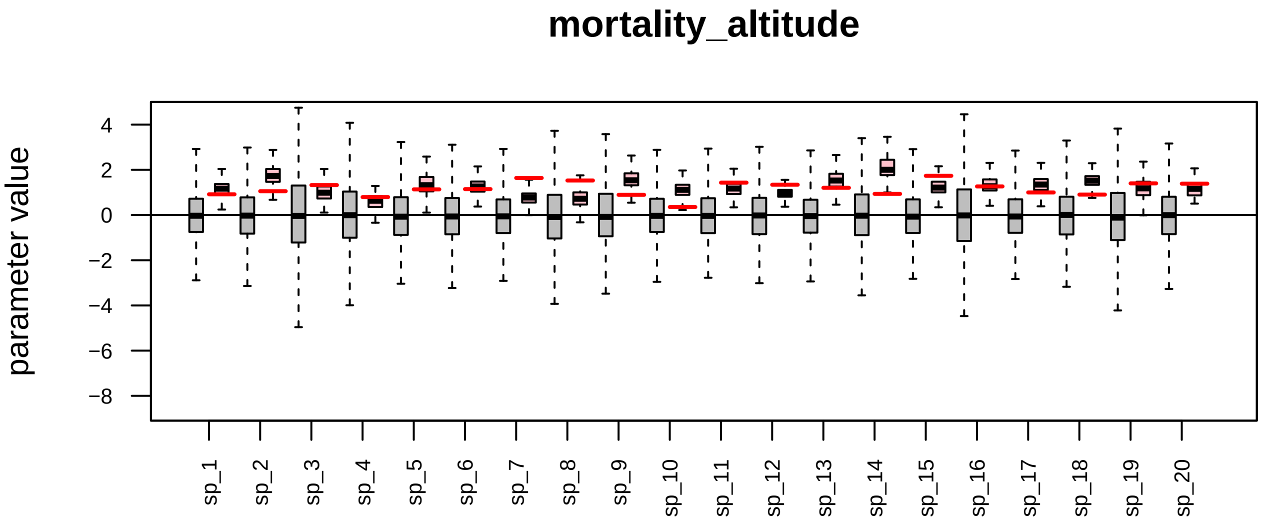

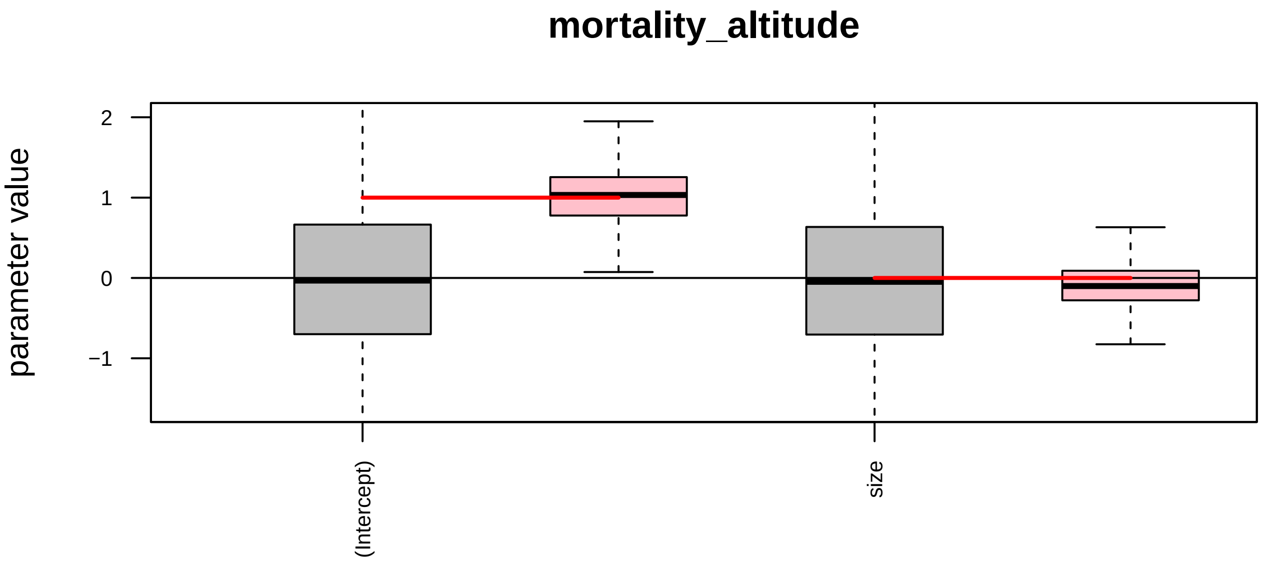

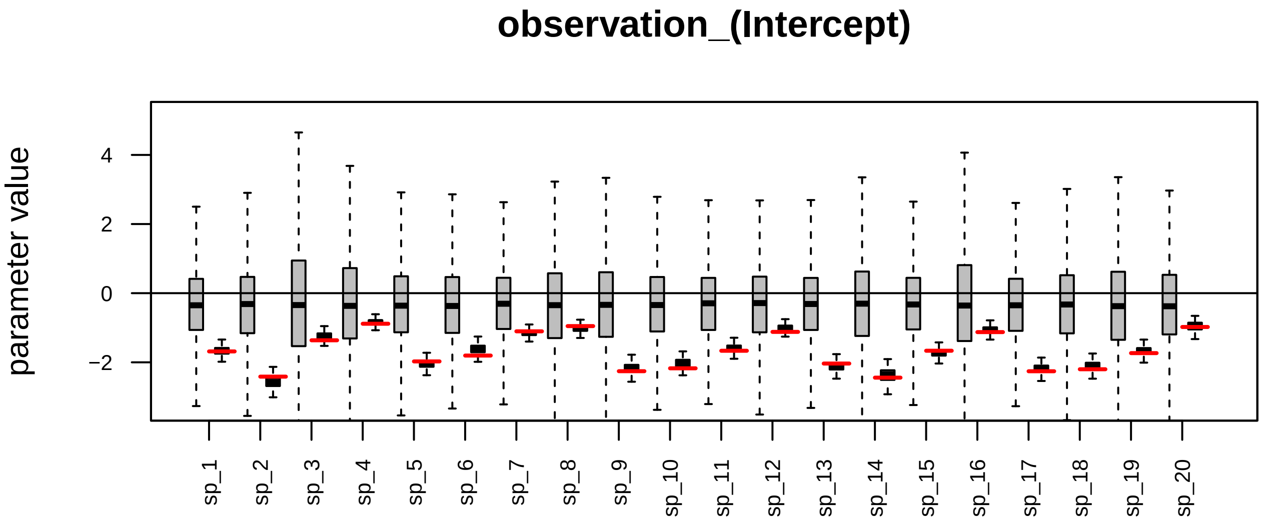

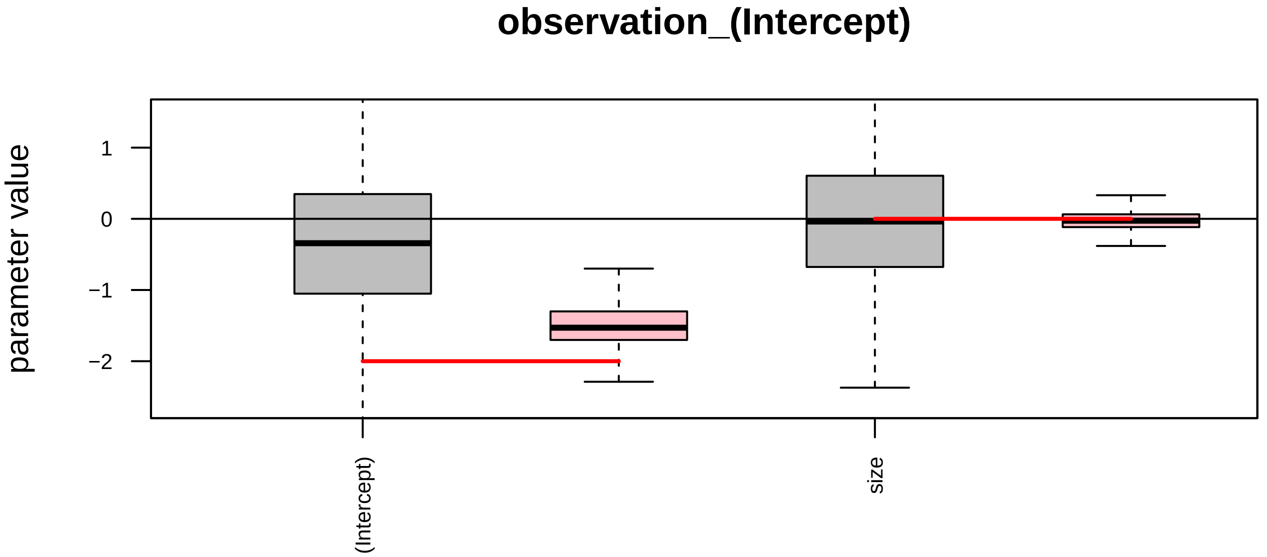

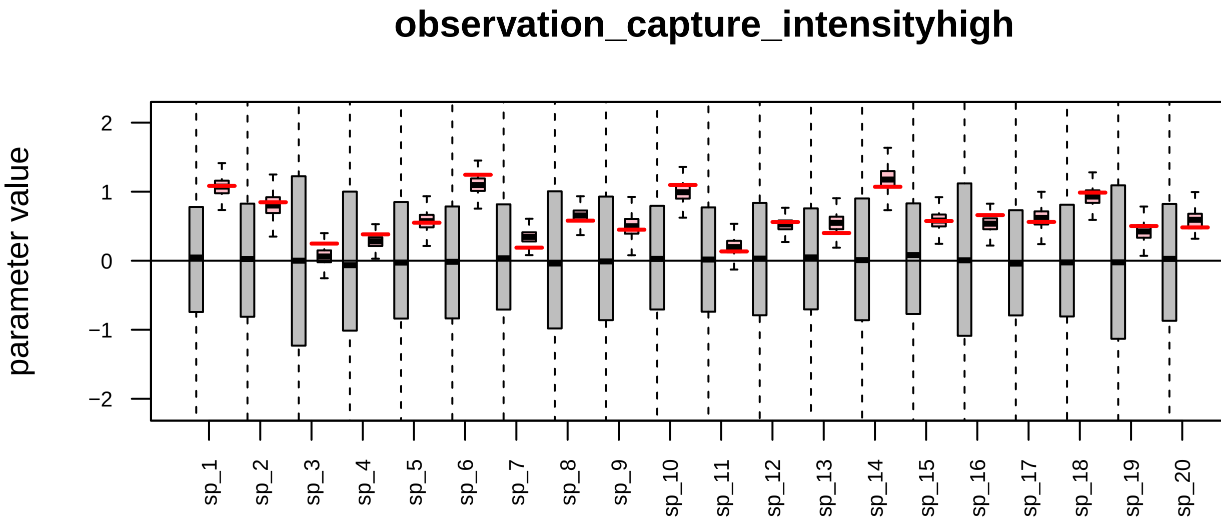

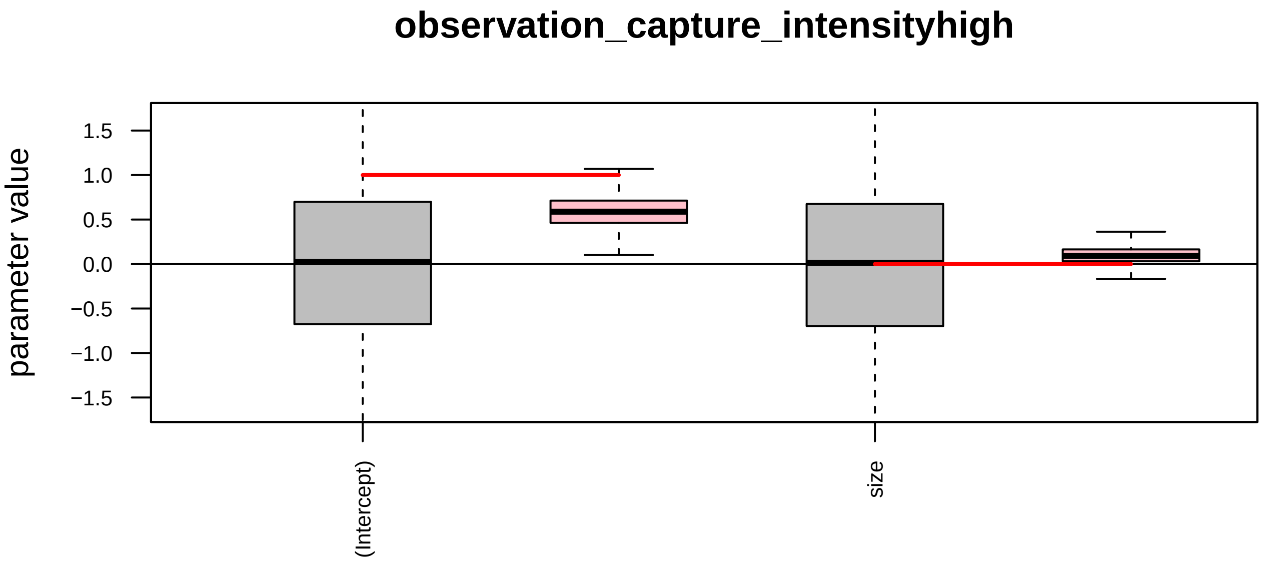

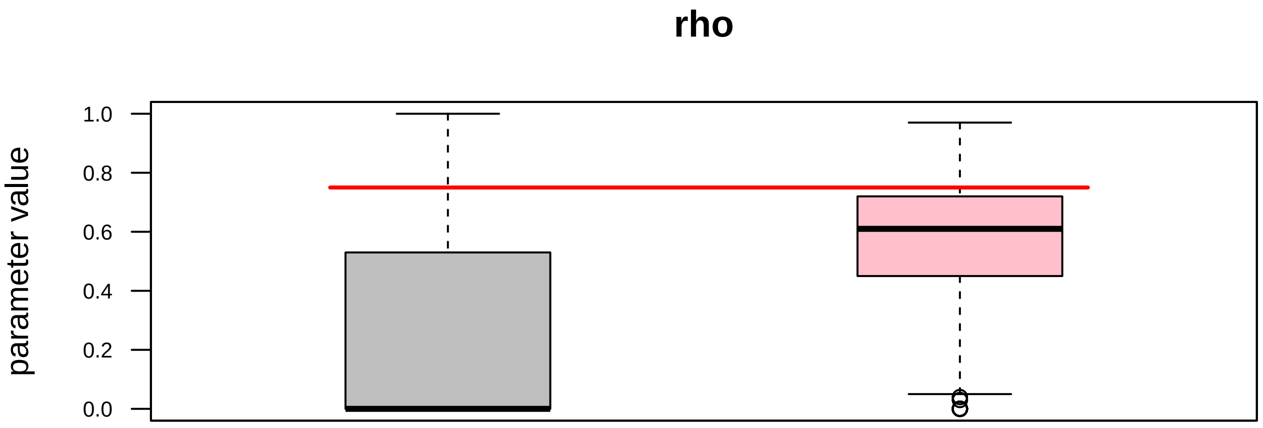

### posteriors_vs_priors_simulated_ICP.pdf

#### diffusion_(Intercept)

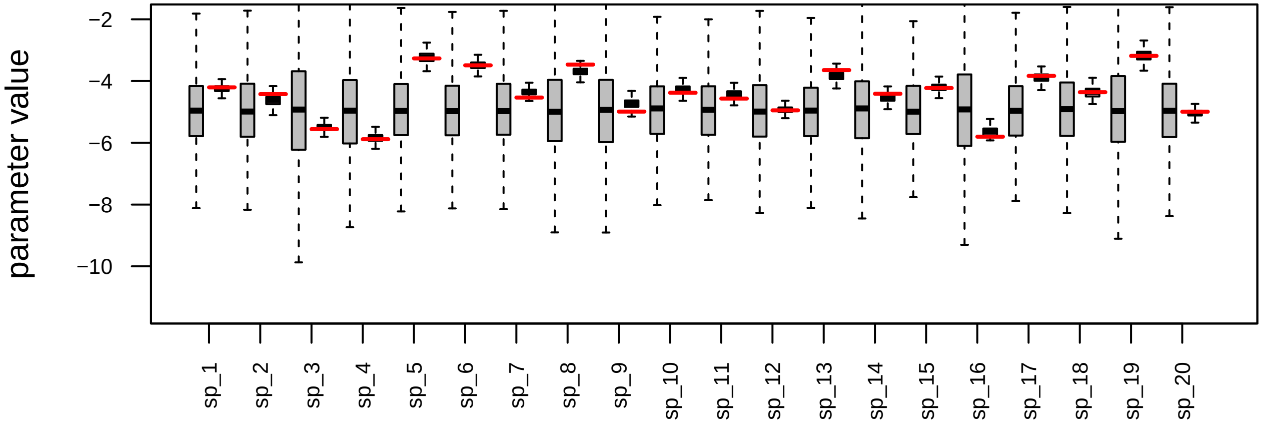


#### diffusion_(Intercept)

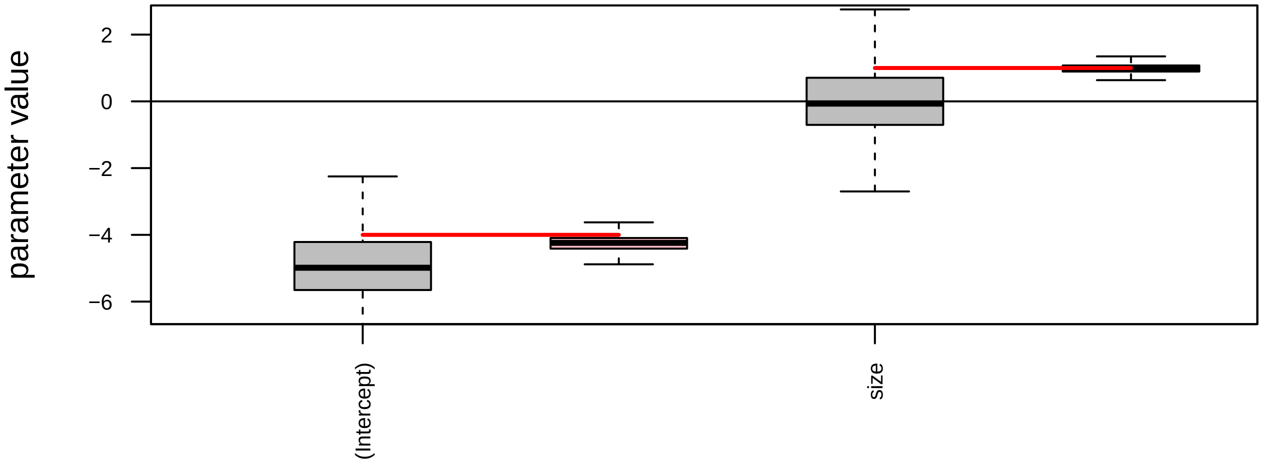

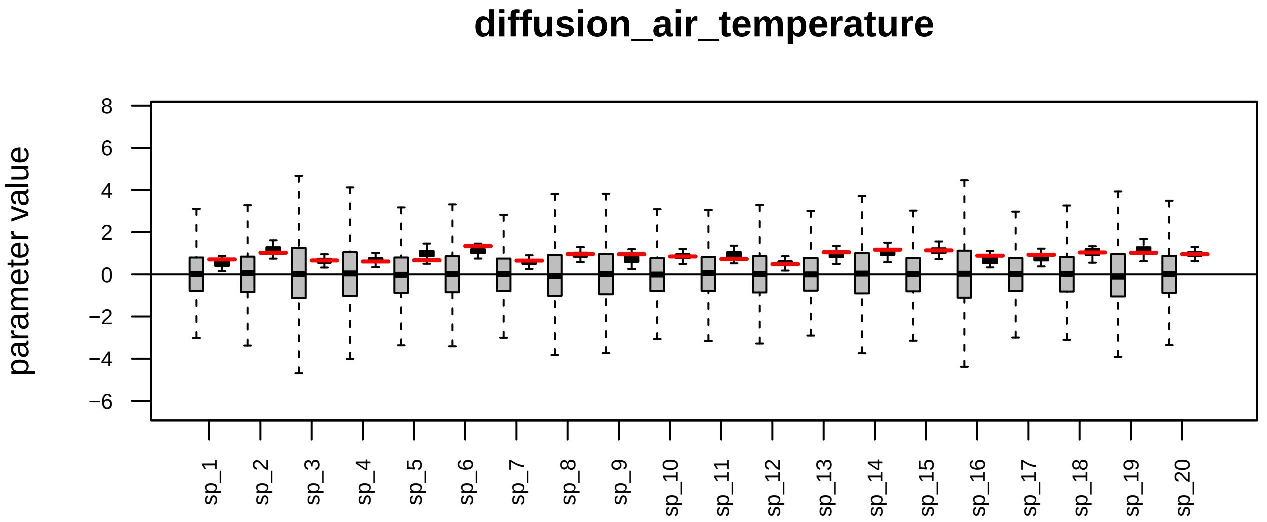

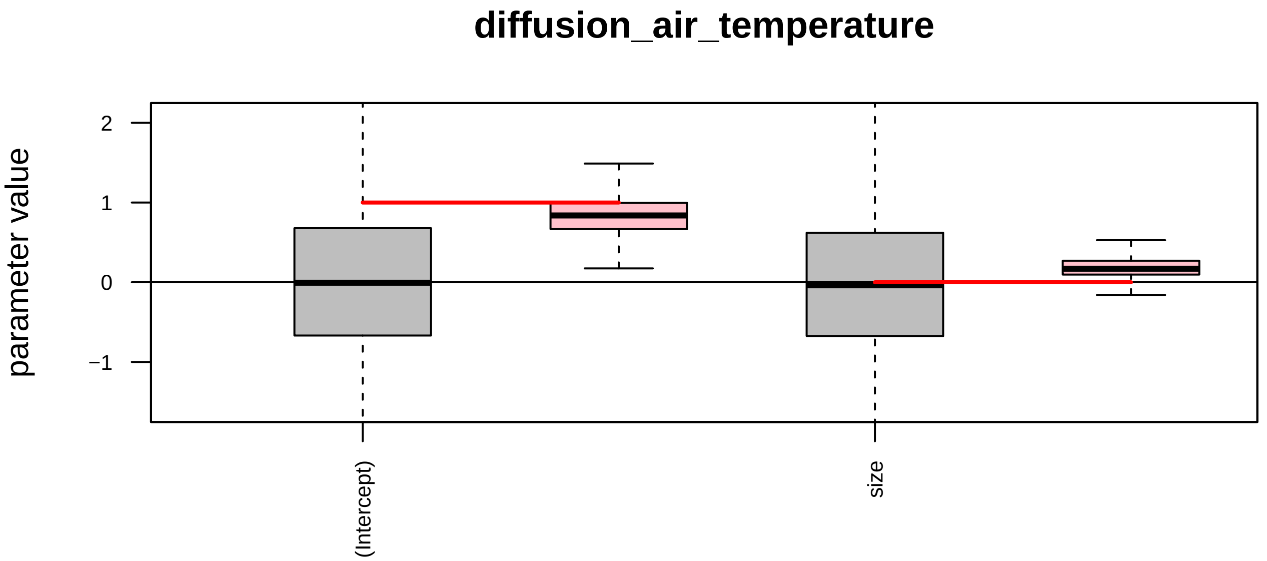

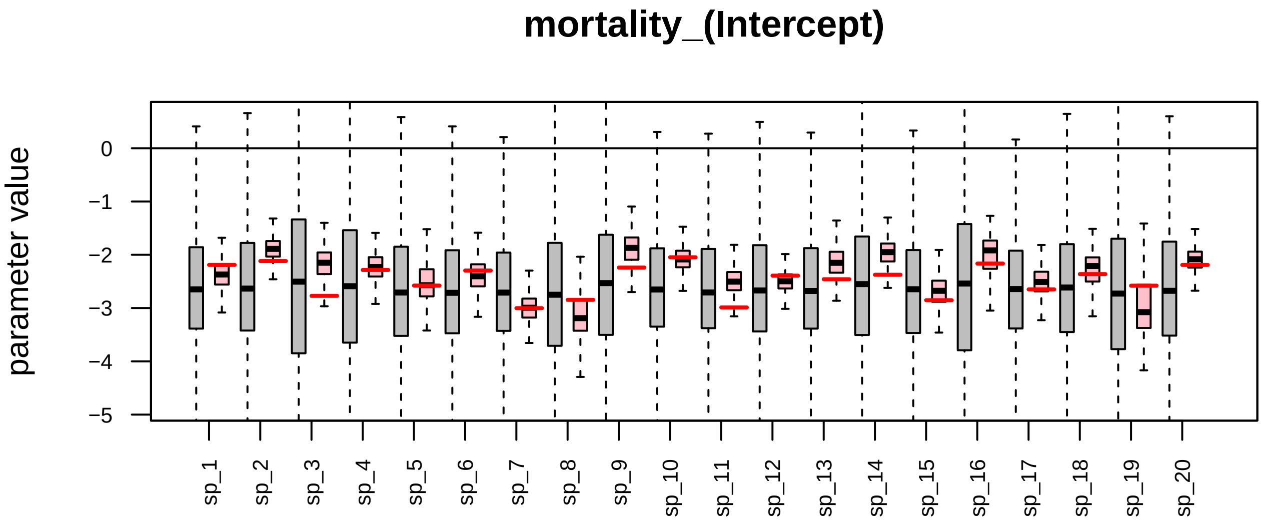

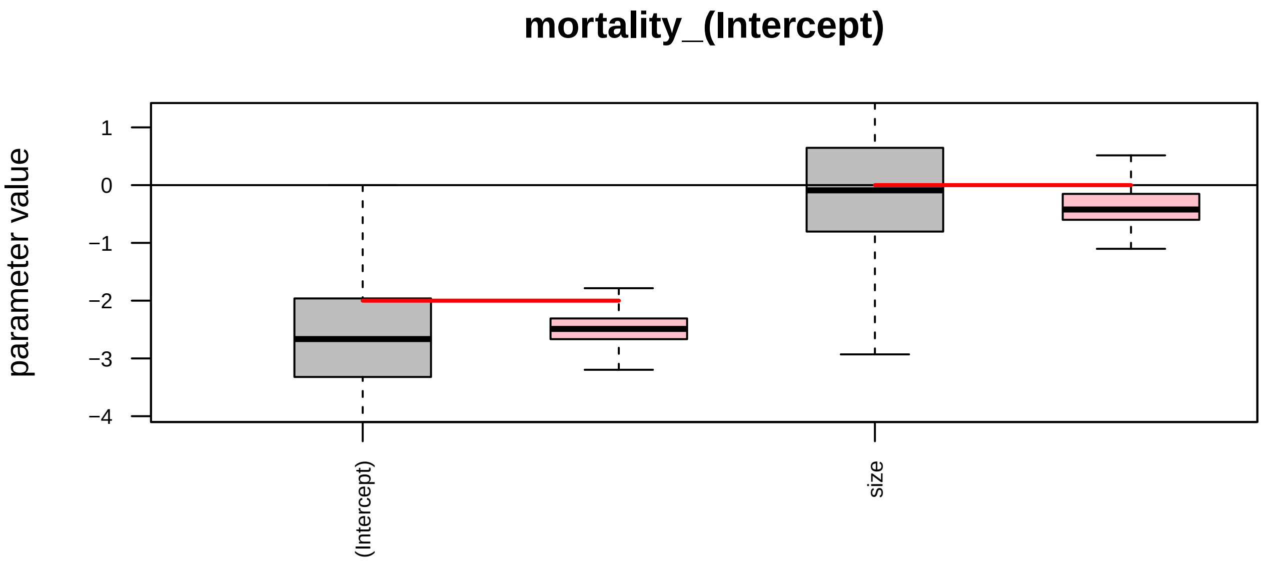

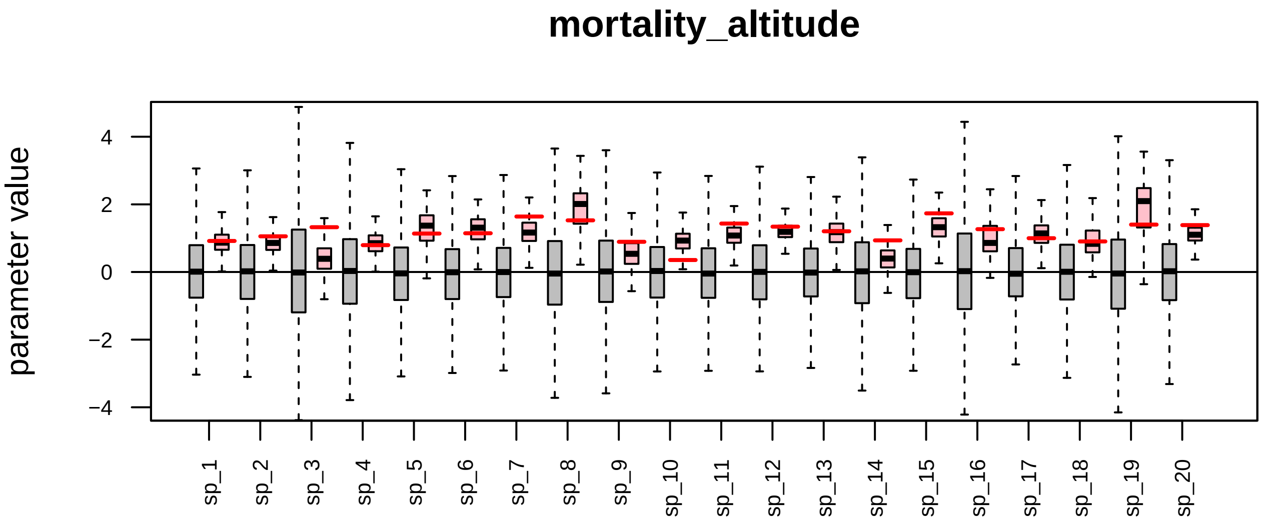

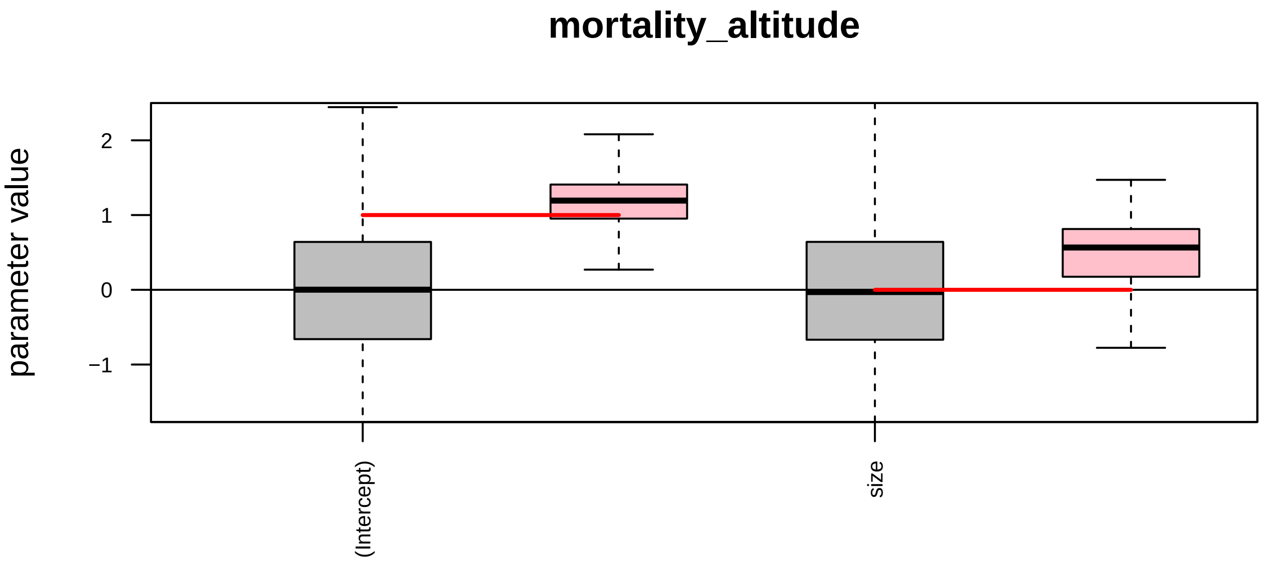

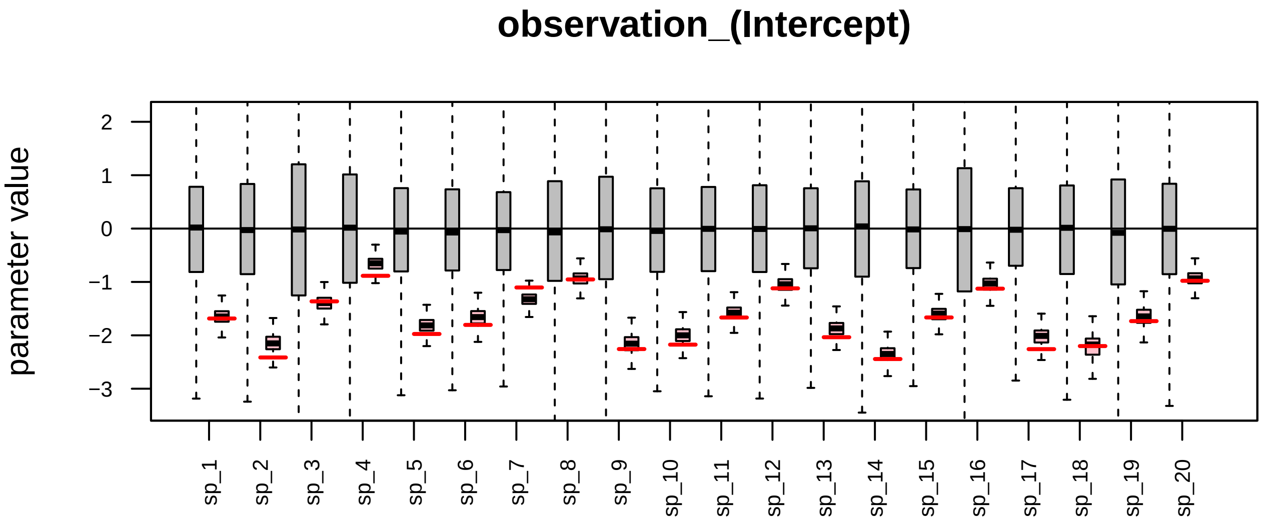

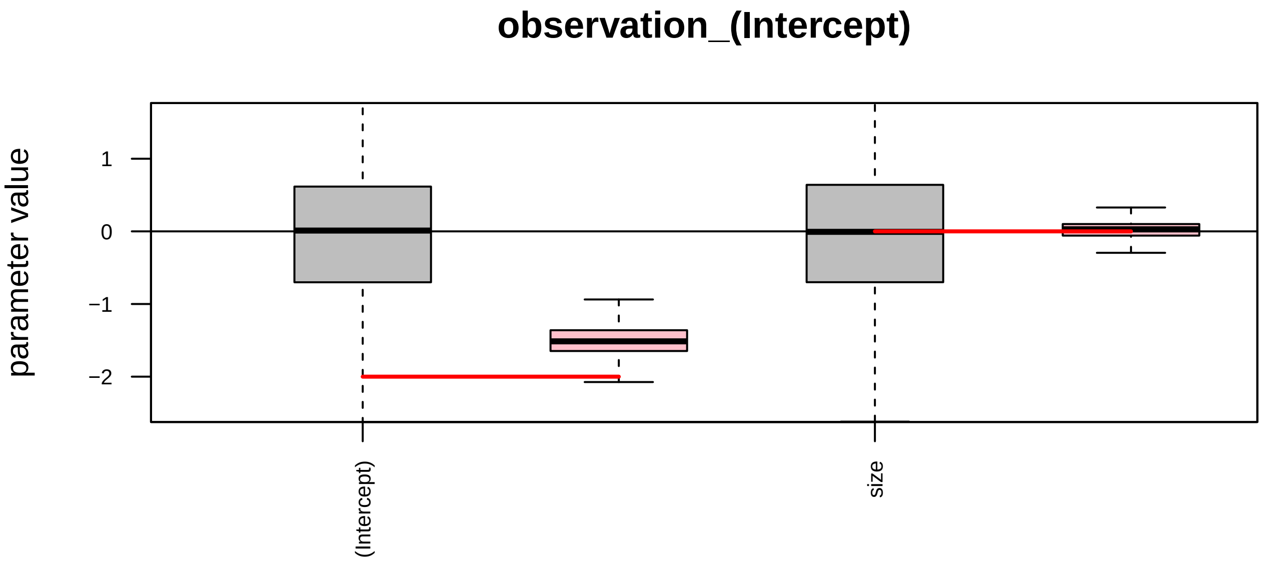

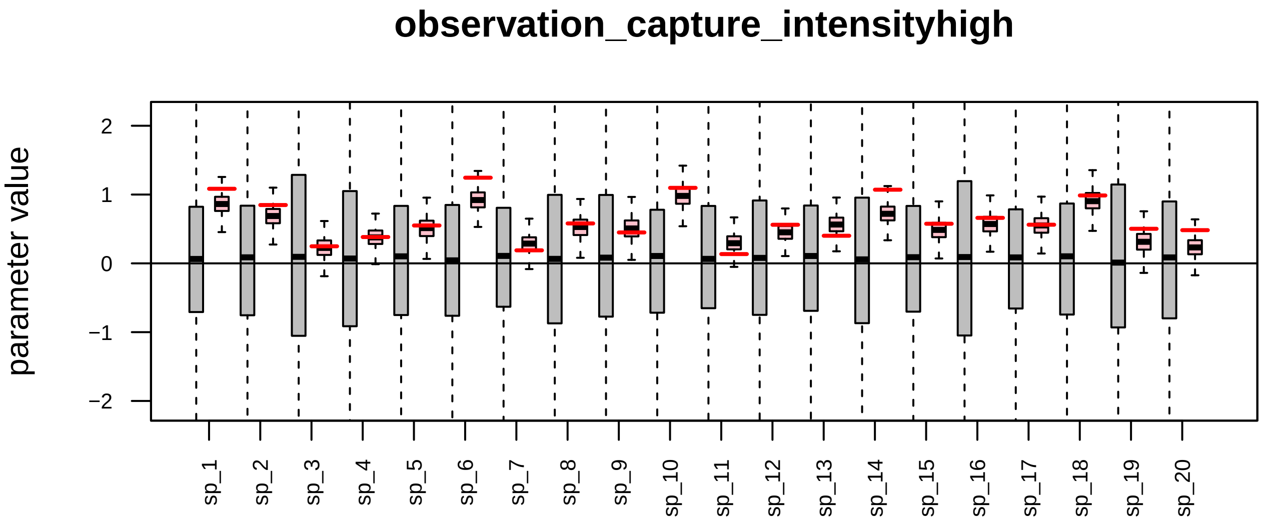

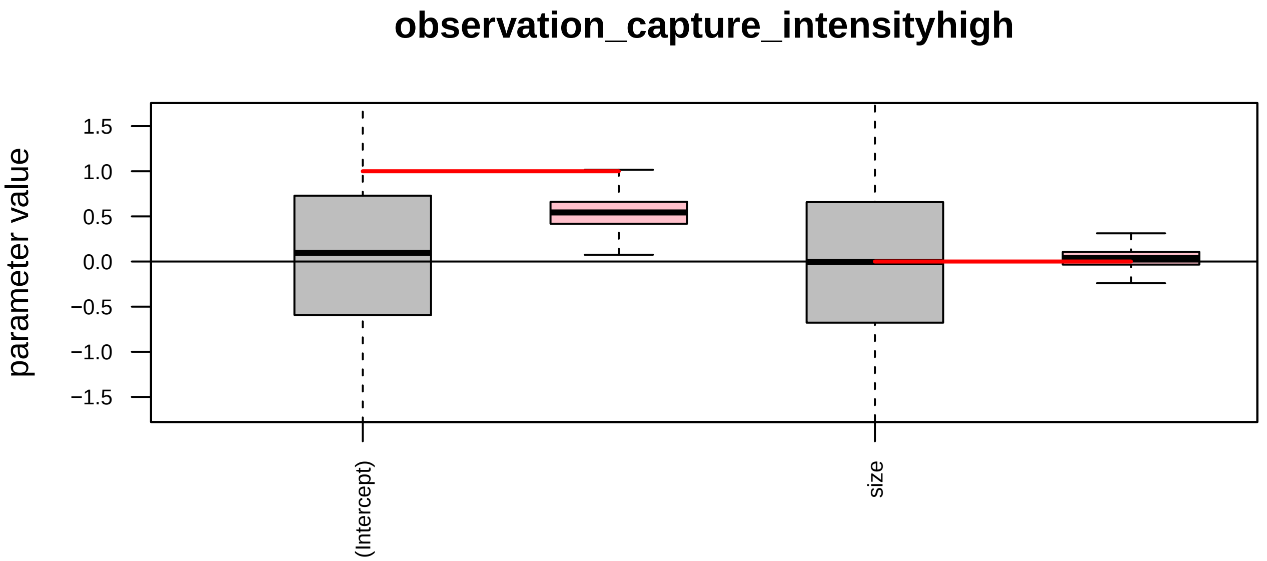

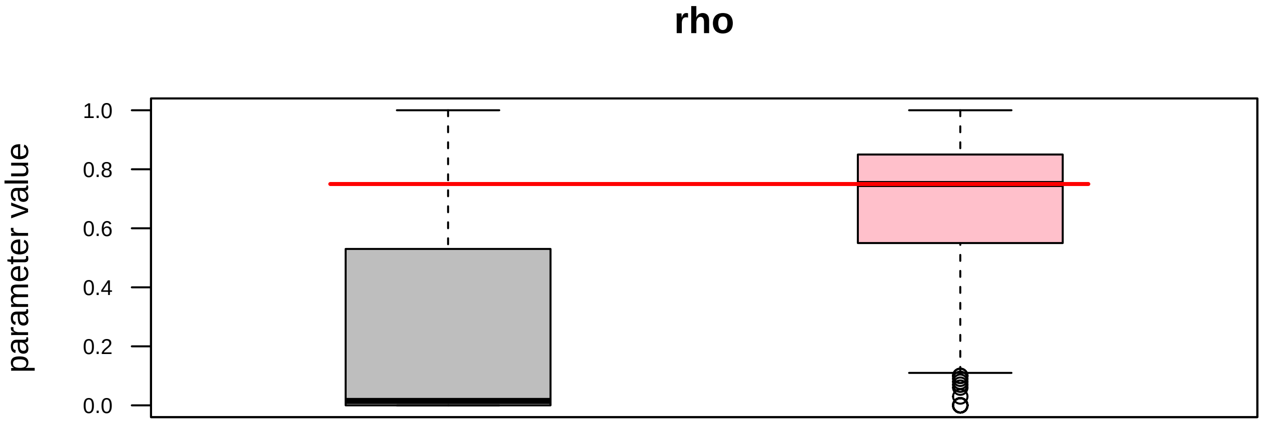
