## Supplementary material for "R-package Jsmm: Joint species movement modelling of mark-recapture data": S3 Additional information on the moths case study

**Supporting Information S3:** Luisa F. Rodriguez and Otso Ovaskainen, 2025. Joint Species Movement Modelling with the R-package Jsmm.

Table 1: Moths species identification. Species name, wingspan (mm) trait, number of marked individuals, and total number of recaptures by species.

| ID | Species name | Wingspan (mm) | N marked individuals | Total n of recaptures |
|---|---|---|---|---|
| 1 | Lymantria monacha | 44 | 418 | 42 |
| 2 | Habrosyne pyritoides | 40 | 164 | 5 |
| 3 | Phalera bucephala | 50 | 48 | 5 |
| 4 | Diachrysia chrysitis | 40 | 151 | 19 |
| 5 | Mythimna ferrago | 38 | 70 | 6 |
| 6 | Eilema lurideola | 32 | 862 | 61 |
| 7 | Craniophora ligustri | 37 | 189 | 5 |
| 8 | Apamea monoglypha | 50 | 623 | 10 |
| 9 | Eilema griseola | 35 | 954 | 35 |
| 10 | Xestia triangulum | 42 | 1022 | 166 |
| 11 | Euthrix potatoria | 55 | 609 | 37 |
| 12 | Cosmia trapezina | 34 | 1316 | 131 |
| 13 | Axylia putris | 34 | 107 | 5 |
| 14 | Agrotis exclamationis | 38 | 786 | 41 |
| 15 | Diarsia mendica | 35 | 176 | 13 |
| 16 | Hydriomena furcata | 32 | 894 | 22 |
| 17 | Noctua pronuba | 55 | 718 | 6 |
| 18 | Campaea margaritata | 40 | 442 | 8 |
| 19 | Laothoe populi | 80 | 159 | 17 |
| 20 | Crocallis elinguaria | 38 | 163 | 8 |
| 21 | Euproctis similis | 33 | 722 | 24 |

Table 2: Model Formulas declarations for the moths example.

| Model parameter | Model formula |
|---|---|
| **`diffusion`** | ~ air_temperature |
| **`advection_1`** | - |
| **`advection_2`** | - |
| **`habitat_preference`** | ~ vegetation_type |
| **`mortality`** | ~ 1 |
| **`observation`** | ~ 1 |
| **`traits`** | ~ wingspan_mm |

Table 3: Covariates for the movement parameters.

| Covariate name | Spatial | Temporal | Spatio-temporal | Type |
|---|---|---|---|---|
| air_temperature | No | Yes | No | Numeric |
| vegetation_type | Yes | No | No | Factor |

Table 4: Air temperature covariate. Values correspond to the mean measurements of the dry air temperature for the hours in which the light traps were active for each of the experiment nights (from dawn to dusk). Temperature measurements were taken from the automatic weather station T08 Wytham (51° 46' 52.86"N, 1°20'9.81"W) from the UK Environmental Change Network (ECN) (Renie et al, 2017)

| Time | Date | Air temperature (°C) | Scaled air temperature |
|---|---|---|---|
| 0 | 2009-06-14 | 12.820 | -0.278 |
| 1 | 2009-06-15 | 12.861 | -0.262 |
| 2 | 2009-06-16 | 10.999 | -0.987 |
| 3 | 2009-06-17 | 11.681 | -0.721 |
| 4 | 2009-06-18 | 10.874 | -1.035 |
| 5 | 2009-06-19 | 10.004 | -1.373 |
| 6 | 2009-06-20 | 10.366 | -1.233 |
| 7 | 2009-06-21 | 11.419 | -0.823 |
| 8 | 2009-06-22 | 11.984 | -0.603 |
| 9 | 2009-06-23 | 15.991 | 0.955 |
| 10 | 2009-06-24 | 12.051 | -0.577 |
| 11 | 2009-06-25 | 12.066 | -0.572 |
| 12 | 2009-06-26 | 15.026 | 0.579 |
| 13 | 2009-06-27 | 14.829 | 0.503 |
| 14 | 2009-06-28 | 15.160 | 0.631 |
| 15 | 2009-06-29 | 18.333 | 1.865 |
| 16 | 2009-06-30 | 19.277 | 2.232 |
| 17 | 2009-07-01 | 20.594 | 2.744 |
| 18 | 2009-07-02 | 18.627 | 1.980 |
| 19 | 2009-07-03 | 19.121 | 2.172 |
| 20 | 2009-07-04 | 14.770 | 0.480 |
| 21 | 2009-07-05 | 14.549 | 0.394 |
| 22 | 2009-07-06 | 13.299 | -0.092 |
| 23 | 2009-07-07 | 12.793 | -0.289 |
| 24 | 2009-07-08 | 11.319 | -0.862 |
| 25 | 2009-07-09 | 11.064 | -0.961 |
| 26 | 2009-07-10 | 9.716 | -1.485 |
| 27 | 2009-07-11 | 13.401 | -0.052 |
| 28 | 2009-07-12 | 13.949 | 0.161 |
| 29 | 2009-07-13 | 13.324 | -0.082 |
| 30 | 2009-07-14 | 13.380 | -0.061 |
| 31 | 2009-07-15 | 13.932 | 0.154 |
| 32 | 2009-07-16 | 13.684 | 0.057 |
| 33 | 2009-07-17 | 13.550 | 0.005 |
| 34 | 2009-07-18 | 12.606 | -0.361 |
| 35 | 2009-07-19 | 12.505 | -0.401 |
| 36 | 2009-07-20 | 11.781 | -0.682 |

| Time | Date | Air temperature (°C) | Scaled air temperature |
|---:|---|---:|---:|
| 37 | 2009-07-21 | 13.361 | -0.068 |
| 38 | 2009-07-22 | 13.319 | -0.084 |
| 39 | 2009-07-23 | 12.409 | -0.438 |
| 40 | 2009-07-24 | 12.178 | -0.528 |

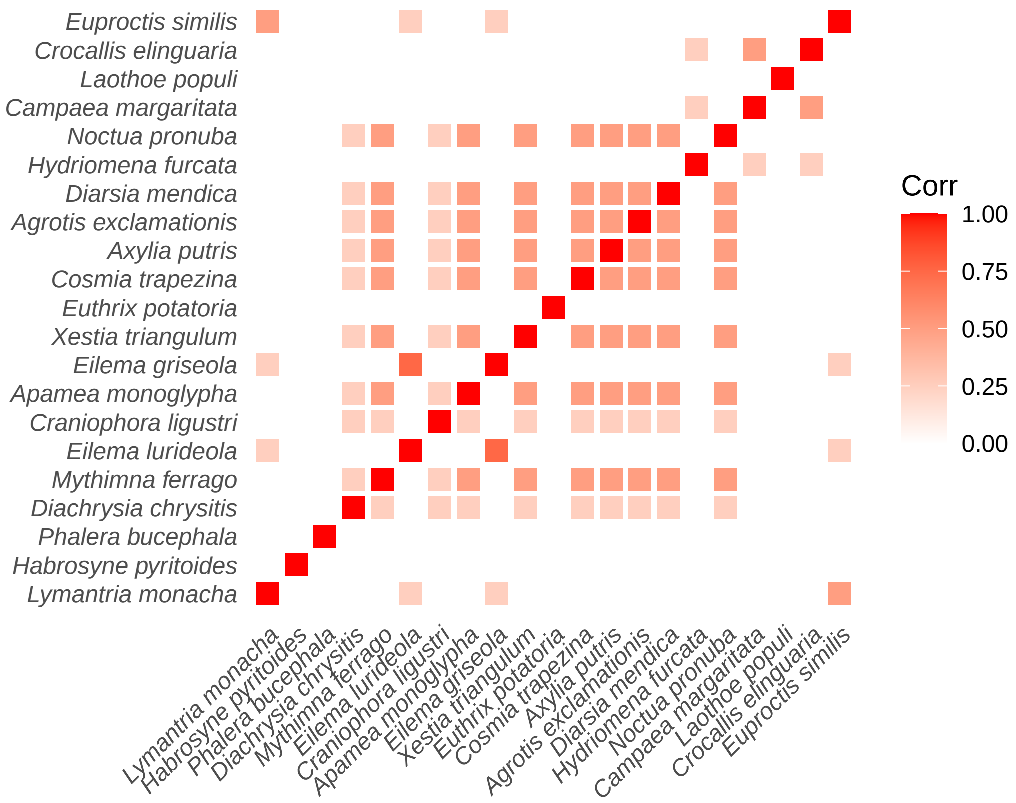


Figure 1: Phylogenetic correlation matrix.

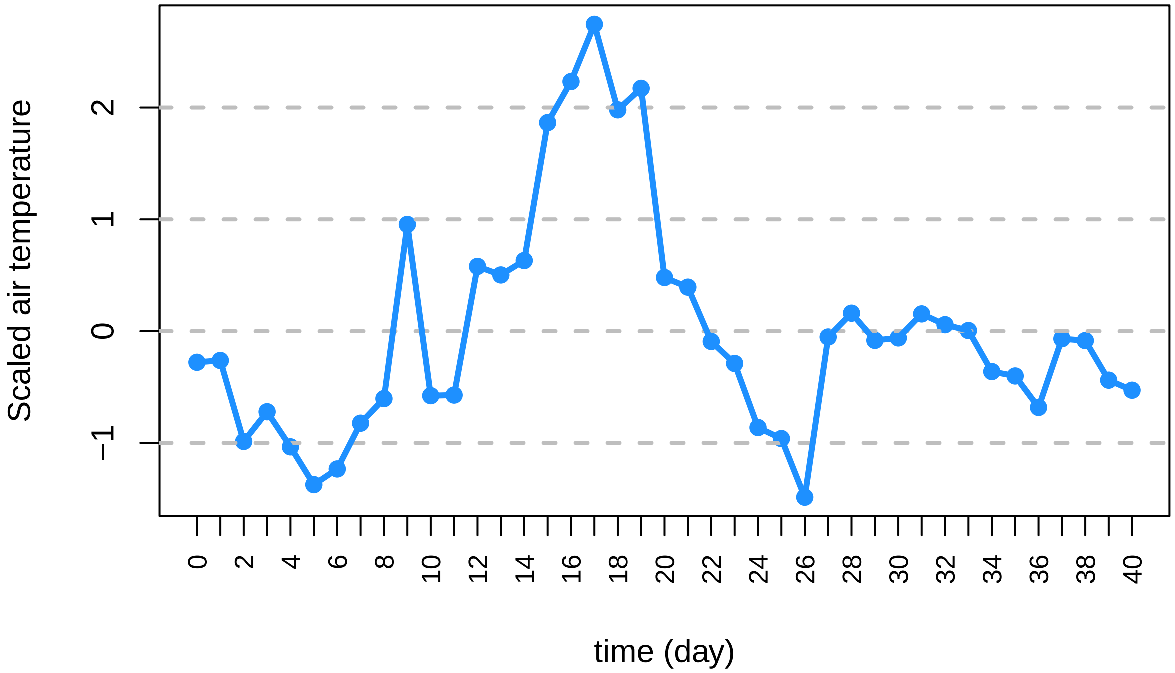


Figure 2: Scaled dry air temperature which was originally in Celcius degrees. Values in Table 2 S 04.

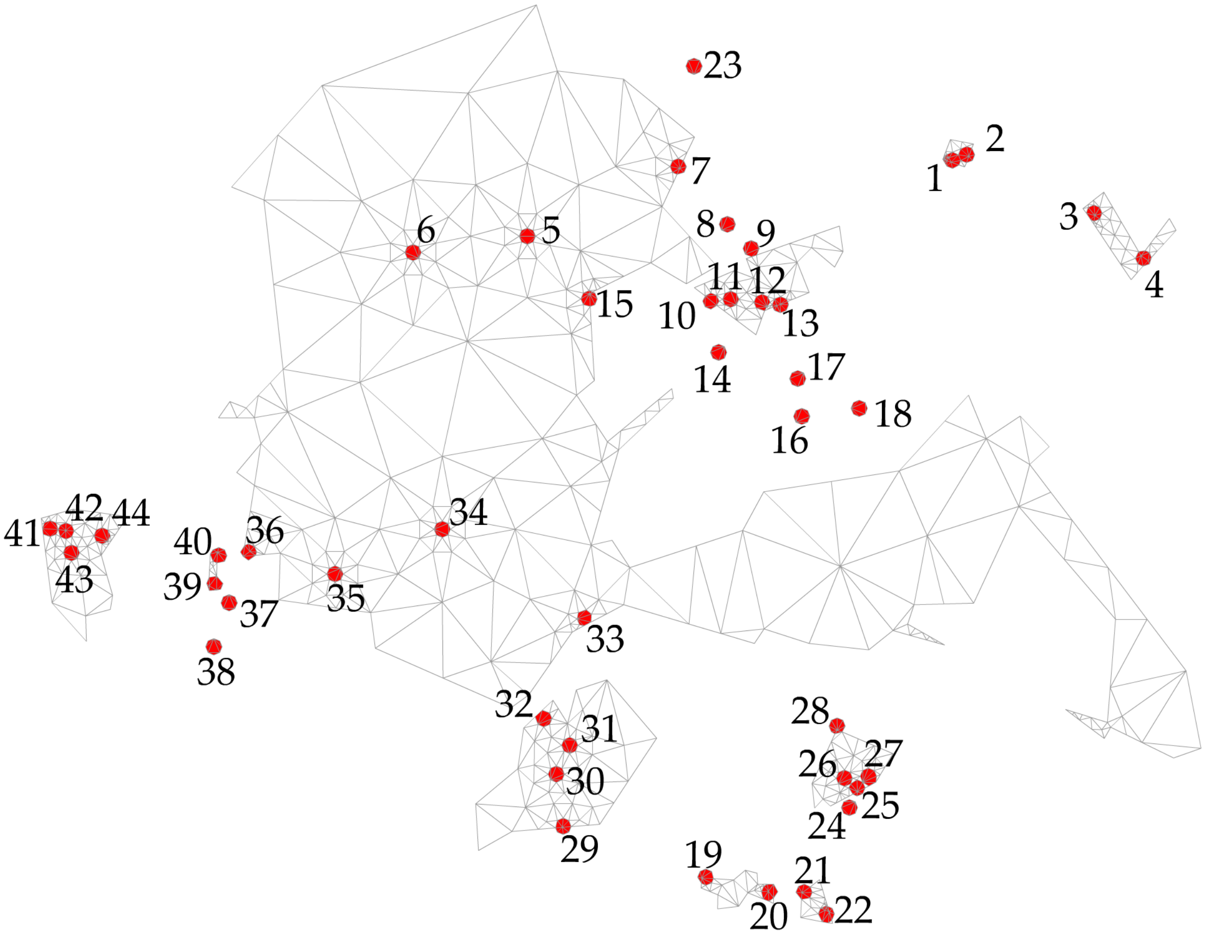


Figure 3: Detail of the 44 6W-actinic light traps locations that correspond to the release and capture sites in which the experiment was conducted.

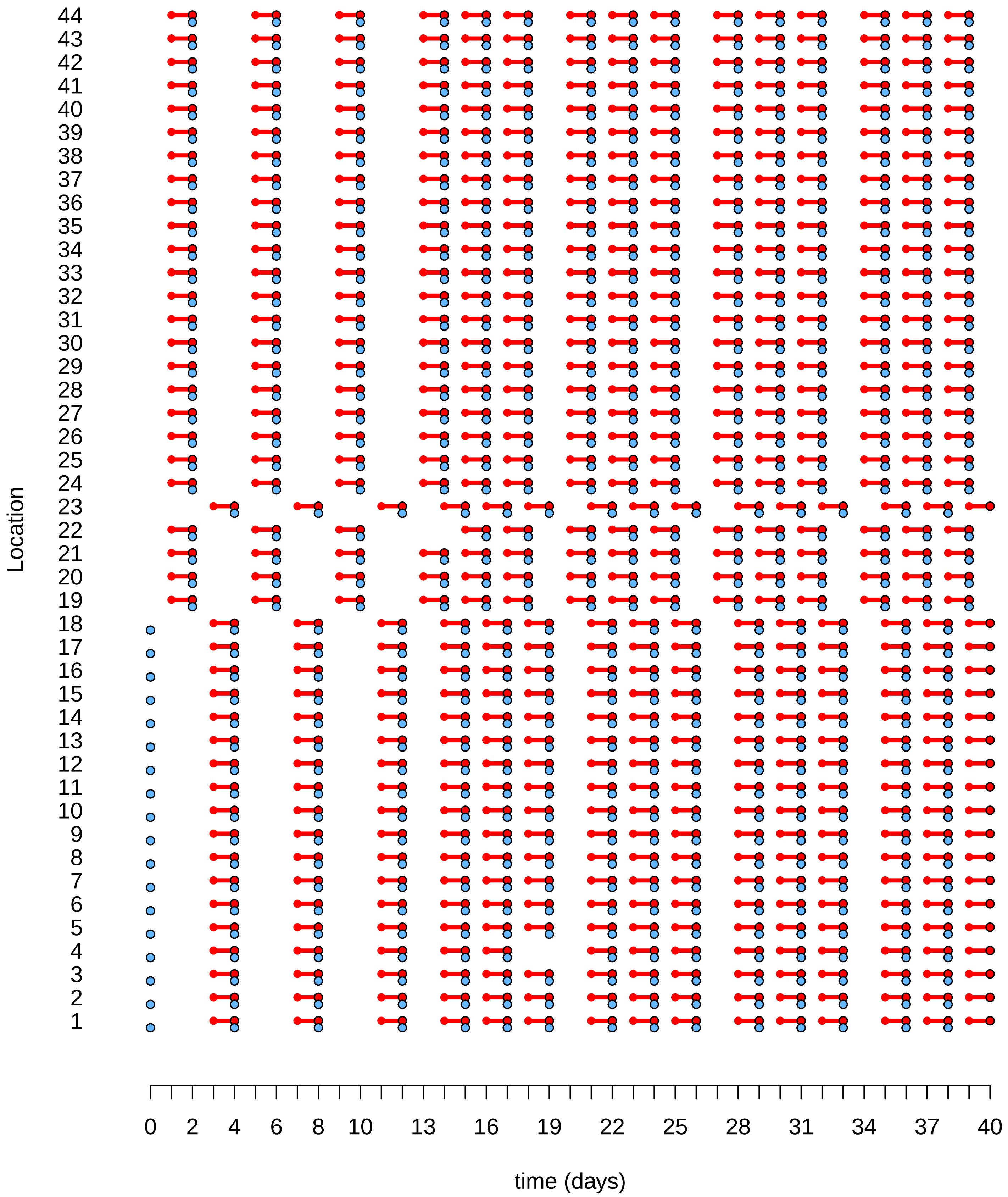


Figure 4: Observation effort scheme for CPP capture process moths example. Times in which releases and captures are possible are represented by blue, and red colors, respectively.
